# Conserved herpesvirus protein kinase (CHPK)-mediated phosphorylation of viral proteins associated with nucleocytoplasmic trafficking during natural infection

**DOI:** 10.64898/2026.07.07.737029

**Authors:** Haji Akbar, Nagendraprabhu Ponnuraj, Bushra Fazal Minhas, Christopher A. Gaulke, Stephen J. Spatz, Keith W. Jarosinski

## Abstract

The conserved herpesvirus protein kinase (CHPK) is encoded by all members of the *Orthoherpesviridae* and contributes to replication in cell culture but is not strictly required. Marek’s disease virus (MDV) CHPK is dispensable for replication in cultured cells yet essential for horizontal transmission in chickens. To elucidate its role during natural infection, we performed RNA sequencing (RNA-seq) and mass spectrometry (MS)-based phosphoproteomics on spleen and feather follicle epithelial skin cells from chickens infected with wild-type or CHPK-null MDV. RNA-seq detected only a limited number of viral transcripts in the spleen—including latency-associated transcripts (LATs) and the major oncogene Meq—with minimal differences between wild-type and CHPK-null infections. In feather follicle epithelial skin cells, the full repertoire of viral genes was expressed, but only seven genes showed differential expression between wild-type and CHPK-null viruses. In striking contrast, MS-based phosphoproteomics identified many differentially phosphorylated proteins, including 21 viral proteins. These findings indicate that CHPK’s critical functions in skin replication and subsequent horizontal transmission are primarily mediated through post-translational modifications (PTMs) rather than transcriptional regulation. Among the CHPK-targeted viral proteins were three MDV-unique proteins, eight conserved within the *Alphaherpesvirinae*, and ten conserved across the *Orthoherpesviridae*. *In silico* analysis revealed that many differentially phosphorylated serine and threonine residues lie near or within predicted nuclear localization signals (NLS) and nuclear export signals (NES). Functional validation confirmed that several of these motifs actively control nucleocytoplasmic shuttling of the respective viral proteins. Collectively, these data suggest that MDV CHPK orchestrates the subcellular localization of multiple viral proteins in epithelial skin cells via phosphorylation, thereby enabling efficient replication and horizontal transmission in the natural host.

**AUTHOR SUMMARY:** Understanding the mechanisms by which herpesviruses replicate and spread within their natural hosts and identifying the viral genes essential for these processes are fundamental to developing effective antiviral strategies. Marek’s disease virus (MDV), a highly contagious alphaherpesvirus, remains a major economic threat to the global poultry industry while serving as a powerful natural animal model for studying herpesvirus pathogenesis and transmission *in vivo*. Using an established *in vivo* enrichment method for infected cells, we conducted a comprehensive analysis of viral gene expression, protein abundance, and post-translational modifications (PTMs) during natural infection. Remarkably, RNA sequencing revealed virtually no differences in viral transcription between wild-type and CHPK-null viruses in either spleen or feather follicle epithelial skin cells. In contrast, phosphoproteomics showed that CHPK extensively regulates the phosphorylation of multiple viral proteins specifically in skin epithelial cells. *In silico* and functional analyses further indicate that these CHPK-mediated phosphorylations occur near or within nuclear localization (NLS) and nuclear export (NES) signals, directly controlling the nucleocytoplasmic shuttling of key viral proteins. This work suggests CHPK as a master regulator of viral protein subcellular localization during replication in the natural host and highlights CHPK orthologs as promising broad-spectrum therapeutic targets against herpesviruses.

## INTRODUCTION

Herpesviruses are ubiquitous in animals and highly adapted to their host species, having evolved together over millions of years [1]. Despite a broad host range and pathogenic potential, a core set of genes is shared by all members of the *Orthoherpesviridae,* including the conserved herpesvirus protein kinase (CHPK). The extent of this conservation suggests a vital role for CHPK in the herpesvirus replication cycle, even though the overall sequence of these proteins can vary considerably. CHPKs are serine/threonine kinases [2] and are more conserved within their respective *Alphaherpesvirinae*, *Betaherpesvirinae*, and *Gammaherpesvirinae* subfamilies [3]. CHPK orthologs are components of the tegument and play crucial roles during viral entry and the initiation of infection [4–6]. They have been linked to multiple processes at the mechanistic level, including nuclear egress of capsids [7–13], tegument association/dissociation [14, 15], viral gene expression [8, 16–20], viral DNA replication [8, 12, 13, 21–26], DNA damage responses [27, 28], cell-cycle regulation [29–32], evasion of the interferon response [33–35], and viral protein expression or stability [20, 36].

CHPK is required for the horizontal transmission of gallid alphaherpesvirus 2 (GaAH2), widely referred to as Marek’s disease virus (MDV) [20, 37, 38] and gallid alphaherpesvirus 3 (GaAHV3) [39] in chickens. MDV causes Marek’s disease (MD), which clinically presents as neurological symptoms and cancerous tumors in the form of transformed T cells. Natural infection with MDV begins with the inhalation of infectious virus previously shed from chicken skin dander, and cytolytic infection initiates in B and T lymphocytes [40]. Infected immune cells subsequently disseminate to lymphoid organs, where the virus is spread through direct cell-to-cell contact. It is believed that some late viral proteins required for cell-free production are suppressed during lymphocyte replication, termed semi-productive replication. To disseminate into the environment, infected immune cells travel to the skin, where productive replication ensues in epithelial cells surrounding feather follicles, termed feather follicle epithelial (FFE) skin cells. In these cells, new infectious particles are produced and shed into the environment, a mode of dissemination similar to that of varicella-zoster virus (VZV), which causes chickenpox and shingles in humans [41].

MDV gene expression follows the typical alphaherpesvirus cascade, with herpes simplex virus (HSV) being the best understood. Following virus infection, immediate-early (IE) genes activate early (E) genes involved in DNA replication, and late (L) structural genes are then expressed [42]. Like other members of the *Orthoherpesviridae* family, MDV can replicate both *in vitro* and *in vivo*. However, MDV maturation is impaired *in vitro*. Nucleocapsids assemble in the nucleus and acquire a primary envelope at the inner nuclear membrane, but secondary envelopment and the production of extracellular cell-free virions are minimal or absent in most standard monolayer cultures [43]. It has been observed that the expression of some late viral structural proteins is absent or dysregulated [44, 45], resulting in progeny viruses that are strictly cell-associated and require direct contact for cell-to-cell spread (sometimes referred to as semi-productive replication). In contrast, all viral proteins required for cell-free virus production (sometimes referred to as fully productive replication) are abundantly expressed in FFE cells in chicken skin, resulting in the shedding of infectious virus and its dissemination within the population [46]. MDV CHPK is critical for the production of cell-free virus in epithelial skin cells, and current data suggest that it is important for late viral gene expression [20].

Many herpesviral proteins are localized to the nucleus and may contribute to host cell functions, such as proliferation and differentiation [47]. These proteins typically encode nuclear localization signals (NLS) and nuclear export signals (NES) that direct their shuttling between the nucleus and cytoplasm. The importation of proteins into the nucleus is mediated by the importin alpha/beta nuclear import pathway, which involves interactions between importins and the NLS sequences [48]. There are three types of NLS: 1) classical monopartite NLS consisting of a single cluster of basic amino acids [K(K/R)X(K/R)], 2) classical bipartite with two separate clusters of basic amino acids [(R/X(X)10-12KRXK), where X is any residue] [49], and 3) non-classical NLS (ncNLS) with no recognizable signal sequence [50–54]. Conversely, the nuclear export signal (NES) mediates the export of proteins from the nucleus through interactions with exportins such as CRM1 [55]. These NES motifs typically contain ordered hydrophobic residues such as Φ1-X2,3-Φ2-X2,3-Φ3-X-Φ4 (where Φn represents L, V, I, F, or M; X denotes any amino acid). Oftentimes, nucleocytoplasmic proteins encode both NLS and NES motifs, allowing them to shuttle between the cytoplasm and the nucleus.

It is well established that protein phosphorylation is a common post-translational modification (PTM) that regulates protein function [56]. For example, phosphorylation of specific serine and threonine residues near functional NLS motifs governs the movement of the HSV proteins UL47_TEG5 and UL54_ICP27 between the cytoplasm and the nucleus [57, 58]. In summation of our work studying MDV UL13_CHPK, we have shown: 1) the expression, localization, and stability of MDV CHPK are regulated through phosphorylation [20], 2) the expression of late viral proteins is severely affected in the absence of CHPK activity [20], 3) there is a tight cooperative regulation of MDV UL13_CHPK and US10, including their subcellular colocalization [36]. While initial data suggested that CHPK regulates late genes post-transcriptionally, more conclusive evidence is needed. Here, we combined RNA sequencing (RNA-seq) and mass spectrometry (MS)-based proteomics to elucidate viral targets of CHPK during fully productive replication in the host.

## RESULTS and DISCUSSION

### Experimental design for combined RNA sequencing and MS-based proteomics

The overall experimental design is shown in Figure 1 and is further described in the Materials and Methods section. Briefly, chickens were experimentally infected with vCHPKwt/10HA or vΔCHPK/10HA, which have been previously published [36]. Both viruses are based on the MDV strain RB-1B, which expresses UL47eGFP [59]. In addition, 3×Flag and 2×HA epitopes were inserted in frame at the C-termini of UL13 (CHPK) and US10, respectively, and UL47eGFP was expressed [36]. For brevity, each virus is referred to as vCHPKwt and vΔCHPK hereafter. Part of this work (vCHPKwt) RNA-seq and MS-based proteomics has been published previously [60].

**Fig 1.**
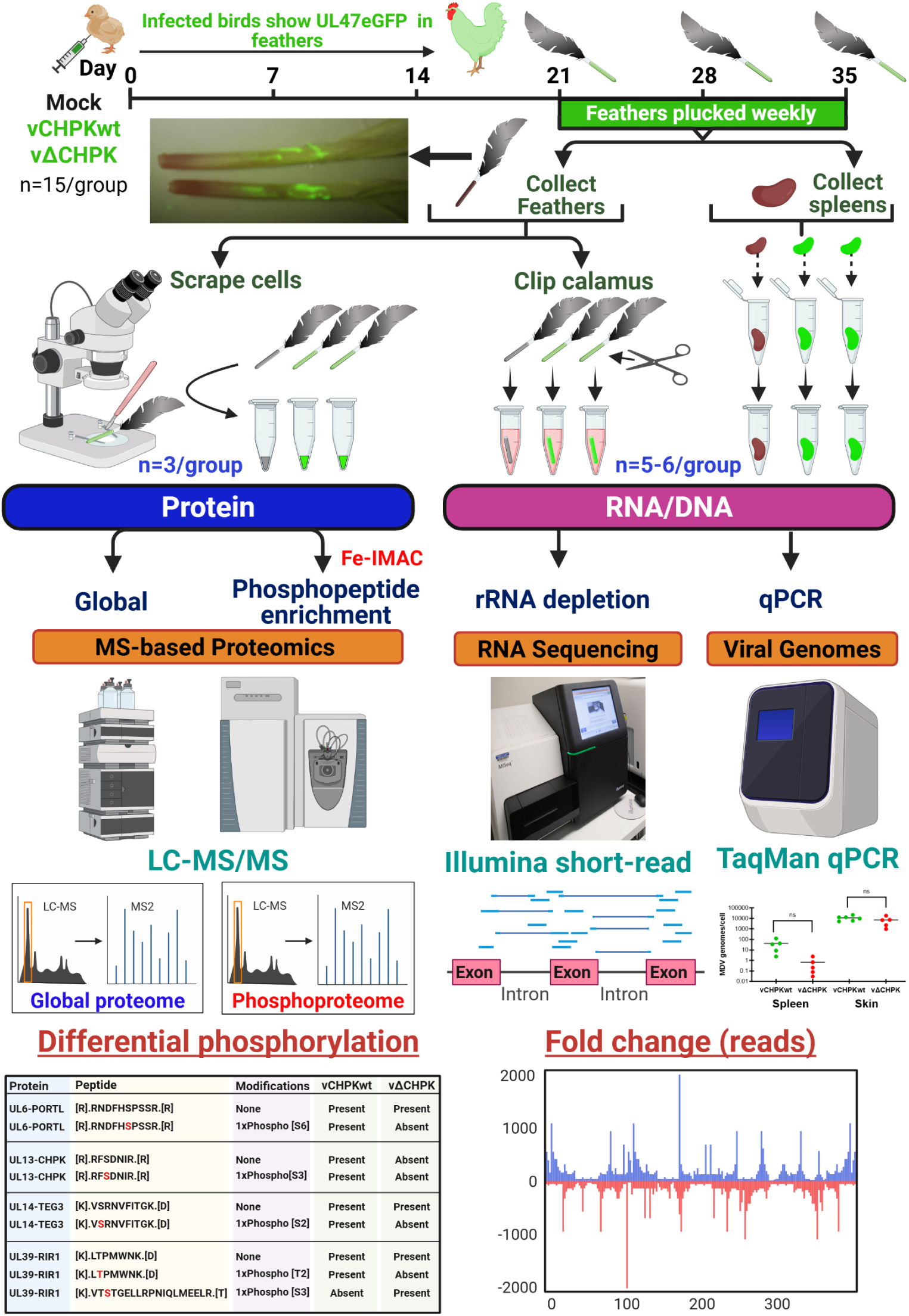
Schematic illustration of the experimental approach. Three-week-old chicks were experimentally infected with vCHPKwt and vΔCHPK or mock-infected (n = 15 per group). Feathers were plucked weekly, and birds with abundant expression of UL47eGFP were humanely euthanized for sample collection. An equal number of age-matched chickens were used as mock-infected controls. Approximately ten to fifteen feathers per bird were directly clipped at the calamus, dropped in ice-cold RNA STAT60, snap-frozen on dry ice, and stored at -80°C until processed for RNA extraction. Spleens were snap-frozen in RNA STAT-60 and stored at -80°C. After sample collection, all samples were processed for RNA extraction. Approximately 3-4 feathers per bird (n=3/group) were used to collect skin protein by scraping fluorescent cells into an Eppendorf tube, and all samples were stored at -80°C until transferred to the University of Illinois Proteomics core for MS-based proteomics (see Materials and Methods). RNA sequencing and LC-MS/MS were performed on samples. Protein expression, phosphoproteomic, and RNA-seq data were analyzed.

Feathers were collected between 21 and 35 days post-infection from chickens expressing abundant UL47eGFP in their feathers, a marker of late viral gene expression, as described in a previous animal experiment [36]. A summary of clinical status, relative UL47eGFP expression, and the days post-infection at which samples were collected is shown in S1 Table. Feathers and spleens were collected from birds one day after evaluation. After humane euthanasia, 20-30 wing and breast feathers were plucked from vCHPKwt- and vΔCHPK-infected chickens, as well as an equal number from mock-infected chickens. RNA (n=5-6/group) was collected from spleen and skin, and protein (n=3/group) from skin surrounding the feather follicle was isolated as previously described [60]. Following RNA extraction from spleen and skin samples, DNA was collected to quantify viral genomes, which showed no significant difference between the two infected groups in either tissue (Fig 2A, B). Overall, gene expression profiles differed significantly in feathers (ANOVA, dev = 1,751, *P* = 0.017) and spleens (dev = 2,934, *P* = 0.013) of animals infected with vCHPKwt and vΔCHPK. These differences were consistent with the separation observed in PCA of variance-stabilized expression values (Fig 2C, D). Viral gene expression across the viral genome was similar between samples within tissues (S1 and S2 Figs)

**Fig 2.**
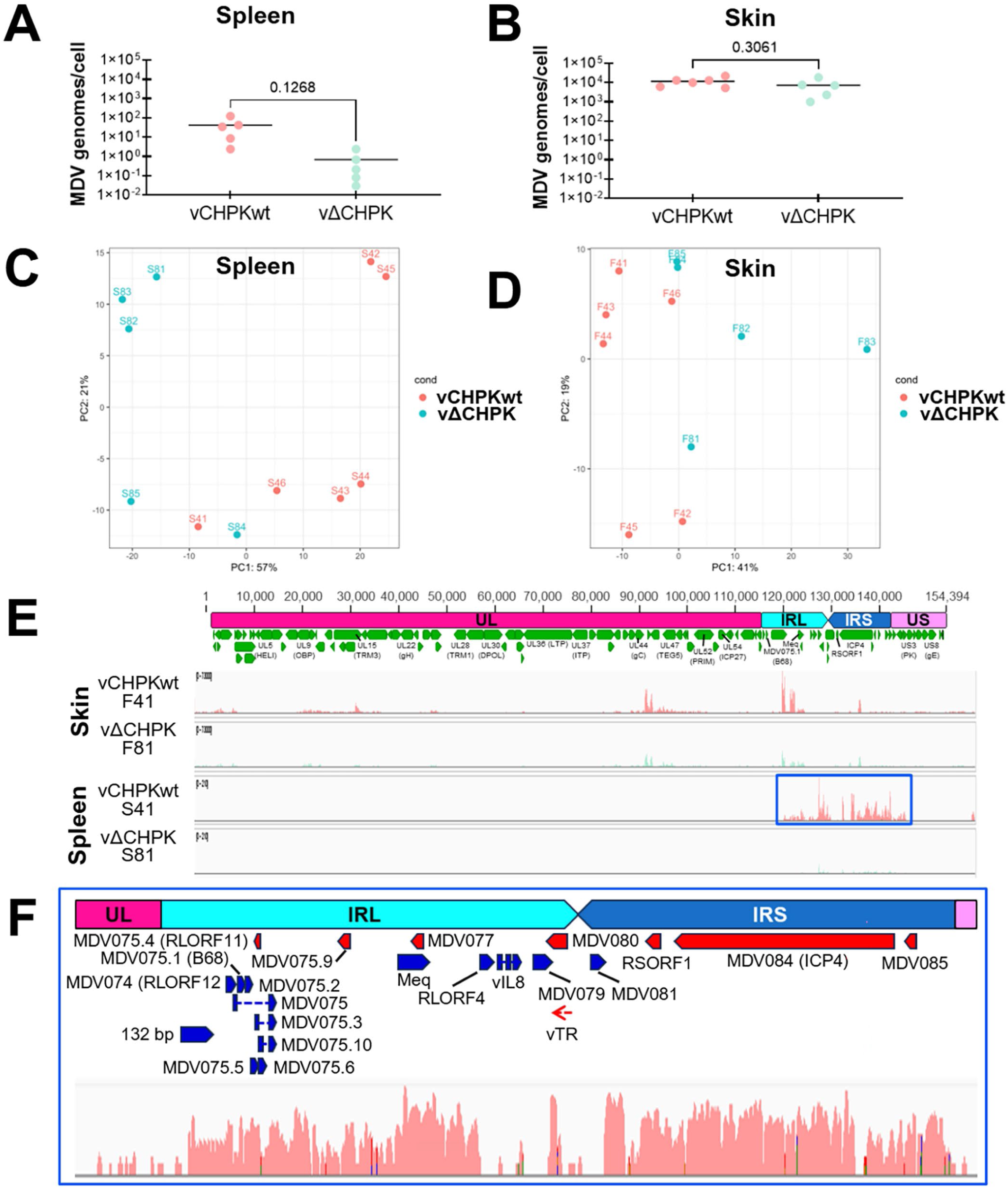
Virus replication and RNA expression in spleen and skin. (A-B) DNA extracted from RNA samples was used in qPCR assays to measure the relative viral genome copy number per cell in spleen (A) and skin (B) samples. No significant differences were determined between vCHPKwt- and vΔCHPK-infected spleen and skin samples using Student’s t-tests. (C-D) Principal component analysis of RNA-seq for spleen (C) and skin (D) samples. (E) Schematic representation of the MDV genome (TRL and TRS trimmed) and read depth of representative skin (F41, F81) and spleen (S41, S81) from RNA sequencing. The boxed blue region was amplified in F to better show read depth within the internal repeat long (IRL) and internal repeat short (IRS) regions in S81 spleen cells.

### Viral gene transcription in the spleen of infected chickens

Overall gene expression was higher in vCHPKwt-infected spleens, but both the vCHPKwt and vΔCHPK groups showed limited gene expression (Fig 2E). RNA expression in the spleen was restricted to the IRL and IRS regions (Fig 2F). Only 29 viral transcripts, out of 19,221 total transcripts, were detected across all samples (S1 File). These viral transcripts included many genes expected to be expressed during latency or transformation, consistent with the time during infection (>28 days PI). These included genes within the repeat regions [viral telomerase RNA (vTR), *MDV005/MDV076* (*RLORF7*, Meq), *MDV002/MDV079* (*RLORF1*, ICP0-like), *MDV084/MDV100 (*ICP4), *MDV083/MDV101* (anti-sense RNA protein), *MDV081/MDV103*, and *MDV003/MDV078* (vIL8v-CXC)]. This region is believed to be critical in maintaining latency and transformation.

Comparing transcripts from CHPKwt- and vΔCHPK-infected spleens, 4,393 genes were differentially expressed with p-adj < 0.05. More than 99% (4,386) of genes were from chicken, while only seven were viral (S3 Fig). These included *MDV002/MDV079* (*RLORF1*-ICP0-like), *MDV003/MDV078* (vCXC-IL8), *MDV005/MDV076* (*RLORF7*-Meq), *MDV010* (*LORF2*, vLIP), *MDV081/MDV103*, *MDV083/MDV101* (anti-sense RNA protein), and *MDV084/MDV100* (ICP4) were downregulated. Host gene responses have been previously reported [61]. These results are consistent with expectations for viral gene expression in the spleen during the latency/transformation phase of infection (>28 days pi).

### Viral gene transcription in the skin of infected chickens

Information on MDV-infected (vCHPKwt) vs control (Mock) in skin has been reported previously [60]. In that report, the topmost expressed viral genes (FDR <0.05 and FC ≤-2 or ≥2) were those involved in viral packaging [e.g., *MDV057*, *MDV057.2* (envelope glycoprotein C, gC), *MDV031* (major capsid protein, UL19), and *MDV069* (tegument protein VP11/12), etc.]. Comparing RNA-seq data from skin cells infected with wild-type (vCHPKwt) or CHPK-null (vΔCHPK) revealed key insights into CHPK’s role in viral gene expression. In contrast to the spleen, most of the viral genome was expressed, consistent with productive replication, with some regions of the genome bounteously transcribed (Fig 2E). For both vCHPKwt- and vΔCHPK-infected skins, *MDV057* (*UL44*, gC) and the region within the IRL encoding the complex 1.8 kb family of transcripts (*MDV075* and its subgenomic RNAs) were abundant. In the combined analysis of skin, 255 host and viral genes were differentially expressed between vΔCHPK- and vCHPK-infected skin at a p-adj < 0.05 (S2 File). Host gene responses have been previously reported [61]. Only two viral genes were downregulated: *MDV025* (*UL13*_CHPK) and *MDV027* (*UL15*_TRM3), the former likely due to 2/3 of the gene missing in vΔCHPK (S4 Fig). It is not known why *UL15* expression was reduced, but it could be due to the removal of regulatory regions for *UL15* transcription.

Overall, these data show that viral gene expression in the spleen was restricted to the repeat regions, whereas in the skin, most viral genes were expressed. In addition, there was little difference between vCHPKwt- and vΔCHPK chickens in both tissues when comparing viral gene expression.

### MS-based proteomic analysis of skin infected with MDV lacking CHPK

RNA-seq analysis revealed only minor changes in viral mRNA expression in the skin, corroborating previous RT-qPCR findings [20] and suggesting a minimal role for CHPK in modulating viral gene transcription. Analysis of global protein expression showed moderate differences in overall protein/peptide levels between the two groups (S3 File). Only 16 viral proteins showed significant differences (q < 0.100), with all but one exhibiting a change of less than 2-fold between the two groups, namely UL13_CHPK (21.95-fold). This large difference is likely due to the deletion of 2/3 of the UL13_CHPK protein in vΔCHPK. These results show that there is overall little difference (<2-fold) in total viral protein expression in the skin of chickens infected with vCHPKwt and vΔCHPK.

### CHPK targeted virus proteins

Next, we analyzed specific PTMs of viral peptides. Since CHPK is a protein kinase, we focused our analysis on differences in the phosphorylation status of specific viral peptides between vCHPKwt- and vΔCHPK-infected skins, following phospho-enrichment of protein samples by LC/MS-MS-based proteomics. For this analysis, a stringent filtering criterion was applied to ensure that, across all three biological replicates, each viral phosphorylated peptide was consistently present or absent when comparing vCHPKwt- and vΔCHPK-infected skin samples. Twenty-one viral proteins with unique phosphopeptides were found in all three vCHPKwt-infected skin cell samples but were absent from all replicates infected with vΔCHPK (Table 1, Fig 3). Of these 21 proteins, three are MDV-specific (MDV012_VG12, MDV069_VG69, and MDV071_VG71), while eight genes (UL3_NP03, UL44_gC, UL45_EV45, UL46_TEG1, UL47_TEG5, ICP4_ICP4, US10_US10, and US2_US02) are conserved within the *Alphaherpesvirinae* subfamily, and ten genes are common among the core *Orthoherpesviridae* family (UL6_PORTL, UL13_CHPK, UL14_TEG3, UL26_SCAF, UL30_DPOL, UL34_NEC2, UL36_LTP, UL39_RIR1, UL51_TEG7, and UL54_ICP27). Interestingly, one protein, UL39_RIR1, exhibited a unique phosphopeptide in both vCHPKwt- and vΔCHPK-infected cells, suggesting a distinct regulatory pattern. Furthermore, analysis based on protein function revealed that eleven of the differentially phosphorylated proteins are structural components like capsid/envelope/tegument proteins (UL6_PORTL, UL14_TEG3, UL26_SCAF, UL36_LTP, UL44_gC, UL45_EV45, UL46_TEG1, UL47_TEG5, UL51_TEG7, US2_US02, and US10_US10), while the remaining six are regulatory proteins (UL3_NP03, UL13_CHPK, UL30_DPOL, UL34_NEC2, UL39_RIR1, UL54_ICP27, and ICP4)

**Fig 3.**
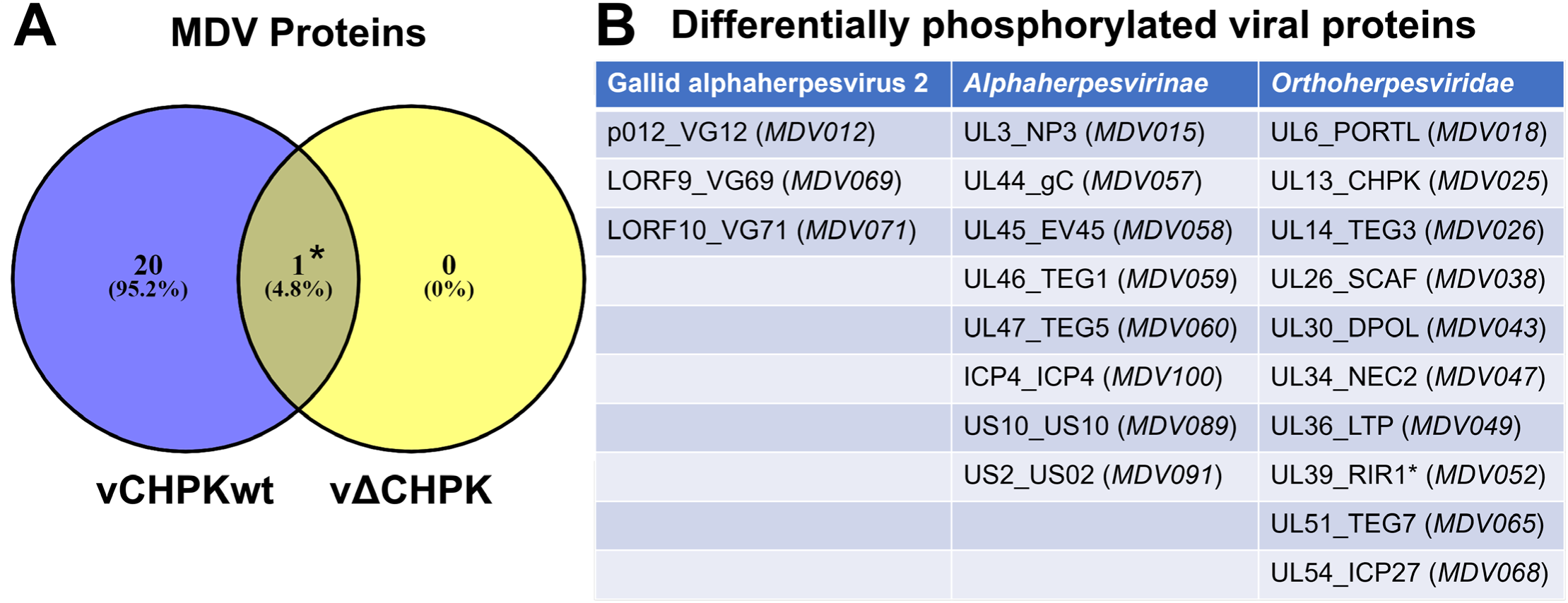
Differential phosphorylation of viral proteins in chicken skin. (A) Venn diagram of viral proteins uniquely phosphorylated between vCHPKwt- and vΔCHPK-infected epithelial skin cells. (B) Viral proteins that are differentially phosphorylated are separated by species, subfamily, and family.

**Table 1.**
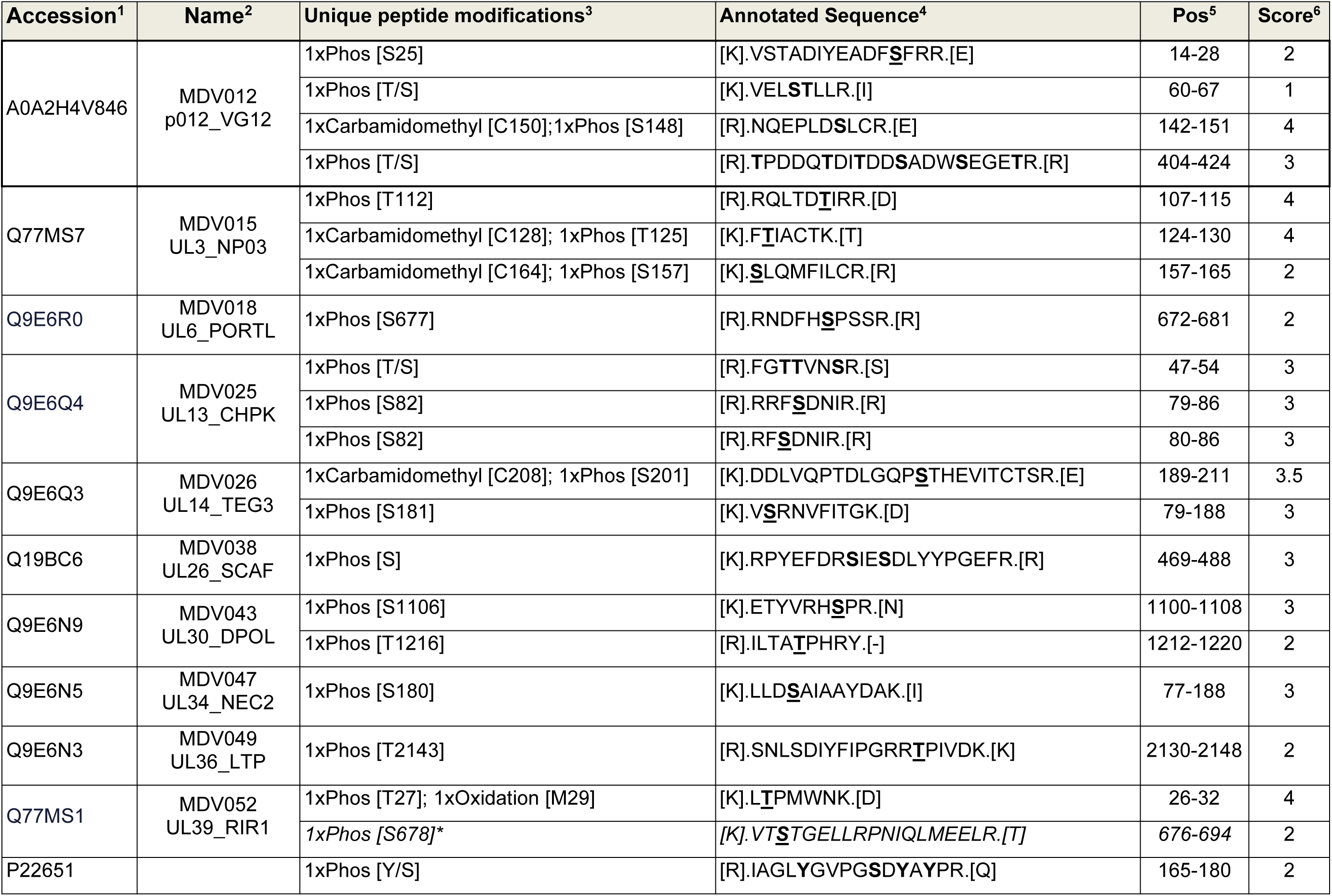
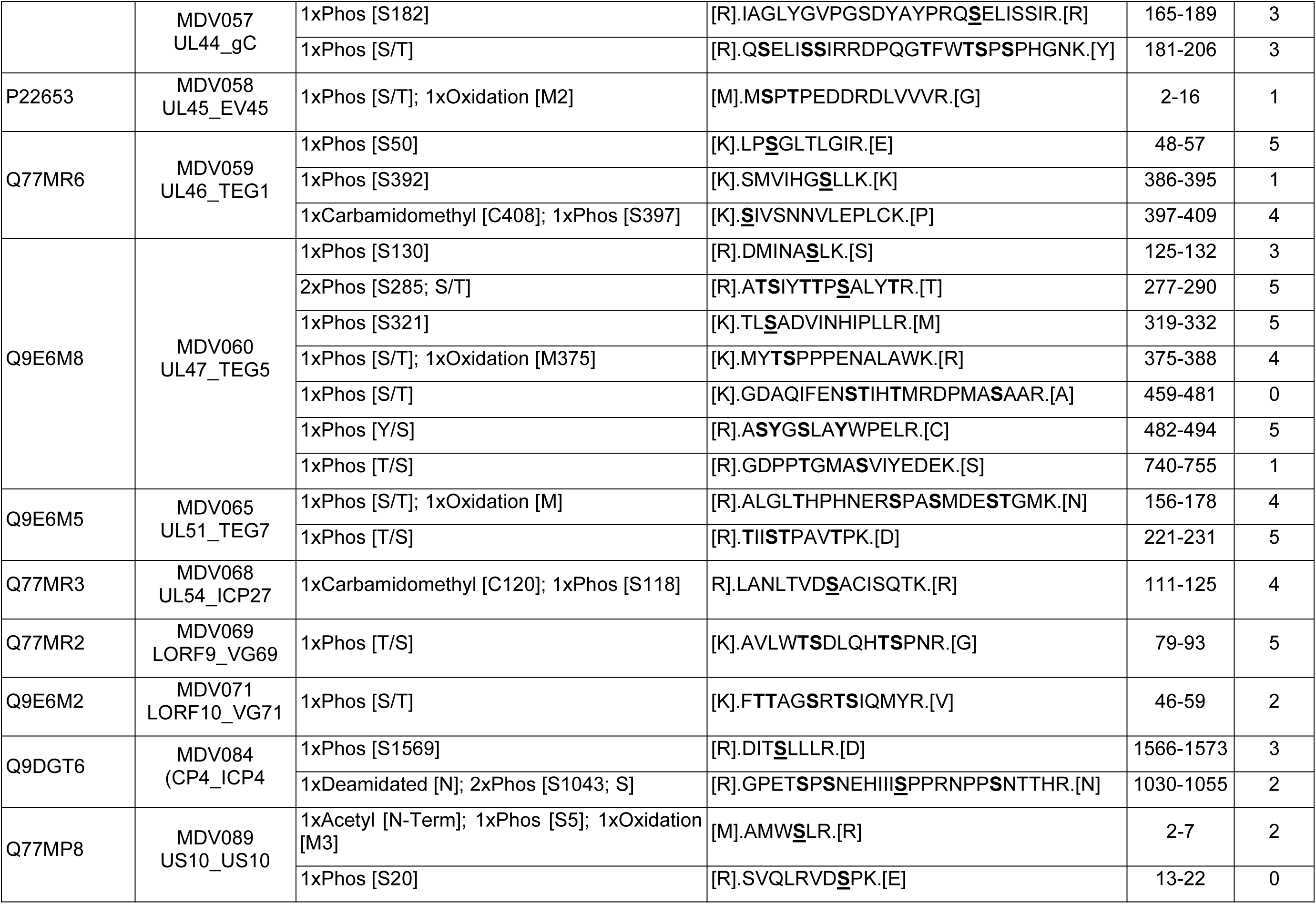
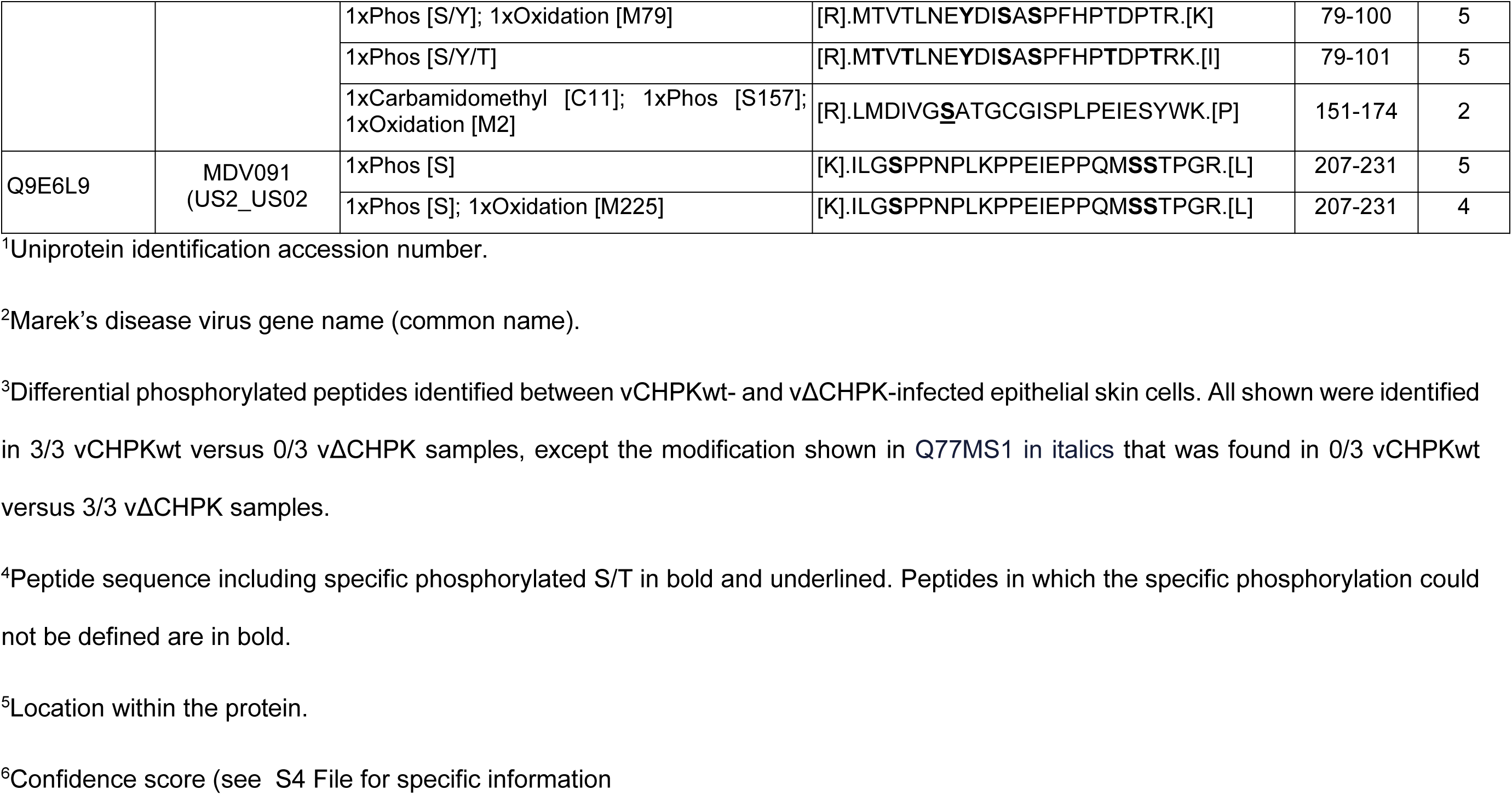
Differential phosphorylation of viral peptides in epithelial skin cells infected with vCHPKwt versus vΔCHPK.

Following initial analyses of the identified peptides, we further evaluated their relative confidence. using three criteria: 1) Is the peptide sequence ID correct? Here, we compared infected and mock samples to determine that the MS1 elution profile was unique to infected samples. 2) Is the unique peptide only found in vCHPKwt or vΔCHPK? Here, we determined the presence or absence based on extracted ion chromatograms of varying clarity. 3) Is the PTM localized to the correct residue on the peptide, where there are multiple possibilities? Rich MS2 spectra with ion peaks from fragmentation on both sides of the phosphosite are needed to do this. Some of the MS2 spectra were insufficient to define the exact peptide sequence, as noted below, but others were sufficiently rich to be confidently identified. The relative confidence level for each peptide is shown in Table 1. Specific peptide profiles are available in the S4 File, and MS2 spectra of unique phosphopeptides are shown in the S5 File.

### Differential phosphorylation of viral proteins near nuclear import and export signals

In previous studies, we observed dysregulated expression and subcellular localization of late MDV proteins during replication in cell culture compared to epithelial skin cells in chickens, including UL46_TEG1, UL47_TEG5, UL48_VP16, US10_US10, and UL54_ICP27 [20, 36, 45, 59, 62]. For example, UL47_TEG5 and UL54_ICP27 are barely detectable and localized to nuclear puncta in cell culture, whereas they are abundantly expressed and nucleocytoplasmic in skin cells [59, 62]. Notably, cell culture models do not support the production of infectious cell-free MDV particles, whereas epithelial skin cells do, linking these late viral proteins to cell-free virion production required for horizontal transmission and natural infection. The underlying mechanisms are unknown; however, we hypothesize that CHPK regulates the expression and subcellular localization of specific viral proteins in these cells.

Phosphorylation events, particularly at or near NLS sequences, have previously been shown to influence the subcellular localization of UL47_TEG5 and UL54_ICP27 homologs [63–65]. Using multiple *in silico* programs, putative NLS and NES motifs were predicted for the 21 differentially phosphorylated viral proteins (see Materials and Methods), yielding NLS and NES motifs in 15 and 12 viral proteins, respectively. Notably, MDV_gC and UL45_EV45 lacked predicted NLS or NES, consistent with their membrane localization via transmembrane domains [66, 67]. Importantly, many of the unique phosphorylated peptides were within or near putative localization motifs, as discussed below. These data suggest CHPK-mediated phosphorylation within or near NLS and NES motifs may be critical for late viral protein expression, localization, and function within cells. The following is a summary of unique peptides detected in our analysis, putative NLS and NES motifs present in these select MDV proteins, and current data on herpesvirus orthologs with respect to their phosphorylation state and cellular localization.

### Differentially phosphorylated MDV-specific proteins

#### LORF2_VG12 (*MDV012*)

LORF2_VG12 (*MDV012*) is conserved among avian herpesviruses. Cell culture studies have demonstrated that the encoded protein LORF2_VG12 is phosphorylated, is essential for replication, and contains a functional NLS with numerous serine and tyrosine residues [68]. However, this study did not investigate the role of phosphorylation in subcellular localization. Here, we identified four unique peptides phosphorylated in vCHPKwt-infected cells that were not detected in vΔCHPK-infected cells (Table 1, Fig 4). Two of these peptides have identifiable phosphorylated residues (S25 and S148), whereas specific residues could not be identified for the remaining two peptides from their MS spectra. The peptide S[R].NQEPLD**<u>S</u>**LCR.[E] containing S148 was detected at a high (score of 4) confidence level (Table 1), while all others were low to moderate (score of ≤ 3 score) based on our confidence criteria (S4 File).

**Fig 4.**
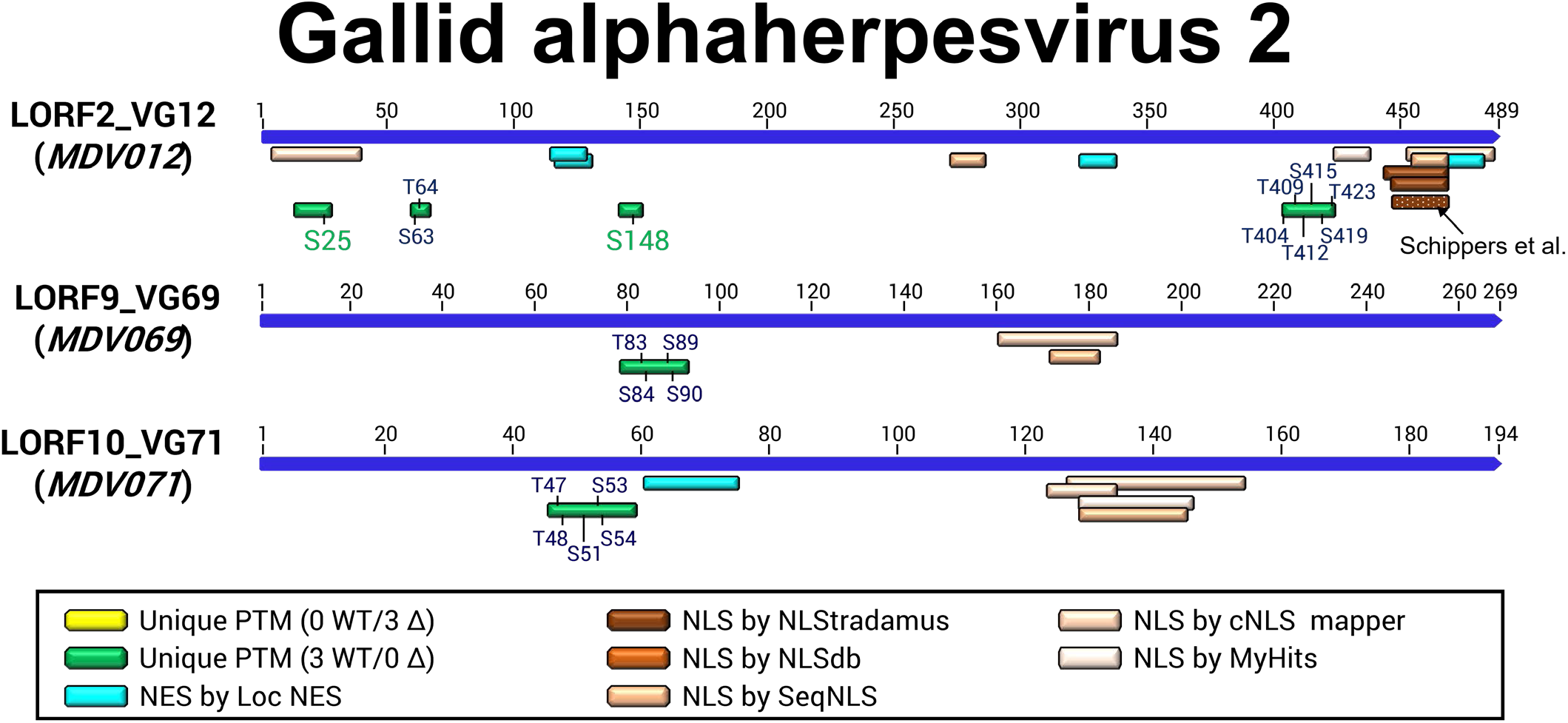
Putative NLS and NES motifs for gallid alphaherpesvirus 2 (MDV) differentially phosphorylated proteins in skins. The three MDV-specific proteins were found to harbor differentially phosphorylated peptides between vCHPKwt- and vΔCHPK-infected skins. Specific phosphorylated residues are shown in green, while potential phosphorylated residues for which the specific residue could not be determined in the peptide are shown in blue. Multiple programs were used to predict NLS or NES sequences (see Materials and Methods). The previously identified NLS in LORF2_VG12 by Schippers *et al.* [68] is shown.

Numerous NLS hits across multiple programs predicted that the C-terminus contained a strong NLS motif (Fig 4), consistent with the previously identified functional NLS [68]. Importantly, a unique peptide containing phosphorylated residues mapped near this region contained a potential CHPK basic motif (^404^TPDDQTDITDDSADWSEGETR^424^) with a moderate confidence score of 3 (Table 1, S4 File). *In silico* analysis using MyHits also predicted a putative NLS at the N-terminus, aligning with S25 identified in our MS-based analysis. Furthermore, LocNES identified four potential NES sequences, with one having a high score of 0.577 (^117^ER**<u>L</u>**ATSW**<u>L</u>**TA**I**R**<u>LIL</u>**^131^). The unique phosphorylated residue S148 is <20 aa from this high-scoring putative NES site. The functionality of these potential NES motifs and the role of CHPK-mediated phosphorylation of LORF2_VG12 in its localization remain to be elucidated *in vivo*. Nuclear export was not addressed previously [68].

#### LORF9_VG69 (*MDV069*) and LORF10_VG71 (*MDV071*)

In contrast to the LORF2_VG12 (*MDV012*), which exhibits conservation across avian alphaherpesviruses [68], LORF9_VG69 (*MDV069*) and LORF10_VG71 (*MDV071*) are found exclusively encoded by MDV. Interestingly, our analysis identified peptides corresponding to the translation products of MDV-specific genes (Table 1, Fig 4). The unique phosphopeptide detected for LORF9_VG69 scored high (score of 5) in confidence, but the exact phosphoresidue could not be determined from the MS2 chromatogram. For LORF10_VG71, the identified phosphopeptide scored low to moderate confidence (score of 2). Thus, further validation is necessary to determine the relevance of this result.

*In silico* analyses using NucPred [69] predicted a low nuclear localization score (0.04) for LORF9_VG69, while LORF10_VG71 exhibited a more neutral score (0.40). However, both proteins possess putative NLS motifs, identified using multiple programs. For LORF9_VG69, both cNLS mapper and SeqNLS identified overlapping putative NLS between aa 116-186,^161^PGI**K**PALLMDP**RR**FLEM**R**DP**RK**IICL^186^ (score 4.0) and ^172^**RR**FLEM**R**DP**RK**^182^ (score 0.433), respectively. An NLS region within aa 124-154 of LORF10_VG71 was predicted by three programs: cNLS mapper predicted two (^127^PP**KR**YHKYHDSSLG**RRR**GISVDRSANTA^154^, score of 3.1; ^124^LNSPP**KR**Y^134^, score of 3.5), while SeqNLS and MyHits predicted similar sequences (^129^**KR**YHKYHDSSLG**RRR**GI^145^, score 0.924) and (^129^**KR**YHKYHDSSLG**RRR**GIS^146^), respectively. A putative NES (^87^GRARTDG**L**TRE**FVIL**^75^) was also predicted by MyHits, but the score was relatively low (0.129). Importantly, the unique phosphopeptide identified in vCHPKwt-and absent in vΔCHPK-infected skin resides near this putative NES. Both LORF9_VG69 and LOR10_VG71 are nonessential for viral replication *in vitro*. However, deletion of *MDV069* (LORF9_VG69) resulted in attenuated MDV *in vivo* [70]. While no known homologs exist for LORF9_VG69, LORF10_VG71 reportedly shares homology with VZV ORF2 [71].

### Differentially phosphorylated Alphaherpesvirus-specific proteins

#### UL3_NP03 (*MDV015*)

UL3_NP03 is conserved among the *Alphaherpesvirinae* and is nonessential for cell-culture replication [72]. While its major function is currently unknown, previous studies have identified UL3_NP03 as a nucleocytoplasmic phosphoprotein [73], and both NLS and NES regions have been formally defined for the related HSV-1 [74]. Our study identified three peptides with unique phosphorylated residues: two with high scores of 4 and one with a low-to-moderate (score of 2) confidence score (Table 1, Fig 5). Two of these peptides reside within or near putative NLS and NES motifs (S5 Fig). Interestingly, UL3_NP03 shows high sequence conservation between HSV- 1 and MDV with 45.9% total amino acid identity. In other phosphoproteomic studies, HSV UL3_NP03 S125 was shown to be phosphorylated during treatment with the DNA inhibitor phosphonoacetic acid (PAA). Moreover, the defined NLS and NES regions of HSV-1 UL3_NP03 are conserved in MDV UL3_NP03 with 100% and 90% identity, respectively. The putative NES motifs align with two uniquely phosphorylated peptides identified in our phosphoproteomic analysis. It is tempting to speculate that CHPK-mediated phosphorylation of UL3_NP03 T112 and T125 may play an important role in localization or protein-protein interactions in epithelial skin cells. However, further investigation is necessary to definitively establish the functional significance of the observations.

**Fig 5.**
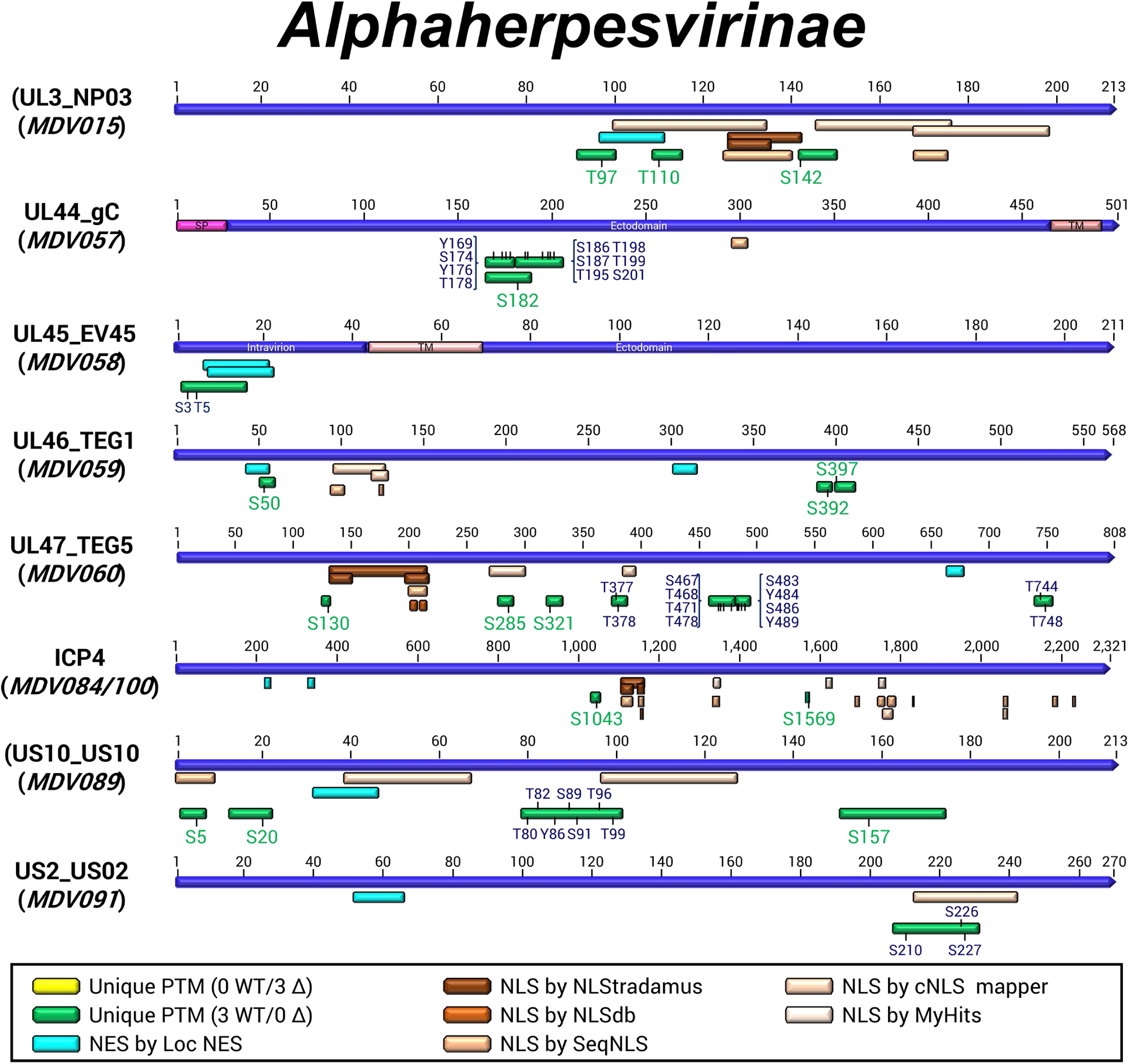
Putative NLS and NES motifs for differentially phosphorylated proteins conserved among the *Alphaherpesvirinae*. Eight proteins conserved among the alphaherpesviruses were found to have differentially phosphorylated peptides between vCHPKwt- and vΔCHPK-infected skins. Specific phosphorylated residues are shown in green, while potential phosphorylated residues for which the specific residue could not be determined in the peptide are shown in blue. Multiple programs were used to predict NLS or NES sequences (see Materials and Methods).

#### UL44_gC (*MDV057*)

Members of the *Alphaherpesvirinae* subfamily encode homologs of glycoprotein C (gC). These heavily glycosylated type 1 membrane proteins play multifaceted roles in the virus’s life cycle, including attachment of cell-free viruses to heparin- and chondroitin sulfate proteoglycans on the surface of cells [75, 76], the final stages of virus egress from infected cultured cells [75, 77], release from the cells [78], and immune evasion functions through binding and inhibiting the complement component C3 [79–81] and chemokine-mediated leukocyte migration for VZV [82]. Interestingly, while gC homologs of *Mardiviruses* are dispensable for *in vitro* and *in vivo* replication during experimental infection, gC of MDV, GaAHV3, and herpesvirus of turkeys (HVT) is crucial for host-to-host transmission or natural infection [37, 83, 84]. This parallels the situation with HSV-1 and VZV, where gC homologs play only a minor role in tissue culture model systems but are critical for replication in human skin cells, as demonstrated in the SCID-hu mouse model [85].

No prior reports have documented the phosphorylation of gC homologs during infection. However, our differential phosphorylation analysis detected three phosphopeptides, but all three had low-to-moderate (scores of 2-3) confidence scores (Table 1, Fig 5). The functional significance of gC phosphorylation during skin replication remains unclear. Nonetheless, detecting differential phosphorylation between vCHPKwt- and vΔCHPK-infected cells is intriguing, considering that both gC and CHPK are required for transmission [37]. This finding represents the first suggestive evidence that their roles in natural infection might converge at the mechanistic level. Further studies are warranted to explore this possibility.

#### UL45_EV45 (*MDV058*)

UL45_EV45 is also a conserved component of the virion particle within members of the *Alphaherpesvirinae* subfamily [86]. HSV-1 UL45_EV45 is expressed as a late gene and functions as a type II membrane protein, dispensable for replication in cell culture [66, 87]. While it interacts with gB and is required for cell-to-cell fusion induced by gB variant syncytial viruses [88], its precise function during herpesvirus replication remains unknown. Homology modeling suggests a C-type lectin fold region in UL45_EV45 homologs [89]. Our MS-based phosphoproteome analysis identified a unique peptide at the N-terminus of MDV UL45_EV45, though this had a low confidence score of 1.

*In silico* analysis predicted two NES with scores of 0.699 (^7^EDDRD**LVVV**RGR**L**R**M**^21^) and 0.480 (^8^DDRD**LVVV**RGR**L**R**MM**^22^) at the N-terminus (Fig 5). However, since these overlapping peptides align with the predicted intra-virion or intra-cellular domain based on HSV-1 UL45_EV45 homology, the functionality of these NES motifs remains uncertain. NLS or NES motifs have not been defined for other alphaherpesvirus UL45_EV45 homologs. Given the low confidence score for this unique phosphopeptide and the low likelihood of nucleocytoplasmic transport of this envelope protein, this result is likely insignificant.

#### UL46_TEG1 (*MDV059*)

UL46_TEG1, also known as VP11/12, is encoded by *UL46* (*MDV059*), another late gene conserved among the *Alphaherpesvirinae*. It is a major tegument protein dispensable for MDV and other alphaherpesviruses in cell culture [90–92]. While its role in the MDV lifecycle remains unclear and underexplored, UL46_TEG1 homologs in HSV-1 and PRV are abundant tegument proteins, found in complexes with UL47_TEG5, UL48_VP16, and UL49_VP22 [93–95], and are heavily phosphorylated by both viral and cellular kinases [96]. Phosphorylation of UL46_TEG1 is mediated by both US3 [97] and CHPK [98]. The localization of UL46_TEG1 homologs is predominantly cytoplasmic or perinuclear [99], including in MDV [45]. However, they are also detected in the nuclei of infected cells during infection [20, 100]. Xu *et al.* [100] identified a potential NLS in PRV UL46_TEG1 at aa 3-18 (**RR**A**R**GT**RR**ASW**K**DAS**R**) and a potential NES at the N-terminus, though the NES is, at best, a non-classical NES between aa 19-48. Putative NLS or NES sequences have not been described for HSV UL46_TEG1 to date. Interestingly, amino acid sequence alignments reveal limited homology between PRV and MDV UL46_TEG1 in the NLS and NES regions (S6 Fig). However, *in silico* analysis predicted numerous NLS motifs in MDV UL46_TEG, along with two NES motifs, the first of which is positioned near the proposed PRV NES.

Our phosphoproteomic analysis identified three unique phosphopeptides in vCHPKwt-infected skin that were absent in vΔCHPK-infected skin. Two phosphopeptides were scored with high confidence (≥4), while the third was scored low at 1. Of the two high-scoring phosphopeptides, the S50 residue resides within the first putative NES and is the phosphorylated residue containing the UL13 (CHPK) basic “PS” motif [101]. Cell culture and epithelial skin cell infections have shown that MDV UL46_TEG1 expression differs considerably between the two cell types. Modification of S50 in UL46_TEG1, potentially via CHPK, may explain the regulation of this protein. Further studies are needed to identify functional NLS and NES motifs and better understand whether CHPK directly regulates the expression and localization of MDV UL46_TEG1.

#### UL47_TEG5 (*MDV060*)

Much work has been conducted on HSV-1 UL47_TEG5, elucidating its functions and the phosphorylation-mediated localization during replication. This protein is a nucleocytoplasmic shuttling protein [64] and is essential for MDV horizontal transmission in chickens [102]. For HSV-1, defined NLS and NES sequences dictate its subcellular location, which is regulated by US3-mediated phosphorylation in cell culture [57]. Specifically, S77 of HSV-1 UL47_TEG5 is important for its localization and overall virus replication, and this residue is near its defined NLS at aa 63- 75 (S7 Fig). HSV US3, a PKA-like kinase (RRXS*/T*), has been shown to phosphorylate UL47_TEG5 at S77, thereby directing its nuclear translocation. While the MDV S130 sequence deviates from the PKA consensus motif, we identified phosphorylation on S151 ([R].NA**<u>S</u>**MHMHFR.[G]) that was phosphorylated in both vCHPKwt and vΔCHPK-infected samples. This is consistent with potential US3-mediated phosphorylation.

Our analysis of MDV UL47_TEG5 predicted numerous potential NLS motifs, some of which aligned well with those identified in the known HSV-1 UL47_TEG5. Interestingly, our proteomics analysis identified a unique peptide in vCHPKwt-infected skin with S130 phosphorylation in MDV UL47_TEG5 located near a putative NLS. This phosphopeptide scored moderate confidence (score of 3). Furthermore, additional phosphopeptides were detected, including high-confidence (score of ≥4) phosphopeptides at positions 277-290, 319-332, 375-388, and 482-494 within or near predicted NLS or NES motifs. Two studies have investigated the export of HSV-1 UL47_TEG5 from the nucleus via two NES sequences, one of which was identified by Williams *et al.* [103] as a CRM1-dependent NES, while Funk *et al.* [104] discovered another NES in the protein (S7 Fig). Similarly, analysis using the LocNES software predicted two NES for MDV UL47_TEG5, one that aligns well with the NES described by Funk *et al.* [104] for HSV-1, while the other resides further towards the C-terminus.

Previous studies have shown that the expression and localization of MDV UL47_TEG5 differ considerably between replication in cell culture and in epithelial skin cells of infected chickens [59]. That is, UL47_TEG5 is barely detectable and localizes primarily to the nucleus in cell culture, whereas in FFE skin cells, it localizes to both the nucleus and the cytoplasm during the productive MDV replication cycle. This differential expression and localization of MDV UL47_TEG5 correlates with the cell-associated vs cell-free stages of MDV infection, suggesting UL47_TEG5 subcellular trafficking plays a functional role in the assembly and release of enveloped cell-free virus (CFV). Thus, it is important to pinpoint the exact NLS and NES motifs in MDV UL47_TEG5 to better understand how phosphorylation within or near these regions regulates its localization/function. Ongoing studies are addressing these questions, and a more detailed analysis of MDV UL47_TEG5 localization is in preparation for future publications.

#### ICP4_ICP4 (*MDV084/100*)

ICP4 is the major regulatory protein for all Alphaherpesviruses. MDV ICP4 predominantly localizes to the nucleus and, to a lesser extent, the cytoplasm [45]. ICP4 homologs exhibit functional complexity and contain numerous domains. *In silico* analyses of MDV ICP4 revealed a high nuclear score (0.96) using NucPred [69], consistent with its known role in the nucleus, where it regulates the transcriptional activity of viral genes. In studies focusing on HSV-1 ICP4, Mullen *et al.* [105] defined ^726^GRKRKSP^732^ as the NLS sequence. *In silico* analysis of the MDV ICP4 protein revealed >20 predicted NLS and two NES sequences. (Fig 5) with some NLS overlapping with the previously described NLS region in HSV-1 ICP4 (S8 Fig). However, none appeared to be in close proximity to unique phosphopeptides in our phosphoproteomic analysis.

HSV-1 ICP4 is phosphorylated on Ser and Thr residues, especially in the serine-rich region, which contains multiple phosphorylation sites [106]. The current model proposes that, following viral infection, ICP4 is first phosphorylated by kinases within its serine-rich region and then performs multiple functions during infection, including regulating the transcriptional activity of immediate-early (IE), early (E), and late (L) genes [107]. Some kinases shown to phosphorylate ICP4 include PKA, PKC, or others [106, 108, 109]. It has been suggested that US3_PK or UL13_CHPK may phosphorylate ICP4, but no direct evidence has supported this idea [110]. In our proteomics analysis, two phosphopeptides with moderate confidence (scores of 2-3) were detected only in vCHPKwt-infected cells, including one near the serine-rich region of MDV ICP4 and just upstream of predicted NLSs (S8 Fig). Although our data here do not directly show that UL13_CHPK phosphorylates ICP4, our phosphoproteomic studies suggest it may. Thus, at least some of the phosphorylation of ICP4 may be mediated by CHPK, at least in the skin, since the non-phosphopeptide was found in the vΔCHPK samples.

#### US10_US10 (*MDV089*)

US10 is a virion protein found in the virus’s tegument and is conserved among the alphaherpesviruses [111]. HSV-1 US10 is dispensable for cell culture replication [112, 113]. Similarly, Ponnuraj *et al.* [36] showed that US10 of MDV is dispensable for cell culture replication, experimental infection, and transmission in chickens, but plays a role in disease induction during natural infection. This report also showed that UL13_CHPK and US10 expression and subcellular localization were linked, with their expression dependent on each other and co-localizing in the cytoplasm and nucleus of cells. Importantly, when UL13_CHPK was absent during infection (vΔCHPK), US10 expression was almost completely abrogated. Conversely, when US10 was absent (vΔUS10), CHPK was almost exclusively localized to the nucleus, implicating that they are both involved in each other’s expression or localization in cells during infection.

Although potential NLS or NES sites in both proteins were not addressed in that report, we now show that MDV US10 harbors three predicted NLS sequences and one predicted NES sequence. A few reports of computer-predicted NLS and NES sites for HSV-1 US10 have been published. Funk *et al.* [104] predicted an NES for HSV-1, as shown in S9 Fig, but, to our knowledge, the regions responsible for protein movement have not been reported. Previous research on HSV-1 US10 suggests it resides primarily within the nucleus in transient transfection assays, where HSV-1 US10 was fused to YFP [114], and a separate study using the Nuclear EXport Trapped by RAPamycin (NEX-TRAP) system, HSV-1 US10 remained in the nucleus, suggesting it lacked nuclear export [104]. In contrast, MDV US10 fused with eGFP appears mainly cytoplasmic during *in vitro* infection [115] and in transfected cells [36].

To further assess the potential role of NLS or NES in MDV US10 subcellular localization, transient transfection assays were performed using the US10eGFP expression construct previously described [36] in the absence or presence of leptomycin B (LMB), which disrupts classical NES function, and cycloheximide (CHX) to block translation. Consistent with prior findings [36, 115], US10 was mostly cytoplasmic with large puncta staining (Fig 6). However, treating cells with LMB led to a broader distribution of US10, with increased nuclear accumulation. These results suggest that US10 may have a functional NES that controls its nuclear export.

**Fig 6.**
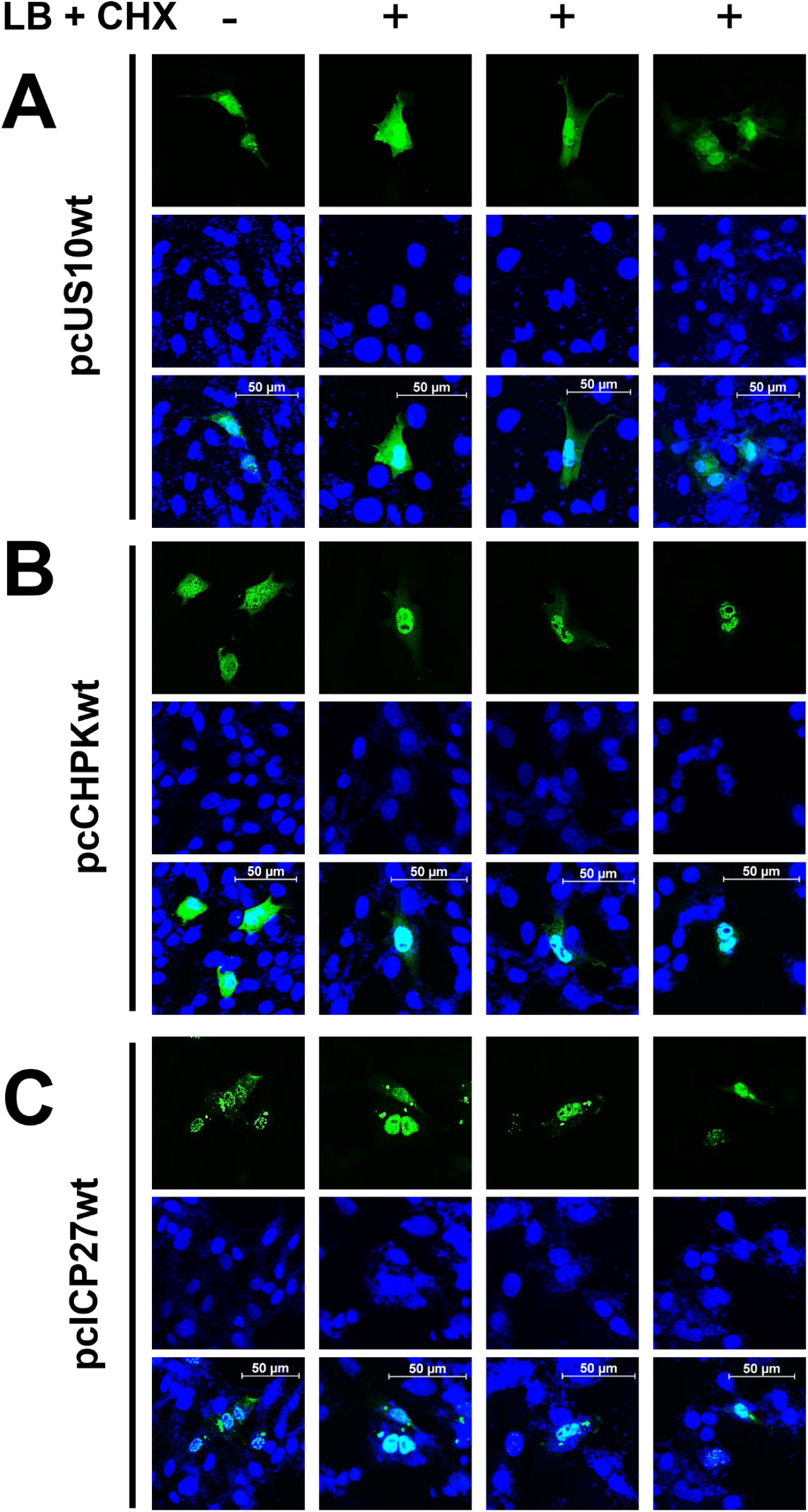
Subcellular localization of MDV US10, CHPK, and ICP27 in cell culture. DF-1 cells were transfected with US10 (A), CHPK (B), or ICP27 (C) expression constructs, then treated with leptomycin B (LMB) and cycloheximide (CHX). Localization of each protein was evaluated using confocal microscopy using direct or indirect fluorescence assays. Untreated (n=1) and treated (n=3) are shown for each.

To test the functionality of the predicted NLS and NES in MDV US10, deletion mutants were generated in the US10-eGFP expression construct [36]. Based on *in silico* analysis, three NLS and one NES were predicted for MDV US10. These regions were removed to create pcUS10ΔNLS1 (aa Δ2-9), pcUS10ΔNLS2 (aa Δ39-67), pcUS10ΔNLS3 (aa Δ97-127), and pcUS10ΔNES (aa Δ32-46), shown in Fig 7A. Following transient transfection in immortalized chicken DF-1 cells, there were no apparent changes in US10-eGFP localization.These data suggest that these predicted sites are not functional, at least in the context of transient transfections in cell culture.

**Fig 7.**
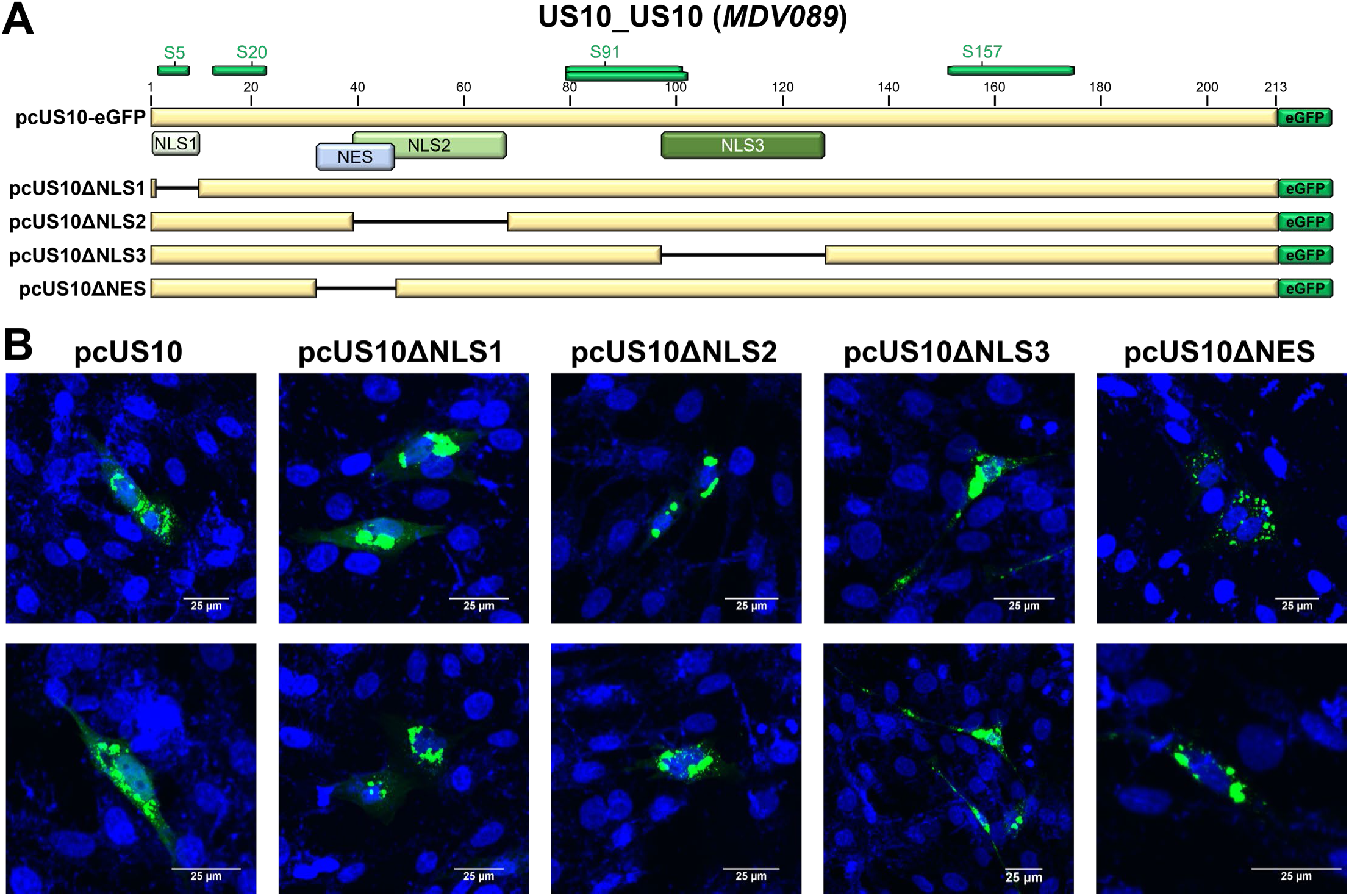
Subcellular localization of MDV US10 NLS and NES mutants. Putative NLS and NES motifs (A) were deleted in the US10-eGFP (pcUS10) expression construct using site-directed mutagenesis, including NLS1 (pcUS10ΔNLS1), NLS2 (pcUS10ΔNLS2), NLS3 (pcUS10ΔN3), and NES (pcUS10ΔNES). (B) Chicken DF-1 cells were transfected, and after 24 h, fixed with 4% PFA and stained with Hoechst 33342 to visualize nuclei. Images were collected using confocal microscopy. Shown are two representative images for each construct. No visual differences in cell localization were observed between pcUS10 and the deletion mutants.

Based on our previous studies, the localization of US10 within the cell likely depends on its interactions with UL13_CHPK and possibly UL55_TEG6. MDV UL13_CHPK and US10 expression and subcellular localization are linked [36]. Thus, when UL13_CHPK was absent during infection (vΔCHPK), US10 expression was completely abrogated. Conversely, when US10 was absent (vΔUS10), CHPK was almost exclusively localized to the nucleus. Recently, it has been demonstrated that HSV-1 UL13_CHPK, US10, and UL55_TEG6 interact and regulate CHPK kinase activity in cell culture [116], suggesting more complex interactions may mediate US10 subcellular localiation. Further studies on MDV UL13_CHPK and UL55_TEG6 are needed to address their potential contribution to US10 localiation and expression in cells.

### US2_US02 (*MDV091*)

US2 is nonessential for cell-culture propagation of all herpesviruses studied but may play a role in virulence during natural infections [38, 117–120]. While most US2 homologs studied localize to the cell and viral membranes [121–123], HSV and duck enteritis virus (DEV) US2 were observed in the cytoplasm, distributed into bright granules near the nucleus during infection [124, 125]. Interestingly, HSV-2 US2 appears solely cytoplasmic, lacking concentrated granules in Vero cells transfected with an expression vector [125]. MDV US2 has a low nuclear prediction score of 0.10 using NucPred [69], consistent with its homologs being localized outside the nucleus. No studies have directly shown that US2 is exported from or imported into the nucleus during infection. A study by Funk *et al.* [104], using the NEX-TRAP assay, suggests that HSV-1 US2 may be exported from the nucleus. The *in silico* analysis of the MDV US2 protein sequence revealed a potential NLS (^213^NPL**<u>K</u>**PPEIEPPQMSSTPG**<u>R</u>**LFCCGKCC**<u>KK</u>**E^242^) and NES (^52^RAAD**<u>LF</u>**R**<u>F</u>**AKP**<u>MLIL</u>**^66^). Notably, two unique phosphopeptides with high confidence (scores ≥ 4) were detected in our proteomics study of vCHPK-infected skin and overlapped with the predicted NLS (Table 1, Fig 5). More research is required to determine whether these putative NLS and NES motifs are functional and whether phosphorylation contributes to US2’s overall function.

### Differentially phosphorylated *Orthoherpesviridae* proteins

#### UL6_PORTL (*MDV018*)

The HSV-1 procapsid comprises the major capsid protein (VP5; UL19_MCP), the triplex proteins (VP19C; UL38_TRX1 and VP23; UL18_TRX2), and the portal ring (UL6_PORTL). UL6_PORTL is essential for viral replication and has been shown to localize to the nucleus [126, 127], but no defined NLS has been described. Cai *et al.* [128] described HSV-1 UL6_PORTL transport to the nucleus through Ran-, transportin-, or importin-dependent nuclear import mechanisms but did not define an NLS. Our analysis identified a unique phosphopeptide with low confidence (score of 2) that maps near a predicted NLS in MDV UL6_PORTL (Fig 8). This region is rich in arginine or lysine residues and contains the HSV-1 CHPK consensus sequence (SP/PS). Interestingly, HSV-1 also contains a similar arginine-rich motif (^671^**<u>RR</u>**DG**<u>RR</u>**^676^) at its C-terminus (S10 Fig). While details on UL6_PORTL phosphorylation are limited, Bell *et al.* [129] reported residue S459 phosphorylation in HSV-infected cells treated with acyclovir. Given the low confidence score of 2 for the single phosphopeptide detected in vCHPKwt but not in vΔCHPK-infected skin cells, further studies are required to determine its importance during infection and UL6_PORTL subcellular localization.

**Fig 8.**
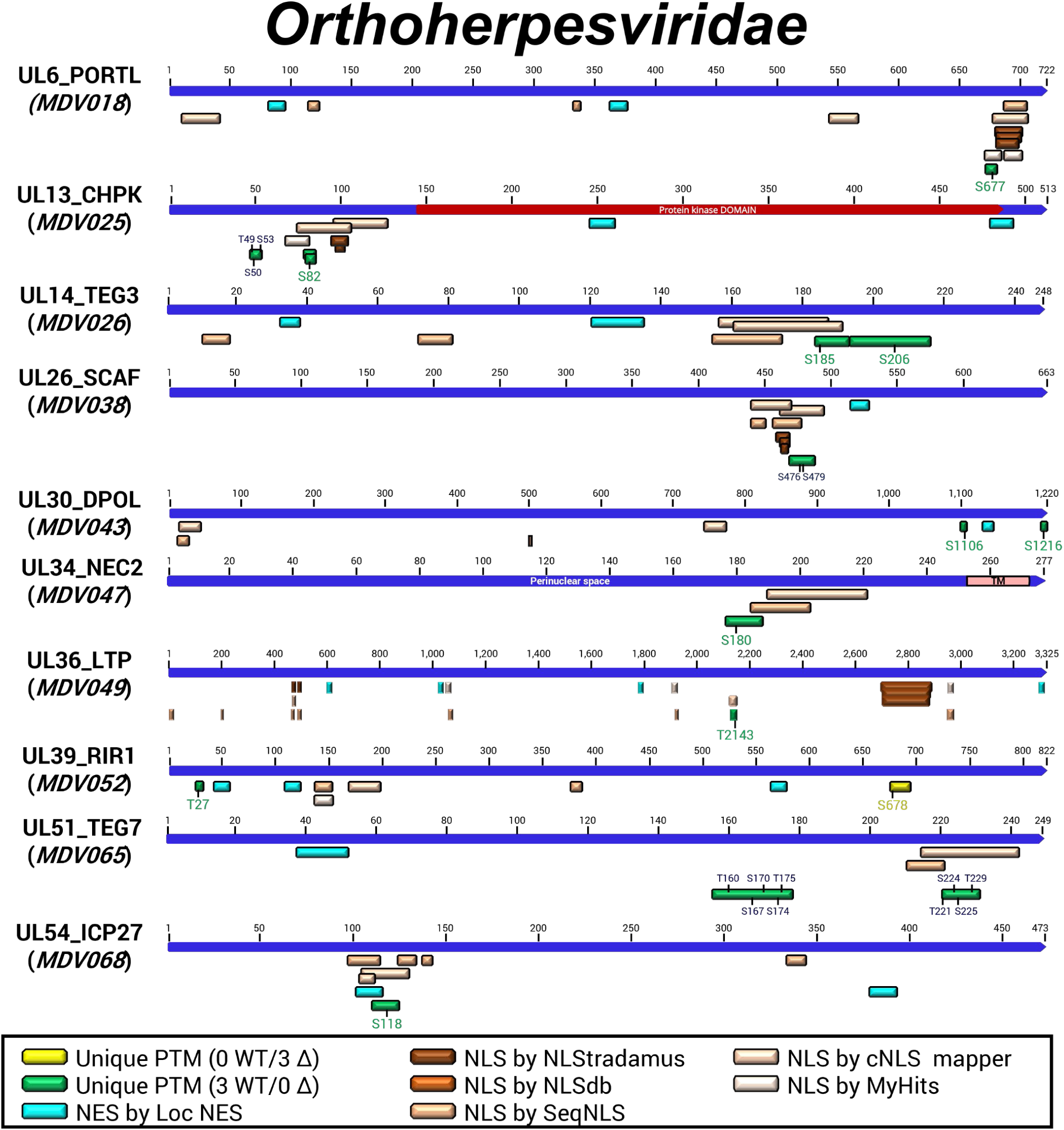
Putative NLS and NES motifs for differentially phosphorylated proteins are conserved among the *Orthoherpesviridae*. Ten proteins conserved in the *Orthoherpesviridae* were found to have differentially phosphorylated peptides between vCHPKwt- and vΔCHPK-infected skins. Specific residue phosphorylated residues are shown in green, while peptides for which the specific residue could not be determined are shown in blue. Multiple programs were used to predict NLS or NES sequences (see Materials and Methods).

#### UL13_CHPK (*MDV025*)

UL13_CHPK is known to autophosphorylate itself, and multiple studies have identified phosphorylated residues, including HSV-1 and HSV-2 [130, 131]. Koyanagi *et al.* [130] identified S11, S18, S47, S50, S90, and S91 phosphorylated in HSV-2 infected cells, while Bell *et al.* [129] identified S18, S91, S109, and S119 phosphorylated in HSV-1 infected cells. Our comparative phosphoproteomic analysis identified six phosphopeptides in the vCHPKwt-infected cells. Three phosphopeptides in our comparison were mapped to the region missing in the vΔCHPK mutant; therefore, they cannot be validated. The remaining three were still encoded in the ΔCHPK with moderate confidence (Table 1). This suggests that at least three sites on UL13_CHPK are differentially phosphorylated in vCHPKwt-infected cells. Corroborating these findings, our preliminary phosphoproteomic analysis using LC-MS/MS of vCHPKwt- and vCHPKmut-infected skin identified the same differentially phosphopeptides [36], potentially representing MDV UL13_CHPK substrates. Interestingly, all three unique phosphopeptides detected solely in vCHPKwt-infected skin mapped within or near predicted NLS (Fig 8).

Our previous work exploring the subcellular localization of MDV UL13_CHPK mutants, either transiently or during infection, suggests an NLS sequence is present within the N-terminal 1/3 of the protein. While diffuse nucleocytoplasmic staining was observed with full-length UL13_CHPK, the truncated protein was almost exclusively nuclear in cells [36]. Although that study did not specifically target NLS or NES sequences, the data suggest an NLS might be located within the N-terminal third of MDV UL13_CHPK or, conversely, an NES in the C-terminal 2/3 of the protein. Intriguingly, no NLS or NES have been defined for other *Orthoherpesviridae* kinases. However, research by Funk *et al.* [104] demonstrated that HSV-1 UL13_CHPK is exported from the nucleus using the NEX-TRAP system and predicted an NES at the C-terminus. This NES shows significant conservation with the predicted NES for MDV UL13_CHPK (S11 Fig). Interestingly, multiple differentially detected peptides map to a region with multiple NLS predictions, including a phosphorylation site at S82 (Table 1). Two NLS motifs have been identified in DEV CHPK, one of which aligns well with the predicted NLS for MDV CHPK [132].

Nuclear export inhibition was used in transient transfection assays to assess the subcellular localization of UL13_CHPK and its potential for nuclear export. Consistent with prior work, UL13_CHPK exhibited a nucleocytoplasmic distribution in DF-1 cells (Fig 6B). However, treatment with leptomycin B and cycloheximide resulted in a predominantly nuclear staining, indicating functional CRM1-mediated nuclear export. These results suggest that pCHPK undergoes active nuclear export. Further studies are in progress to address MDV CHPK localization in cells and will be published in more detail in subsequent studies.

#### UL14_TEG3 (*MDV026*)

HSV UL14_TEG3, a minor tegumental protein of ∼32 kDa, is expressed late during infection[133], promotes secondary envelopment [133], and shares characteristics with heat shock proteins (HSPs) or molecular chaperones [134]. HSV-2 UL14_TEG3 is cytoplasmic at physiological temperature (37°C), but shuttles to the nucleus upon heat stress (43°C). However, no defined NLS or NES motifs have been described for HSV-1 or HSV-2. Li *et al.* [54] demonstrated that the N-terminal 98 aa of DEV UL14_TEG3 contains a non-classical NLS (^8^**<u>R</u>**Q**<u>RR</u>**L**<u>R</u>**^13^), which might mediate nuclear entry. Notably, MDV UL14_TEG3 exhibits significant conservation in this region and harbors a predicted NLS (S12 Fig). Our proteomic analysis identified two unique phosphopeptides with moderate (≤ 3.5) confidence (Table 1); both map near the predicted C-terminal NLS of MDV UL14_TEG3 (S12 Fig). One of the phosphorylated sites (S201) conforms to the consensus CHPK motif (SP/PS). This motif is absent in the DEV UL14_TEG3 homolog. Further characterization of MDV UL14_TEG3, including its cellular localization and potential regulation by CHPK, may be warranted, since little is known about its role in MDV replication in chickens.

#### UL26_SCAF (*MDV038*)

UL26_SCAF and UL26.5_ICP35 of HSV-1 are encoded as a single open reading frame. *UL26* encodes the essential viral serine protease essential for virus replication, while *UL26.5* encodes ICP35, the major scaffolding protein that forms a scaffold around which the capsid shell assembles [126]. During procapsid maturation, this serine protease undergoes autocleavage, generating two proteins, VP24 (containing the protease) and VP21, which is almost identical to ICP35 except for an N-terminal extension of 59 amino acids [135–137]. In addition to acting as the molecular scaffold, UL26_SCAF functions as a nuclear import chaperone for the major capsid protein (MCP) in NLS-mediated translocation. While Gao *et al.* [126] showed independent nuclear localization of the protease and ICP35, defined NLS sequences have not been established for either HSV-1 protein. In contrast, experimental validation of an NLS has been achieved for UL80_SCAF of cytomegalovirus (CMV) [138]. Of note, the UniProt entry for HSV-1 UL26_SCAF (P10210) denotes an NLS based on sequence similarity to UL80_SCAF.

Our phosphoproteomic analysis revealed a unique phosphopeptide with moderate confidence (score of 3) in vCHPKwt-infected skin (Table 1), mapping to UL26_SCAF. The identified phosphorylation site resides upstream of a region predicted to contain both NLS and NES motifs (Fig 8). While NLS or NES motifs have not been documented for HSV or other alphaherpesvirus viral protease and scaffold proteins, Plafker *et al.* [138] described two functional NLS sequences (NLS1 and NLS2) in simian and human CMV. Interestingly, a predicted NLS in MDV UL26_SCAF aligns with NSL1 in SCAF homologs of HSV-1 and HCMV, and importantly, the unique phosphopeptide falls within this NLS1 region (S13 Fig). Although there is high sequence conservation within NLS1 between HSV, HCMV, and MDV, there were few similarities among aligned amino acids at the NLS2 site. Despite the high conservation of NLS1 across members of *Orthoherpesviridae*, there is no documented evidence that this region is involved in phosphorylation-mediated regulation. However, Plafker *et al.* [139] identified phosphorylated sites on simian CMV homolog UL80_SCAF and proposed a CKII motif in which two serines were phosphorylated. Our MS-based proteomics analysis, together with these findings, suggests that CHPK mediates phosphorylation of MDV UL26_SCAF in the skin.

#### UL30_DPOL (*MDV043*)

All herpesviruses encode their own DNA polymerase composed of two subunits, a catalytic subunit (DPOL) encoded by HSV *UL30* and a processivity factor encoded by *UL42* that interacts at the extreme C-terminus of DPOL. The C-terminus of HSV-1 UL30_DPOL harbors a well-defined NLS [140], guiding the protein into the nucleus for viral DNA synthesis. Our analysis of vCHPKwt-infected skin cells revealed two unique peptides mapping to the C-terminus of MDV UL30_DPOL. Although prediction programs did not identify this region as a potential NLS, sequence alignments show high conservation at the C-terminus, suggesting a likely functional NLS in MDV UL30_DPOL (S14 Fig). Additionally, Alvisi *et al.* [141] identified a bipartite NLS in HSV UL30_DPOL at positions 1114-1136 that corresponds to the map position of a single phosphopeptide we detected in our analysis with a moderate confidence score of 3 (Table 1). This position also aligned with the phosphorylation sites (S1113) of UL30_DPOL in HSV-1-infected cells, as reported by Bell *et al.* [129]. Despite our *in silico* analysis’s failure to predict an NLS sequence within this region, the conserved nature of the C-terminus of DPOL homologs, as first reported by Loregian *et al.* [140], suggests this region likely encodes a functional NLS for MDV UL30_DPOL. Further studies are needed to better define the NLS of MDV UL30_DPOL and to assess the potential role of phosphorylation in its nuclear translocation, thereby mediating viral DNA replication.

#### UL34_NEC2 (*MDV047*)

The core of the nuclear egress machinery comprises two components, nuclear egress complexes 1 and 2 (NEC1 and NEC2), encoded by *UL31* and *UL34*, respectively, in HSV-1 and other alphaherpesviruses. Together, these form a heterodimer essential for the egress of newly assembled viral particles from the nucleus. A hallmark of egress by many DNA viruses packaged in the nucleus is the phosphorylation of lamin proteins, which can disrupt the strong interactions between Lamin proteins, allowing for an “escape route” through a weakened lamina meshwork. For HSV-1, US3_PK phosphorylates both NEC1 and NEC2, and is important but not necessary for nuclear egress [142]. Lamin phosphorylation by CHPK has been reported for HCMV, HSV-2, and EBV [143–145]. At least for HCMV, UL97_CHPK phosphorylates UL53_NEC1 and UL50_NEC2, specifically at S19 and S216, respectively, which is important for the localization of the heterodimer [143].

Our study detected a unique phosphopeptide with a moderate confidence score of 3 (Table 1) near a predicted NLS. Translocation of HCMV UL50_NEC2 is mediated through an NLS at the N-terminus [146], and it was suggested that other herpesvirus NEC2 homologs utilize NLS based on predicted NLS sequences at this end. Interestingly, other NLS sequences could also be predicted within the C-terminal 1/3 of the protein, including HSV-1 UL34_NEC2 (S15 Fig). However, it is unknown whether the predicted MDV or HSV-1 UL34_NEC2 NLS motif is functional within the alphaherpesvirus system. The unique MDV UL34_NEC2 phosphopeptide in our study was near a predicted NLS, consistent with reports for HSV-1 NEC2. In addition, Bell *et al.* [129] detected S198 phosphorylated near a predicted NLS sequence of UL34_NEC2 in HSV-1-infected cells during PAA treatment. Further investigations are warranted to elucidate the function of the identified NLS sequence and determine whether alphaherpesvirus UL13_CHPK phosphorylates UL34_NEC2, as CMV does. However, our current data suggest that MDV UL13_CHPK may phosphorylate UL34_NEC2 in epithelial skin cells.

#### UL36_LTP (*MDV049*)

The large tegument protein (LTP), encoded by *UL36* for most alphaherpesviruses, has been studied primarily in HSV-1 and PRV, where it is essential for replication [147, 148] and is involved in capsid trafficking to the nucleus [149–151], capsid docking at the nuclear pore [152, 153], and the release of the viral DNA into the nucleus [154]. Structurally, it comprises folded domains interspersed with unstructured regions that link the capsid to the tegument layer [155]. Expression of UL36_LTP during transient transfection is localized to both the nucleus and cytoplasm, while it is primarily cytoplasmic during infection [156]. It is a phosphoprotein [90] and phosphorylated on serine residues [157]. NLS sequences have been described for HSV [158], PRV [159], and CMV [160] that appear conserved across all *Orthoherpesviridae*. Our *in silico* analysis predicted numerous NLS and NES motifs for MDV, including multiple predictions within the conserved NLS previously identified for HSV, PRV, and CMV (Fig 8). Very little is known about the phosphorylation of UL36_LTP, but Bell *et al.* [129] detected phosphorylated T702 in HSV-1-infected cells, and S832 and T1618 were phosphorylated when HSV-1-infected cells were treated with the DNA replication inhibitor PAA. A single phosphopeptide with a low-to-moderate confidence score of 2 was detected in vCHPKwt-infected skin (Table 1). Interestingly, this peptide was located within another predicted bipartite NLS (S16 Fig). Further studies are required to determine whether this NLS is functional and whether CHPK modification of UL36_LTP affects its function.

#### UL39_RIR1 (*MDV053*)

Most herpesviruses encode ribonucleotide reductase (RR) that catalyzes the conversion of ribonucleotides to deoxyribonucleotides [161]. The active form of HSV RR is a tetramer comprising homodimeric RR1 and RR2 subunits encoded by *UL39* (RIR1) and *UL40* (RIR2). Although not essential for *in vitro* replication, RR is important for the virulence of PRV in pigs and MDV in chickens [162, 163]. HSV-1 and MDV UL39_RIR1 are localized to cytoplasmic foci around the nucleus during infection in cell culture [164, 165]. Numerous NLS and NES sequences were predicted in our *in silico* analysis (Fig 8), but as far as we know, no NLS or NES have been described for RIR1 orthologs. One unique phosphopeptide with moderate confidence in our analysis mapped near a predicted NES in UL39_RIR1 of MDV (S17 Fig). For cellular RR homologs, both RIR1 and RIR2 localize in the cytoplasm [166], and phosphorylation of *Saccharomyces cerevisiae* Rnr1 (RIR1 homolog) directs its localization during S phase [167]. Under genotoxic and replicational stress, RR subunits redistribute in yeast. During normal cell cycles, Rnr2 and Rnr4 are predominantly in the nucleus. However, upon stress, these subunits shuttle to the cytoplasm in a checkpoint-dependent manner [168]. Thus, phosphorylation near the predicted NES of UL39_RIR1 homologs may help regulate its subcellular localization during stress induced by infection.

Interestingly, UL39_RIR1 is the only viral protein in our analysis to have a unique phosphopeptide with a low-to-moderate confidence score of 2 in vΔCHPK-infected cells. The phosphorylation of RR helps regulate its catalytic activity. For example, phosphorylation by CDK2/cyclin A during the S/G2 phase enhances cellular RR enzymatic activity [169]. Previously, it was thought that the N-terminus of UL39_RIR1 of HSV-1 possesses kinase activity involved in autophosphorylation. However, follow-up experiments demonstrated that this activity was due to co-purifying kinases [170–172]. Nonetheless, this does provide evidence that RR may be a substrate for phosphorylation. Using an MS-based proteomics approach, Bell *et al.* [129] showed that HSV-1 UL39_R1R1 was phosphorylated. Interestingly, T999 was phosphorylated following PAA treatment, which is near the unique phosphoserine identified in the vΔCHPK-infected skin group (S17 Fig). It is tempting to speculate that a kinase other than CHPK mediates RR activity in differentiated skin to compensate for the lack of cellular DNA replication.

#### UL51_TEG7 (*MDV065*)

TEG7, encoded by *UL51* (*MDV065*) in the MDV genome, is a minor tegument phosphoprotein [173]. While UL51_TEG7 homologs are important, but dispensable for the replication of HSV [174, 175], PRV [176], and CMV [177], ORF7_TEG7 is essential for VZV replication in human skin [178] and has recently been shown to be crucial for MDV replication *in vitro* [179]. It plays a role in the secondary envelopment of viral particles and exhibits cell-type-specific functions through interactions with UL7_CEP1, mediating cell-to-cell spread [175]. TEG7 proteins are typically localized to cytoplasmic membranes, in particular the Golgi [177, 180, 181]. MDV UL51_TEG7 exhibits distinct localization patterns. When expressed independently, it targets the mitochondria [179]. However, co-expression with its binding partner UL7_CEP1 results in a nucleocytoplasmic distribution. These findings suggest that interacting proteins may regulate the localization of UL51_TEG7.

Several studies have focused on TEG7 phosphorylation. Using MS-based proteomics, Bell *et al.* [129] identified phosphorylated residues during infection and treatment with DNA replication inhibitors, while Kato *et al.* [182] detected five phosphorylated sites on HSV-1 UL51_TEG7 during transient transfection assays. Notably, mutation of S184 significantly reduced replication and cell-to-cell spread *in vitro*, as well as mortality in mice. In our analysis of vCHPKwt-infected skin, we detected two unique peptides mapping to MDV UL51_TEG7, with high confidence scores ≥ 4 (S18 Fig). Alignment with HSV-1 UL51_TEG7 revealed that both peptides map near S184 and T190, which have been previously shown to be phosphorylated [129, 182]; however, due to weak MS2 data, we could not pinpoint the exact phosphorylated residue(s) within the two MDV peptides. Interestingly, one peptide mapped near a predicted NES, and the other near a predicted NLS. The functionality of these NLS and NES sequences and their potential role in MDV UL51_TEG7 remains to be investigated.

#### UL54_ICP27 (*MDV068*)

ICP27 is an immediate early (IE) multifunctional regulatory protein involved in the regulation of delayed-early (DE) and late (L) gene expression, including promoting transcription through interacting with the C-terminus of RNA polymerase II [183, 184]; blocking mRNA splicing through interaction with splicing factors [185–188]; and exporting viral mRNAs from the nucleus to the cytoplasm through binding to G/C-rich RNA sequences [189]. HSV-1 UL54_ICP27 predominantly resides in the nucleus but can shuttle to the cytoplasm via specific NES sites [65, 190–193]. Similar to HSV-1, MDV UL54_ICP27 is predominantly nuclear during *in vitro* infection, but diffuse nucleocytoplasmic staining is observed in infected skin *in vivo* [62].

Phosphorylation plays a vital role in ICP27 function, mediating interactions with cellular proteins [194, 195] and subcellular localization [58]. For example, specific serine residue mutations of S16, S18, and S114 in HSV-1 UL54_ICP27 severely impact replication and protein localization [195]. While S16 and S18 are thought to regulate the N-terminal NES, S114 appears to modulate the NLS, ultimately reducing nuclear import. Bell *et al.* [129] also identified that residues S114, S116, and T132 of ICP7 are phosphorylated in HSV-1-infected cells, further supporting this notion. Our *in silico* analysis suggests that MDV UL54_ICP27 encodes an N-terminal NES/NLS (Fig 8), and our MS-based proteomics analysis identified a single unique phosphopeptide within the predicted NES/NLS region with high confidence (S19 Fig). This region is also close to the described NLS of HSV UL54_ICP27, regulated by phosphorylation at S114 via the specific kinases PKA or US3_PK [58]. Interestingly, the presence of a phosphorylated S118 in our study suggests a potential role for CHPK in this modification. The proximity of phosphorylated S118 within this complex region, which contains putative NLS and NES sites, is particularly intriguing and warrants ongoing investigation.

To begin characterizing putative NLS and NES motifs in MDV UL54_ICP27, we assessed the functionality of the predicted NLS and NES sequences using transient transfection assays. Similar to our studies with US10_US10, we examined the subcellular localization of UL54_ICP27 by treating cells transiently transfected with an MDV UL54_ICP27 expression vector with leptomycin B and cycloheximide. This treatment caused a clear shift of UL54_ICP7 from a nucleocytoplasmic to a predominantly nuclear distribution (Fig 6C). These findings agree with those of other ICP27 homologs [190:Lengyel, 2002 #18000].

Next, we tested whether the predicted NLS and NES motifs were functional in MDV UL54_ICP27. To do this, we generated single- and double-deletion mutants: pcICP27ΔNES2 (aa Δ379-394), pcICP27ΔNLS1/NES1 (aa Δ98-115), pcICP27ΔNLS2 (aa Δ116-134), as shown in Fig 9A. In DF-1 cells transfected with pcICP27 or pcICP27ΔNLS1/NES1, no change in N/C ratio was measured (Fig 9BC). However, both pcICP27ΔNLS2 and pcICP27ΔNES2 led to a significant increase in nuclear localization, suggesting that these regions are involved in nucleocytoplasmic shuttling. Removal of NES2 led to a significant increase in nuclear staining, as expected for the removal of an NES. However, removal of NLS2 also significantly increased MDV ICP27 nuclear localization, contrary to the expected result for an NLS removal. It is possible that deleting NLS2 also removed an essential residue in the predicted NES1, since this deletion removed part of the NES1 sequence. Additionally, this deletion removed the S118 phosphorylation site identified in our phosphoproteomic study with infected skin (S19 Fig). However, the observed changes in subcellular localization upon deletion of the translocation signal strongly suggest their role in regulating UL54_ICP27’s movement within the cell, consistent with other ICP27 homologs [195]. These findings highlight the need for further studies to define putative NLS and NES for MDV UL54_ICP27. More defined studies examining the regulation of MDV UL54_ICP27 by CHPK are ongoing in our laboratory.

**Fig 9.**
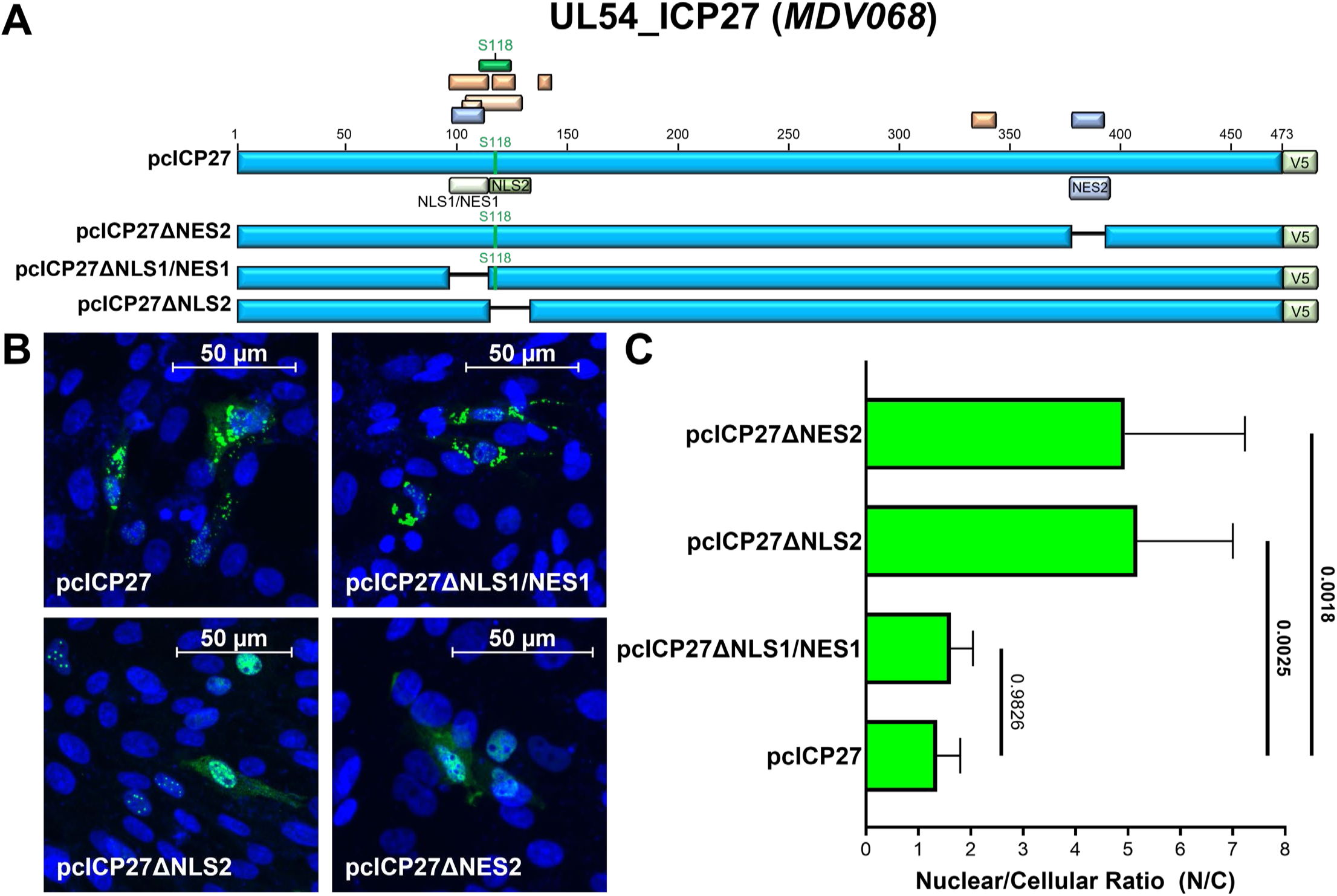
Subcellular localization of MDV UL54_ICP27 NLS and NES mutants. Putative NLS and NES motifs (A) were deleted in the MDV ICP27 expression vector previously published [62], using site-directed mutagenesis, including NLS1 (pcUS10ΔNLS1), NLS2 (pcUS10ΔNLS2), NLS3 (pcUS10ΔN3), and NES (pcUS10ΔNES). (B) Representative images of chicken DF-1 cells transfected, fixed at 24 h, and stained with a primary anti-V5 antibody and a secondary anti-mouse IgG Alexa Fluor 488 antibody. (C) Nuclear and cytoplasmic ROIs were generated using ImageJ version 2.16.0 for individual cells (n = 12 per construct), and Alexa Fluor 488 fluorescence was measured in the nucleus and cytoplasm. Mean nuclear-to-cytoplasmic (N/C) ratios were computed, and the mean with standard deviation is shown. Normality of the data was assessed using GraphPad Prism’s built-in normality test, followed by one-way ANOVA with Dunnett’s multiple post hoc comparison test. Adjusted P values are shown, with significant differences (p < 0.05) shown in bold.

### CHPK as a master regulator of viral protein expression and localization in skin cells

Although substantial additional experimental work is required to confirm which specific viral proteins are directly targeted by UL13_CHPK *in vivo*, Figure 10 presents our current working model of its role in fully productive replication within epithelial skin cells. Our phosphoproteomic analysis identified 21 proteins that were differentially phosphorylated in skin cells infected with vCHPKwt versus vΔCHPK. While some unique peptides still require validation, several candidates have been identified with reasonable confidence using our scoring criteria.

**Fig 10.**
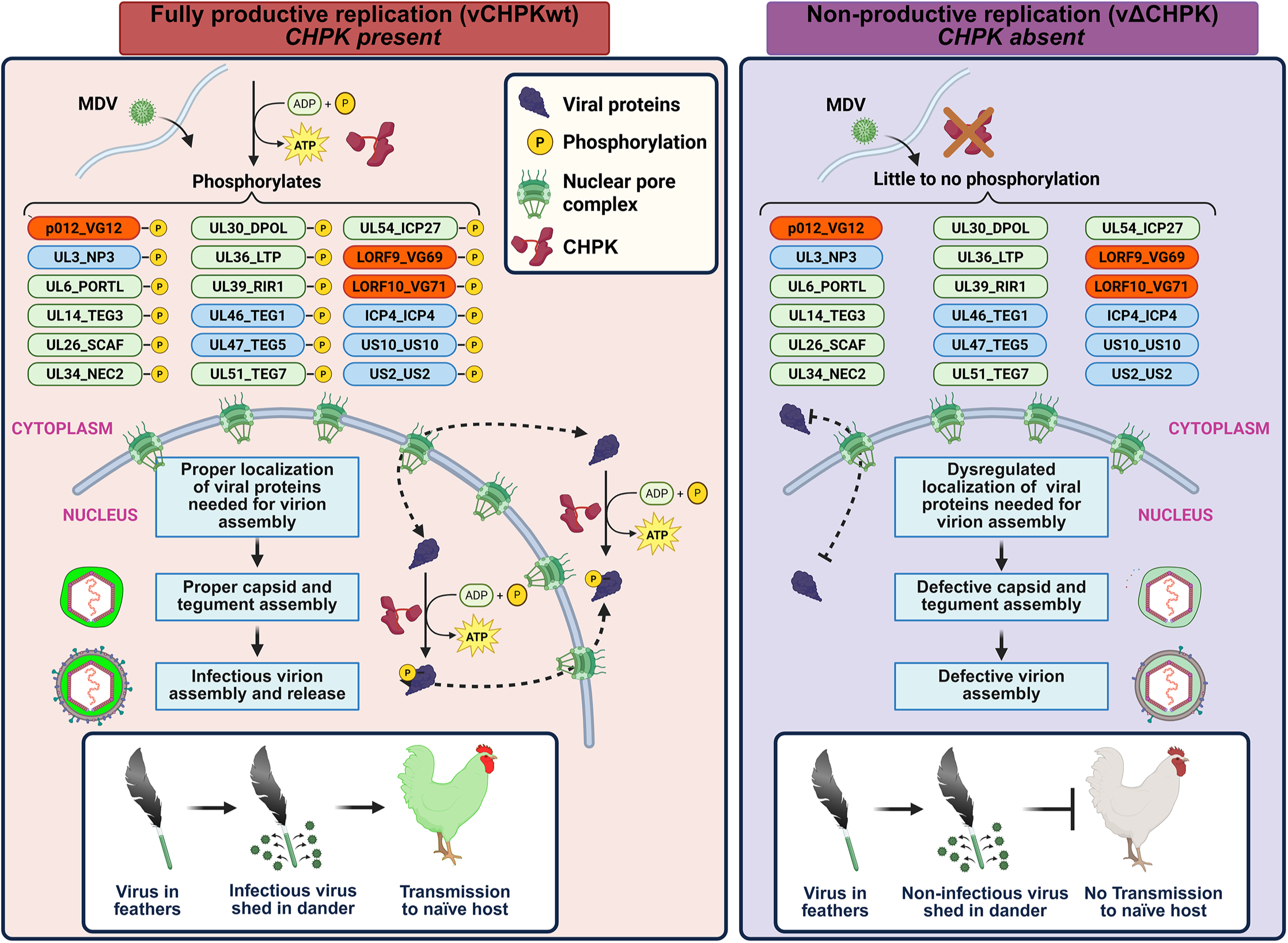
Working hypothesis for viral protein phosphorylation in the presence (vCHPKwt) and absence (vΔCHPK) during MDV replication in epithelial skin cells. Briefly, UL13_CHPK-mediated phosphorylation on viral proteins important for virion assembly is required for their proper expression and subcellular localization in cells. Without UL13_CHPK, virion assembly is defective, leading to the production of non-infectious virions in chickens and an inability to transmit to naïve chickens. Since UL13_CHPK is found in the nucleus and cytoplasm, it may play a role in both the import of viral proteins into the nucleus and the export of these proteins through phosphorylation. Colors of viral proteins are based on their conservation within the *Mardivirus* genus (orange), the *Alphaherpesvirinae* subfamily (blue), and the *Orthoherpesviridae* family (green). UL44_gC and UL45_EV45 are excluded from the figure since they are membrane proteins and not expected to be exported or imported to the nucleus.

Previous studies have established that MDV proteins UL44_gC, UL47_TEG5, and UL54_ICP27 are essential for horizontal transmission in chickens [38, 62, 102]. MDV US10 enhances disease induction during natural infection but is not strictly required for transmission [36], while UL51_TEG7 contributes to experimental infection and disease severity [179]. Deletion of LORF9_VG69 significantly reduces mortality and tumor formation in experimentally infected birds, with impaired replication specifically in FFE skin cells [70]. All six of these proteins were detected in our phosphoproteomic dataset (Fig 3), suggesting that UL13_CHPK-mediated phosphorylation may modulate their functions. The roles of the remaining MDV proteins identified in our study have not yet been characterized, to our knowledge.

Although the functional consequences of phosphorylation are likely protein-specific, a common theme may be the regulation of expression levels and subcellular localization of structural and regulatory proteins by UL13_CHPK in epithelial skin cells. In particular, UL47_TEG5 and UL54_ICP27 are known nucleocytoplasmic shuttling proteins required for horizontal transmission of MDV and whose activities are modulated by phosphorylation in other herpesviruses. Further studies are underway to determine whether UL13_CHPK directly regulates the phosphorylation, localization, and function of UL47_TEG5 and UL54_ICP27 during CFV production and horizontal transmission in chickens.

### Limitations

One limitation of this study was the absence of MS-based proteomics analysis on spleen samples. This decision was driven by several factors. First, spleen tissue was collected primarily as a control, given that the critical role of UL13_CHPK in MDV transmission is believed to occur in skin cells. Consistent with previous reports [36], deletion of UL13_CHPK does not significantly alter viral replication or oncogenesis, but specifically impairs chicken-to-chicken transmission. Second, viral replication in the spleen was limited at the sampled time points (21–35 days post-infection), as confirmed by RNA-seq data (Fig 2). This low abundance made reliable detection of viral proteins by LC-MS/MS technically challenging. Importantly, our RNA-seq results aligned with established MDV pathogenesis, revealing only a small number of viral transcripts, all of which corresponded to genes involved in latency and transformation.

A second limitation is the lack of direct evidence demonstrating phosphorylation of viral proteins by MDV UL13_CHPK. Such experiments are technically demanding and costly in the chicken model and have not yet been performed. Our own attempts to express functional MDV UL13_CHPK *in vitro* for kinase assays have also been unsuccessful to date. Future work should therefore focus on clarifying whether UL13_CHPK modifies target viral proteins directly or indirectly during replication. Nevertheless, the phosphoproteomic data presented here offer a focused, hypothesis-driven roadmap for addressing these questions.

## CONCLUSIONS

UL13_CHPK is essential for the horizontal transmission of MDV and GaHV-3 in chickens [Jarosinski, 2007 #18251; Jarosinski, 2010 #6858; Jarosinski, 2017 #17244; Blondeau, 2007 #18252; Krieter, 2020 #19285]. This function is likely mediated through CHPK-dependent regulation of viral and cellular proteins in vivo, consistent with the role of the related VZV ORF47_CHPK in human skin cells [Moffat, 1998 #18292].

To define the contribution of MDV UL13_CHPK during natural infection, we employed a top-down systems approach focused on the skin—the critical site of infectious cell-free virus production and environmental shedding. While RNA-seq and global proteomics indicated only a modest role for CHPK in viral gene transcription, MS-based phosphoproteomics revealed a far more substantial impact. Deletion of UL13_CHPK caused widespread changes in the viral phosphoproteome, affecting not only MDV-specific proteins (n=3) but also conserved Alphaherpesvirinae proteins (n=8) and core Orthoherpesviridae proteins (n=10).

A central finding of this study is the striking discordance between transcript and protein phosphorylation levels: UL13_CHPK deletion produced only minor changes in viral mRNA abundance yet triggered extensive alterations in protein phosphorylation, particularly at or near predicted NLS and NES motifs. Functional validation of select candidates supports the hypothesis that CHPK-mediated phosphorylation modulates nucleocytoplasmic trafficking of key viral proteins in epithelial skin cells. Ongoing studies are investigating these trafficking events and their importance for productive replication.

Collectively, our data establish that UL13_CHPK functions primarily as a post-translational regulator during lytic replication in skin epithelial cells, with limited influence on viral transcription. By phosphorylating multiple viral proteins—many at sites that govern subcellular localization—CHPK likely orchestrates the coordinated steps required for efficient assembly and release of cell-free virions, thereby enabling horizontal transmission (Fig 10).

Integrating these phosphoproteomic findings with existing literature on herpesviral protein localization and *in silico* NLS/NES predictions yields a coherent model: CHPK regulates the subcellular trafficking of a broad subset of viral proteins in the natural host. While this study identified functional NLS and NES motifs for several proteins and linked CHPK activity to their localization *in vivo*, it represents only an initial framework. Substantial additional work is needed to map the full repertoire of CHPK targets and to define its effects on both viral and cellular proteins during replication in epithelial cells. Nevertheless, this work provides a solid foundation for future mechanistic studies of CHPK in herpesvirus replication within its natural host and highlights its critical role in the viral life cycle.

## MATERIALS AND METHODS

### Ethics statement

All animal experimentation was conducted in accordance with national regulations. The animal care facilities and programs at UIUC meet all requirements of the law (89–544, 91–579, 94–276) and NIH regulations on laboratory animals, and comply with the Animal Welfare Act, PL 279. The UIUC Animal Care Program is accredited by the Association for Assessment and Accreditation of Laboratory Animal Care (AAALAC). All experimental procedures were conducted in compliance with approved Institutional Animal Care and Use Committee protocols. Water and food were provided to chickens *ad libitum*.

### Recombinant (r) MDV

For RNS-seq and MS data, samples were collected from a previously published experiment [36]. Briefly, MDV RB-1B strain-based vCHPKwt/10HA (VCHPKwt) and vΔCHPK/10HA (vΔCHPK) were used, in which 3×Flag and 2×HA epitopes were inserted in frame at the C-termini of *UL13* (CHPK) and *US10*, respectively, in addition to expressing UL47eGFP.

### Experimental design and tissue sampling

Pure Columbian chicks were purchased from the University of Illinois Poultry Farm (Urbana, IL). Unsexed chicks were experimentally infected (n=15/group) at three days of age with 2,000 plaque-forming units (PFU) of cell-associated MDV by intra-abdominal inoculation (Fig 1). Since both rMDVs express UL47eGFP, infection in the skin was monitored by plucking actively growing feathers that showed fluorescent viral protein expression [59]. Based on the relative level of UL47eGFP expression in feathers, the heavily infected chickens were euthanized the next day via CO2 asphyxiation to collect 20-30 actively growing wing or breast feathers and the spleen as a control tissue. For RNA collection, the calamus of the feathers (∼10 per bird) and spleen were immediately transferred to RNA STAT-60 (Tel-Test, Inc., Friendswood, TX), snap-frozen, and stored at -80°C. For protein collection, approximately 3-4 feathers per bird (n=3 birds/group) were used to scrape fluorescent cells from the calamus using a scalpel, and the cells were placed into an Eppendorf tube containing ice-cold PBS. Cells were pelleted at 2,000 × *g* for 10 min; the supernatant was removed, and all samples were stored at -80°C until transferred to the University of Illinois Proteomics core for MS-based proteomics.

### Extraction of nucleic acids from the calamus of feathers and whole spleens

RNA and DNA were extracted from FFE skin cells and matched spleens of birds infected with vCHPKwt, vΔCHPK, or mock-inoculated using the manufacturer’s instructions for RNA STAT-60 and DNA STAT-60 (Tel-Test, Inc.). Briefly, samples frozen in RNA STAT-60 were thawed, and 1.0 ml was removed for nucleic acid isolation. After chloroform was added to the thawed samples and the mixture was centrifuged, the aqueous phase was removed and processed for RNA as previously reported by Volkening *et al.* [60]. To isolate DNA from the phenol phase after RNA extraction, DNA STAT-60 and chloroform were added according to the following ratio of starting RNA STAT-60/DNA STAT-60/Chloroform (10/8/1.6) to the phenol phase. After extraction and centrifugation, the resulting aqueous phase containing DNA was extracted twice with phenol/chloroform and once with chloroform. The DNA was then precipitated with ammonium acetate and isopropanol. The quantities of RNA and DNA were determined using Qubit Broad-range Fluorometer assays (Thermo Fisher). The quality of the RNA was assessed using an RNA Nano chip on a Bioanalyzer 2100. Only RNA samples with RIN values > 7.0 were used in downstream Illumina-based sequencing applications.

### Viral genome levels using qPCR assays

MDV DNA copy numbers were detected using qPCR as previously described using primers and probes specific for the MDV-infected cell protein 4 (*ICP4)* locus [196] that were normalized to cellular genome copies of chicken α-collagen [197].

### RNA sequencing (RNA-seq)

The procedure for sample collection, RNA extraction, and RNA sequencing was recently reported [60]. RNA integrity was determined using a Bioanalyzer 2100 (Agilent, Santa Clara, CA, USA). RNA samples with RIN values >7.0 were depleted of rRNAs using QIAseq FastSelect –rRNA HMR kit (Qiagen, Germantown, MD, USA) in combination with the KAPA stranded mRNA seq kit (Kapa Biosystems, Wilmington, MA, USA). The high-quality ribosome-depleted RNA was reverse transcribed using random hexamer priming, end-repaired, and indexed with individual adaptors. Libraries were quantified using a Qubit fluorometer and analyzed on a Bioanalyzer 2100 to determine their size distribution. Pooling cDNAs with fragments (200–300 bp) was done using qPCR concentrations. The quality of the final pool was determined using Qubit, a fragment analyzer, and qPCR. RNA libraries were prepared for sequencing on the Illumina NextSeq 500 instrument using Illumina’s dilute and denature protocol. Pooled libraries were diluted to 2nM and denatured with NaOH. The denatured libraries were further diluted to 2.2 pM, and PhiX was added at 1% of the library volume. Data were demultiplexed, adapter sequences were trimmed, and barcoded sequences were uploaded to the BaseSpace Sequencing Hub.

### RNA-Seq data analysis

Short-read RNA-seq data analysis was previously published for vCHPKwt-infected skin cells [60]. Here, we analyzed RNAseq data on vCHPKwt- and vΔCHPK-infected spleen and skin samples. Paired-end RNA sequencing reads were processed using a combined host-virus analysis framework. Raw FASTQ reads were first filtered using fastp to remove low-quality bases and reads below the minimum threshold using options --detect_adapter_for_pe, --qualified_quality_phred score of 20, and minimum length_requied of 30 [198]. FastQC was used to quality check the trimmed data. Filtered reads were then aligned to a combined reference genome consisting of Gallus_gallus bGalGal1.mat.broiler.GRCg7b and the viral genome using the STAR aligner [199].

Gene-level read quantification was performed using featureCounts (subread package) with a combined GTF annotation using both chicken and viral gene models [200]. Reads were assigned to genomic features annotated as exons and summarized at the gene level from the annotation file. DESeq2 was used on the resulting count matrix to perform differential expression analysis between vCHPKwt and vΔCHPK infections using a negative binomial model, and log2 fold changes were calculated [201]. P-values were adjusted for multiple testing to control the false discovery rate (FDR).

An mvabund multivariate test (mvabund::mvabund; mvabund v4.2.8) [202] was used to identify associations between gene expression profiles and conditions. Features for multivariate tests were selected by first computing variable stabilizing transformation values for all genes in DESeq2. Raw counts for each of the top 500 most variable genes were retained, and negative binomial generalized linear models were fit for each gene using condition (i.e.,wt vs mutant) as a predictor. Multivariate *P* values were calculated using PIT-trap resampling (nBoot=999). Deviance tests were used to assess deviations in gene expression between vCHPKwt- and vΔCHPK-infected samples.

### Proteomics and phosphopeptide analyses

The complete procedure for protein extraction, processing, and analysis has recently been published [60]. Briefly, proteins were used for sequential proteolytic digestion by LysC (1:100 w/w enzyme: substrate; Wako Chemicals) for 4 h at 30°C and trypsin (1:50 w/w; Pierce) overnight at 37°C, followed by desalting using Sep-Pak C18 columns (Waters, Milford, MA) and dried in a vacuum centrifuge. For phosphor-proteomic analysis, phosphopeptides were enriched by iron-immobilized metal ion affinity chromatography (Fe-IMAC) in a microtip format before being desalted once more using StageTips [203].

Digested peptides were analyzed using a Thermo UltiMate 3000 UHPLC system coupled to a high-resolution Thermo Q Exactive HF-X mass spectrometer and were separated by reversed-phase chromatography. The MS was operated in a data-dependent manner where precursor scans from 350 to 1500 m/z (120,000 resolution) were followed by higher-energy collisional dissociation (HCD) of the 15 most abundant ions. MS2 scans were acquired at a resolution of 15,000 with a precursor isolation window of 1.2 m/z and a dynamic exclusion window of 60 s.

#### Differential protein analysis

The raw LC-MS/MS data were analyzed using the Byonic peptide search algorithm (Protein Metrics) integrated into Proteome Discoverer 2.4 (Thermo Scientific) against the UniProt GaHV2 database (taxon 10390; 1300 sequences). Main search settings were initially determined with Byonic Preview (Protein Metrics) and included a peptide precursor mass tolerance of 8 ppm with a fragment mass tolerance of 20 ppm. A maximum of two missed tryptic digestion cleavages was specified. Tryptic digestion was specified with a maximum of two missed cleavages, while peptide and fragment mass tolerances were set to 10 ppm and 0.6 Da, respectively. Variable modifications included oxidation of methionine, acetylation of protein N-termini, and phosphorylation of serine, threonine, and tyrosine; a fixed modification to account for cysteine carbamidomethylation was also added to the search. The false discovery rate was set to 0.01 using a decoy database strategy, and any sequences that did not meet this cutoff were discarded. Phosphorylation sites were localized using ptmRS [204].

Differential expression analyses were performed as previously reported [205]. First, the identified protein data were corrected for nonspecific background. The normalized LFQ data were uploaded to LFQ-Analyst (a web-based tool) and processed for statistical analysis to perform pair-wise comparison between vCHPKwt, vΔCHPK, and uninfected control (i.e., vCHPKwt vs. uninfected, vΔCHPK vs. vCHPKwt) protein data. Significantly differentially expressed proteins between the comparisons were declared at the threshold cutoff of FDR (adj- p-value) < = 0.05, and linear FC ≤-2 or ≥2. Among the replicates, outliers were removed based on correlation and PCA analysis. The differentially expressed proteins were used for GO enrichment analysis. The false discovery rate was set to 0.01 using a decoy database strategy, and any sequences that did not meet this cutoff were discarded. Phosphorylation sites were localized using ptmRS [204].

### Plasmids

Expression plasmids for MDV US10 and ICP27 have been previously described [36, 62]. Site-directed mutagenesis was used to delete putative NLS and NES motifs in plasmids using primers shown in S2 Table. Briefly, respective plasmids pcCHPKwt (with 3×Flag epitope tag), pcUS10wt (with eGFP tag), and pcICP27wt (with V5 epitope tag) were PCR amplified using Thermo Scientific Phusion Flash High-Fidelity PCR Master Mix, gel purified, and then ligated using T4 DNA ligase (Invitrogen). Deleted sequences were verified using Sanger sequencing at the University of Illinois DNA Services core facility.

### Laser scanning confocal microscopy

Immortalized chicken embryonic fibroblast DF-1 cells (ATCC CRL-12203) were seeded onto sterile glass coverslips in 6-well tissue culture dishes and transfected with expression plasmids using Lipofectamine 2000 (Invitrogen) according to the manufacturer’s instructions. DF-1 cells were maintained in Corning Dulbecco’s modified essential medium (Fisher Scientific) supplemented with 10% fetal bovine serum and 1% penicillin-streptomycin antibiotics (100 U/ml penicillin and 100 µg/ml streptomycin) and used in transient expression assays. After 24 h, cells were fixed and permeabilized with PFA buffer (2% paraformaldehyde, 0.05% Triton X-100) for 15 min, then washed twice with PBS. For some experiments, nuclear inhibition assays were used in transient transfection assays. To do this, DF-1 cells transfected with expression plasmids were treated with 20 nM leptomycin B (Thermo Scientific Chemicals, Ward Hill, MA, USA) for 6 h to stop nuclear export, plus 50 µg/ml cycloheximide (Millipore Sigma, St. Lois, MO, USA) for the last 1.5 h of treatment to stop protein translation before fixation. Viral protein localization was either directly visualized using fluorescent tags or stained with antibodies to epitope tags, as previously described [20]. Mouse anti-Flag M2 (Sigma-Aldrich), or anti-V5 E10/V4RR (Thermo Fisher Scientific, Waltham, MA) mAbs were used to detect CHPK and ICP27, respectively. A Nikon A1 Confocal Laser Microscope with the NIS-Elements C platform was used to capture images, and images were compiled using Adobe® Photoshop® version 21.0.1.

### NLS and NES predictions

Multiple prediction models were used for all viral proteins identified in this report. For NLS predictions, Nostradamus [206], NLSdb [207], SeqNLS [208], cNLS mapper [209], and MyHits [210] were used, while LocNES [211] was used to predict NES motifs. Since all prediction programs have their own strengths and weaknesses, we used the lowest stringency when running the programs, but included only the three highest-scoring sequences for each protein and prediction program in our analyses.

### Statistical analysis

All statistical analyses were performed using GraphPad Prism version 10.6.1 (GraphPad Software). Data are presented as Mean ± SEM unless otherwise indicated. The statistical test used for each dataset is specified in the corresponding figure legend.

## DATA AVAILABILITY STATEMENT

Raw transcriptome sequencing data are available in NCBI under BioProject PRJNA1484153 (SRA). The LC-MS data have been deposited with the ProteomeXchange Consortium via the jPOST partner repository (https://jpostdb.org) with the dataset identifier (To be deposited).

## AUTHOR CONTRIBUTIONS

**Conceptualization:** Keith W. Jarosinski, Stephen J. Spatz

**Data Curation:** Keith W. Jarosinski, Bushra Fazal Minhas, Christopher A. Gaulk

**Formal Analysis:** Haji Akbar, Keith W. Jarosinski, Bushra Fazal Minhas, Christopher A. Gaulk

**Funding Acquisition:** Keith W. Jarosinski, Stephen J. Spatz

**Investigation:** Haji Akbar, Keith W. Jarosinski, Nagendraprabhu Ponnuraj

**Methodology:** Haji Akbar, Keith W. Jarosinski, Nagendraprabhu Ponnuraj

**Project Administration:** Keith W. Jarosinski

**Resources:** Keith W. Jarosinski, Bushra Fazal Minhas, Christopher A. Gaulk

**Supervision:** Keith W. Jarosinski, Christopher A. Gaulk

**Validation:** Haji Akbar, Keith W. Jarosinski, Christopher A. Gaulk

**Visualization:** Haji Akbar, Keith W. Jarosinski, Nagendraprabhu Ponnuraj, Bushra Fazal Minhas

**Writing—original draft preparation:** Keith W. Jarosinski

**Writing—review and editing:** Haji Akbar, Keith W. Jarosinski, Bushra Fazal Minhas, Christopher A. Gaulk, Stephen J. Spatz

All authors have read and agreed to the published version of the manuscript.

## FUNDING

This report was supported by Agriculture and Food Research Initiative Competitive Grant nos. 2013-67015-26787, 2016-67015-26777, and 2020-67015-21399 from the USDA National Institute of Food and Agriculture to KWJ, and USDA-ARS NACA agreements nos. 58-6040-8-037 and 58-6040-0-015 to KWJ and SJS. The funders had no role in study design, data collection and analysis, decision to publish, or preparation of the manuscript.

## ACKNOWLEDGEMENTS

The authors would like to thank Dr. Justine Arrington at the University of Illinois Protein Sciences core for her help in performing and analyzing proteomics data and Dr. Jeremy Volkening at Base2Bio for consultation on RNA sequencing and proteomics analyses. This work made use of the equipment, software, and facilities provided by the University of Illinois Urbana-Champaign College of Veterinary Medicine Shared Equipment Program’s Biocomputing Shared Resource (BioShaRe). The College of Veterinary Medicine BioShaRe is housed in the Illinois Campus Cluster, a computing resource operated by the Illinois Campus Cluster Program (ICCP) in conjunction with the National Center for Supercomputing Applications (NCSA), and supported by funds from the University of Illinois at Urbana-Champaign.

## SUPPORTING INFORMATION CAPTIONS

**S1 Fig.**
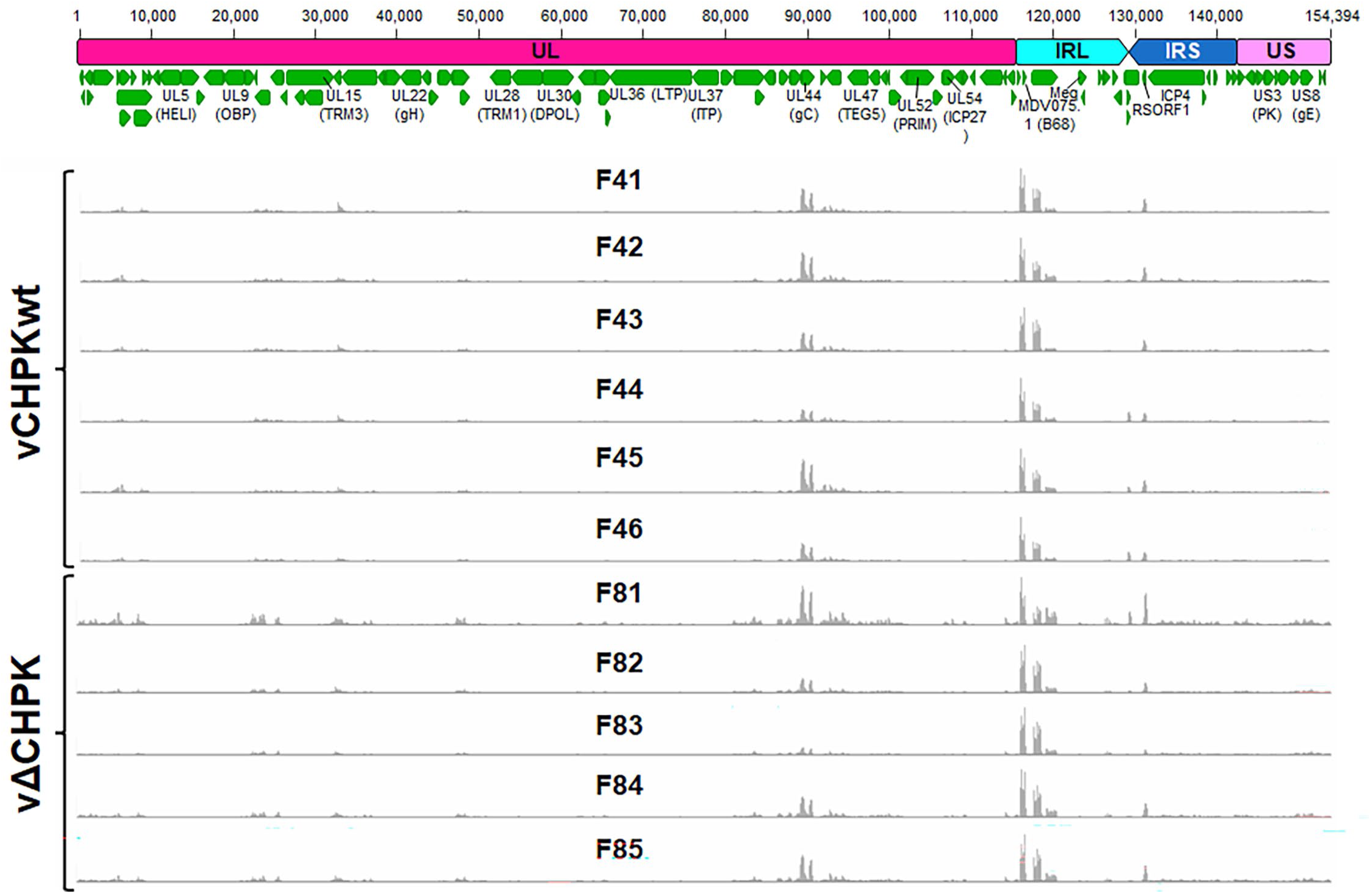
Schematic representation of the MDV genome (TRL and TRS trimmed) and RNA-seq read depth from vCHPKwt- (F41-46) and vΔCHPK- (F81-85) infected skin tissues. Read depth tracks were generated using the Integrative Genomics Viewer (IGV) version 2.19.7 [212] for all infected skin (feather) samples.

**S2 Fig.**
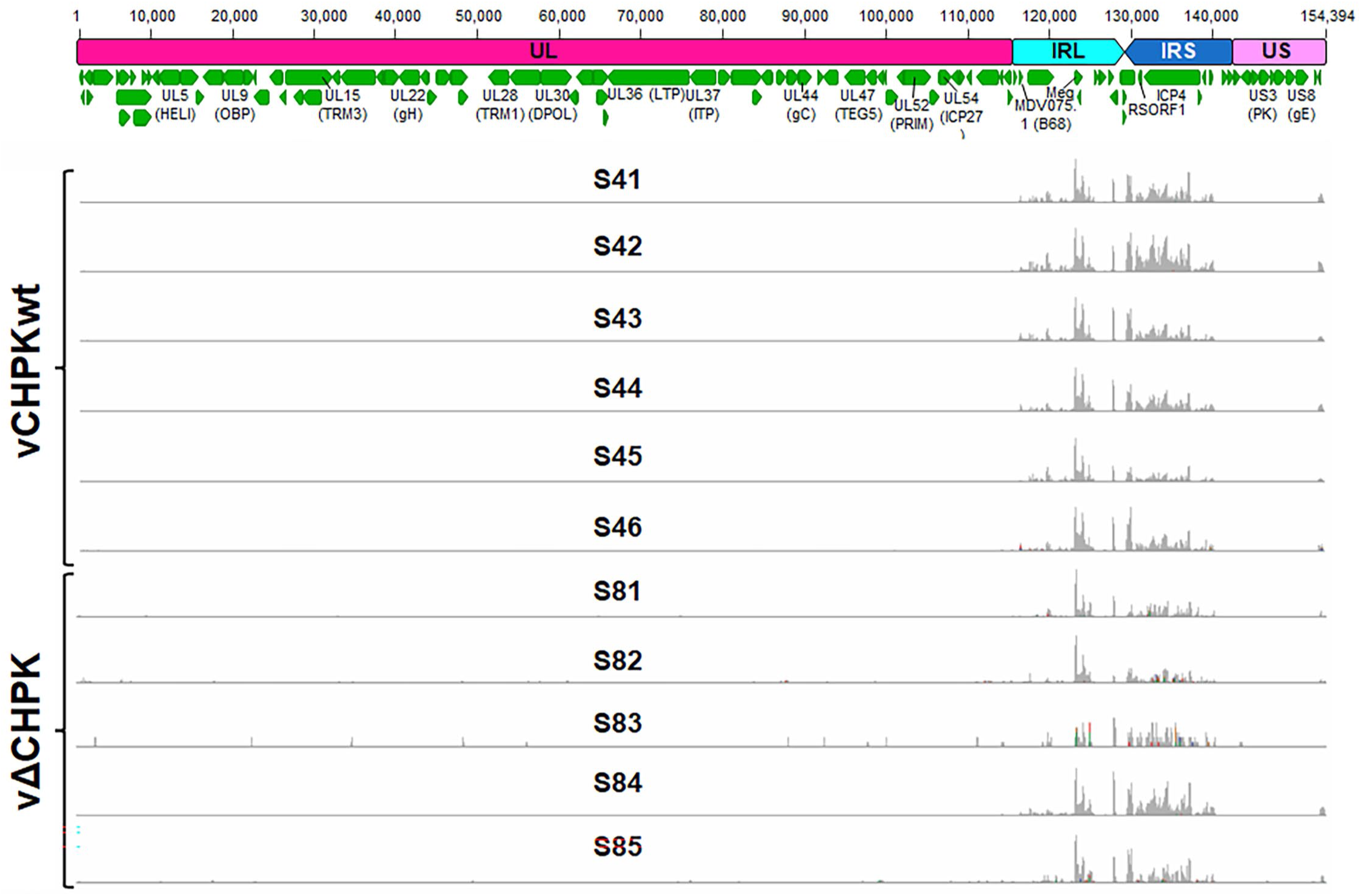
Schematic representation of the MDV genome (TRL and TRS trimmed) and RNA-seq read depth from vCHPKwt- (S41-46) and vΔCHPK- (S81-S85) infected spleens. Read depth tracks were generated using the Integrative Genomics Viewer version 2.19.7 [212] for all spleen samples.

**S3 Fig.**
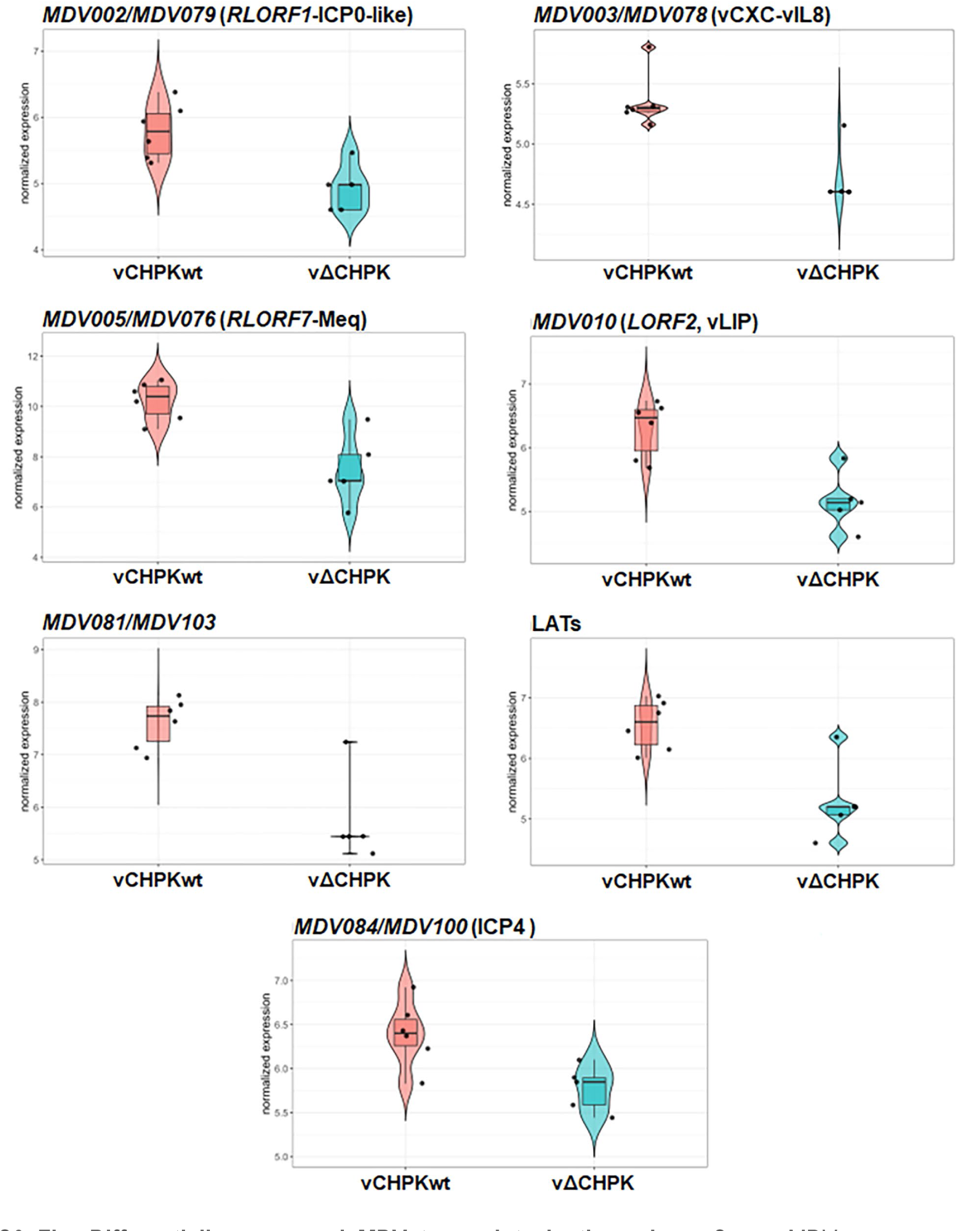
Differentially expressed MDV transcripts in the spleen. Seven MDV genes were differentially expressed between vCHPKwt and vΔCHPK-infected spleens at a p-adj value of < 0.05.

**S4 Fig.**
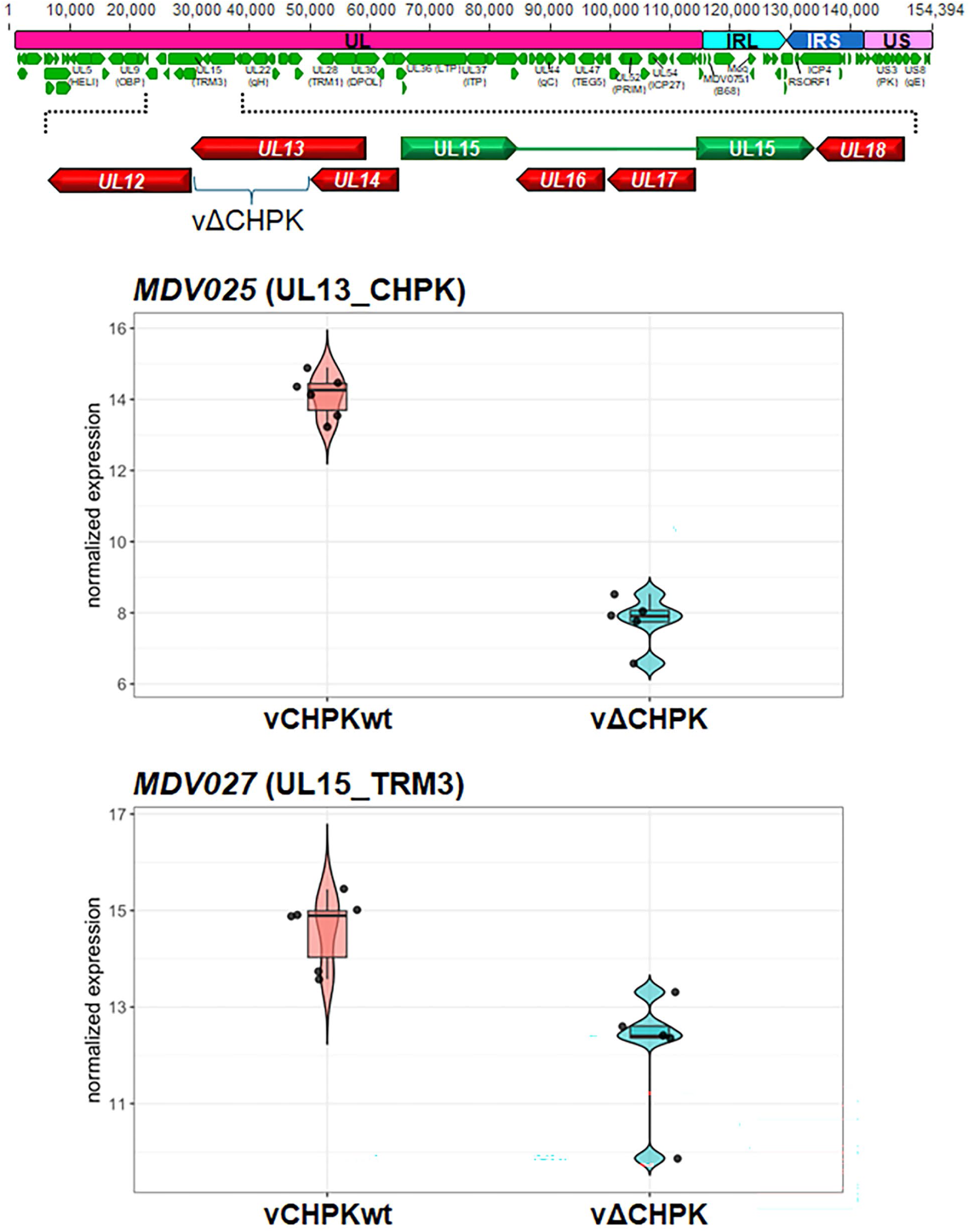
Differentially expressed MDV genes in the skin. (A) Schematic representation of the MDV genome (with TRL and TRS trimmed) and an expanded view of the region encoding UL12-UL18. (B) Normalized gene expression of the two MDV genes differentially transcribed between vCHPKwt- and vΔCHPK-infected skins, with p-adj values <0.05.

**S5 Fig.**
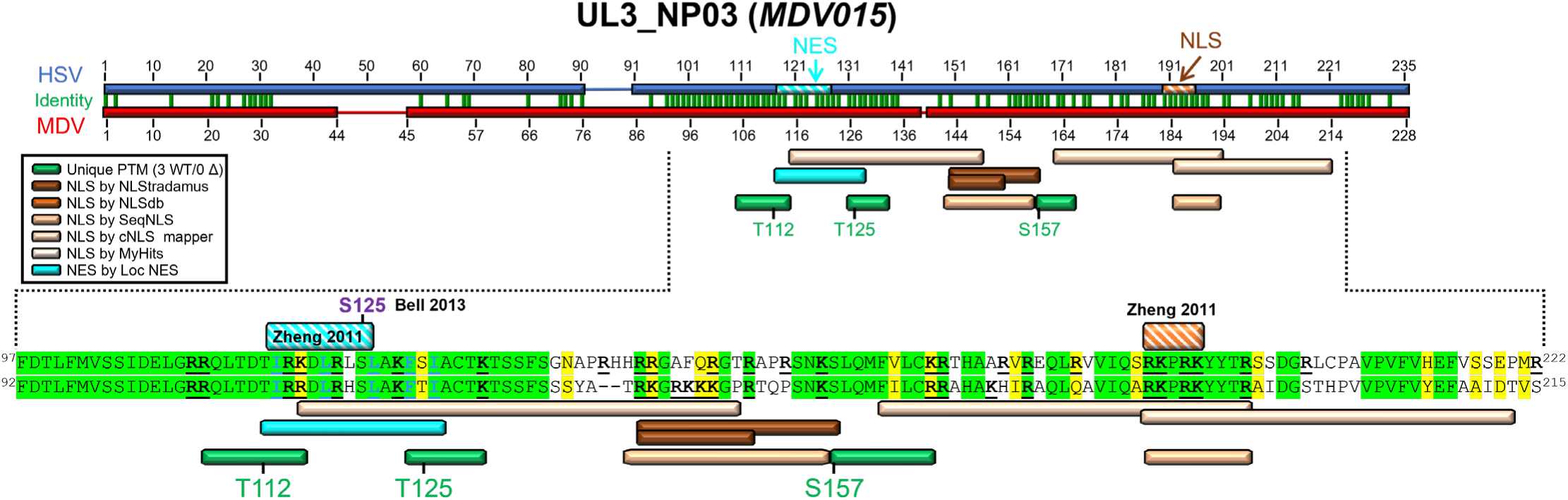
Alignment of HSV (Q1XBW5) and MDV (Q77MS7) UL3_NP03 proteins from the UniProt Consortium [213] using MUSCLE Alignment in Geneious Prime. Regions of note are expanded to the residue level. Specific residues identified as phosphorylated are shown in green text. Residues important for NLS are bolded and underlined in black. Residues in red align with the predicted consensus site for HSV CHPK phosphorylation (SP/PS). Conserved residues are based on BLOSUM62 matrix scores, with green residues indicating 80-100% similarity and yellow residues indicating 60-80% similarity. Residue previously shown to be phosphorylated by Bell *et al.* [129] is noted in purple text. NES and NLS, as identified by Zheng *et al.* [74], are shown.

**S6 Fig.**
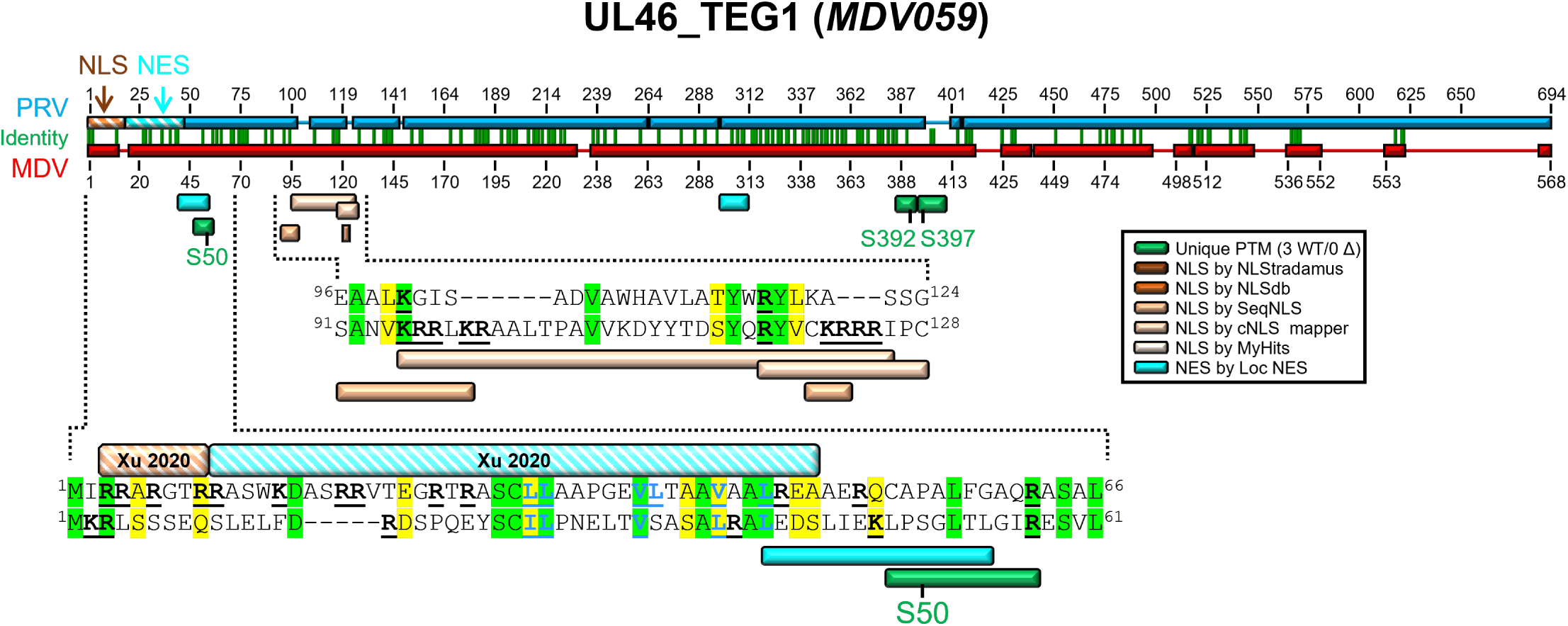
Alignment of PRV (A0A1W5LAD9) and MDV (Q77MR6) UL46_TEG1 proteins from the UniProt Consortium [213] using MUSCLE Alignment in Geneious Prime. Regions of note are expanded to the residue level. Published and predicted NLS and NES are shown along with unique peptides identified in this report. Specific residues identified as phosphorylated are shown in green text. Residues important for NES are bolded and underlined in blue, while residues important for NLS are bolded and underlined in black. Conserved residues are based on BLOSUM62 matrix scores, with green residues indicating 80-100% similarity and yellow residues indicating 60-80% similarity. NES and NLS for PRV UL46_TEG1have been functionally characterized by Xu *et al.* [100].

**S7 Fig.**
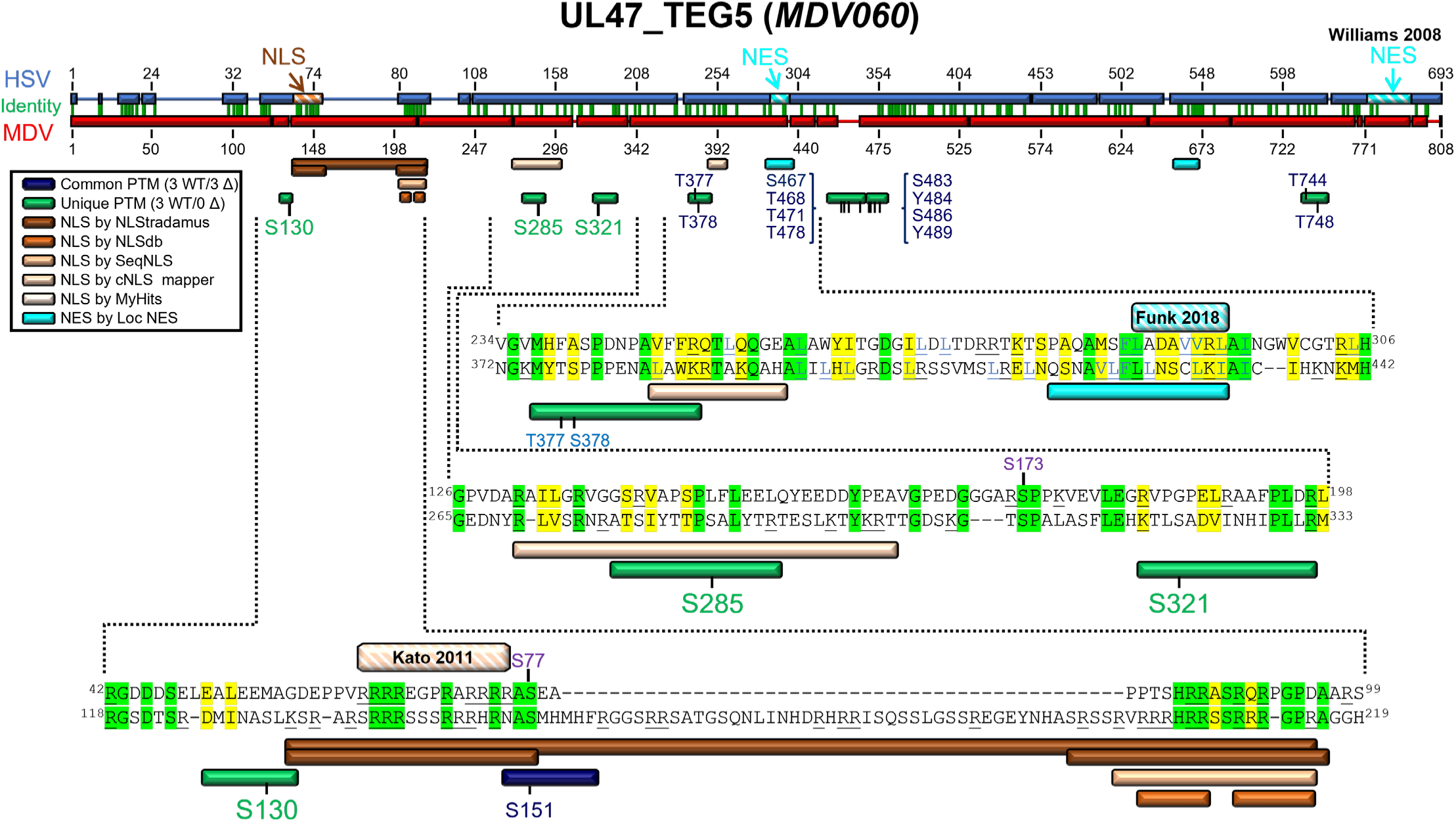
Alignment of HSV (P10231) and MDV (Q9E6M8) UL47_TEG5 proteins from the UniProt Consortium [213] using MUSCLE Alignment in Geneious Prime. Regions of note are expanded to the residue level. Published and predicted NLS and NES are shown along with unique peptides identified in this report. Specific residues identified as phosphorylated are shown in green text. Residues important for NES are bolded and underlined in blue, while residues important for NLS are bolded and underlined in black. Conserved residues are based on BLOSUM62 matrix scores, with green residues indicating 80-100% similarity and yellow residues indicating 60-80% similarity. An NLS for HSV UL47_TEG5 has been functionally characterized by Kato *et al.* [57] and NES motifs were defined by Williams *et al.* [103] and Funk *et al.* [104].

**S8 Fig.**
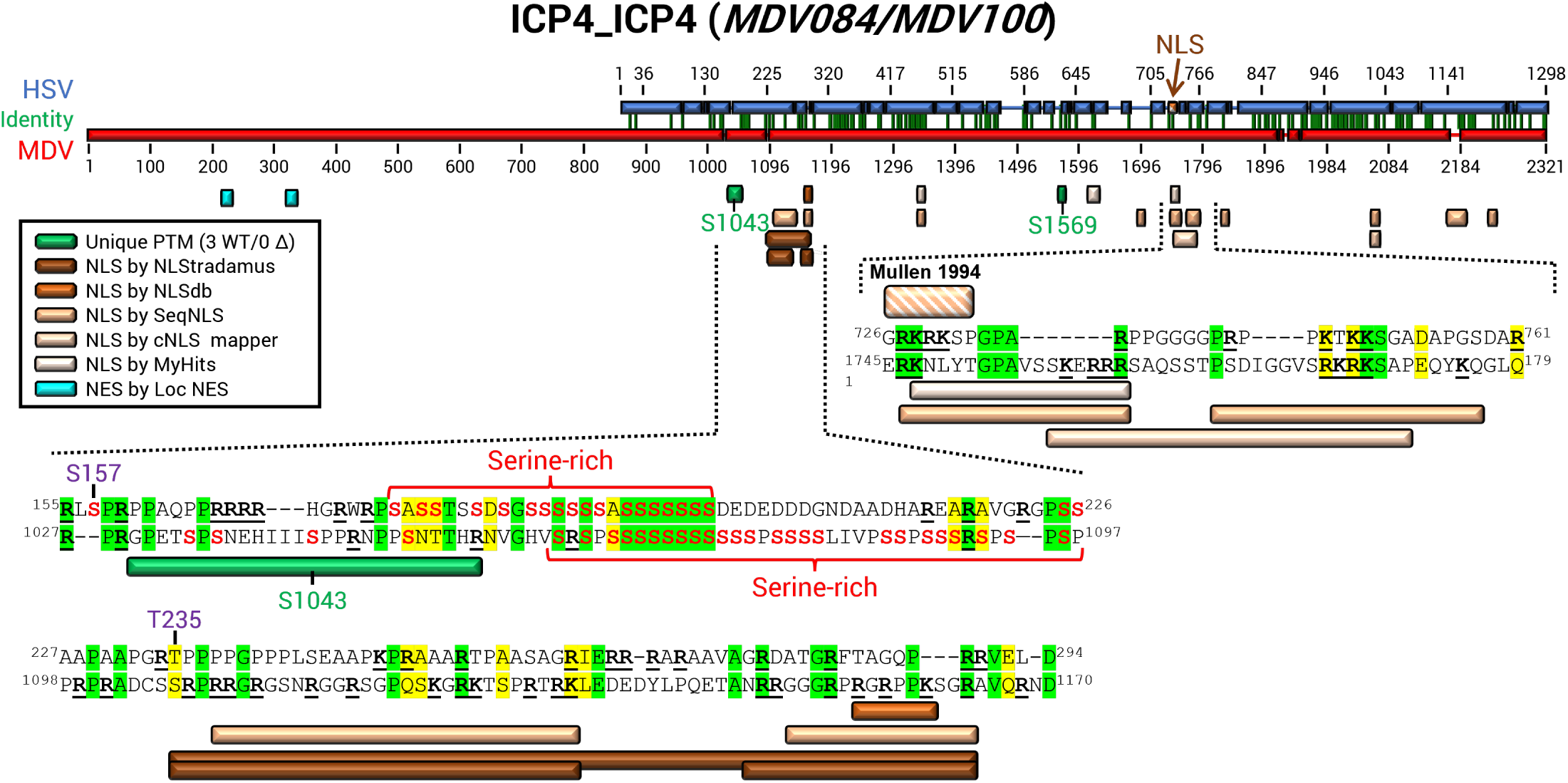
Alignment of HSV (P08392) and MDV (Q9DGT6) ICP4 proteins from the UniProt Consortium [213] using MUSCLE Alignment in Geneious Prime. Regions of note are expanded to the residue level. Published and predicted NLS and NES are shown along with unique peptides identified in this report. Specific residues identified as phosphorylated in this report are shown in green text. Residues important for NLS are bolded and underlined in black. Conserved residues are based on BLOSUM62 matrix scores, with green residues indicating 80-100% similarity and yellow residues indicating 60-80% similarity. An NLS for HSV ICP4 has been previously described by Mullen *et al.* [105]. Bell *et al.* [129] identified phosphorylated residues annotated in purple here.

**S9 Fig.**
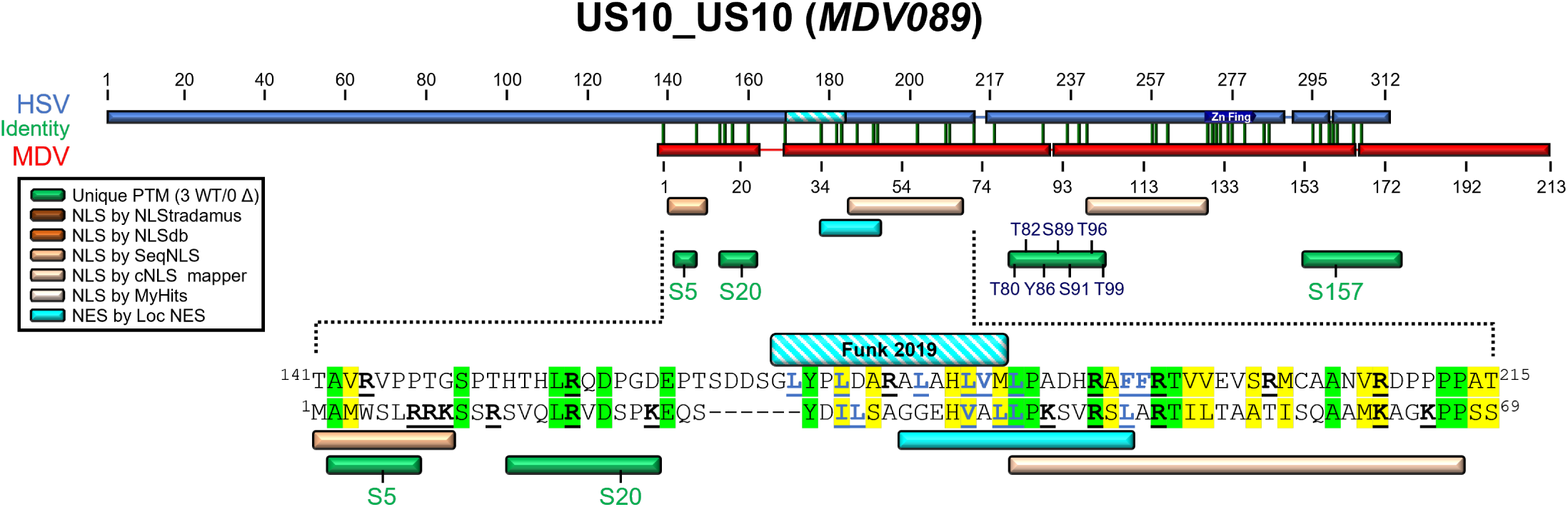
Alignment of HSV (P06486) and MDV (Q77MP8) US10 proteins from the UniProt Consortium [213] using MUSCLE Alignment in Geneious Prime. Regions of note are expanded to the residue level. Published and predicted NLS and NES are shown along with unique peptides identified in this report. Specific residues identified as phosphorylated are shown in green text. Residues important for NLS are bolded and underlined in black, while residues important for NES are bolded and underlined in blue. Conserved residues are based on BLOSUM62 matrix scores, with green residues indicating 80-100% similarity and yellow residues indicating 60-80% similarity. An NES was predicted for HSV US10 by Funk *et al.* [104], which aligns closely with the predicted MDV US10 NES.

**S10 Fig.**
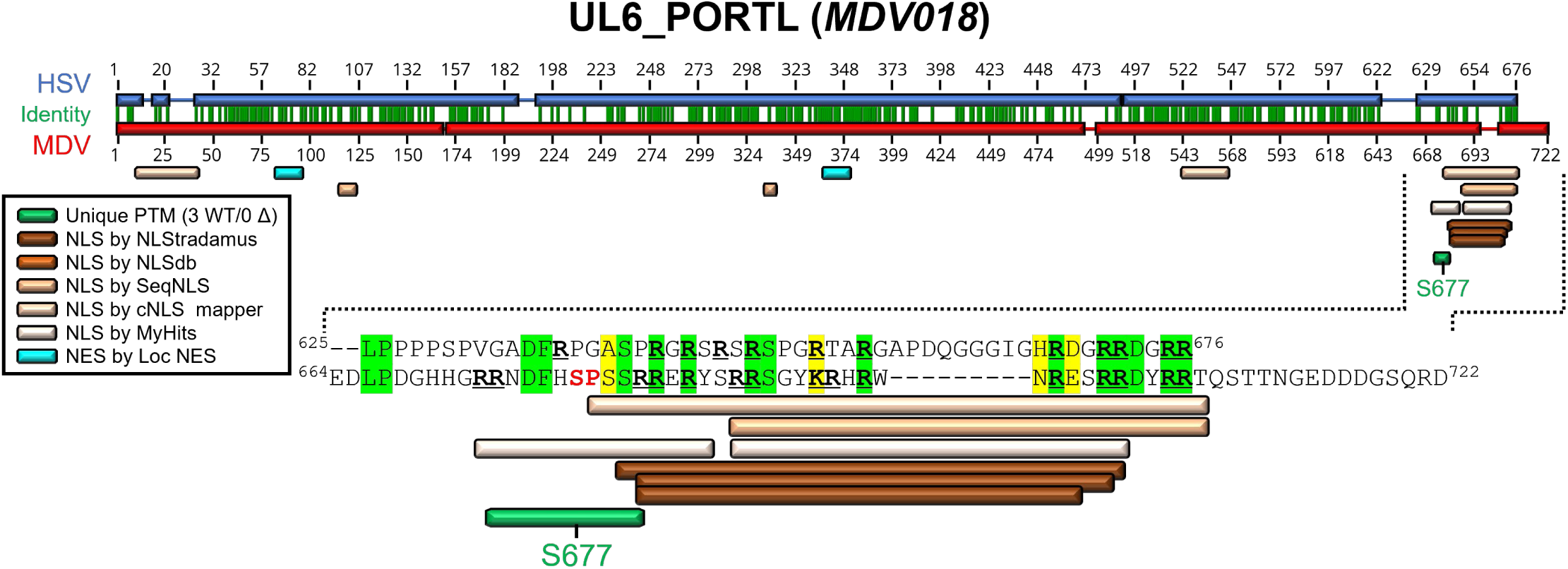
Alignment of HSV (P10190) and MDV (Q9E6R0) UL6_PORTL proteins from the UniProt Consortium [213] using MUSCLE Alignment in Geneious Prime. Regions of note are expanded to the residue level. Specific residues identified as phosphorylated are shown in green text. Residues important for NLS are bolded and underlined in black. Residues in red align with the predicted consensus site for HSV CHPK phosphorylation (SP/PS). Conserved residues are based on BLOSUM62 matrix scores, with green residues indicating 80-100% similarity and yellow residues indicating 60-80% similarity.

**S11 Fig.**
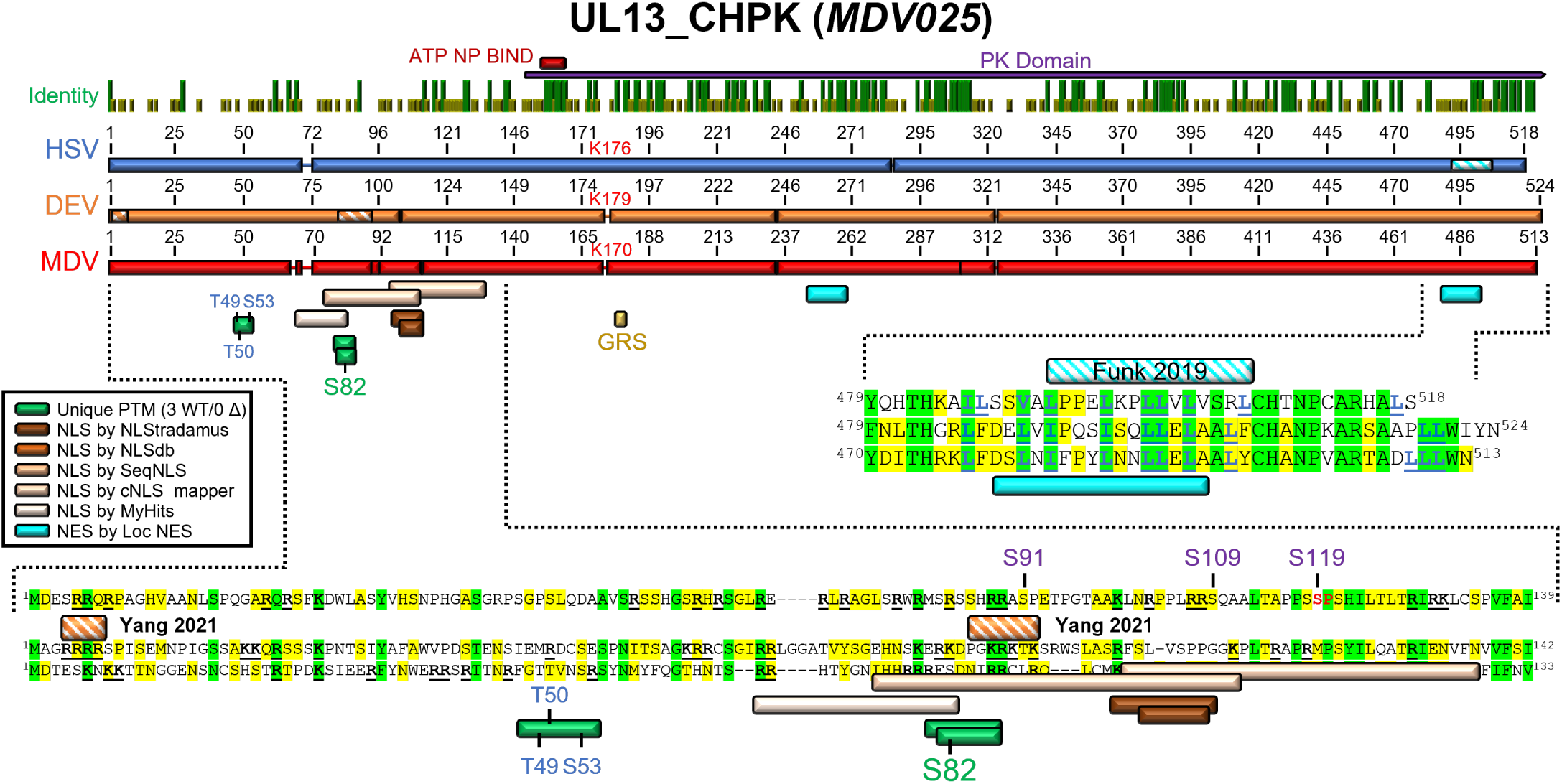
Alignment of HSV (P04290), DEV (A5A413_9ALPH), and MDV (Q9E6Q4) CHPK proteins from the UniProt Consortium [213] using MUSCLE Alignment in Geneious Prime. Regions of note are expanded to the residue level. Published and predicted NLS and NES are shown along with unique peptides identified in this report. Specific residues identified as phosphorylated are shown in green text. Residues important for NES are bolded and underlined in blue, while residues important for NLS are bolded and underlined in black. Conserved residues are based on BLOSUM62 matrix scores, with green residues indicating 80-100% similarity and yellow residues indicating 60-80% similarity. Funk *et al.* [104] predicted an NES for HSV UL13_CHPK, while Yang *et al.* [132] identified two NLS in DEV CHPK. Residues previously shown to be phosphorylated by Koyanagi *et al.* [130] and Bell *et al.* [129] are noted in purple text.

**S12 Fig.**
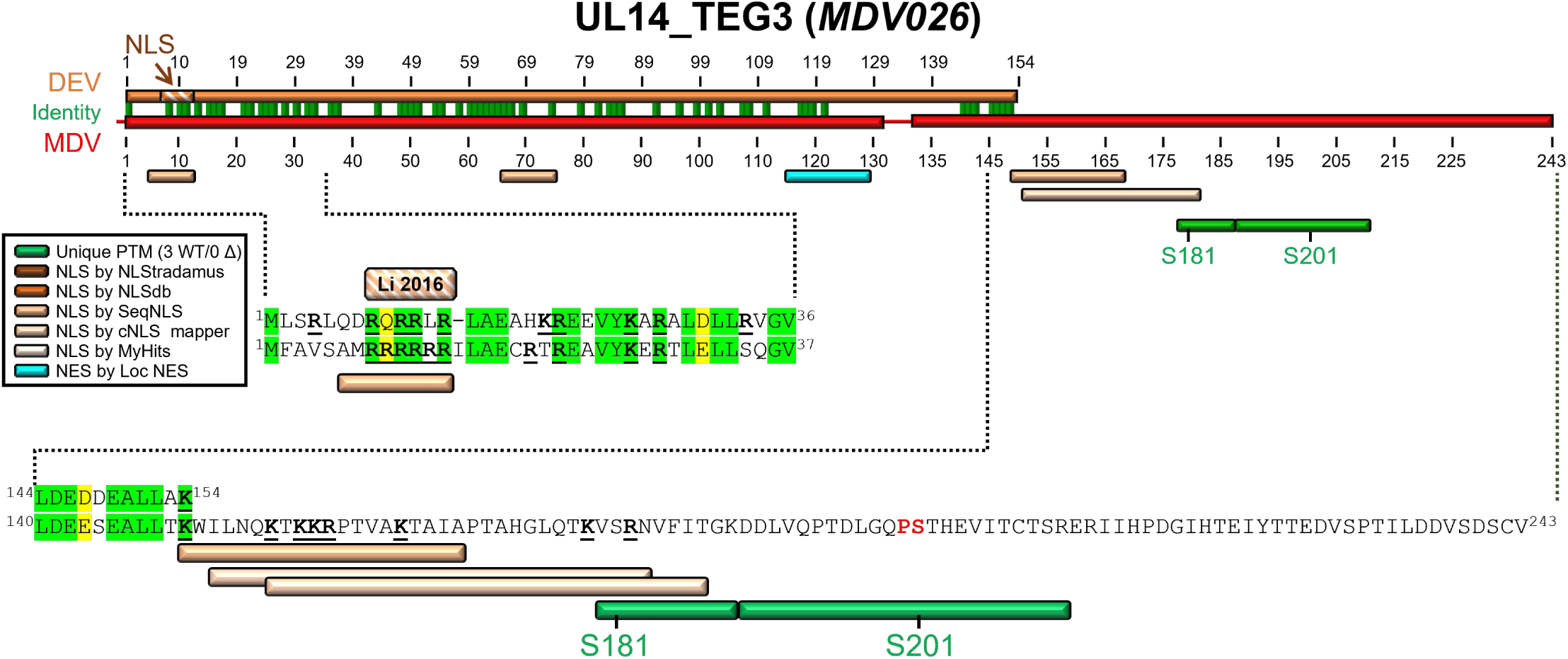
Alignment of DEV (A5A414) and MDV (Q9E6Q3) TEG3 proteins from the UniProt Consortium [213] using MUSCLE Alignment in Geneious Prime. Regions of note are expanded to the residue level. Published and predicted NLS and NES are shown along with unique peptides identified in this report. Specific residues identified as phosphorylated are shown in green text. Residues important for NLS are bolded and underlined in black. Residues in red align with the predicted consensus site for HSV CHPK phosphorylation (SP/PS). Conserved residues are based on BLOSUM62 matrix scores, with green residues indicating 80-100% similarity and yellow residues indicating 60-80% similarity. An NLS for DEV has been characterized by Li *et al.* [54].

**S13 Fig.**
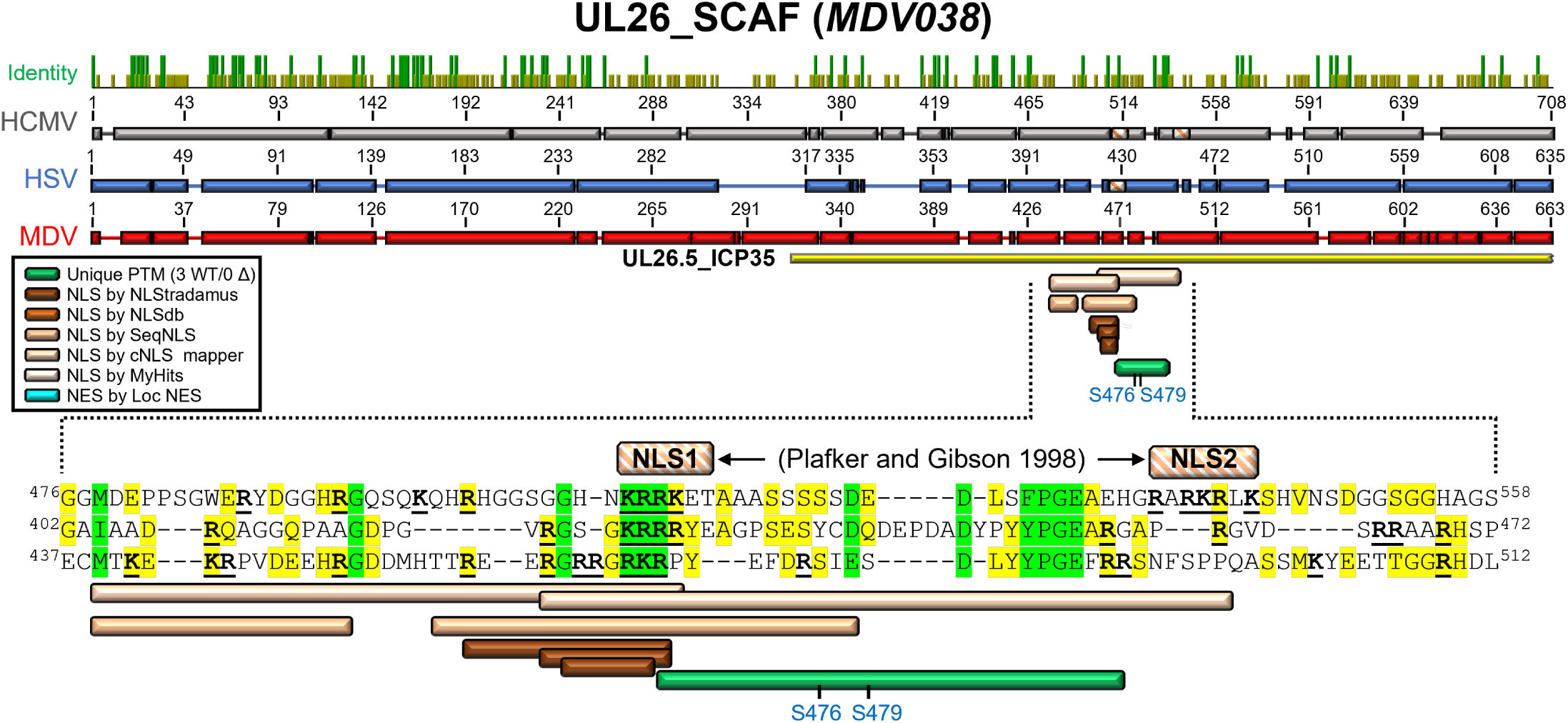
Alignment of HCMV (Q6SW62), HSV (P10210), and MDV (Q9E6P2) SCAF proteins from the UniProt Consortium [213] using MUSCLE Alignment in Geneious Prime. Regions of note are expanded to the residue level. Published and predicted NLS are shown along with unique peptides identified in this report. Peptides in which the exact residue phosphorylated could not be determined are in blue. Residues important for NLS are bolded and underlined in black. Conserved residues are based on BLOSUM62 matrix scores, with green residues indicating 80-100% similarity and yellow residues indicating 60-80% similarity. CMV SCAF NLS motifs as previously identified by Plafker and Gibson [138] are shown.

**S14 Fig.**
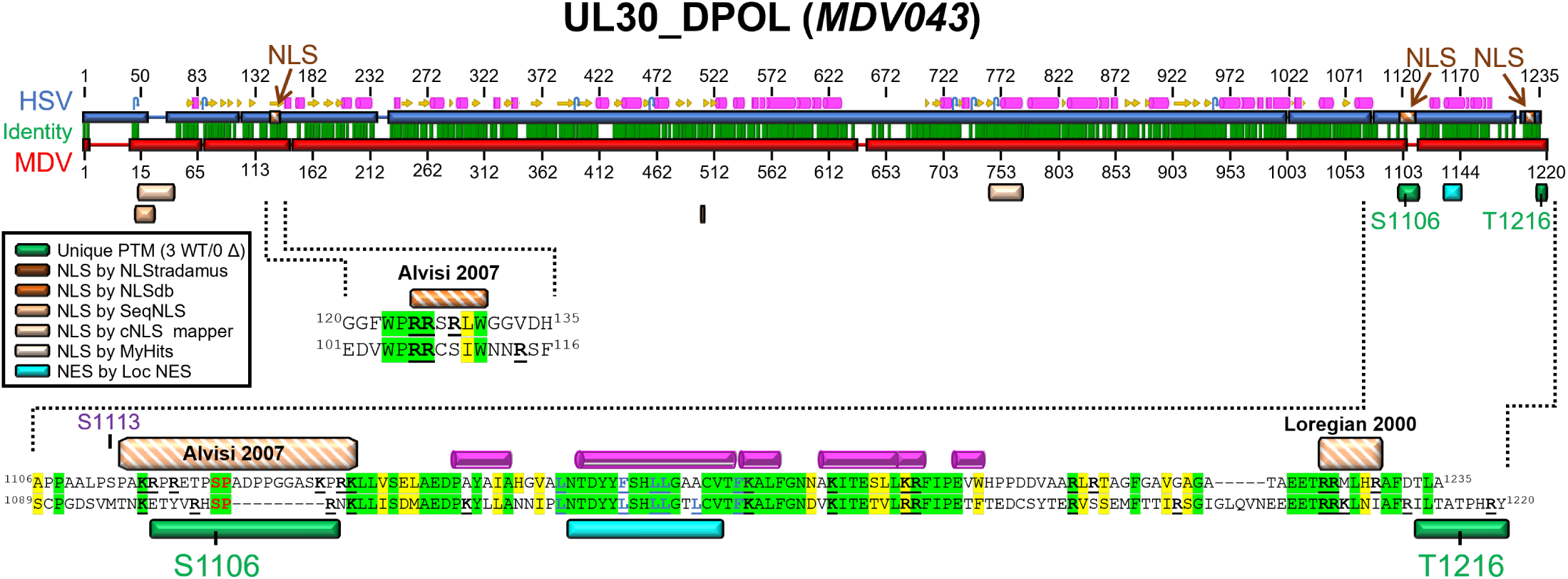
Alignment of HSV (P04293) and MDV (Q9E6N9) UL30_DPOL proteins from the UniProt Consortium [213] using MUSCLE Alignment in Geneious Prime. Regions of note are expanded to the residue level. Published and predicted NLS and NES are shown along with unique peptides identified in this report. Specific residues identified as phosphorylated are shown in green text. Residues important for NES are bolded and underlined in blue, while residues important for NLS are bolded and underlined in black. Conserved residues are based on BLOSUM62 matrix scores, with green residues indicating 80-100% similarity and yellow residues indicating 60-80% similarity. Residues in red align with the predicted consensus site for HSV CHPK phosphorylation (SP/PS). An NLS for HSV has been functionally characterized by Alvisi *et al.* [141] and Loregian *et al.* [140]. Bell *et al.* [129] identified S113 (shown in purple) as phosphorylated in HSV-1-infected cells.

**S15 Fig.**
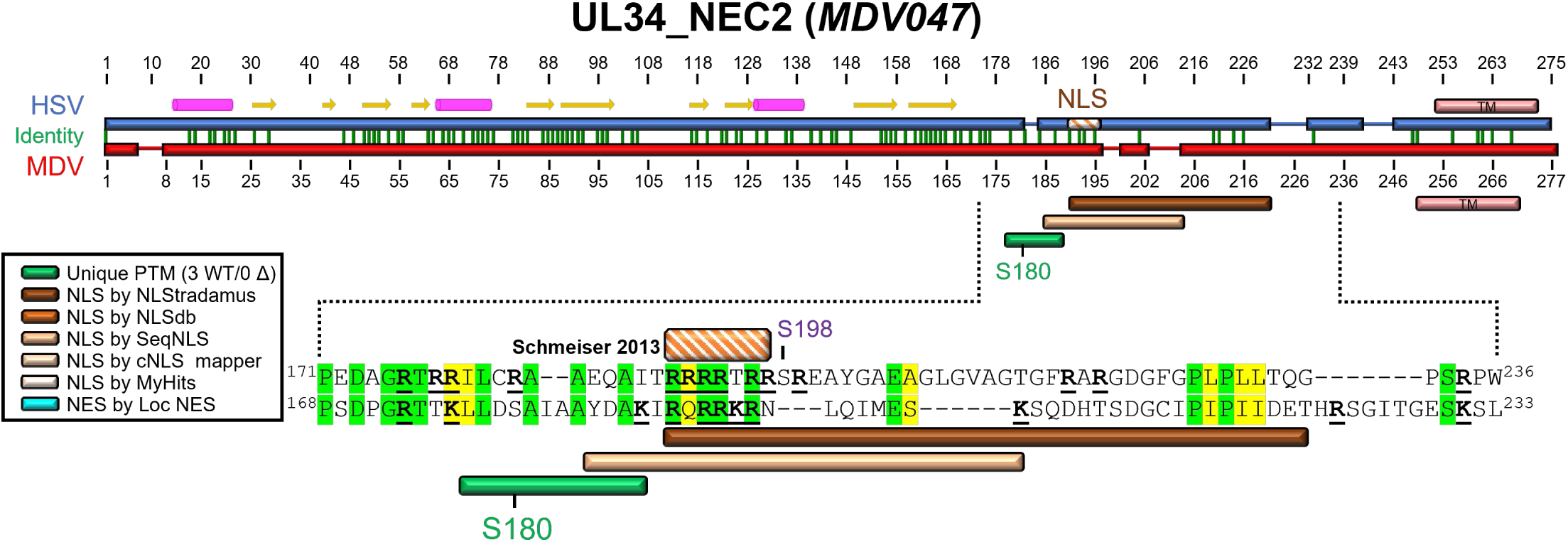
Alignment of HSV (P10218) and MDV (Q9E6N5) UL34_NEC2 proteins from the UniProt Consortium [213] using MUSCLE Alignment in Geneious Prime. Regions of note are expanded to the residue level. Published and predicted NLS and NES are shown along with unique peptides identified in this report. Specific residues identified as phosphorylated are shown in green text. Residues important for NES are bolded and underlined in blue, while residues important for NLS are bolded and underlined in black. Conserved residues are based on BLOSUM62 matrix scores, with green residues indicating 80-100% similarity and yellow residues indicating 60-80% similarity. An NLS for HSV has been predicted by Schmeiser *et al.* [146]. Bell *et al.* [129] reported that S198 (shown in purple) of HSV NEC2 was phosphorylated in infected cells, both in the absence and following PAA treatment.

**S16 Fig.**
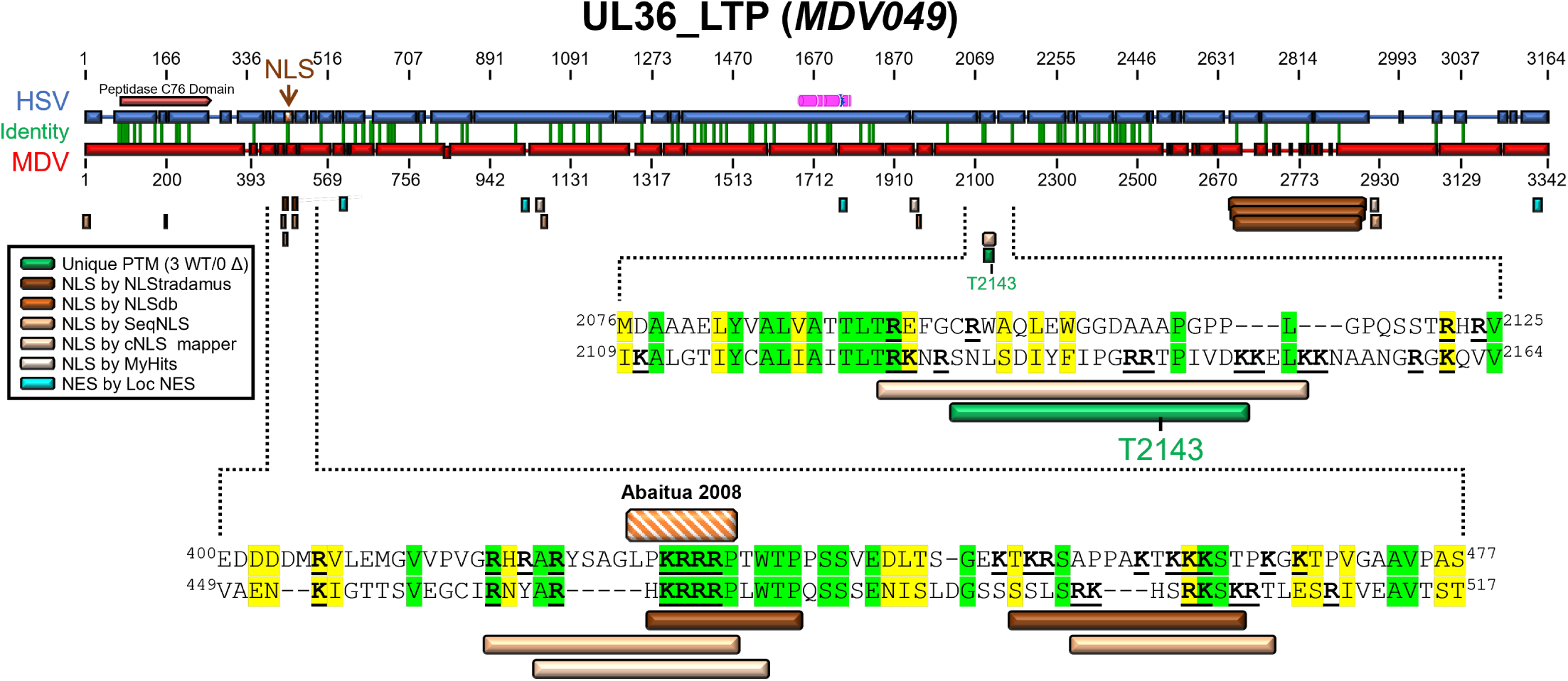
Alignment of HSV (P10220) and MDV (Q9E6N3) UL36_LTP proteins from the UniProt Consortium [213] using MUSCLE Alignment in Geneious Prime. Regions of note are expanded to the residue level. Published and predicted NLS and NES are shown along with unique peptides identified in this report. Specific residues identified as phosphorylated are shown in green text. Residues important for NLS are bolded and underlined in black. Conserved residues are based on BLOSUM62 matrix scores, with green residues indicating 80-100% similarity and yellow residues indicating 60-80% similarity. An NLS for HSV has been functionally characterized by Abaitua *et al.* [158].

**S17 Fig.**
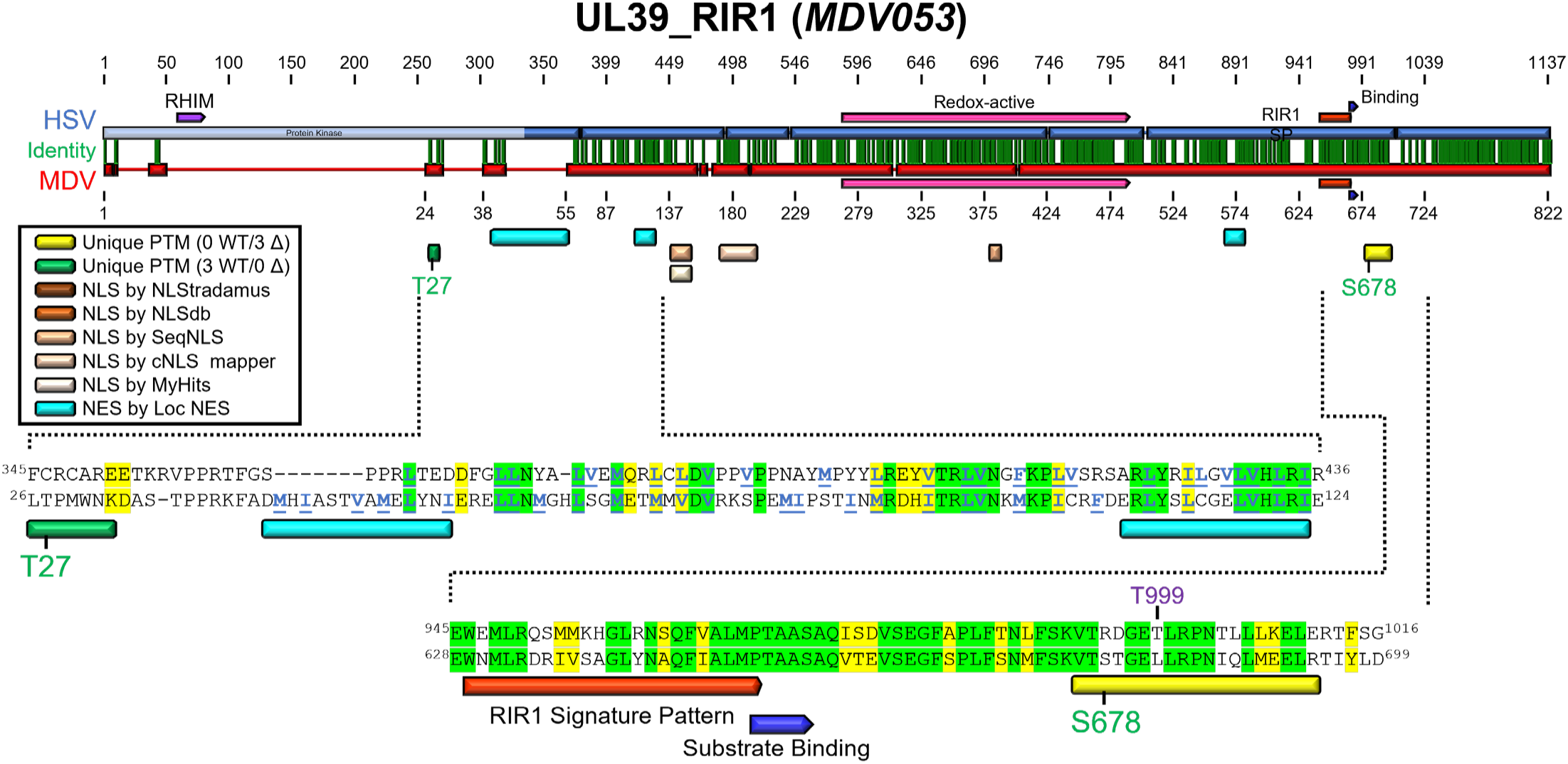
Alignment of HSV (P08543) and MDV (Q77MS1) RIR1 proteins from the UniProt Consortium [213] using MUSCLE Alignment in Geneious Prime. Regions of note are expanded to the residue level. Published and predicted NLS and NES are shown along with unique peptides identified in this report. Specific residues identified as phosphorylated in this report are shown in green text. Residues important for NES are bolded and underlined in blue. Conserved residues are based on BLOSUM62 matrix scores, with green residues indicating 80-100% similarity and yellow residues indicating 60-80% similarity. Bell *et al.* [129] identified that T999 (shown in purple) of HSV UL39_RIR1 was phosphorylated following PAA treatment.

**S18 Fig.**
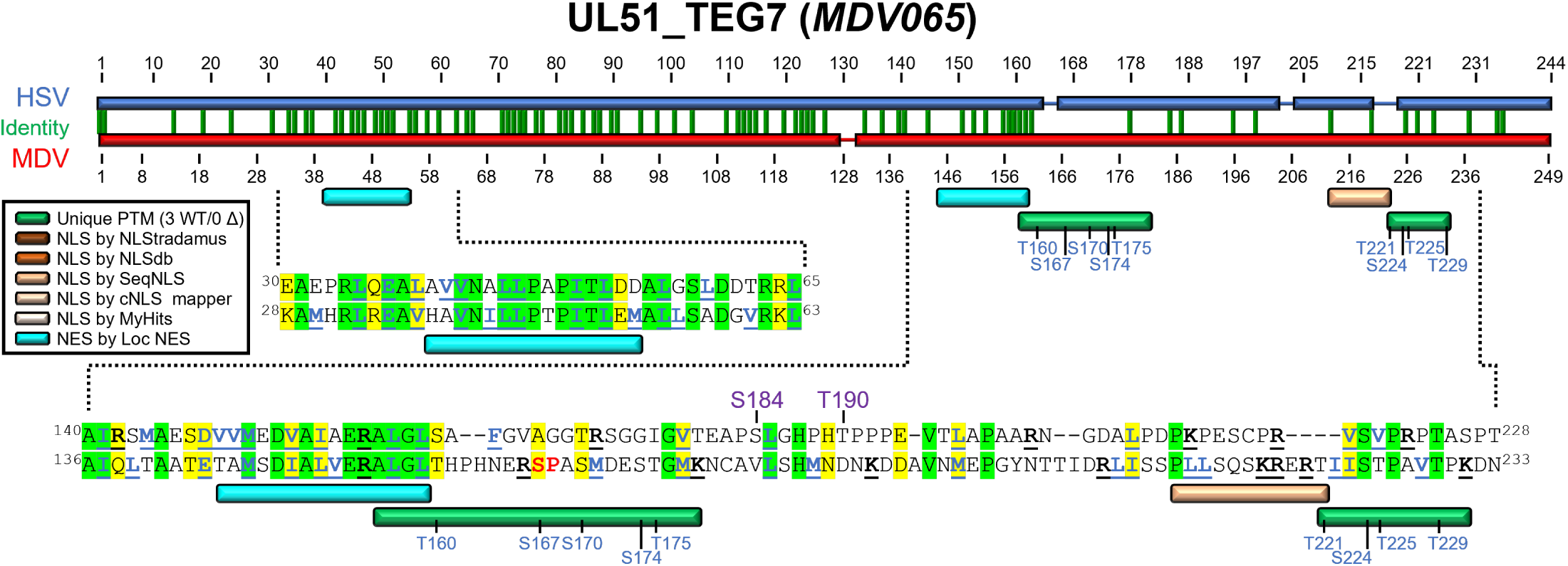
Alignment of HSV (P10235) and MDV (Q9E6M5) UL51_TEG7 proteins from the UniProt Consortium [213] using MUSCLE Alignment in Geneious Prime. Regions of note are expanded to the residue level. Predicted NLS and NES are shown along with unique peptides identified in this report. Specific residues identified as phosphorylated in this report are shown in green text, and peptides in which the exact residue phosphorylated could not be determined are in blue. Residues important for NES are bolded and underlined in blue, while residues important for NLS are bolded and underlined in black. Residues in red align with the predicted consensus site for HSV CHPK phosphorylation (SP/PS). Conserved residues are based on BLOSUM62 matrix scores, with green residues indicating 80-100% similarity and yellow residues indicating 60-80% similarity. Bell *et al.* [129] and Kato *et al.* [182] identified S184 and T190 (shown in purple) as phosphorylated in HSV-infected cells.

**S19 Fig.**
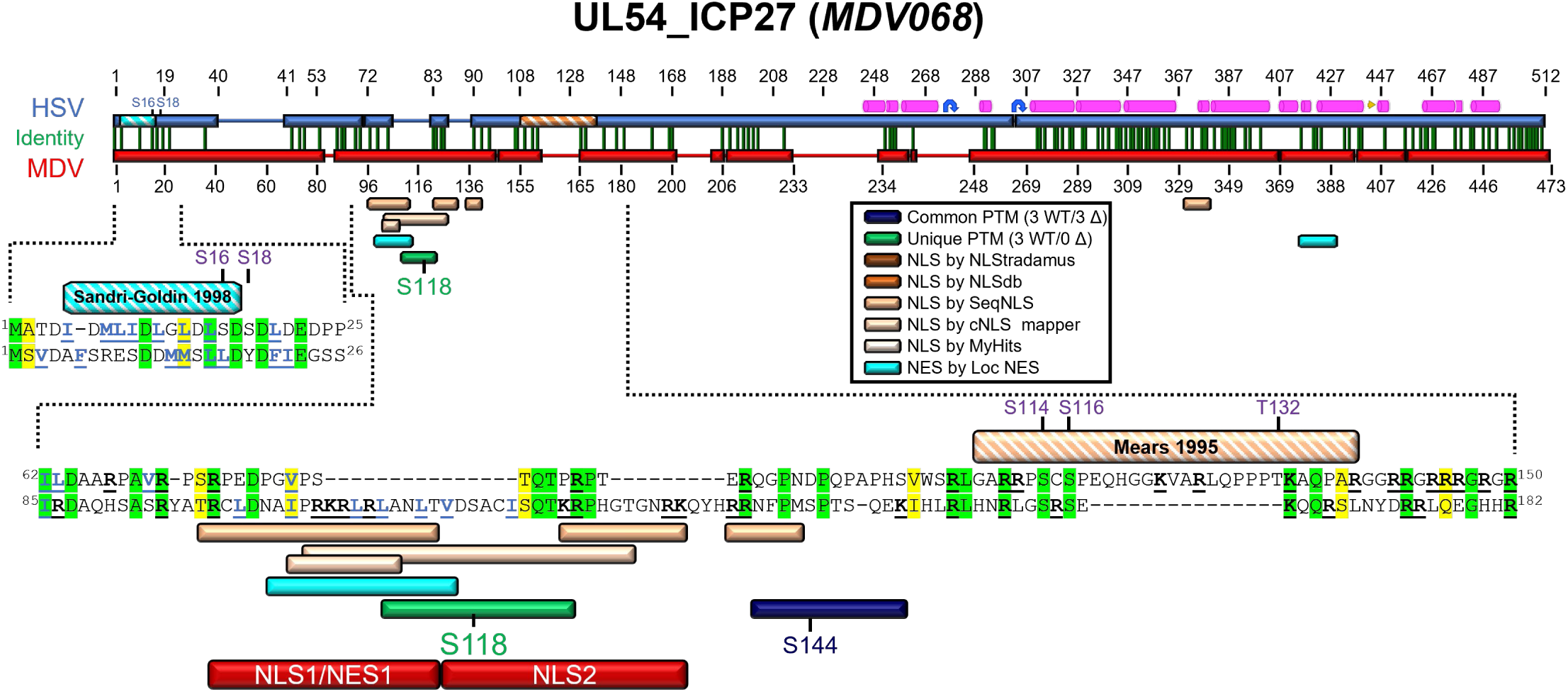
Alignment of HSV (P10238) and MDV (Q77MR3) UL54_ICP27 proteins from the UniProt Consortium [213] using MUSCLE Alignment in Geneious Prime. Regions of note are expanded to the residue level. Published and predicted NLS and NES are shown along with unique peptides identified in this report. Specific residues phosphorylated are shown in green, while peptides in which the exact residue phosphorylated could be determined are in blue, and all potential residues are shown. Residues important for NES are bolded and underlined in blue, while residues important for NLS are bolded and underlined in black. Conserved residues are based on BLOSUM62 matrix scores, with green residues indicating 80-100% similarity and yellow residues indicating 60-80% similarity. Multiple papers have described the phosphorylation of HSV ICP27, as indicated by the purple lettering. Also shown are regions deleted in subcellular localization studies. An NLS for HSV UL54_ICP27 has been functionally characterized by Mears *et al.* [63], and an NES was defined by Sandri-Goldin *et al.* [190].

**S1 Table.**
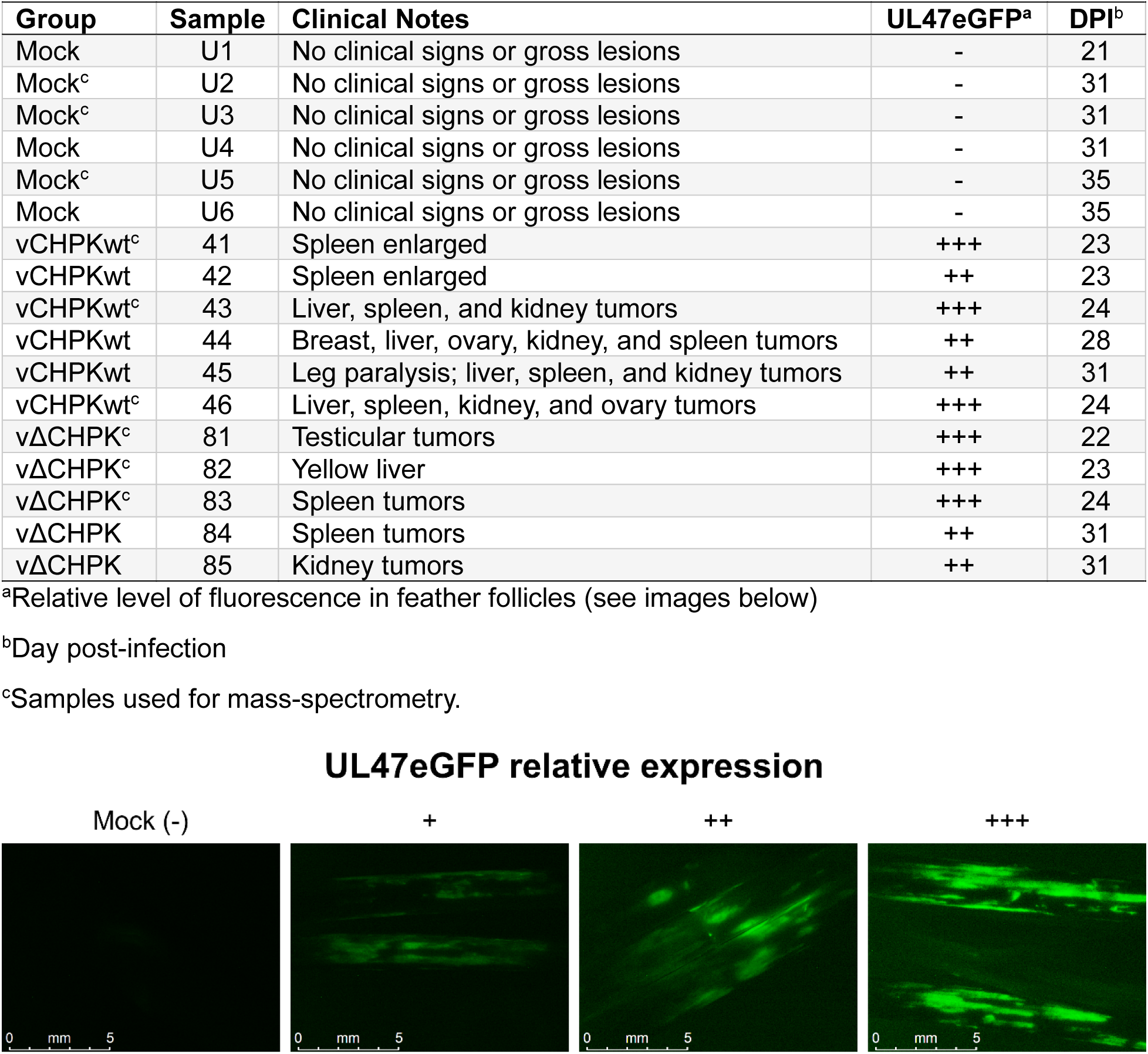
Summary of relative infection level in feather follicles used in the experimental plan.

**S2 Table.**
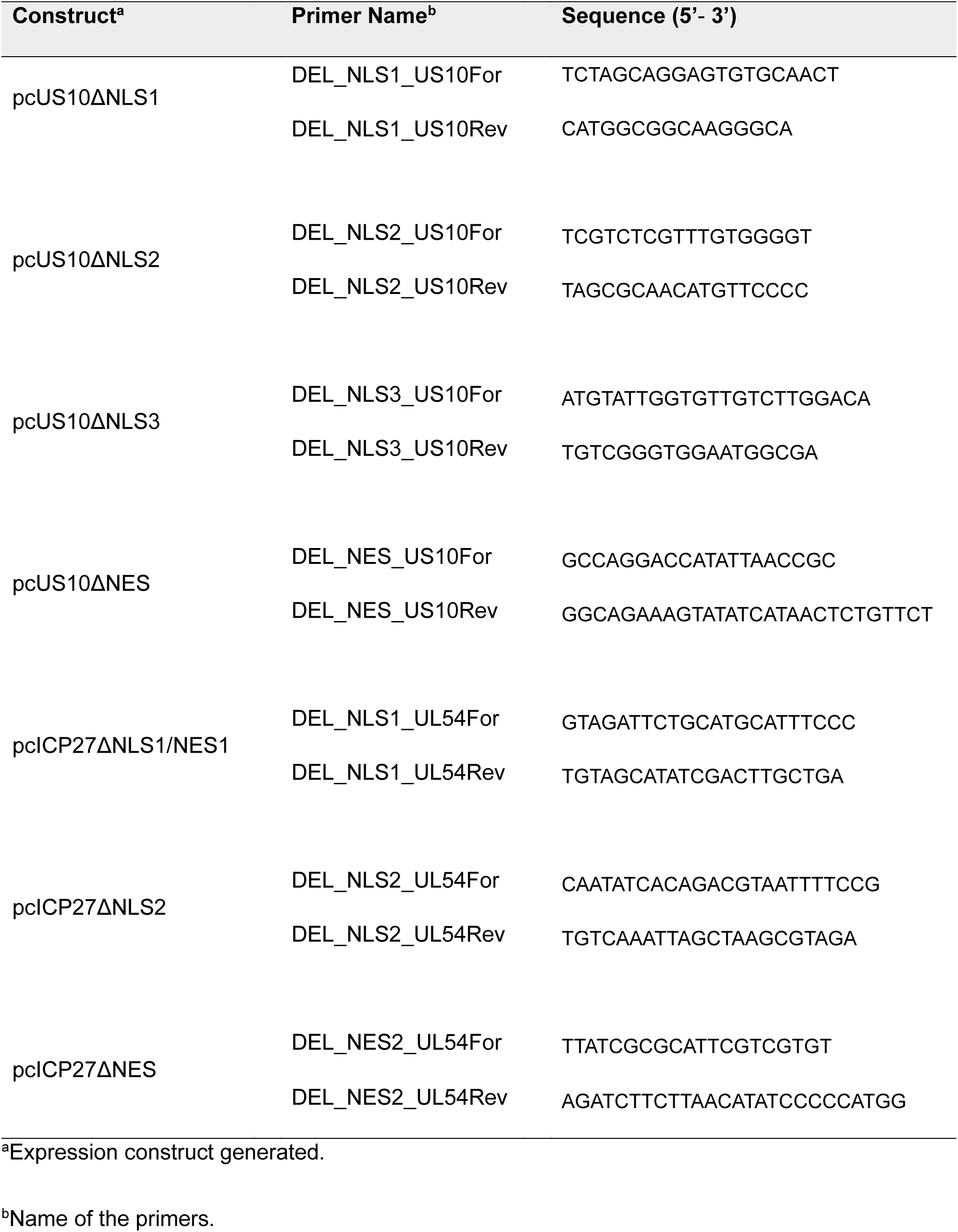
Primers used to generate mutations in expression plasmids.

**S1 File. RNA-seq data from the spleen.**

**S2 File. RNA-seq data from epithelial skin cells.**

**S3 File. LC/MS-MS data from epithelial skin cells.**

**S4 File. Specific peptide profiles and their confidence scores.**

**S5 File. Unique phosphopeptide MS2 spectra.**

## Notes

### Competing Interest Statement

The authors have declared no competing interest.

