## Supplemental Figures for "Conserved herpesvirus protein kinase (CHPK)-mediated phosphorylation of viral proteins associated with nucleocytoplasmic trafficking during natural infection"

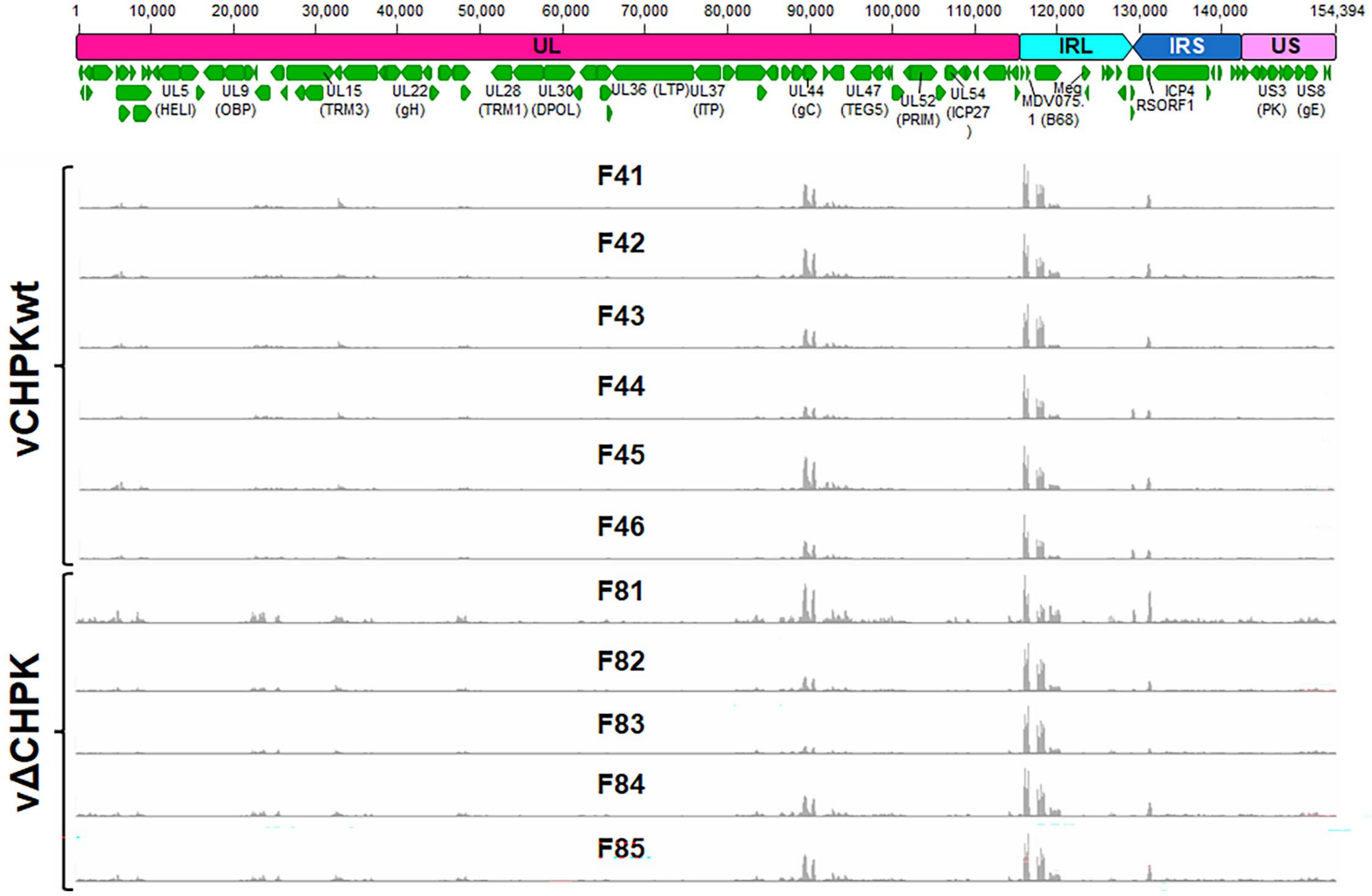


**S1 Fig. Schematic representation of the MDV genome (TRL and TRS trimmed) and RNA-seq read depth from vCHPKwt- (F41-46) and vΔCHPK- (F81-85) infected skin tissues.** Read depth tracks were generated using the Integrative Genomics Viewer (IGV) version 2.19.7 [212] for all infected skin (feather) samples.

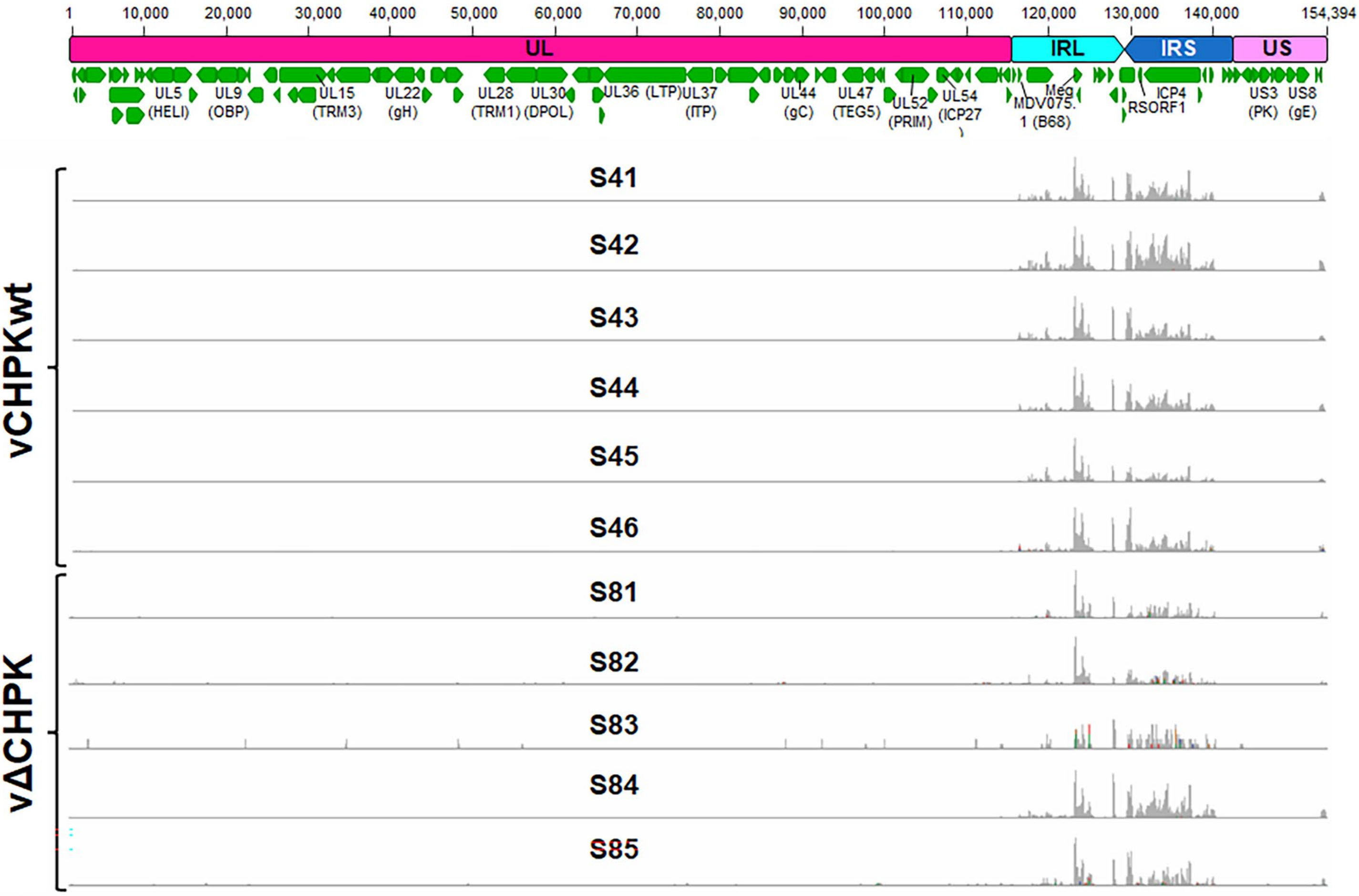


**S2 Fig. Schematic representation of the MDV genome (TRL and TRS trimmed) and RNA-seq read depth from vCHPKwt- (S41-46) and vΔCHPK- (S81-S85) infected spleens.** Read depth tracks were generated using the Integrative Genomics Viewer version 2.19.7 [212] for all spleen samples.

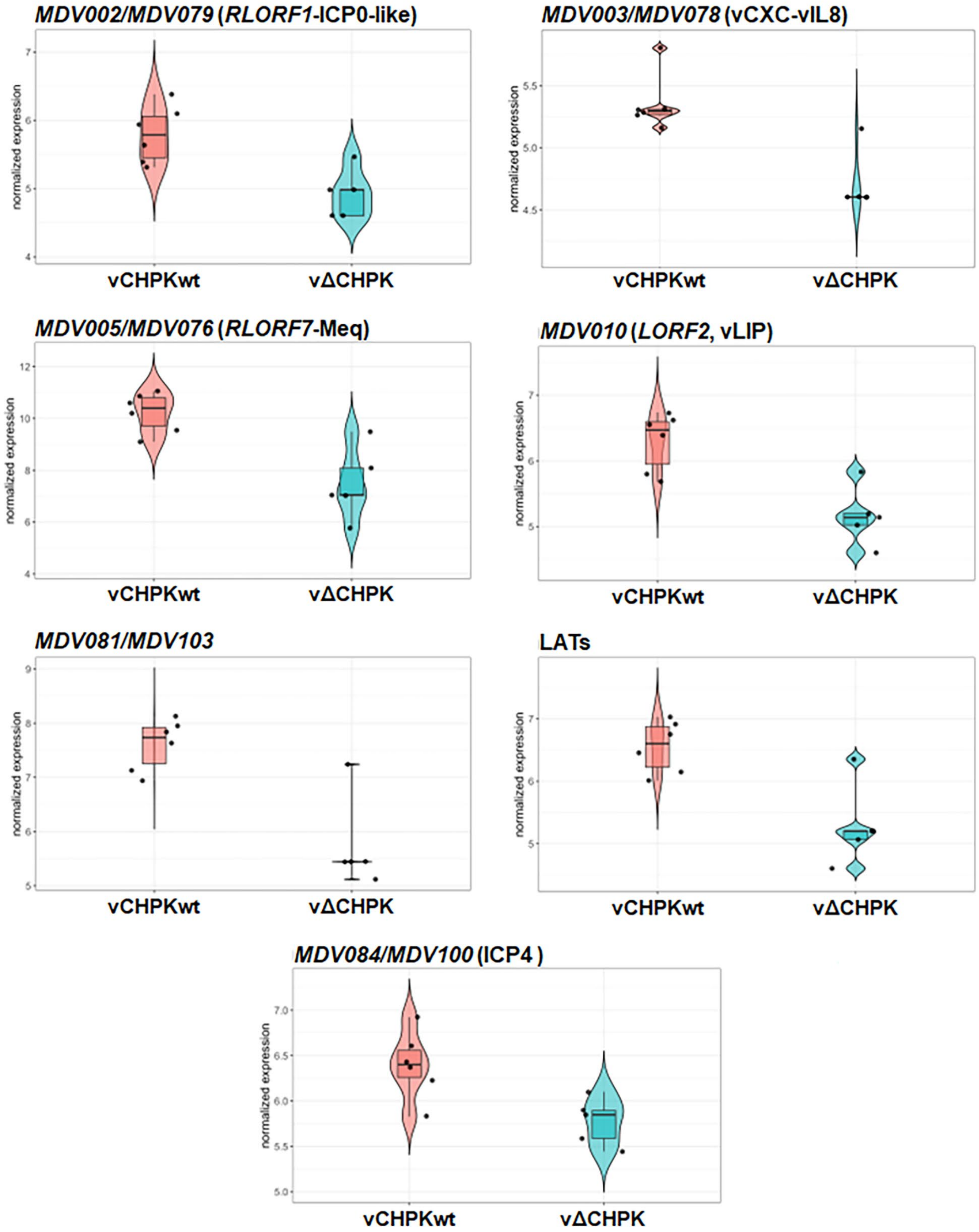


**S3 Fig. Differentially expressed MDV transcripts in the spleen.** Seven MDV genes were differentially expressed between vCHPKwt and vΔCHPK-infected spleens at a p-adj value of <0.05.

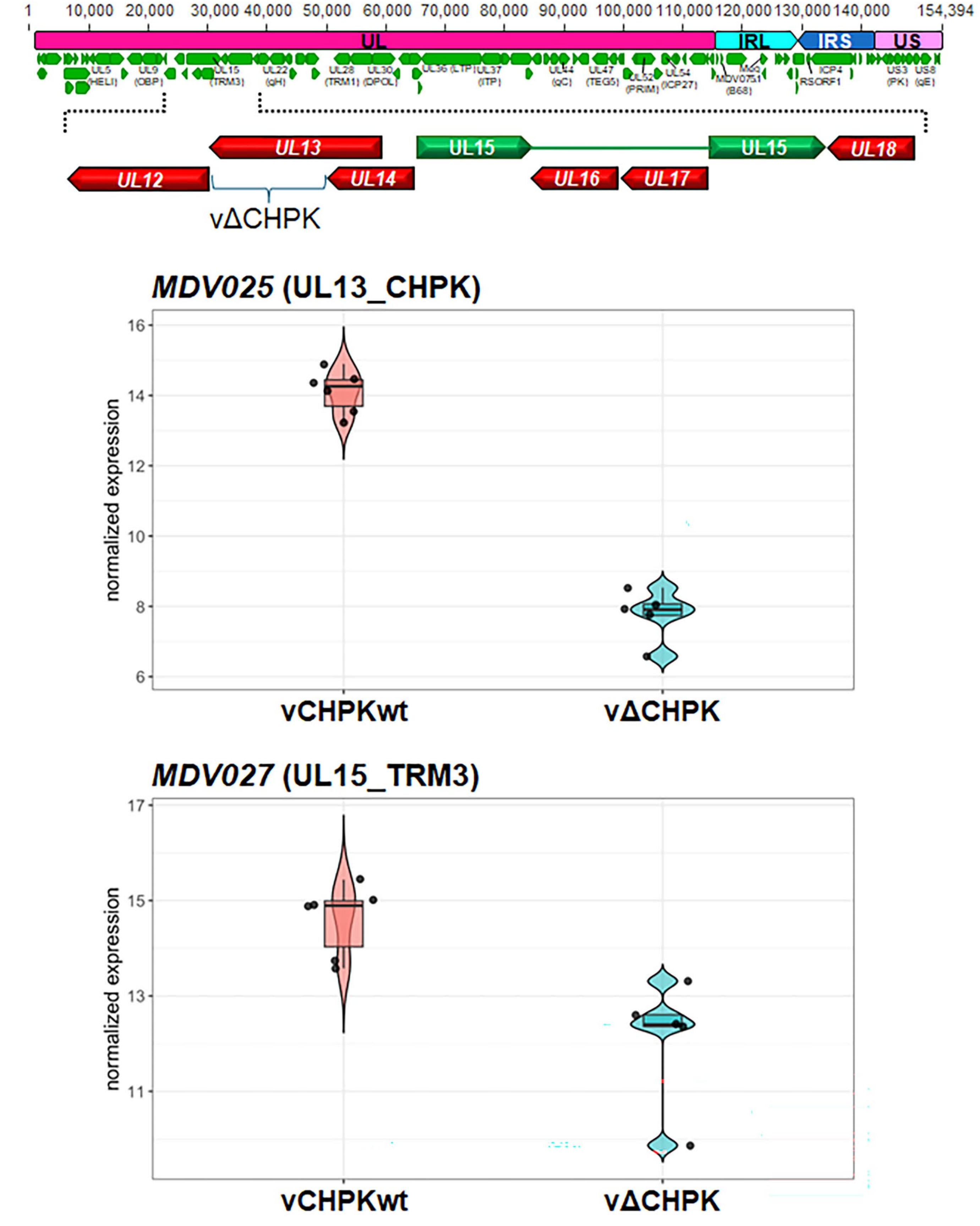


**S4 Fig. Differentially expressed MDV genes in the skin.** (A) Schematic representation of the MDV genome (with TRL and TRS trimmed) and an expanded view of the region encoding UL12-UL18. (B) Normalized gene expression of the two MDV genes differentially transcribed between vCHPKwt- and vΔCHPK-infected skins, with p-adj values < 0.05.

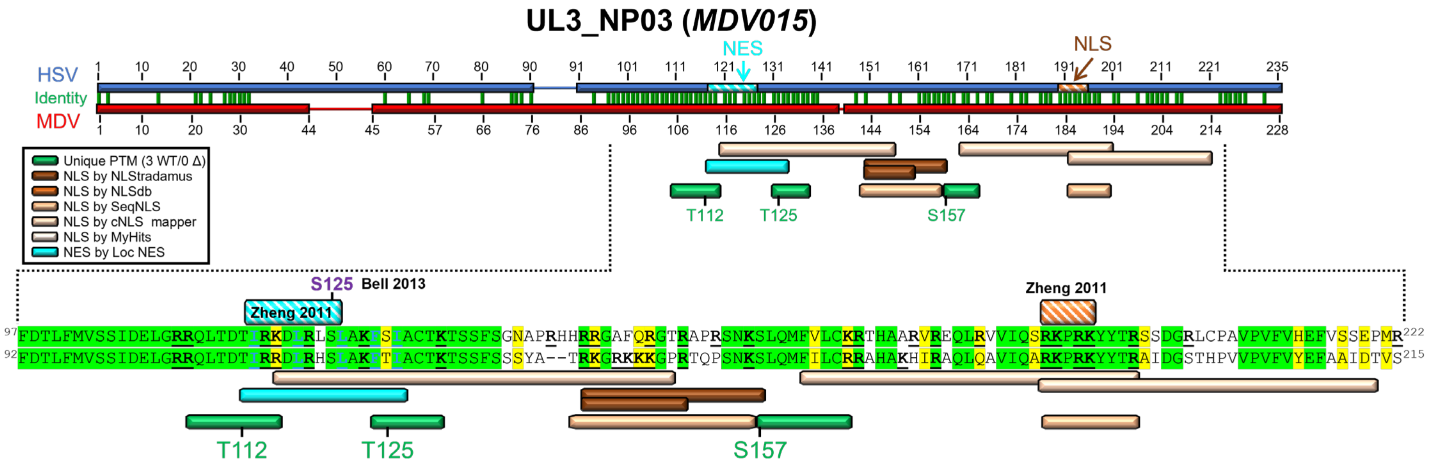


**S5 Fig. Alignment of HSV (Q1XBW5) and MDV (Q77MS7) UL3_NP03 proteins from the UniProt Consortium [213] using MUSCLE Alignment in Geneious Prime.** Regions of note are expanded to the residue level. Specific residues identified as phosphorylated are shown in green text. Residues important for NLS are bolded and underlined in black. Residues in red align with the predicted consensus site for HSV CHPK phosphorylation (SP/PS). Conserved residues are based on BLOSUM62 matrix scores, with green residues indicating 80-100% similarity and yellow residues indicating 60-80% similarity. The residue previously shown to be phosphorylated by Bell *et al.* [129] is noted in purple text. NES and NLS, as identified by Zheng *et al.* [74], are shown.

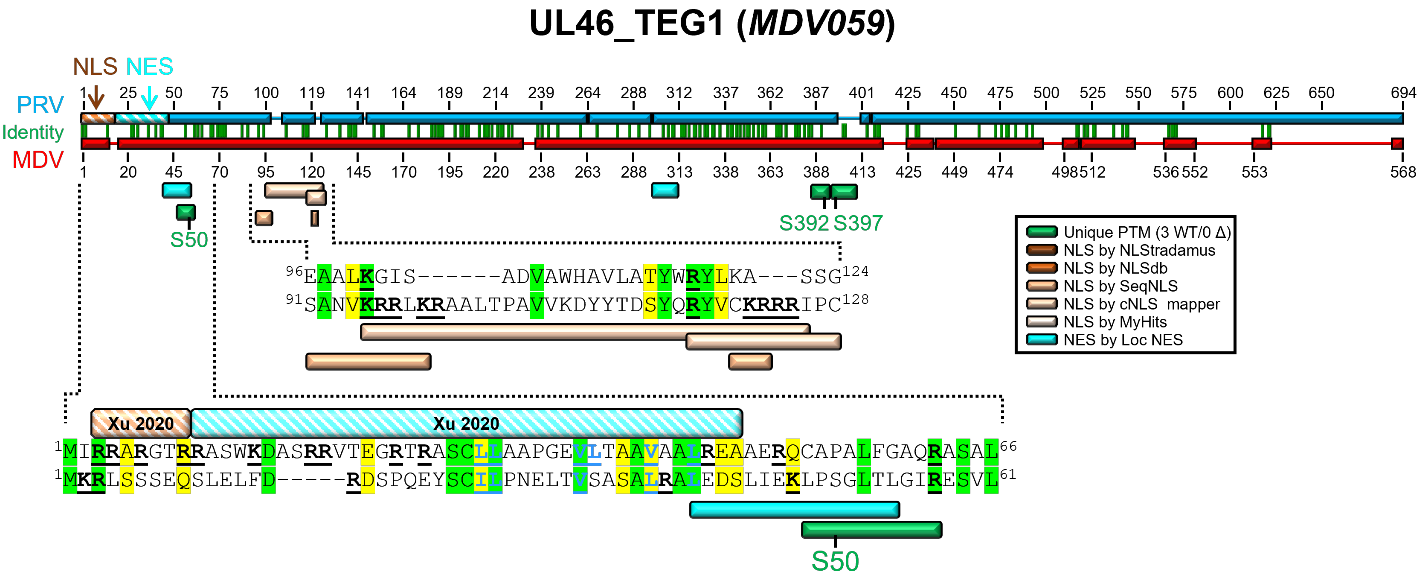


**S6 Fig. Alignment of PRV (A0A1W5LAD9) and MDV (Q77MR6) UL46_TEG1 proteins from the UniProt Consortium [213] using MUSCLE Alignment in Geneious Prime.** Regions of note are expanded to the residue level. Published and predicted NLS and NES are shown along with unique peptides identified in this report. Specific residues identified as phosphorylated are shown in green text. Residues important for NES are bolded and underlined in blue, while residues important for NLS are bolded and underlined in black. Conserved residues are based on BLOSUM62 matrix scores, with green residues indicating 80-100% similarity and yellow residues indicating 60-80% similarity. NES and NLS for PRV UL46_TEG1have been functionally characterized by Xu *et al.* [100].

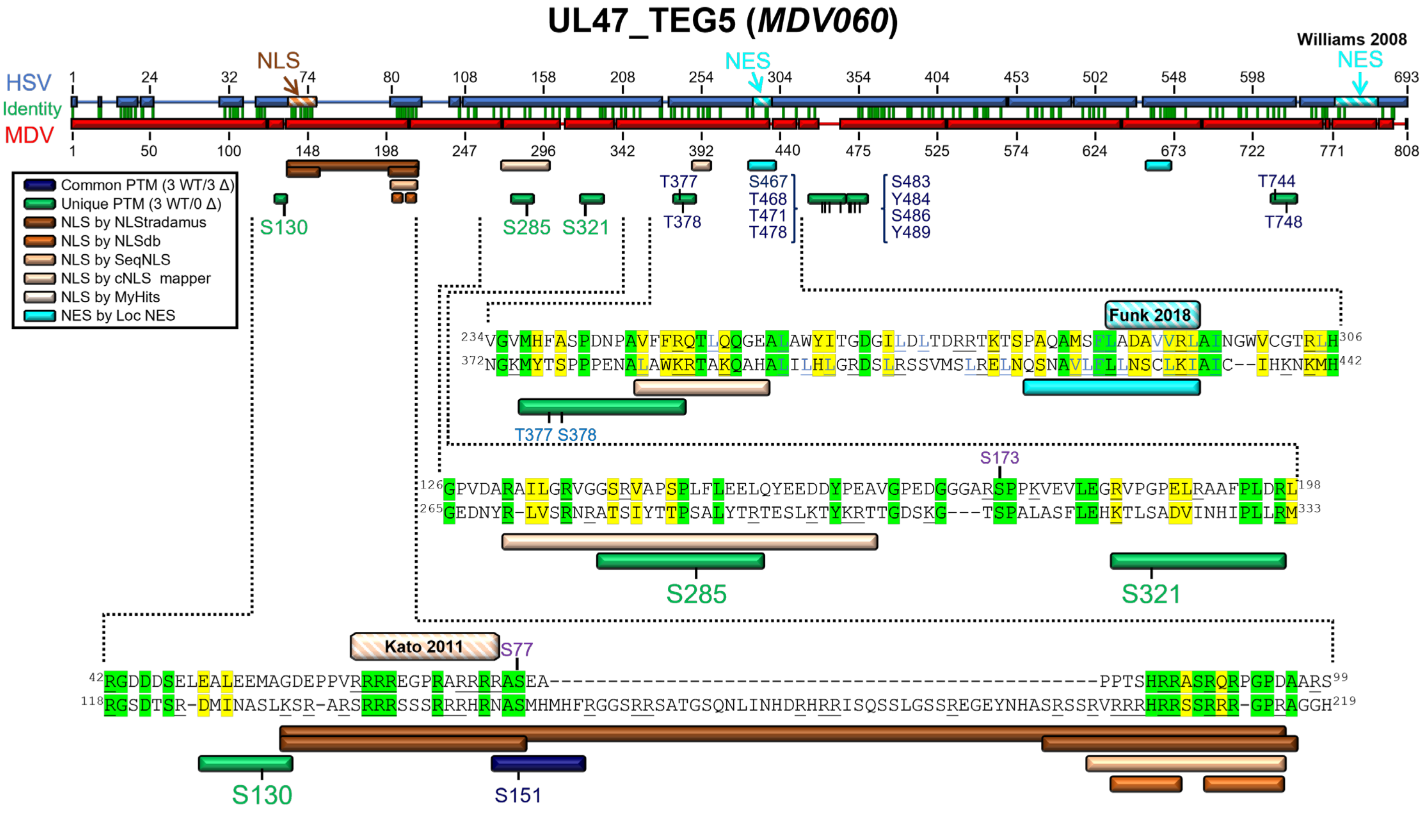


**S7 Fig. Alignment of HSV (P10231) and MDV (Q9E6M8) UL47_TEG5 proteins from the UniProt Consortium [213] using MUSCLE Alignment in Geneious Prime.** Regions of note are expanded to the residue level. Published and predicted NLS and NES are shown along with unique peptides identified in this report. Specific residues identified as phosphorylated are shown in green text. Residues important for NES are bolded and underlined in blue, while residues important for NLS are bolded and underlined in black. Conserved residues are based on BLOSUM62 matrix scores, with green residues indicating 80-100% similarity and yellow residues indicating 60-80% similarity. An NLS for HSV UL47_TEG5 has been functionally characterized by Kato *et al.* [57] and NES motifs were defined by Williams *et al.* [103] and Funk *et al*. [104].

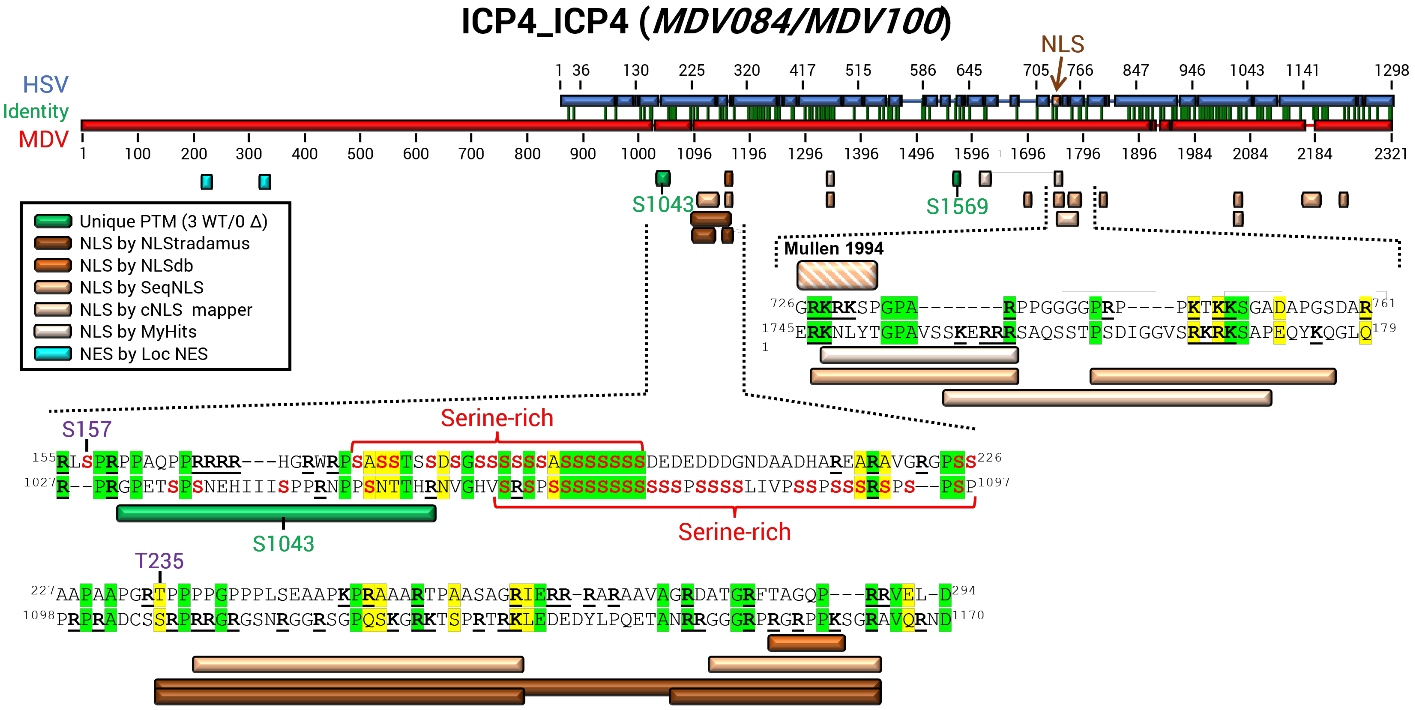


**S8 Fig Alignment of HSV (P08392) and MDV (Q9DGT6) ICP4 proteins from the UniProt Consortium [213] using MUSCLE Alignment in Geneious Prime.** Regions of note are expanded to the residue level. Published and predicted NLS and NES are shown along with unique peptides identified in this report. Specific residues identified as phosphorylated in this report are shown in green text. Residues important for NLS are bolded and underlined in black. Conserved residues are based on BLOSUM62 matrix scores, with green residues indicating 80-100% similarity and yellow residues indicating 60-80% similarity. An NLS for HSV ICP4 has been previously described by Mullen *et al.* [105]. Bell *et al.* [129] identified phosphorylated residues annotated in purple here.

### US10_US10 (*MDV089*)

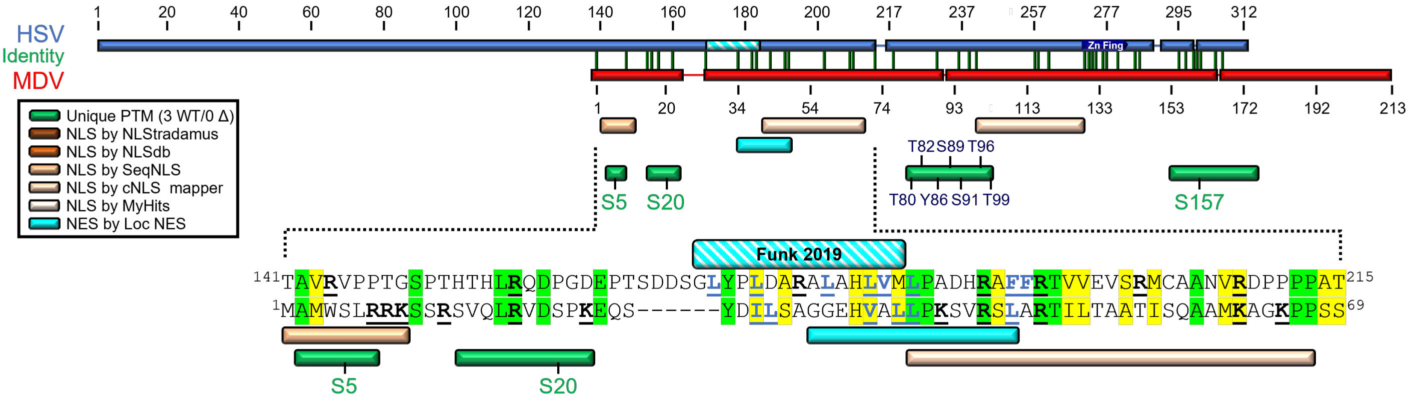


**S9 Fig. Alignment of HSV (P06486) and MDV (Q77MP8) US10 proteins from the UniProt Consortium [213] using MUSCLE Alignment in Geneious Prime.** Regions of note are expanded to the residue level. Published and predicted NLS and NES are shown along with unique peptides identified in this report. Specific residues identified as phosphorylated are shown in green text. Residues important for NLS are bolded and underlined in black, while residues important for NES are bolded and underlined in blue. Conserved residues are based on BLOSUM62 matrix scores, with green residues indicating 80-100% similarity and yellow residues indicating 60-80% similarity. An NES was predicted for HSV US10 by Funk *et al.* [104], which aligns closely with the predicted MDV US10 NES.

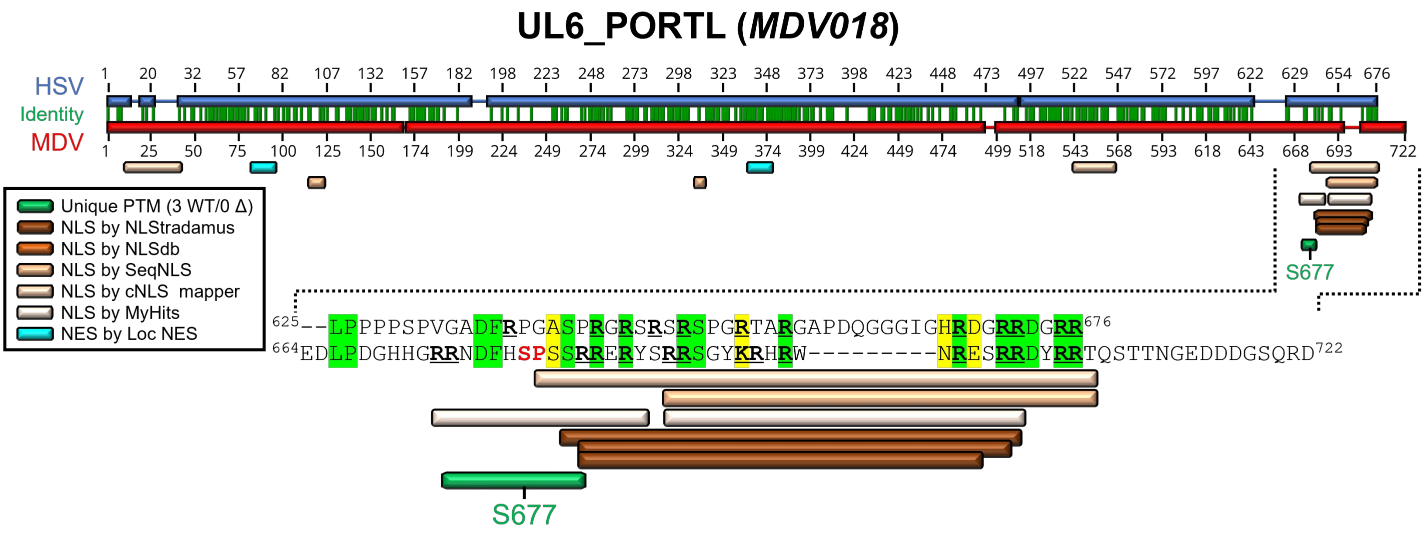


**S10 Fig. Alignment of HSV (P10190) and MDV (Q9E6R0) UL6_PORTL proteins from the UniProt Consortium [213] using MUSCLE Alignment in Geneious Prime.** Regions of note are expanded to the residue level. Specific residues identified as phosphorylated are shown in green text. Residues important for NLS are bolded and underlined in black. Residues in red align with the predicted consensus site for HSV CHPK phosphorylation (SP/PS). Conserved residues are based on BLOSUM62 matrix scores, with green residues indicating 80-100% similarity and yellow residues indicating 60-80% similarity.

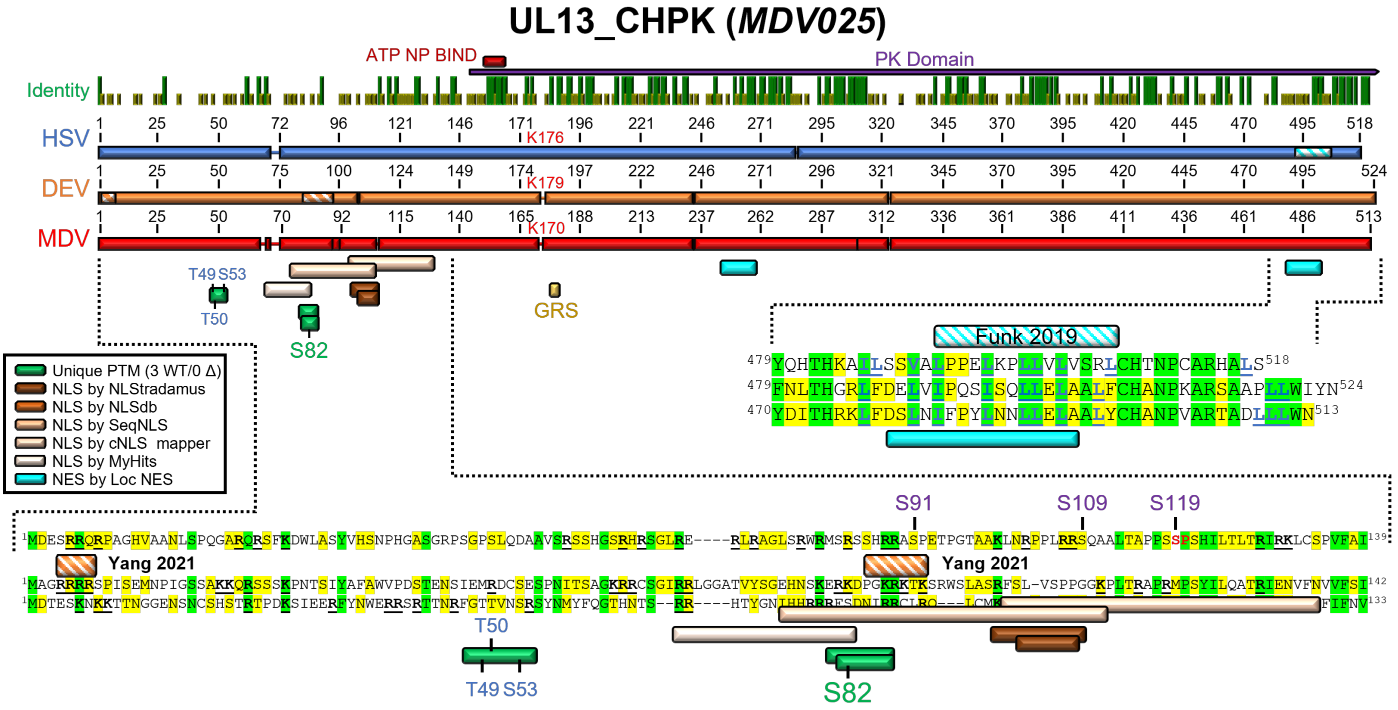


**S11 Fig. Alignment of HSV (P04290), DEV (A5A413_9ALPH), and MDV (Q9E6Q4) CHPK proteins from the UniProt Consortium [213] using MUSCLE Alignment in Geneious Prime.** Regions of note are expanded to the residue level. Published and predicted NLS and NES are shown along with unique peptides identified in this report. Specific residues identified as phosphorylated are shown in green text. Residues important for NES are bolded and underlined in blue, while residues important for NLS are bolded and underlined in black. Conserved residues are based on BLOSUM62 matrix scores, with green residues indicating 80-100% similarity and yellow residues indicating 60-80% similarity. Funk *et al.* [104] predicted an NES for HSV UL13_CHPK, while Yang *et al.* [132] identified two NLS in DEV CHPK. Residues previously shown to be phosphorylated by Koyanagi *et al.* [130] and Bell *et al.* [129] are noted in purple text.

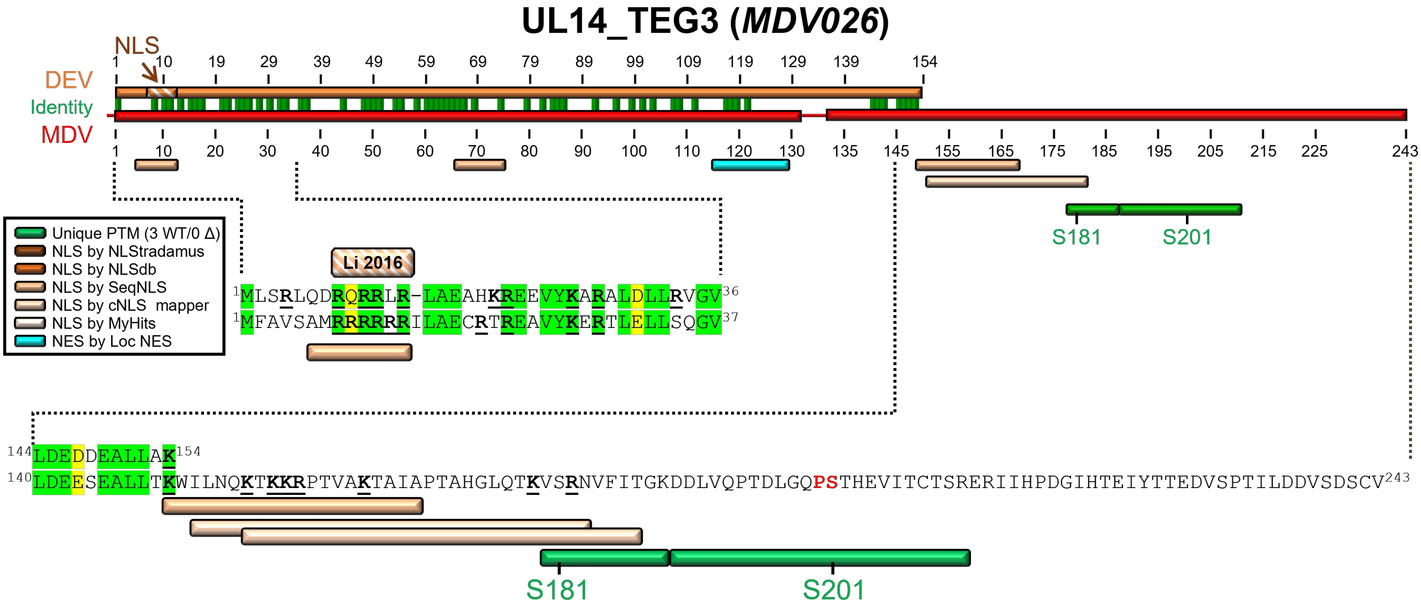


**S12 Fig. Alignment of DEV (A5A414) and MDV (Q9E6Q3) TEG3 proteins from the UniProt Consortium [213] using MUSCLE Alignment in Geneious Prime.** Regions of note are expanded to the residue level. Published and predicted NLS and NES are shown along with unique peptides identified in this report. Specific residues identified as phosphorylated are shown in green text. Residues important for NLS are bolded and underlined in black. Residues in red align with the predicted consensus site for HSV CHPK phosphorylation (SP/PS). Conserved residues are based on BLOSUM62 matrix scores, with green residues indicating 80-100% similarity and yellow residues indicating 60-80% similarity. An NLS for DEV has been characterized by Li *et al.* [54].

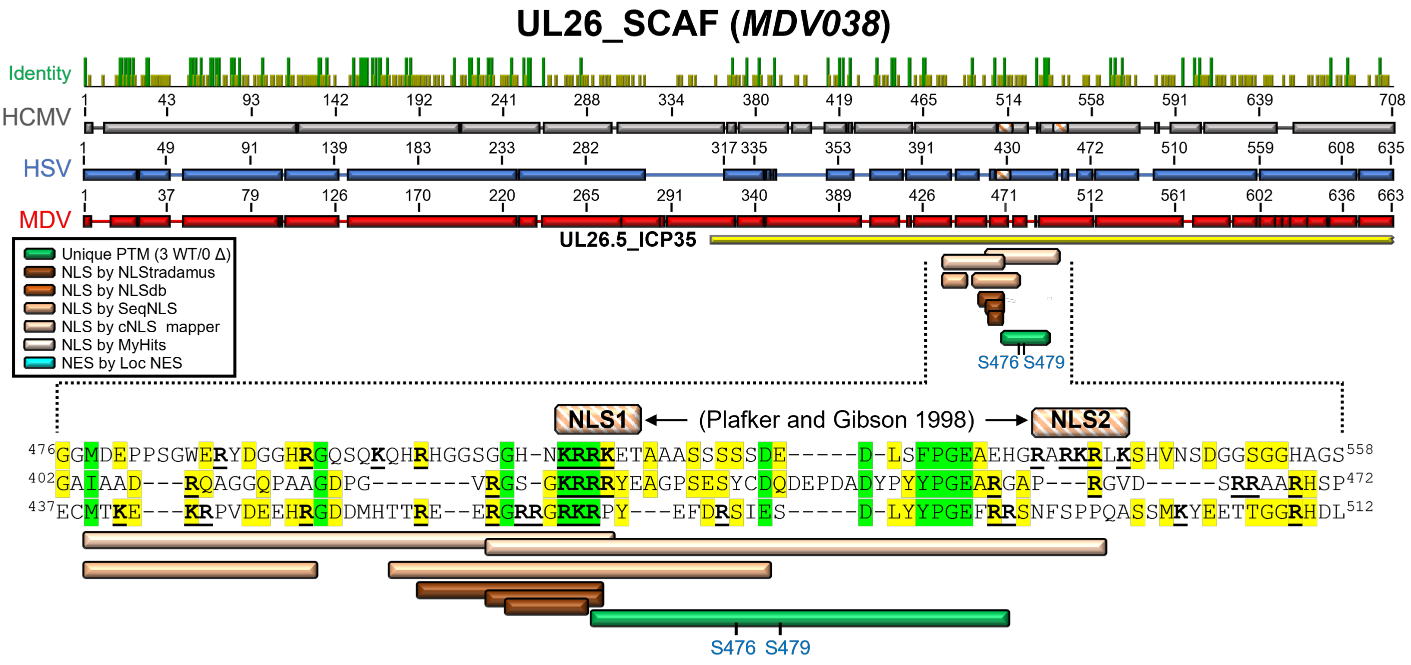


**S13 Fig. Alignment of HCMV (Q6SW62), HSV (P10210), and MDV (Q9E6P2) SCAF proteins from the UniProt Consortium [213] using MUSCLE Alignment in Geneious Prime.** Regions of note are expanded to the residue level. Published and predicted NLS are shown along with unique peptides identified in this report. Peptides in which the exact residue phosphorylated could not be determined are in blue. Residues important for NLS are bolded and underlined in black. Conserved residues are based on BLOSUM62 matrix scores, with green residues indicating 80-100% similarity and yellow residues indicating 60-80% similarity. CMV SCAF NLS motifs, as previously identified by Plafker and Gibson [138], are shown.

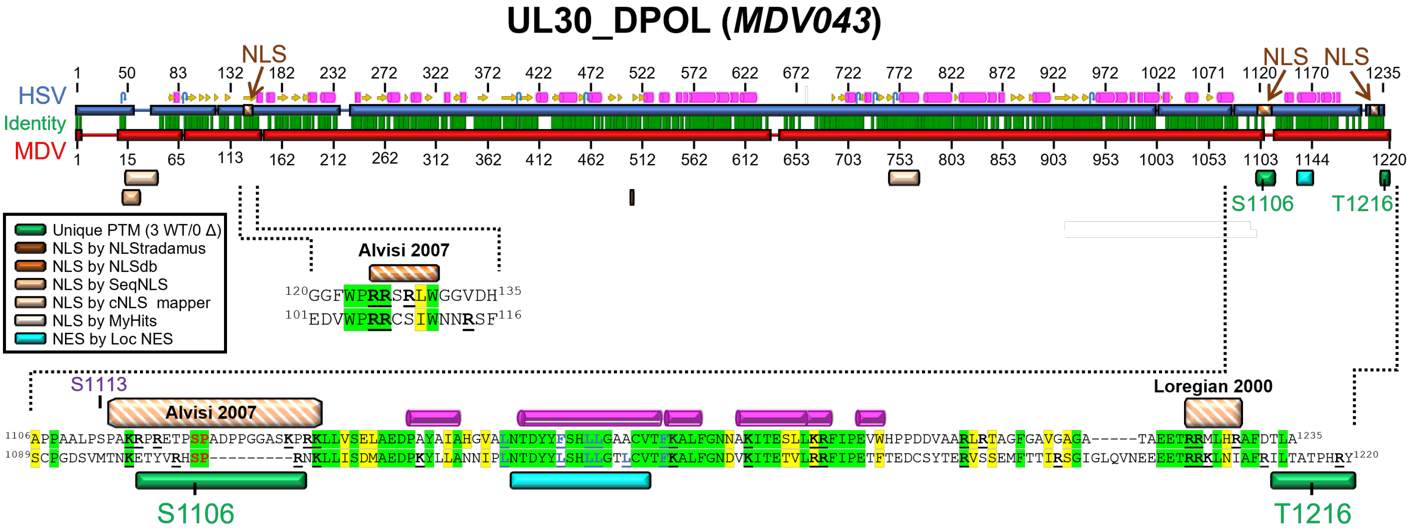


**S14 Fig. Alignment of HSV (P04293) and MDV (Q9E6N9) UL30_DPOL proteins from the UniProt Consortium [213] using MUSCLE Alignment in Geneious Prime.** Regions of note are expanded to the residue level. Published and predicted NLS and NES are shown along with unique peptides identified in this report. Specific residues identified as phosphorylated are shown in green text. Residues important for NES are bolded and underlined in blue, while residues important for NLS are bolded and underlined in black. Conserved residues are based on BLOSUM62 matrix scores, with green residues indicating 80-100% similarity and yellow residues indicating 60-80% similarity. Residues in red align with the predicted consensus site for HSV CHPK phosphorylation (SP/PS). An NLS for HSV has been functionally characterized by Alvisi *et al.* [141] and Loregian *et al.* [140]. Bell *et al.* [129] identified S113 (shown in purple) as phosphorylated in HSV-1-infected cells.

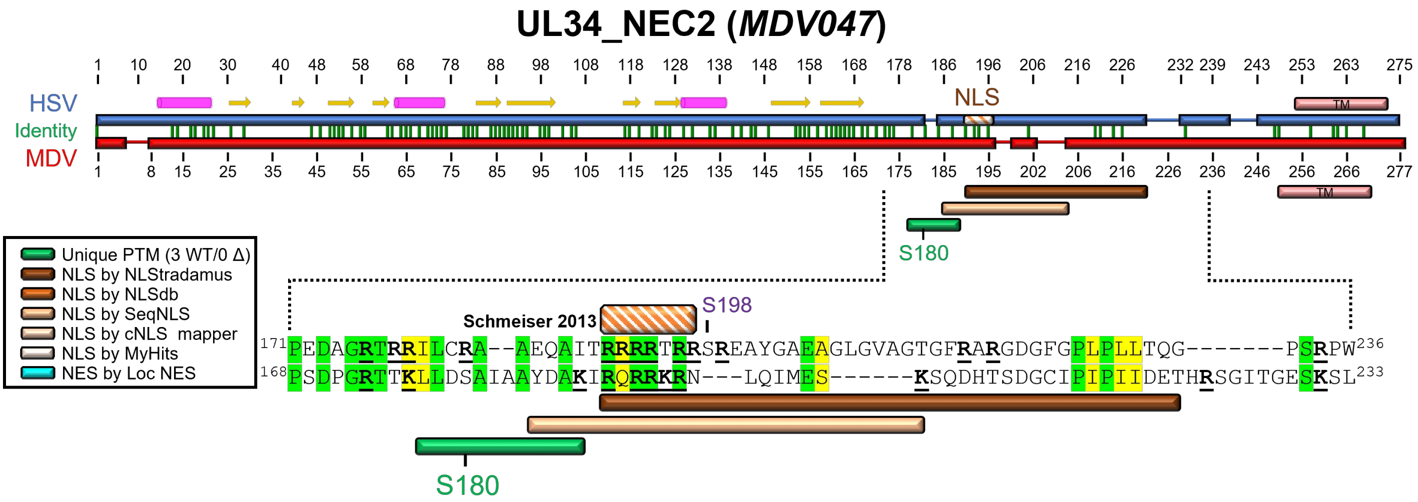


**S15 Fig. Alignment of HSV (P10218) and MDV (Q9E6N5) UL34_NEC2 proteins from the UniProt Consortium [213] using MUSCLE Alignment in Geneious Prime.** Regions of note are expanded to the residue level. Published and predicted NLS and NES are shown along with unique peptides identified in this report. Specific residues identified as phosphorylated are shown in green text. Residues important for NES are bolded and underlined in blue, while residues important for NLS are bolded and underlined in black. Conserved residues are based on BLOSUM62 matrix scores, with green residues indicating 80-100% similarity and yellow residues indicating 60-80% similarity. An NLS for HSV has been predicted by Schmeiser *et al.* [146]. Bell *et al.* [129] reported that S198 (shown in purple) of HSV NEC2 was phosphorylated in infected cells, both in the absence and following PAA treatment.

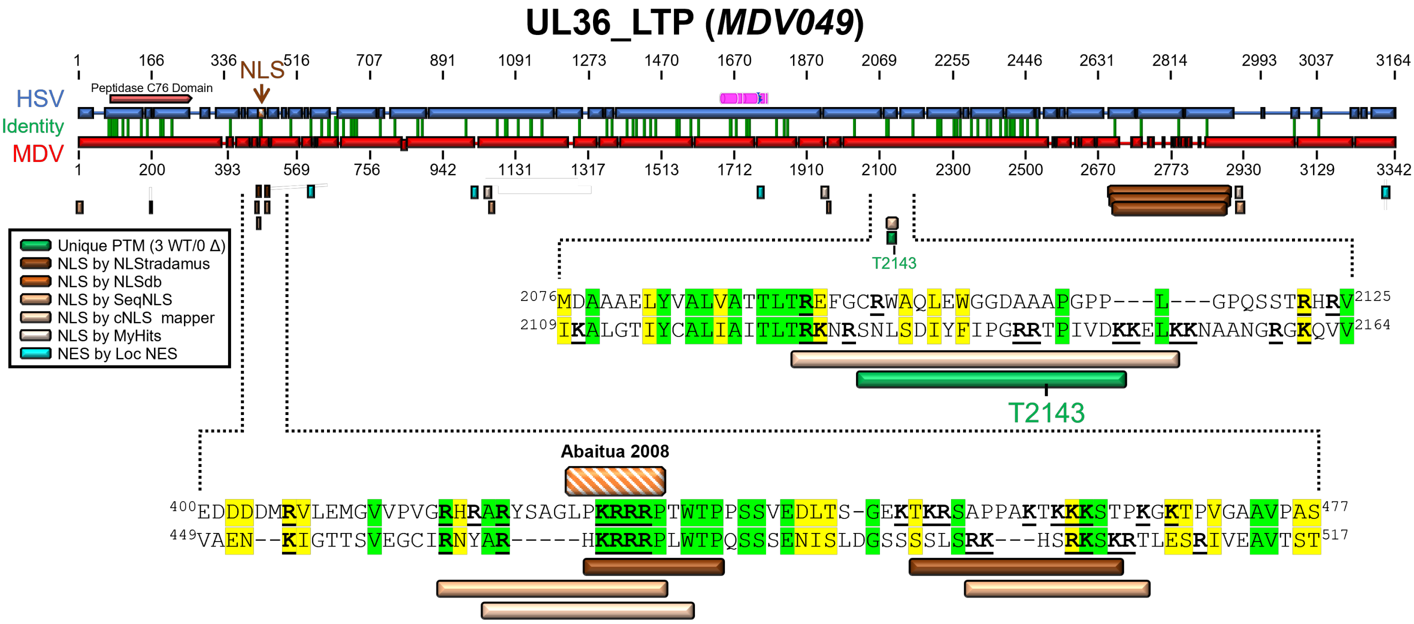


**S16 Fig. Alignment of HSV (P10220) and MDV (Q9E6N3) UL36_LTP proteins from the UniProt Consortium [213] using MUSCLE Alignment in Geneious Prime.** Regions of note are expanded to the residue level. Published and predicted NLS and NES are shown along with unique peptides identified in this report. Specific residues identified as phosphorylated are shown in green text. Residues important for NLS are bolded and underlined in black. Conserved residues are based on BLOSUM62 matrix scores, with green residues indicating 80-100% similarity and yellow residues indicating 60-80% similarity. An NLS for HSV has been functionally characterized by Abaitua *et al.* [158].

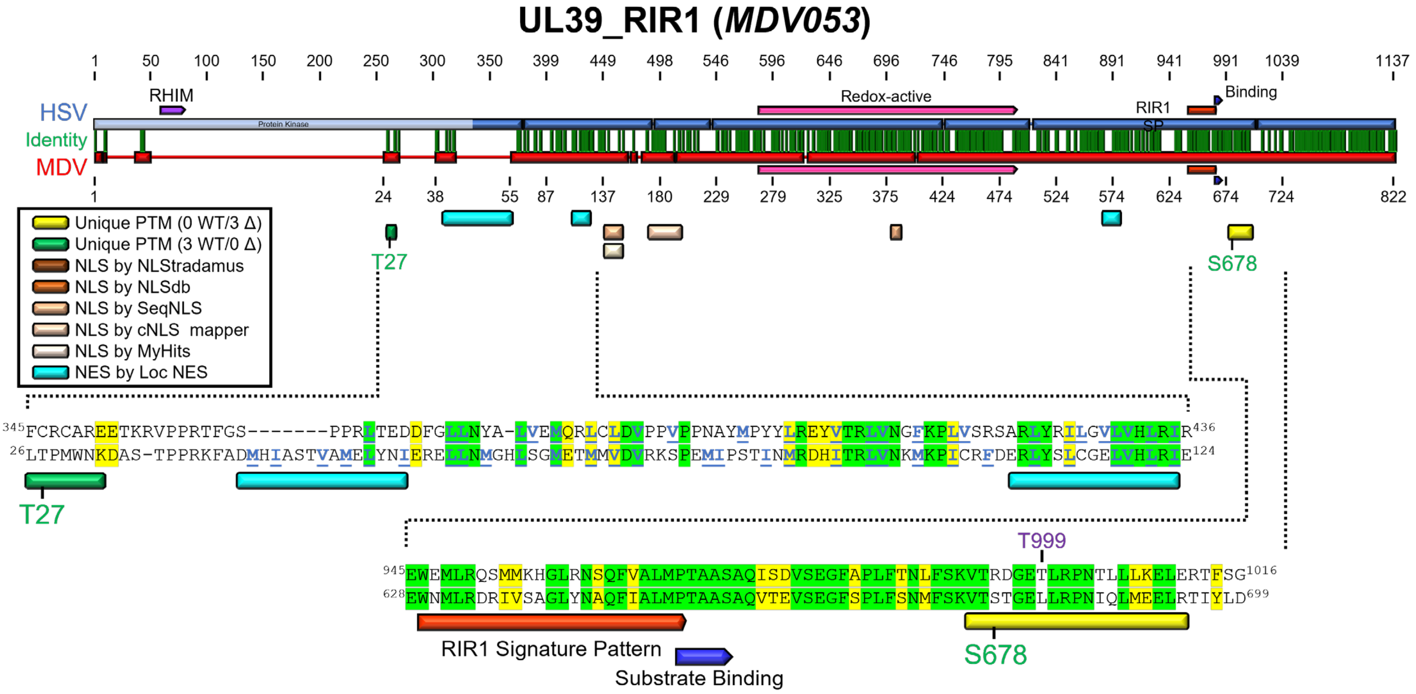


**S17 Fig. Alignment of HSV (P08543) and MDV (Q77MS1) RIR1 proteins from the UniProt Consortium [213] using MUSCLE Alignment in Geneious Prime.** Regions of note are expanded to the residue level. Published and predicted NLS and NES are shown along with unique peptides identified in this report. Specific residues identified as phosphorylated in this report are shown in green text. Residues important for NES are bolded and underlined in blue. Conserved residues are based on BLOSUM62 matrix scores, with green residues indicating 80-100% similarity and yellow residues indicating 60-80% similarity. Bell *et al.* [129] identified that T999 (shown in purple) of HSV UL39_RIR1 was phosphorylated following PAA treatment.

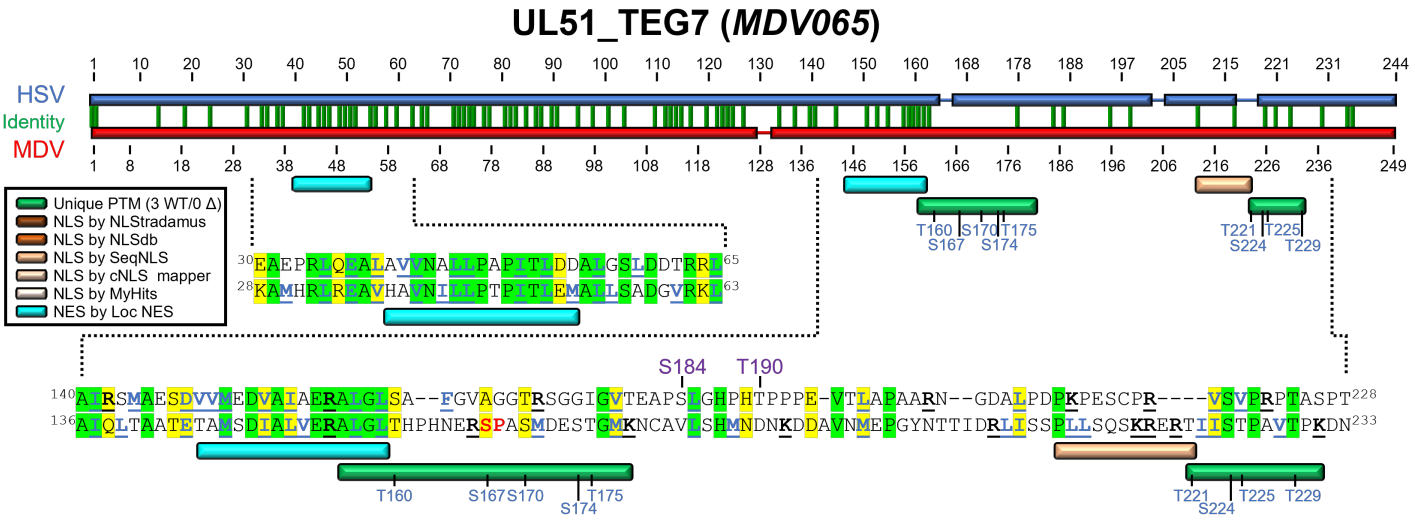


**S18 Fig. Alignment of HSV (P10235) and MDV (Q9E6M5) UL51_TEG7 proteins from the UniProt Consortium [213] using MUSCLE Alignment in Geneious Prime.** Regions of note are expanded to the residue level. Predicted NLS and NES are shown along with unique peptides identified in this report. Specific residues identified as phosphorylated in this report are shown in green text, and peptides in which the exact residue phosphorylated could not be determined are in blue. Residues important for NES are bolded and underlined in blue, while residues important for NLS are bolded and underlined in black. Residues in red align with the predicted consensus site for HSV CHPK phosphorylation (SP/PS). Conserved residues are based on BLOSUM62 matrix scores, with green residues indicating 80-100% similarity and yellow residues indicating 60-80% similarity. Bell *et al.* [129] and Kato *et al.* [182] identified S184 and T190 (shown in purple) as phosphorylated in HSV-infected cells.

### UL54_ICP27 (*MDV068*)

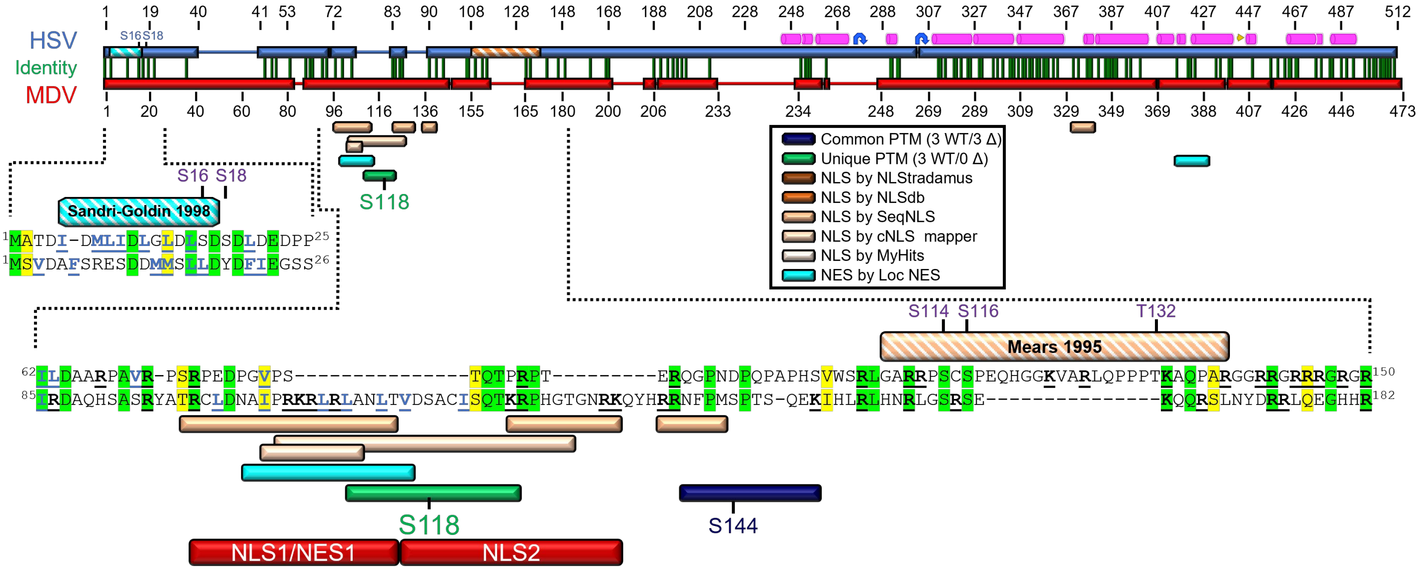


**S19 Fig. Alignment of HSV (P10238) and MDV (Q77MR3) UL54_ICP27 proteins from the UniProt Consortium [213] using MUSCLE Alignment in Geneious Prime.** Regions of note are expanded to the residue level. Published and predicted NLS and NES are shown along with unique peptides identified in this report. Specific residues phosphorylated are shown in green, while peptides in which the exact residue phosphorylated could be determined are in blue, and all potential residues are shown. Residues important for NES are bolded and underlined in blue, while residues important for NLS are bolded and underlined in black. Conserved residues are based on BLOSUM62 matrix scores, with green residues indicating 80-100% similarity and yellow residues indicating 60-80% similarity. Multiple papers have described the phosphorylation of HSV ICP27, as indicated by the purple lettering. Also shown are regions deleted in subcellular localization studies. An NLS for HSV UL54_ICP27 has been functionally characterized by Mears *et al.* [63], and an NES was defined by Sandri-Goldin *et al.* [190].
