## Supplementary figures and images for "Conserved herpesvirus protein kinase (CHPK)-mediated phosphorylation of viral proteins associated with nucleocytoplasmic trafficking during natural infection"

### DMINASLK.global.pdf

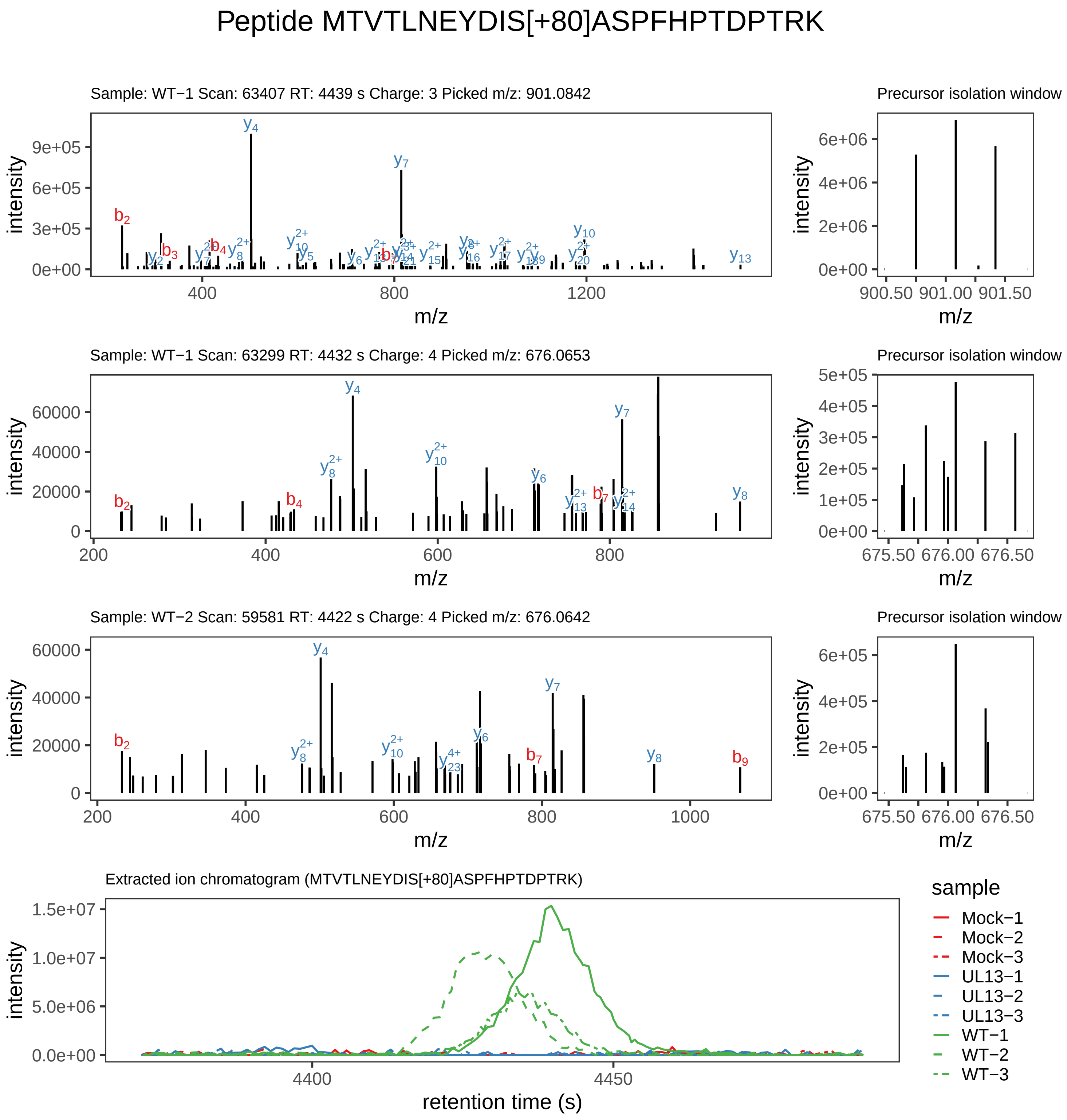

Supplement: Supplemental File 5 [file 737029_file08.zip › phosphopeptide_validation_figs_20240531/MTVTLNEYDISASPFHPTDPTRK/MTVTLNEYDIS[+80]ASPFHPTDPTRK.phospho.pdf]

#### Peptide FT[+80]IAC[+57]TK

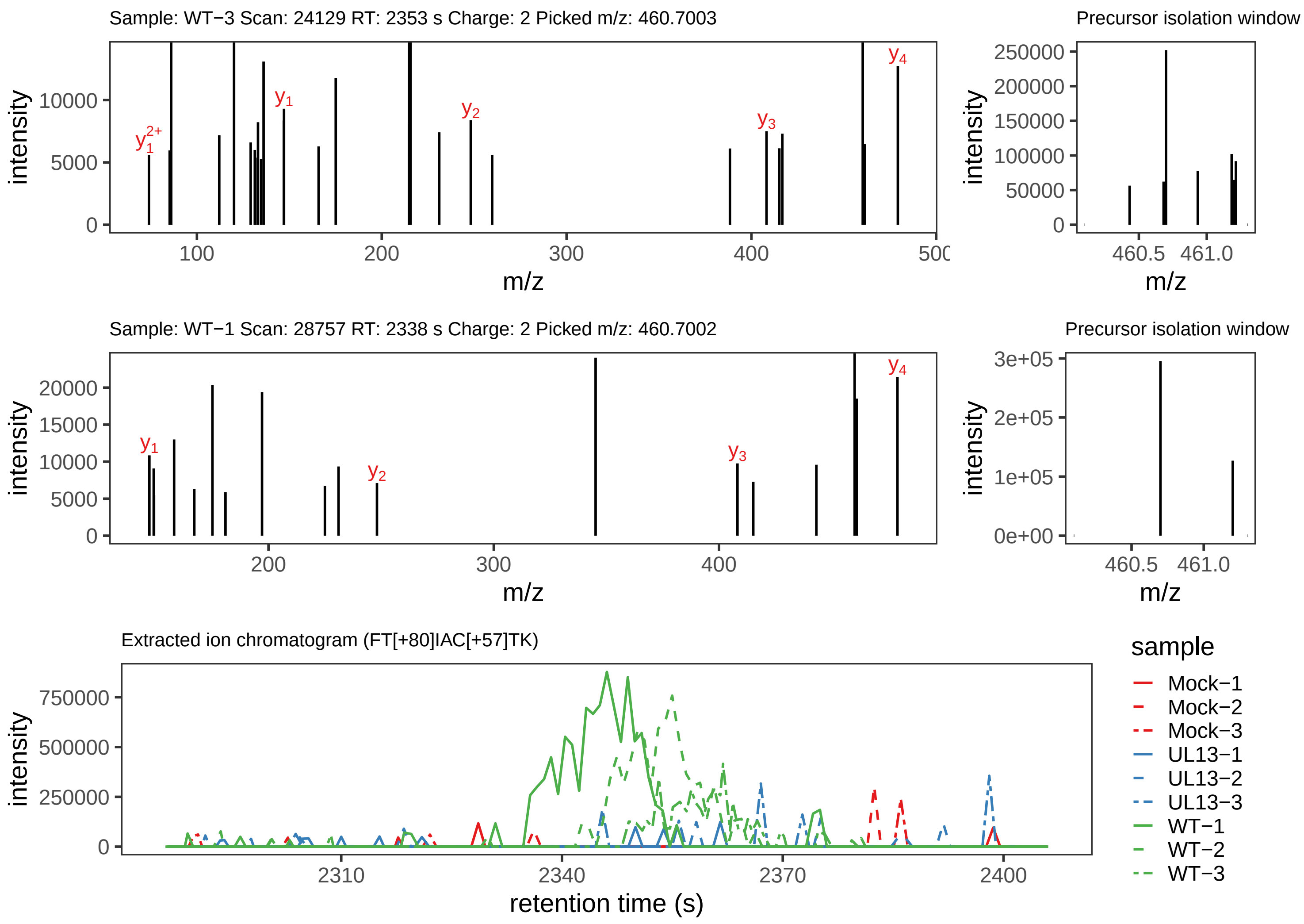

Supplement: Supplemental File 5 [file 737029_file08.zip › phosphopeptide_validation_figs_20240531/FTIACTK/FT[+80]IAC[+57]TK.phospho.pdf]

#### Peptide FTIAC[+57]TK

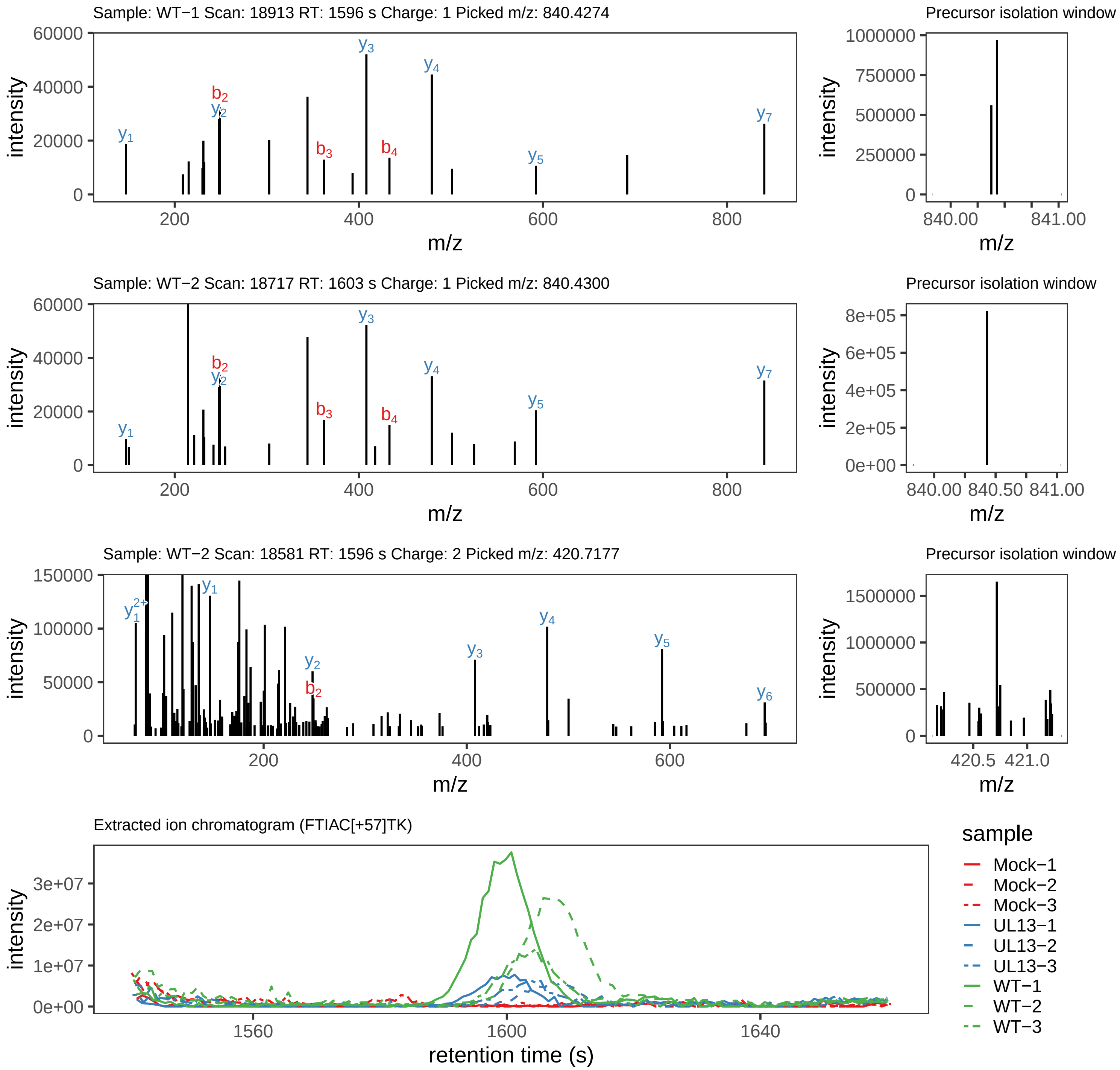

Supplement: Supplemental File 5 [file 737029_file08.zip › phosphopeptide_validation_figs_20240531/FTIACTK/FTIAC[+57]TK.global.pdf]

#### Peptide ALGLTHPHN[+1]ERS[+80]PASMDESTGM[+16]K

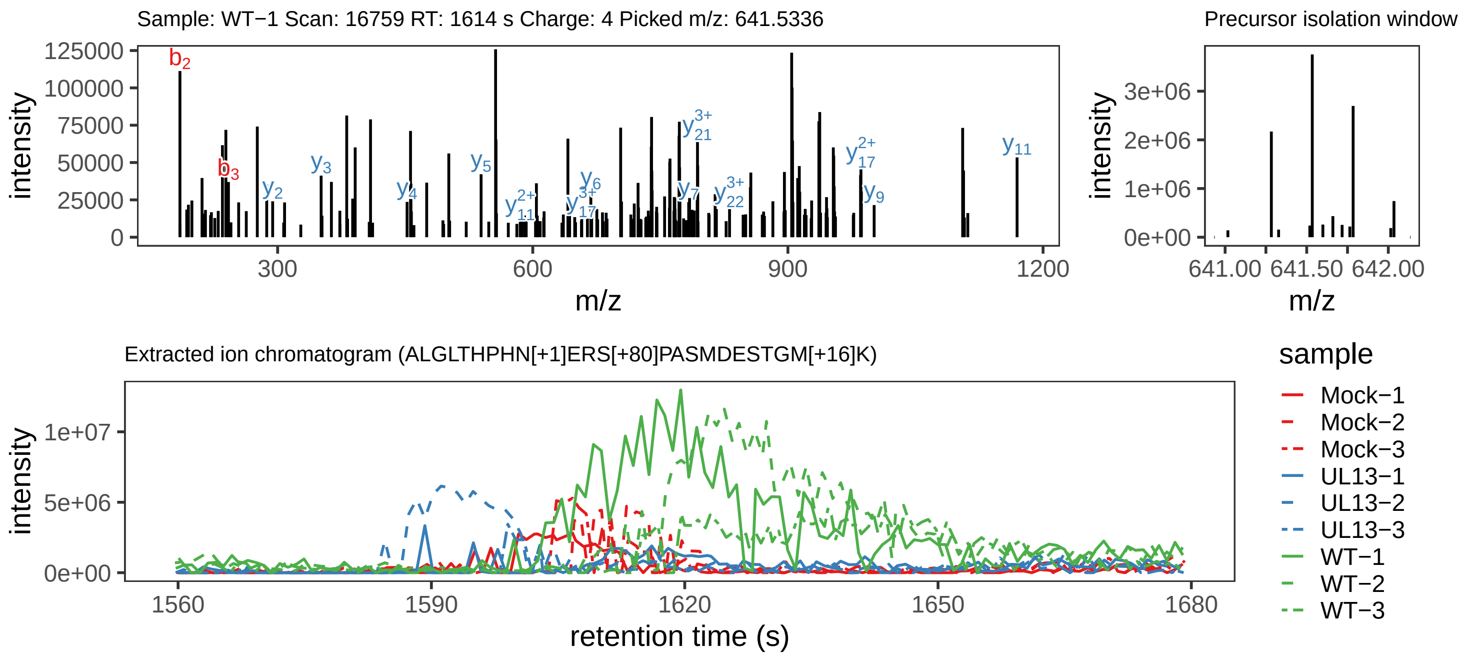

Supplement: Supplemental File 5 [file 737029_file08.zip › phosphopeptide_validation_figs_20240531/ALGLTHPHNERSPASMDESTGMK/ALGLTHPHN[+1]ERS[+80]PASMDESTGM[+16]K.phospho.pdf]

#### Peptide ALGLTHPHNERSPAS[+80]M[+16]DESTGMK

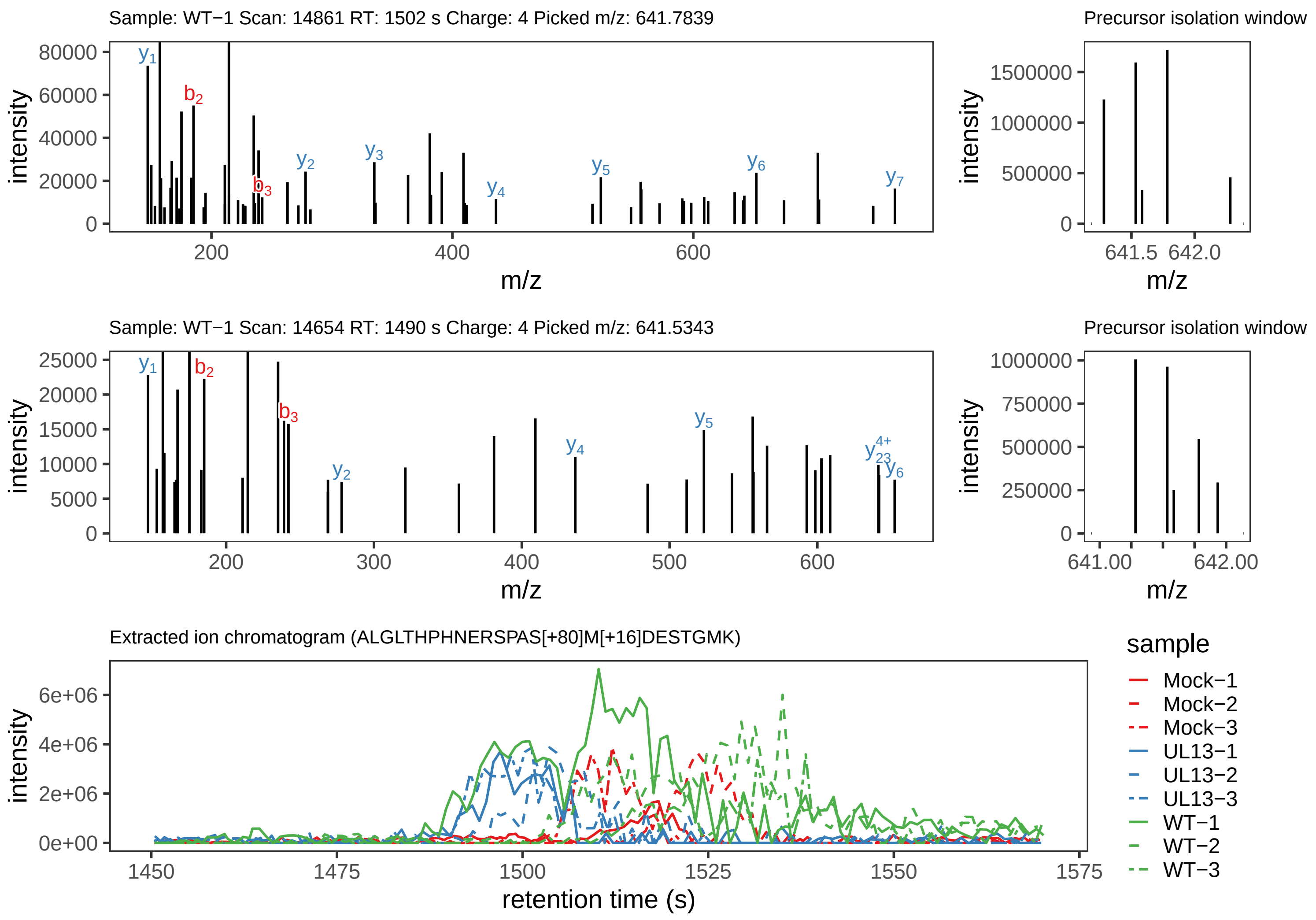

Supplement: Supplemental File 5 [file 737029_file08.zip › phosphopeptide_validation_figs_20240531/ALGLTHPHNERSPASMDESTGMK/ALGLTHPHNERSPAS[+80]M[+16]DESTGMK.phospho.pdf]

#### Peptide ALGLTHPHNERSPASMDES[+80]TGMK

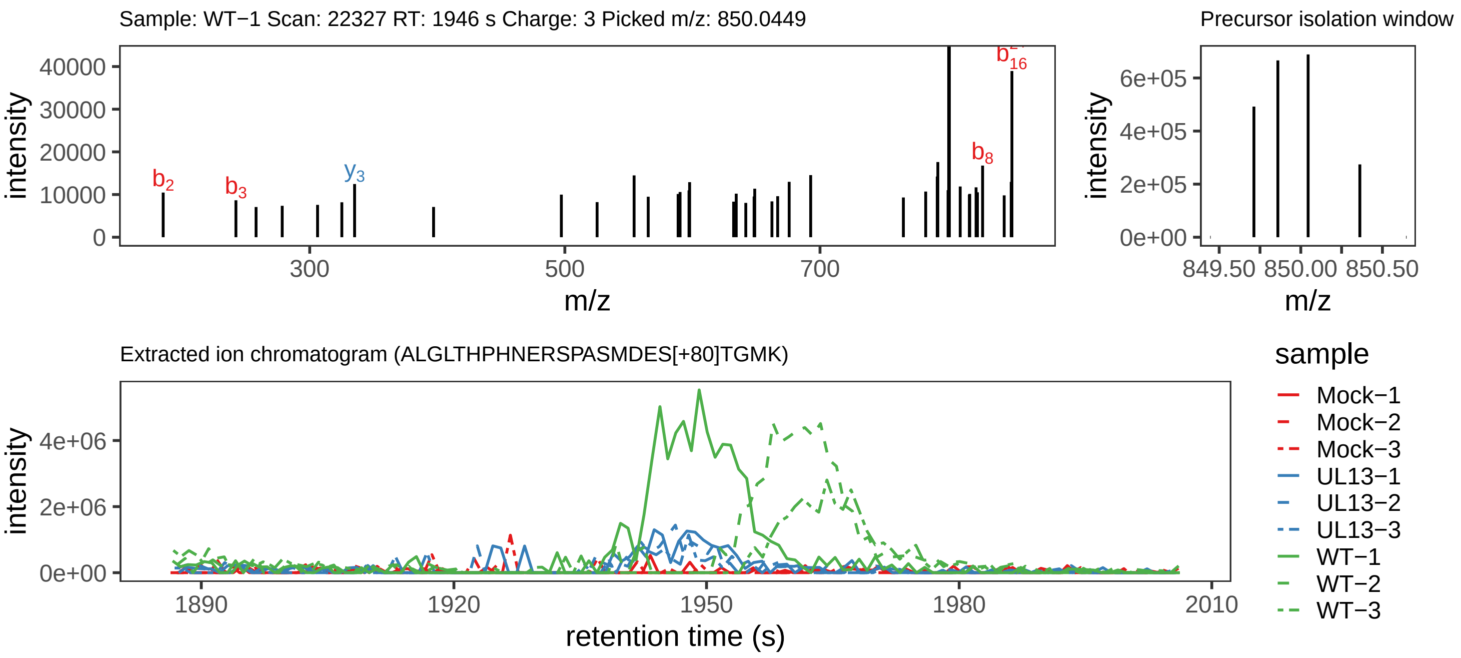

Supplement: Supplemental File 5 [file 737029_file08.zip › phosphopeptide_validation_figs_20240531/ALGLTHPHNERSPASMDESTGMK/ALGLTHPHNERSPASMDES[+80]TGMK.phospho.pdf]

Peptide ALGLTHPHNERS[+80]PASMDESTGM[+16]K

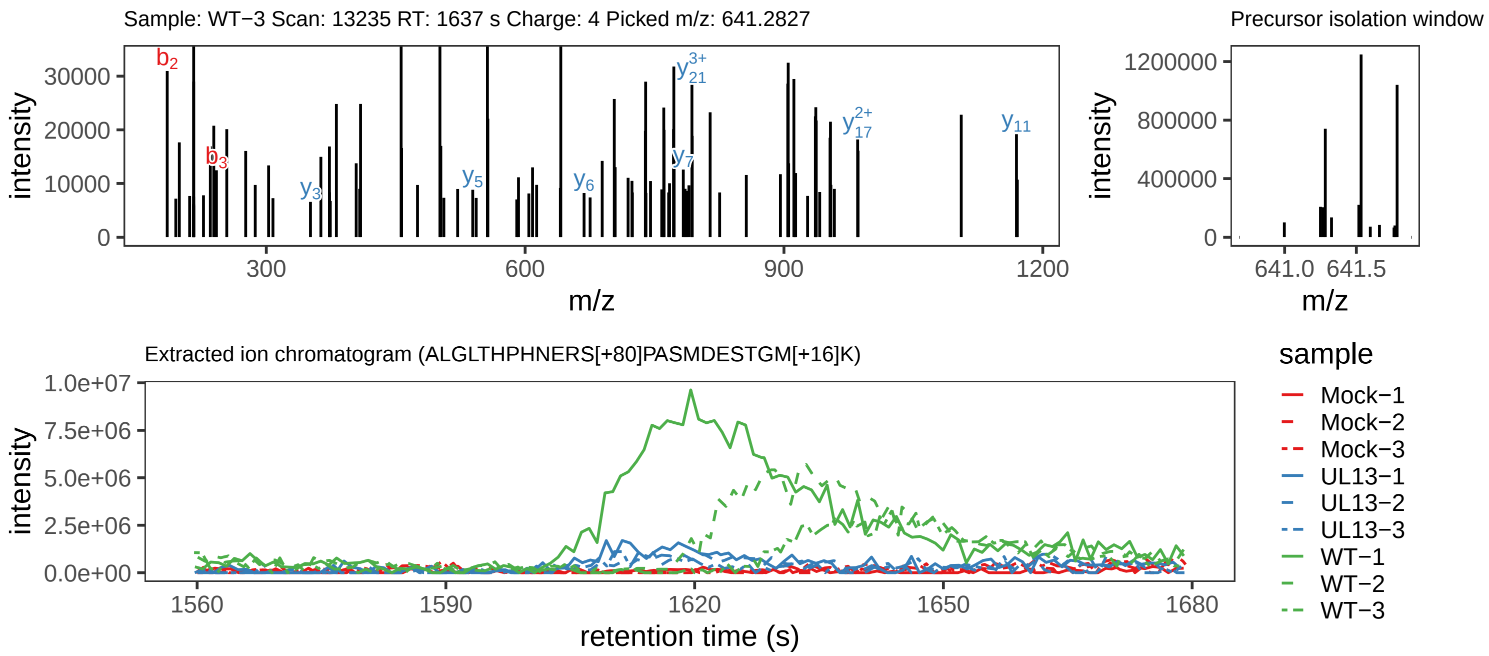

Supplement: Supplemental File 5 [file 737029_file08.zip › phosphopeptide_validation_figs_20240531/ALGLTHPHNERSPASMDESTGMK/ALGLTHPHNERS[+80]PASMDESTGM[+16]K.phospho.pdf]

#### Peptide ALGLTHPHNERS[+80]PASM[+16]DESTGMK

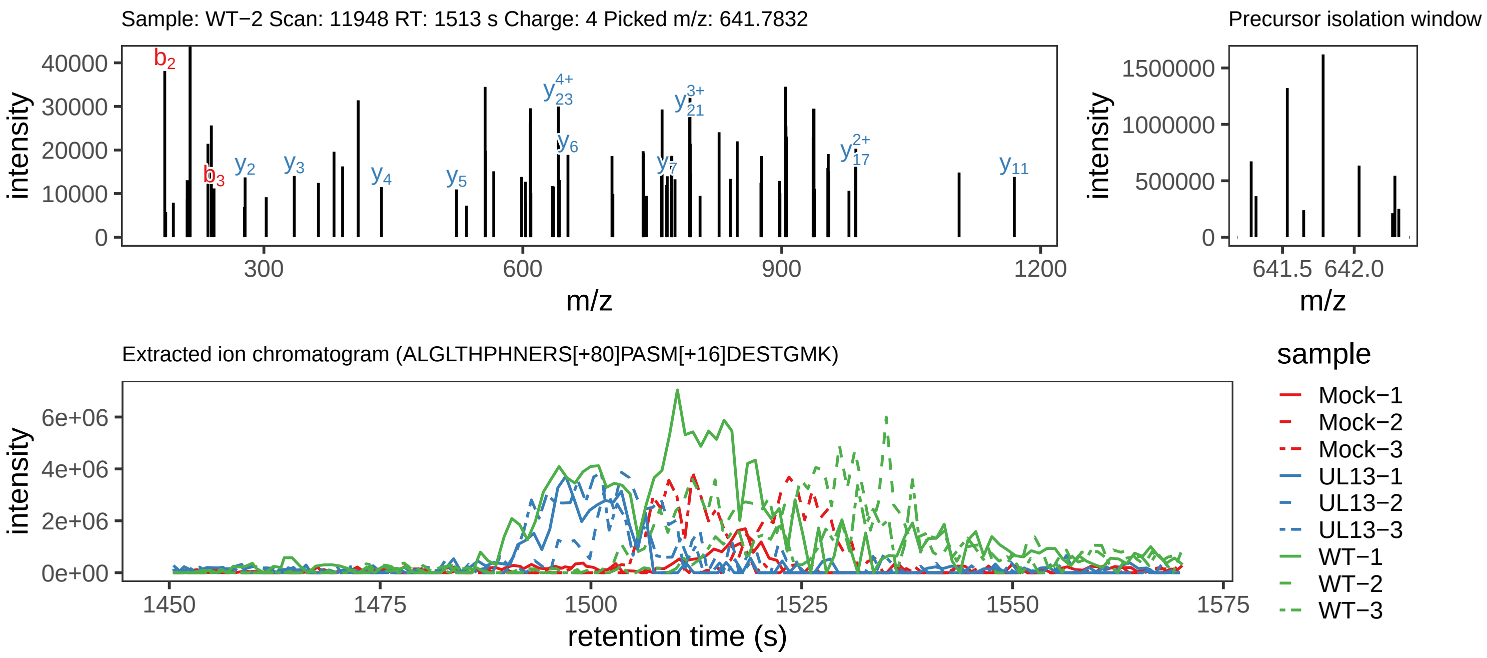

Supplement: Supplemental File 5 [file 737029_file08.zip › phosphopeptide_validation_figs_20240531/ALGLTHPHNERSPASMDESTGMK/ALGLTHPHNERS[+80]PASM[+16]DESTGMK.phospho.pdf]

#### Peptide ALGLTHPHNERS[+80]PASMDESTGMK

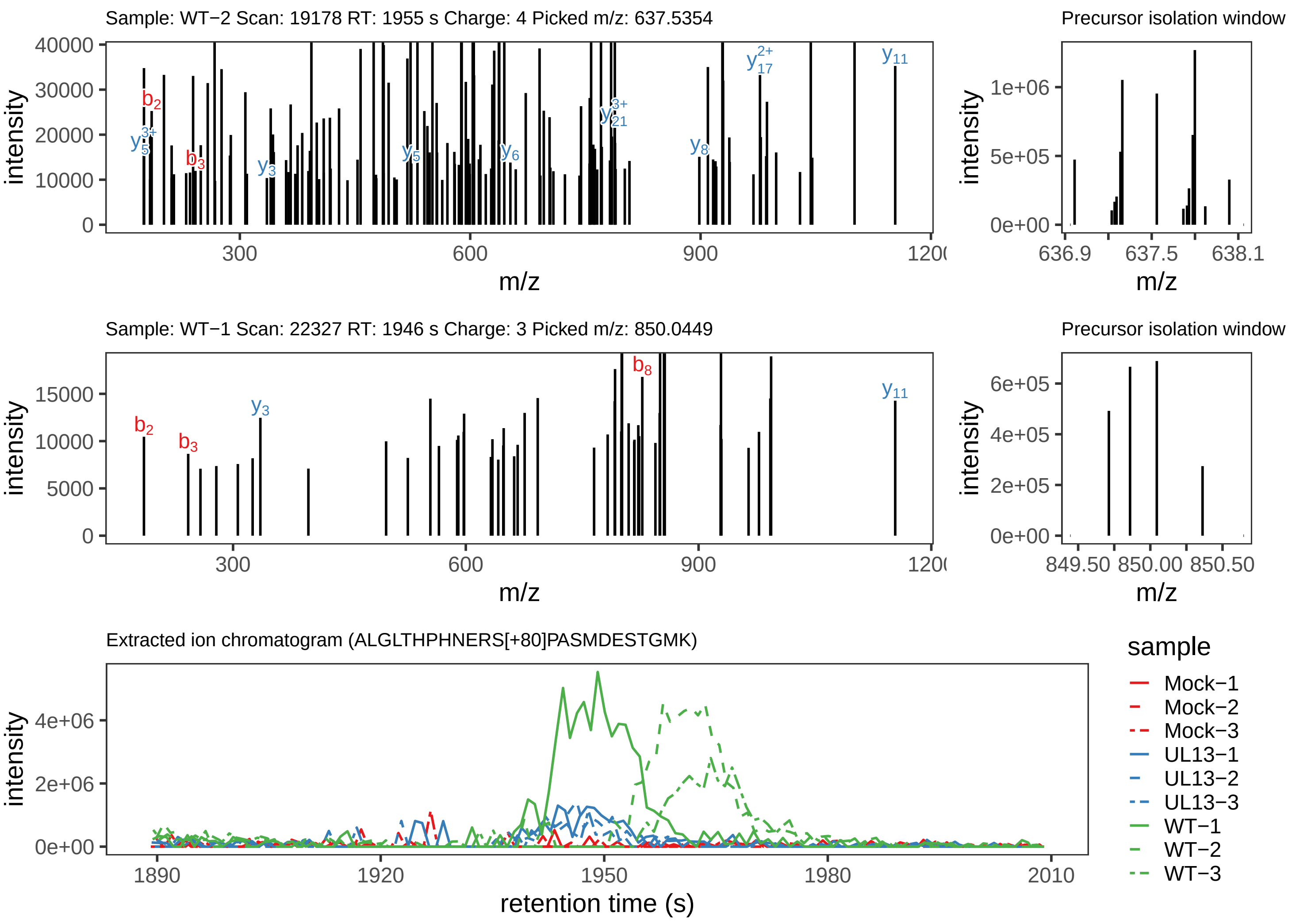

Supplement: Supplemental File 5 [file 737029_file08.zip › phosphopeptide_validation_figs_20240531/ALGLTHPHNERSPASMDESTGMK/ALGLTHPHNERS[+80]PASMDESTGMK.phospho.pdf]

#### Peptide ALGLTHPHN[+1]ERSPAS[+80]M[+16]DESTGMK

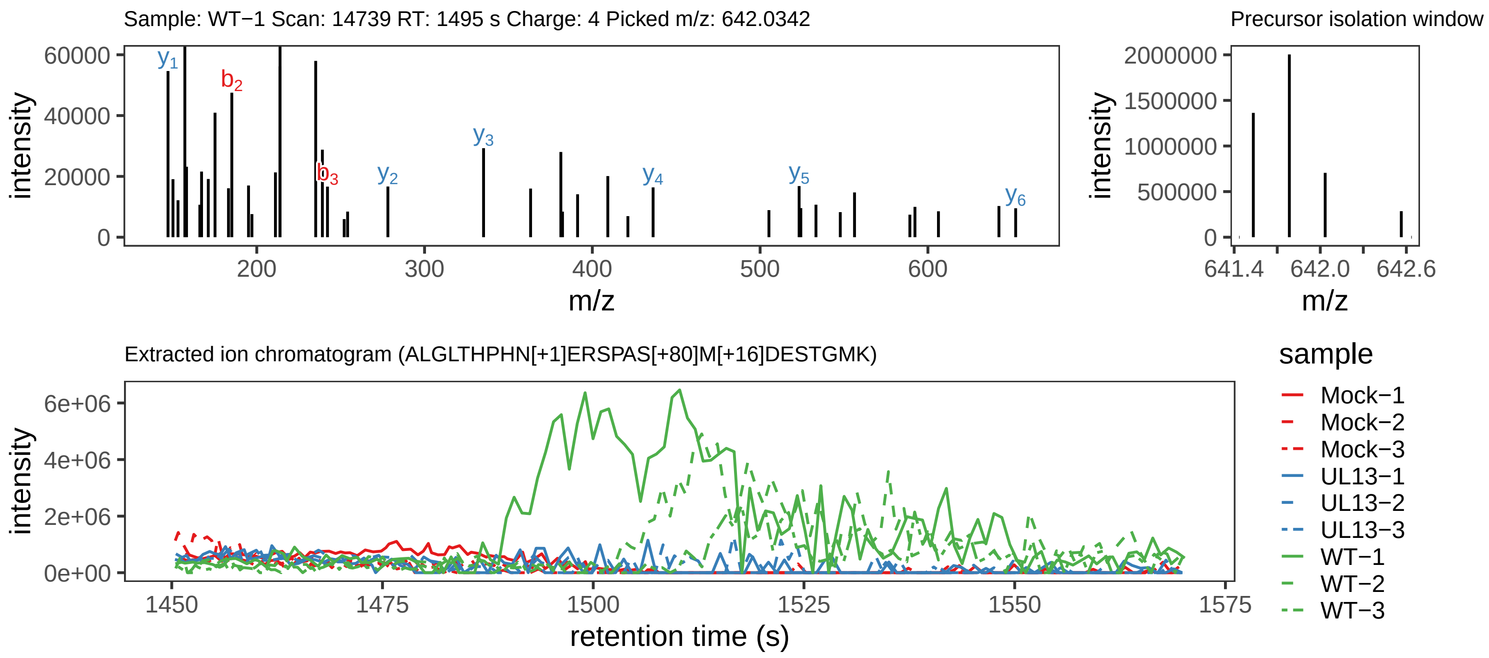

Supplement: Supplemental File 5 [file 737029_file08.zip › phosphopeptide_validation_figs_20240531/ALGLTHPHNERSPASMDESTGMK/ALGLTHPHN[+1]ERSPAS[+80]M[+16]DESTGMK.phospho.pdf]

#### Peptide DM[+16]INAS[+80]LK

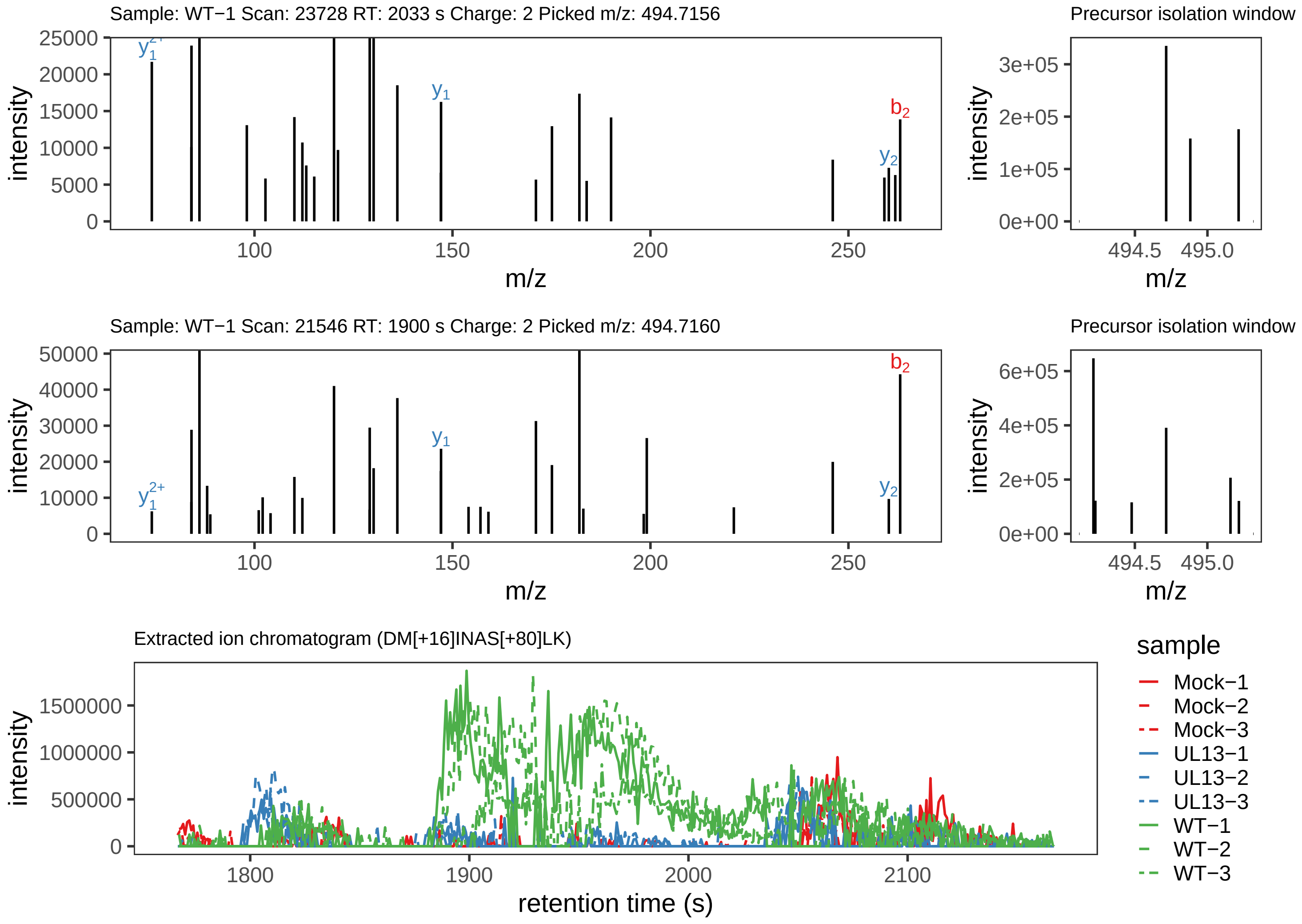

Supplement: Supplemental File 5 [file 737029_file08.zip › phosphopeptide_validation_figs_20240531/DMINASLK/DM[+16]INAS[+80]LK.phospho.pdf]

#### Peptide DMINASLK

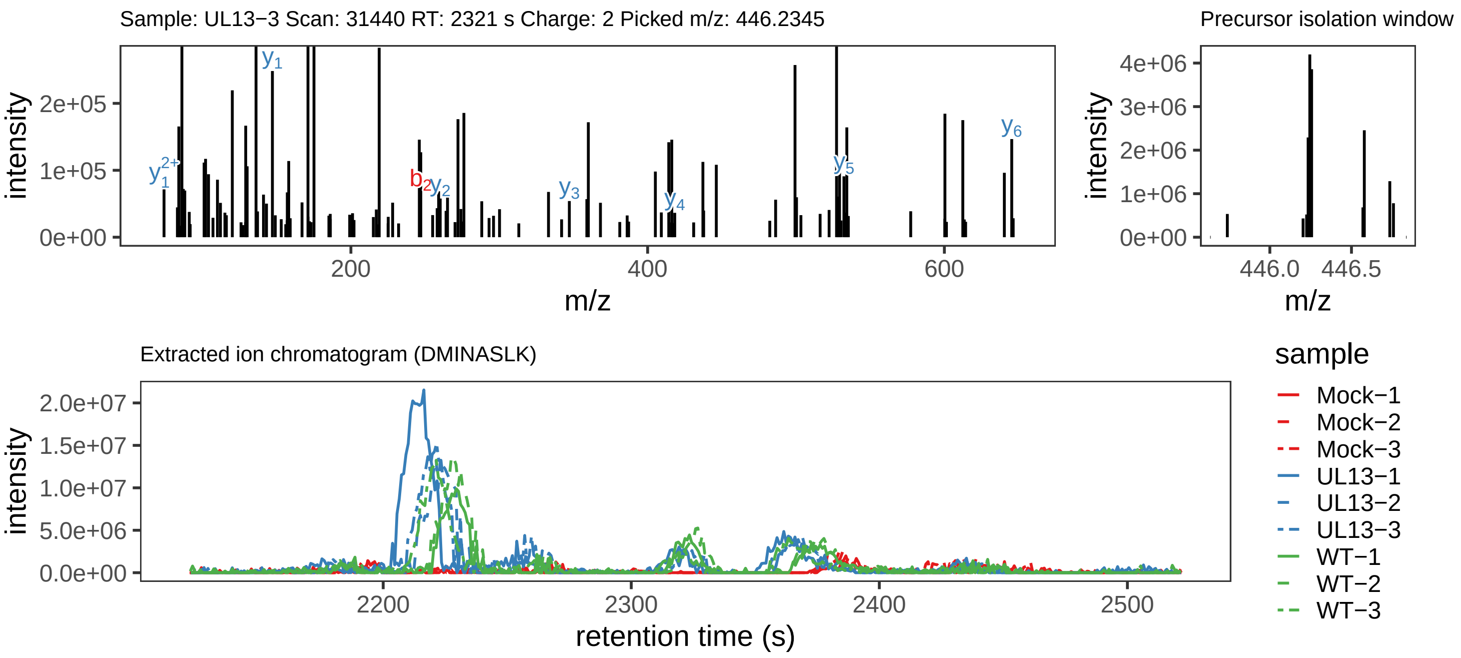

### GDPPTGMASVIYEDEK.global.pdf

#### Peptide ILGSPPNPLKPPEIEPPQMS[+80]STPGR

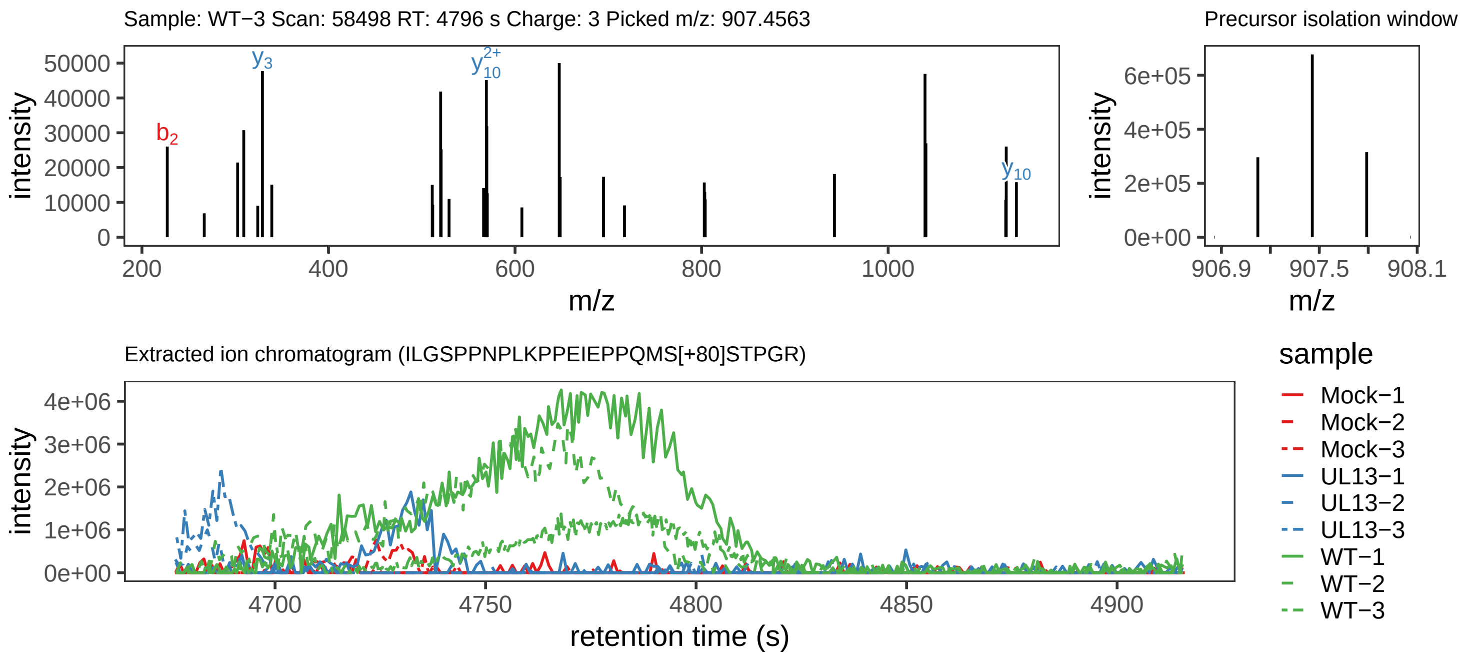

Supplement: Supplemental File 5 [file 737029_file08.zip › phosphopeptide_validation_figs_20240531/ILGSPPNPLKPPEIEPPQMSSTPGR/ILGSPPNPLKPPEIEPPQMS[+80]STPGR.phospho.pdf]

#### Peptide ILGSPPN[+1]PLKPPEIEPPQM[+16]SST[+80]PGR

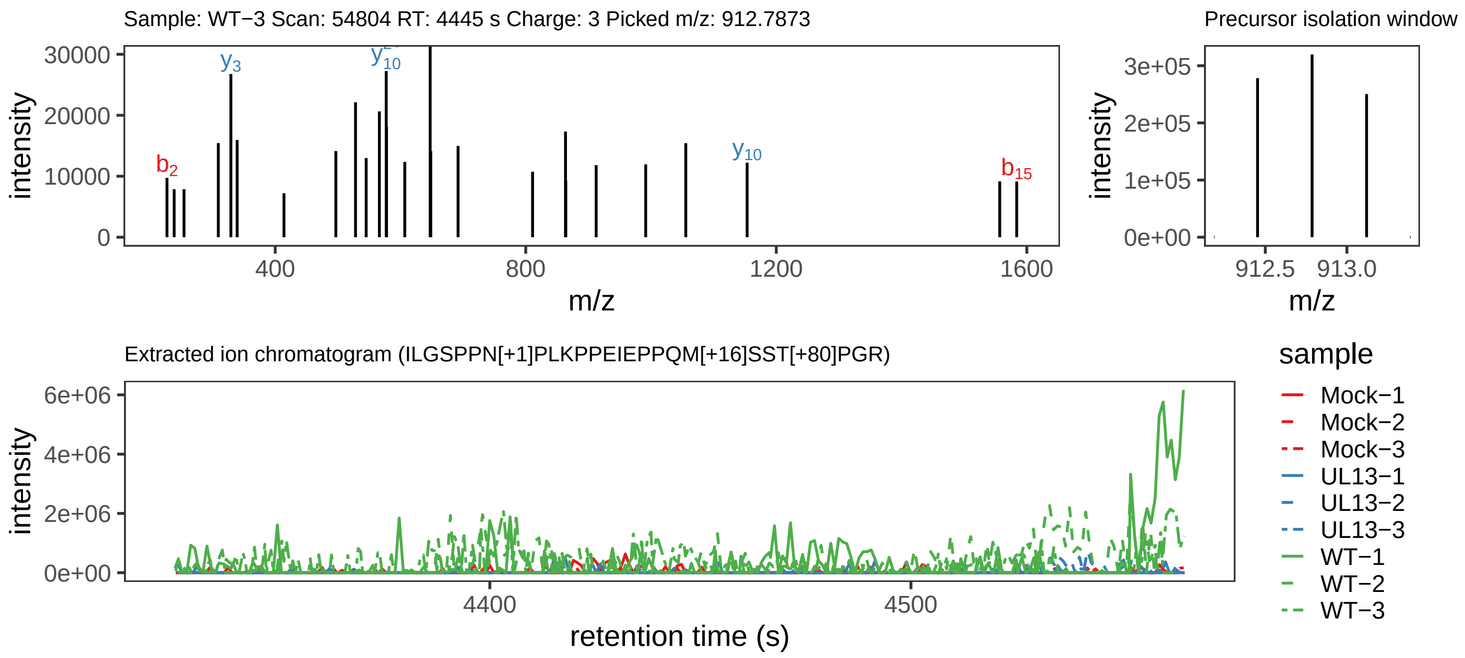

Supplement: Supplemental File 5 [file 737029_file08.zip › phosphopeptide_validation_figs_20240531/ILGSPPNPLKPPEIEPPQMSSTPGR/ILGSPPN[+1]PLKPPEIEPPQM[+16]SST[+80]PGR.phospho.pdf]

#### Peptide ILGSPPNPLKPPEIEPPQM[+16]SS[+80]TPGR

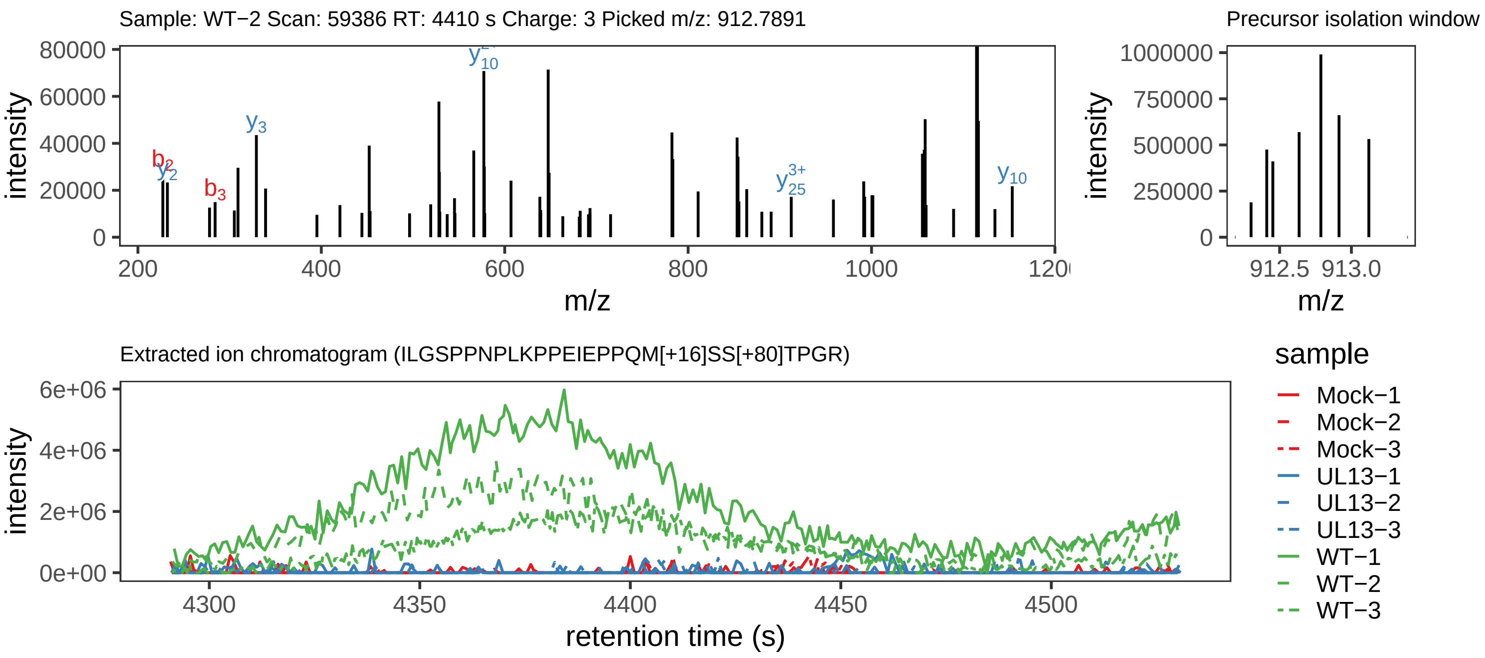

Supplement: Supplemental File 5 [file 737029_file08.zip › phosphopeptide_validation_figs_20240531/ILGSPPNPLKPPEIEPPQMSSTPGR/ILGSPPNPLKPPEIEPPQM[+16]SS[+80]TPGR.phospho.pdf]

#### Peptide ILGS[+80]PPNPLKPPEIEPPQ[+1]M[+16]SSTPGR

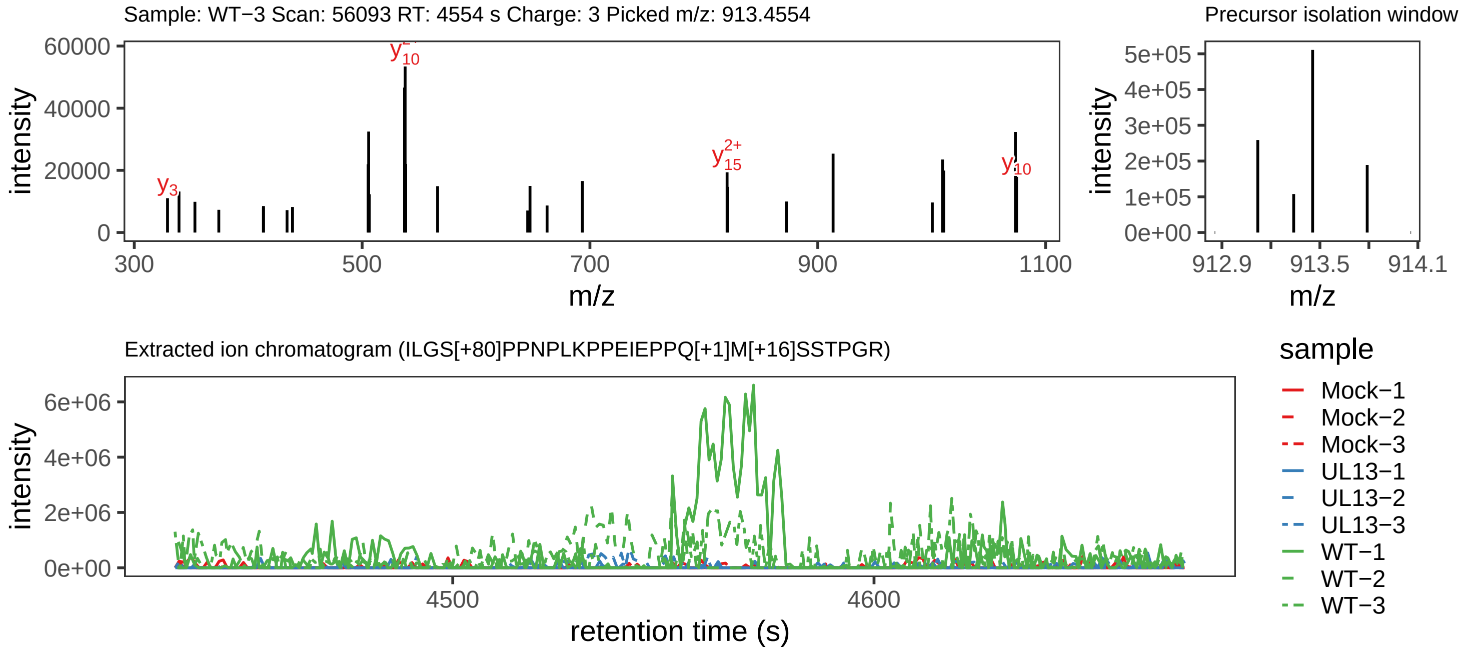

Supplement: Supplemental File 5 [file 737029_file08.zip › phosphopeptide_validation_figs_20240531/ILGSPPNPLKPPEIEPPQMSSTPGR/ILGS[+80]PPNPLKPPEIEPPQ[+1]M[+16]SSTPGR.phospho.pdf]

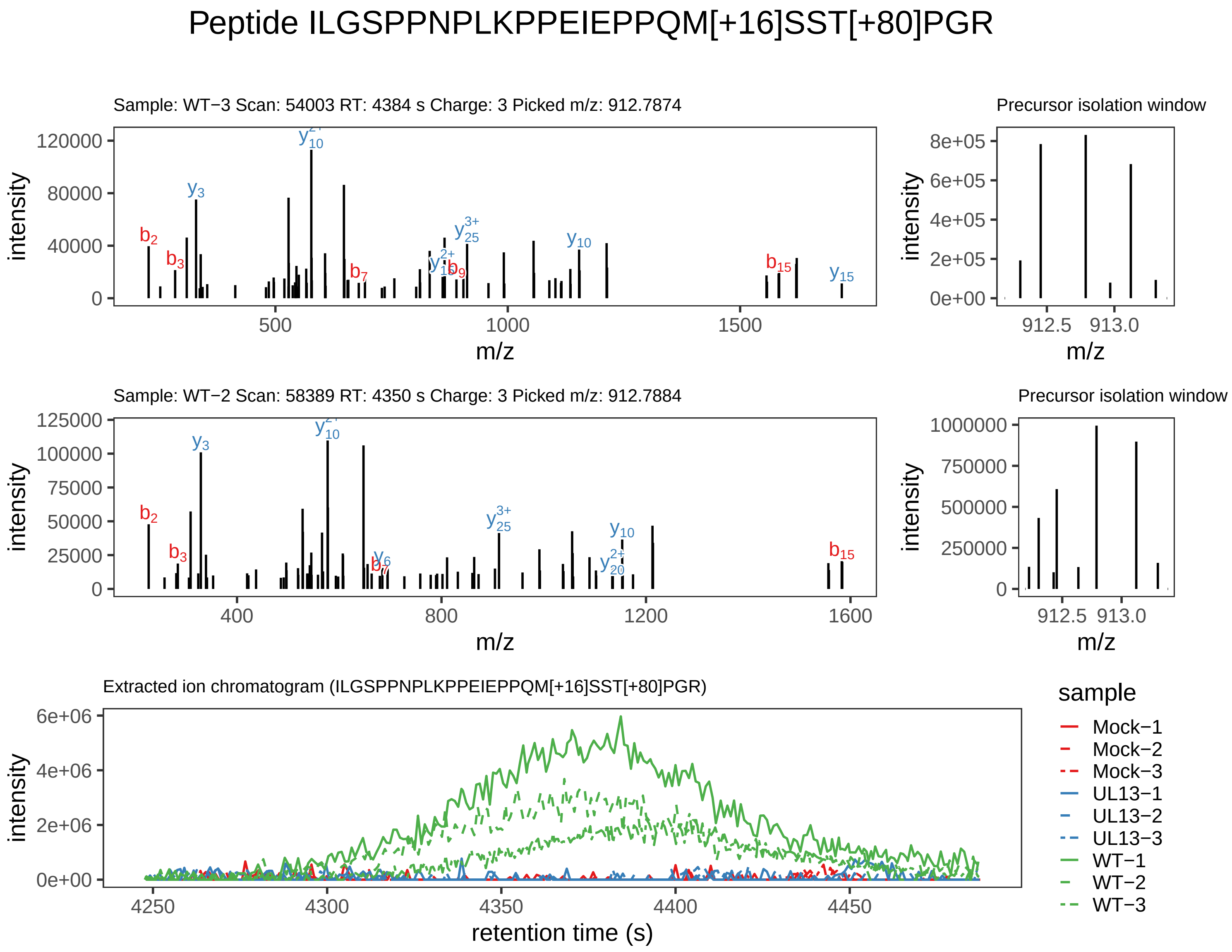

Supplement: Supplemental File 5 [file 737029_file08.zip › phosphopeptide_validation_figs_20240531/ILGSPPNPLKPPEIEPPQMSSTPGR/ILGSPPNPLKPPEIEPPQM[+16]SST[+80]PGR.phospho.pdf]

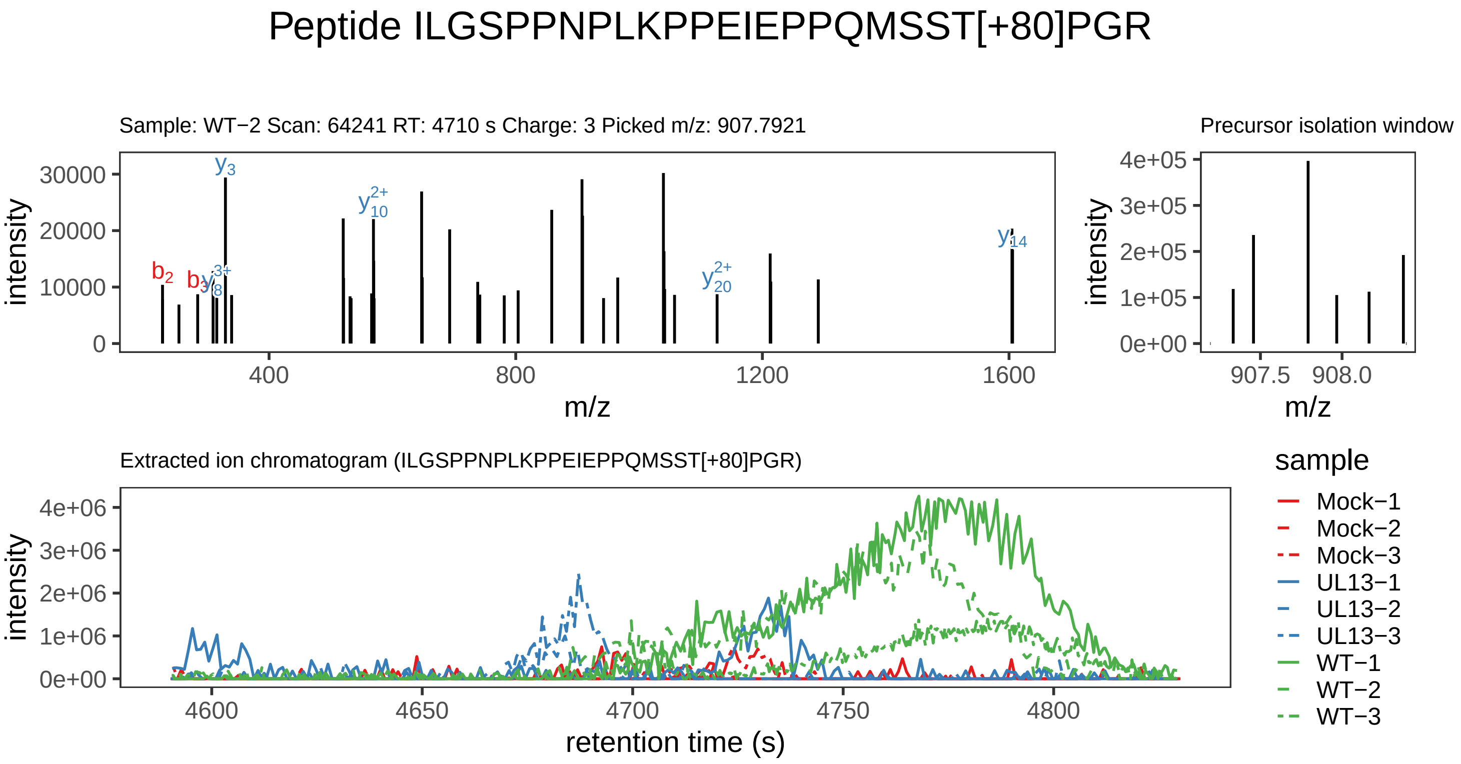

Supplement: Supplemental File 5 [file 737029_file08.zip › phosphopeptide_validation_figs_20240531/ILGSPPNPLKPPEIEPPQMSSTPGR/ILGSPPNPLKPPEIEPPQMSST[+80]PGR.phospho.pdf]

#### Peptide [+42]AM[+16]WS[+80]LR

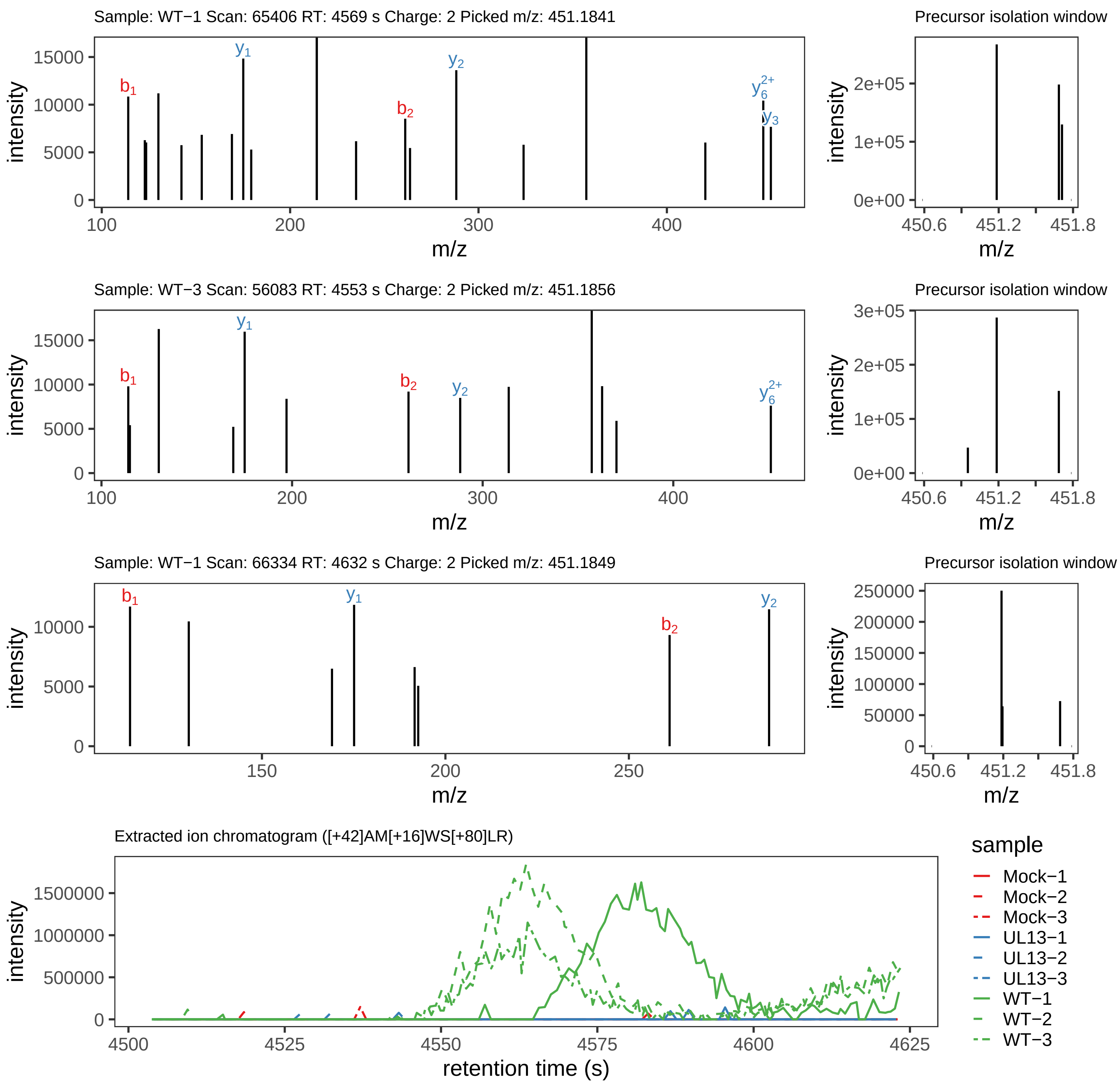

Supplement: Supplemental File 5 [file 737029_file08.zip › phosphopeptide_validation_figs_20240531/AMWSLR/[+42]AM[+16]WS[+80]LR.phospho.pdf]

#### Peptide A[+42]M[+16]WS[+80]LR

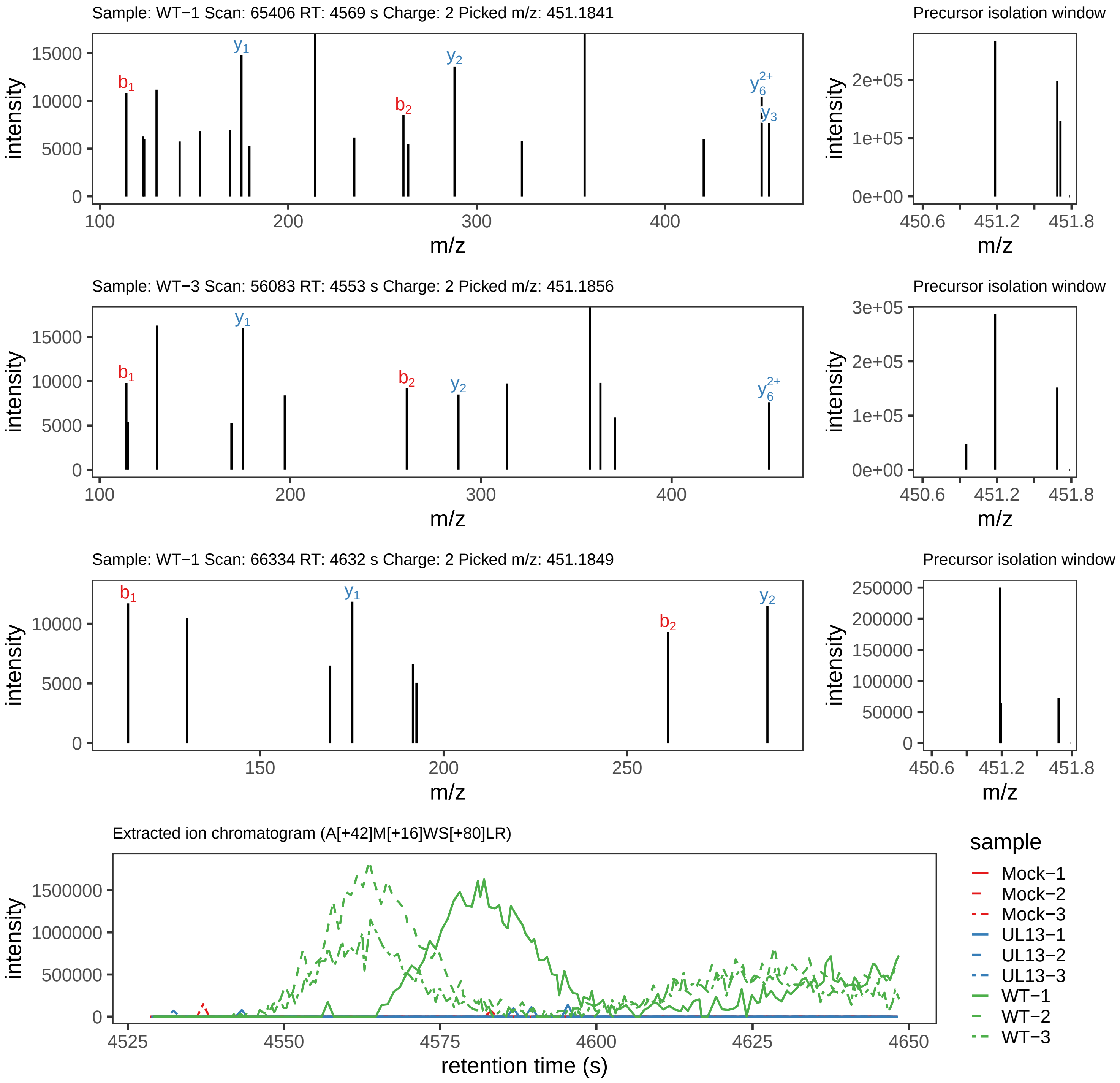

Supplement: Supplemental File 5 [file 737029_file08.zip › phosphopeptide_validation_figs_20240531/AMWSLR/A[+42]M[+16]WS[+80]LR.phospho.pdf]

#### Peptide [+42]AM[+16]WSLR

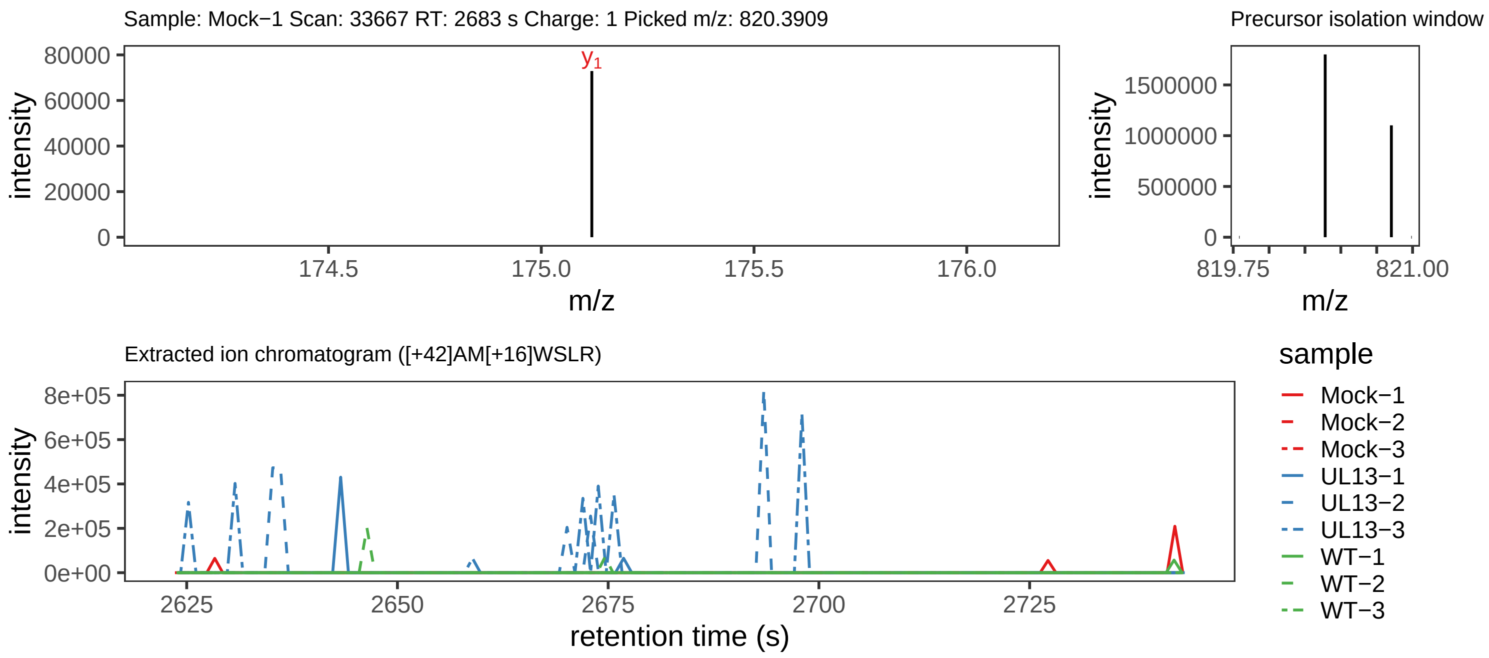

Supplement: Supplemental File 5 [file 737029_file08.zip › phosphopeptide_validation_figs_20240531/AMWSLR/[+42]AM[+16]WSLR.phospho.pdf]

#### Peptide AMWS[+80]LR

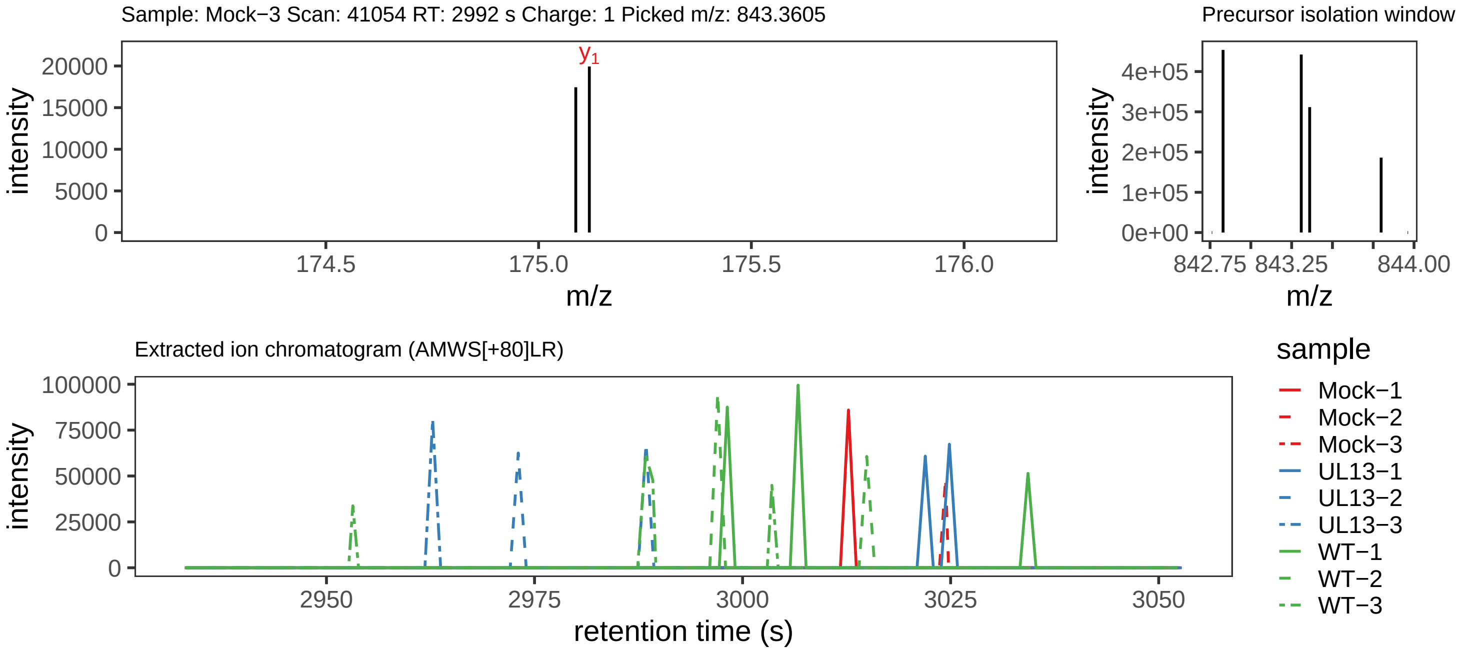

Supplement: Supplemental File 5 [file 737029_file08.zip › phosphopeptide_validation_figs_20240531/AMWSLR/AMWS[+80]LR.phospho.pdf]

#### Peptide QSELISSIRRDPQGT[+80]FWTSPSPHGNK

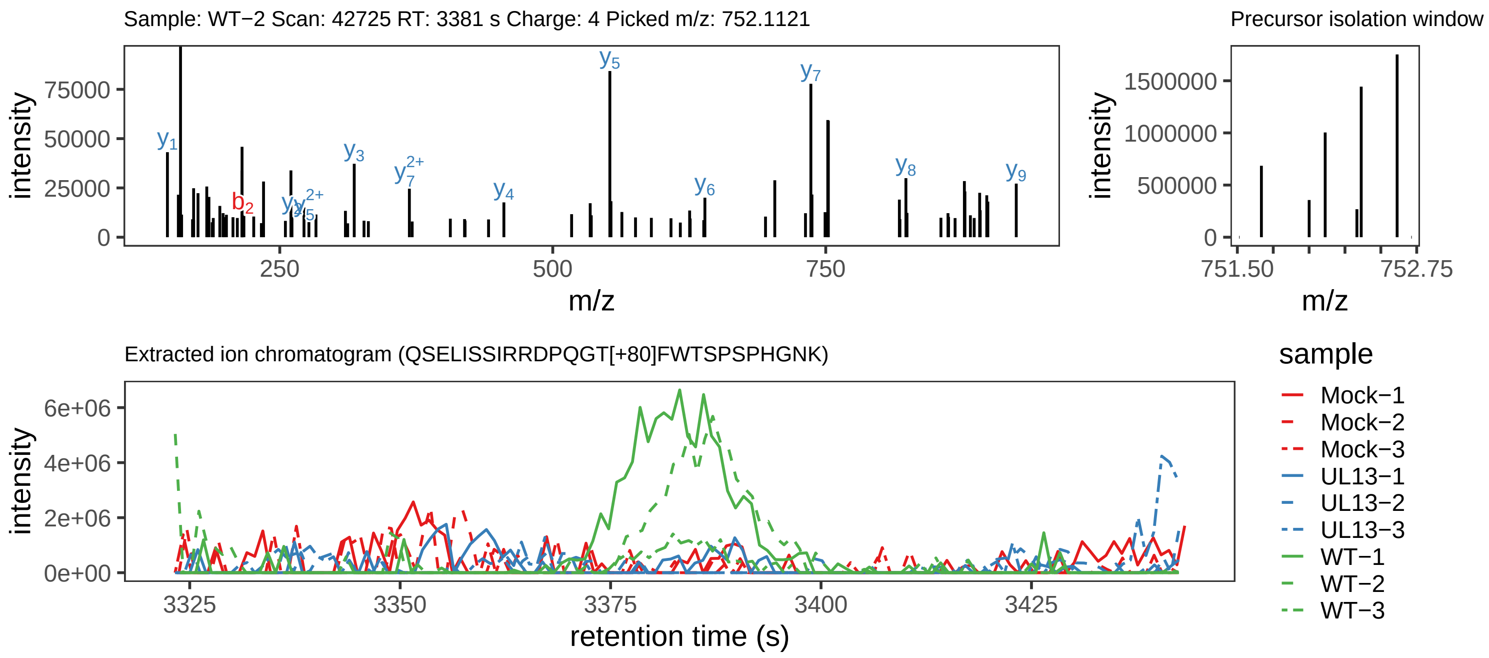

Supplement: Supplemental File 5 [file 737029_file08.zip › phosphopeptide_validation_figs_20240531/QSELISSIRRDPQGTFWTSPSPHGNK/QSELISSIRRDPQGT[+80]FWTSPSPHGNK.phospho.pdf]

#### Peptide QSELIS[+80]SIRRDPQGTFWTSPSPHGNK

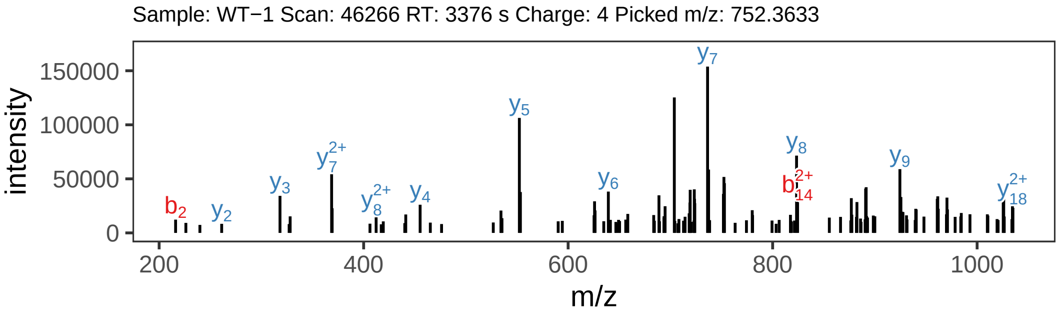


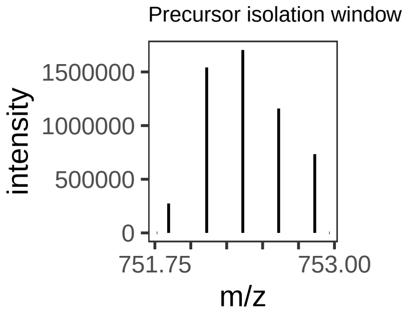


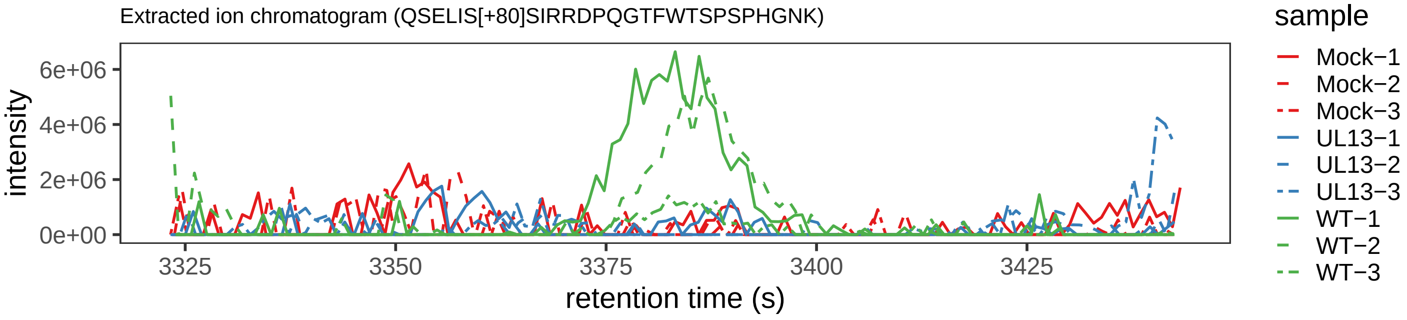

Supplement: Supplemental File 5 [file 737029_file08.zip › phosphopeptide_validation_figs_20240531/QSELISSIRRDPQGTFWTSPSPHGNK/QSELIS[+80]SIRRDPQGTFWTSPSPHGNK.phospho.pdf]

#### Peptide QS[+80]ELISSIRRDPQ[+1]GTFWTSPSPHGNK

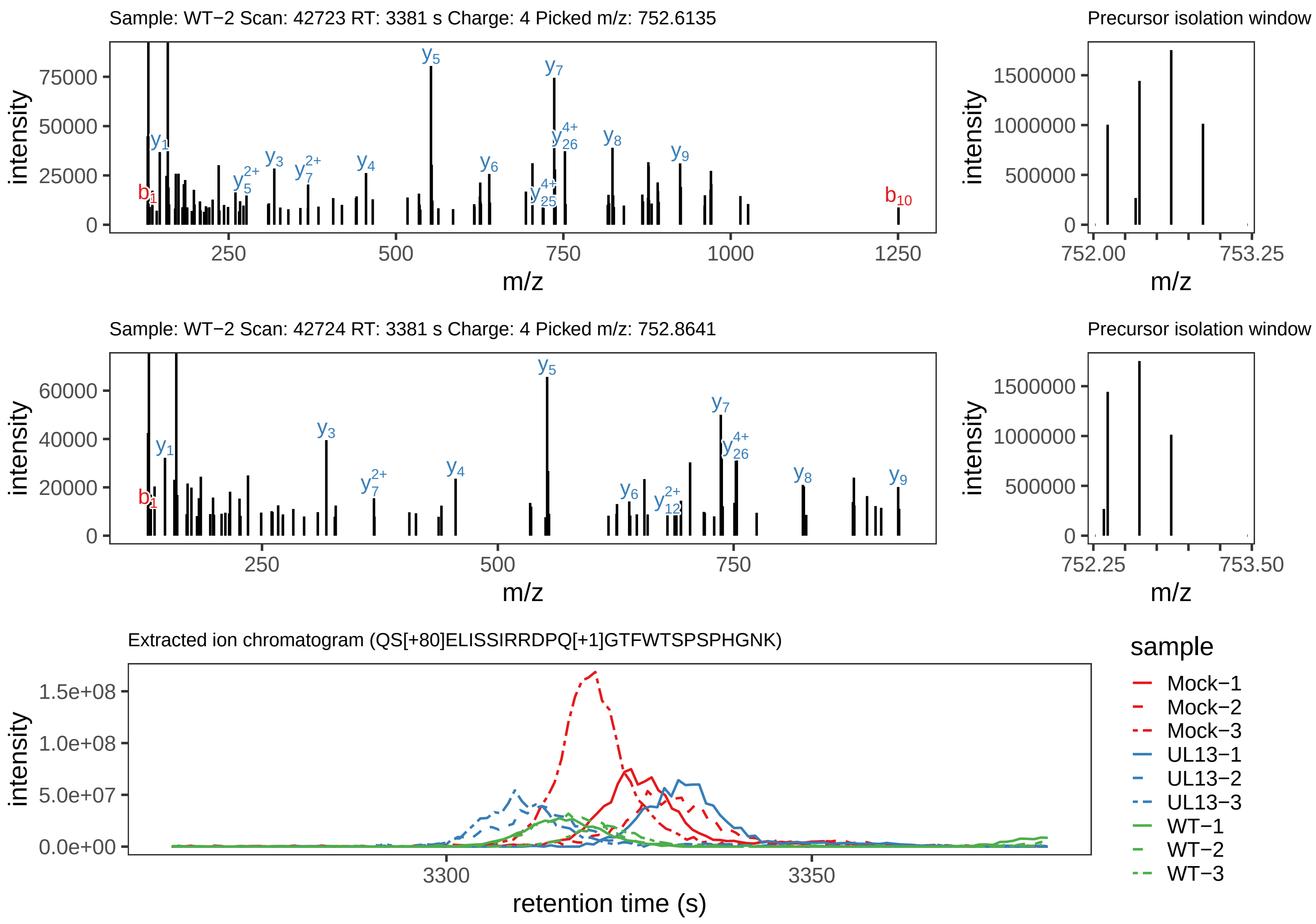

Supplement: Supplemental File 5 [file 737029_file08.zip › phosphopeptide_validation_figs_20240531/QSELISSIRRDPQGTFWTSPSPHGNK/QS[+80]ELISSIRRDPQ[+1]GTFWTSPSPHGNK.phospho.pdf]

#### Peptide LM[+16]DIVGSATGC[+57]GIS[+80]PLPEIESYWK

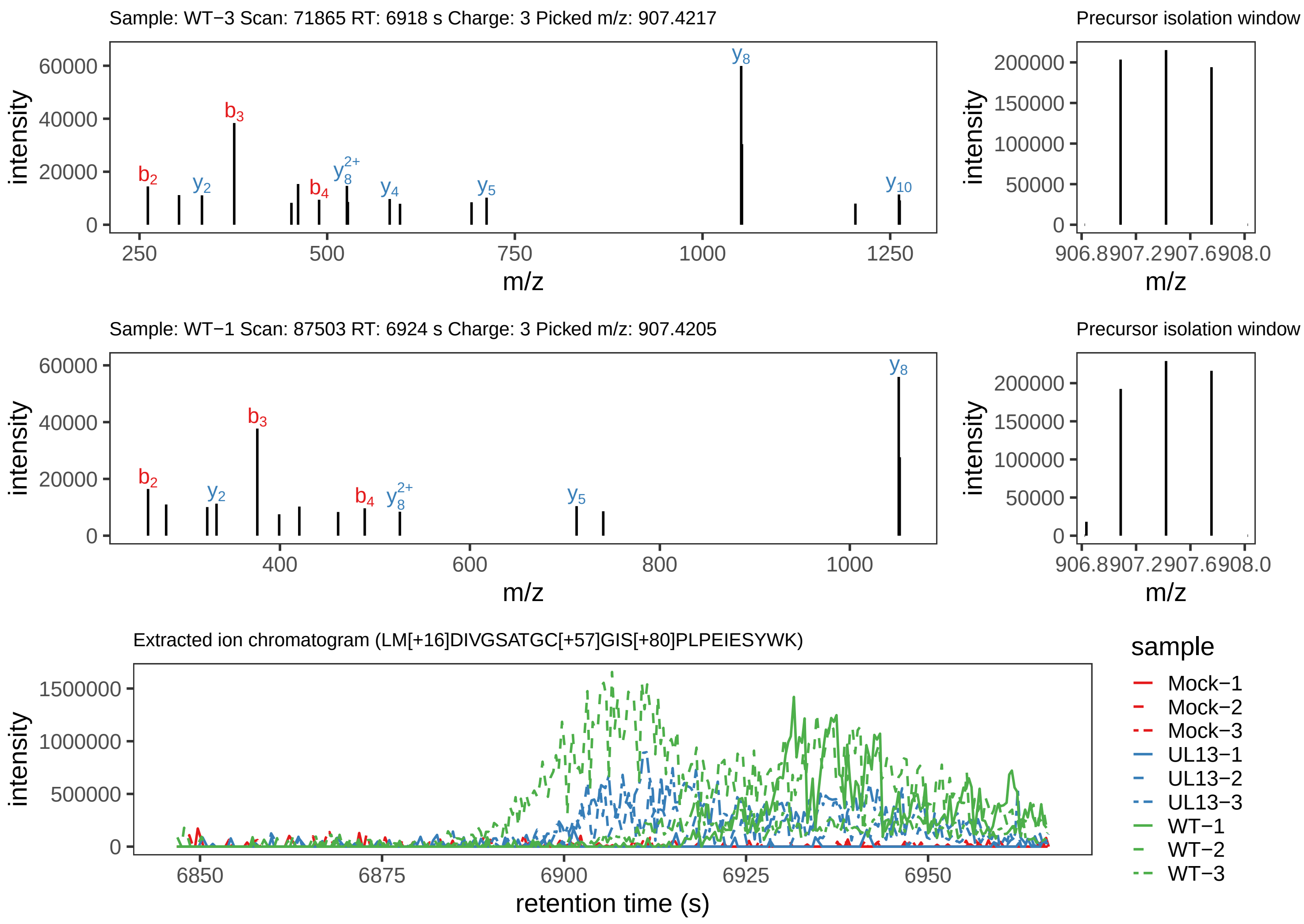

Supplement: Supplemental File 5 [file 737029_file08.zip › phosphopeptide_validation_figs_20240531/LMDIVGSATGCGISPLPEIESYWK/LM[+16]DIVGSATGC[+57]GIS[+80]PLPEIESYWK.phospho.pdf]

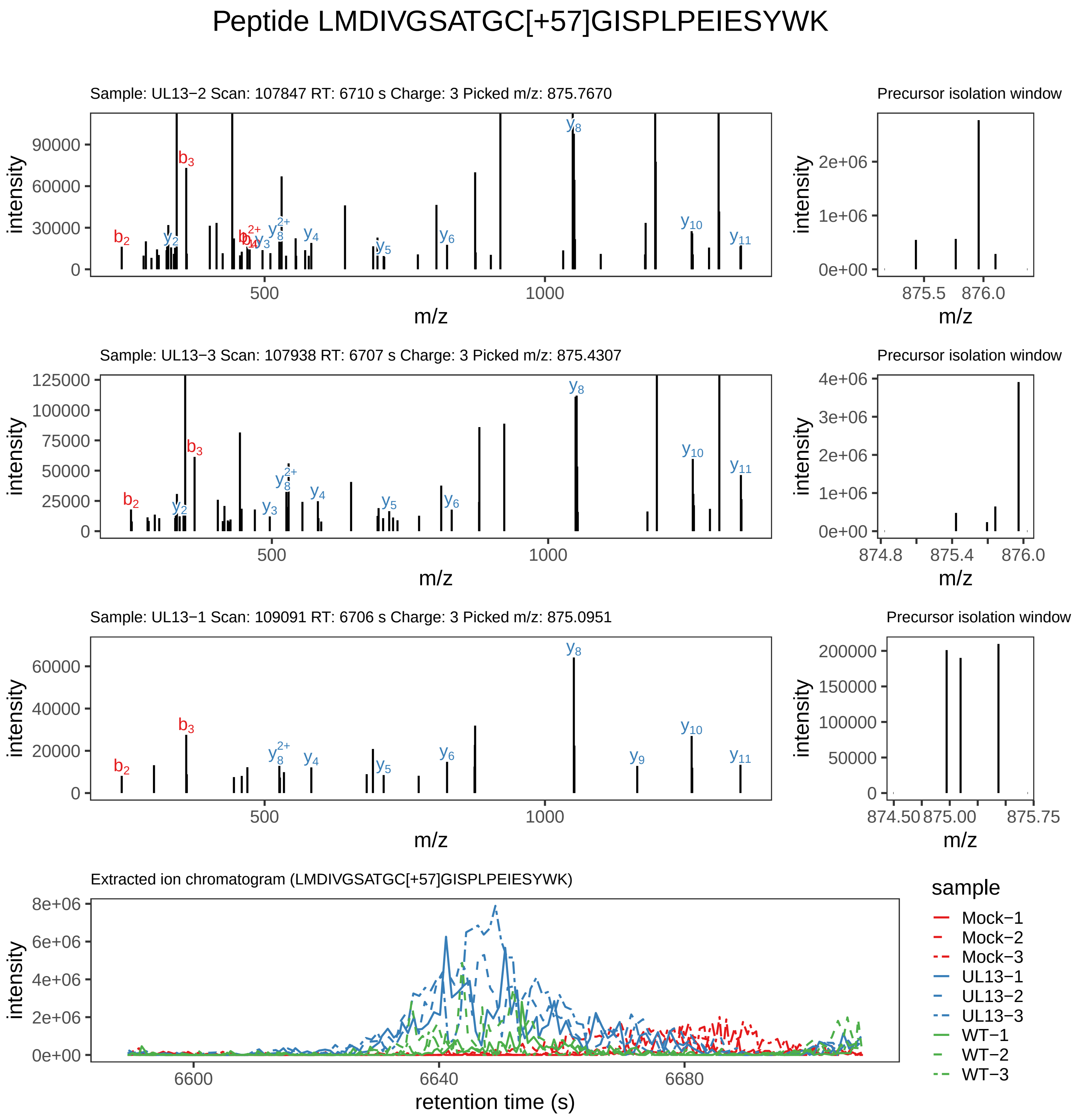

Supplement: Supplemental File 5 [file 737029_file08.zip › phosphopeptide_validation_figs_20240531/LMDIVGSATGCGISPLPEIESYWK/LMDIVGSATGC[+57]GISPLPEIESYWK.global.pdf]

#### Peptide LMDIVGSATGC[+57]GIS[+80]PLPEIESYWK

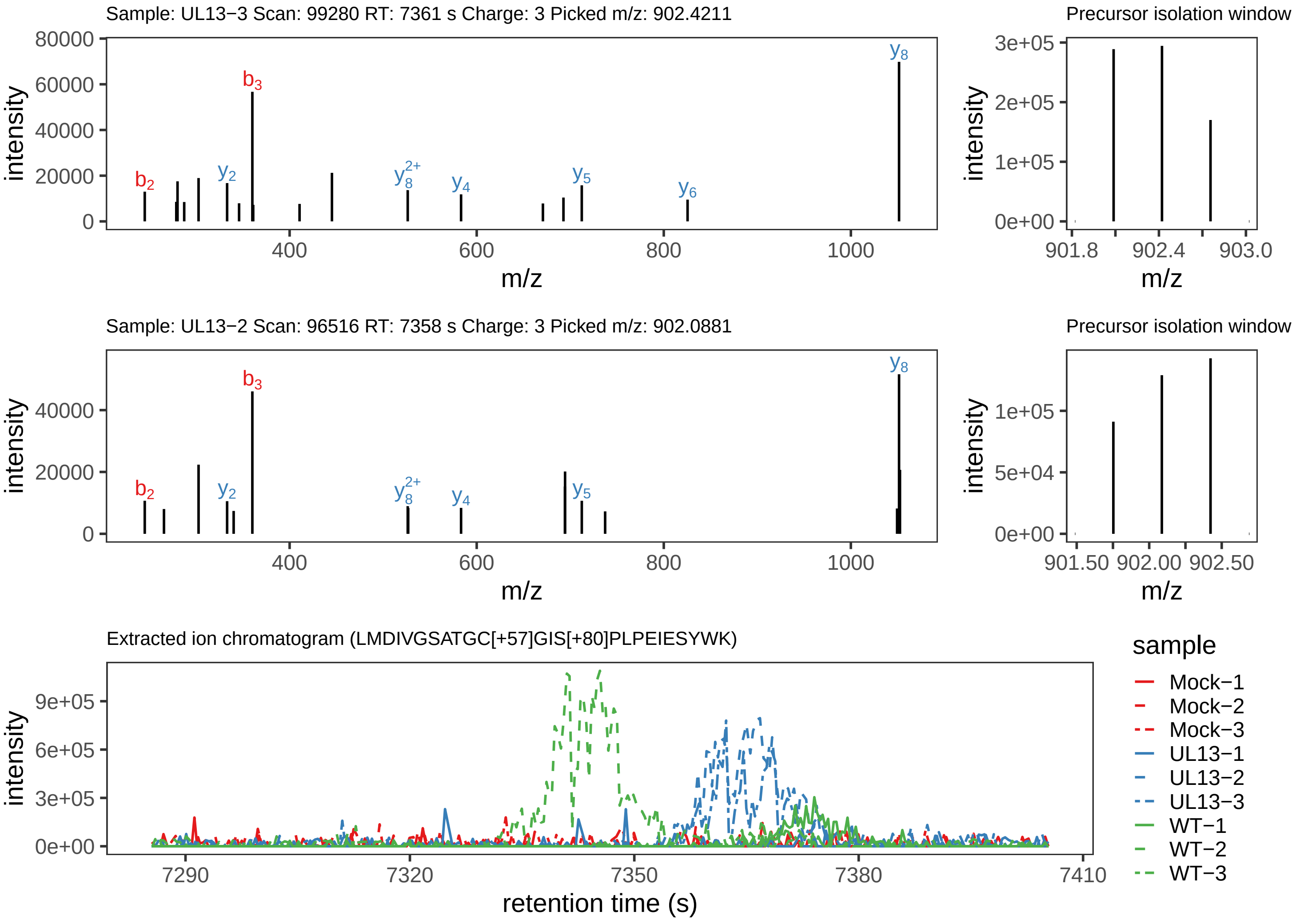

Supplement: Supplemental File 5 [file 737029_file08.zip › phosphopeptide_validation_figs_20240531/LMDIVGSATGCGISPLPEIESYWK/LMDIVGSATGC[+57]GIS[+80]PLPEIESYWK.phospho.pdf]

Peptide LM[+16]DIVGSATGC[+57]GISPLPEIESYWK

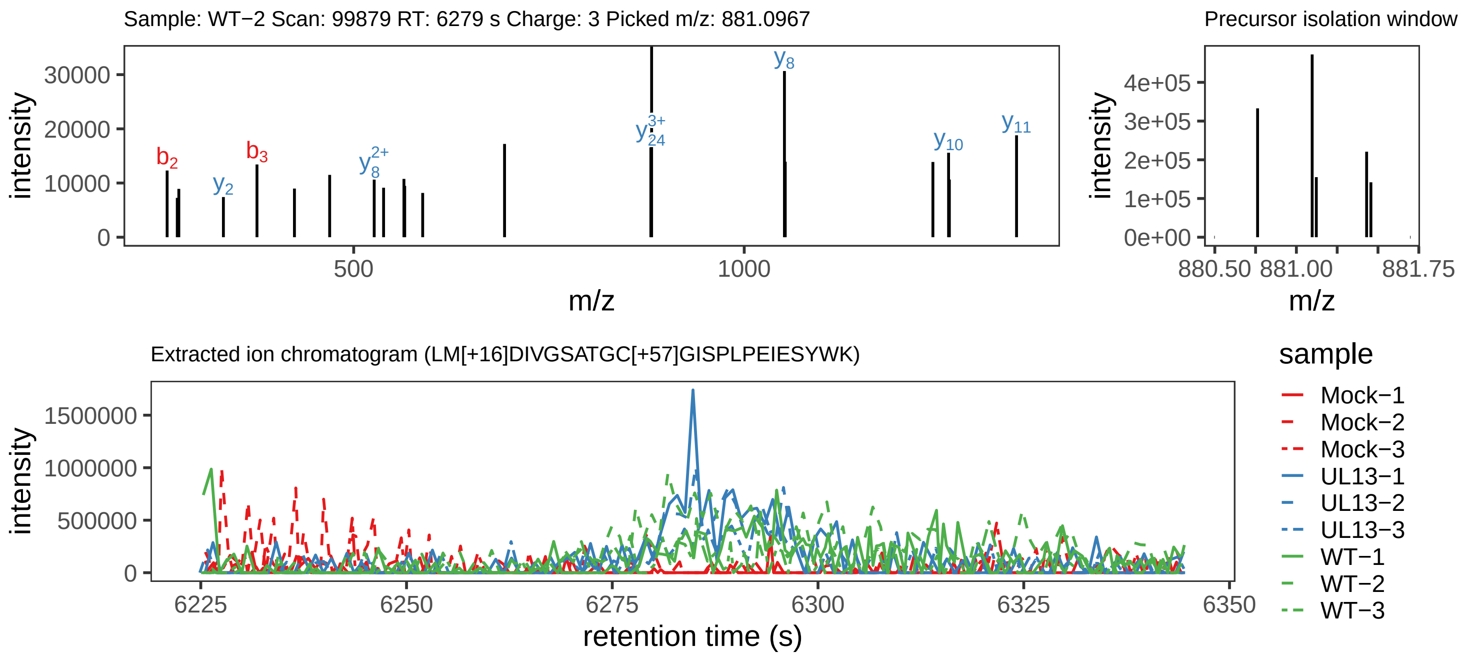

Supplement: Supplemental File 5 [file 737029_file08.zip › phosphopeptide_validation_figs_20240531/LMDIVGSATGCGISPLPEIESYWK/LM[+16]DIVGSATGC[+57]GISPLPEIESYWK.global.pdf]

#### Peptide GDPPTGM[+16]AS[+80]VIYEDEK

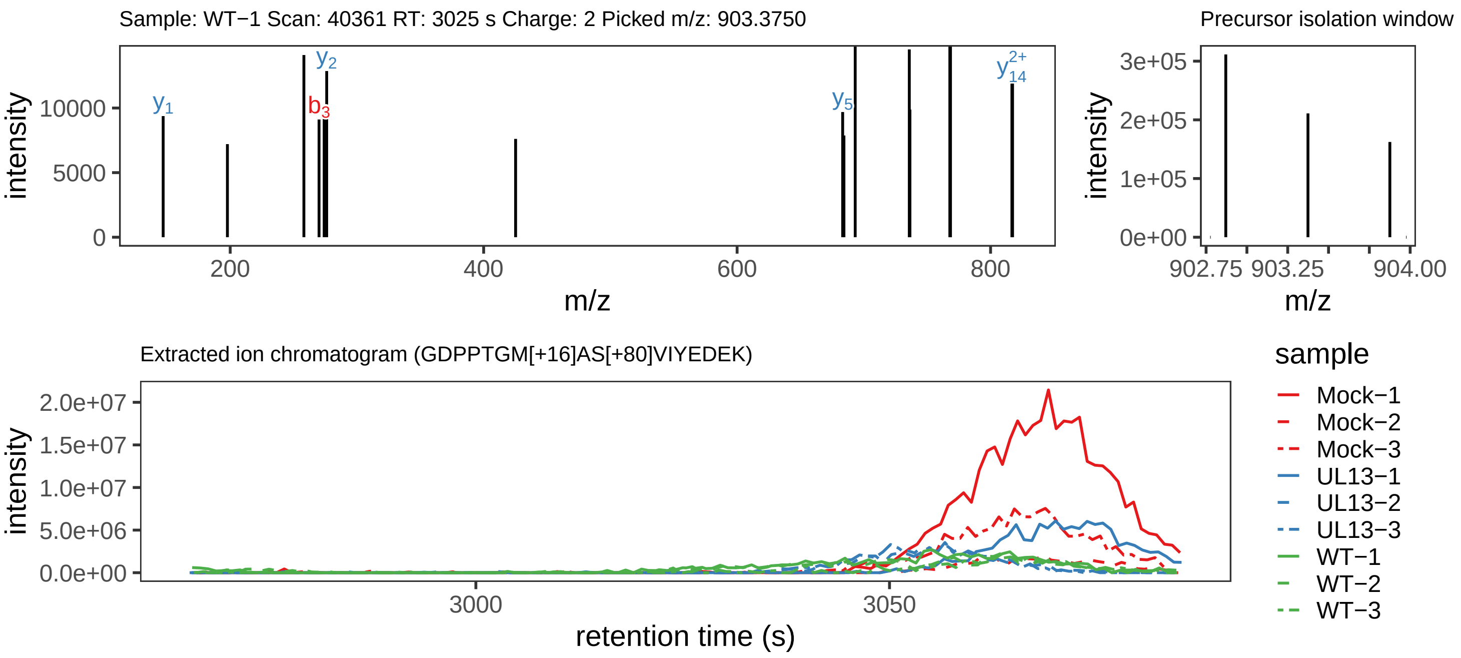

Supplement: Supplemental File 5 [file 737029_file08.zip › phosphopeptide_validation_figs_20240531/GDPPTGMASVIYEDEK/GDPPTGM[+16]AS[+80]VIYEDEK.phospho.pdf]

#### Peptide GDPPTGM[+16]ASVIYEDEK

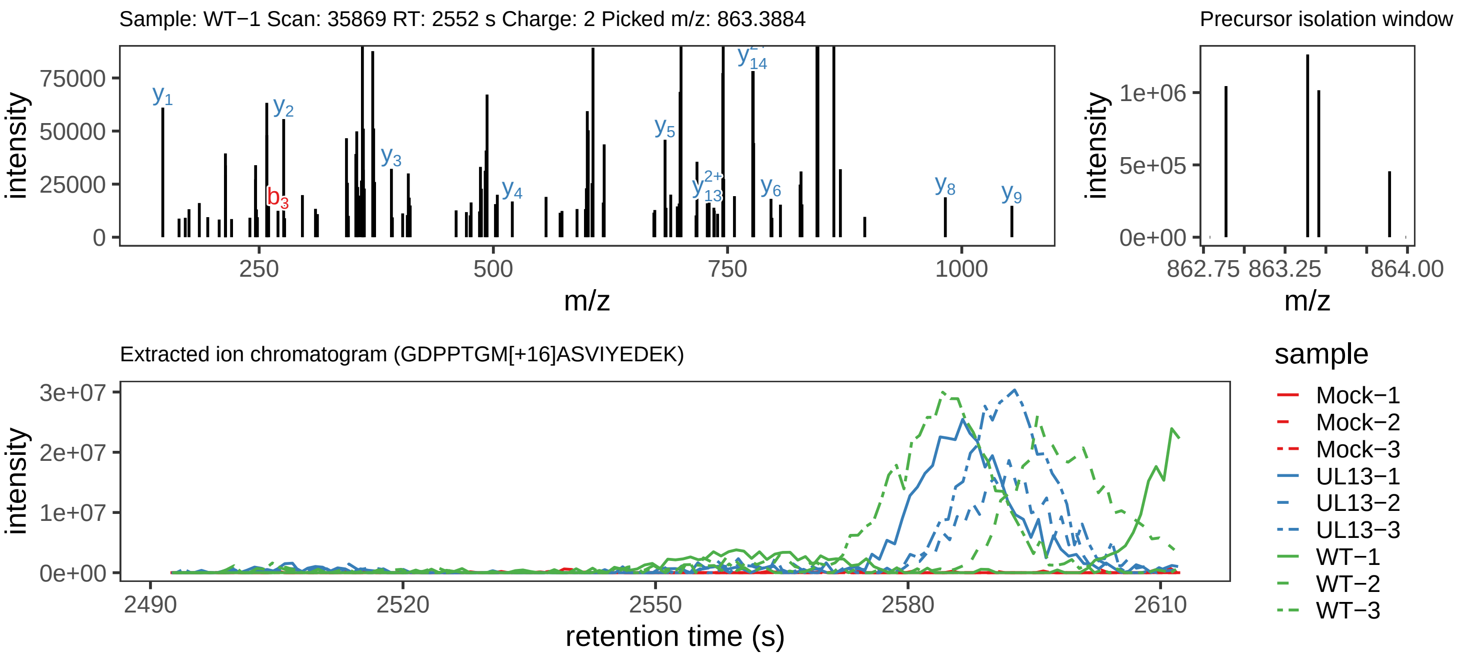

Supplement: Supplemental File 5 [file 737029_file08.zip › phosphopeptide_validation_figs_20240531/GDPPTGMASVIYEDEK/GDPPTGM[+16]ASVIYEDEK.global.pdf]

#### Peptide GDPPTGMASVIYEDEK

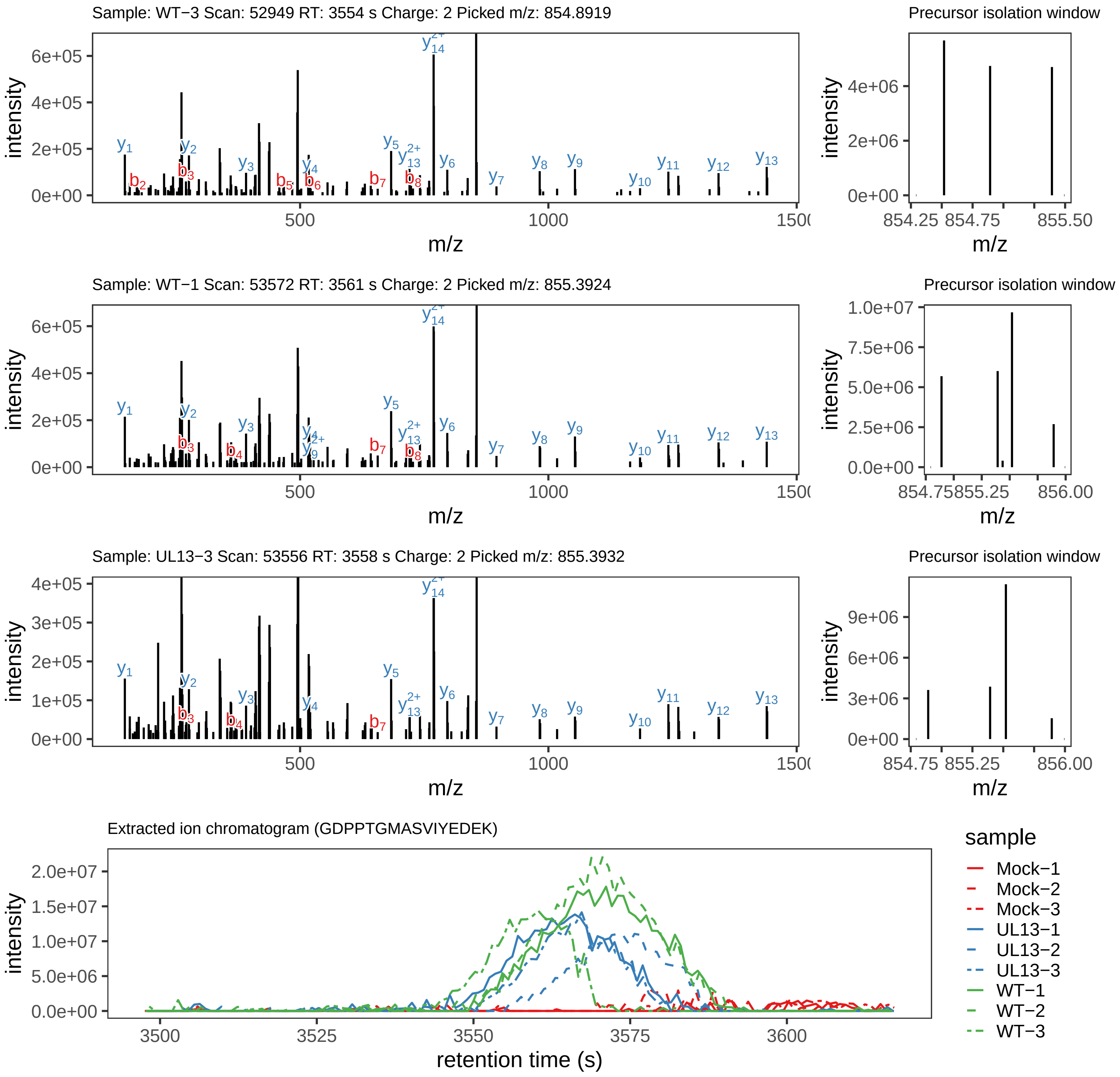

### ILGSPPNPLKPPEIEPPQMSSTPGR.global.pdf

#### Peptide ILGS[+80]PPN[+1]PLKPPEIEPPQM[+16]SSTPGR

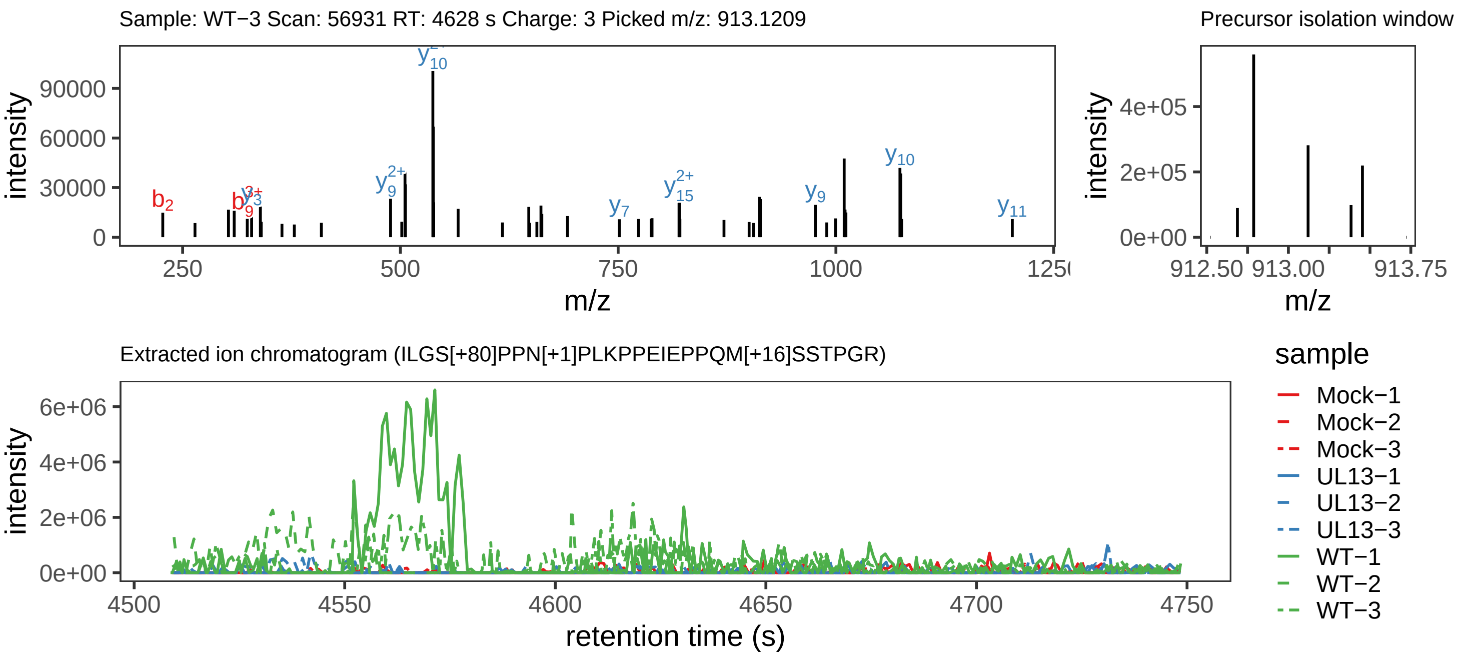

Supplement: Supplemental File 5 [file 737029_file08.zip › phosphopeptide_validation_figs_20240531/ILGSPPNPLKPPEIEPPQMSSTPGR/ILGS[+80]PPN[+1]PLKPPEIEPPQM[+16]SSTPGR.phospho.pdf]

#### Peptide ILGS[+80]PPN[+1]PLKPPEIEPPQMSSTPGR

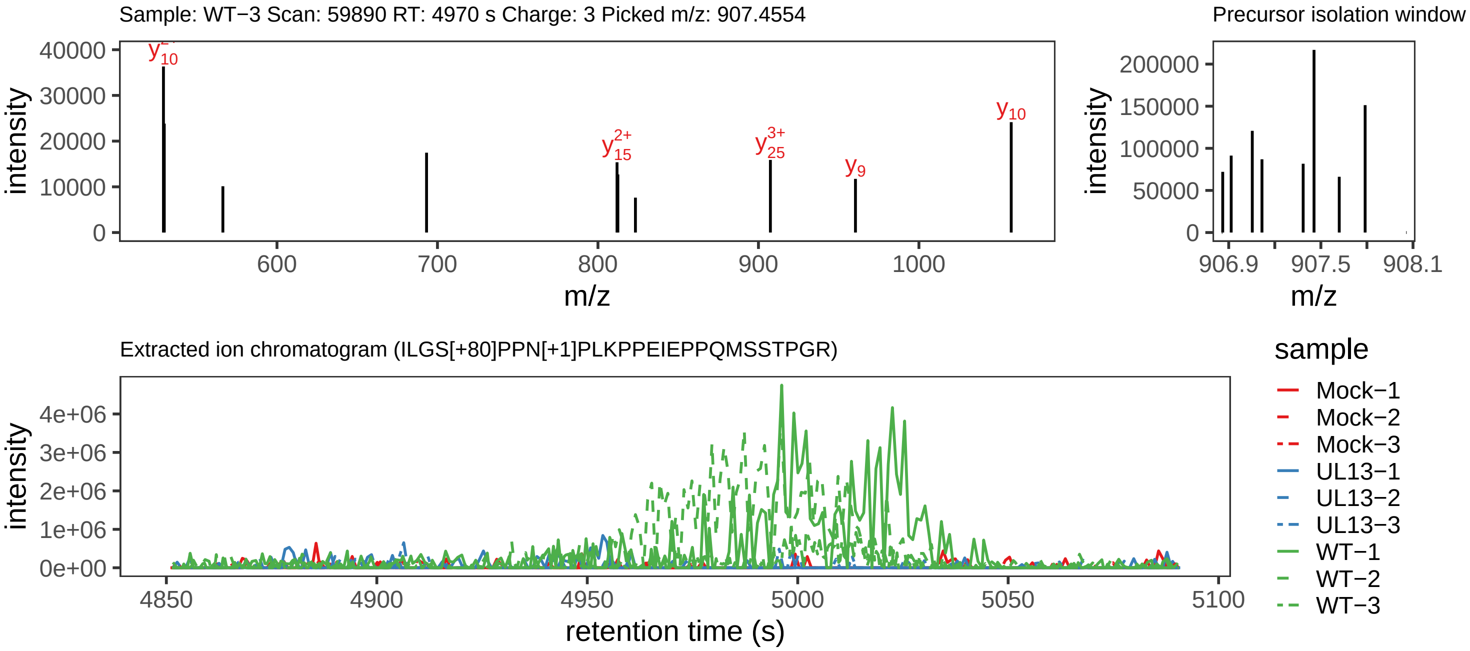

Supplement: Supplemental File 5 [file 737029_file08.zip › phosphopeptide_validation_figs_20240531/ILGSPPNPLKPPEIEPPQMSSTPGR/ILGS[+80]PPN[+1]PLKPPEIEPPQMSSTPGR.phospho.pdf]

#### Peptide ILGS[+80]PPNPLKPPEIEPPQM[+16]SSTPGR

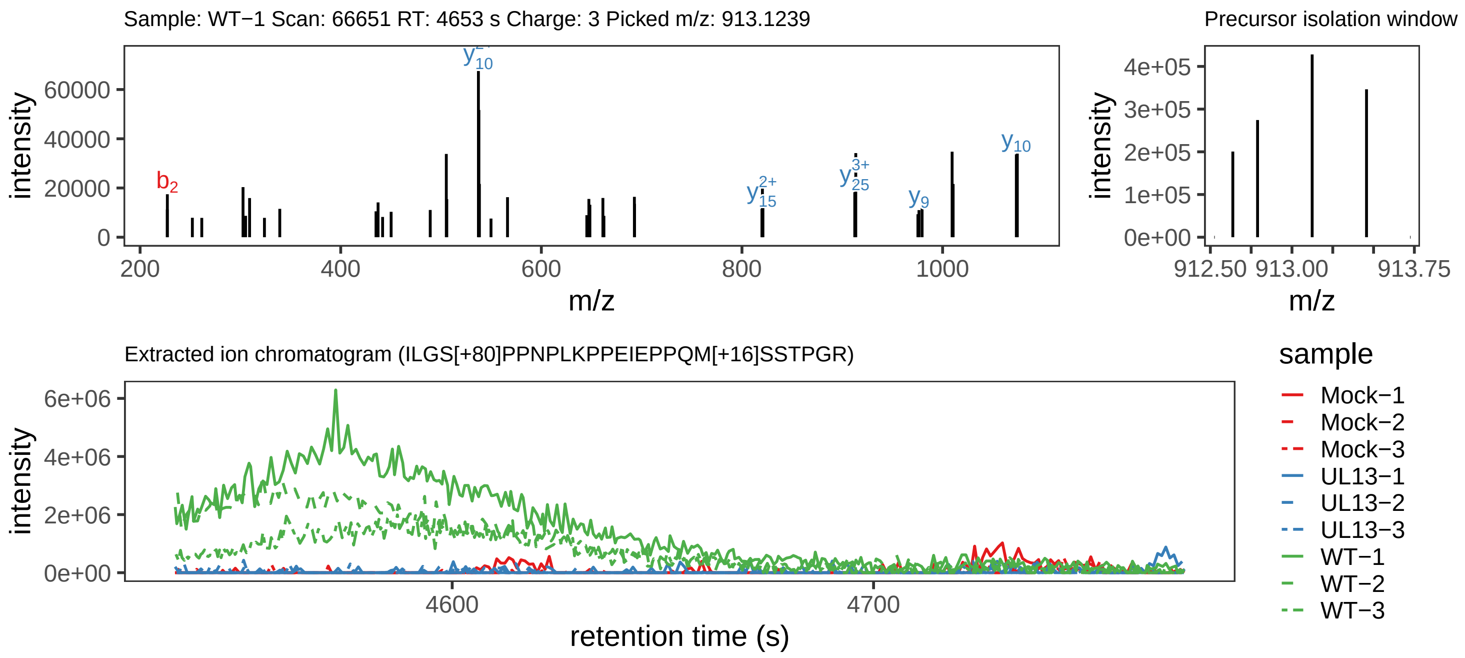

Supplement: Supplemental File 5 [file 737029_file08.zip › phosphopeptide_validation_figs_20240531/ILGSPPNPLKPPEIEPPQMSSTPGR/ILGS[+80]PPNPLKPPEIEPPQM[+16]SSTPGR.phospho.pdf]

#### Peptide ILGS[+80]PPNPLKPPEIEPPQMSSTPGR

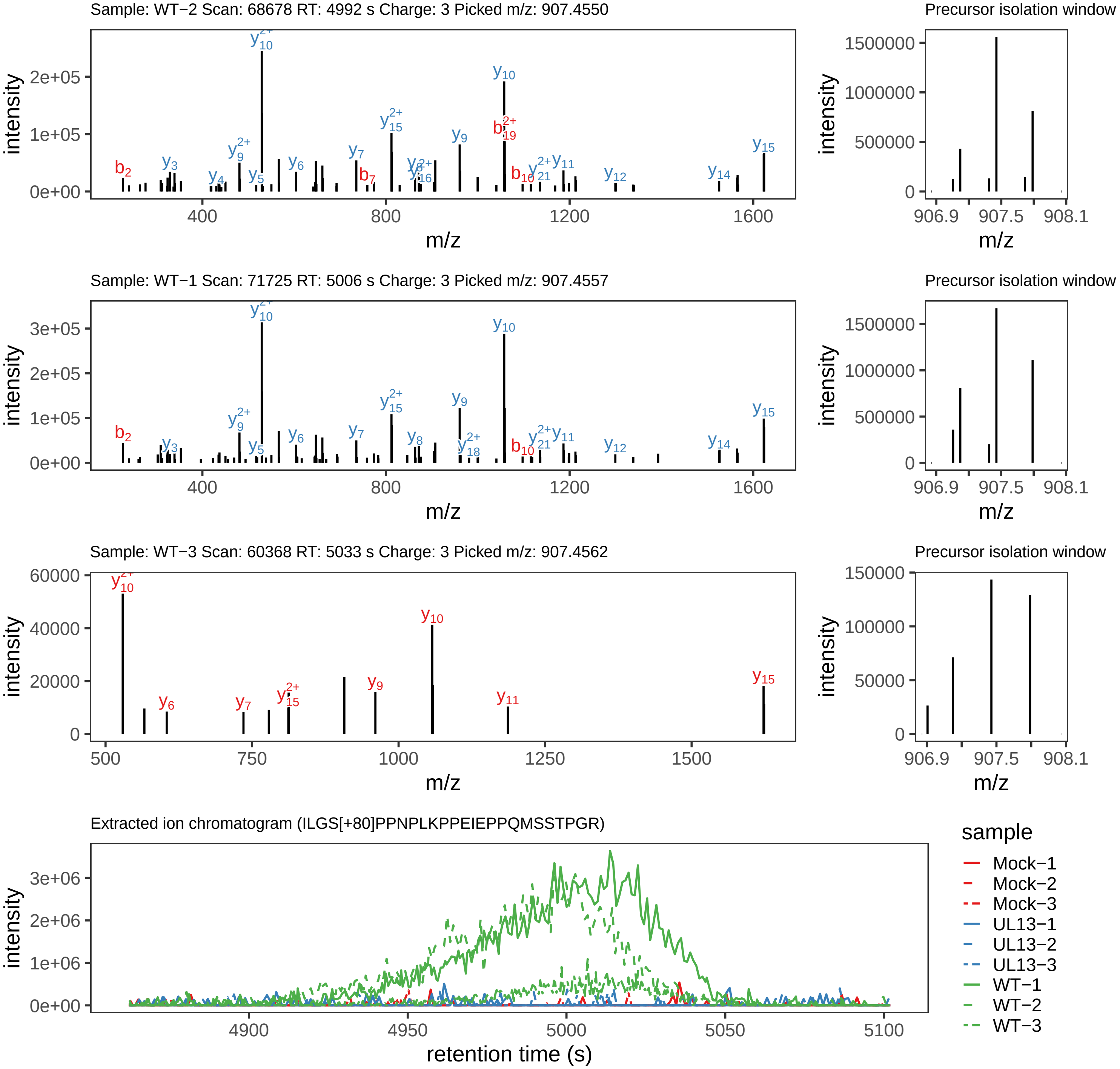

Supplement: Supplemental File 5 [file 737029_file08.zip › phosphopeptide_validation_figs_20240531/ILGSPPNPLKPPEIEPPQMSSTPGR/ILGS[+80]PPNPLKPPEIEPPQMSSTPGR.phospho.pdf]

#### Peptide ILGSPPN[+1]PLKPPEIEPPQMSSTPGR

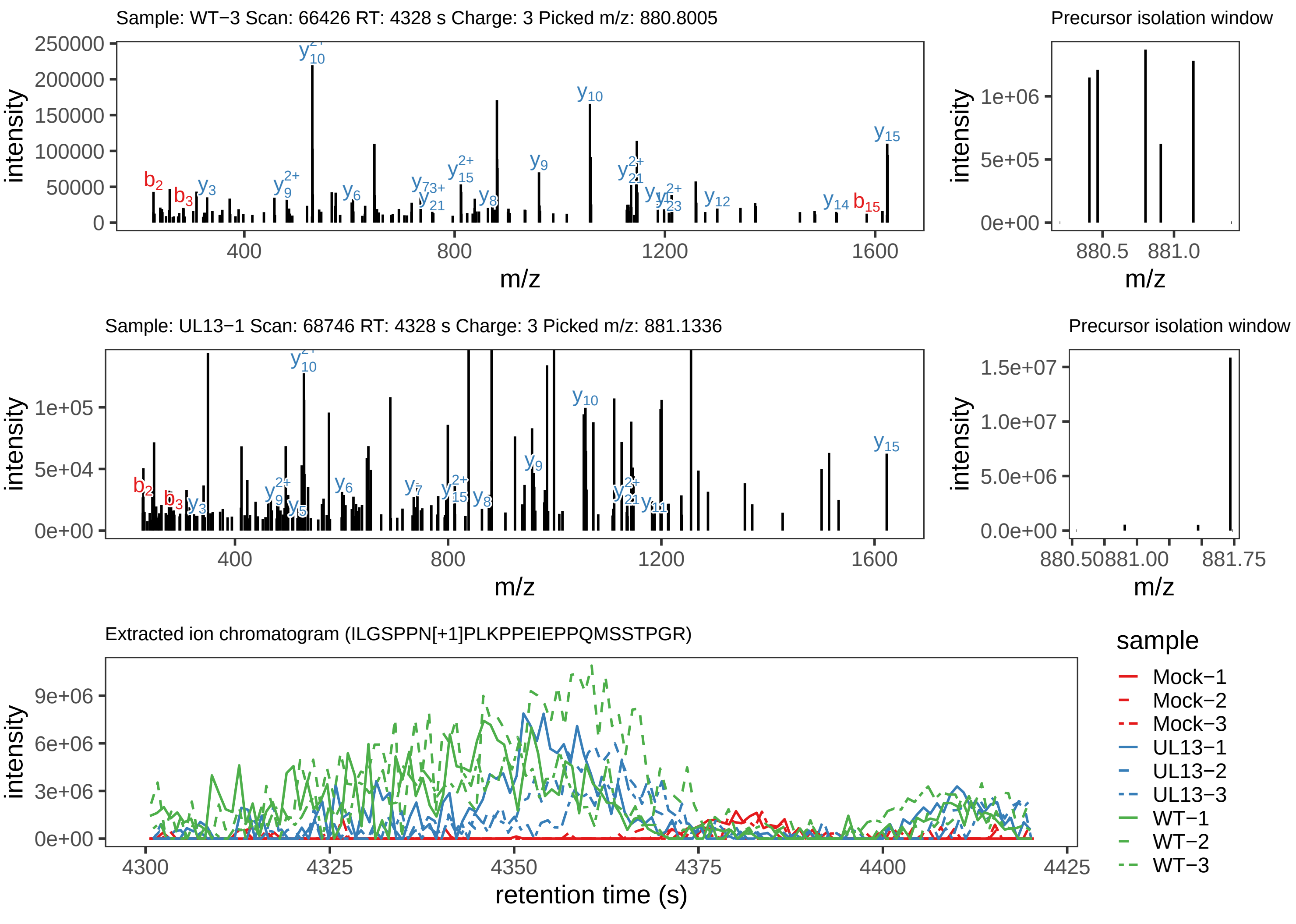

Supplement: Supplemental File 5 [file 737029_file08.zip › phosphopeptide_validation_figs_20240531/ILGSPPNPLKPPEIEPPQMSSTPGR/ILGSPPN[+1]PLKPPEIEPPQMSSTPGR.global.pdf]

#### Peptide ILGSPPNPLKPPEIEPPQMSSTPGR

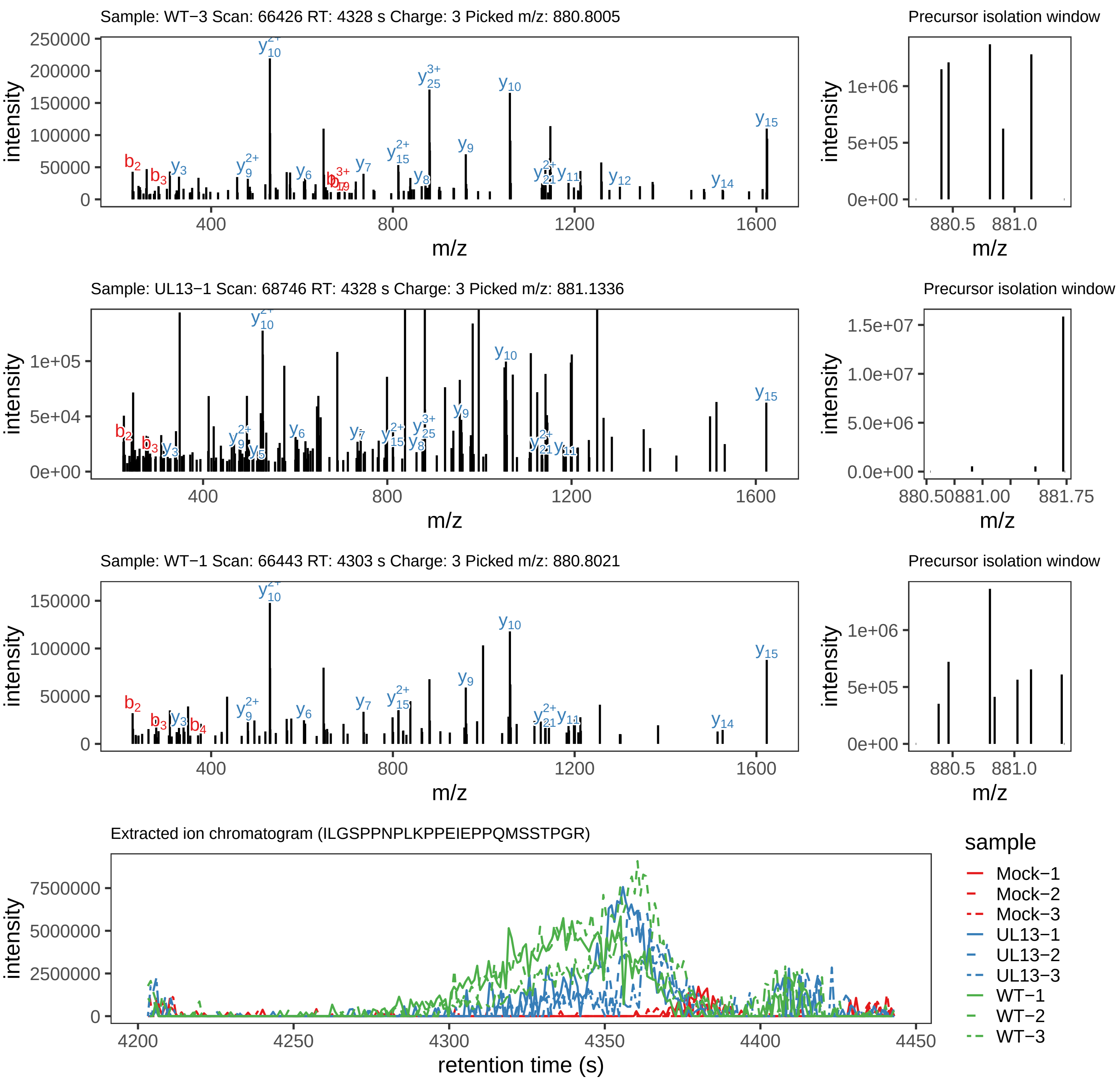

### LTPMWNK.global.pdf

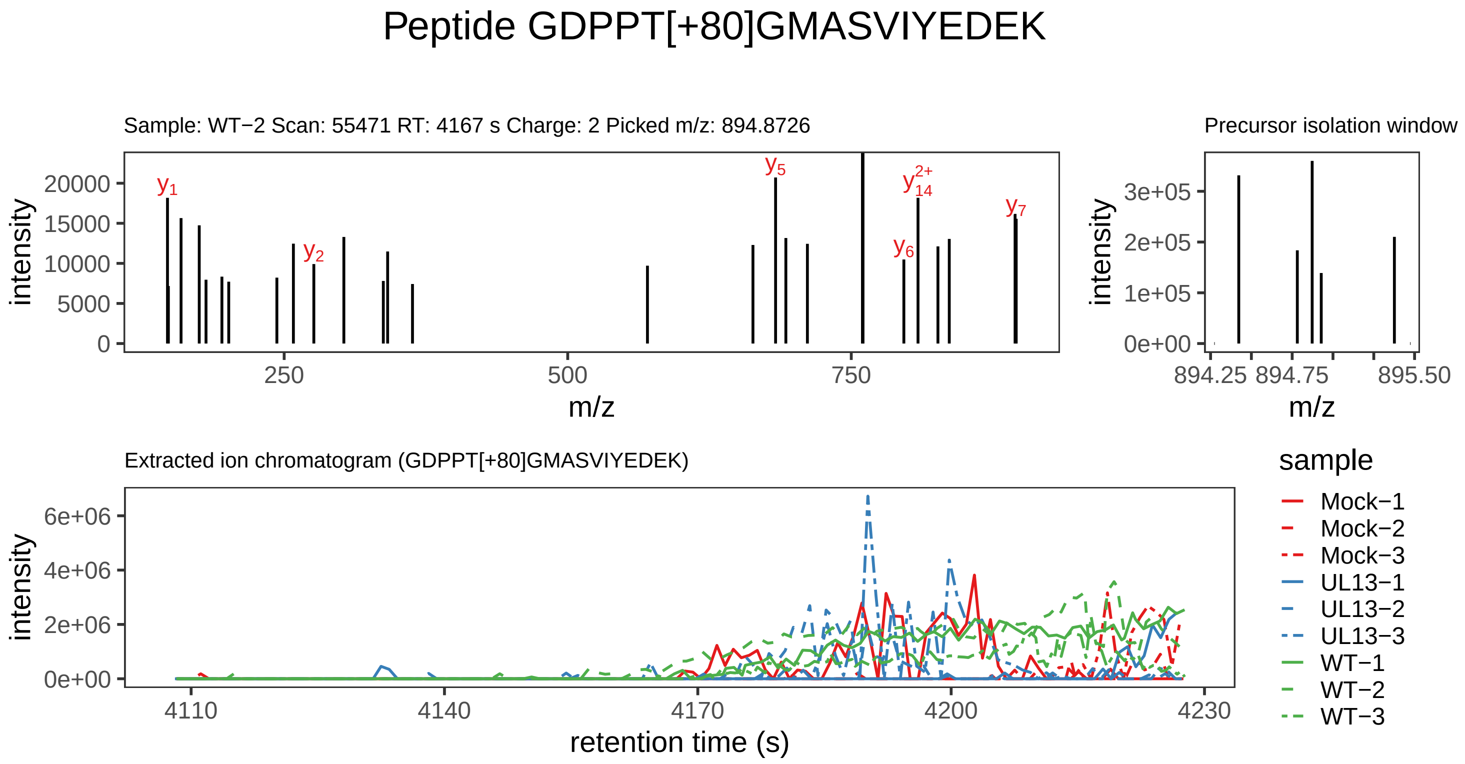

Supplement: Supplemental File 5 [file 737029_file08.zip › phosphopeptide_validation_figs_20240531/GDPPTGMASVIYEDEK/GDPPT[+80]GMASVIYEDEK.phospho.pdf]

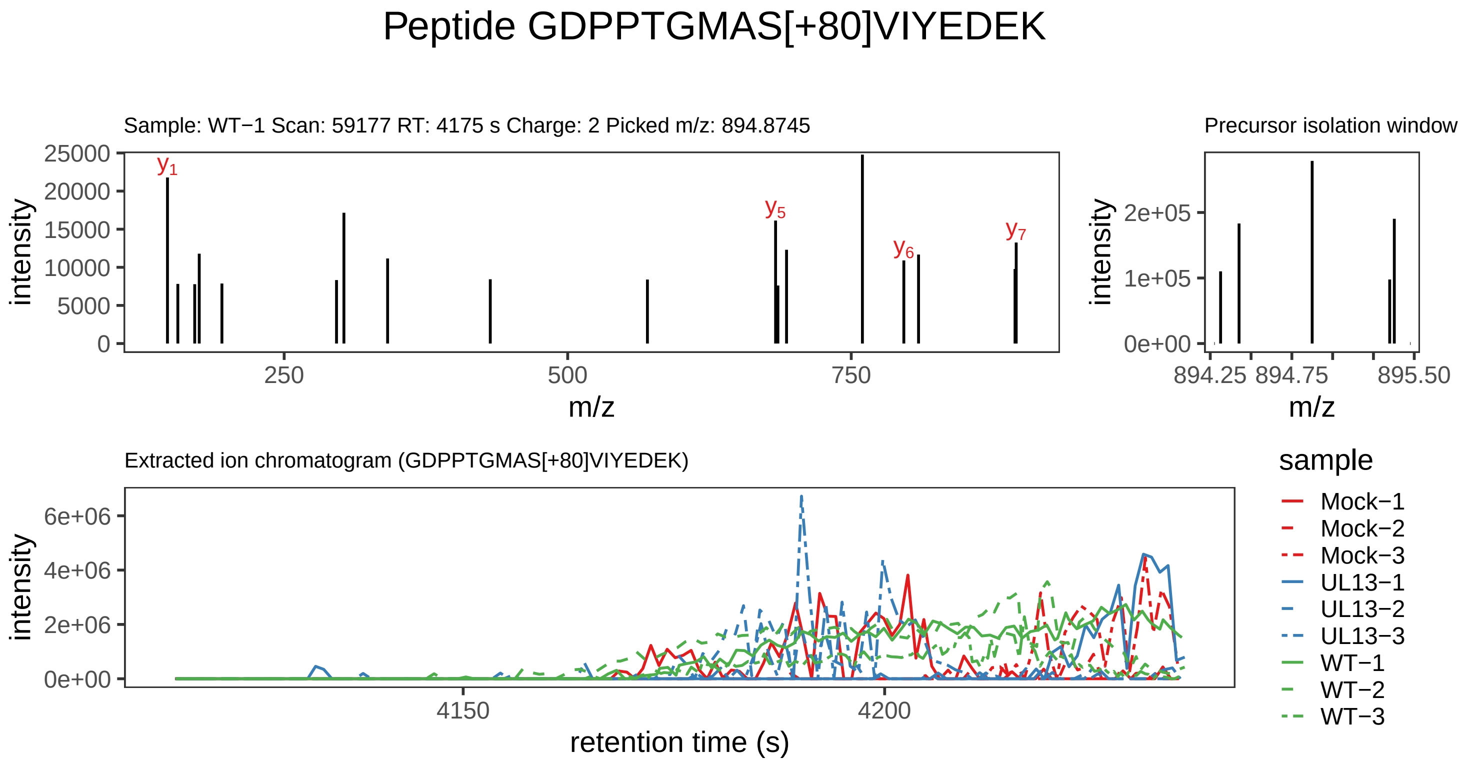

Supplement: Supplemental File 5 [file 737029_file08.zip › phosphopeptide_validation_figs_20240531/GDPPTGMASVIYEDEK/GDPPTGMAS[+80]VIYEDEK.phospho.pdf]

#### Peptide LTPM[+16]WNK

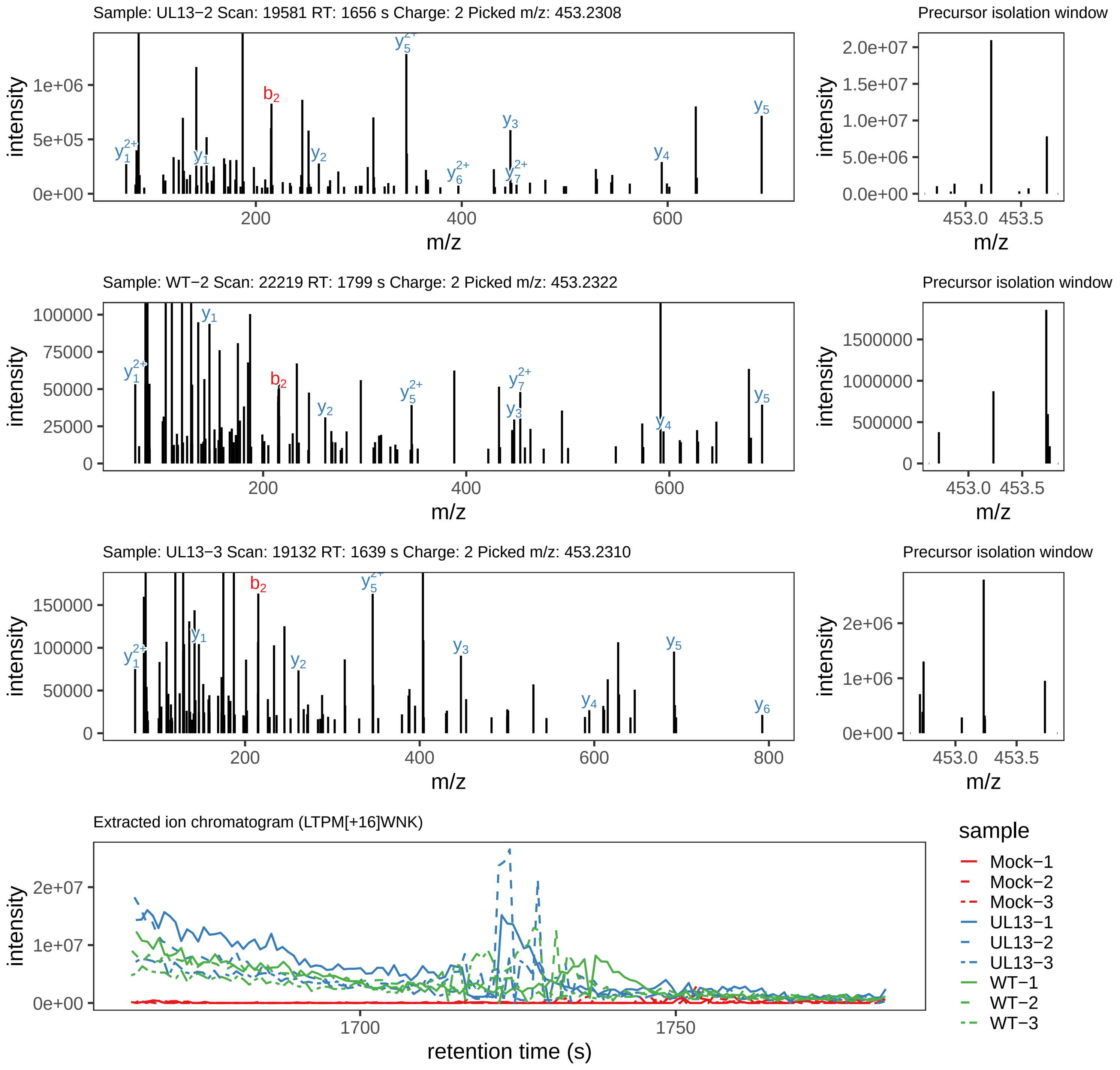

Supplement: Supplemental File 5 [file 737029_file08.zip › phosphopeptide_validation_figs_20240531/LTPMWNK/LTPM[+16]WNK.global.pdf]

#### Peptide LTPMWNK

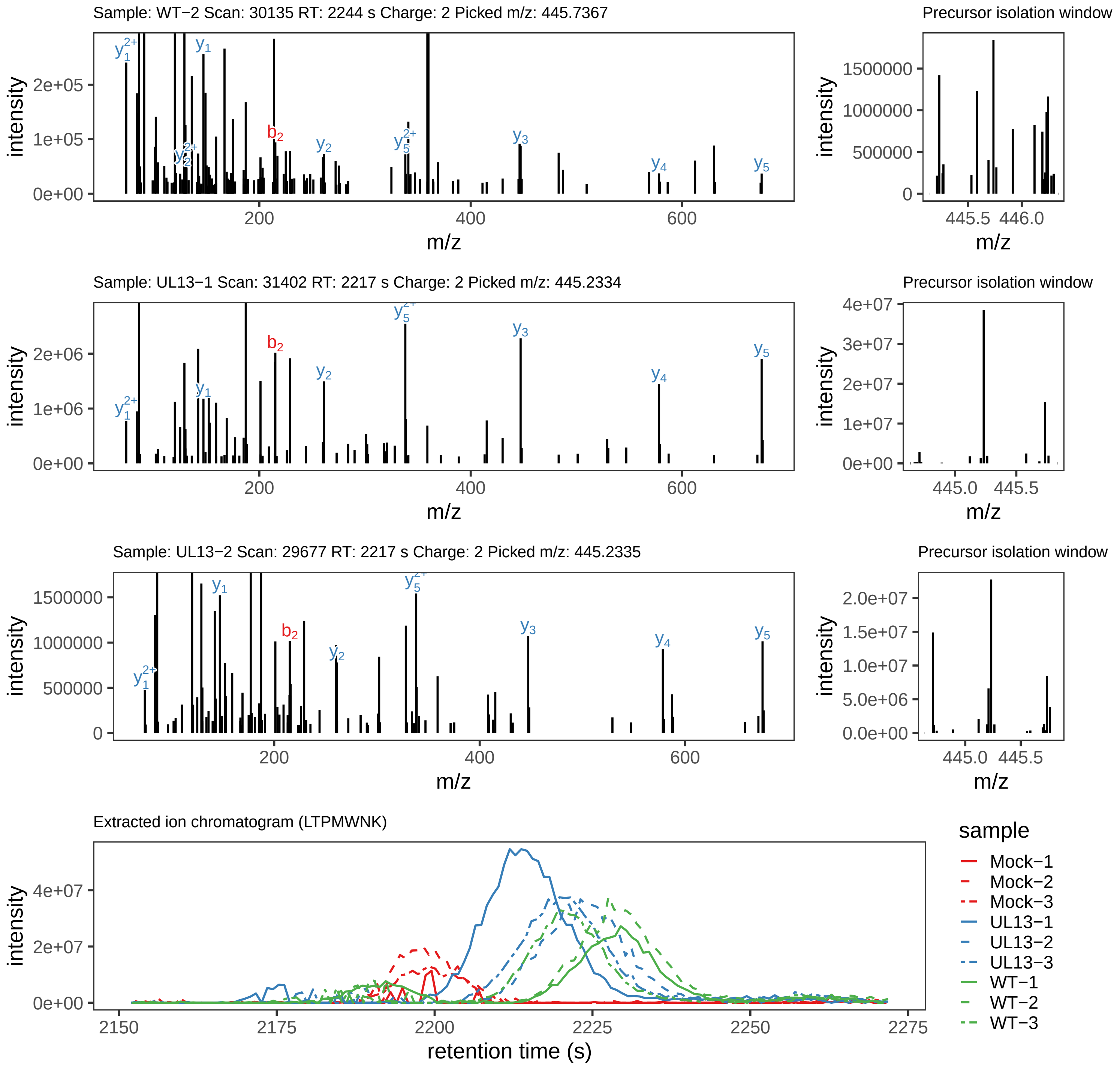

### MTVTLNEYDISASPFHPTDPTRK.global.pdf

#### Peptide LT[+80]PM[+16]WNK

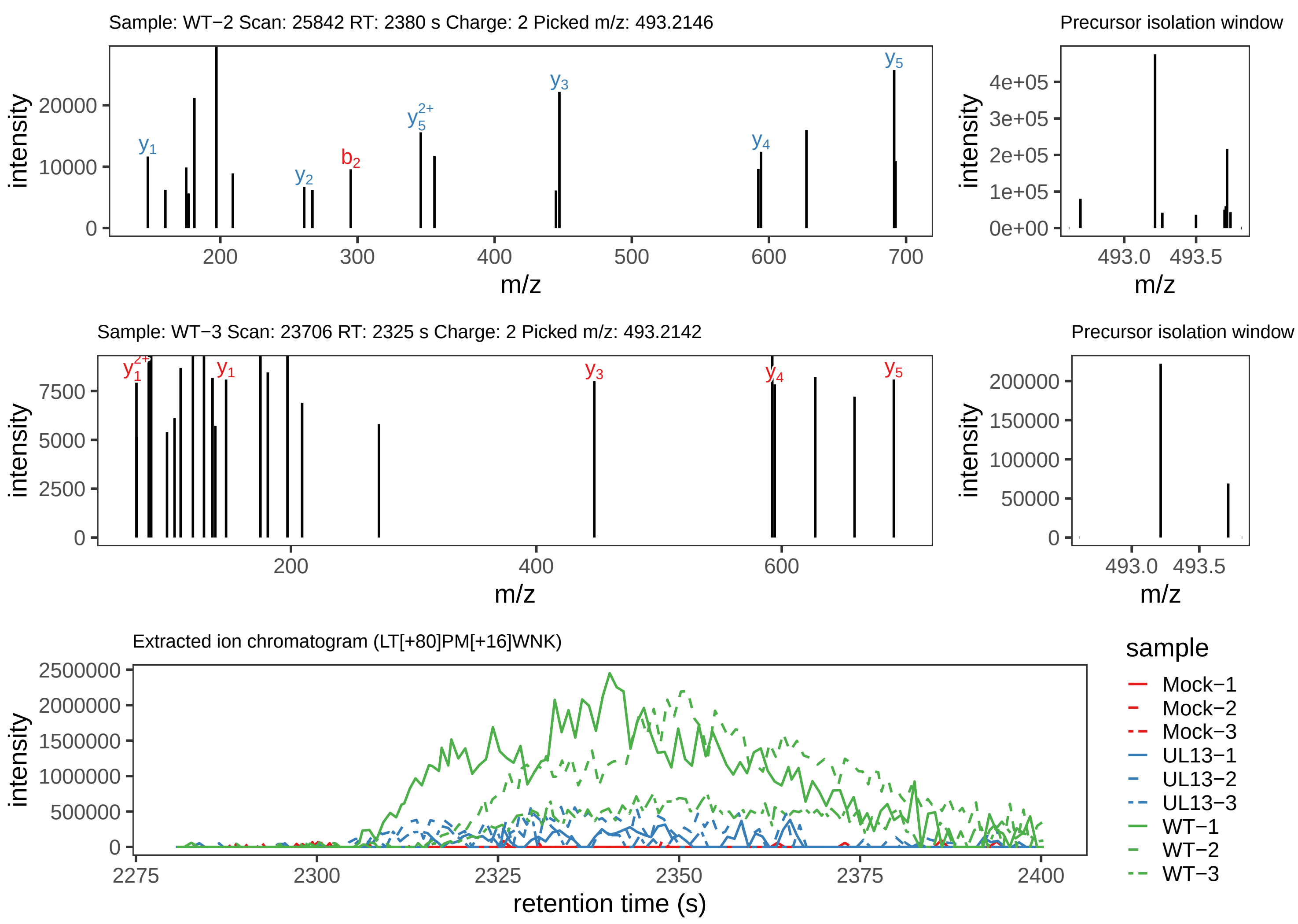

Supplement: Supplemental File 5 [file 737029_file08.zip › phosphopeptide_validation_figs_20240531/LTPMWNK/LT[+80]PM[+16]WNK.phospho.pdf]

#### Peptide M[+16]YTSPPPENALAWK

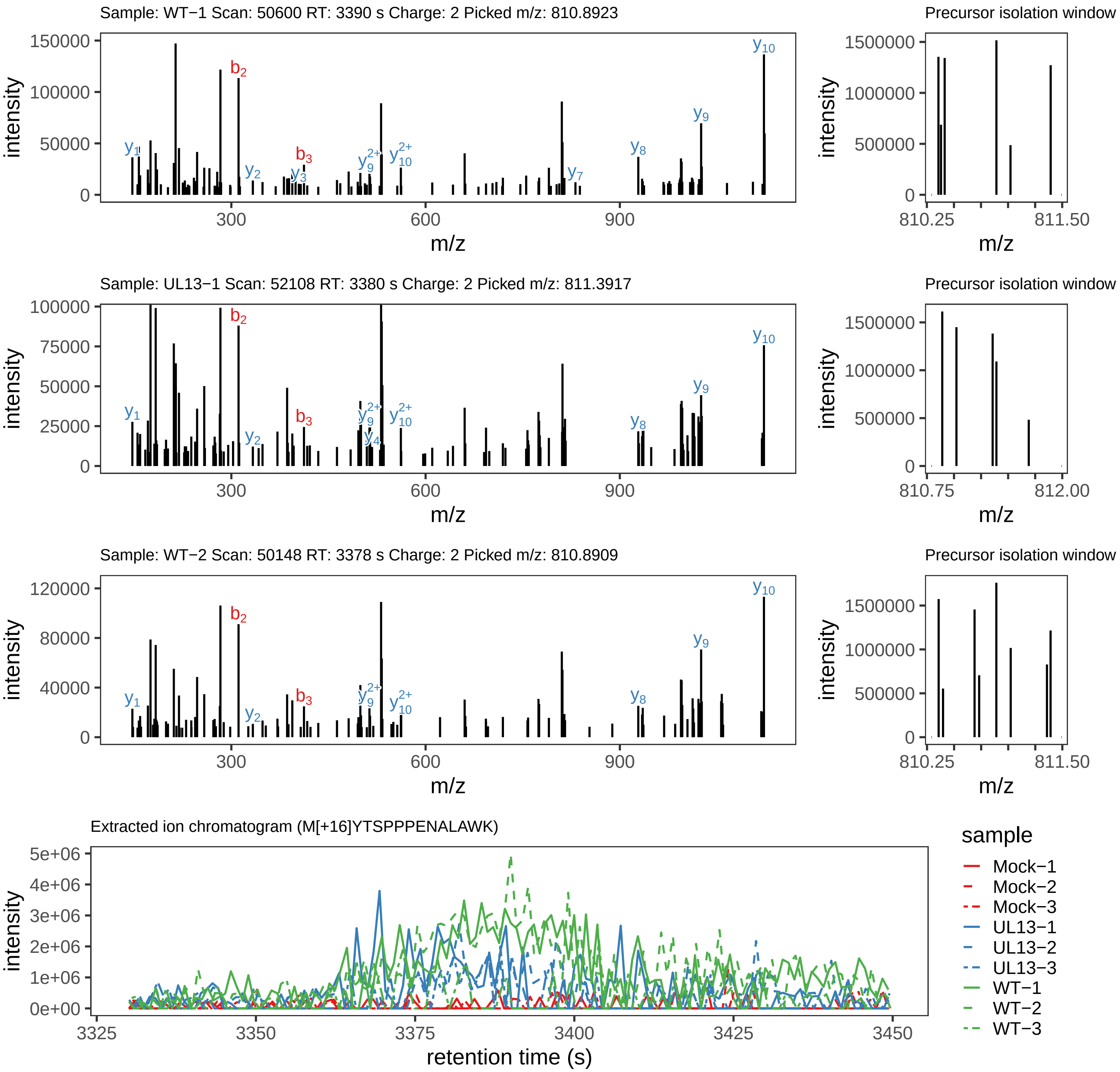

Supplement: Supplemental File 5 [file 737029_file08.zip › phosphopeptide_validation_figs_20240531/MYTSPPPENALAWK/M[+16]YTSPPPENALAWK.global.pdf]

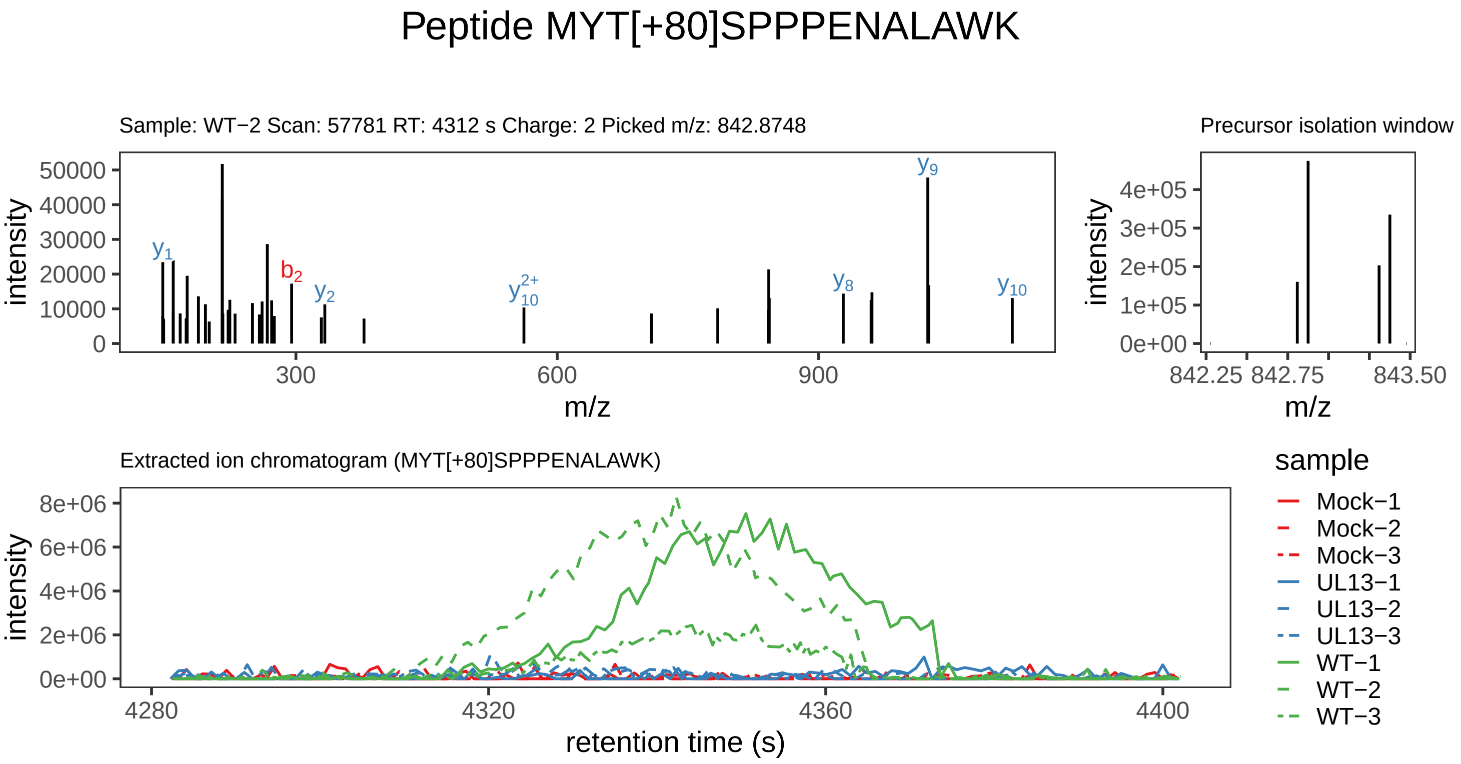

Supplement: Supplemental File 5 [file 737029_file08.zip › phosphopeptide_validation_figs_20240531/MYTSPPPENALAWK/MYT[+80]SPPPENALAWK.phospho.pdf]

#### Peptide M[+16]YT[+80]SPPPENALAWK

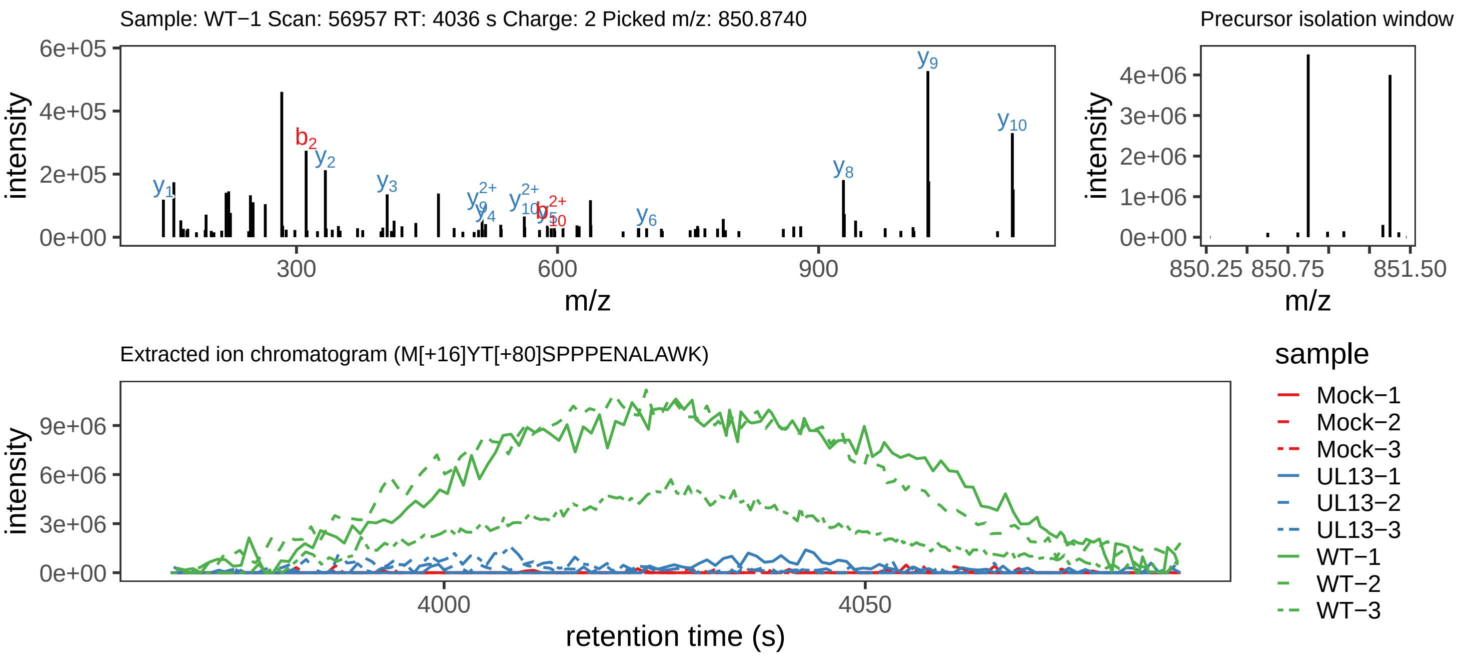

Supplement: Supplemental File 5 [file 737029_file08.zip › phosphopeptide_validation_figs_20240531/MYTSPPPENALAWK/M[+16]YT[+80]SPPPENALAWK.phospho.pdf]

#### Peptide M[+16]YTS[+80]PPPENALAWK

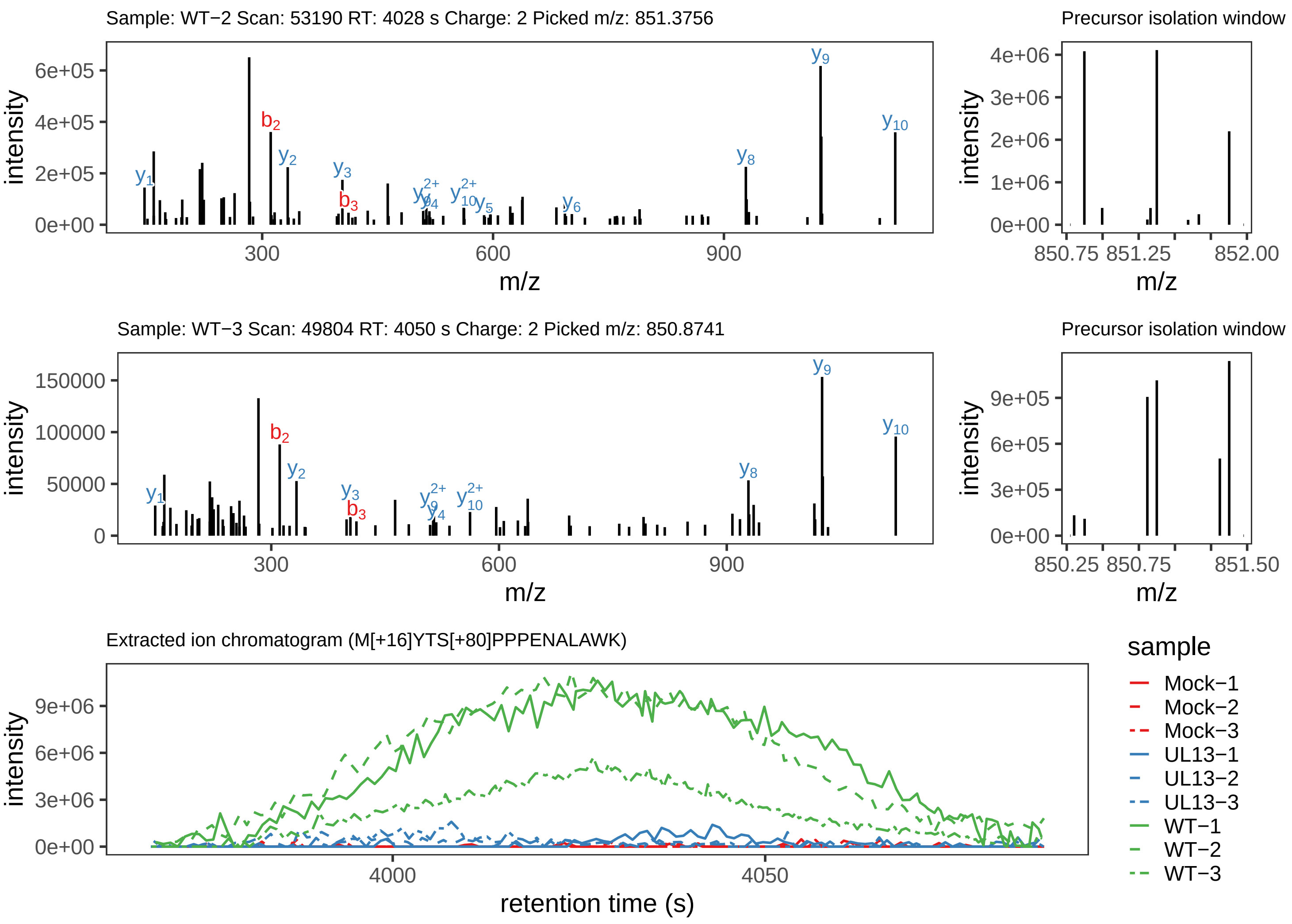

Supplement: Supplemental File 5 [file 737029_file08.zip › phosphopeptide_validation_figs_20240531/MYTSPPPENALAWK/M[+16]YTS[+80]PPPENALAWK.phospho.pdf]

#### Peptide MYTS[+80]PPPENALAWK

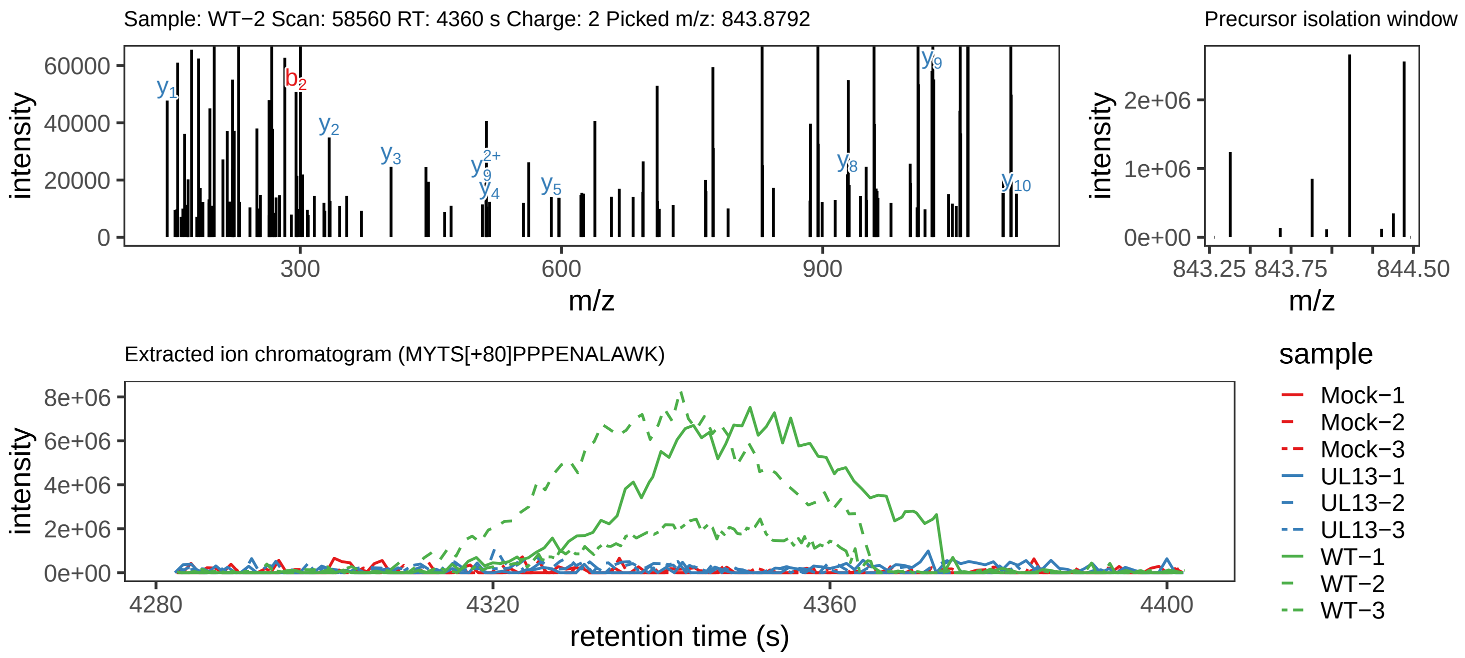

Supplement: Supplemental File 5 [file 737029_file08.zip › phosphopeptide_validation_figs_20240531/MYTSPPPENALAWK/MYTS[+80]PPPENALAWK.phospho.pdf]

#### Peptide GPETSPS[+80]NEHIIISPPRNPPSNTTHR

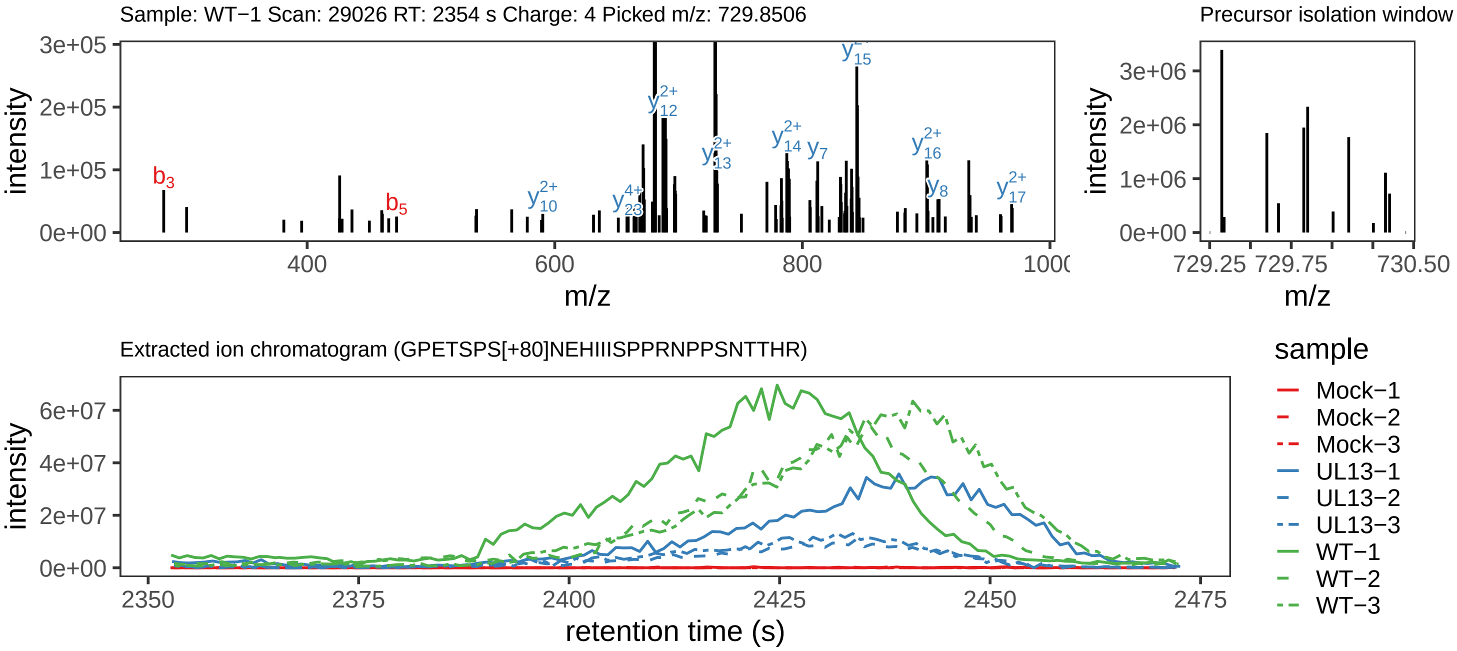

Supplement: Supplemental File 5 [file 737029_file08.zip › phosphopeptide_validation_figs_20240531/GPETSPSNEHIIISPPRNPPSNTTHR/GPETSPS[+80]NEHIIISPPRNPPSNTTHR.phospho.pdf]

#### Peptide GPETS[+80]PSNEHIIIS[+80]PPRNPPSNTTHR

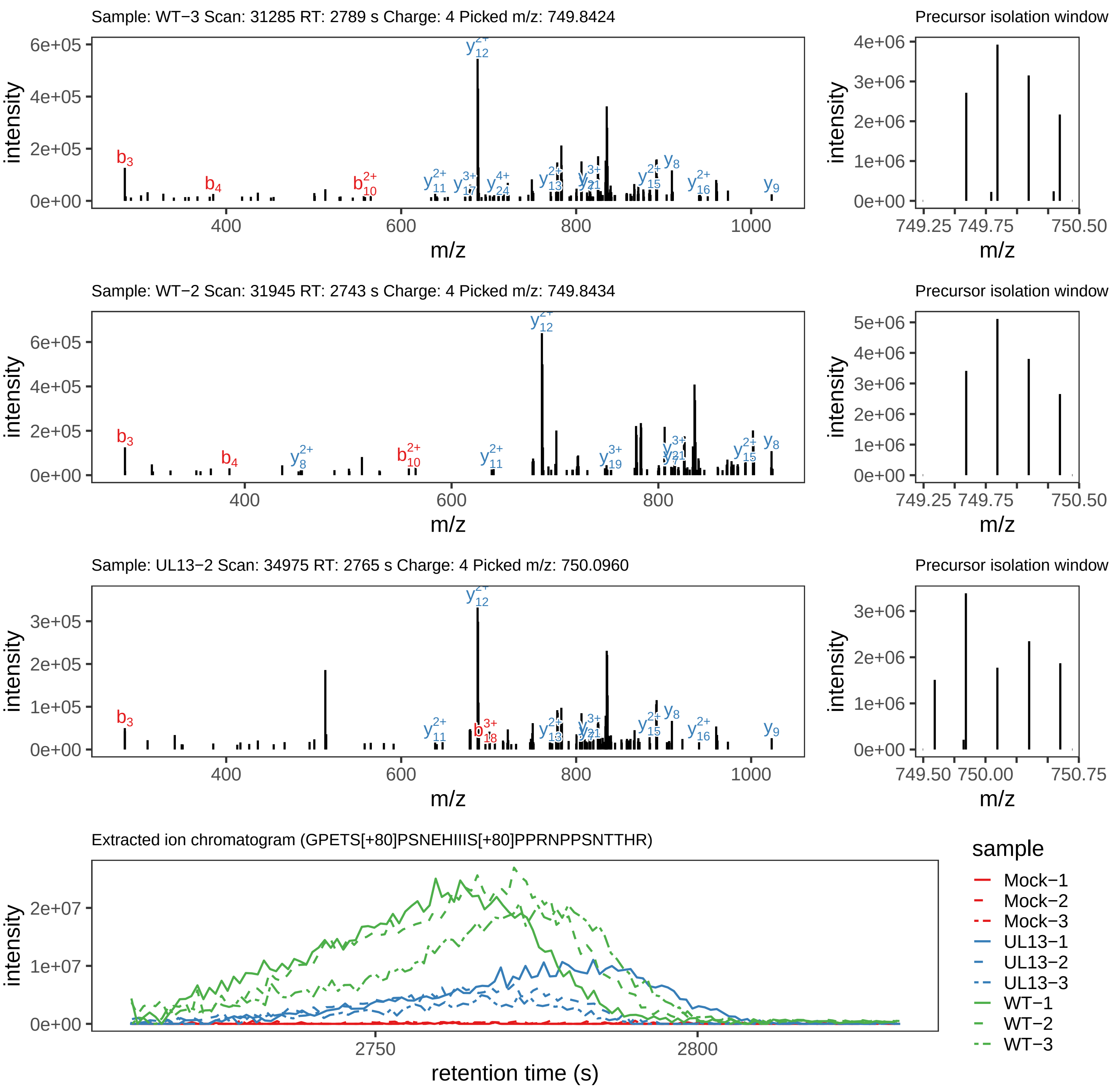

Supplement: Supplemental File 5 [file 737029_file08.zip › phosphopeptide_validation_figs_20240531/GPETSPSNEHIIISPPRNPPSNTTHR/GPETS[+80]PSNEHIIIS[+80]PPRNPPSNTTHR.phospho.pdf]

#### Peptide GPET[+80]SPSNEHIIIS[+80]PPRNPPSNTTHR

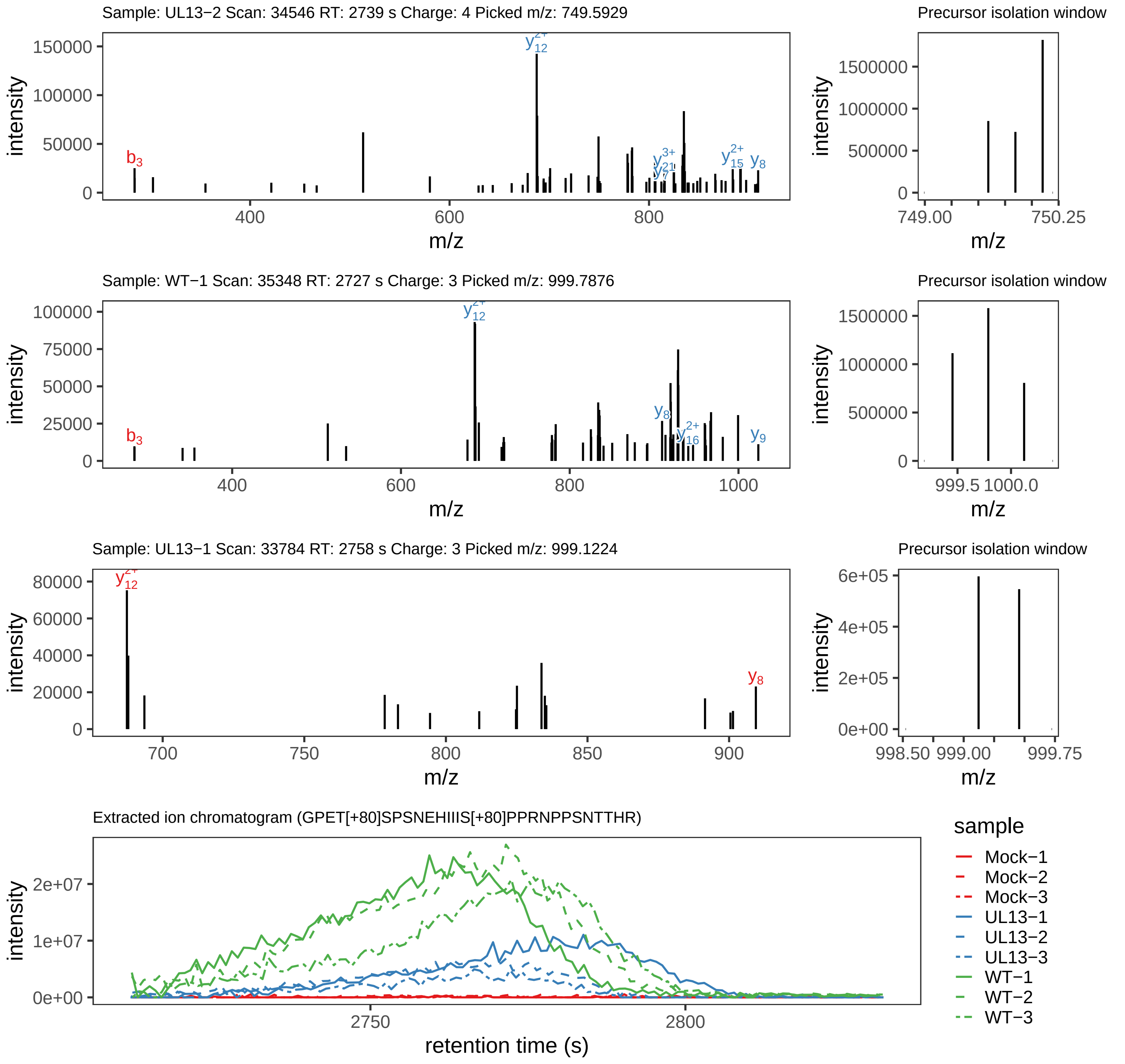

Supplement: Supplemental File 5 [file 737029_file08.zip › phosphopeptide_validation_figs_20240531/GPETSPSNEHIIISPPRNPPSNTTHR/GPET[+80]SPSNEHIIIS[+80]PPRNPPSNTTHR.phospho.pdf]

#### Peptide GPET[+80]SPSNEHIIIS[+80]PPRNPPSN[+1]TTHR

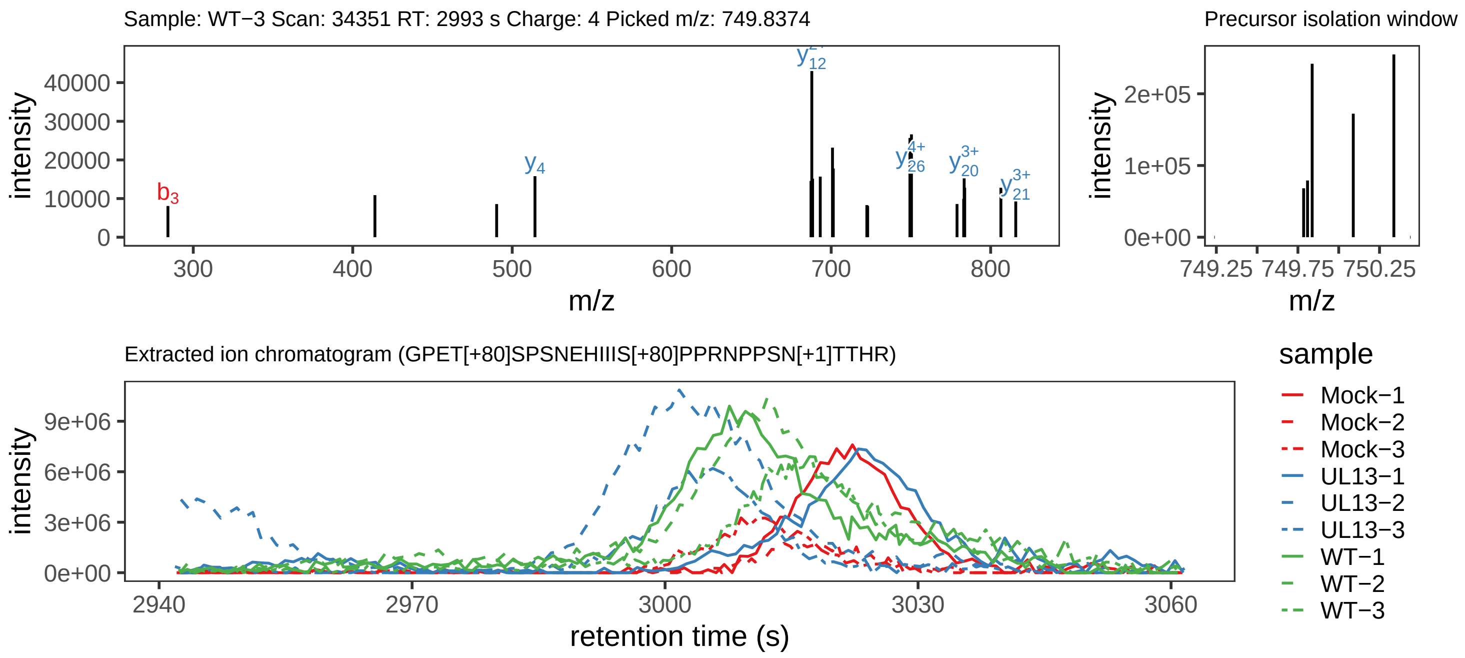

Supplement: Supplemental File 5 [file 737029_file08.zip › phosphopeptide_validation_figs_20240531/GPETSPSNEHIIISPPRNPPSNTTHR/GPET[+80]SPSNEHIIIS[+80]PPRNPPSN[+1]TTHR.phospho.pdf]

#### Peptide GPETSPS[+80]NEHIIISPPRN[+1]PPSNTTHR

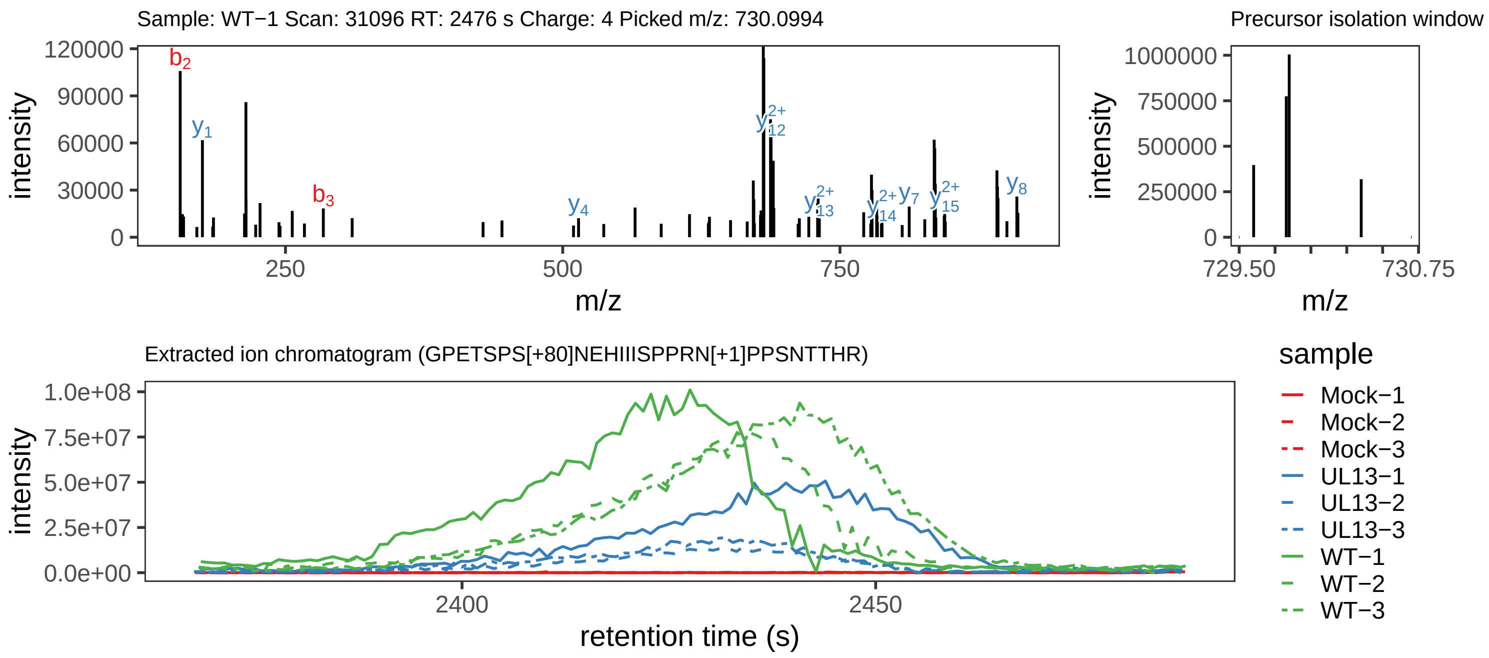

Supplement: Supplemental File 5 [file 737029_file08.zip › phosphopeptide_validation_figs_20240531/GPETSPSNEHIIISPPRNPPSNTTHR/GPETSPS[+80]NEHIIISPPRN[+1]PPSNTTHR.phospho.pdf]

#### Peptide GPETSPSNEHIIIS[+80]PPRN[+1]PPSNTTHR

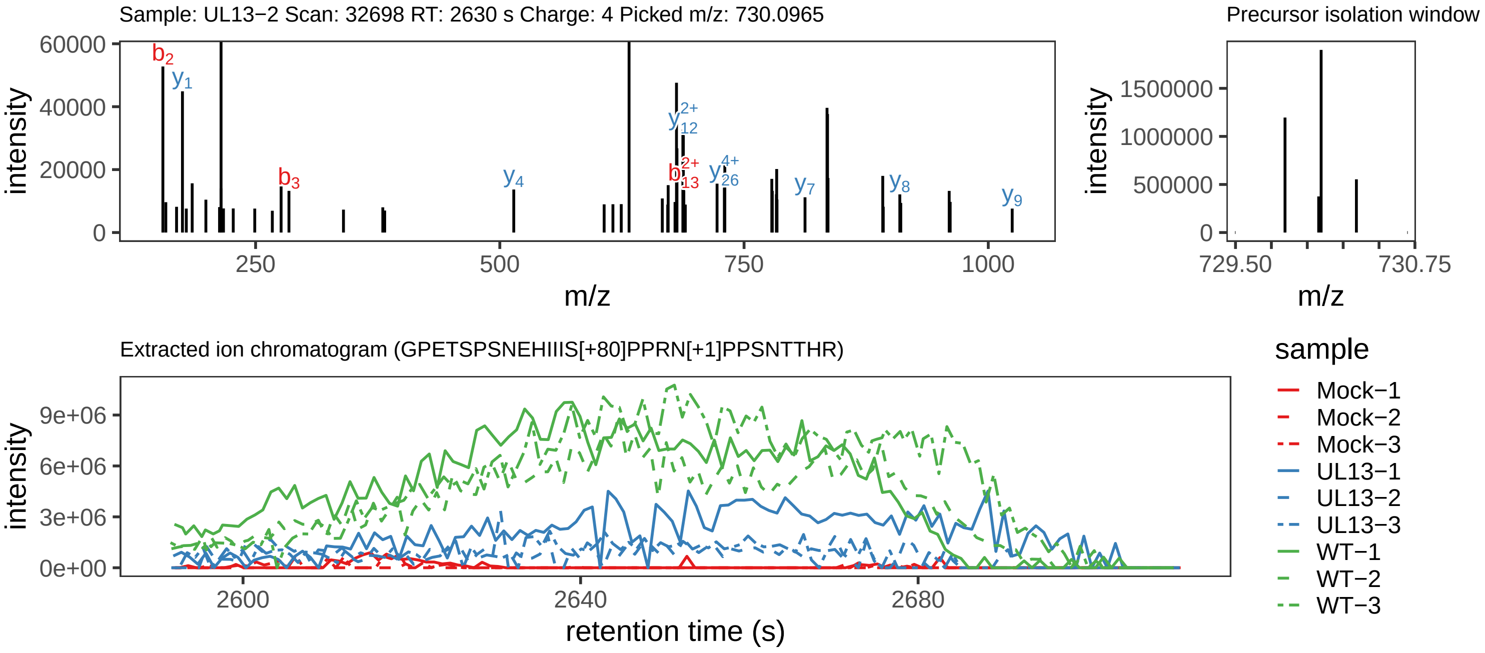

Supplement: Supplemental File 5 [file 737029_file08.zip › phosphopeptide_validation_figs_20240531/GPETSPSNEHIIISPPRNPPSNTTHR/GPETSPSNEHIIIS[+80]PPRN[+1]PPSNTTHR.phospho.pdf]

#### Peptide GPETSPSNEHIIIS[+80]PPRNPPSNTTHR

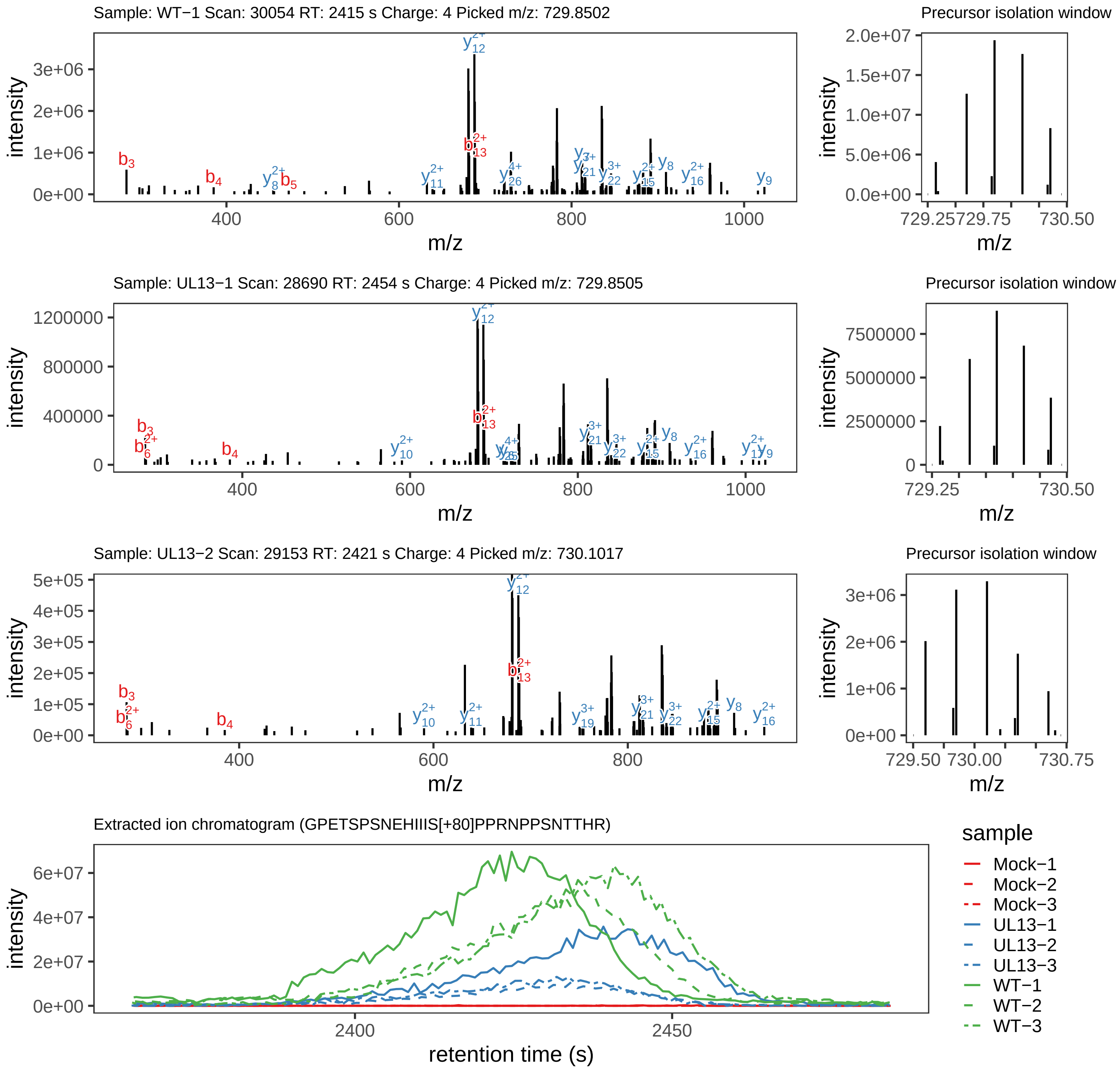

Supplement: Supplemental File 5 [file 737029_file08.zip › phosphopeptide_validation_figs_20240531/GPETSPSNEHIIISPPRNPPSNTTHR/GPETSPSNEHIIIS[+80]PPRNPPSNTTHR.phospho.pdf]

#### Peptide GPETSPS[+80]NEHIIISPPRNPPSN[+1]TTHR

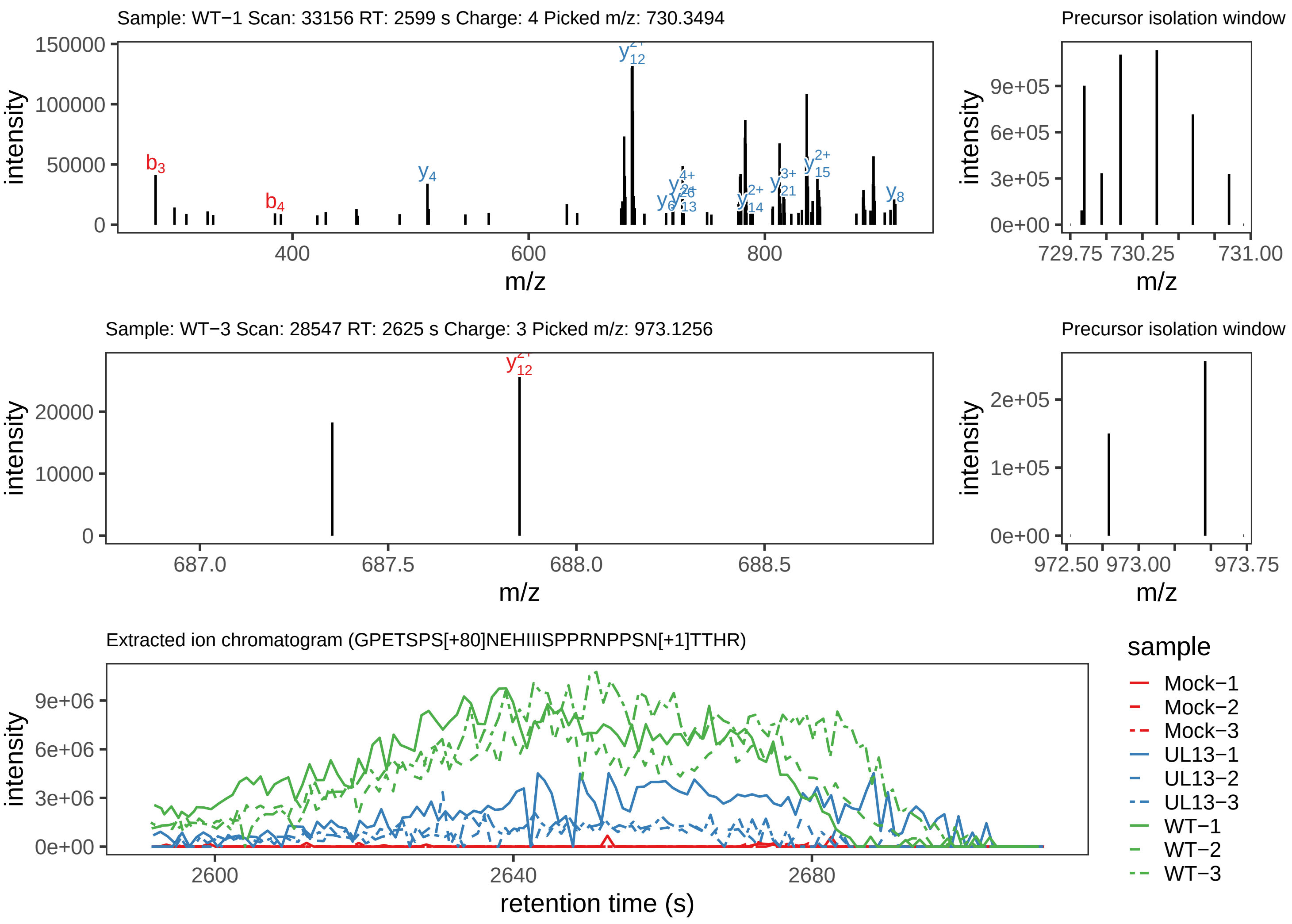

Supplement: Supplemental File 5 [file 737029_file08.zip › phosphopeptide_validation_figs_20240531/GPETSPSNEHIIISPPRNPPSNTTHR/GPETSPS[+80]NEHIIISPPRNPPSN[+1]TTHR.phospho.pdf]

#### Peptide GPETSPSNEHIIIS[+80]PPRNPPSN[+1]TTHR

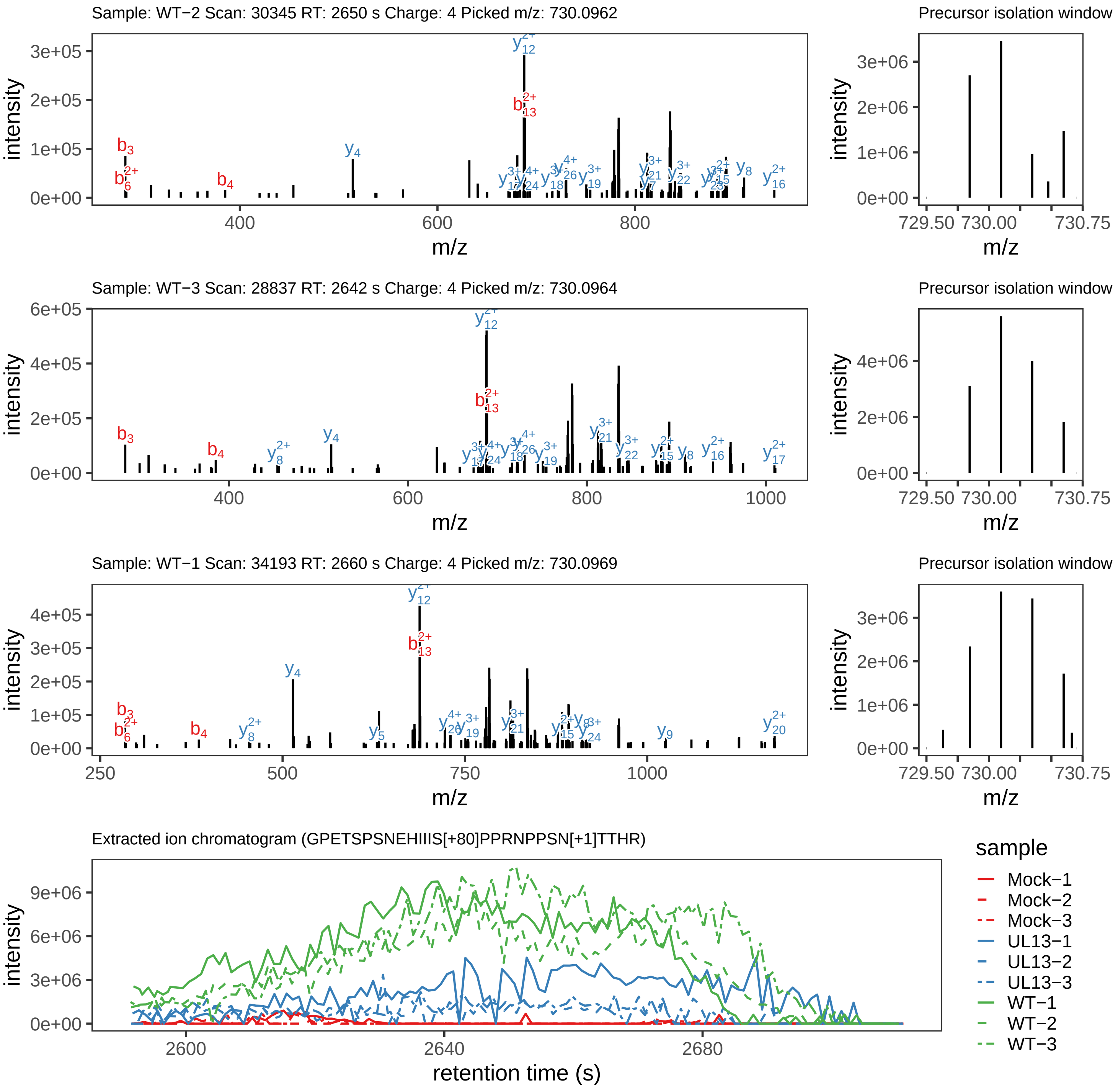

Supplement: Supplemental File 5 [file 737029_file08.zip › phosphopeptide_validation_figs_20240531/GPETSPSNEHIIISPPRNPPSNTTHR/GPETSPSNEHIIIS[+80]PPRNPPSN[+1]TTHR.phospho.pdf]

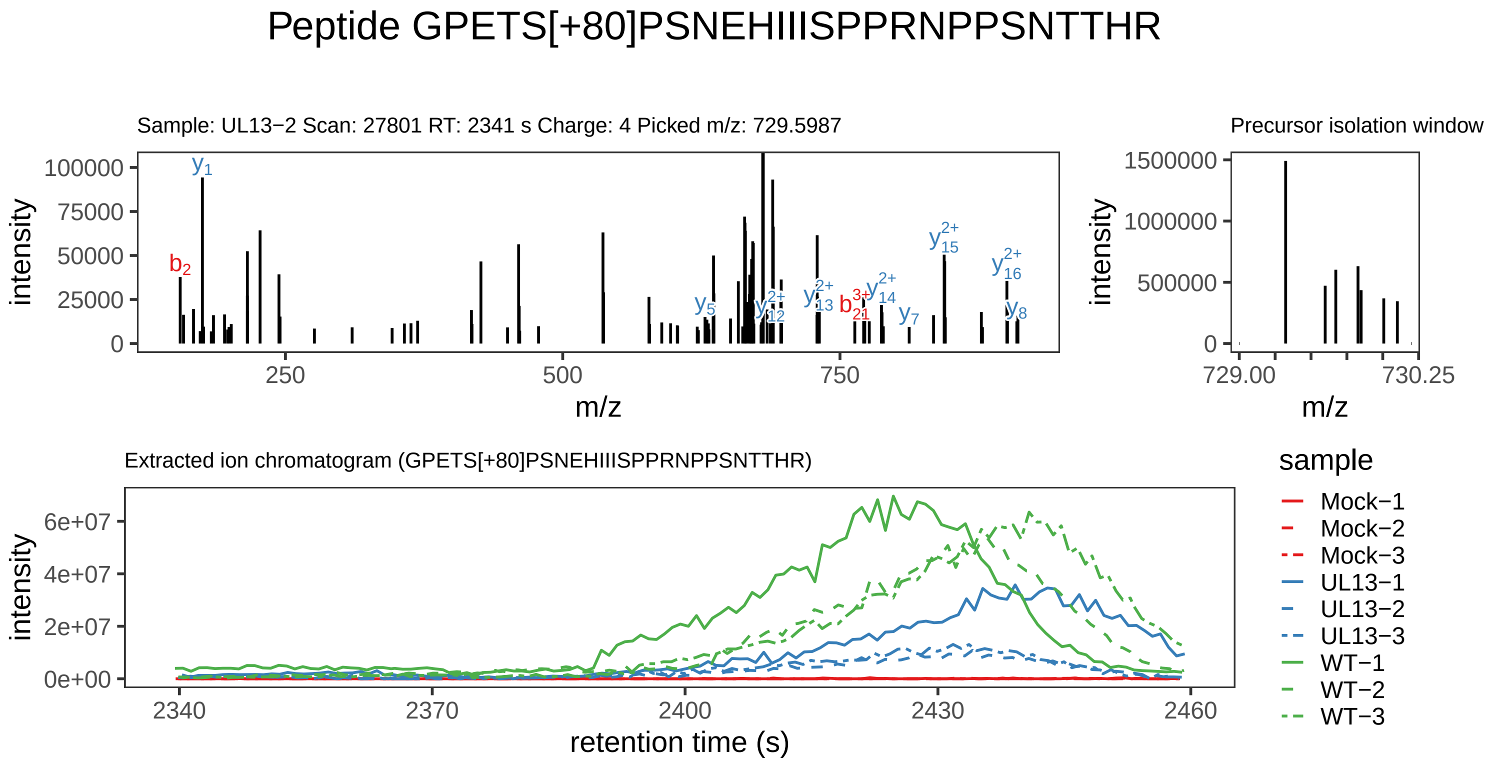

Supplement: Supplemental File 5 [file 737029_file08.zip › phosphopeptide_validation_figs_20240531/GPETSPSNEHIIISPPRNPPSNTTHR/GPETS[+80]PSNEHIIISPPRNPPSNTTHR.phospho.pdf]

#### Peptide GPETS[+80]PSNEHIIIS[+80]PPRNPPSN[+1]TTHR

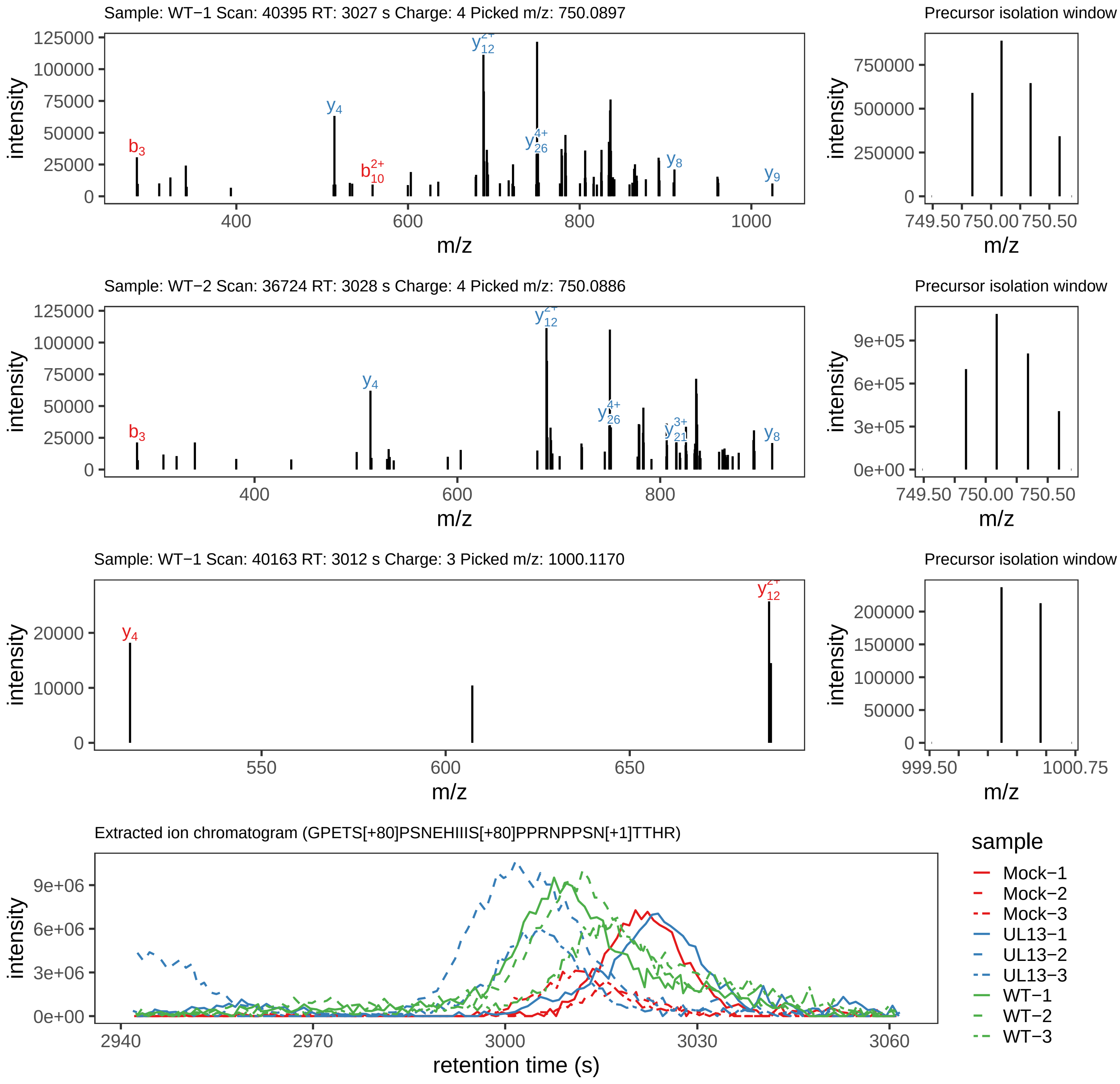

Supplement: Supplemental File 5 [file 737029_file08.zip › phosphopeptide_validation_figs_20240531/GPETSPSNEHIIISPPRNPPSNTTHR/GPETS[+80]PSNEHIIIS[+80]PPRNPPSN[+1]TTHR.phospho.pdf]

#### Peptide GPETS[+80]PSNEHIIIS[+80]PPRN[+1]PPSNTTHR

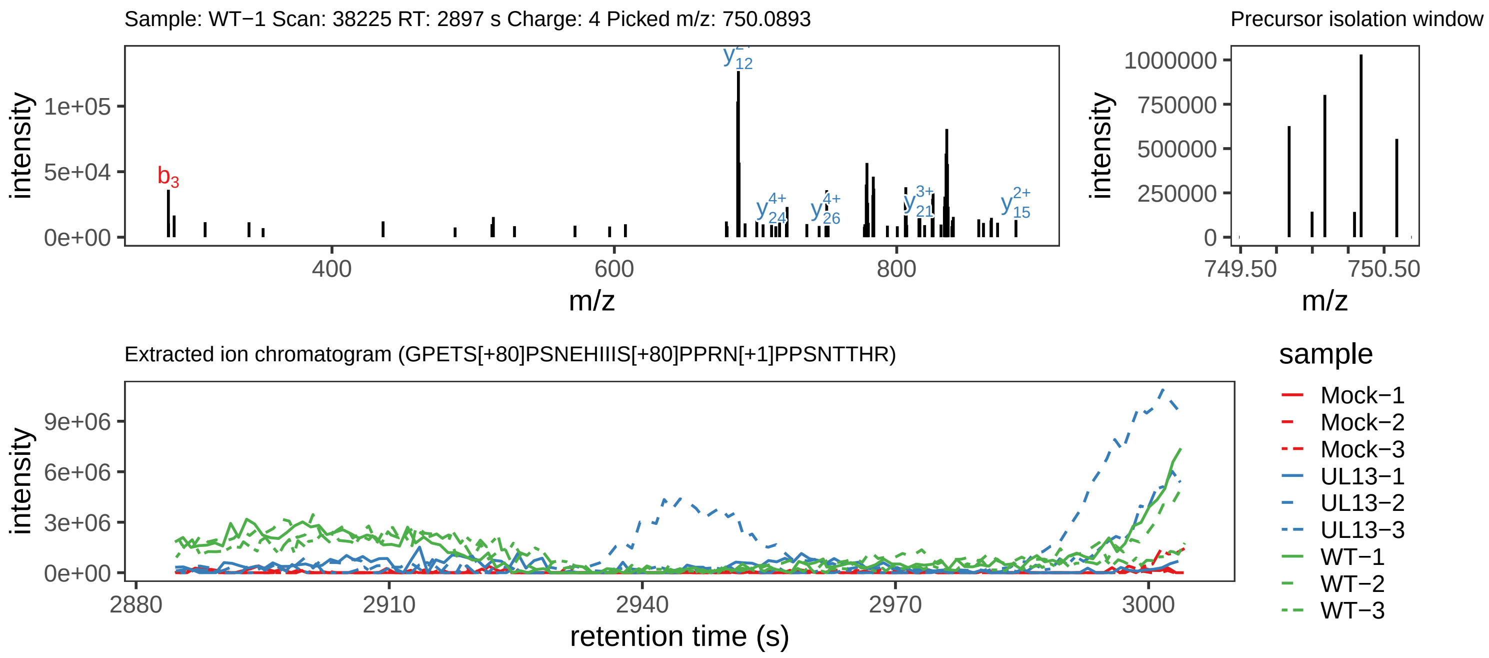

Supplement: Supplemental File 5 [file 737029_file08.zip › phosphopeptide_validation_figs_20240531/GPETSPSNEHIIISPPRNPPSNTTHR/GPETS[+80]PSNEHIIIS[+80]PPRN[+1]PPSNTTHR.phospho.pdf]

#### Peptide GPET[+80]SPSNEHIIISPPRNPPSNTTHR

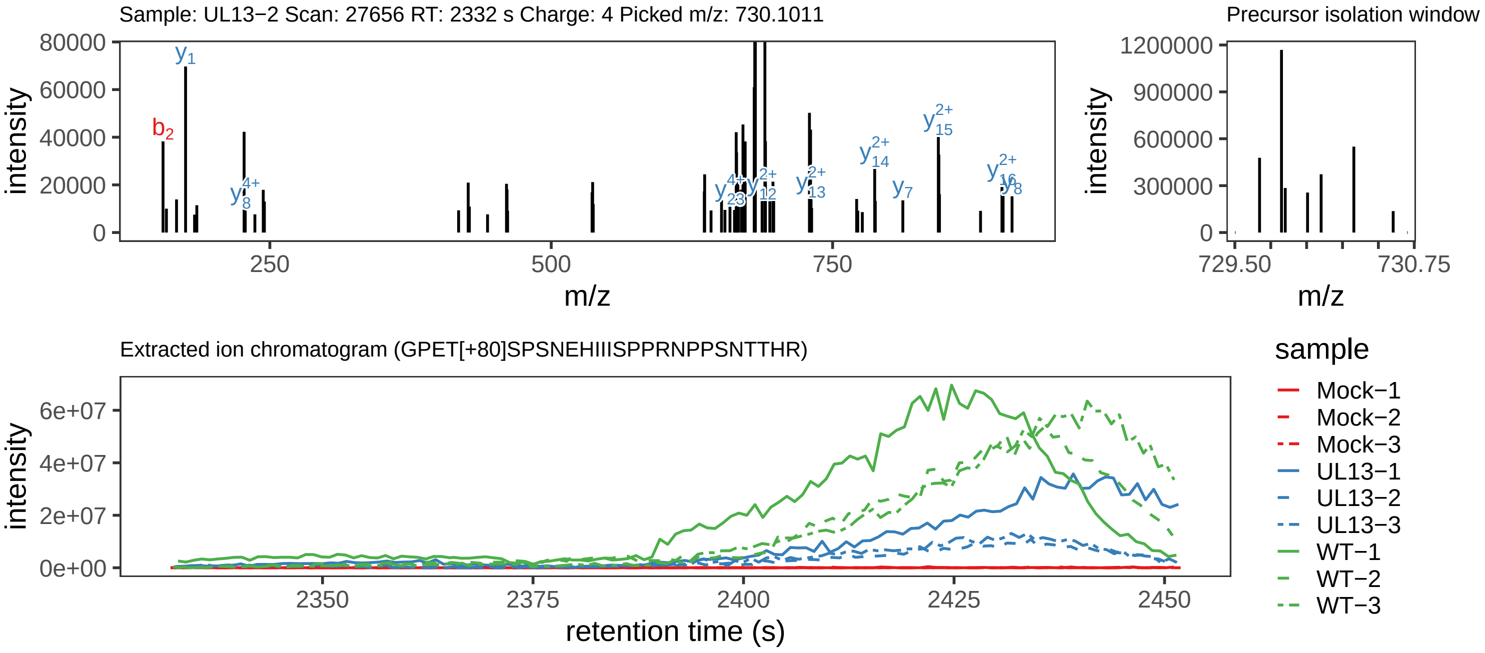

Supplement: Supplemental File 5 [file 737029_file08.zip › phosphopeptide_validation_figs_20240531/GPETSPSNEHIIISPPRNPPSNTTHR/GPET[+80]SPSNEHIIISPPRNPPSNTTHR.phospho.pdf]

#### Peptide SIVSN[+1]NVLEPLC[+57]K

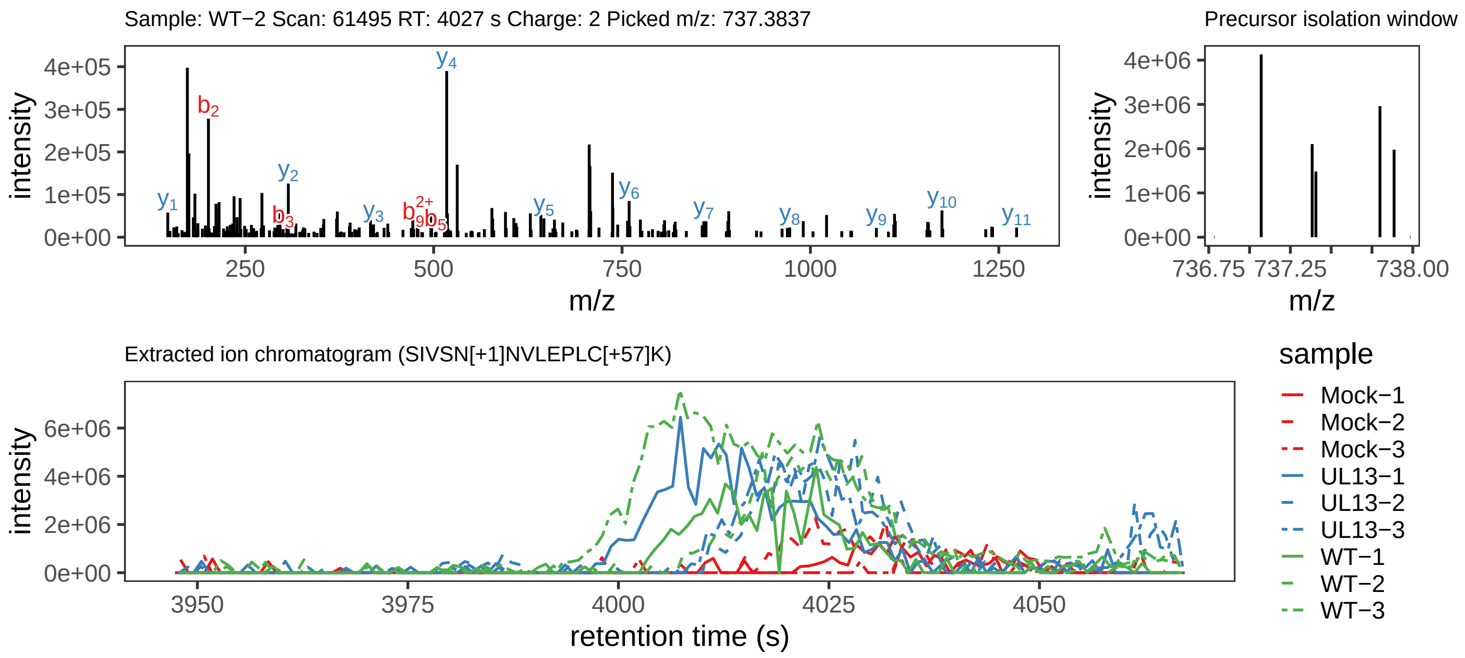

Supplement: Supplemental File 5 [file 737029_file08.zip › phosphopeptide_validation_figs_20240531/SIVSNNVLEPLCK/SIVSN[+1]NVLEPLC[+57]K.global.pdf]

#### Peptide SIVSNNVLEPLC[+57]K

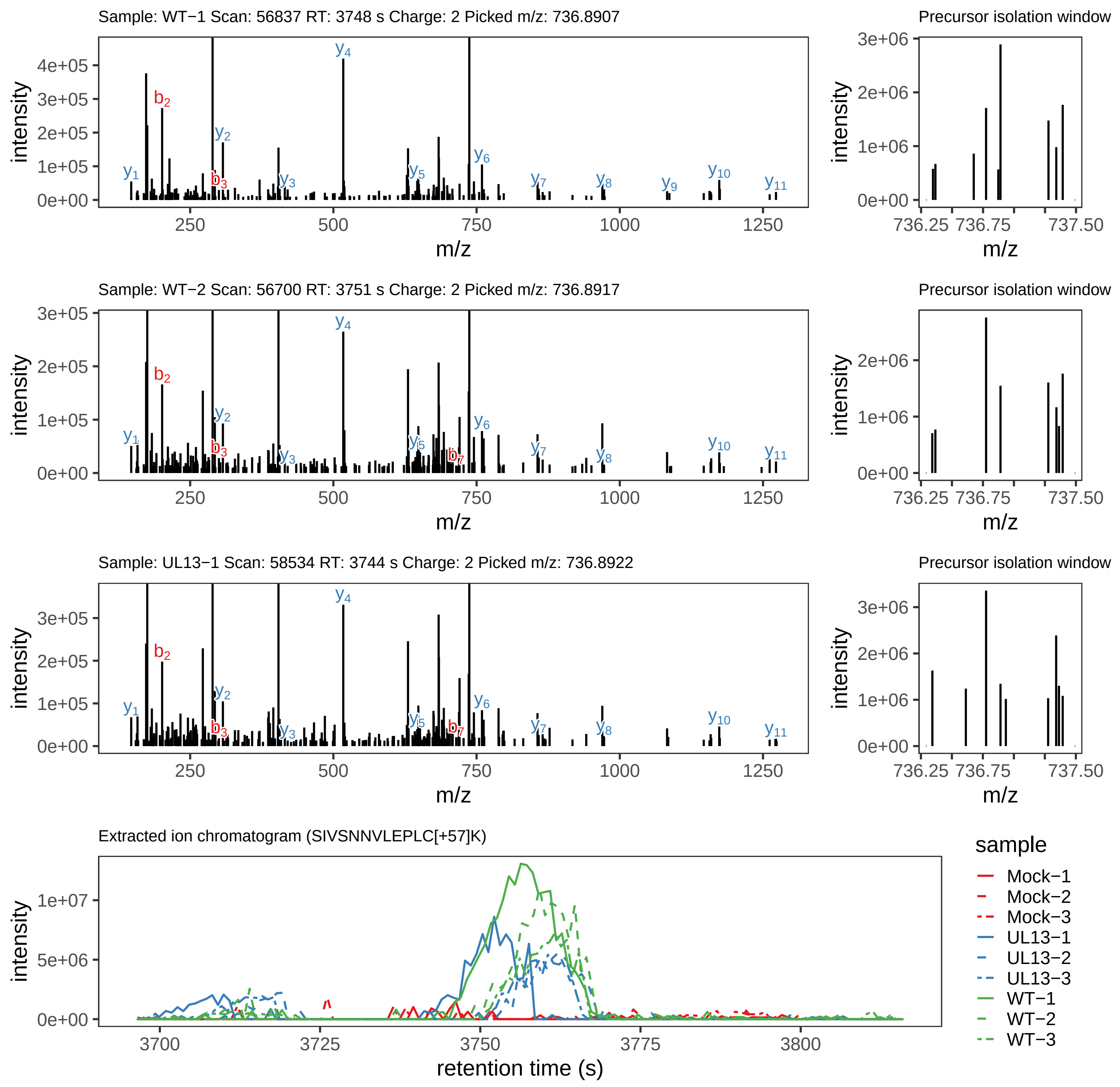

Supplement: Supplemental File 5 [file 737029_file08.zip › phosphopeptide_validation_figs_20240531/SIVSNNVLEPLCK/SIVSNNVLEPLC[+57]K.global.pdf]

#### Peptide S[+80]IVSNNVLEPLC[+57]K

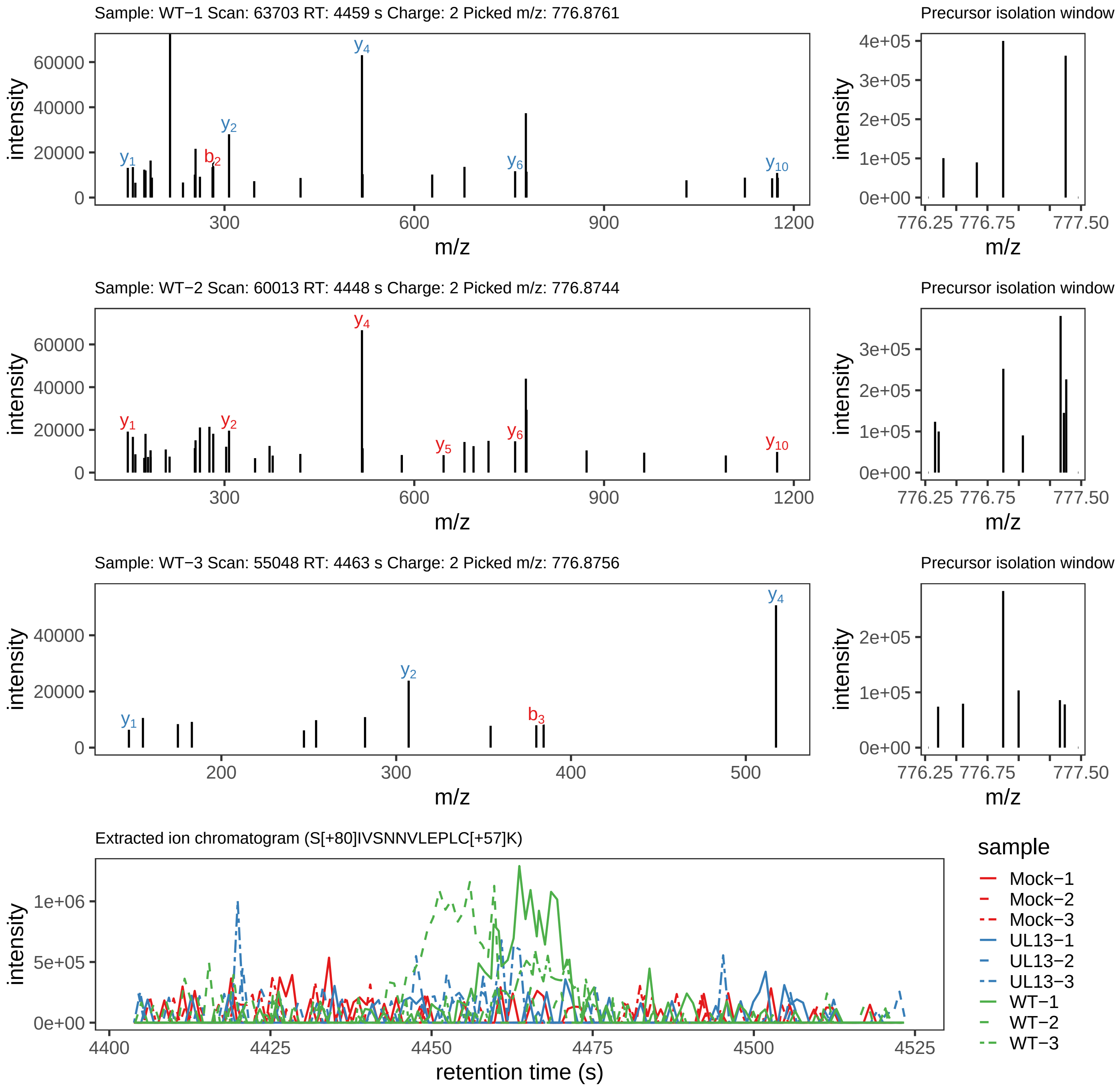

Supplement: Supplemental File 5 [file 737029_file08.zip › phosphopeptide_validation_figs_20240531/SIVSNNVLEPLCK/S[+80]IVSNNVLEPLC[+57]K.phospho.pdf]

#### Peptide SVQ[+1]LRVDS[+80]PK

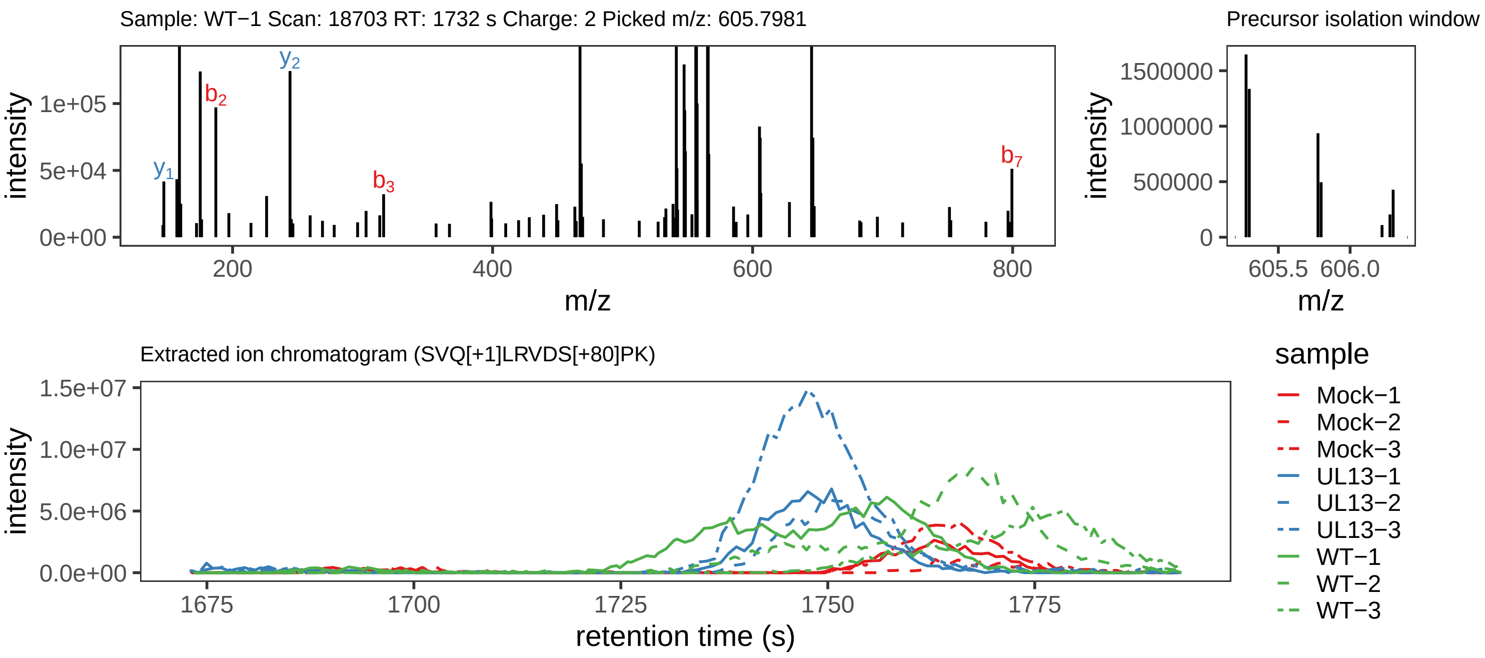

Supplement: Supplemental File 5 [file 737029_file08.zip › phosphopeptide_validation_figs_20240531/SVQLRVDSPK/SVQ[+1]LRVDS[+80]PK.phospho.pdf]

#### Peptide RFS[+80]DNIR

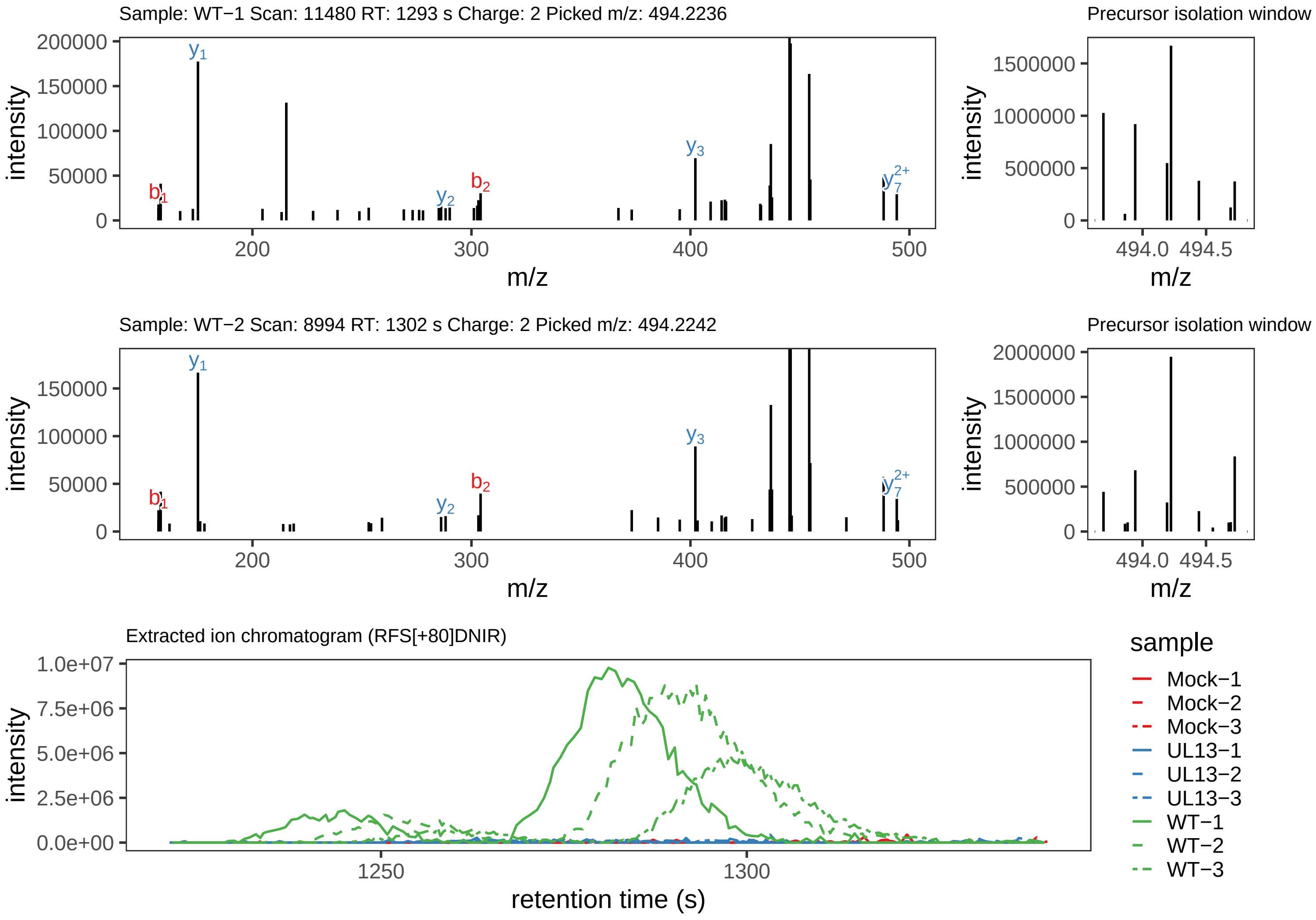

Supplement: Supplemental File 5 [file 737029_file08.zip › phosphopeptide_validation_figs_20240531/RFSDNIR/RFS[+80]DNIR.phospho.pdf]

#### Peptide MTVTLN[+1]EYDIS[+80]ASPFHPTDPTRK

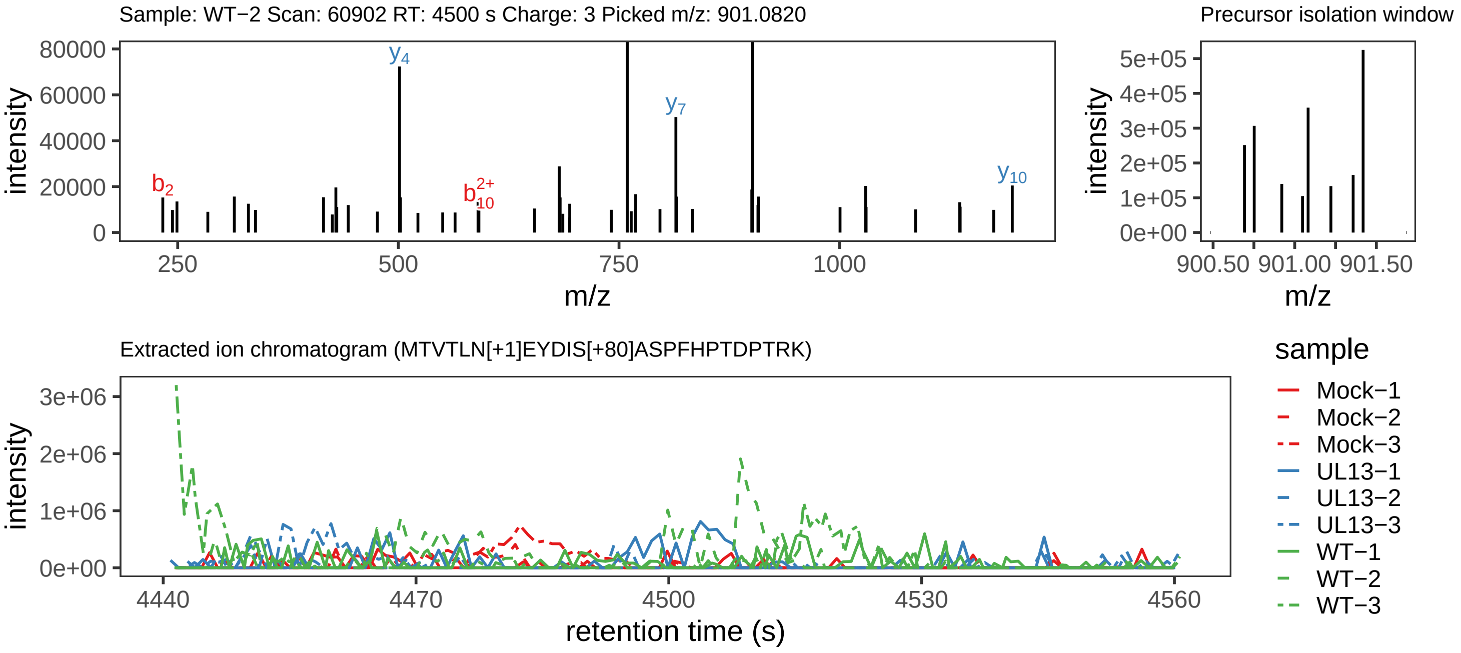

Supplement: Supplemental File 5 [file 737029_file08.zip › phosphopeptide_validation_figs_20240531/MTVTLNEYDISASPFHPTDPTRK/MTVTLN[+1]EYDIS[+80]ASPFHPTDPTRK.phospho.pdf]

#### Peptide M[+16]TVT[+80]LN[+1]EYDISASPFHPTDPTRK

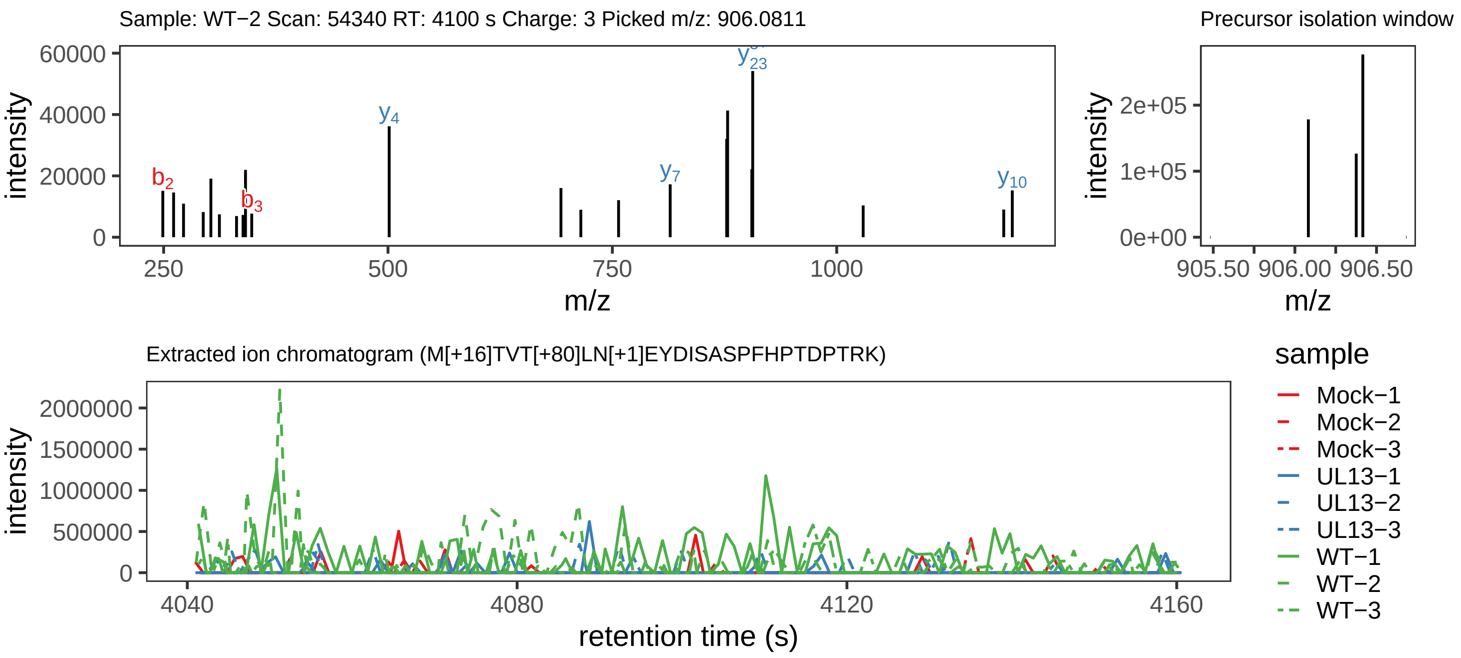

Supplement: Supplemental File 5 [file 737029_file08.zip › phosphopeptide_validation_figs_20240531/MTVTLNEYDISASPFHPTDPTRK/M[+16]TVT[+80]LN[+1]EYDISASPFHPTDPTRK.phospho.pdf]

#### Peptide MT[+80]VTLNEYDISASPFHPTDPTRK

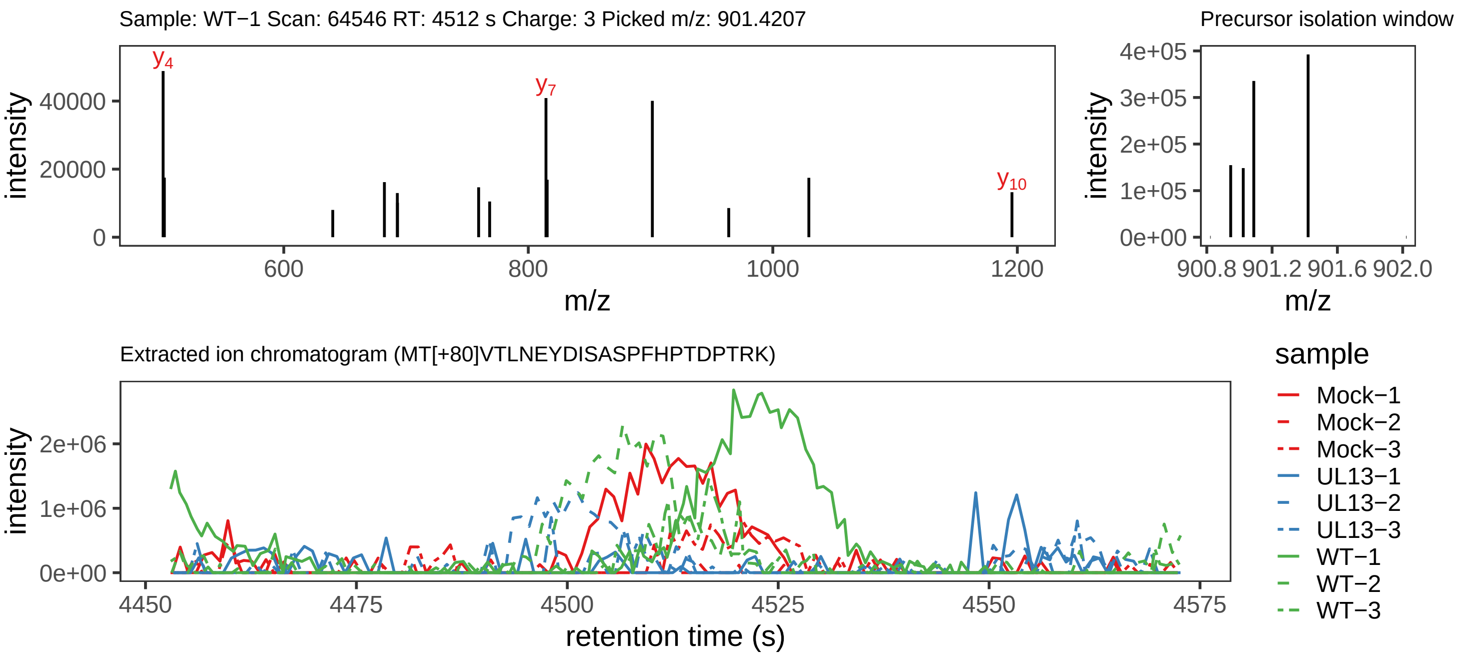

Supplement: Supplemental File 5 [file 737029_file08.zip › phosphopeptide_validation_figs_20240531/MTVTLNEYDISASPFHPTDPTRK/MT[+80]VTLNEYDISASPFHPTDPTRK.phospho.pdf]

#### Peptide M[+16]TVT[+80]LNEYDISASPFHPTDPTRK

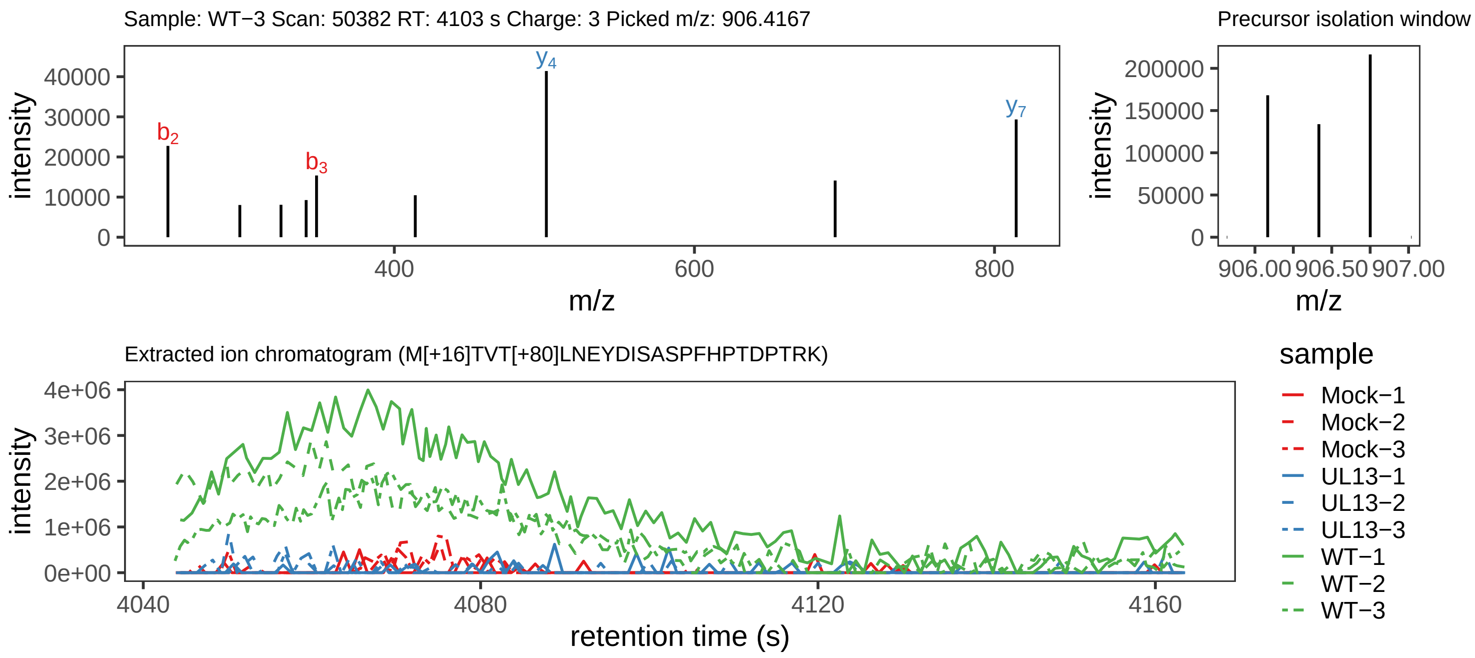

Supplement: Supplemental File 5 [file 737029_file08.zip › phosphopeptide_validation_figs_20240531/MTVTLNEYDISASPFHPTDPTRK/M[+16]TVT[+80]LNEYDISASPFHPTDPTRK.phospho.pdf]

#### Peptide MTVTLNEY[+80]DISASPFHPTDPTRK

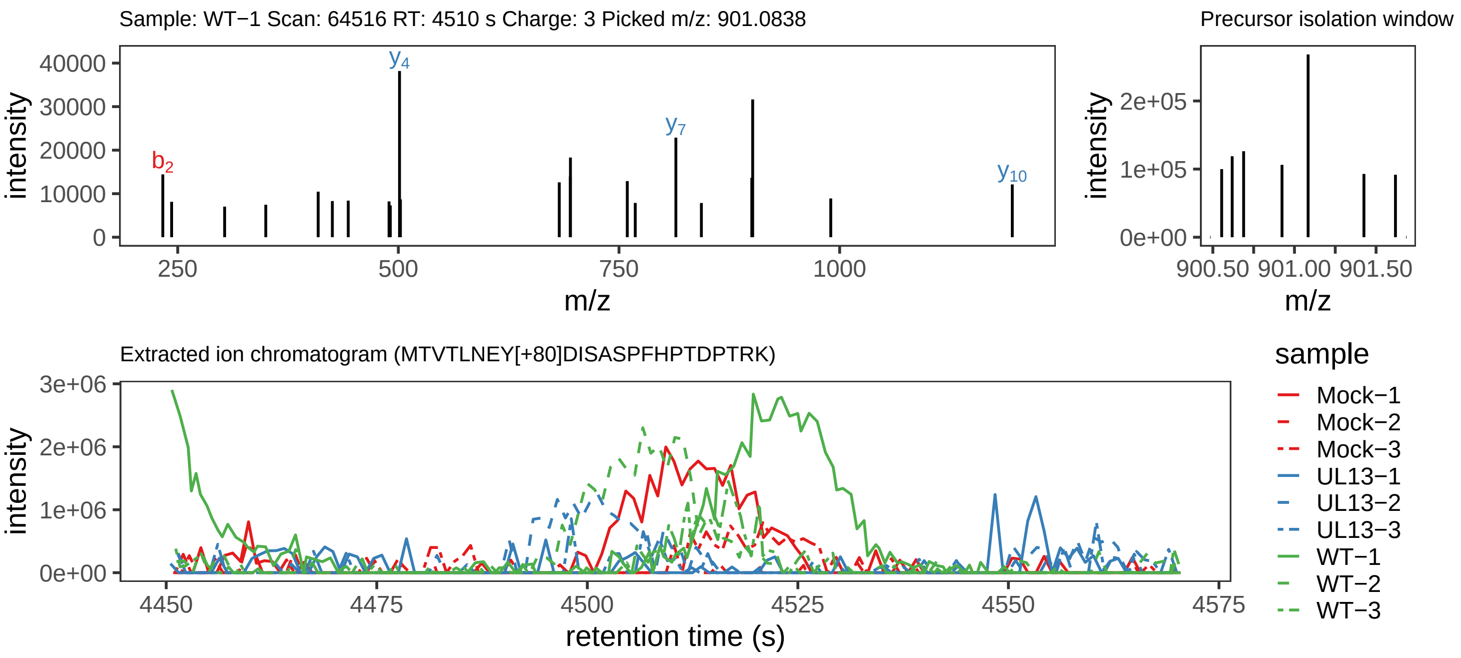

Supplement: Supplemental File 5 [file 737029_file08.zip › phosphopeptide_validation_figs_20240531/MTVTLNEYDISASPFHPTDPTRK/MTVTLNEY[+80]DISASPFHPTDPTRK.phospho.pdf]

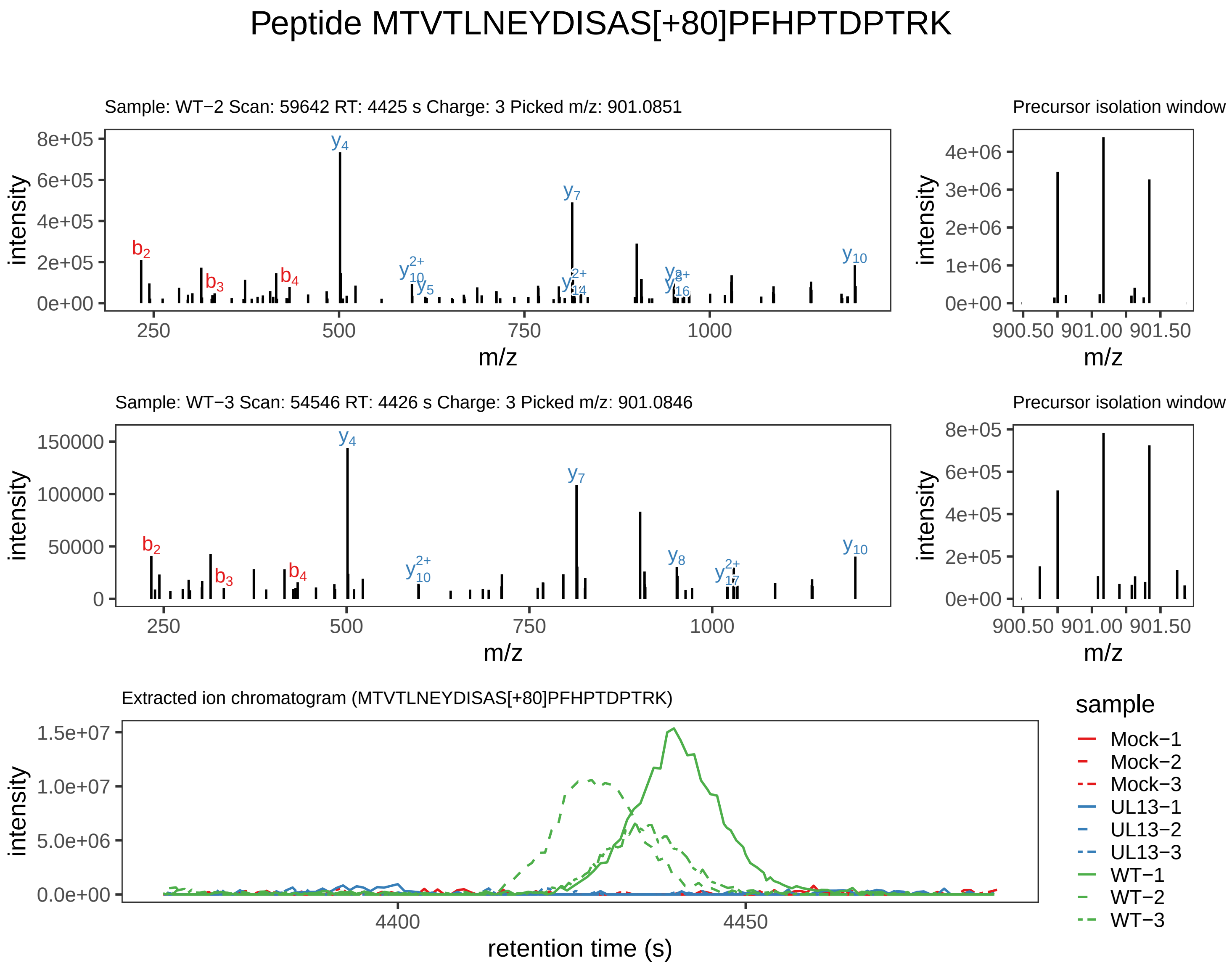

Supplement: Supplemental File 5 [file 737029_file08.zip › phosphopeptide_validation_figs_20240531/MTVTLNEYDISASPFHPTDPTRK/MTVTLNEYDISAS[+80]PFHPTDPTRK.phospho.pdf]

#### Peptide MTVTLN[+1]EY[+80]DISASPFHPTDPTRK

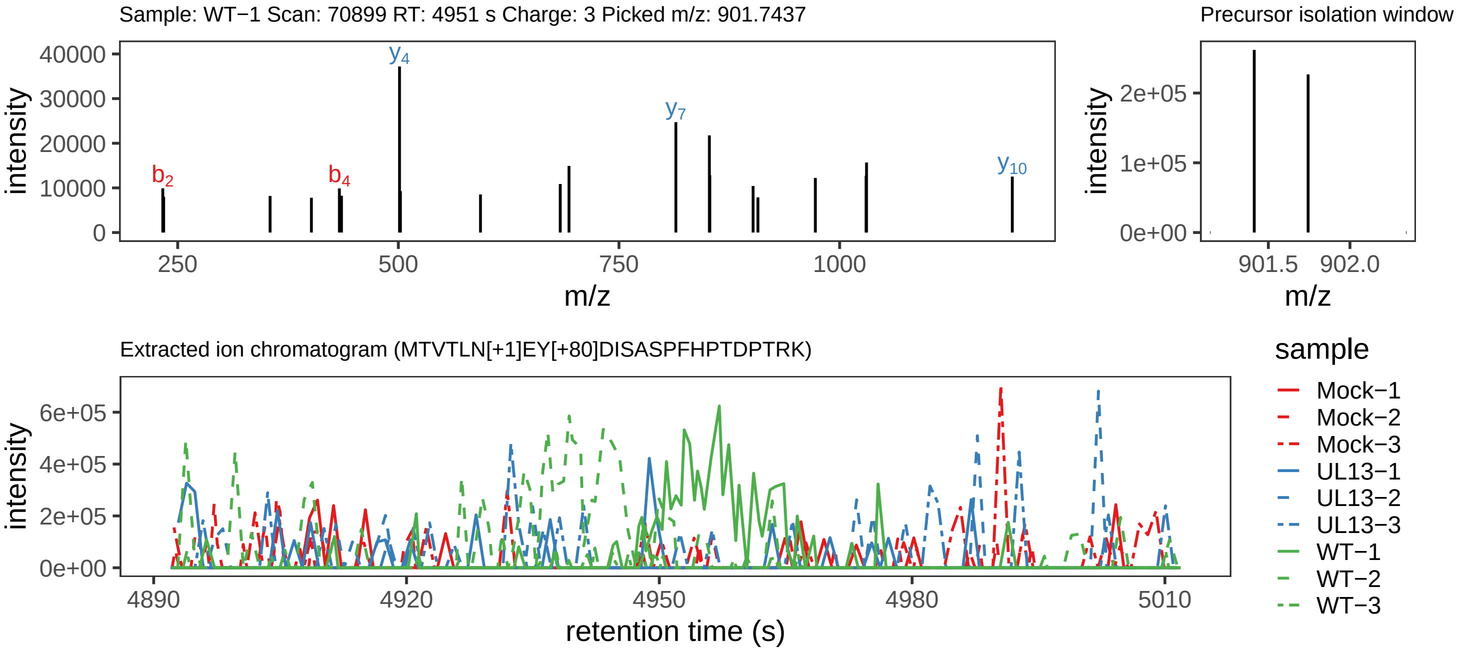

Supplement: Supplemental File 5 [file 737029_file08.zip › phosphopeptide_validation_figs_20240531/MTVTLNEYDISASPFHPTDPTRK/MTVTLN[+1]EY[+80]DISASPFHPTDPTRK.phospho.pdf]

#### Peptide M[+16]TVTLNEYDISAS[+80]PFHPTDPTRK

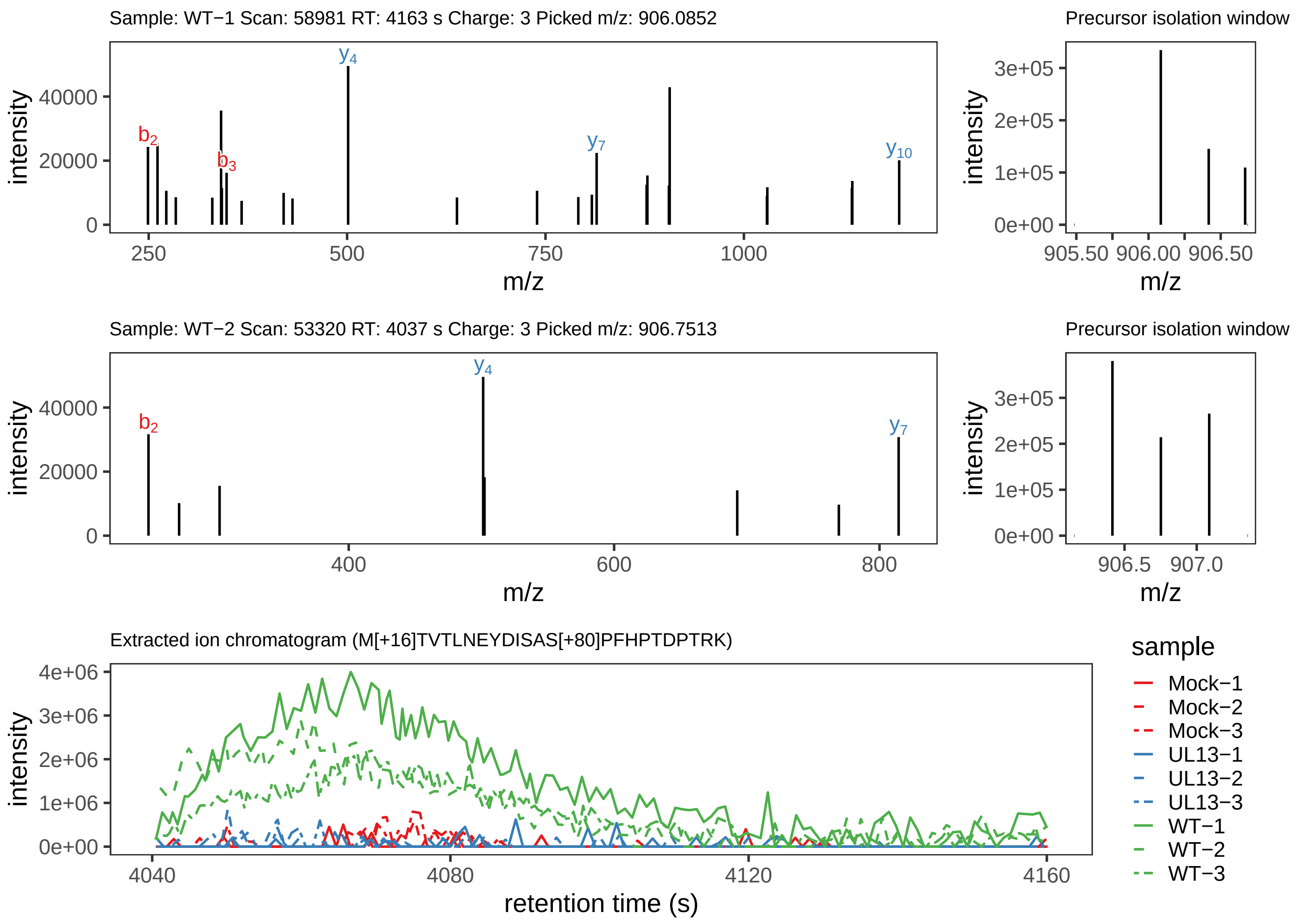

Supplement: Supplemental File 5 [file 737029_file08.zip › phosphopeptide_validation_figs_20240531/MTVTLNEYDISASPFHPTDPTRK/M[+16]TVTLNEYDISAS[+80]PFHPTDPTRK.phospho.pdf]

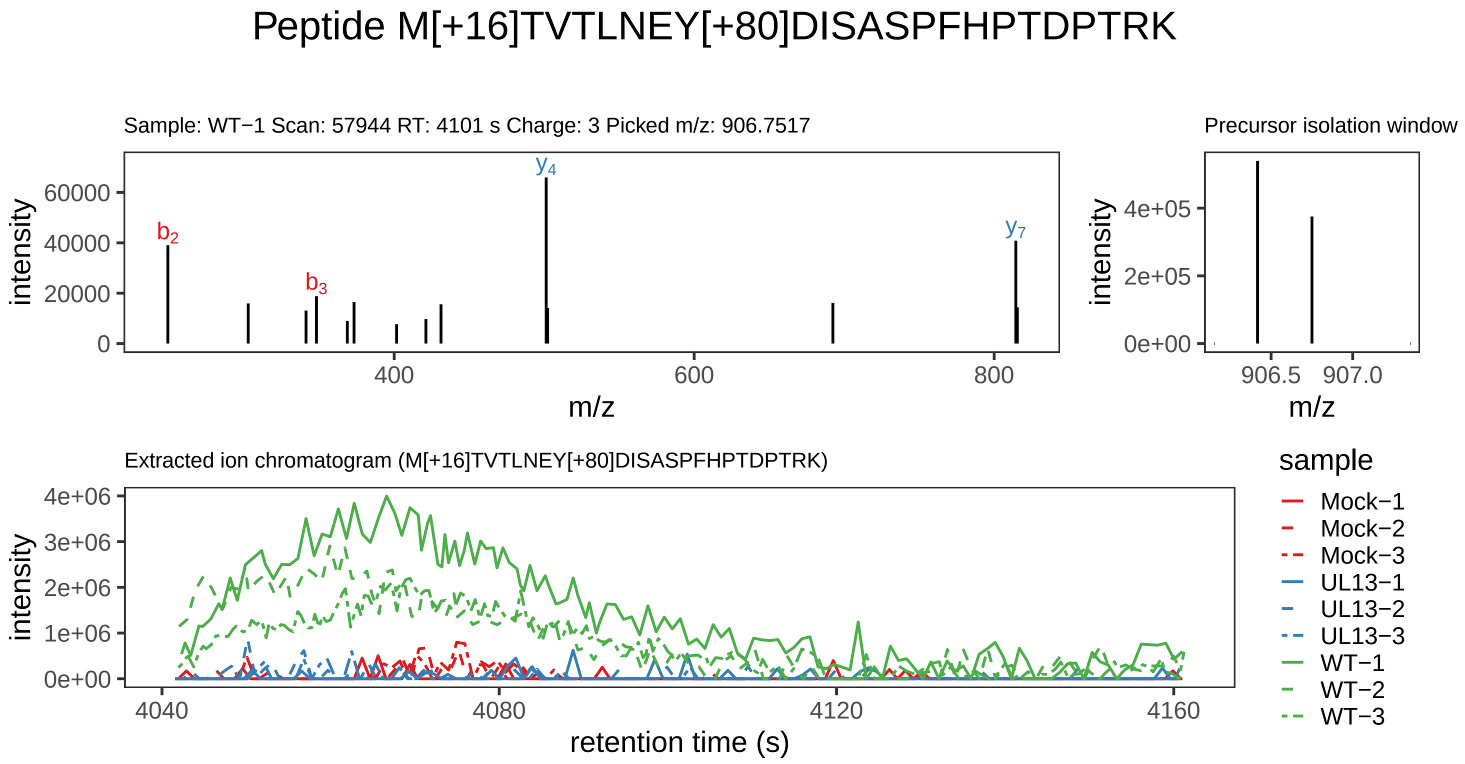

Supplement: Supplemental File 5 [file 737029_file08.zip › phosphopeptide_validation_figs_20240531/MTVTLNEYDISASPFHPTDPTRK/M[+16]TVTLNEY[+80]DISASPFHPTDPTRK.phospho.pdf]

#### Peptide MTVTLNEYDISASPFHPTDPTRK

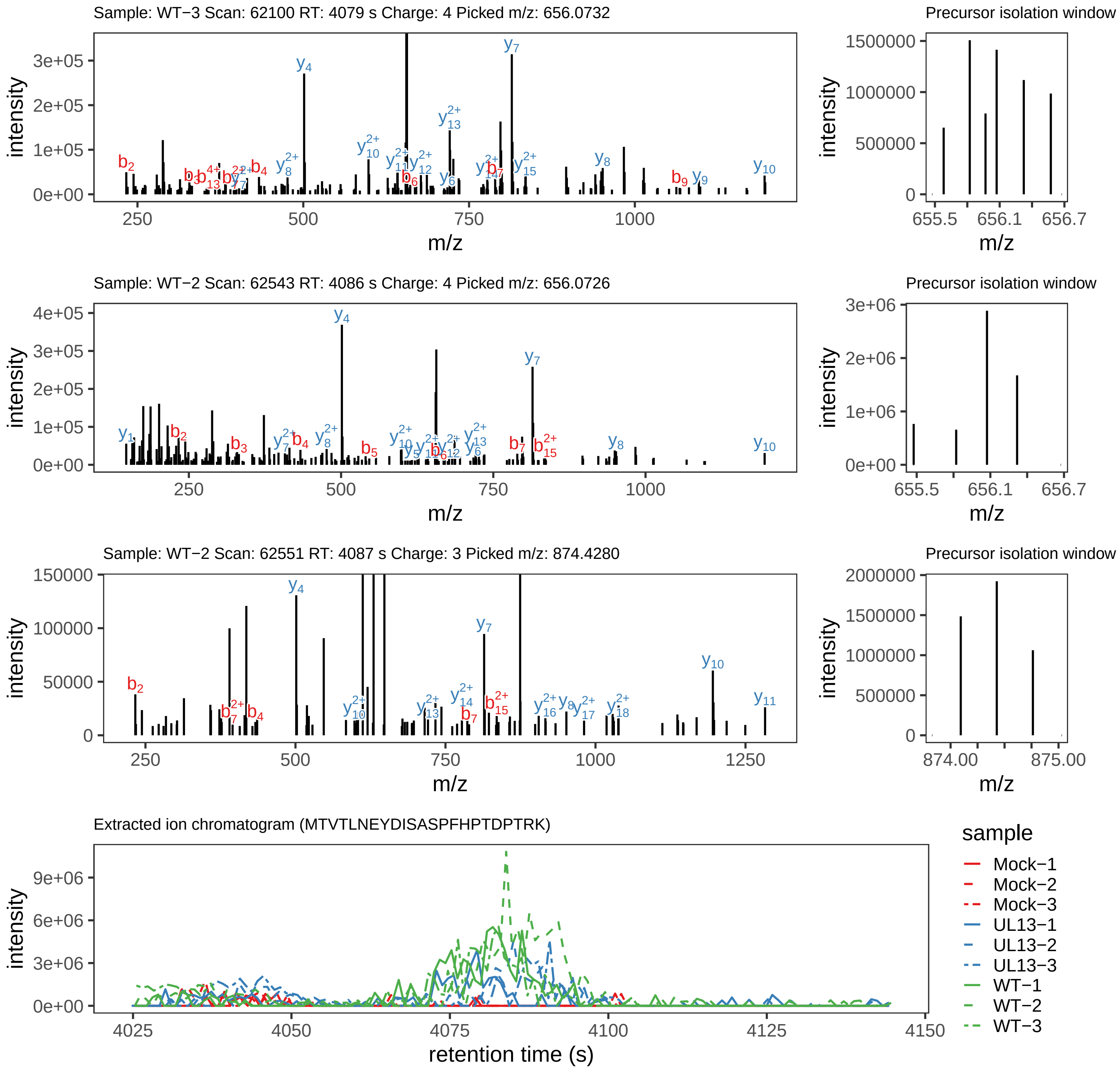
