## Supplementary figures and images for "Conserved herpesvirus protein kinase (CHPK)-mediated phosphorylation of viral proteins associated with nucleocytoplasmic trafficking during natural infection"

### Peptide SMVIHGSLLK

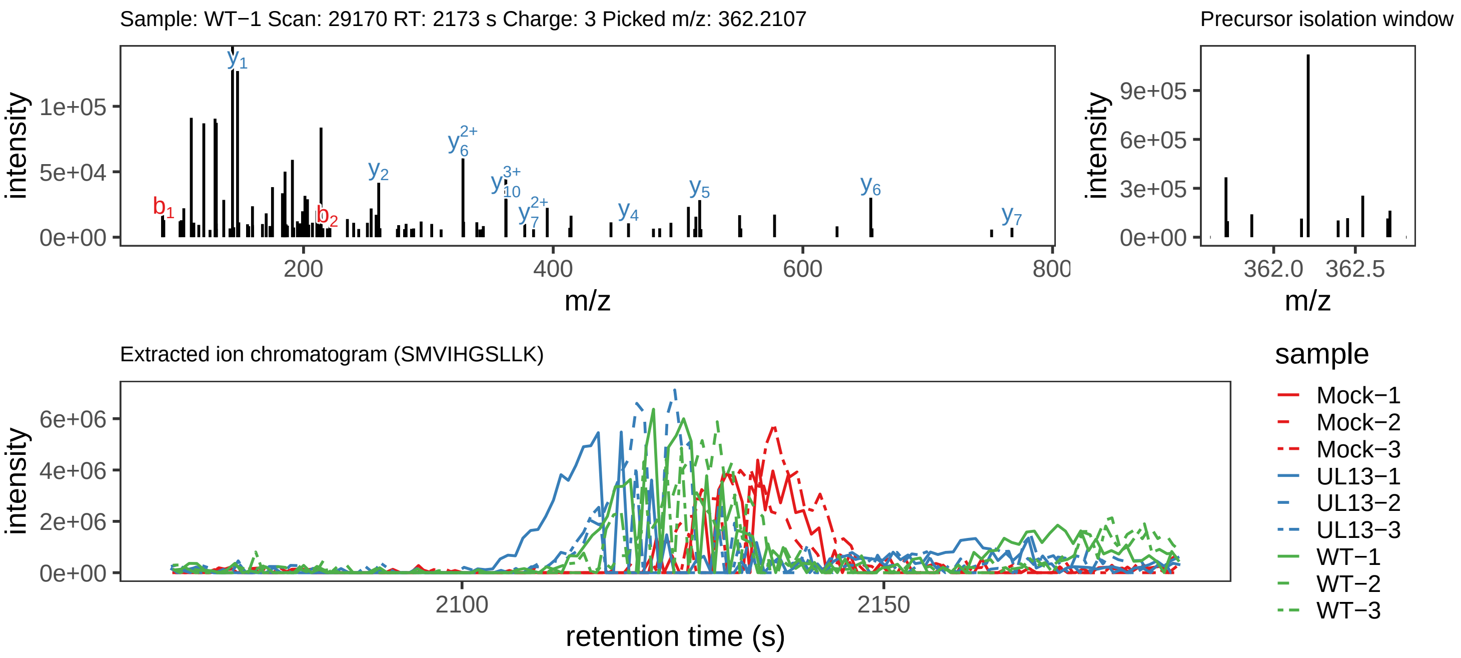

Supplement: Supplemental File 5 [file 737029_file08.zip › phosphopeptide_validation_figs_20240531/SMVIHGSLLK/SMVIHGSLLK.global.pdf]

### Peptide DIT[+80]SLLLR

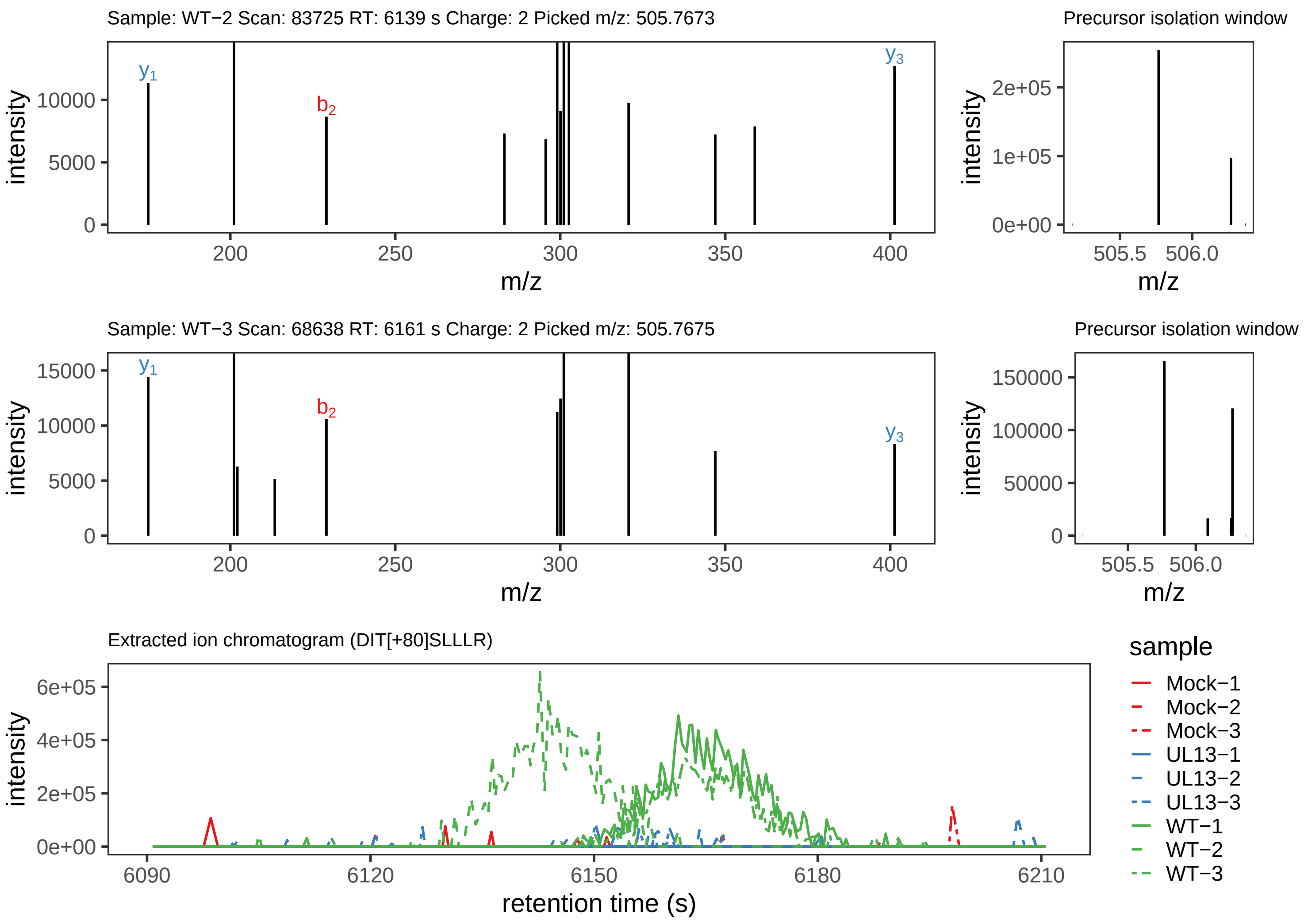

Supplement: Supplemental File 5 [file 737029_file08.zip › phosphopeptide_validation_figs_20240531/DITSLLLR/DIT[+80]SLLLR.phospho.pdf]

### Peptide DITSLLLR

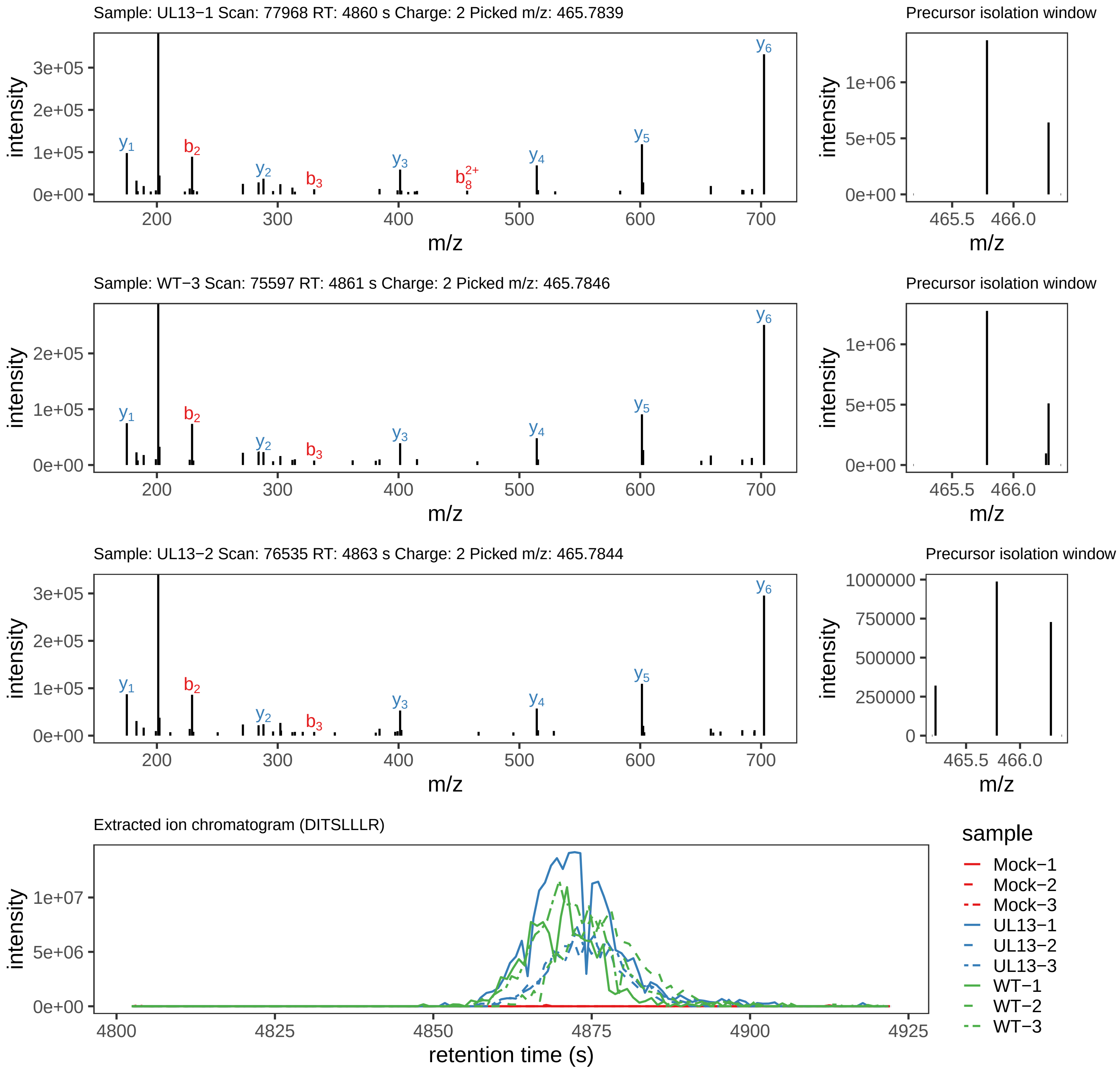

Supplement: Supplemental File 5 [file 737029_file08.zip › phosphopeptide_validation_figs_20240531/DITSLLLR/DITSLLLR.global.pdf]

### Peptide VTST[+80]GELLRPNIQLMEELR

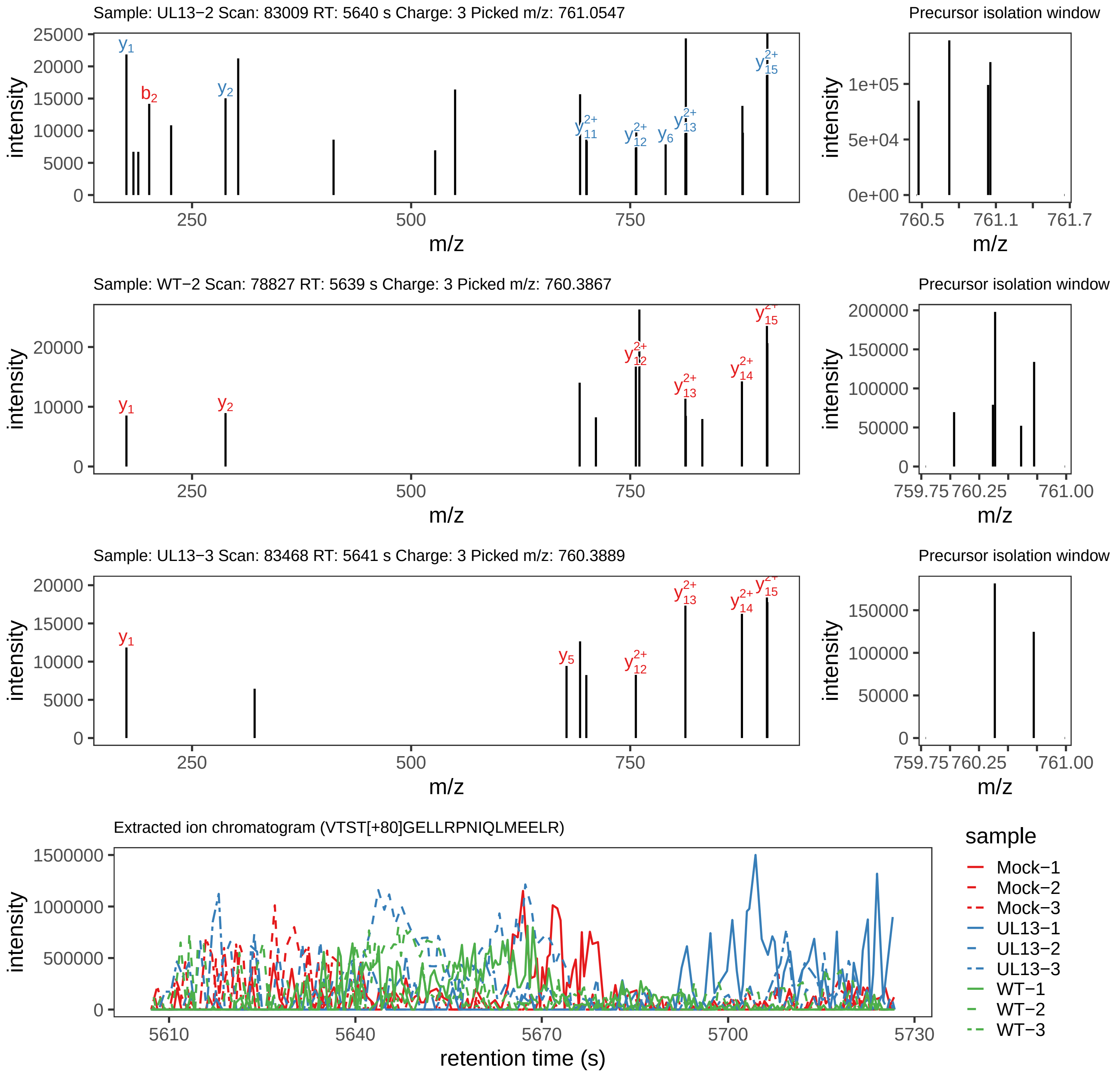

Supplement: Supplemental File 5 [file 737029_file08.zip › phosphopeptide_validation_figs_20240531/VTSTGELLRPNIQLMEELR/VTST[+80]GELLRPNIQLMEELR.phospho.pdf]

### Peptide VTSTGELLRPNIQLMEELR

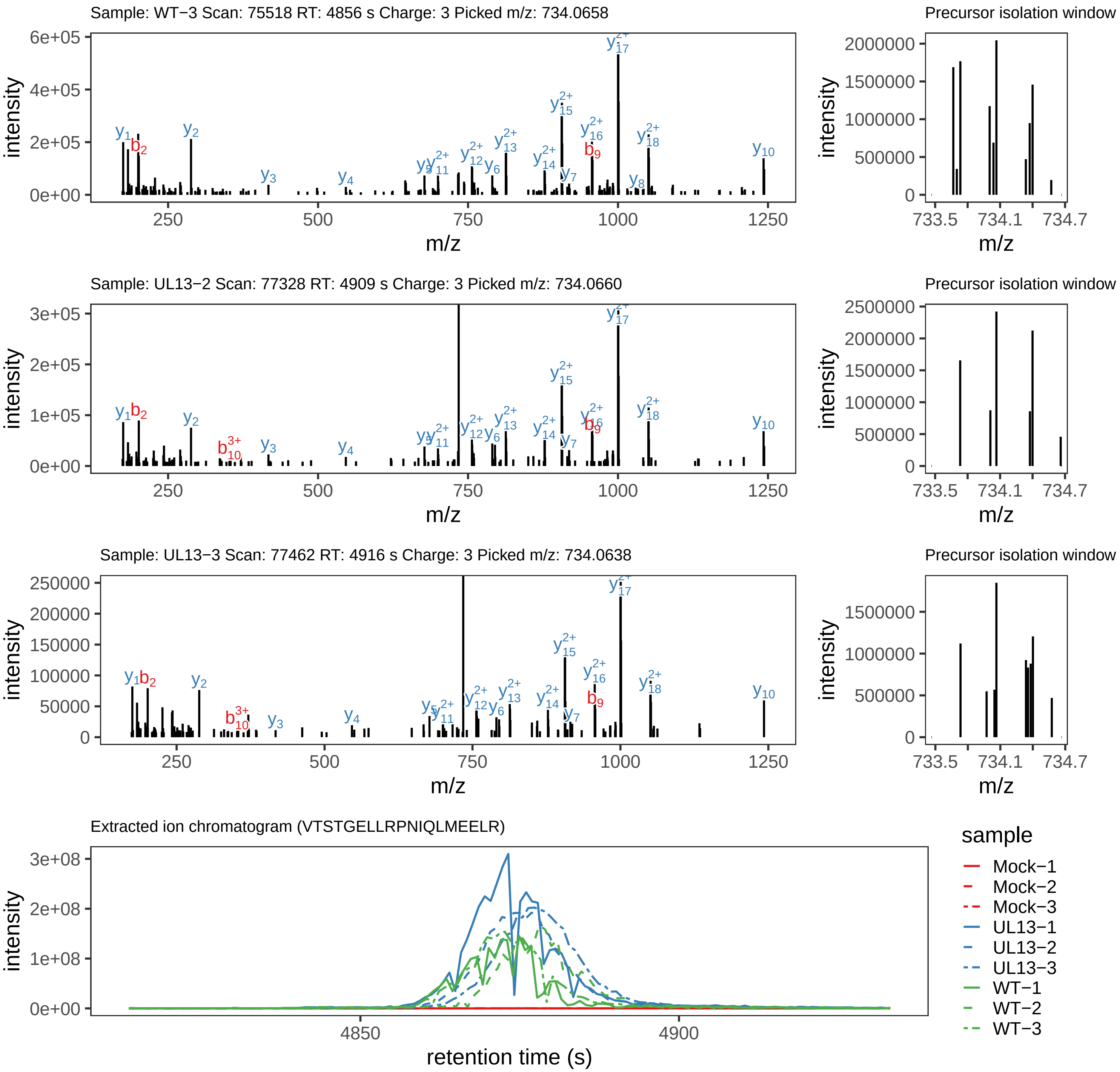

Supplement: Supplemental File 5 [file 737029_file08.zip › phosphopeptide_validation_figs_20240531/VTSTGELLRPNIQLMEELR/VTSTGELLRPNIQLMEELR.global.pdf]

### Peptide VTSTGELLRPNIQLMEELR

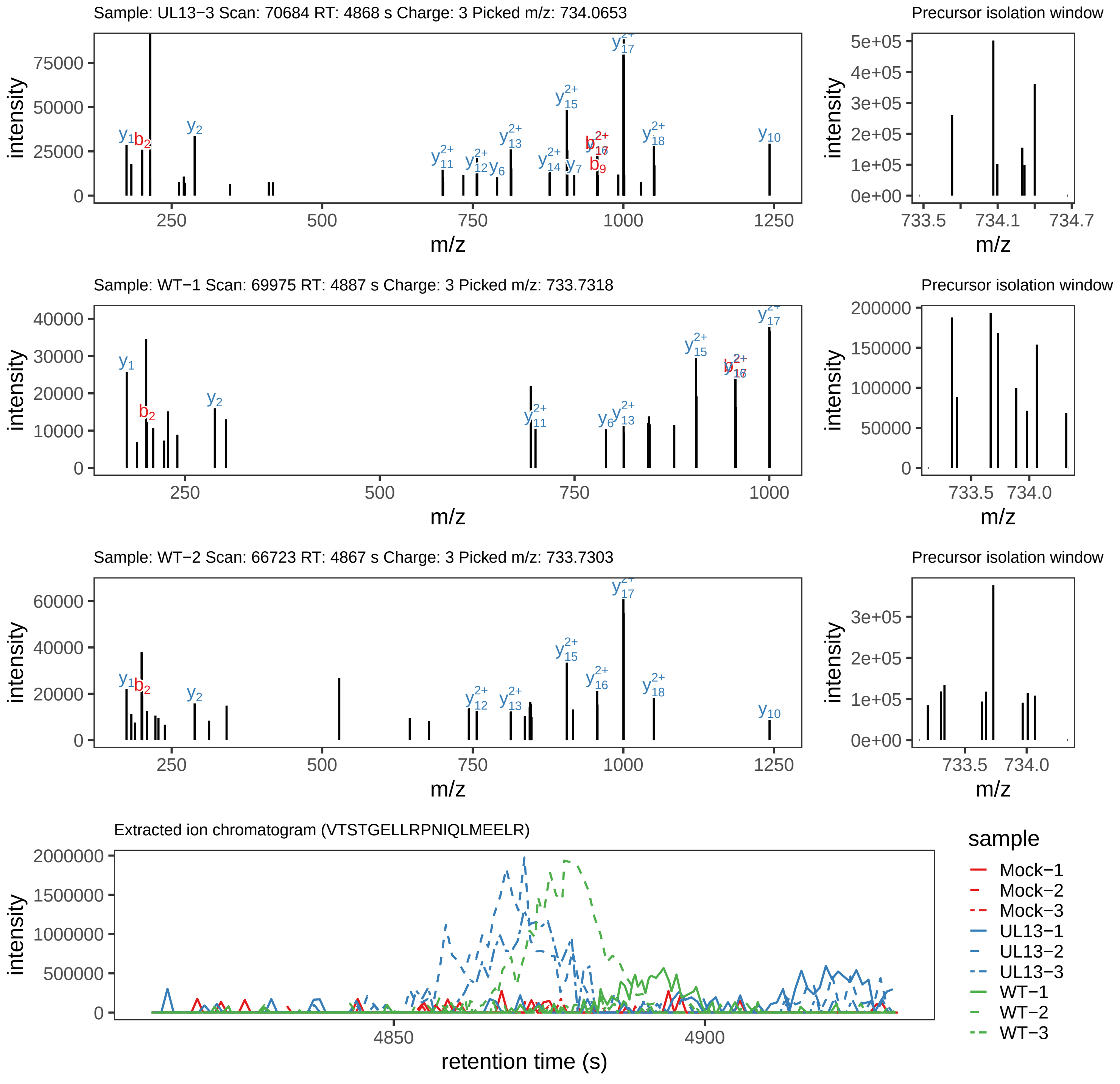

Supplement: Supplemental File 5 [file 737029_file08.zip › phosphopeptide_validation_figs_20240531/VTSTGELLRPNIQLMEELR/VTSTGELLRPNIQLMEELR.phospho.pdf]

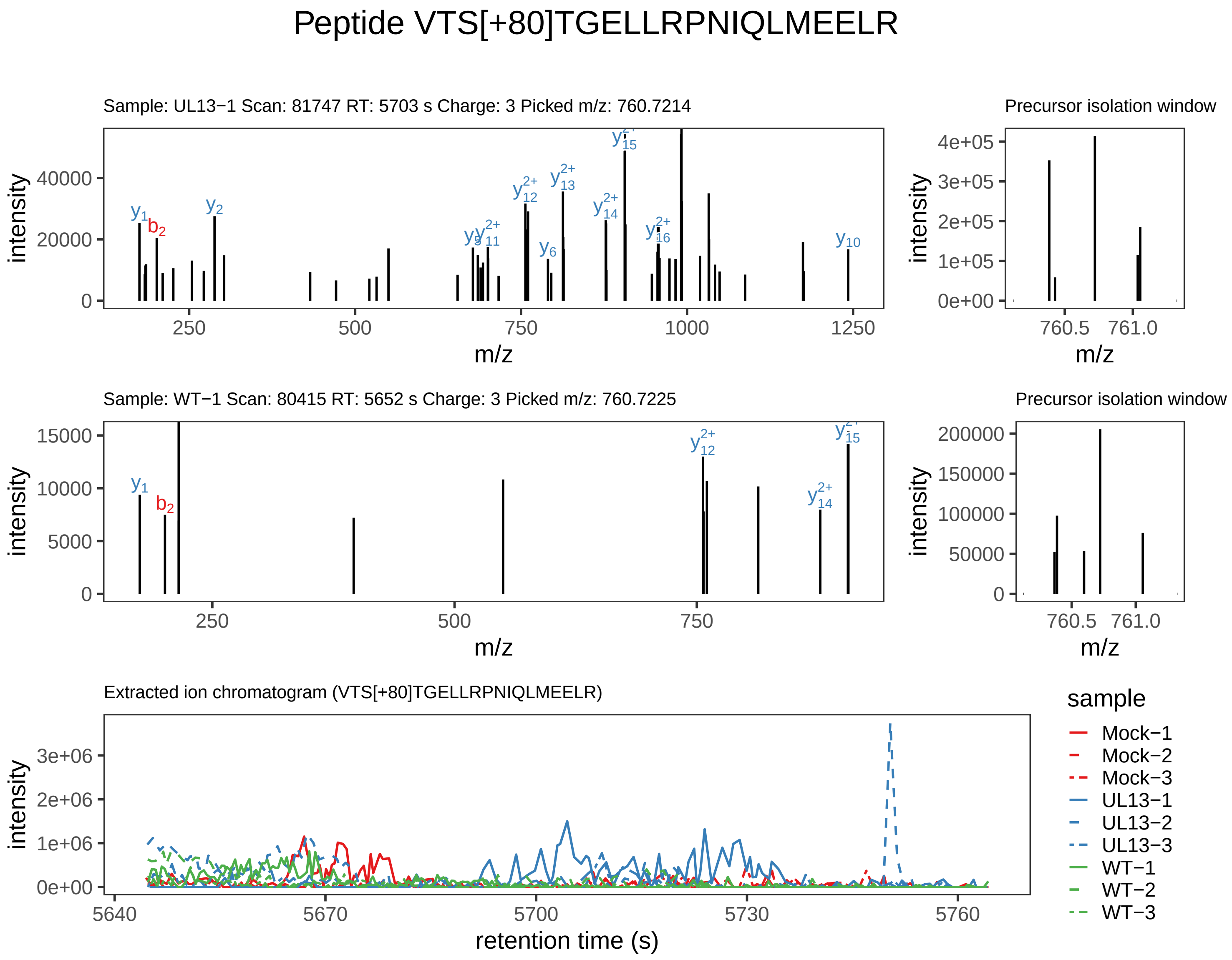

Supplement: Supplemental File 5 [file 737029_file08.zip › phosphopeptide_validation_figs_20240531/VTSTGELLRPNIQLMEELR/VTS[+80]TGELLRPNIQLMEELR.phospho.pdf]

### Peptide TPDDQTDITDDS[+80]ADWSEGETR

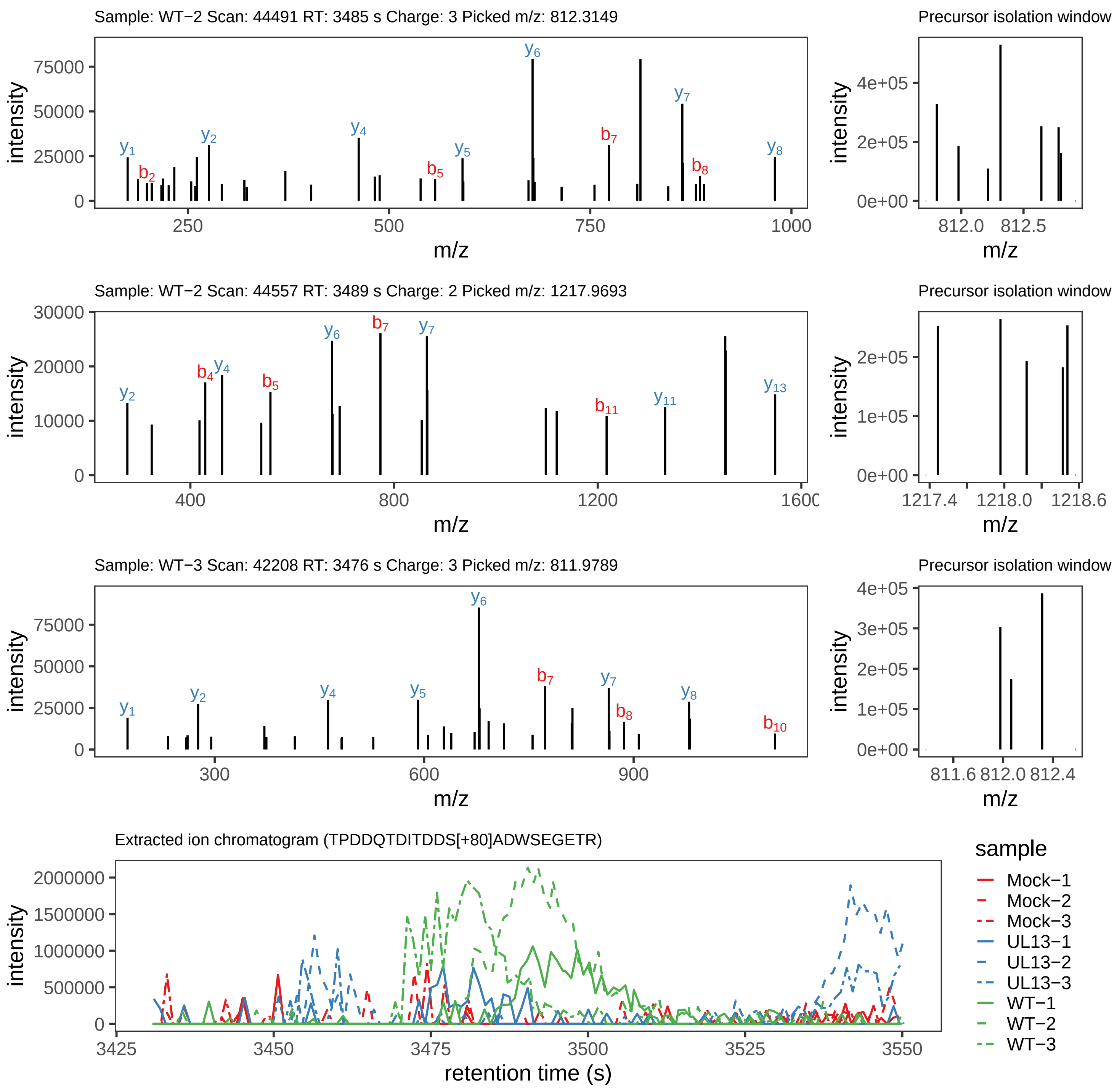

Supplement: Supplemental File 5 [file 737029_file08.zip › phosphopeptide_validation_figs_20240531/TPDDQTDITDDSADWSEGETR/TPDDQTDITDDS[+80]ADWSEGETR.phospho.pdf]

### Peptide TPDDQTDITDDSADWS[+80]EGETR

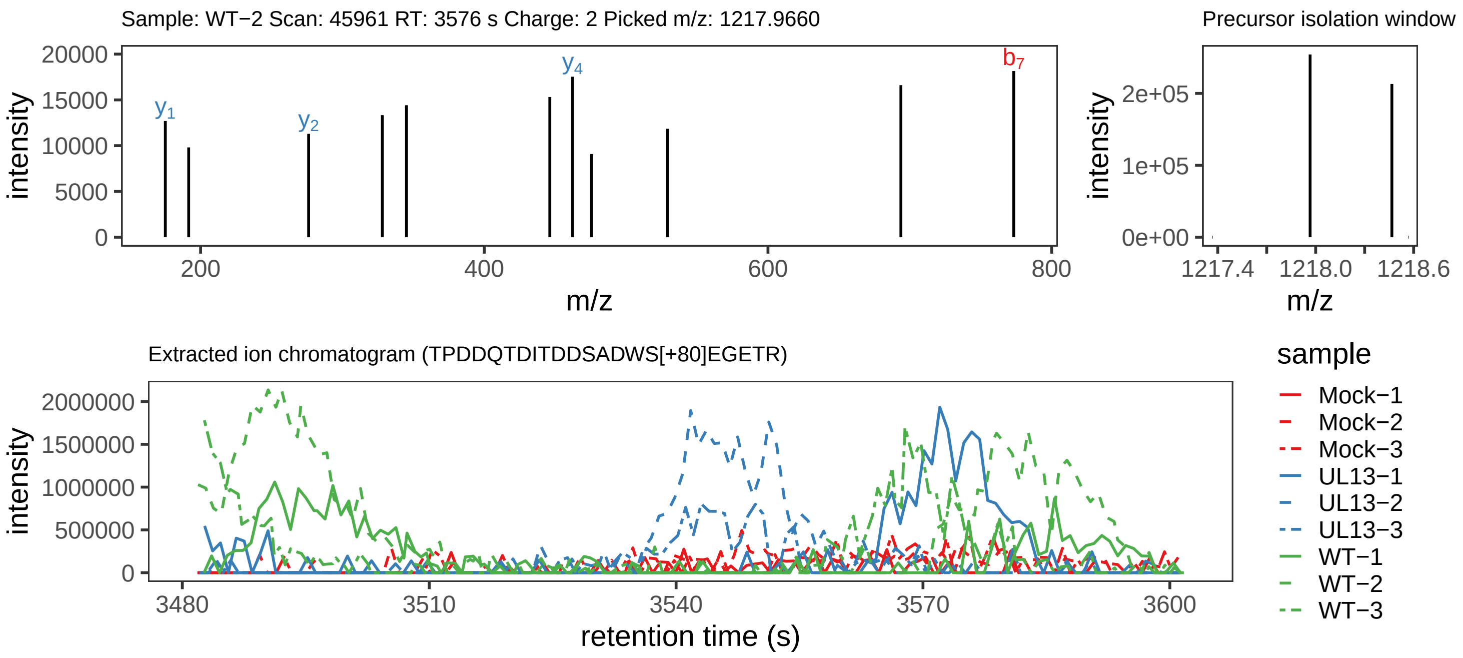

Supplement: Supplemental File 5 [file 737029_file08.zip › phosphopeptide_validation_figs_20240531/TPDDQTDITDDSADWSEGETR/TPDDQTDITDDSADWS[+80]EGETR.phospho.pdf]

### Peptide TPDDQTDIT[+80]DDSADWSEGETR

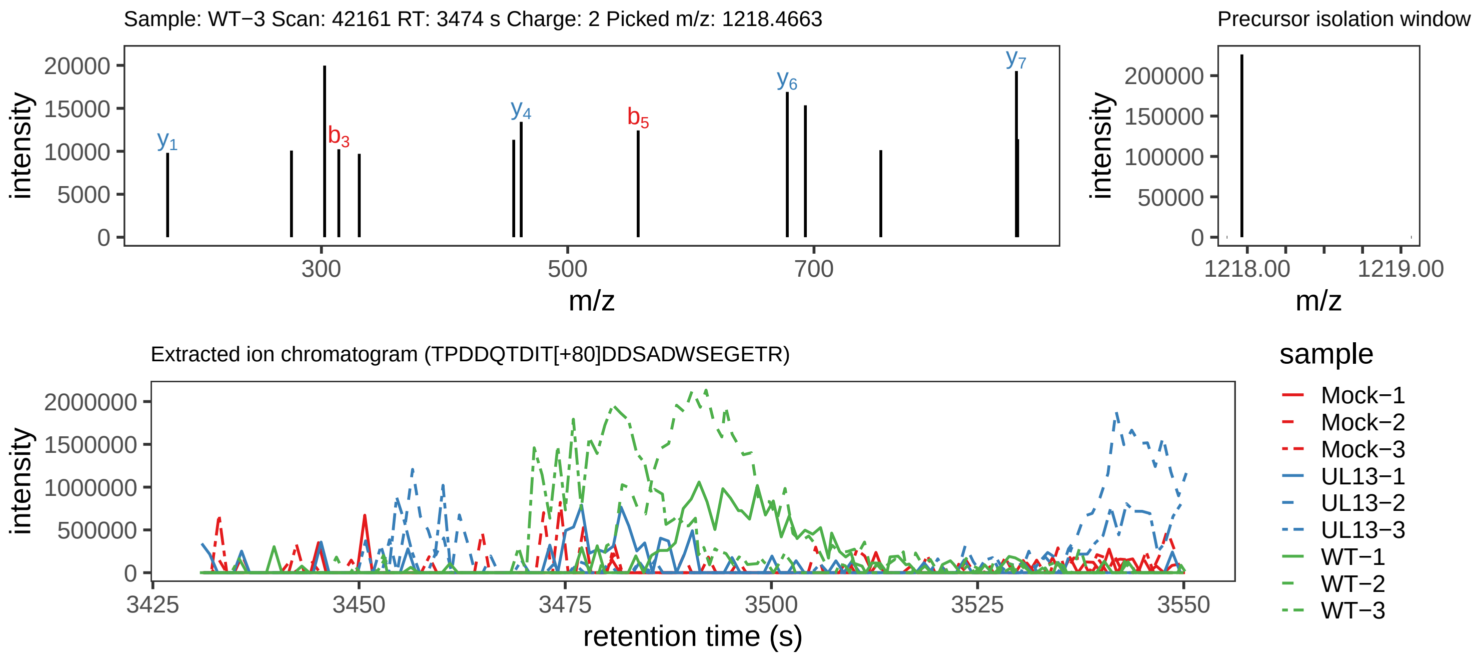

Supplement: Supplemental File 5 [file 737029_file08.zip › phosphopeptide_validation_figs_20240531/TPDDQTDITDDSADWSEGETR/TPDDQTDIT[+80]DDSADWSEGETR.phospho.pdf]

### Peptide LPS[+80]GLTLGIR

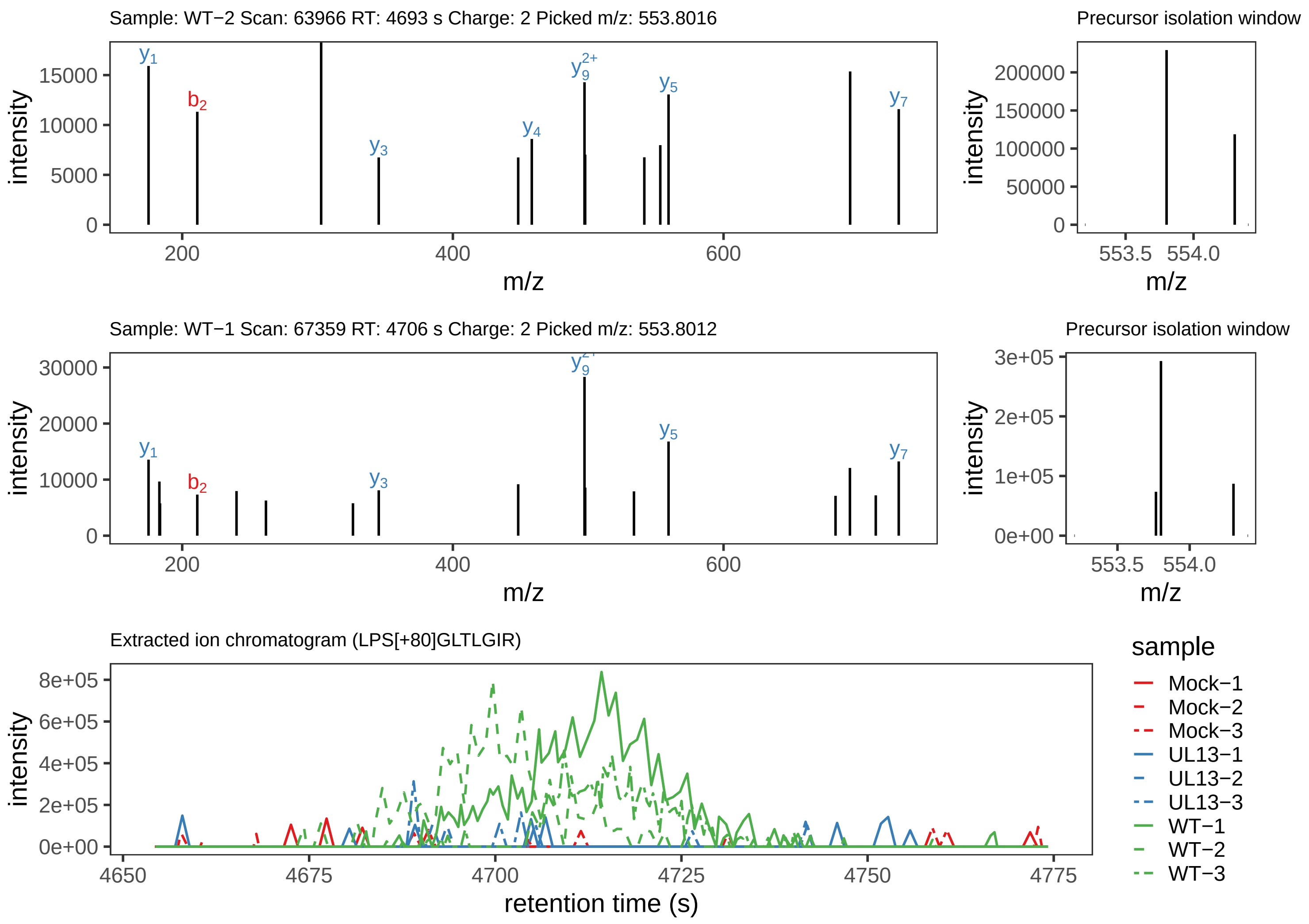

Supplement: Supplemental File 5 [file 737029_file08.zip › phosphopeptide_validation_figs_20240531/LPSGLTLGIR/LPS[+80]GLTLGIR.phospho.pdf]

### Peptide LPSGLTLGIR

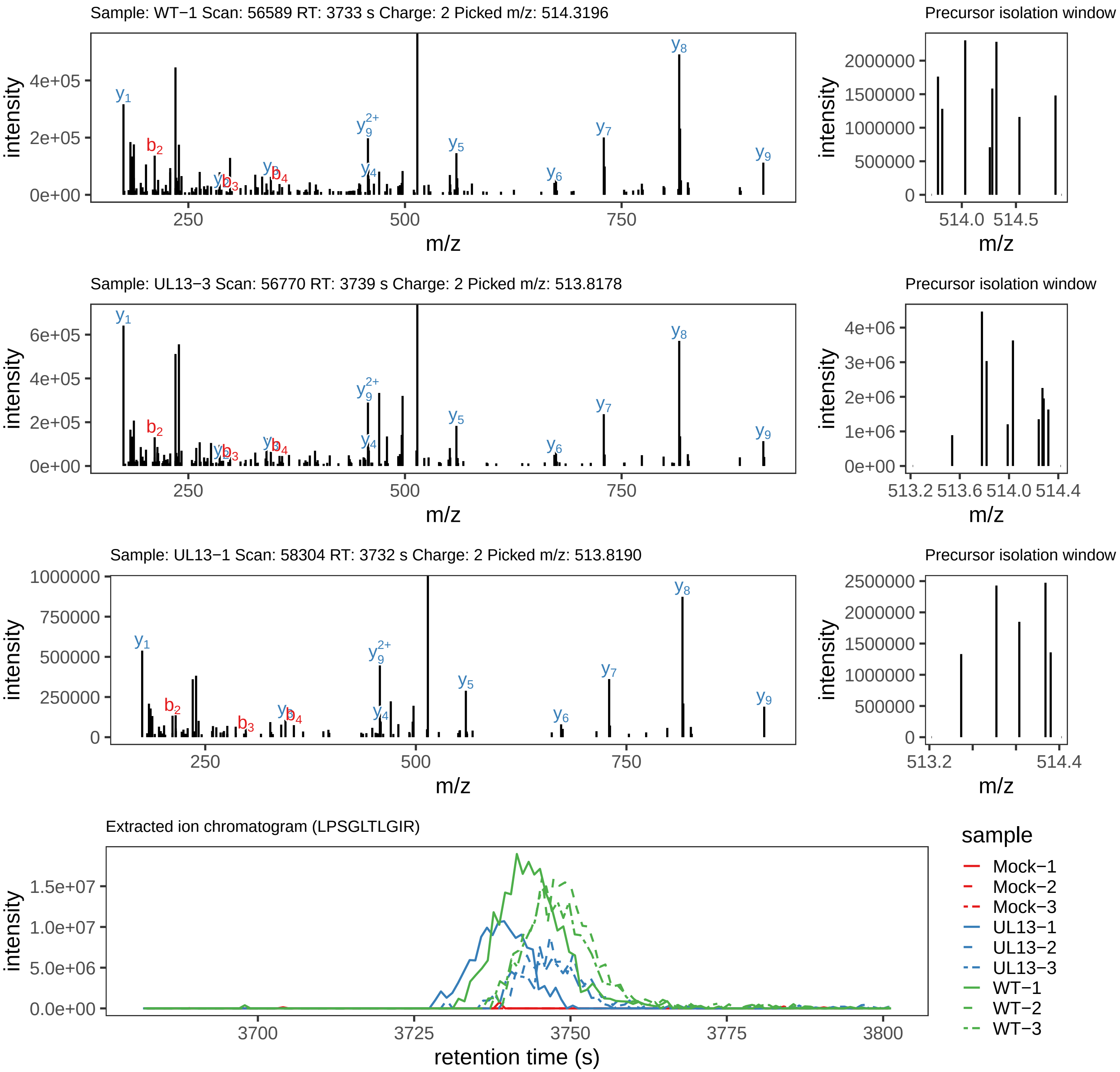

Supplement: Supplemental File 5 [file 737029_file08.zip › phosphopeptide_validation_figs_20240531/LPSGLTLGIR/LPSGLTLGIR.global.pdf]

### Peptide LANLTVDSAC[+57]ISQTK

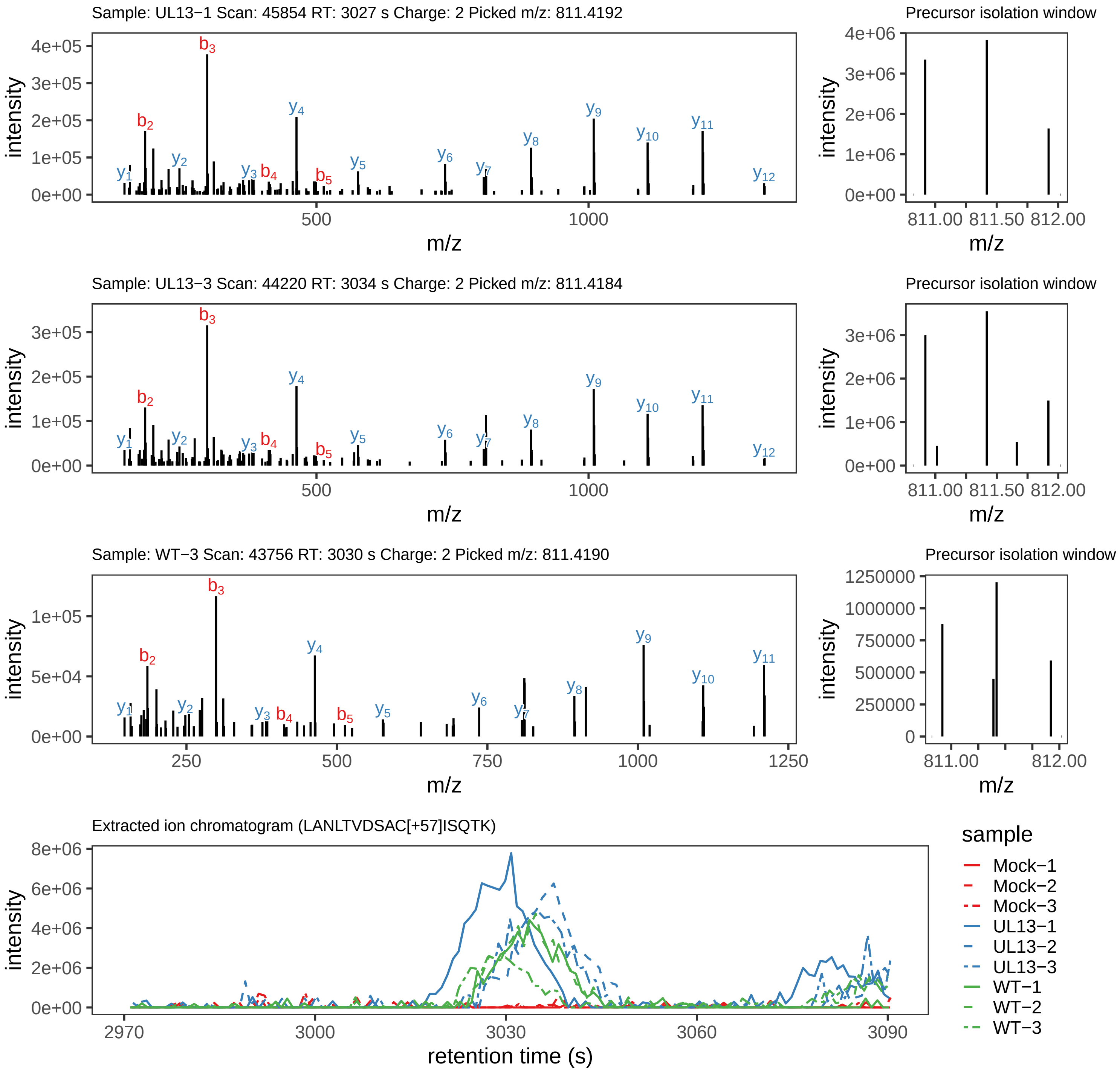

Supplement: Supplemental File 5 [file 737029_file08.zip › phosphopeptide_validation_figs_20240531/LANLTVDSACISQTK/LANLTVDSAC[+57]ISQTK.global.pdf]

### Peptide LANLTVDSAC[+57]ISQTK

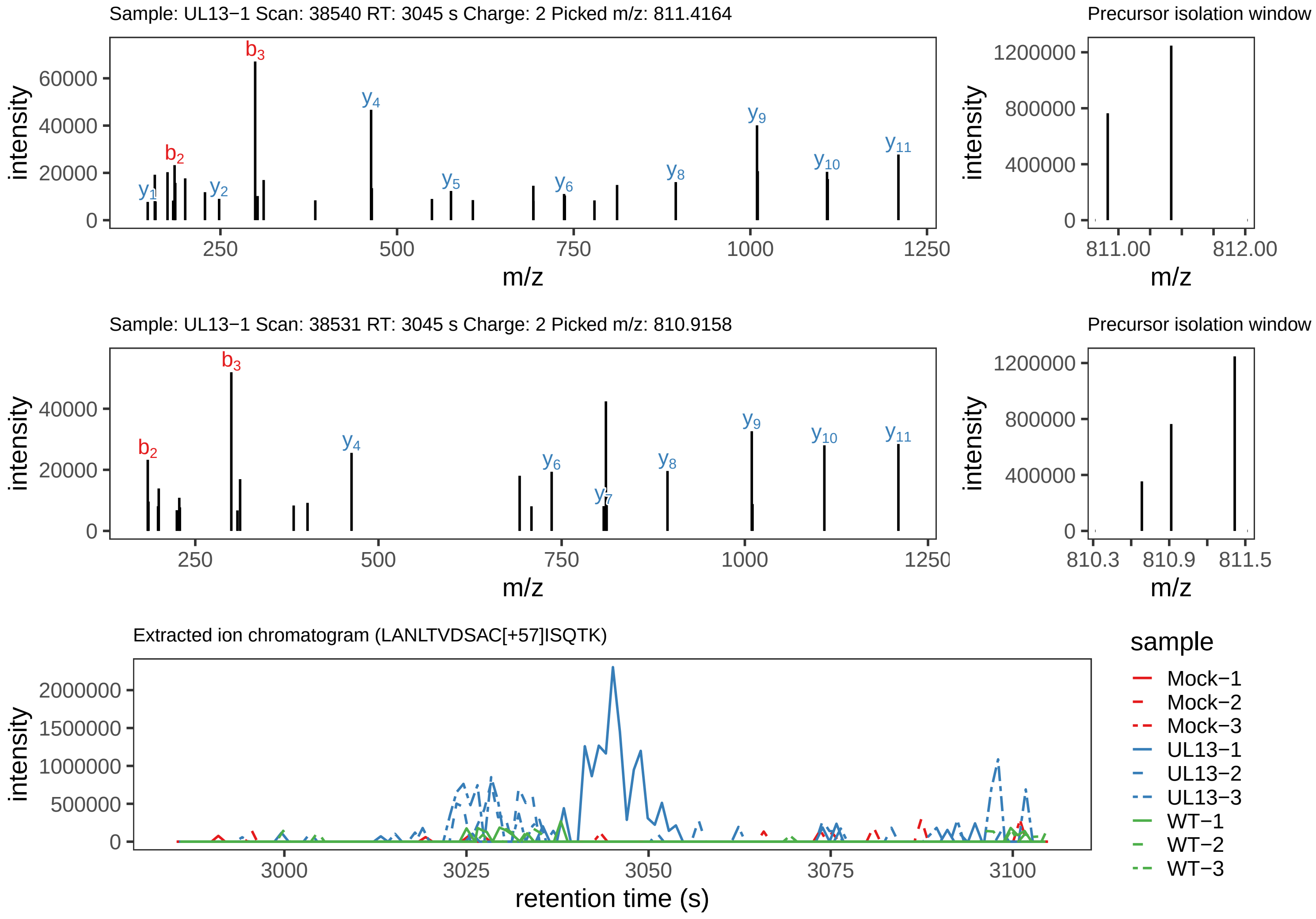

Supplement: Supplemental File 5 [file 737029_file08.zip › phosphopeptide_validation_figs_20240531/LANLTVDSACISQTK/LANLTVDSAC[+57]ISQTK.phospho.pdf]

### Peptide LANLT[+80]VDSAC[+57]ISQTK

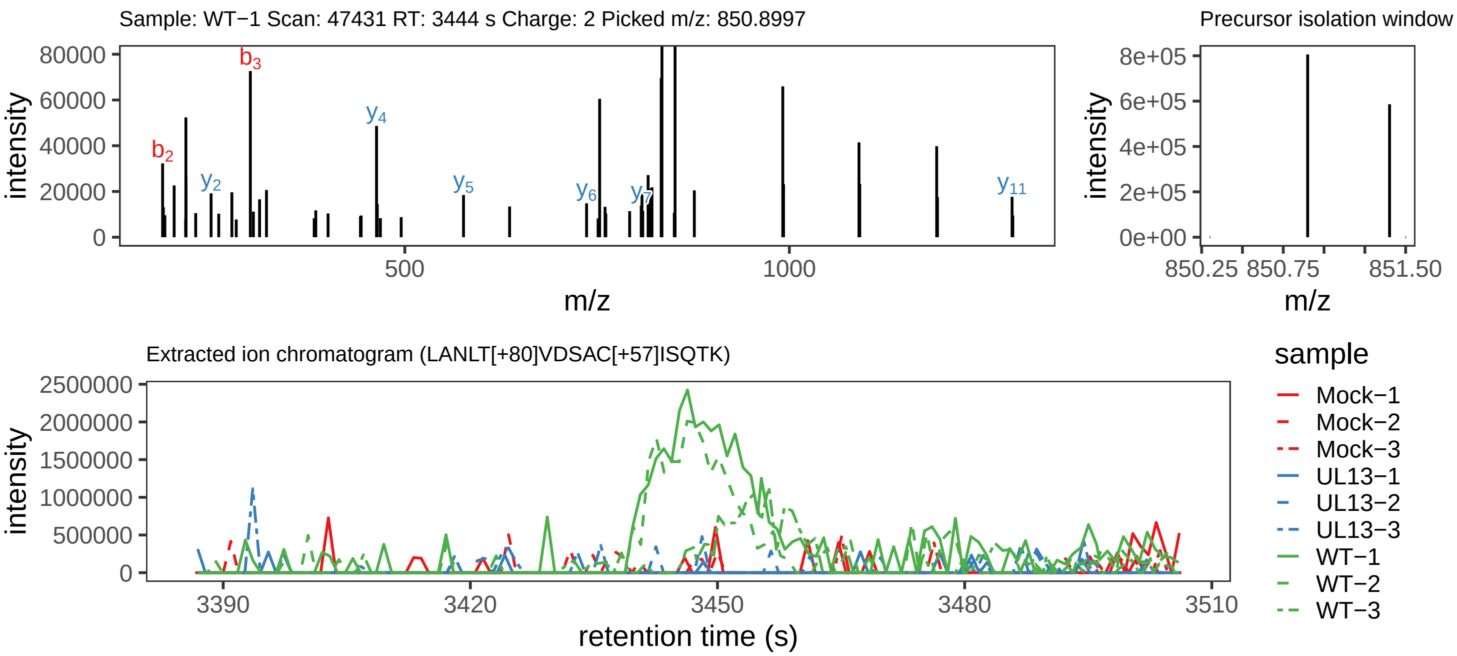

Supplement: Supplemental File 5 [file 737029_file08.zip › phosphopeptide_validation_figs_20240531/LANLTVDSACISQTK/LANLT[+80]VDSAC[+57]ISQTK.phospho.pdf]

### Peptide LANLTVDS[+80]AC[+57]ISQTK

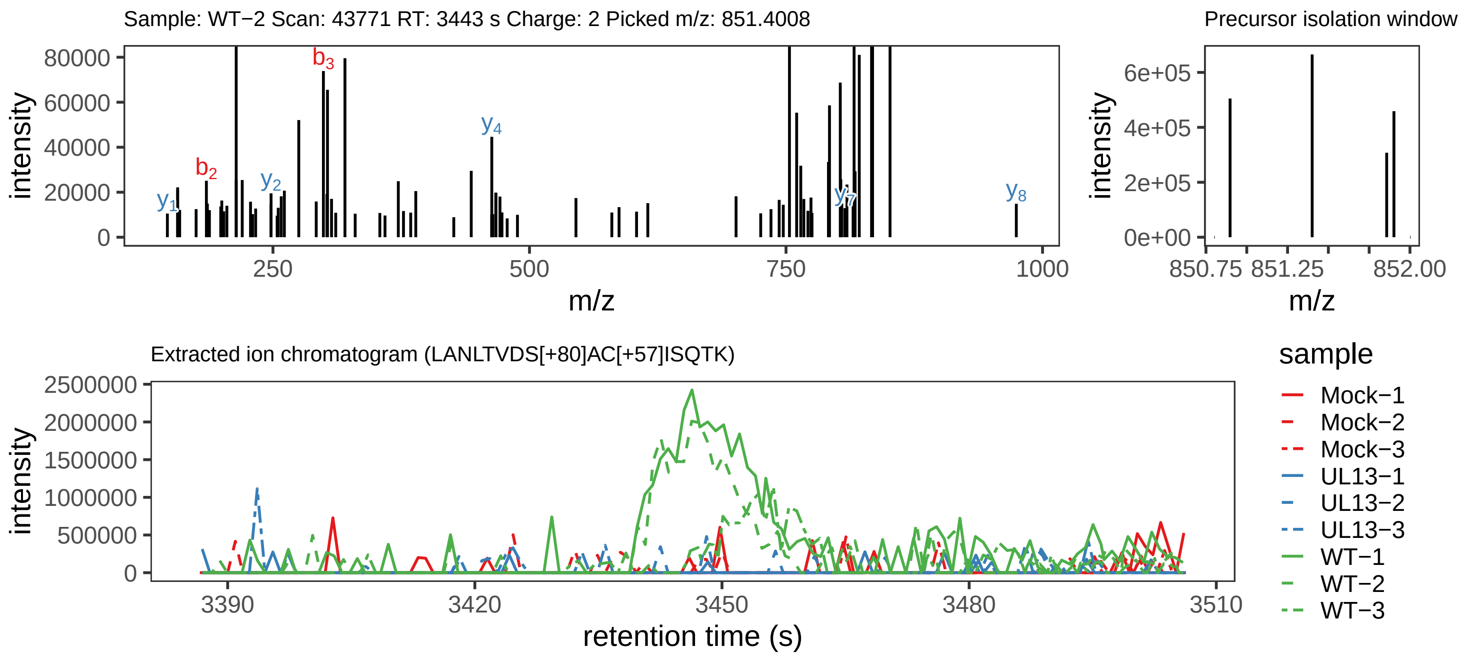

Supplement: Supplemental File 5 [file 737029_file08.zip › phosphopeptide_validation_figs_20240531/LANLTVDSACISQTK/LANLTVDS[+80]AC[+57]ISQTK.phospho.pdf]

### Peptide IAGLY[+80]GVPGSDYAYPR

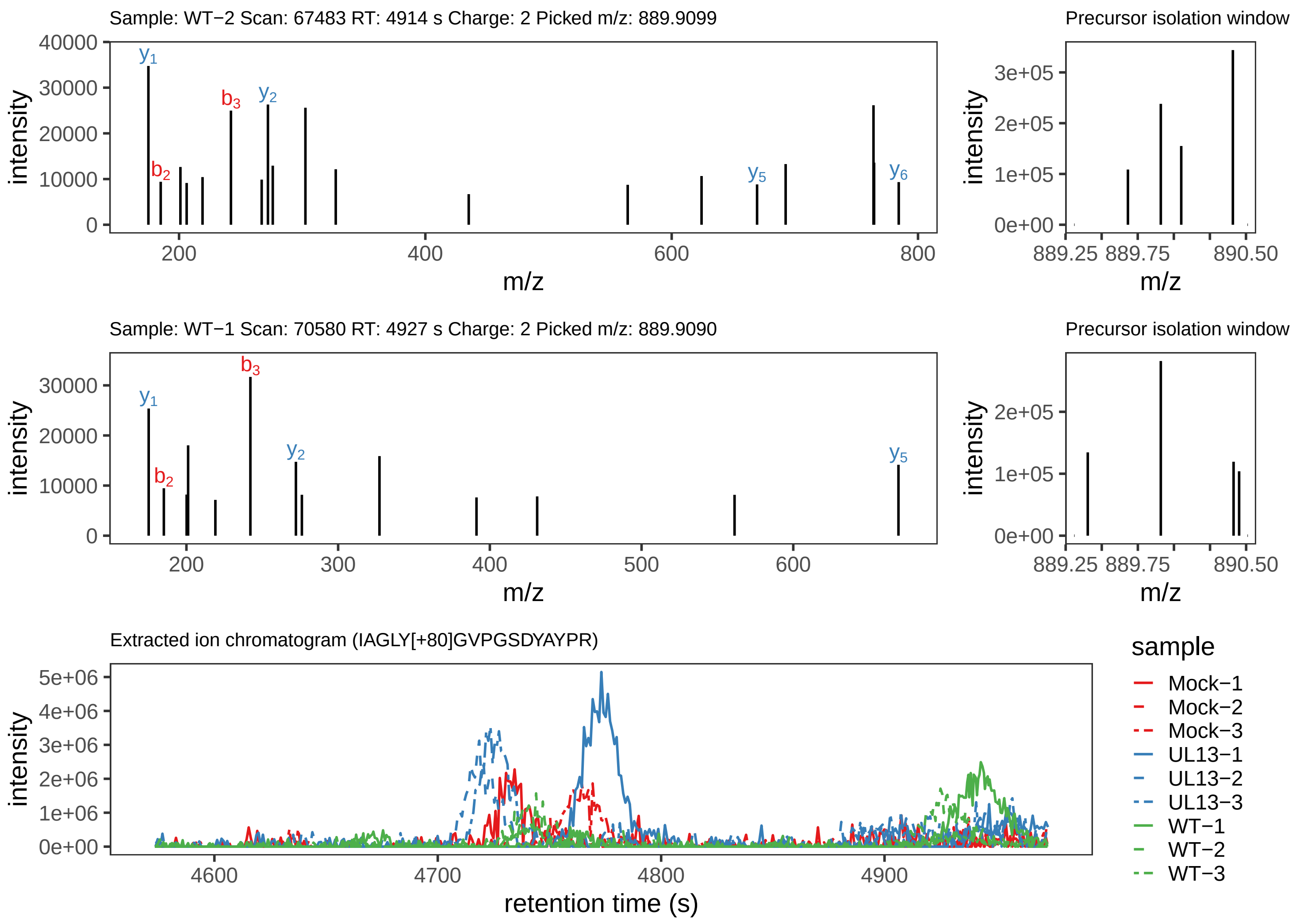

Supplement: Supplemental File 5 [file 737029_file08.zip › phosphopeptide_validation_figs_20240531/IAGLYGVPGSDYAYPR/IAGLY[+80]GVPGSDYAYPR.phospho.pdf]

### Peptide IAGLYGVPGSDYAYPR

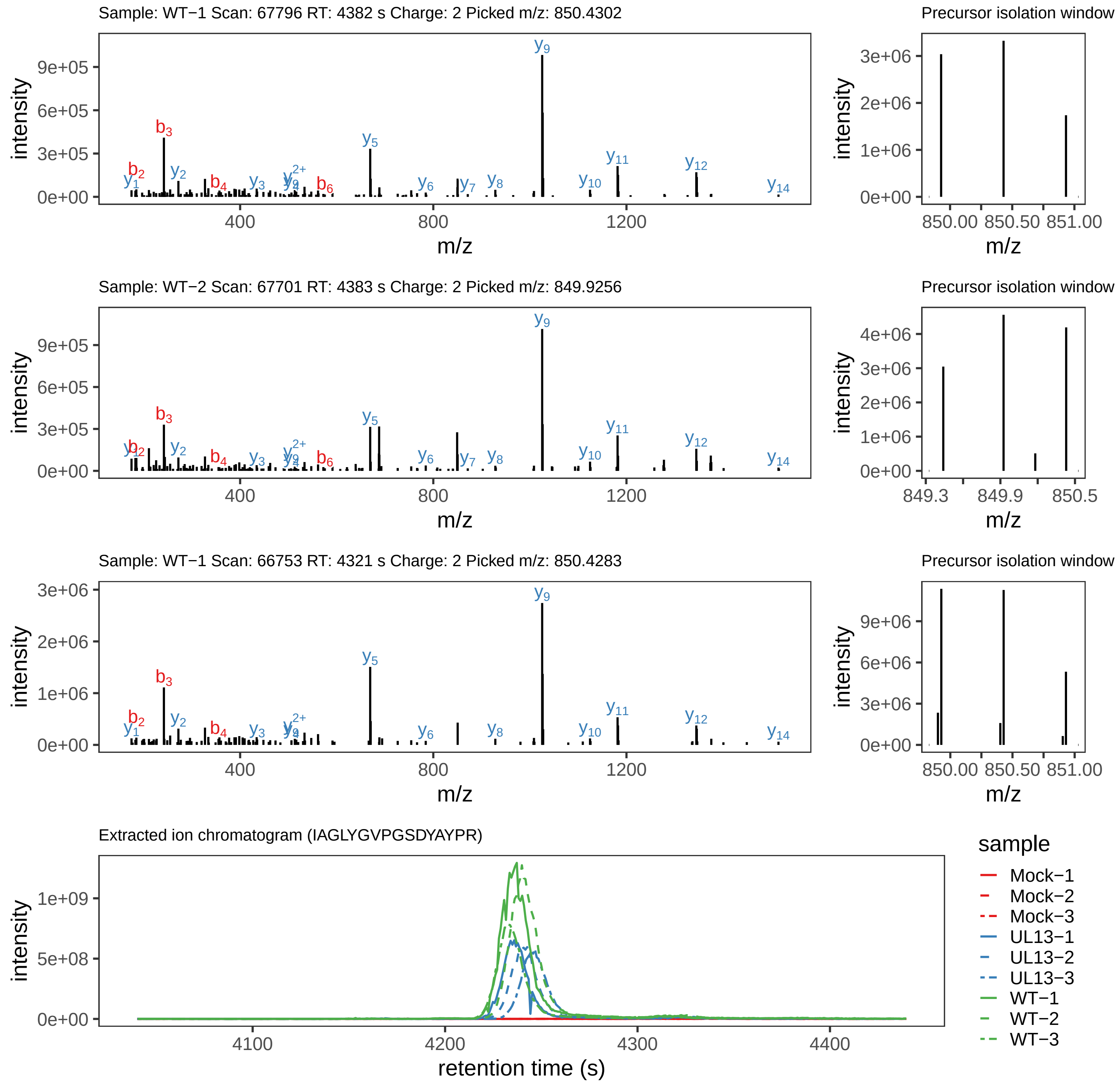

Supplement: Supplemental File 5 [file 737029_file08.zip › phosphopeptide_validation_figs_20240531/IAGLYGVPGSDYAYPR/IAGLYGVPGSDYAYPR.global.pdf]

### Peptide IAGLYGVPGSDYAYPR

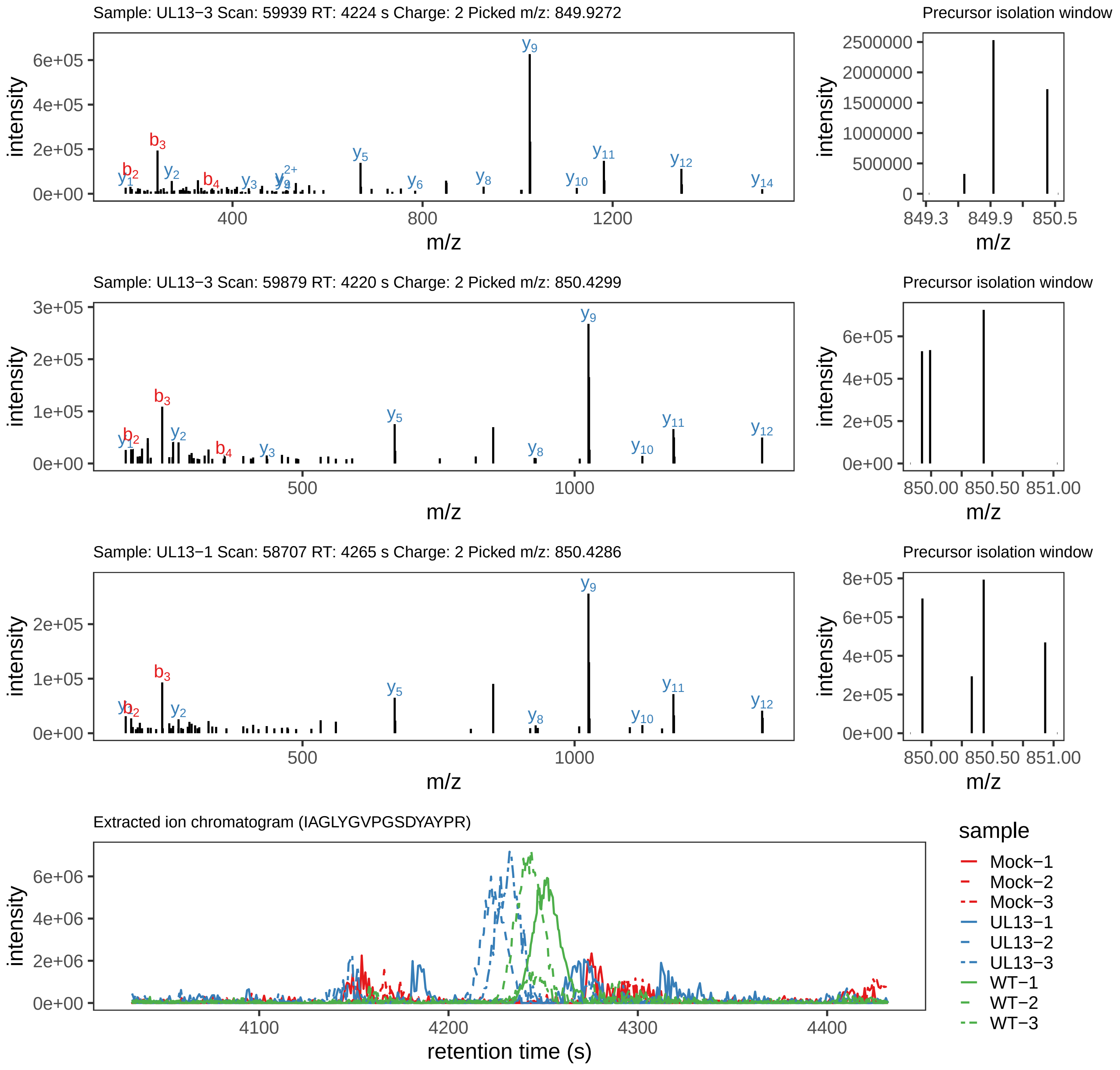

Supplement: Supplemental File 5 [file 737029_file08.zip › phosphopeptide_validation_figs_20240531/IAGLYGVPGSDYAYPR/IAGLYGVPGSDYAYPR.phospho.pdf]

### Peptide IAGLYGVPGS[+80]DYAYPR

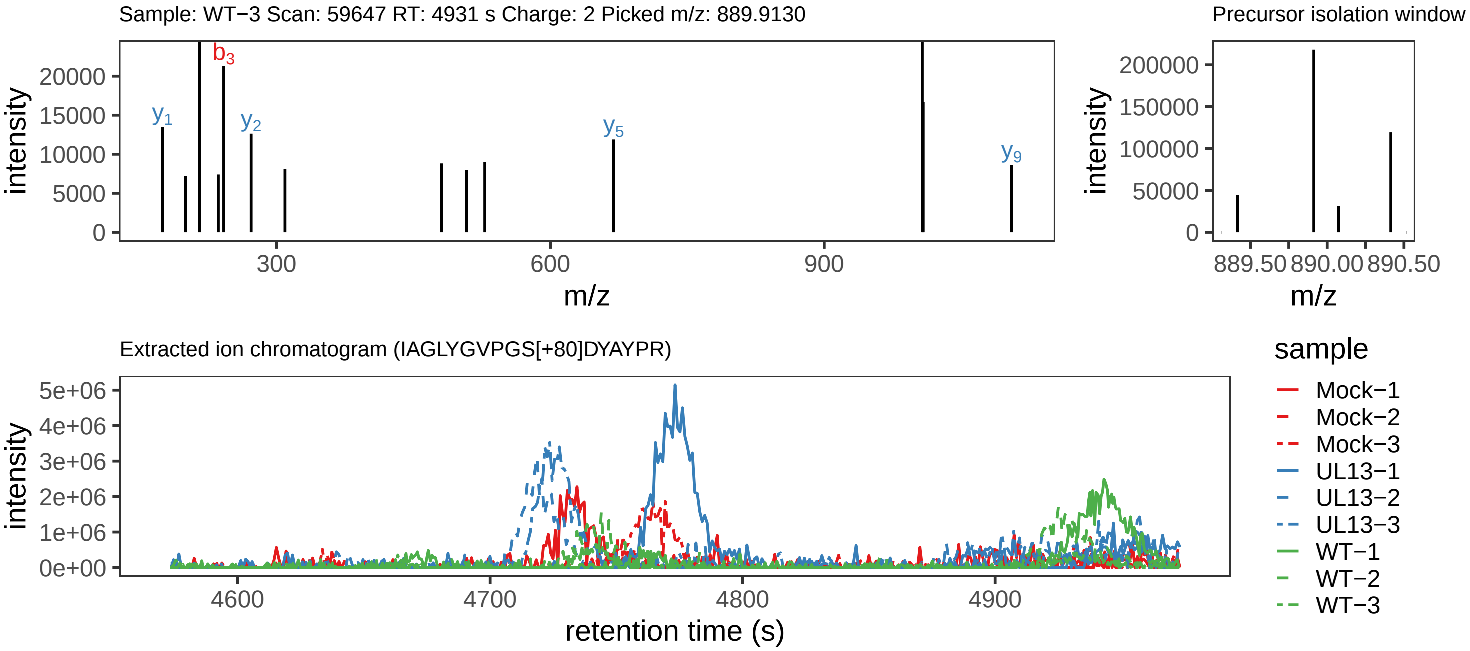

Supplement: Supplemental File 5 [file 737029_file08.zip › phosphopeptide_validation_figs_20240531/IAGLYGVPGSDYAYPR/IAGLYGVPGS[+80]DYAYPR.phospho.pdf]

### Peptide RPYEFDRS[+80]IESDLYYPGEFR

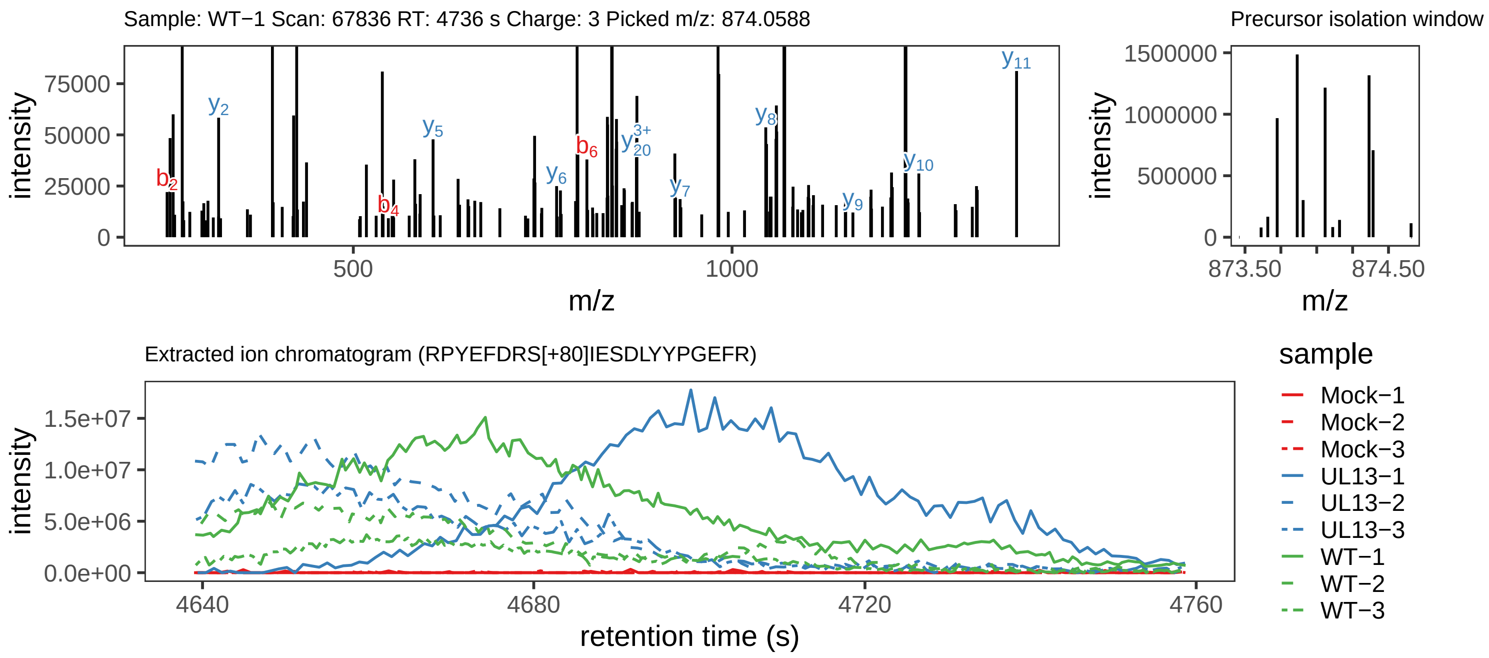

Supplement: Supplemental File 5 [file 737029_file08.zip › phosphopeptide_validation_figs_20240531/RPYEFDRSIESDLYYPGEFR/RPYEFDRS[+80]IESDLYYPGEFR.phospho.pdf]

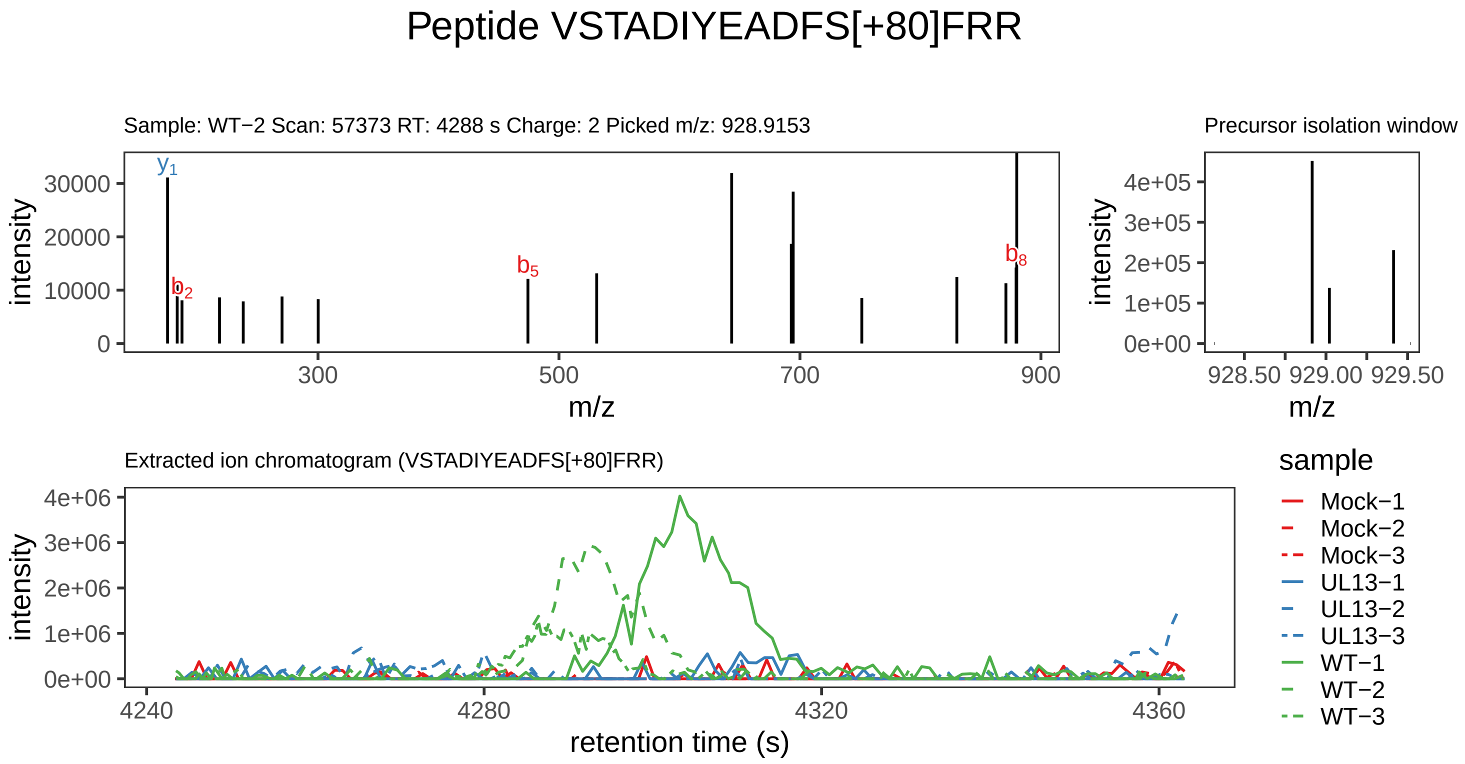

Supplement: Supplemental File 5 [file 737029_file08.zip › phosphopeptide_validation_figs_20240531/VSTADIYEADFSFRR/VSTADIYEADFS[+80]FRR.phospho.pdf]

### Peptide FGTTVNSR

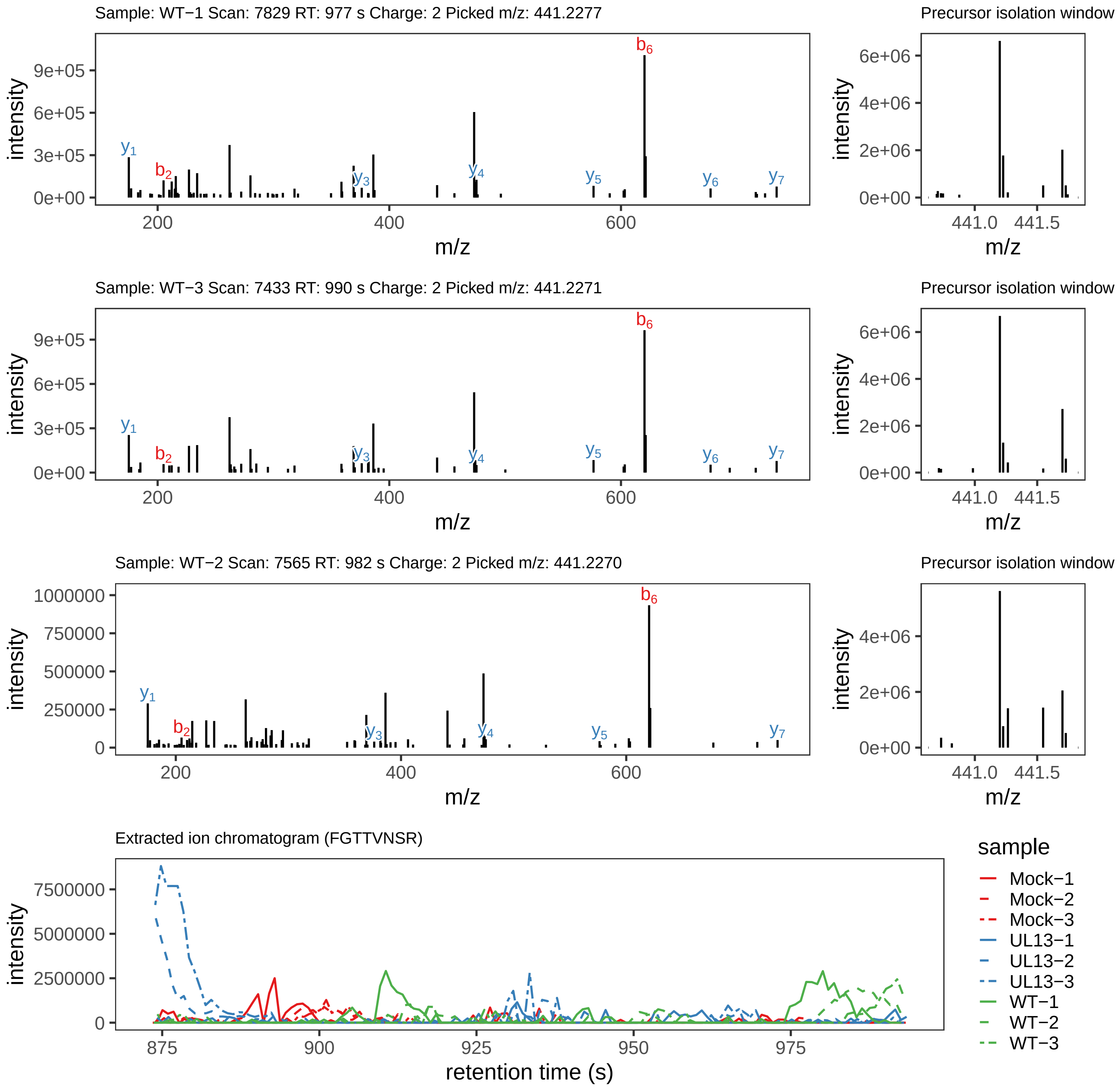

Supplement: Supplemental File 5 [file 737029_file08.zip › phosphopeptide_validation_figs_20240531/FGTTVNSR/FGTTVNSR.global.pdf]

### Peptide FGT[+80]TVNSR

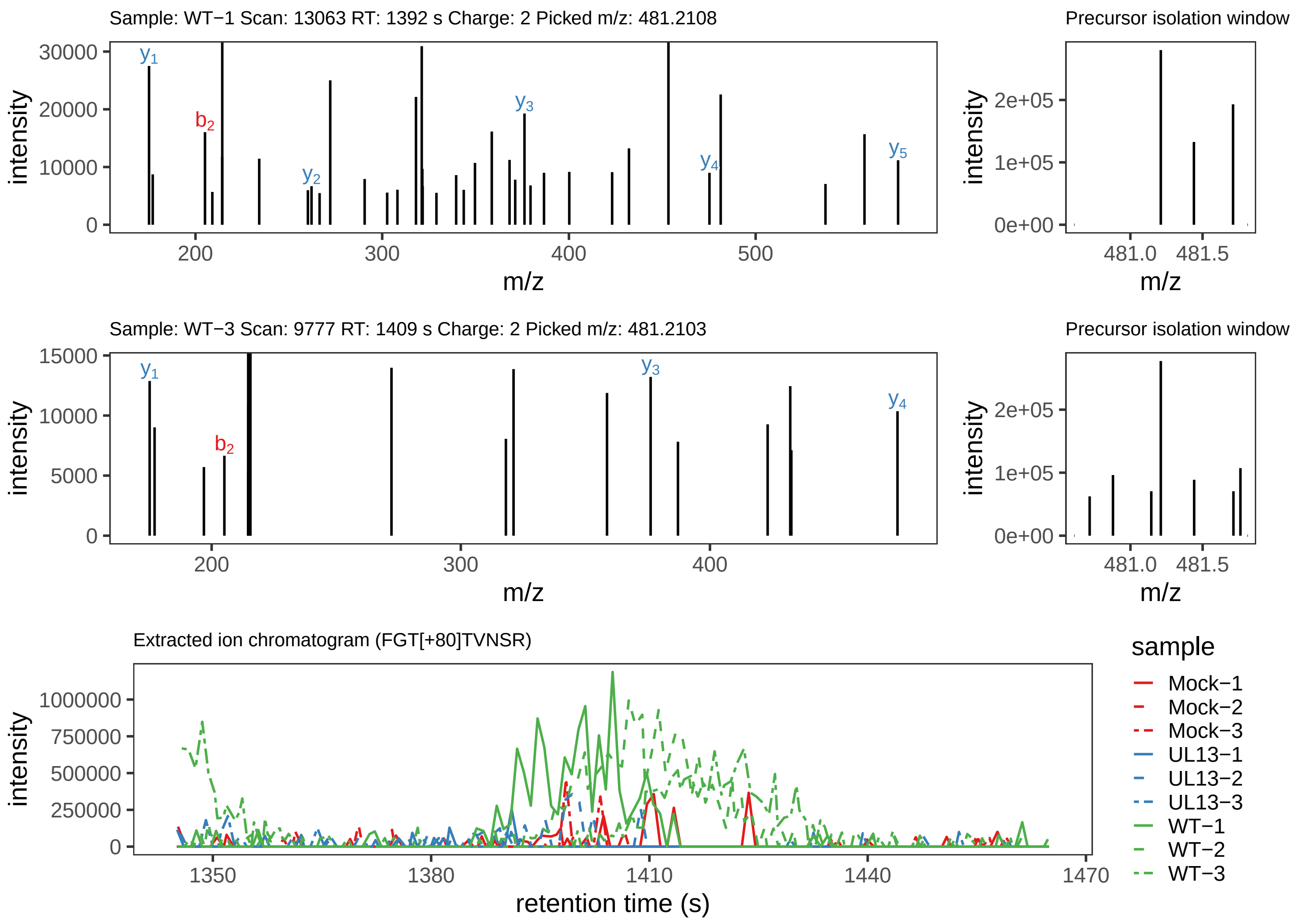

Supplement: Supplemental File 5 [file 737029_file08.zip › phosphopeptide_validation_figs_20240531/FGTTVNSR/FGT[+80]TVNSR.phospho.pdf]

### Peptide FGTTVN[+1]SR

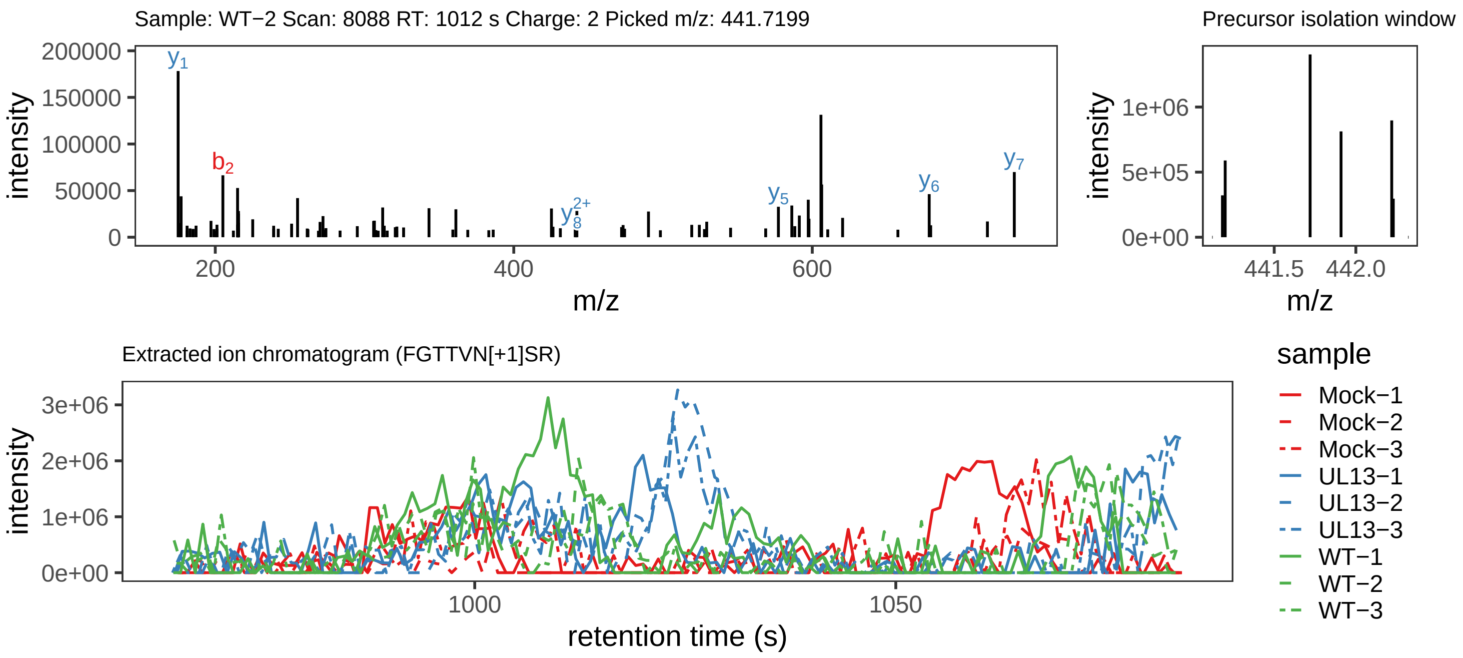

Supplement: Supplemental File 5 [file 737029_file08.zip › phosphopeptide_validation_figs_20240531/FGTTVNSR/FGTTVN[+1]SR.global.pdf]

### Peptide ILT[+80]ATPHRY

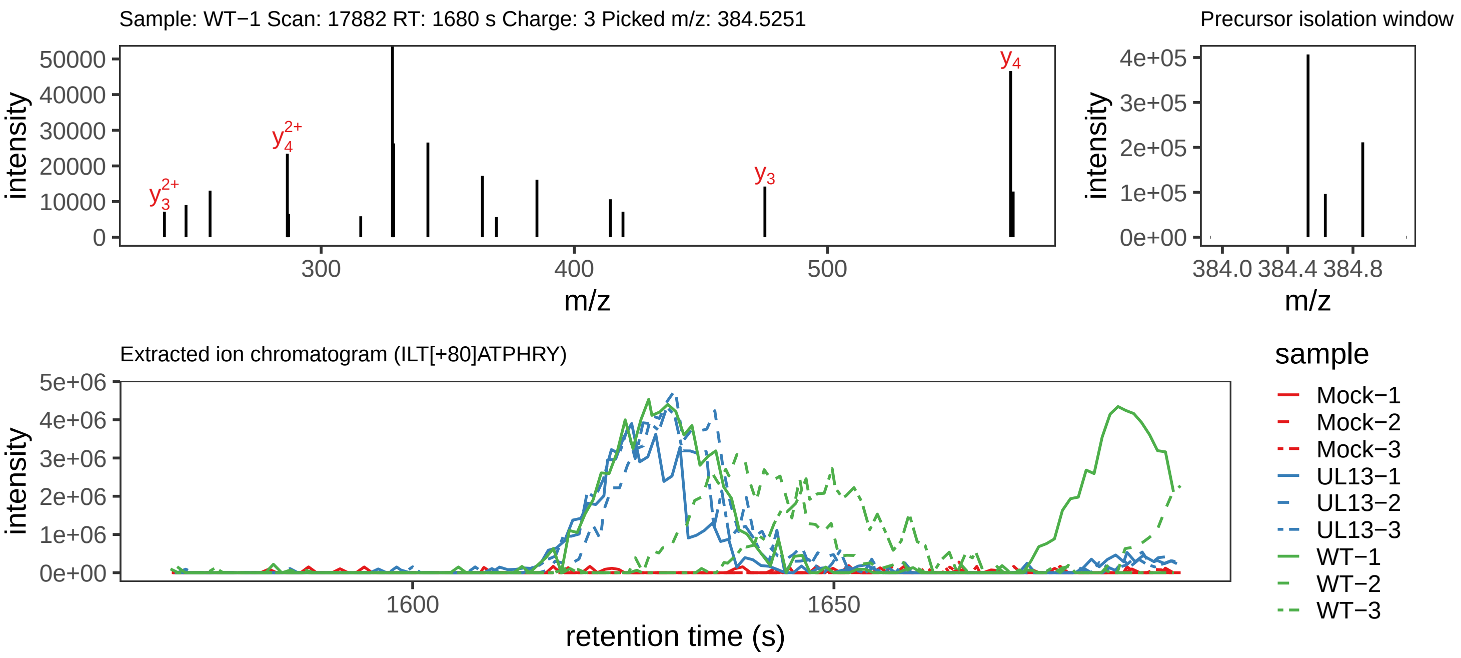

Supplement: Supplemental File 5 [file 737029_file08.zip › phosphopeptide_validation_figs_20240531/ILTATPHRY/ILT[+80]ATPHRY.phospho.pdf]

### Peptide ILTAT[+80]PHRY

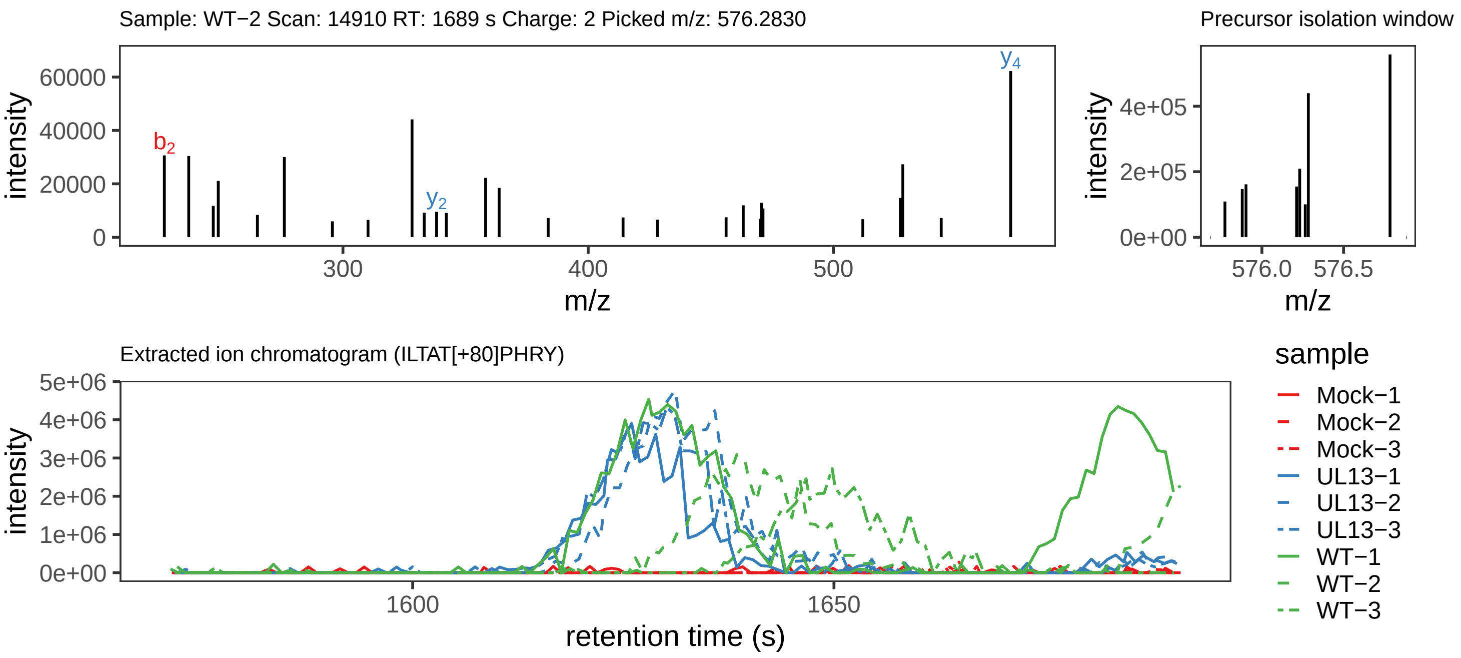

Supplement: Supplemental File 5 [file 737029_file08.zip › phosphopeptide_validation_figs_20240531/ILTATPHRY/ILTAT[+80]PHRY.phospho.pdf]

### Peptide TIIS[+80]TPAVTPK

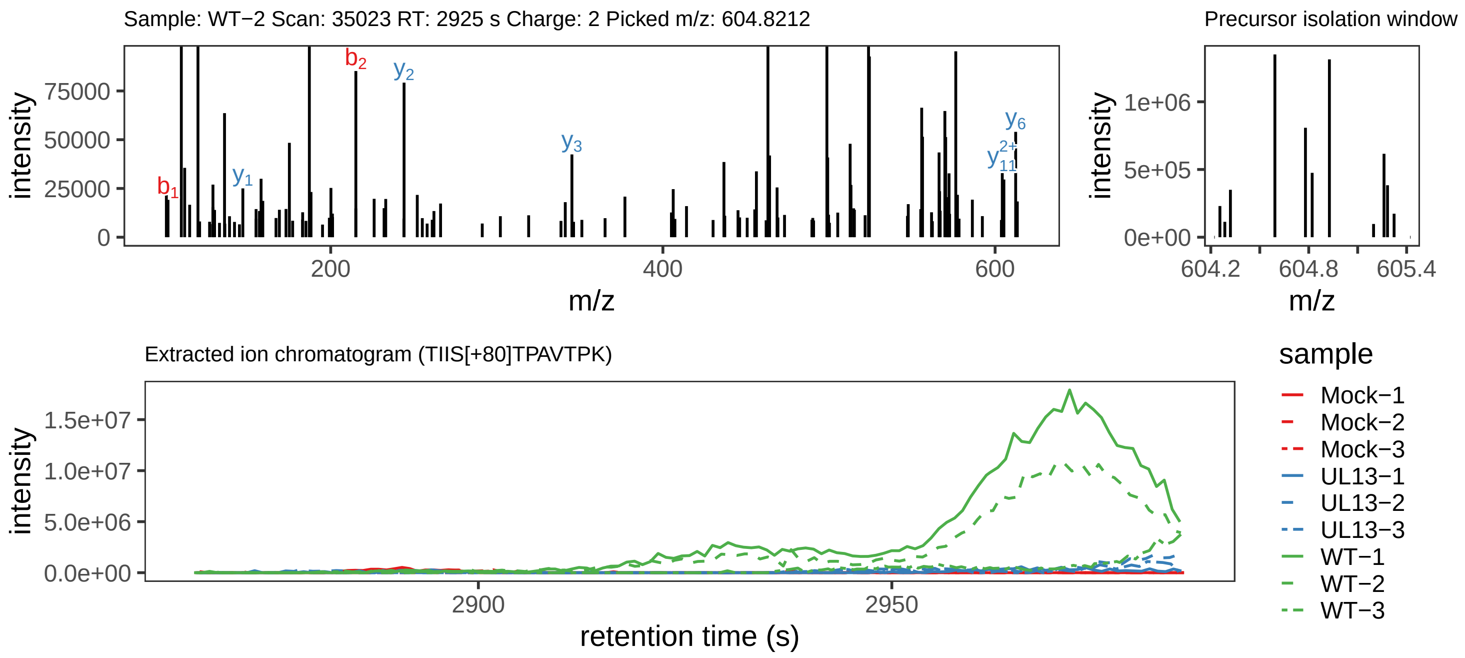

Supplement: Supplemental File 5 [file 737029_file08.zip › phosphopeptide_validation_figs_20240531/TIISTPAVTPK/TIIS[+80]TPAVTPK.phospho.pdf]

### Peptide T[+80]IISTPAVTPK

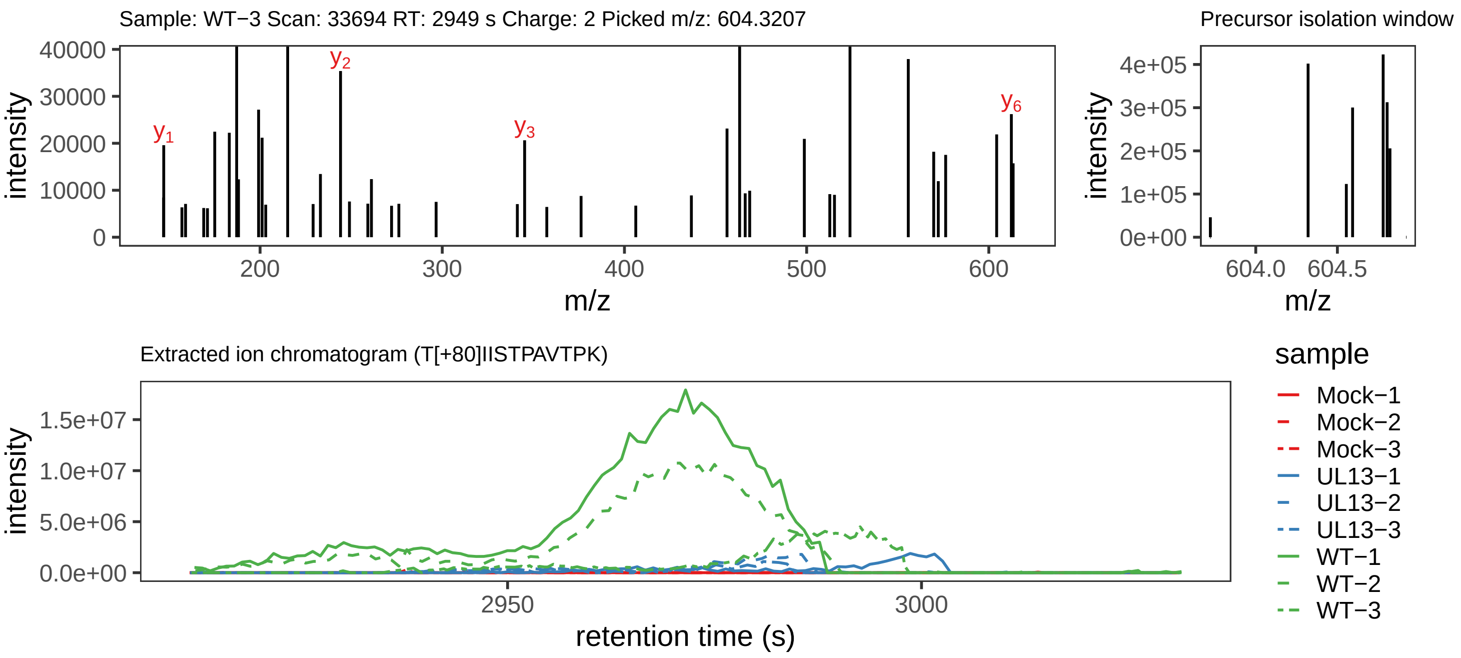

Supplement: Supplemental File 5 [file 737029_file08.zip › phosphopeptide_validation_figs_20240531/TIISTPAVTPK/T[+80]IISTPAVTPK.phospho.pdf]

### Peptide TIIST[+80]PAVT[+80]PK

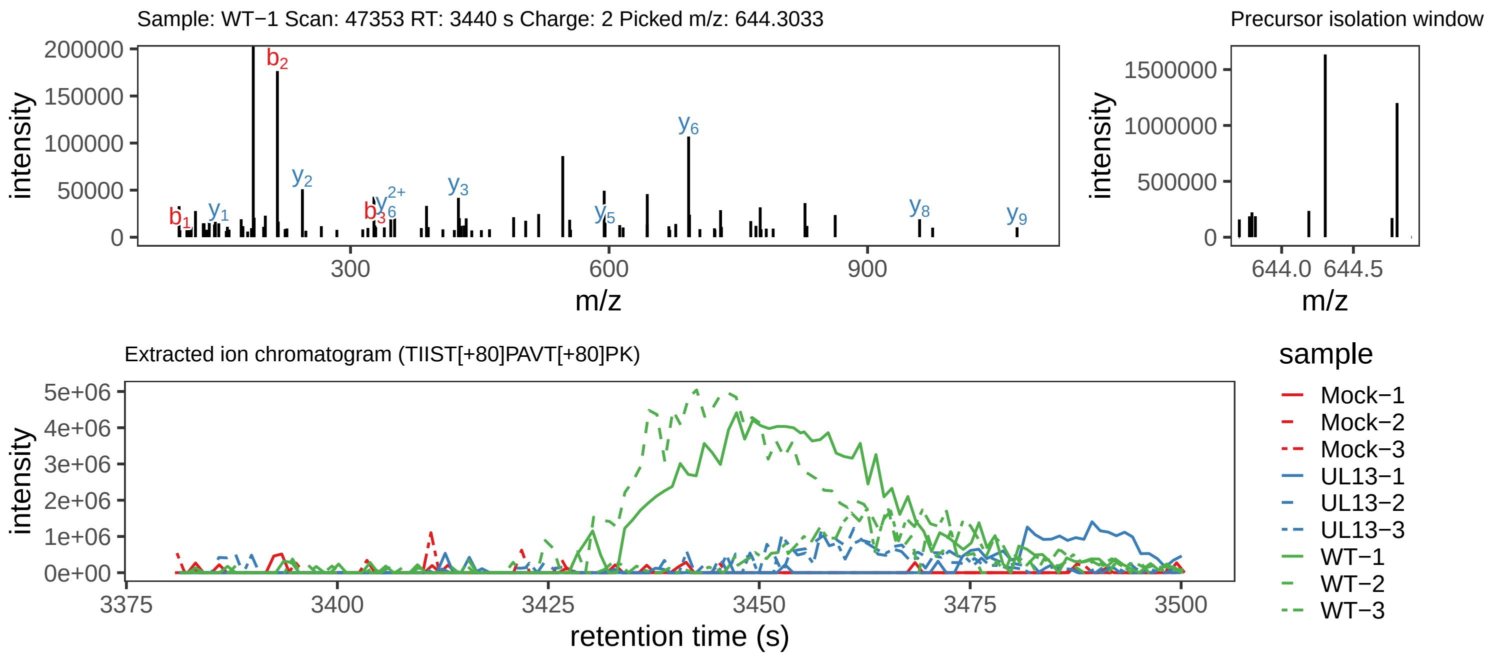

Supplement: Supplemental File 5 [file 737029_file08.zip › phosphopeptide_validation_figs_20240531/TIISTPAVTPK/TIIST[+80]PAVT[+80]PK.phospho.pdf]

### Peptide TIIST[+80]PAVTPK

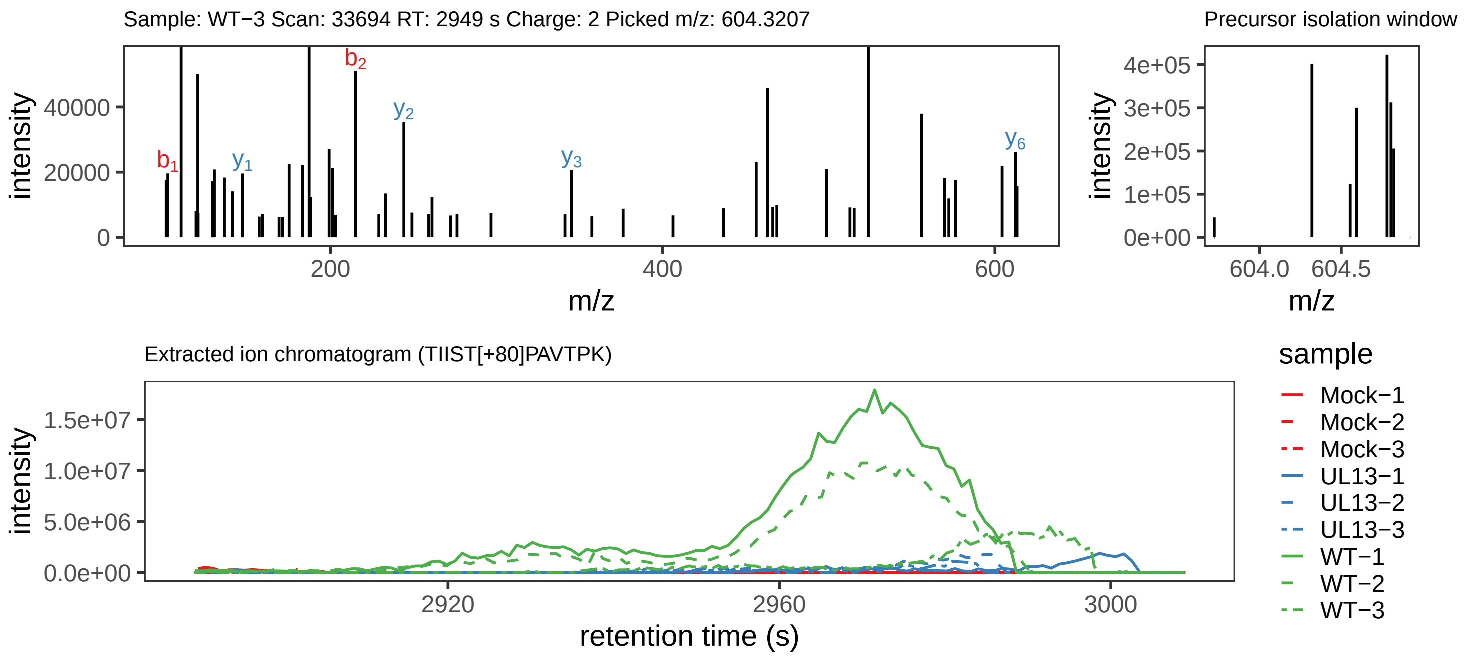

Supplement: Supplemental File 5 [file 737029_file08.zip › phosphopeptide_validation_figs_20240531/TIISTPAVTPK/TIIST[+80]PAVTPK.phospho.pdf]

### Peptide TIISTPAVT[+80]PK

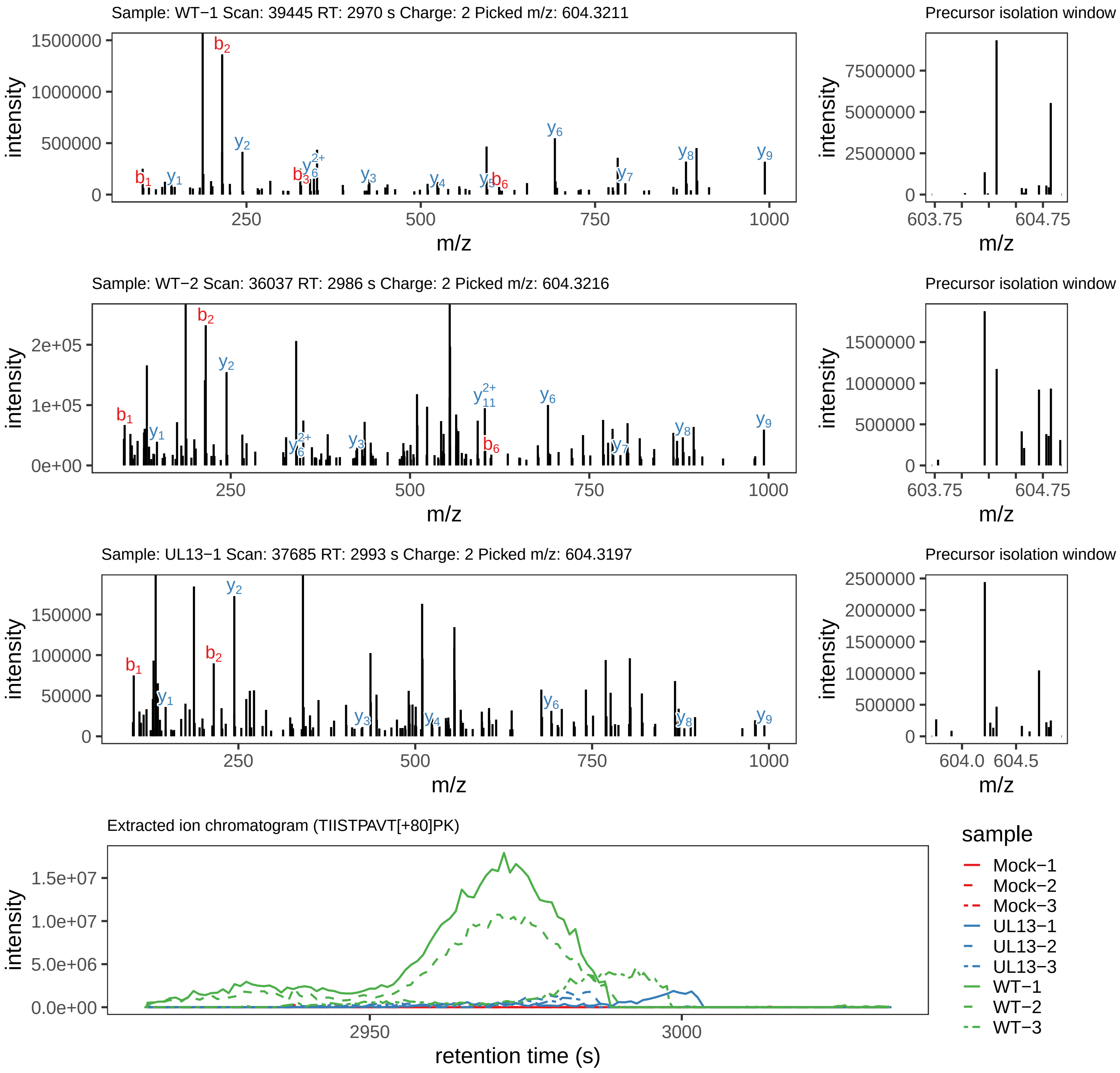

Supplement: Supplemental File 5 [file 737029_file08.zip › phosphopeptide_validation_figs_20240531/TIISTPAVTPK/TIISTPAVT[+80]PK.phospho.pdf]

### Peptide TIISTPAVTPK

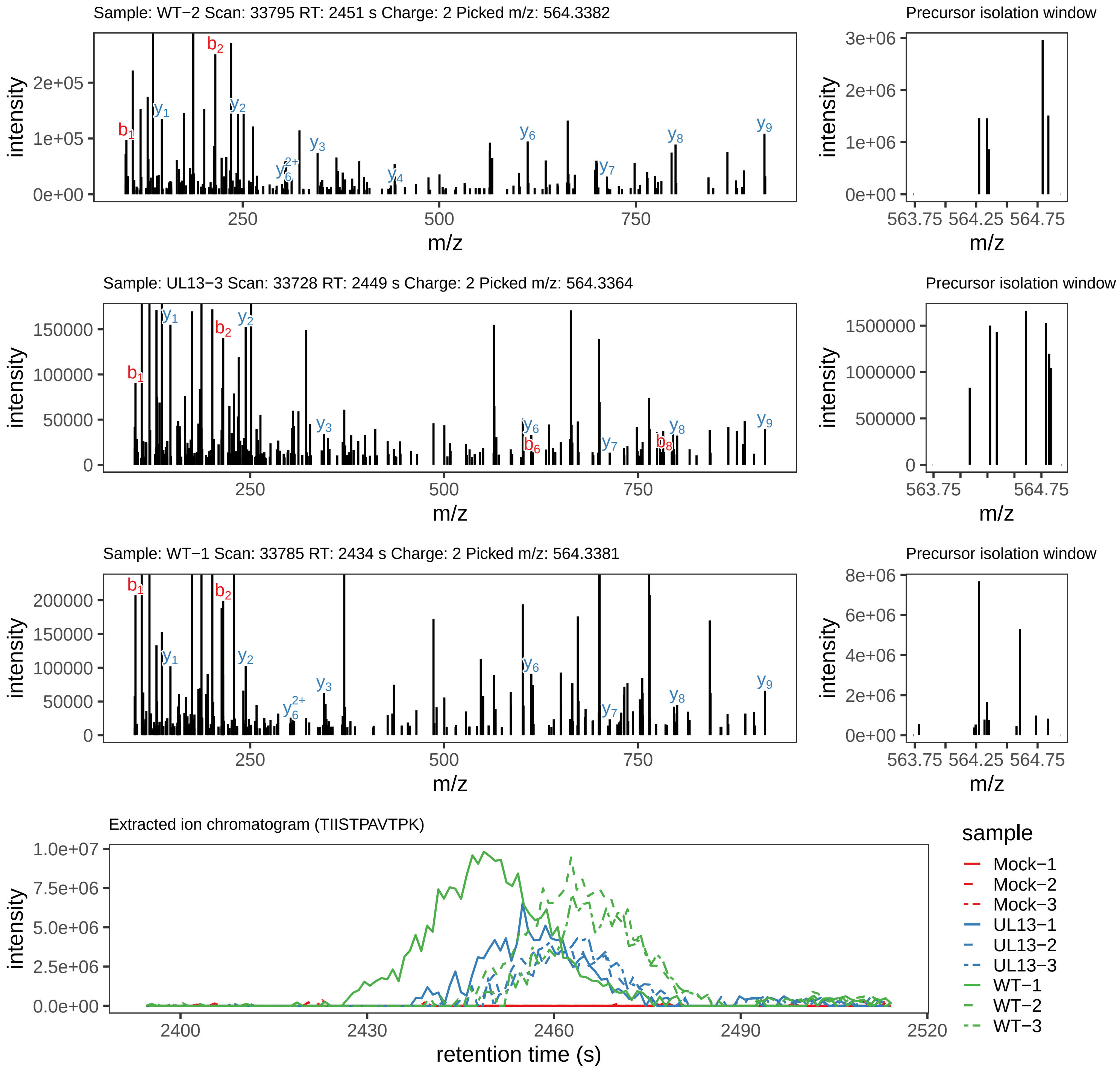

Supplement: Supplemental File 5 [file 737029_file08.zip › phosphopeptide_validation_figs_20240531/TIISTPAVTPK/TIISTPAVTPK.global.pdf]

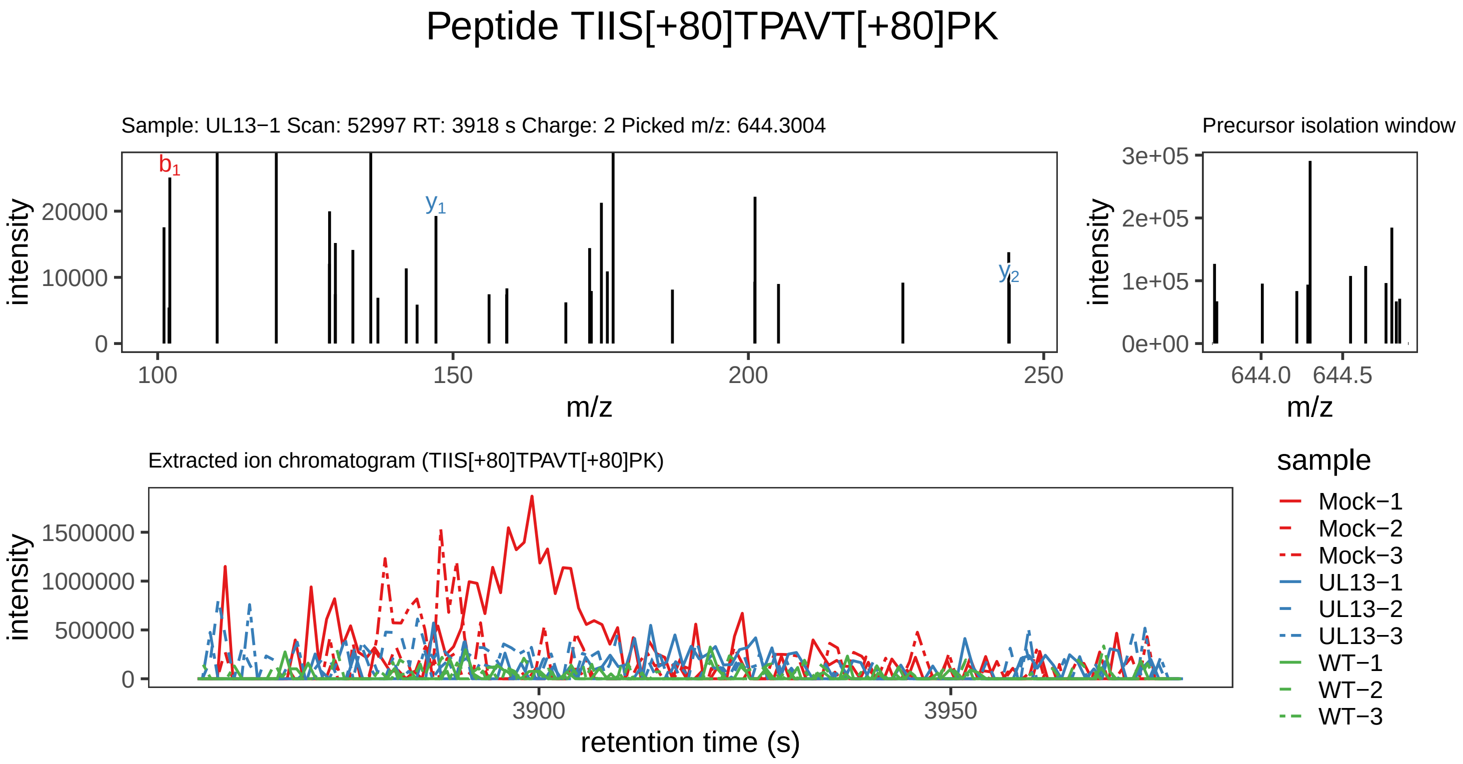

Supplement: Supplemental File 5 [file 737029_file08.zip › phosphopeptide_validation_figs_20240531/TIISTPAVTPK/TIIS[+80]TPAVT[+80]PK.phospho.pdf]

### Peptide RQLTDT[+80]IRR

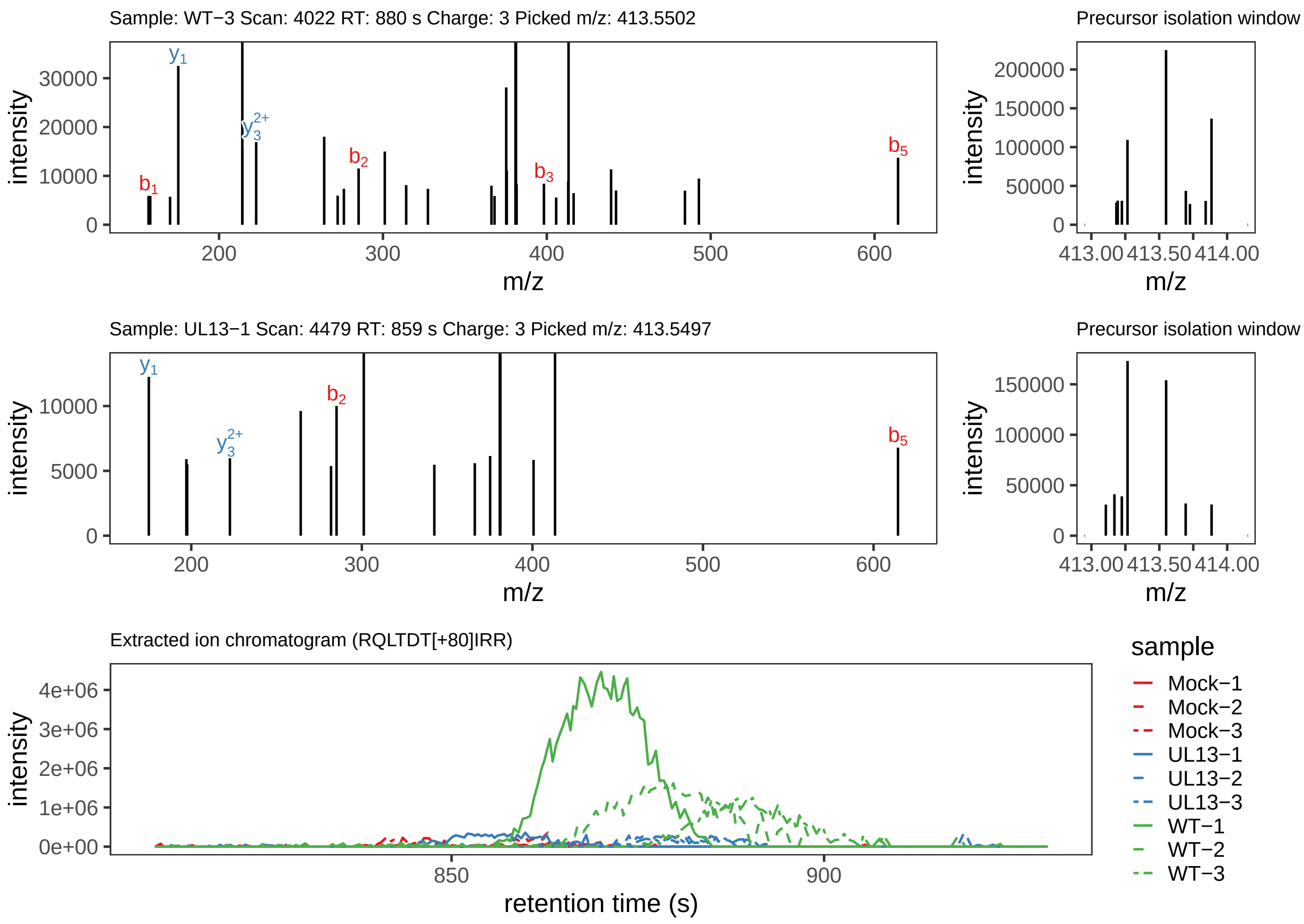

Supplement: Supplemental File 5 [file 737029_file08.zip › phosphopeptide_validation_figs_20240531/RQLTDTIRR/RQLTDT[+80]IRR.phospho.pdf]

### Peptide RQLTDTIRR

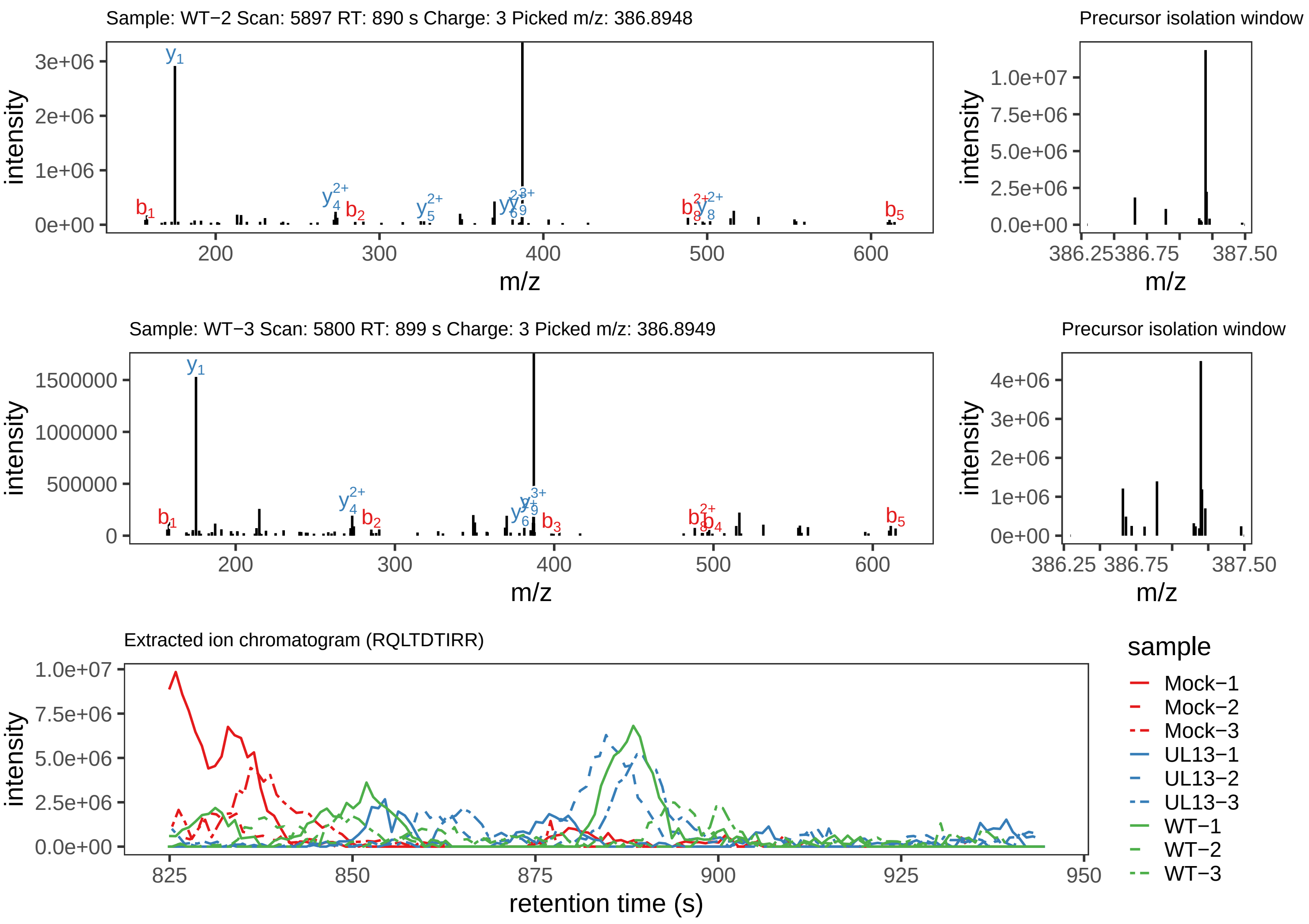

Supplement: Supplemental File 5 [file 737029_file08.zip › phosphopeptide_validation_figs_20240531/RQLTDTIRR/RQLTDTIRR.global.pdf]

### Peptide AVLWTSDLQHTSPNR

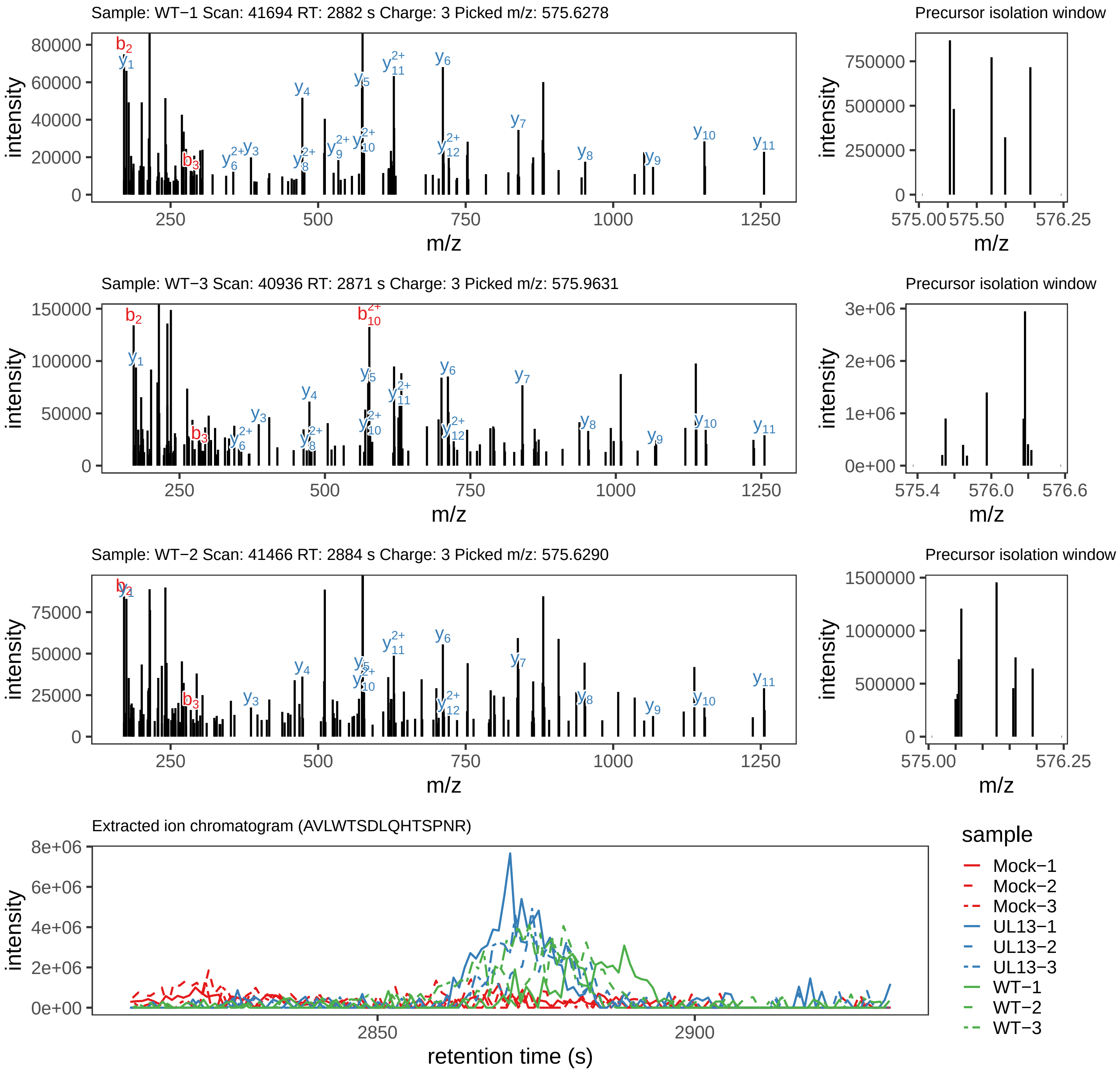

Supplement: Supplemental File 5 [file 737029_file08.zip › phosphopeptide_validation_figs_20240531/AVLWTSDLQHTSPNR/AVLWTSDLQHTSPNR.global.pdf]

### Peptide AVLWTSDLQHTS[+80]PNR

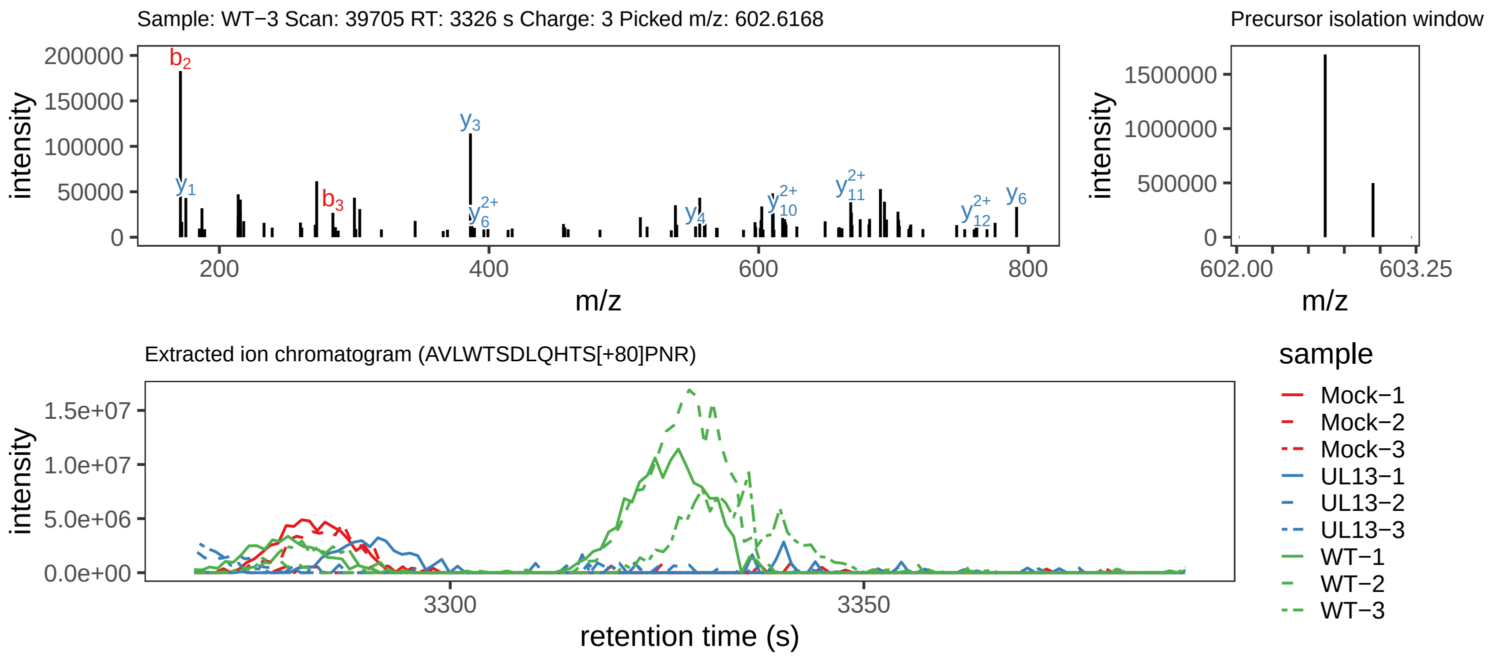

Supplement: Supplemental File 5 [file 737029_file08.zip › phosphopeptide_validation_figs_20240531/AVLWTSDLQHTSPNR/AVLWTSDLQHTS[+80]PNR.phospho.pdf]

### Peptide AVLWTSDLQHT[+80]SPNR

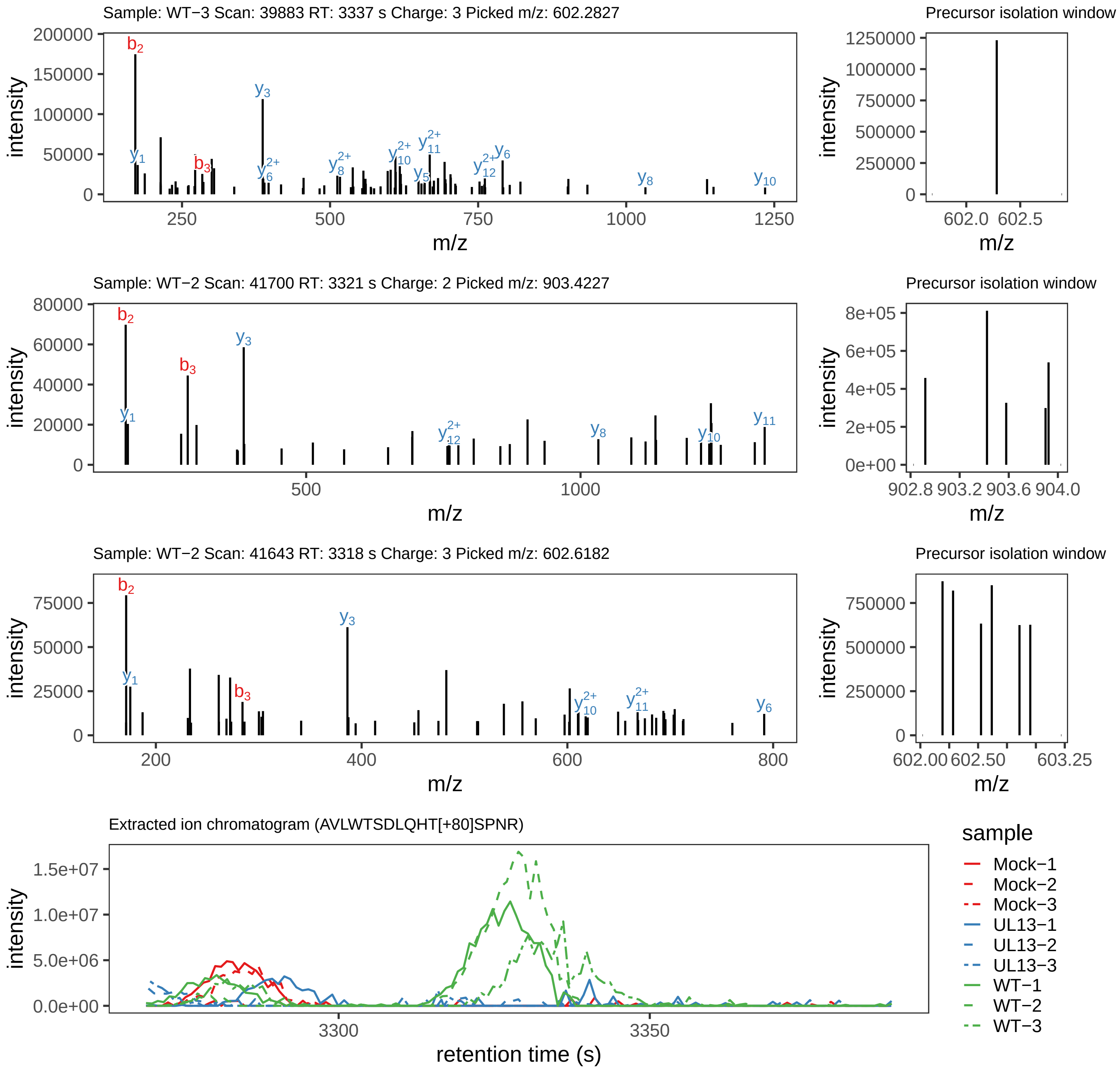

Supplement: Supplemental File 5 [file 737029_file08.zip › phosphopeptide_validation_figs_20240531/AVLWTSDLQHTSPNR/AVLWTSDLQHT[+80]SPNR.phospho.pdf]

### Peptide SNLS[+80]DIYFIPGRRTPIVDK

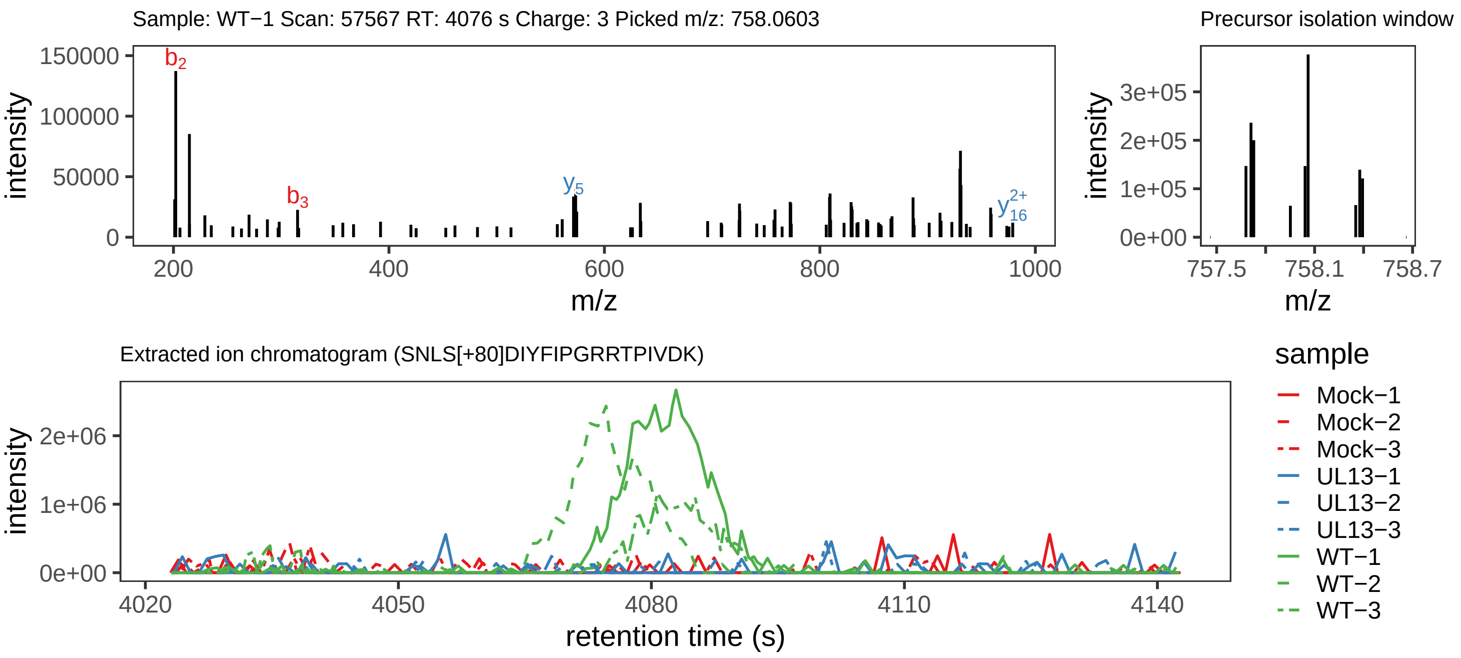

Supplement: Supplemental File 5 [file 737029_file08.zip › phosphopeptide_validation_figs_20240531/SNLSDIYFIPGRRTPIVDK/SNLS[+80]DIYFIPGRRTPIVDK.phospho.pdf]

### Peptide M[+16]TVTLNEYDISAS[+80]PFHPTDPTR

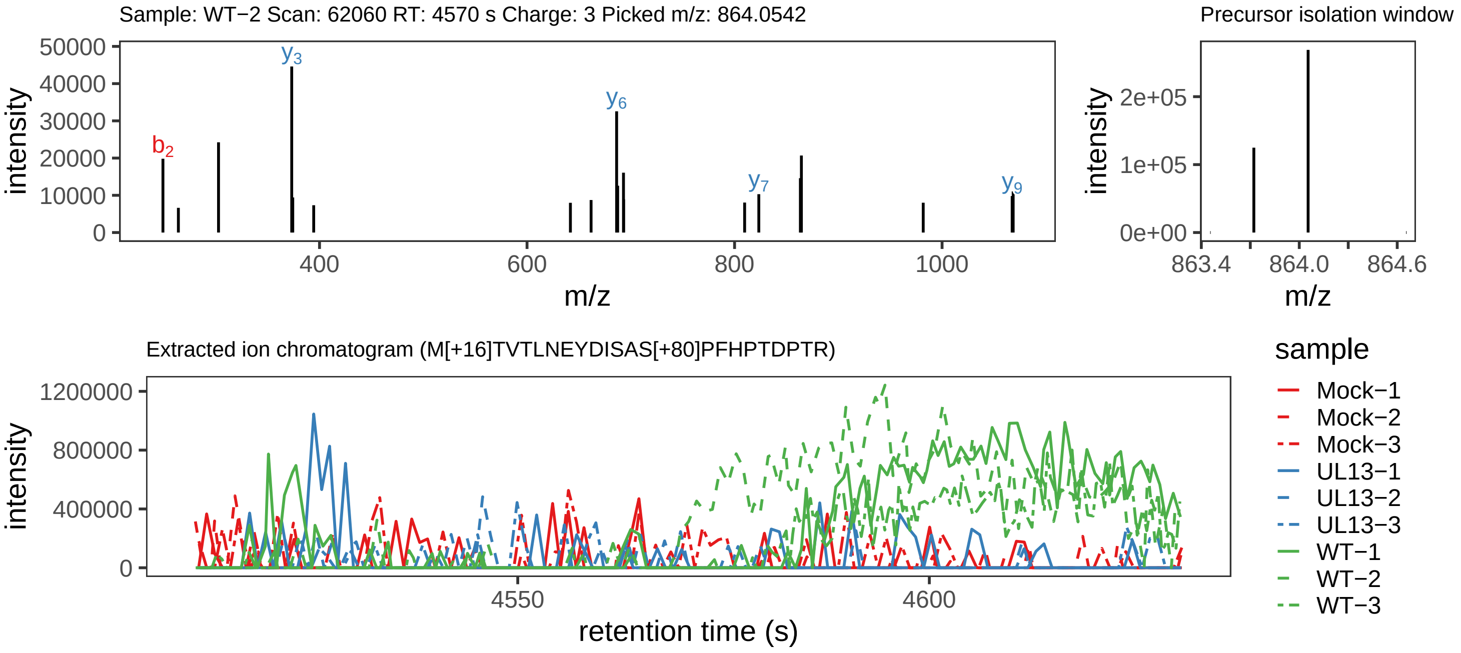

Supplement: Supplemental File 5 [file 737029_file08.zip › phosphopeptide_validation_figs_20240531/MTVTLNEYDISASPFHPTDPTR/M[+16]TVTLNEYDISAS[+80]PFHPTDPTR.phospho.pdf]

### Peptide MTVTLNEYDISAS[+80]PFHPTDPTR

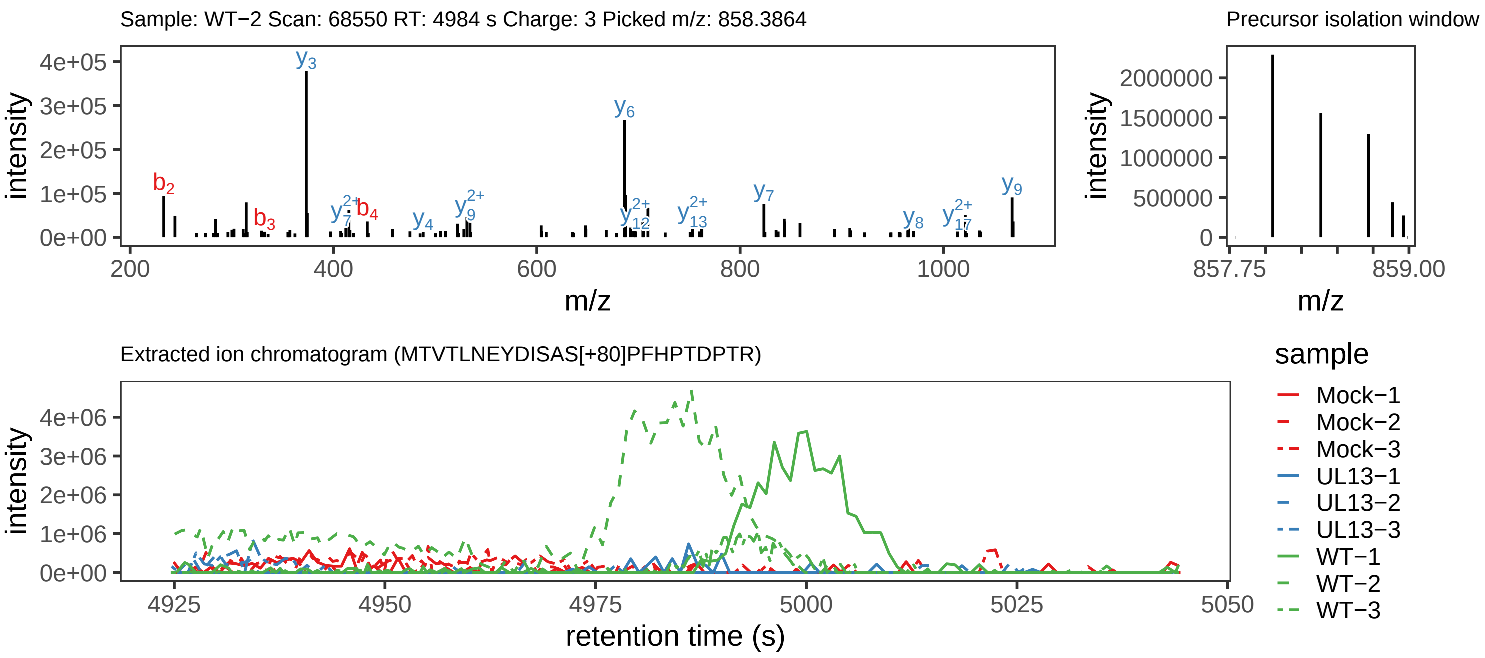

Supplement: Supplemental File 5 [file 737029_file08.zip › phosphopeptide_validation_figs_20240531/MTVTLNEYDISASPFHPTDPTR/MTVTLNEYDISAS[+80]PFHPTDPTR.phospho.pdf]

### Peptide MT[+80]VTLN[+1]EYDISASPFHPTDPTR

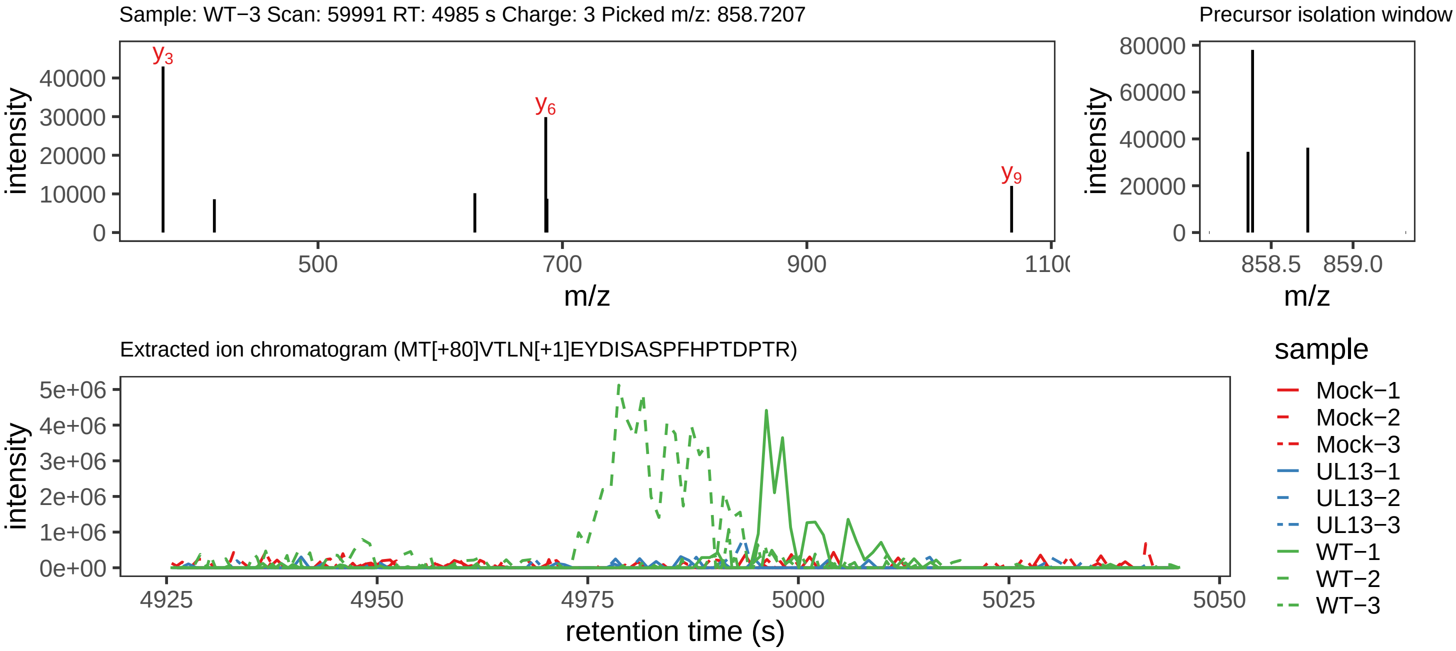

Supplement: Supplemental File 5 [file 737029_file08.zip › phosphopeptide_validation_figs_20240531/MTVTLNEYDISASPFHPTDPTR/MT[+80]VTLN[+1]EYDISASPFHPTDPTR.phospho.pdf]

### Peptide MTVTLNEYDISASPFHPTDPTR

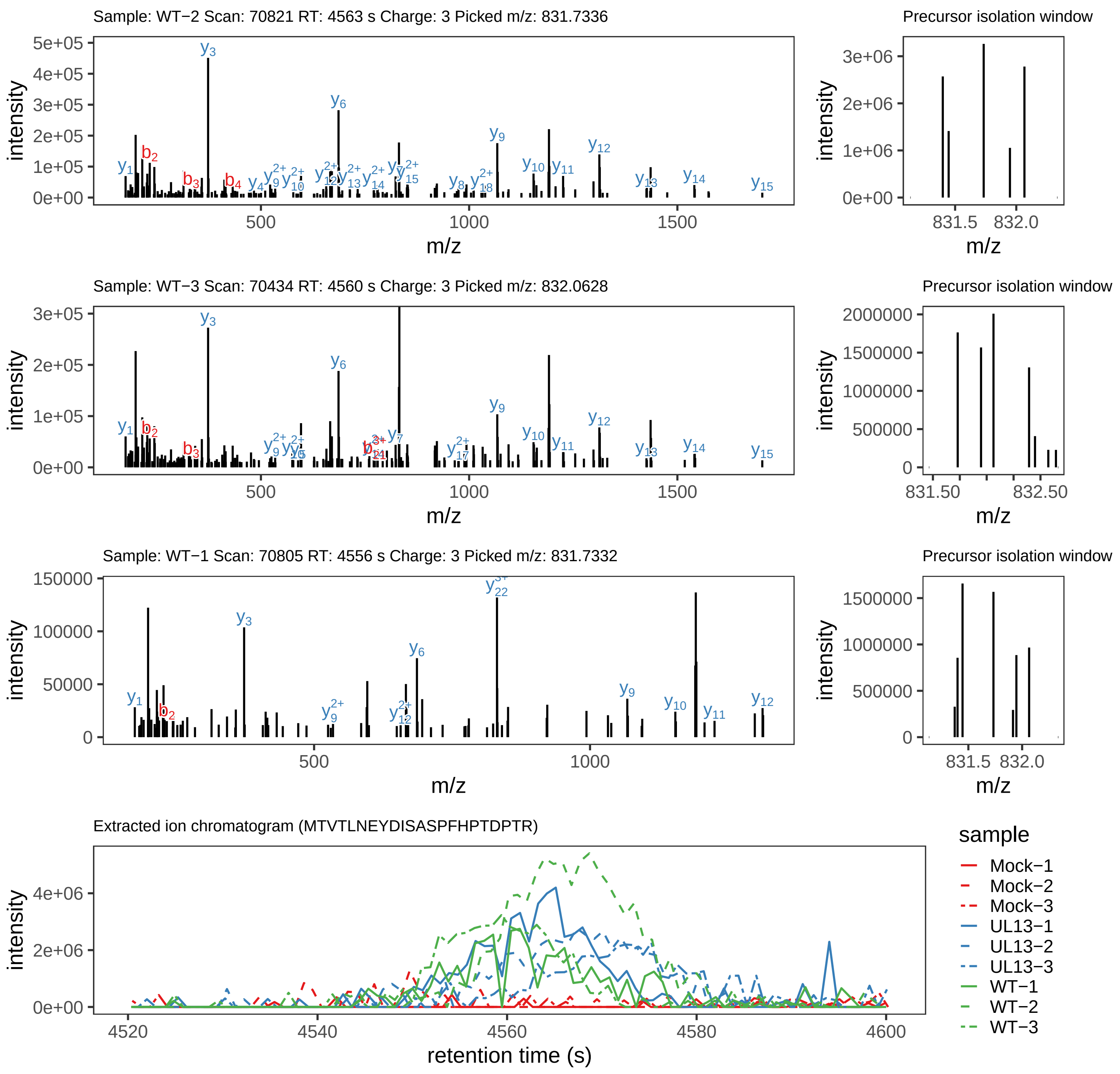

Supplement: Supplemental File 5 [file 737029_file08.zip › phosphopeptide_validation_figs_20240531/MTVTLNEYDISASPFHPTDPTR/MTVTLNEYDISASPFHPTDPTR.global.pdf]

### Peptide MTVTLNEY[+80]DISASPFHPTDPTR

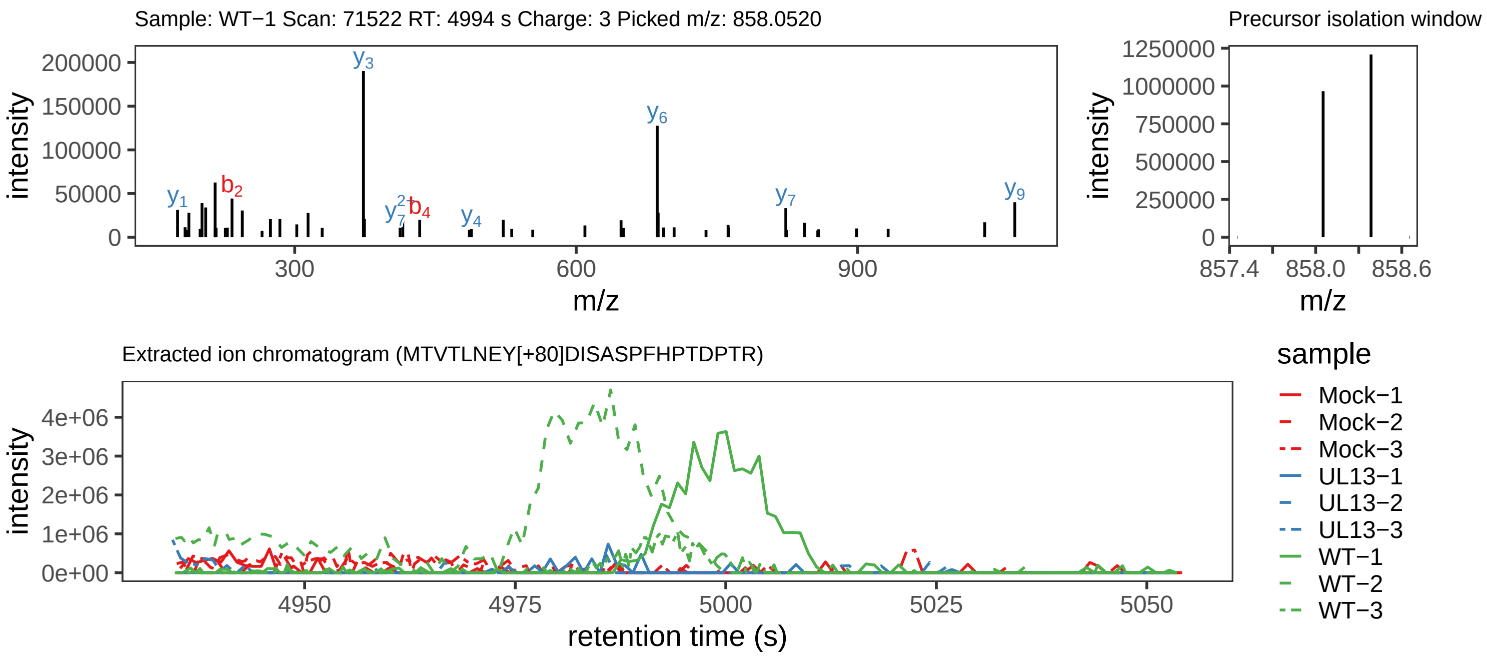

Supplement: Supplemental File 5 [file 737029_file08.zip › phosphopeptide_validation_figs_20240531/MTVTLNEYDISASPFHPTDPTR/MTVTLNEY[+80]DISASPFHPTDPTR.phospho.pdf]

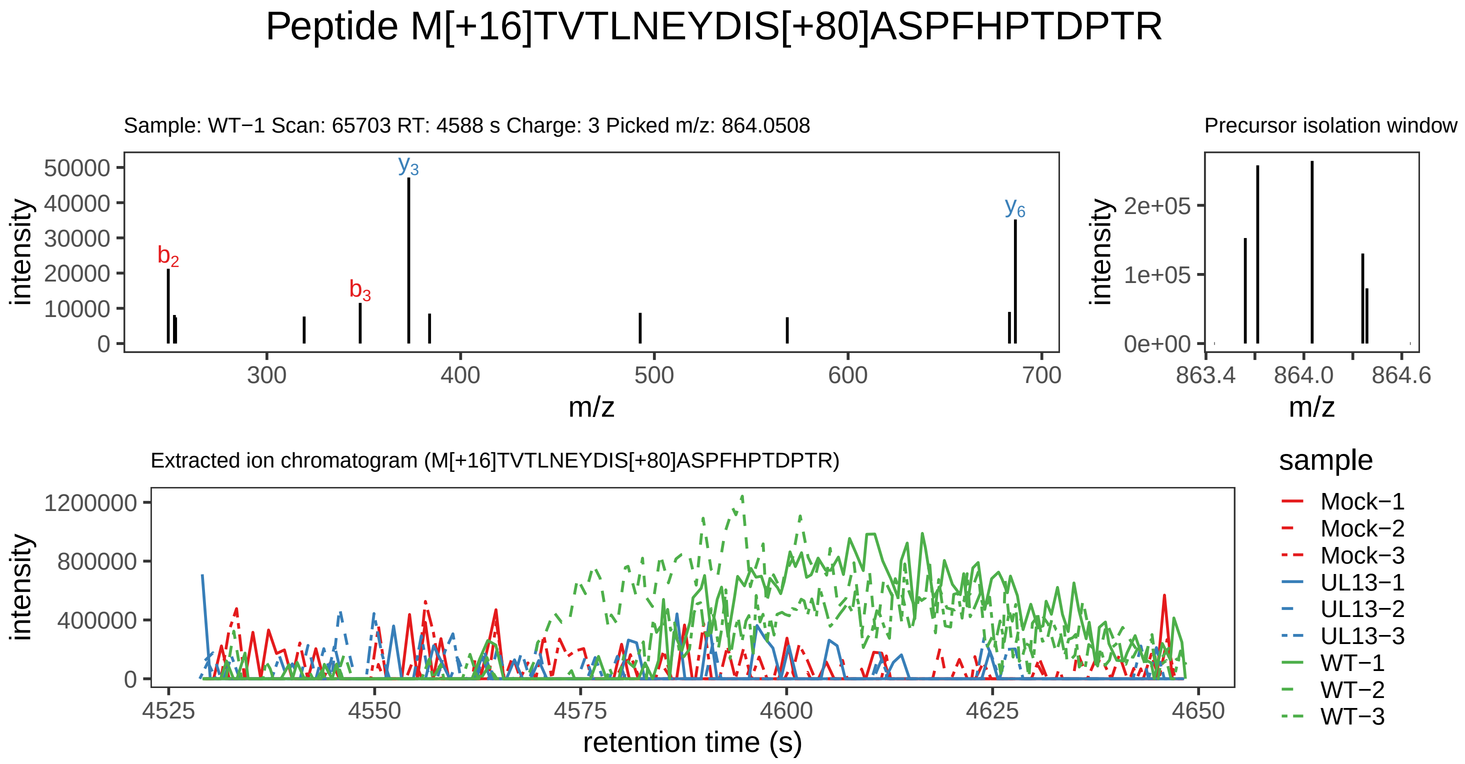

Supplement: Supplemental File 5 [file 737029_file08.zip › phosphopeptide_validation_figs_20240531/MTVTLNEYDISASPFHPTDPTR/M[+16]TVTLNEYDIS[+80]ASPFHPTDPTR.phospho.pdf]

### Peptide MTVTLNEYDIS[+80]ASPFHPTDPTR

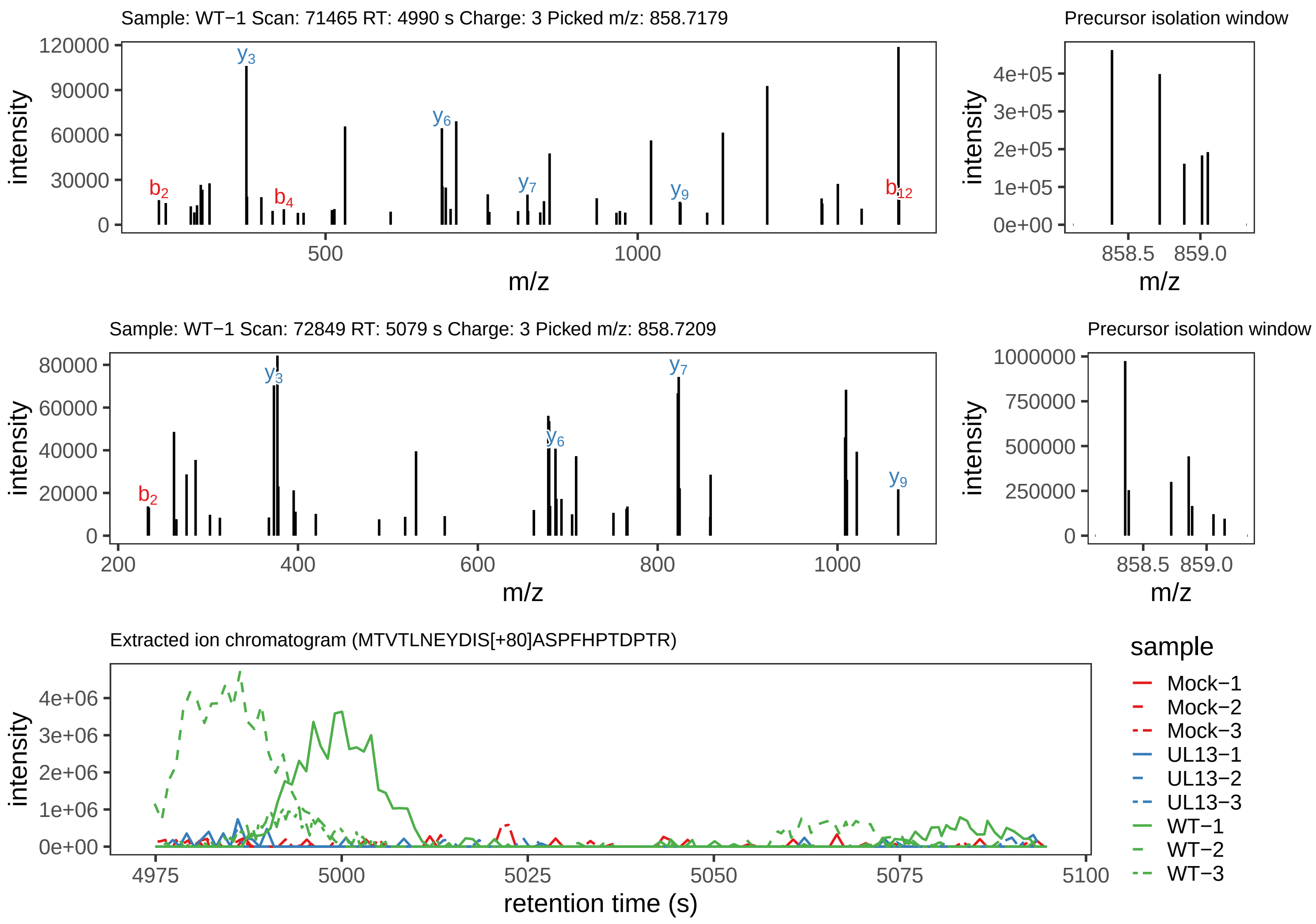

Supplement: Supplemental File 5 [file 737029_file08.zip › phosphopeptide_validation_figs_20240531/MTVTLNEYDISASPFHPTDPTR/MTVTLNEYDIS[+80]ASPFHPTDPTR.phospho.pdf]

### Peptide M[+16]TVTLNEY[+80]DISASPFHPTDPTR

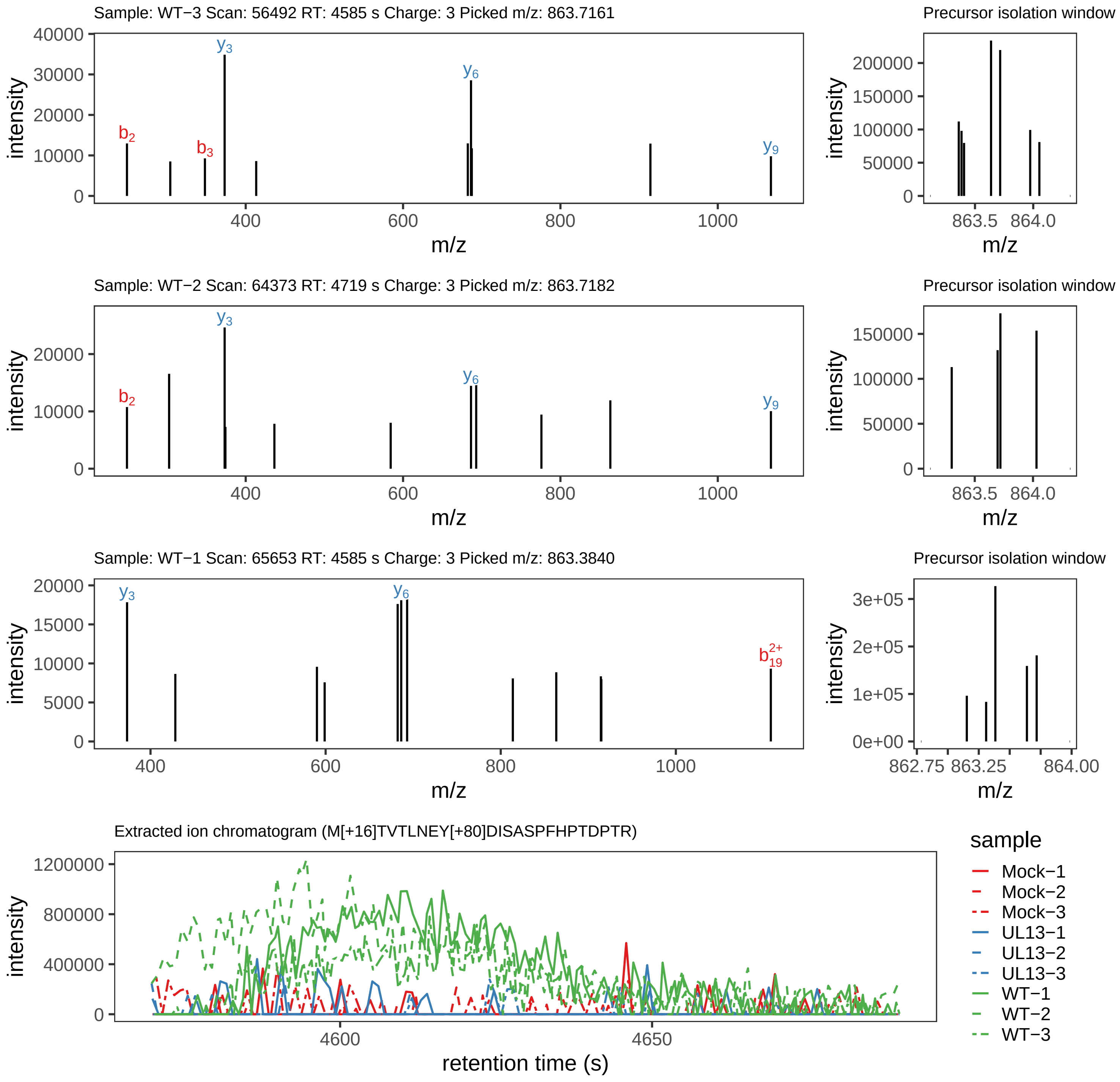

Supplement: Supplemental File 5 [file 737029_file08.zip › phosphopeptide_validation_figs_20240531/MTVTLNEYDISASPFHPTDPTR/M[+16]TVTLNEY[+80]DISASPFHPTDPTR.phospho.pdf]

### Peptide TLSADVINHIPLLR

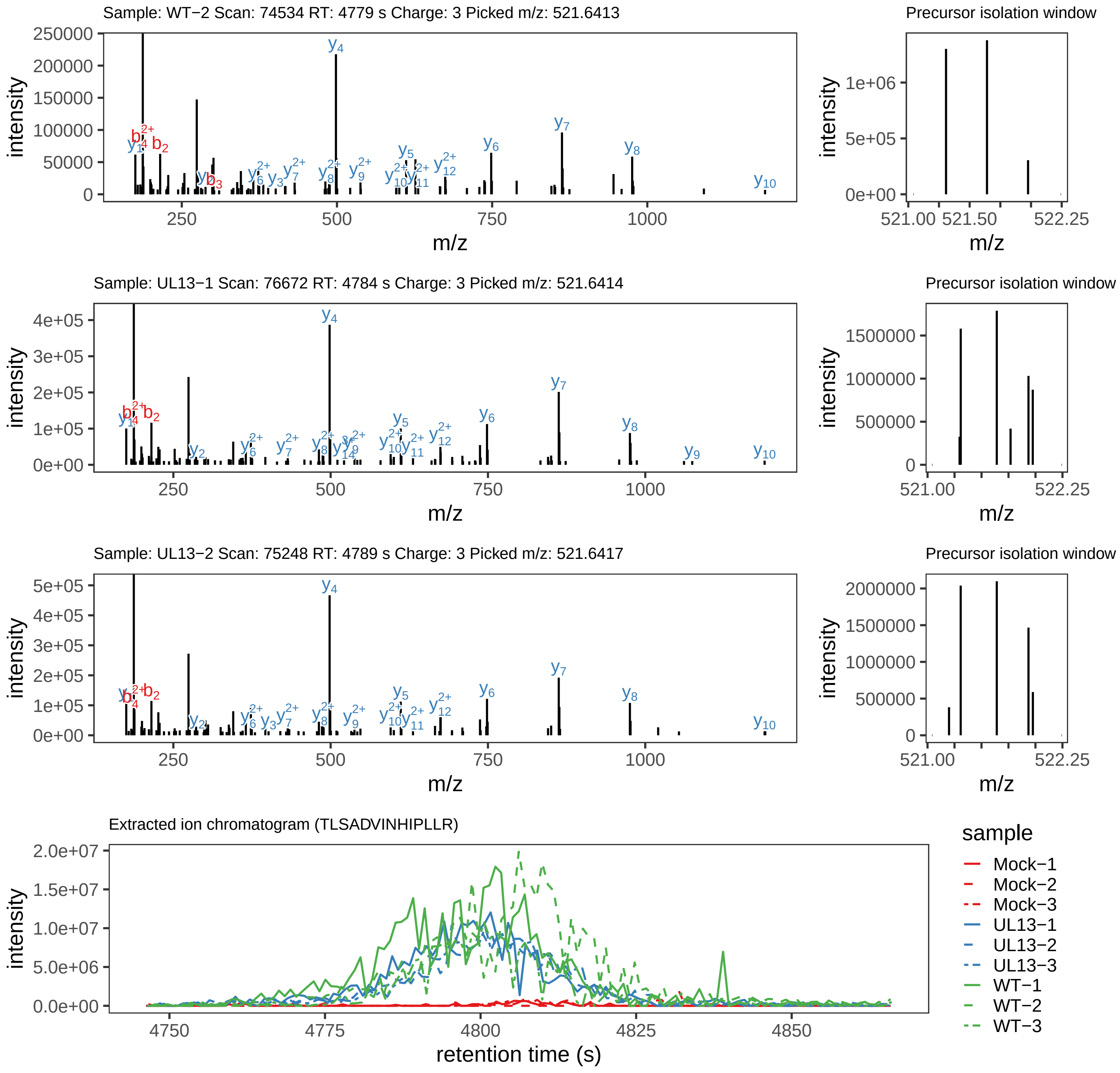

Supplement: Supplemental File 5 [file 737029_file08.zip › phosphopeptide_validation_figs_20240531/TLSADVINHIPLLR/TLSADVINHIPLLR.global.pdf]

### Peptide TLS[+80]ADVINHIPLLR

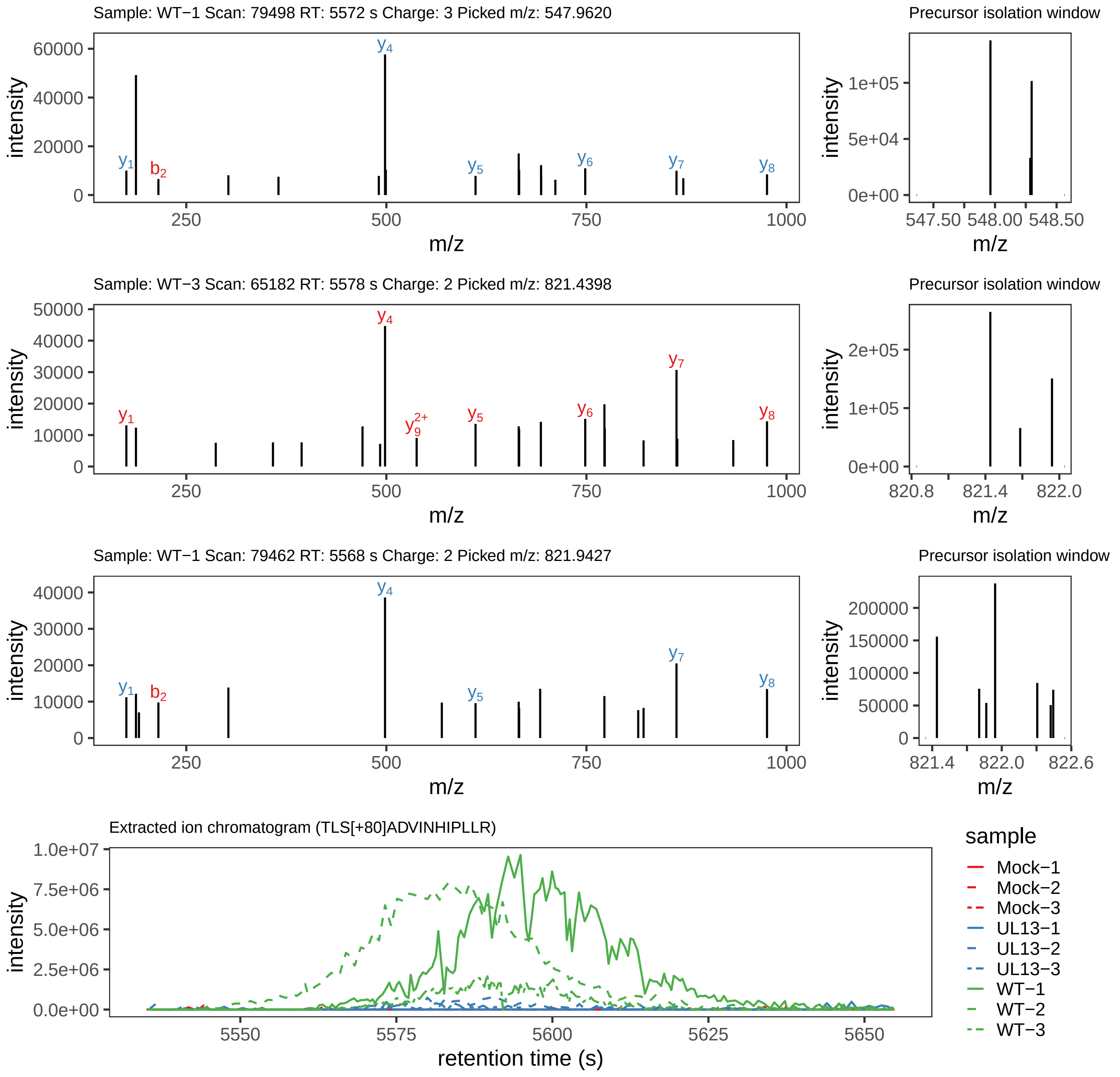

Supplement: Supplemental File 5 [file 737029_file08.zip › phosphopeptide_validation_figs_20240531/TLSADVINHIPLLR/TLS[+80]ADVINHIPLLR.phospho.pdf]

### Peptide IAGLYGVPGSDYAYPRQS[+80]ELISSIR

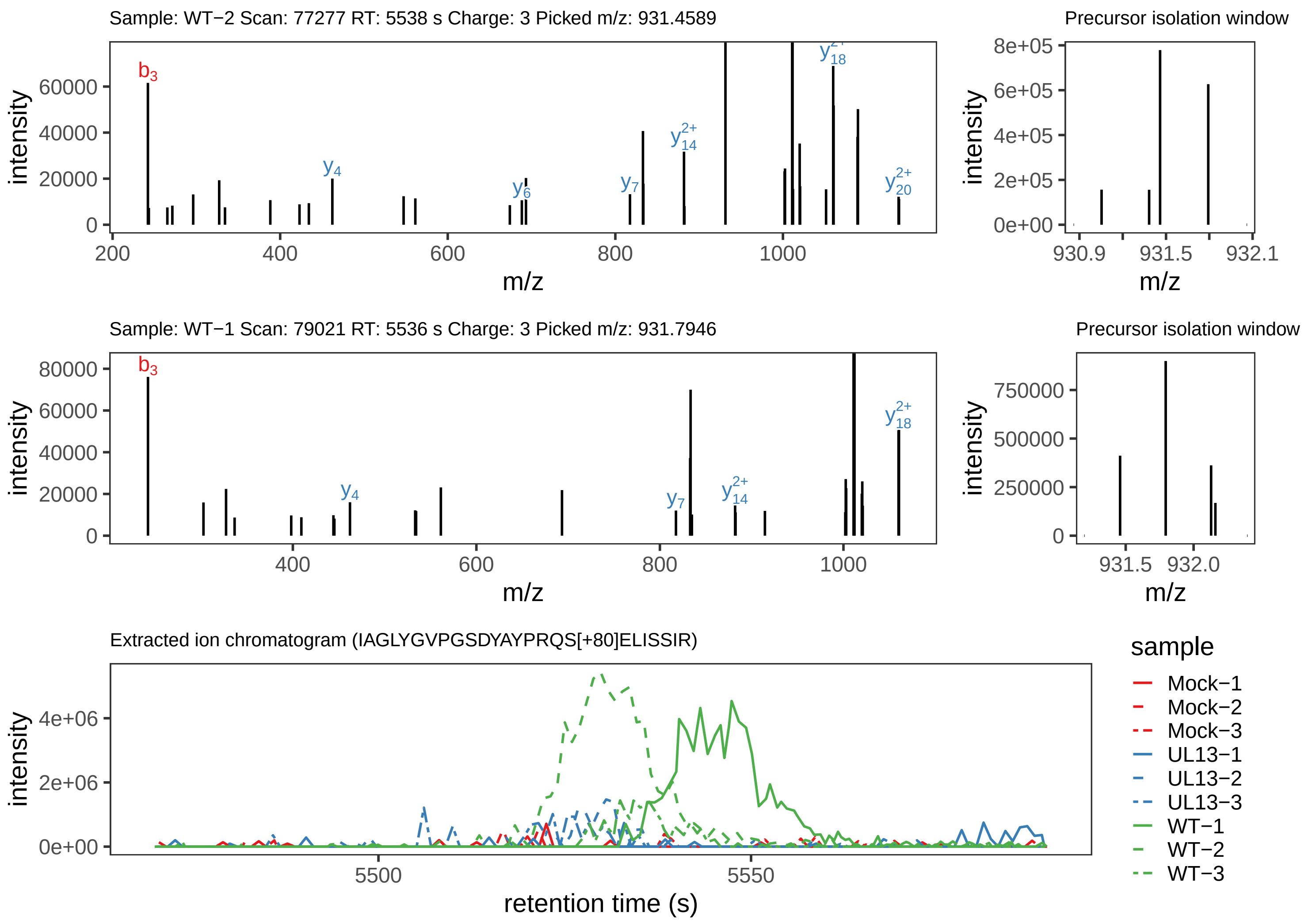

Supplement: Supplemental File 5 [file 737029_file08.zip › phosphopeptide_validation_figs_20240531/IAGLYGVPGSDYAYPRQSELISSIR/IAGLYGVPGSDYAYPRQS[+80]ELISSIR.phospho.pdf]

### Peptide IAGLYGVPGSDY[+80]AYPRQSELISSIR

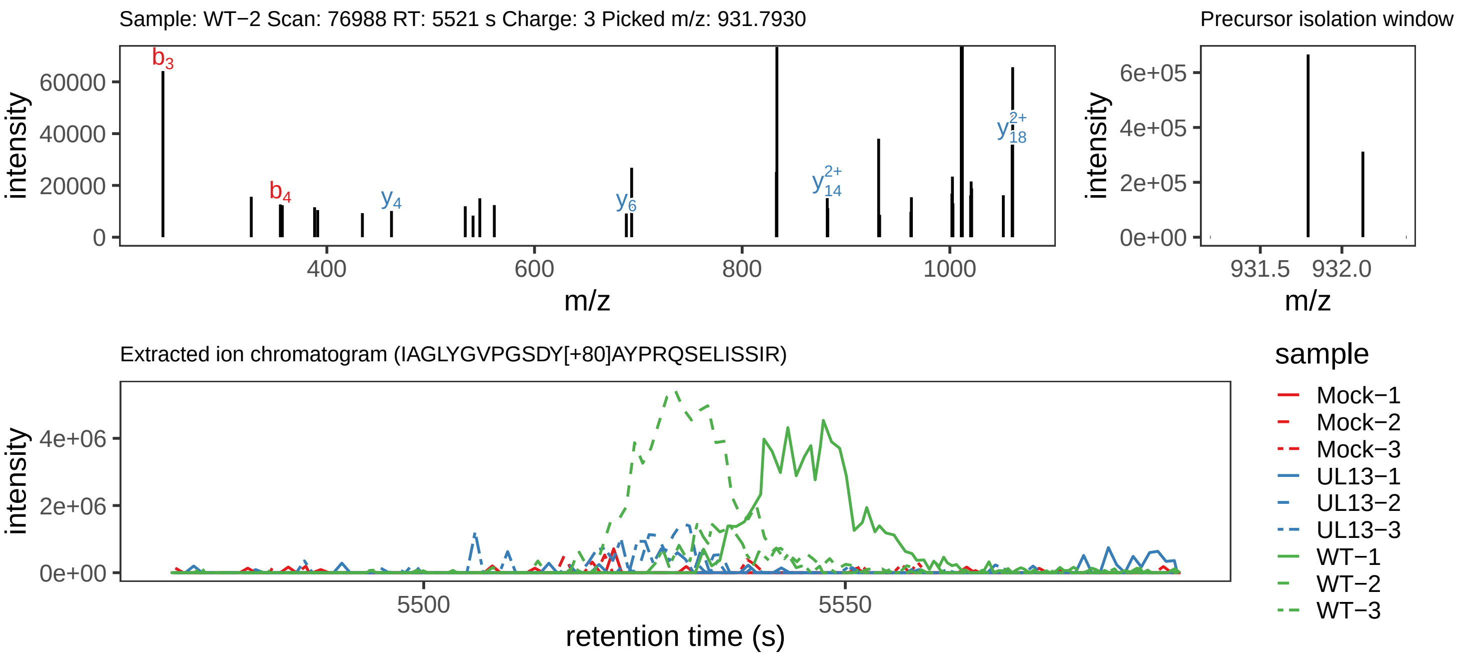

Supplement: Supplemental File 5 [file 737029_file08.zip › phosphopeptide_validation_figs_20240531/IAGLYGVPGSDYAYPRQSELISSIR/IAGLYGVPGSDY[+80]AYPRQSELISSIR.phospho.pdf]

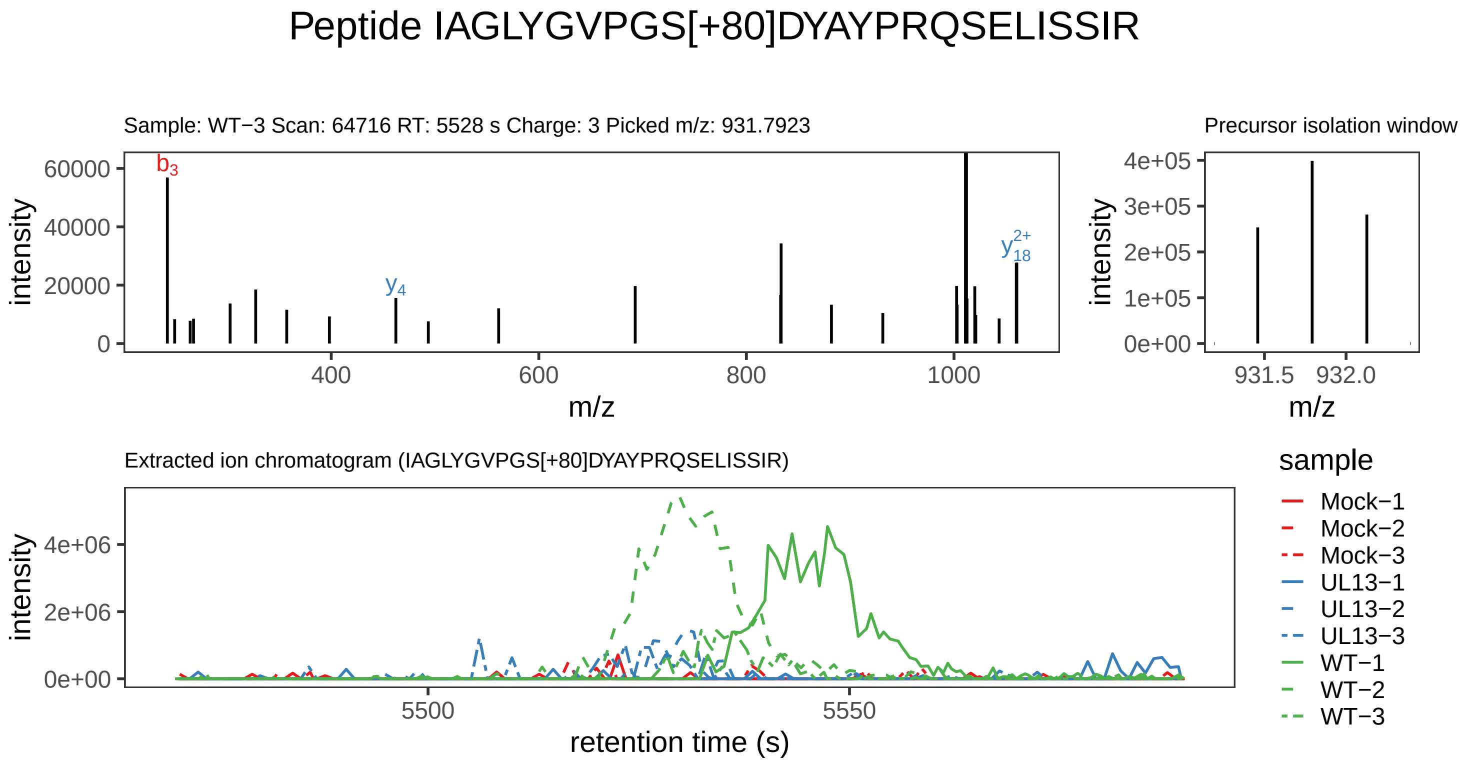

Supplement: Supplemental File 5 [file 737029_file08.zip › phosphopeptide_validation_figs_20240531/IAGLYGVPGSDYAYPRQSELISSIR/IAGLYGVPGS[+80]DYAYPRQSELISSIR.phospho.pdf]

### Peptide RRFS[+80]DNIR

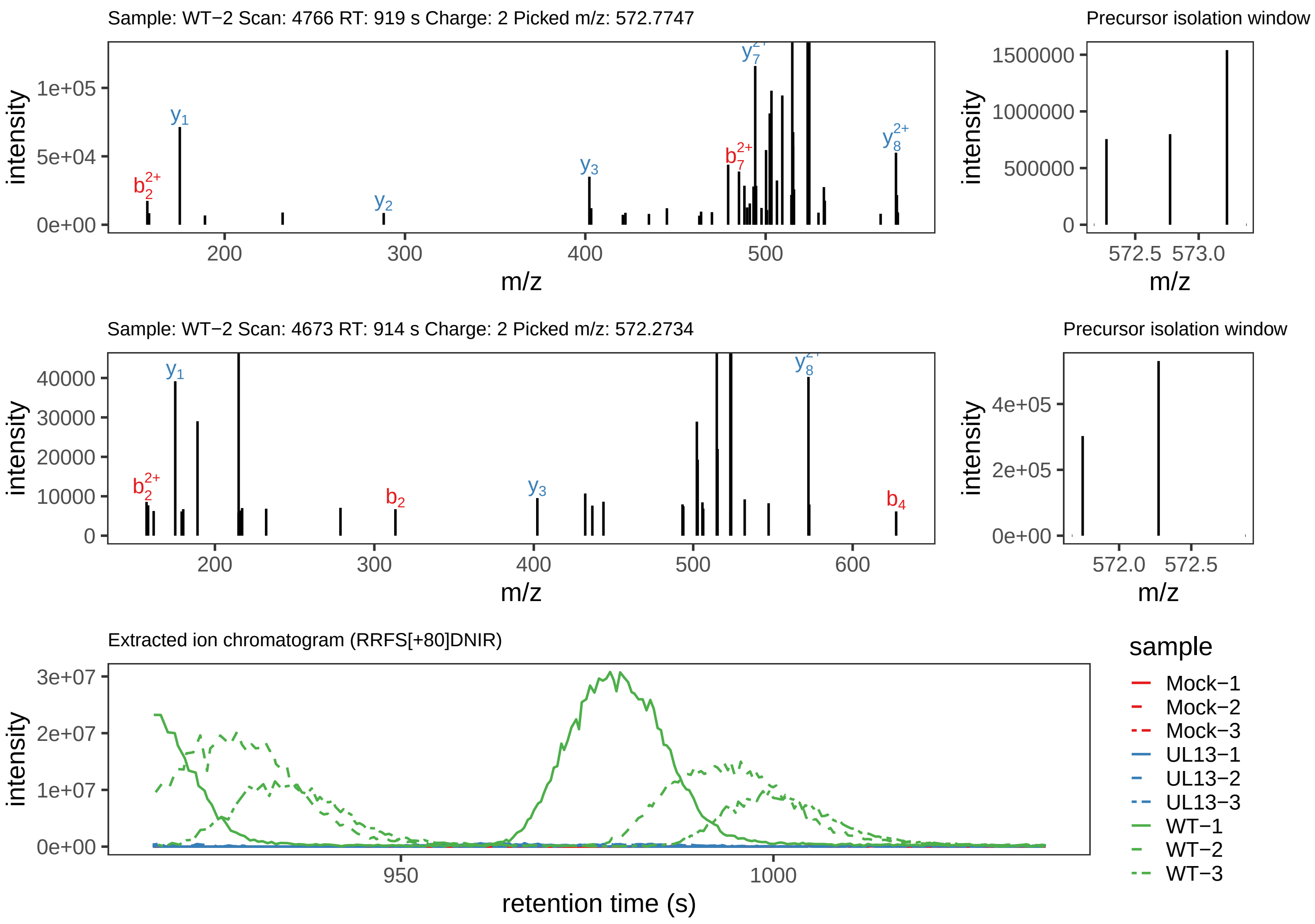

Supplement: Supplemental File 5 [file 737029_file08.zip › phosphopeptide_validation_figs_20240531/RRFSDNIR/RRFS[+80]DNIR.phospho.pdf]

### Peptide NQEPLDSLCR

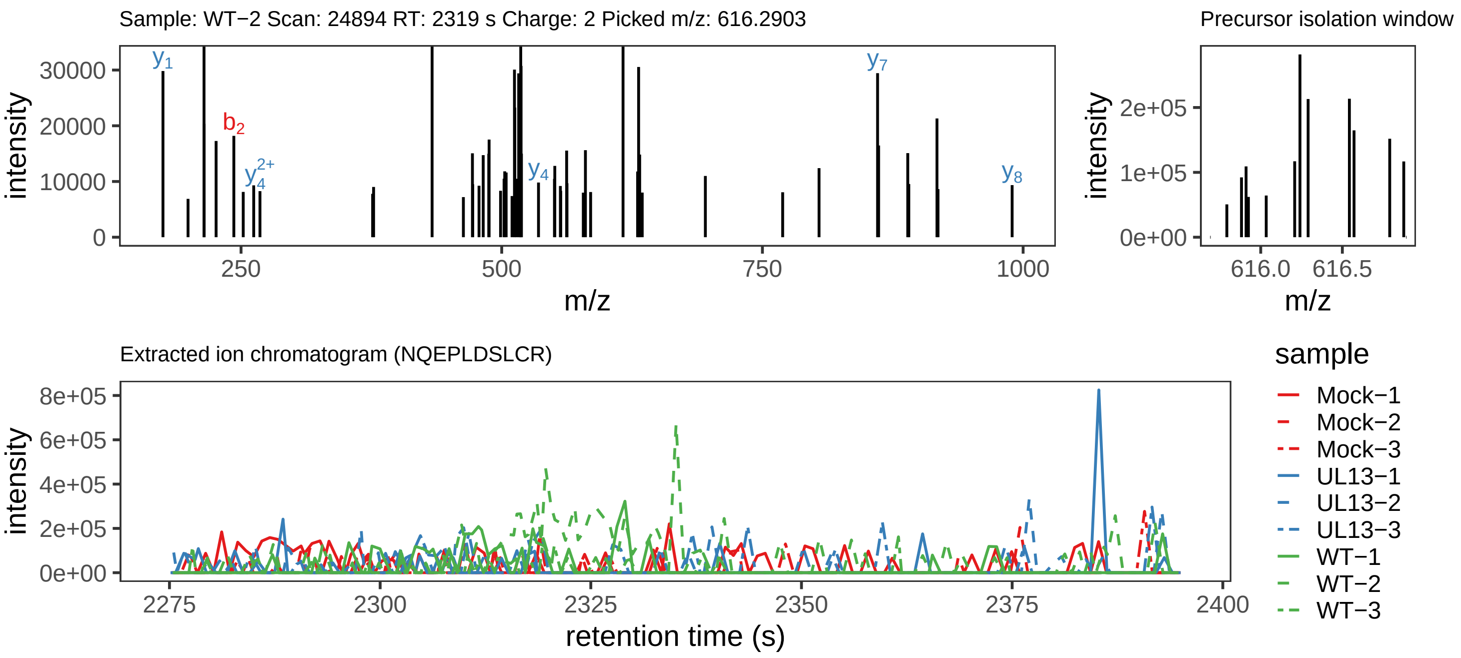

Supplement: Supplemental File 5 [file 737029_file08.zip › phosphopeptide_validation_figs_20240531/NQEPLDSLCR/NQEPLDSLCR.phospho.pdf]

### Peptide N[+1]Q[+1]EPLDSLCR

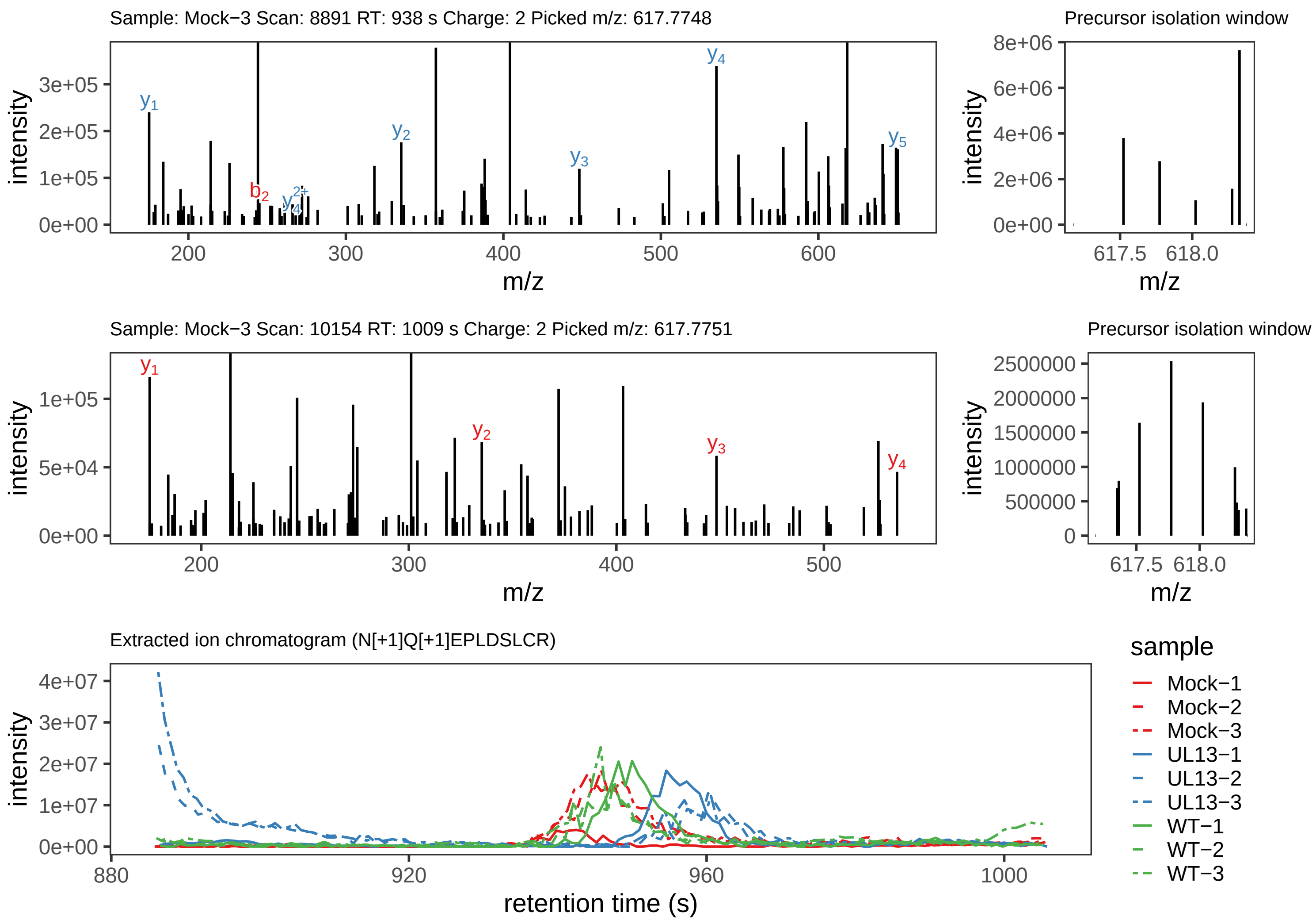

Supplement: Supplemental File 5 [file 737029_file08.zip › phosphopeptide_validation_figs_20240531/NQEPLDSLCR/N[+1]Q[+1]EPLDSLCR.global.pdf]

### Peptide NQEPLDS[+80]LCR

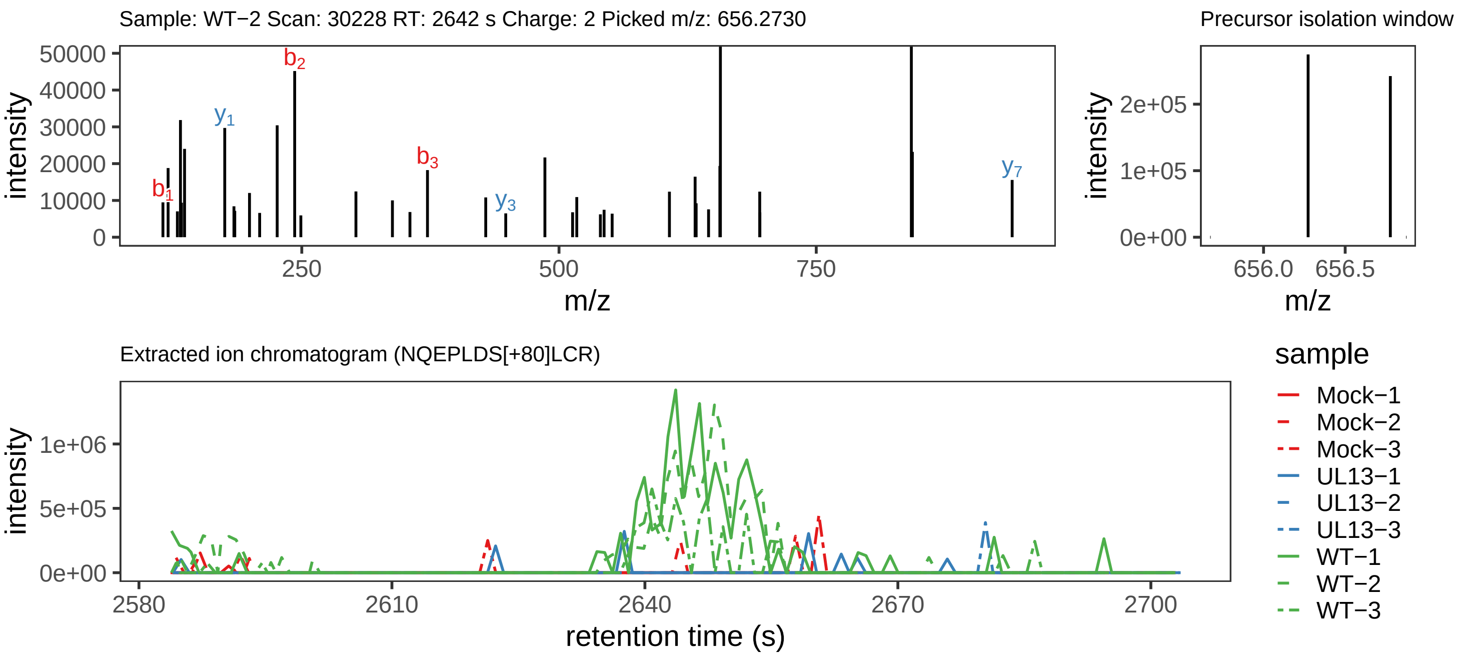

Supplement: Supplemental File 5 [file 737029_file08.zip › phosphopeptide_validation_figs_20240531/NQEPLDSLCR/NQEPLDS[+80]LCR.phospho.pdf]

### Peptide NQEPLDSLCR

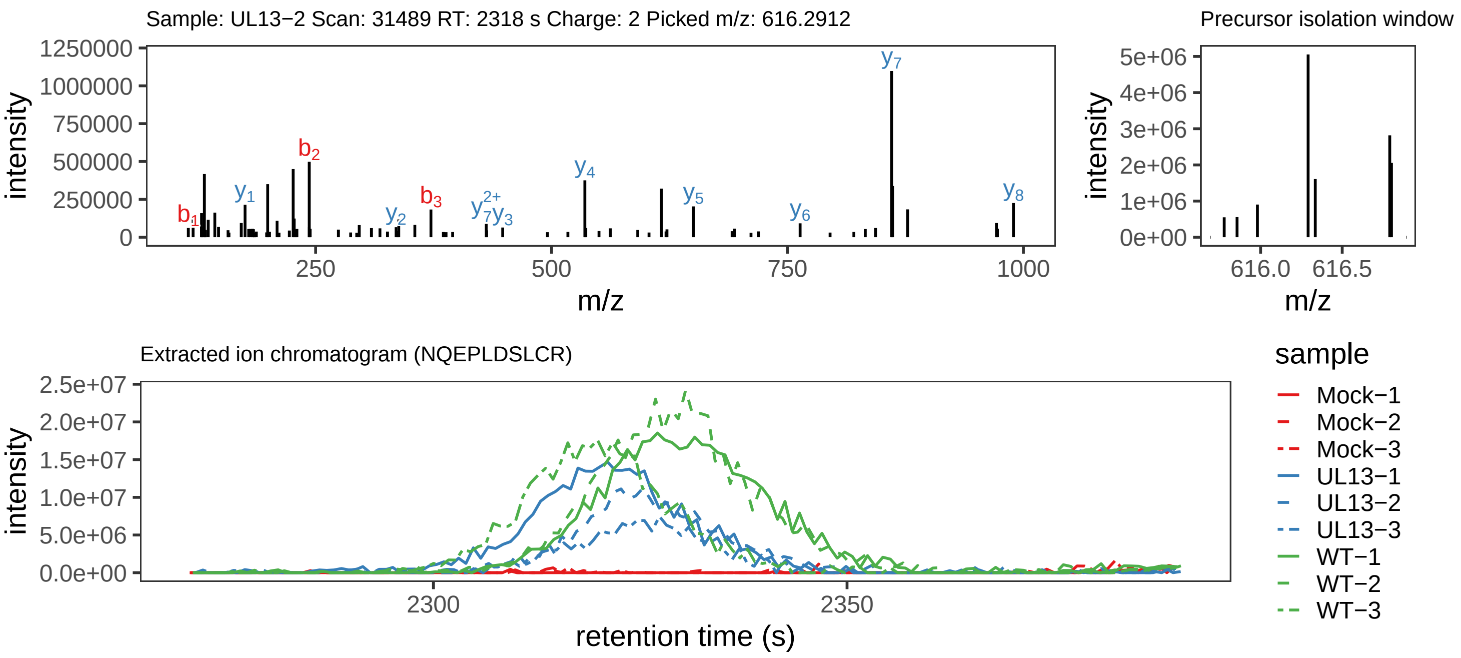

Supplement: Supplemental File 5 [file 737029_file08.zip › phosphopeptide_validation_figs_20240531/NQEPLDSLCR/NQEPLDSLCR.global.pdf]

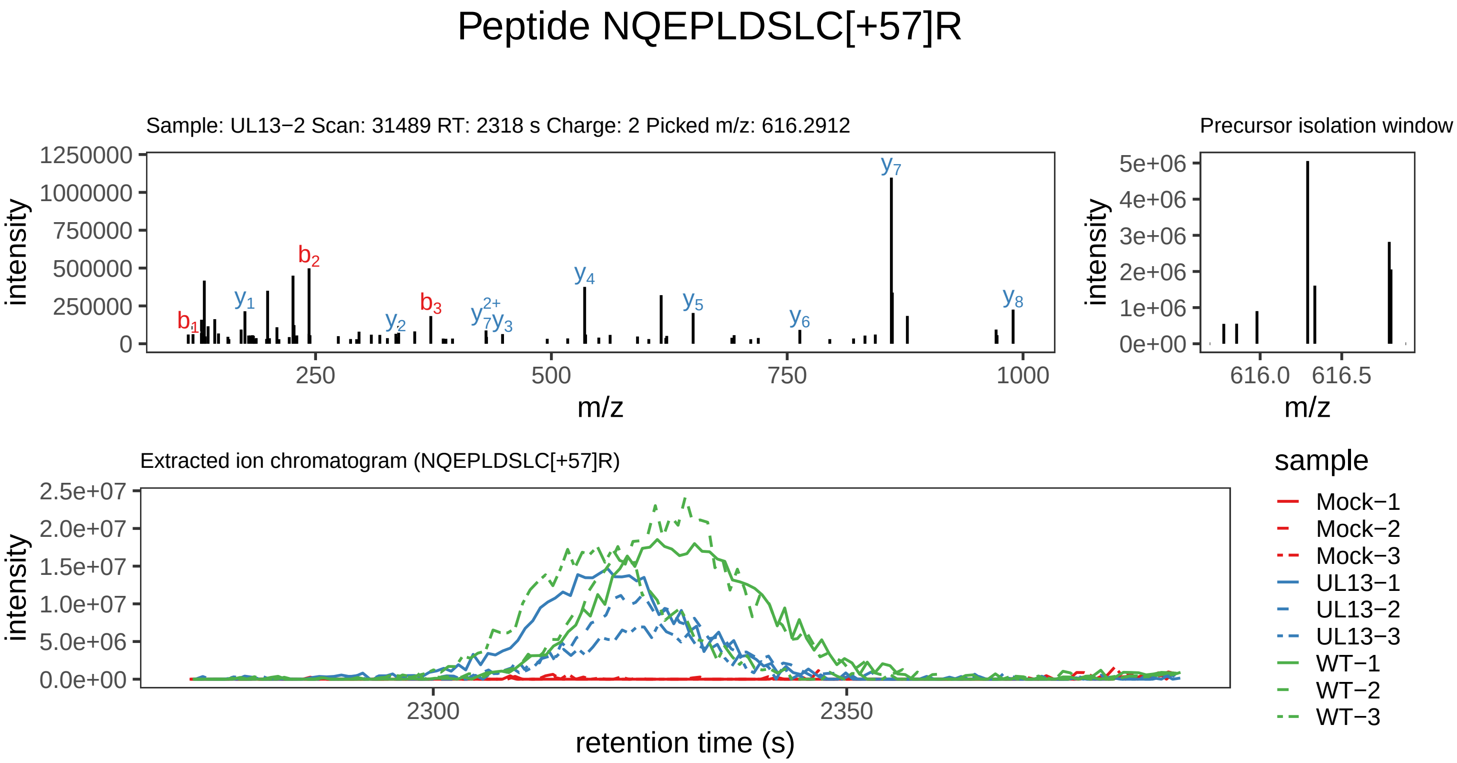

Supplement: Supplemental File 5 [file 737029_file08.zip › phosphopeptide_validation_figs_20240531/NQEPLDSLCR/NQEPLDSLC[+57]R.global.pdf]

### Peptide ASY[+80]GSLAYWPELR

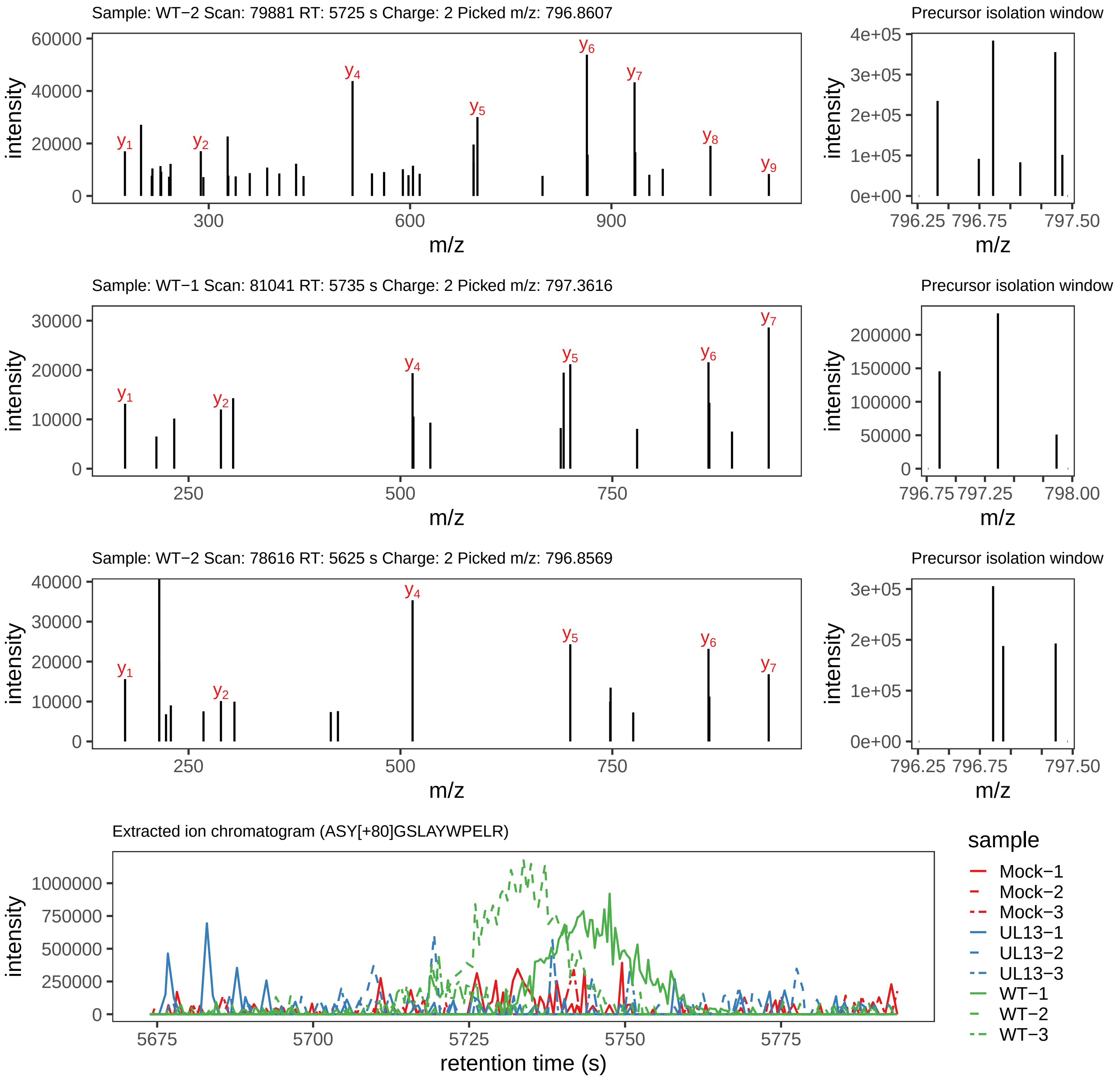

Supplement: Supplemental File 5 [file 737029_file08.zip › phosphopeptide_validation_figs_20240531/ASYGSLAYWPELR/ASY[+80]GSLAYWPELR.phospho.pdf]

### Peptide ASYGSLAYWPELR

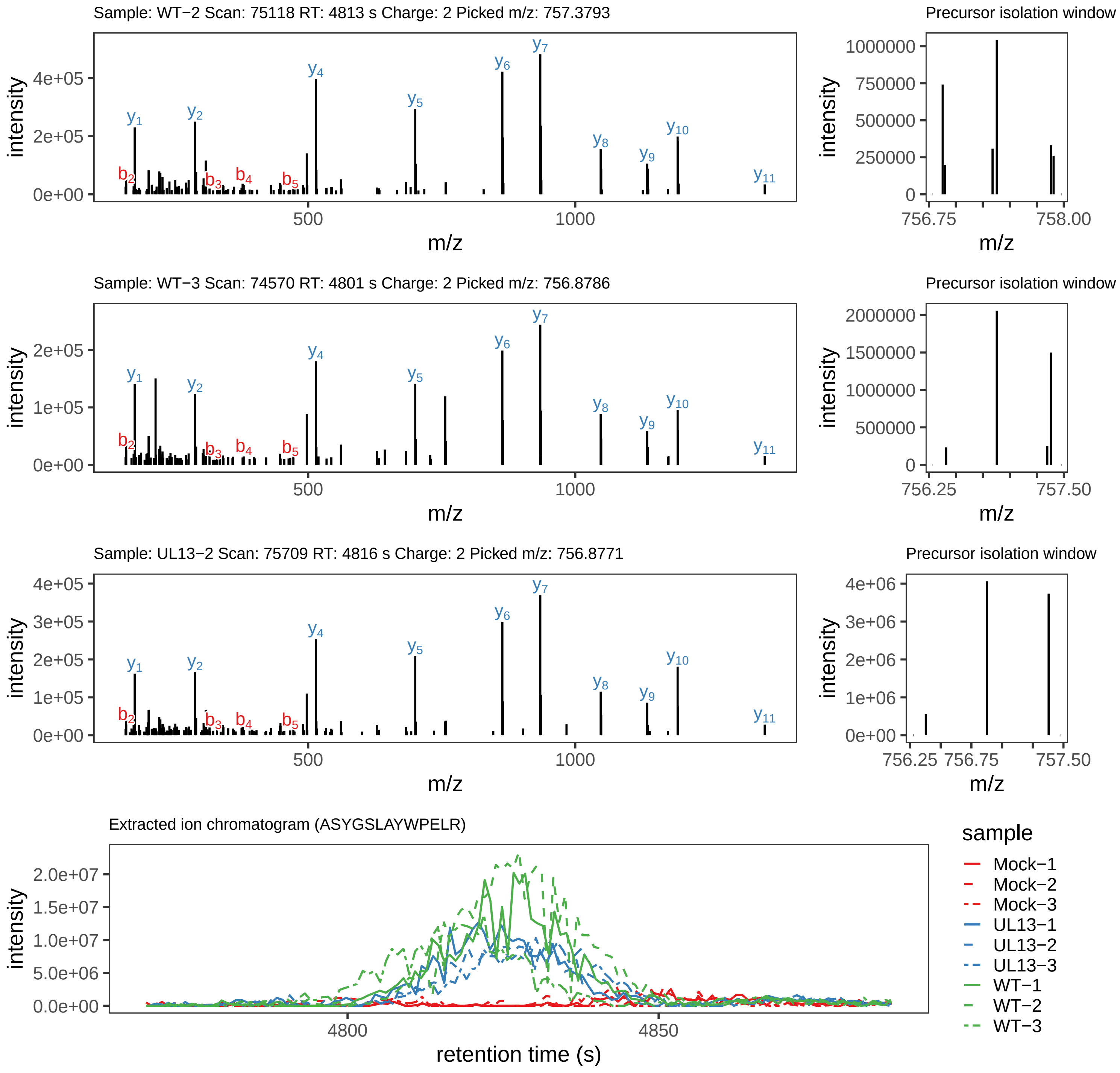

Supplement: Supplemental File 5 [file 737029_file08.zip › phosphopeptide_validation_figs_20240531/ASYGSLAYWPELR/ASYGSLAYWPELR.global.pdf]

### Peptide AS[+80]YGSLAYWPELR

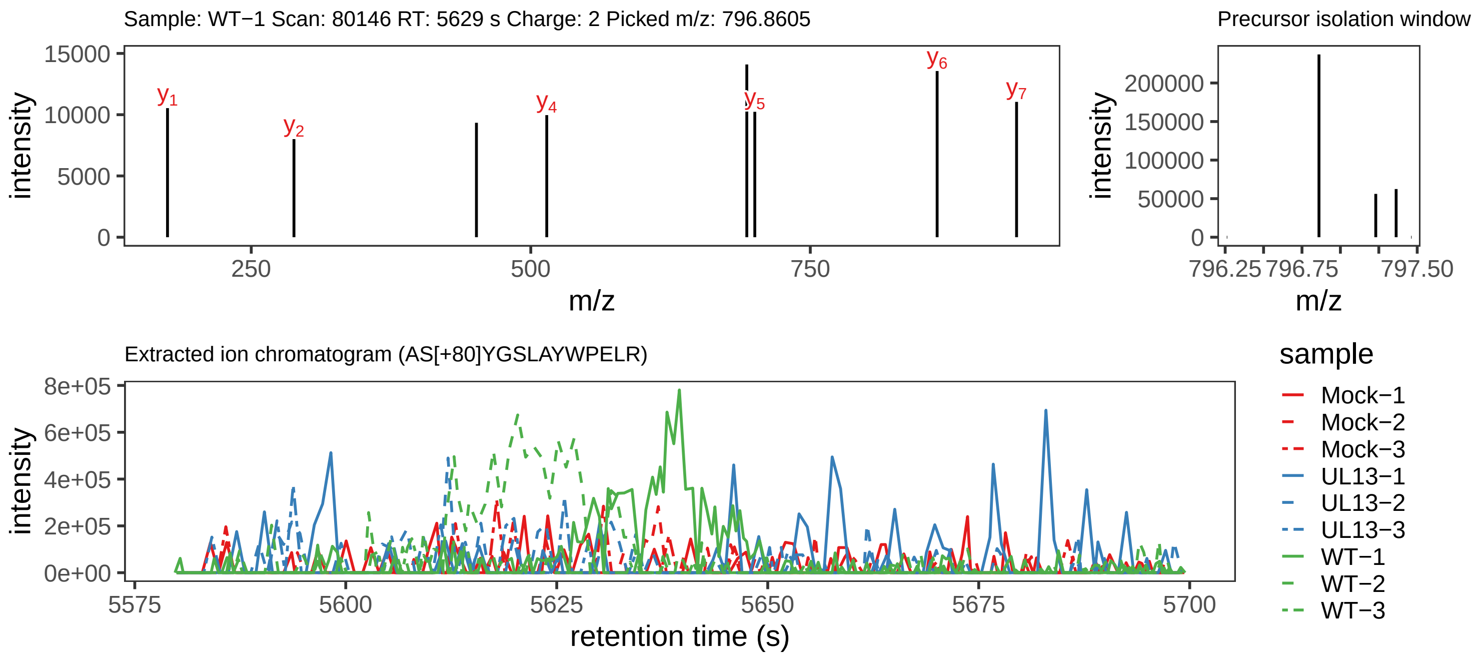

Supplement: Supplemental File 5 [file 737029_file08.zip › phosphopeptide_validation_figs_20240531/ASYGSLAYWPELR/AS[+80]YGSLAYWPELR.phospho.pdf]

### Peptide LLDS[+80]AIAAYDAK

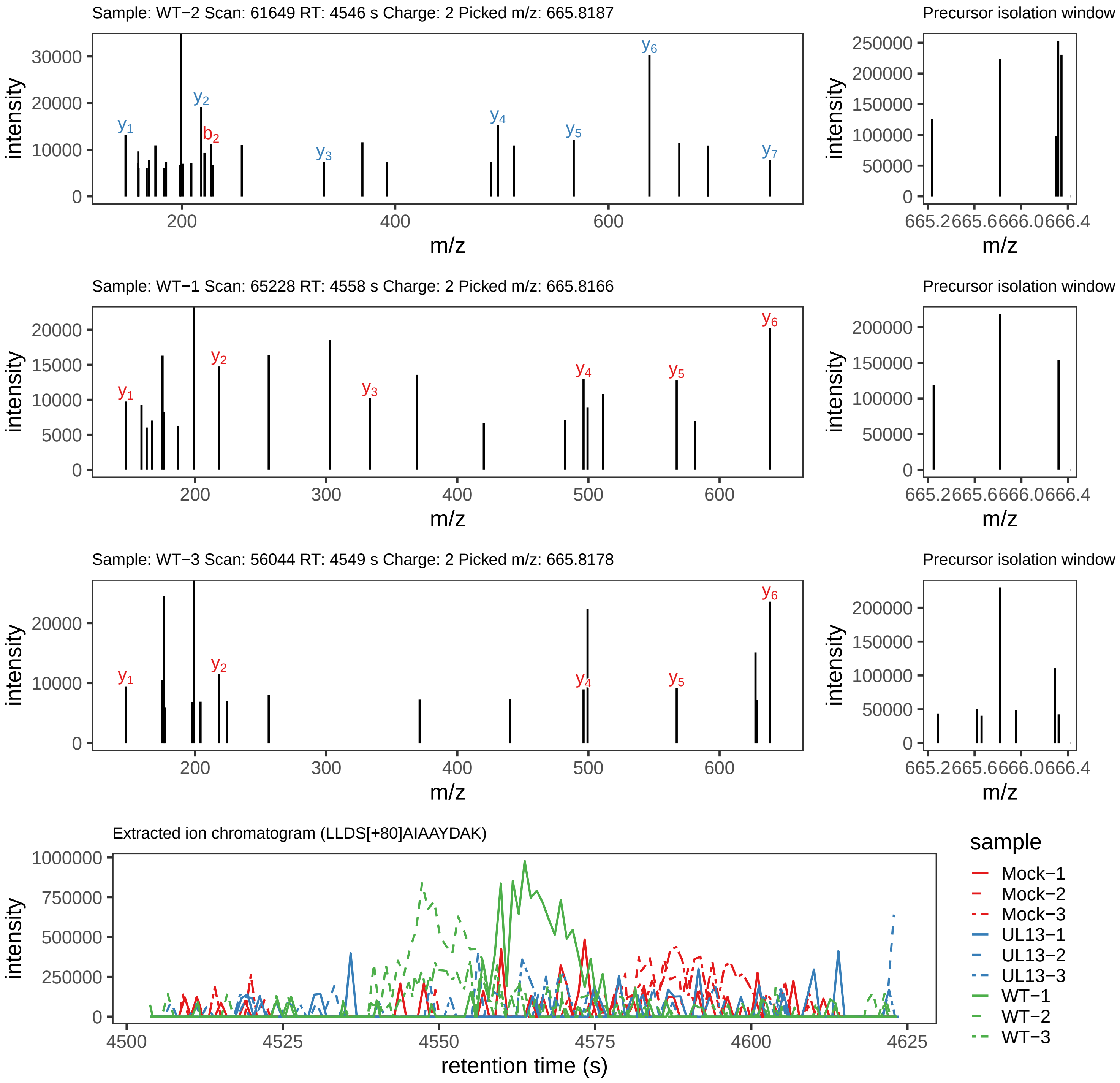

Supplement: Supplemental File 5 [file 737029_file08.zip › phosphopeptide_validation_figs_20240531/LLDSAIAAYDAK/LLDS[+80]AIAAYDAK.phospho.pdf]

### Peptide LLDSAIAAYDAK

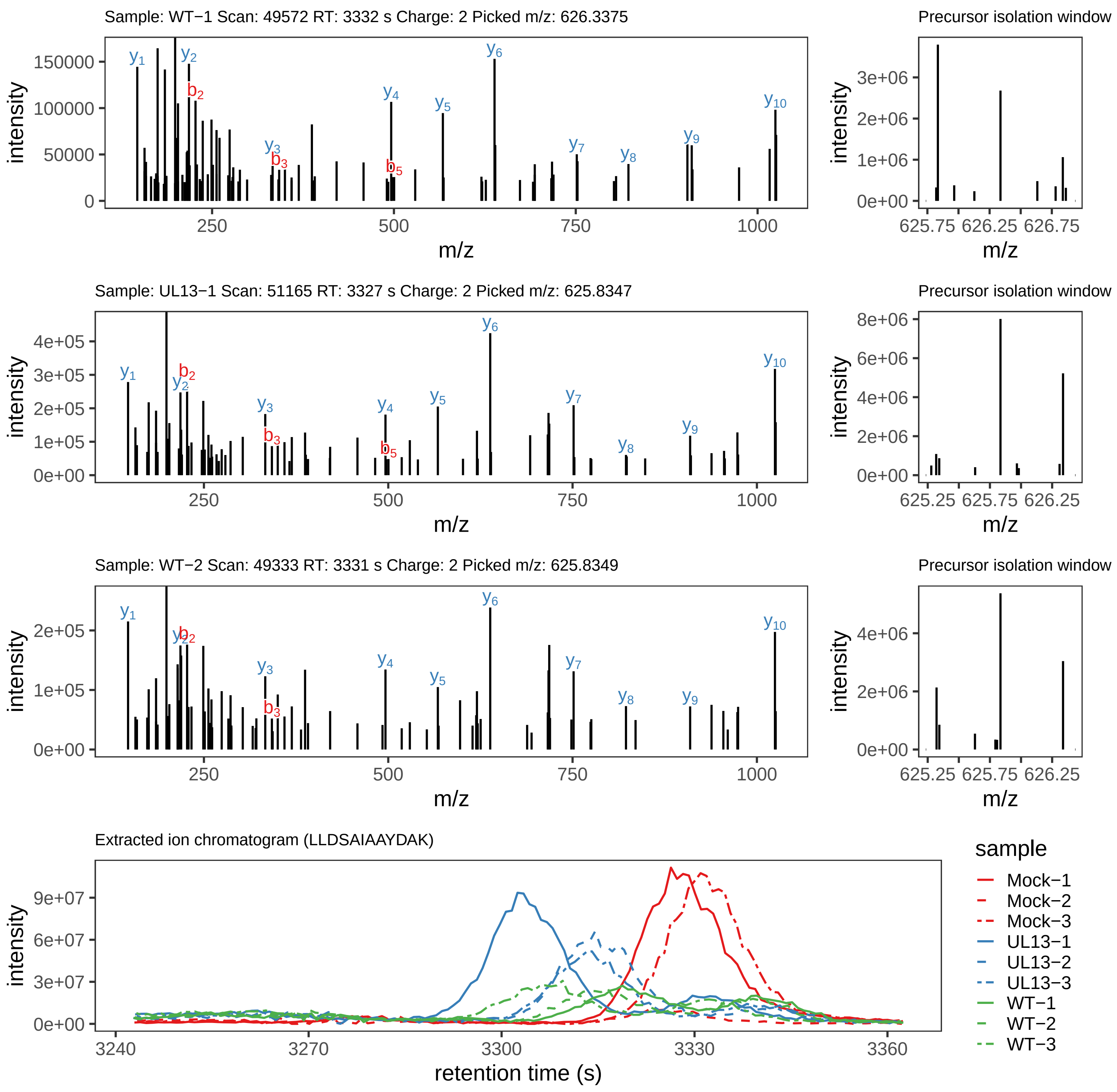

Supplement: Supplemental File 5 [file 737029_file08.zip › phosphopeptide_validation_figs_20240531/LLDSAIAAYDAK/LLDSAIAAYDAK.global.pdf]

### Peptide FTTAGSRTS[+80]IQMYR

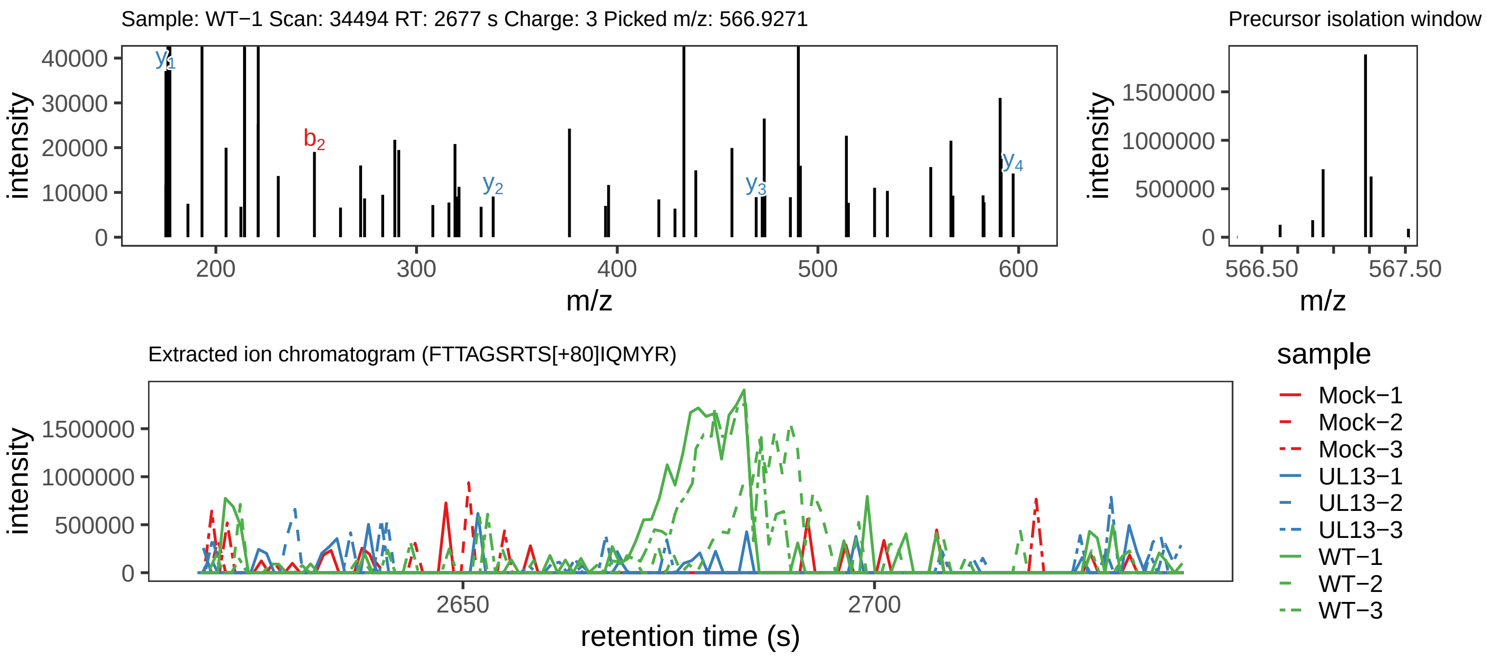

Supplement: Supplemental File 5 [file 737029_file08.zip › phosphopeptide_validation_figs_20240531/FTTAGSRTSIQMYR/FTTAGSRTS[+80]IQMYR.phospho.pdf]

### Peptide ATS[+80]IYTTPSALYTR

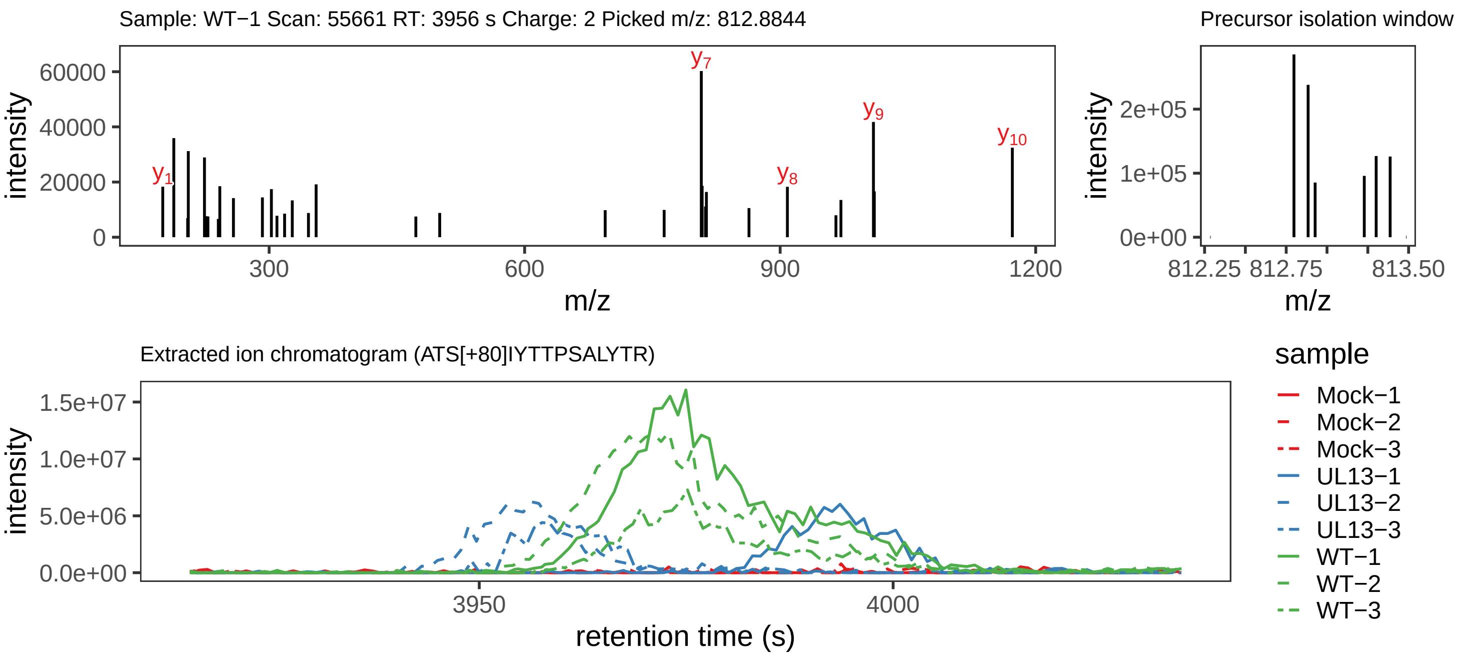

Supplement: Supplemental File 5 [file 737029_file08.zip › phosphopeptide_validation_figs_20240531/ATSIYTTPSALYTR/ATS[+80]IYTTPSALYTR.phospho.pdf]

### Peptide ATSIYTTPSALYTR

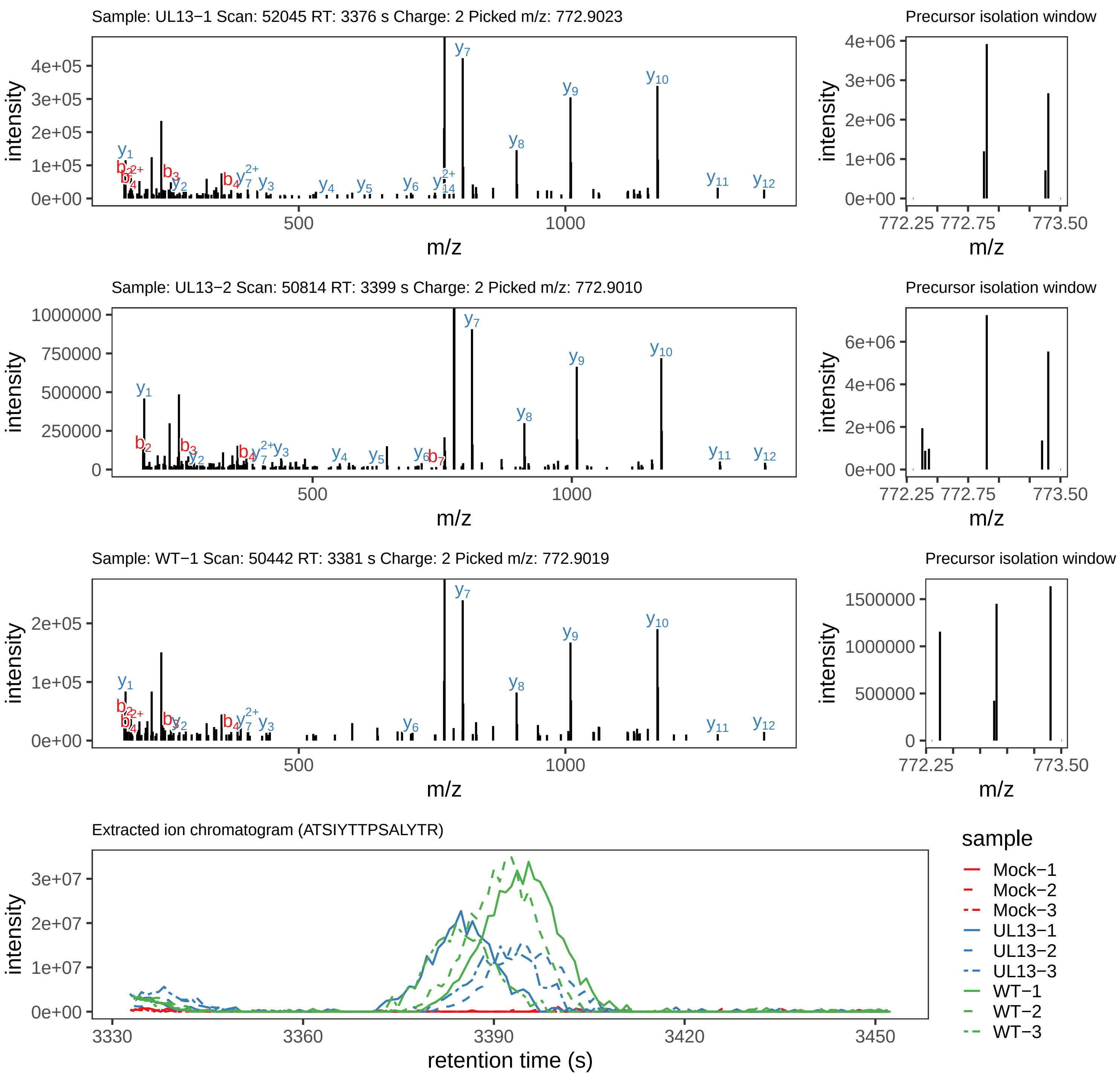

Supplement: Supplemental File 5 [file 737029_file08.zip › phosphopeptide_validation_figs_20240531/ATSIYTTPSALYTR/ATSIYTTPSALYTR.global.pdf]

### Peptide ATSIYT[+80]TPSALYTR

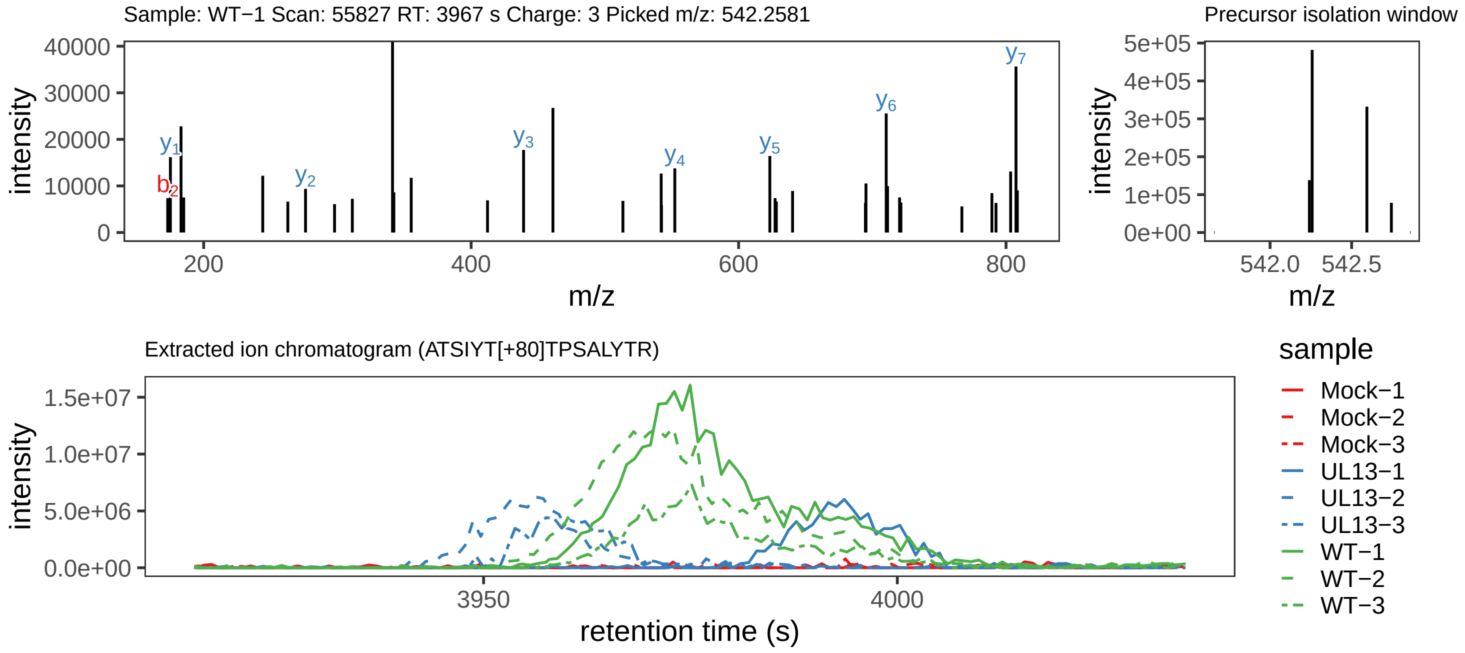

Supplement: Supplemental File 5 [file 737029_file08.zip › phosphopeptide_validation_figs_20240531/ATSIYTTPSALYTR/ATSIYT[+80]TPSALYTR.phospho.pdf]

### Peptide AT[+80]SIYTTPSALYTR

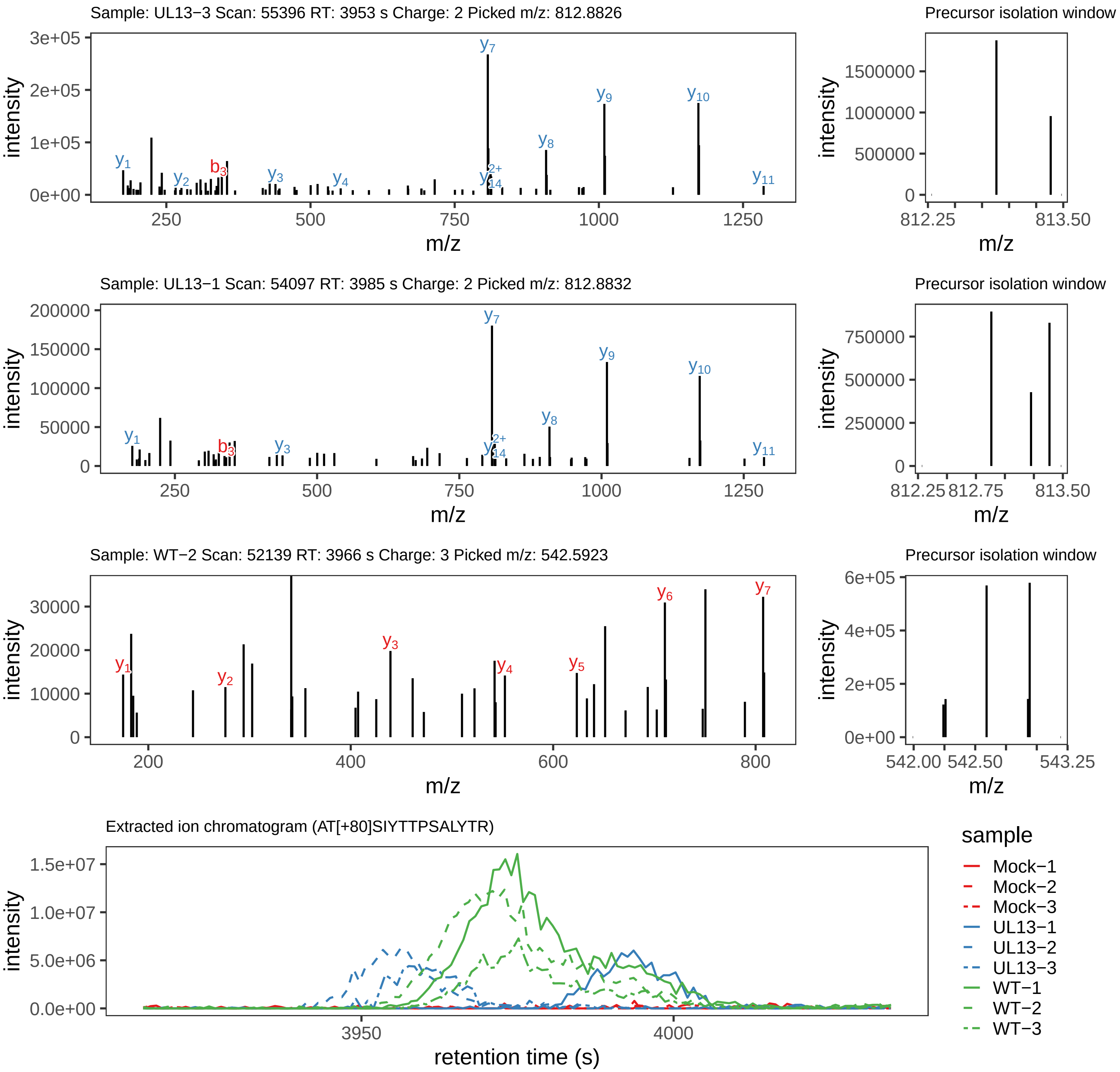

Supplement: Supplemental File 5 [file 737029_file08.zip › phosphopeptide_validation_figs_20240531/ATSIYTTPSALYTR/AT[+80]SIYTTPSALYTR.phospho.pdf]

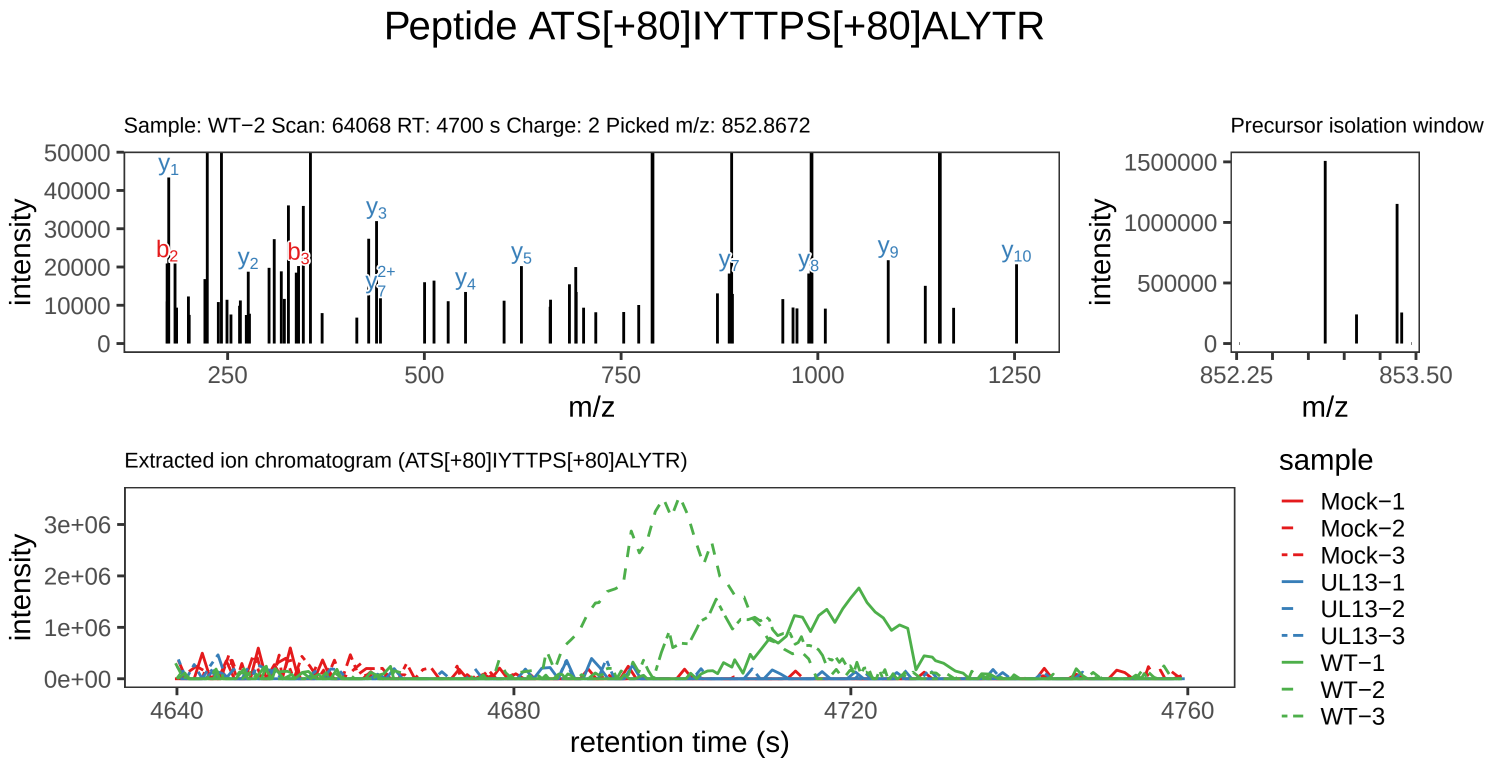

Supplement: Supplemental File 5 [file 737029_file08.zip › phosphopeptide_validation_figs_20240531/ATSIYTTPSALYTR/ATS[+80]IYTTPS[+80]ALYTR.phospho.pdf]

### Peptide [+42]MSPT[+80]PEDDRDLVVVR

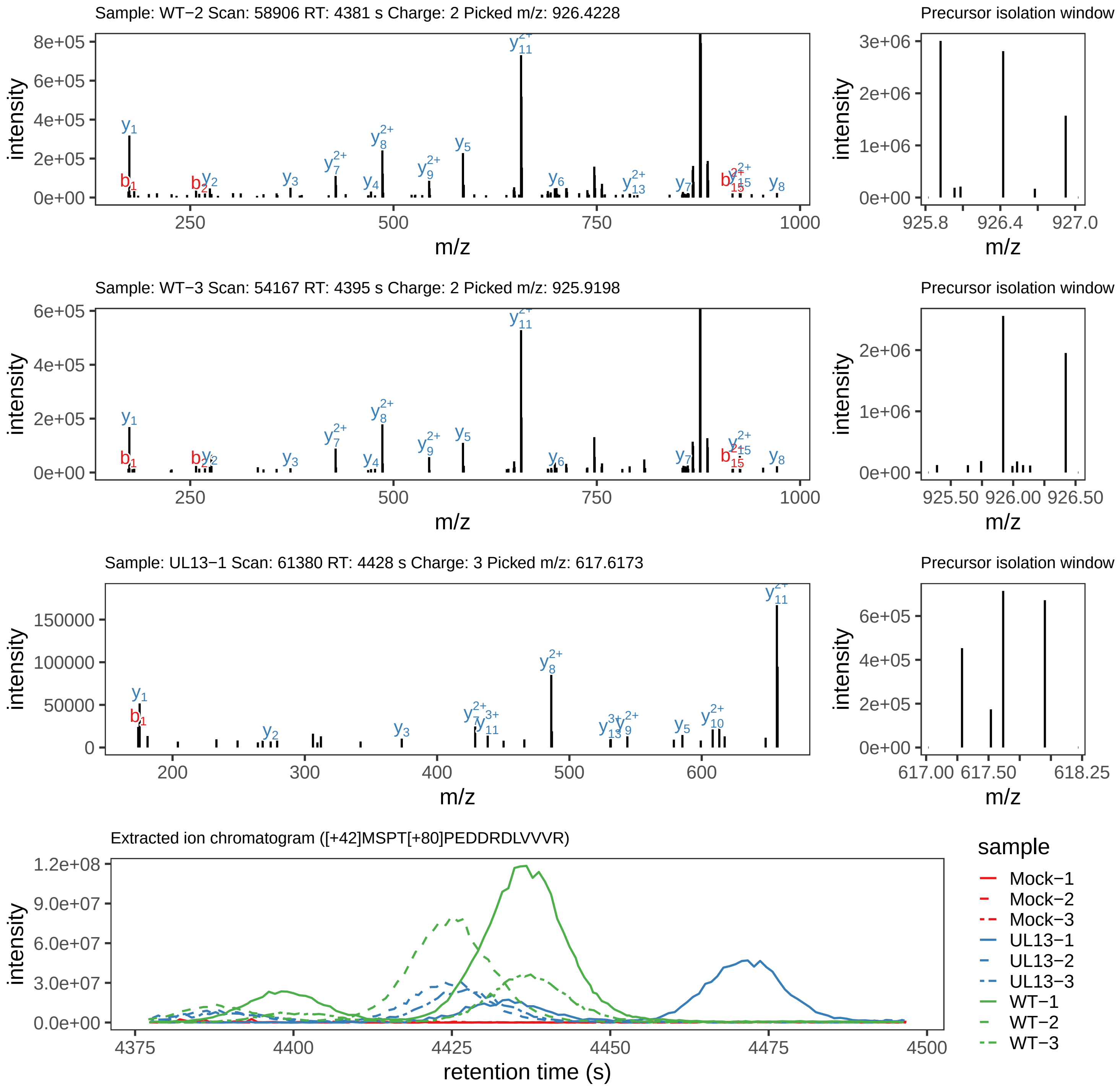

Supplement: Supplemental File 5 [file 737029_file08.zip › phosphopeptide_validation_figs_20240531/MSPTPEDDRDLVVVR/[+42]MSPT[+80]PEDDRDLVVVR.phospho.pdf]

### Peptide [+42]MS[+80]PTPEDDRDLVVVR

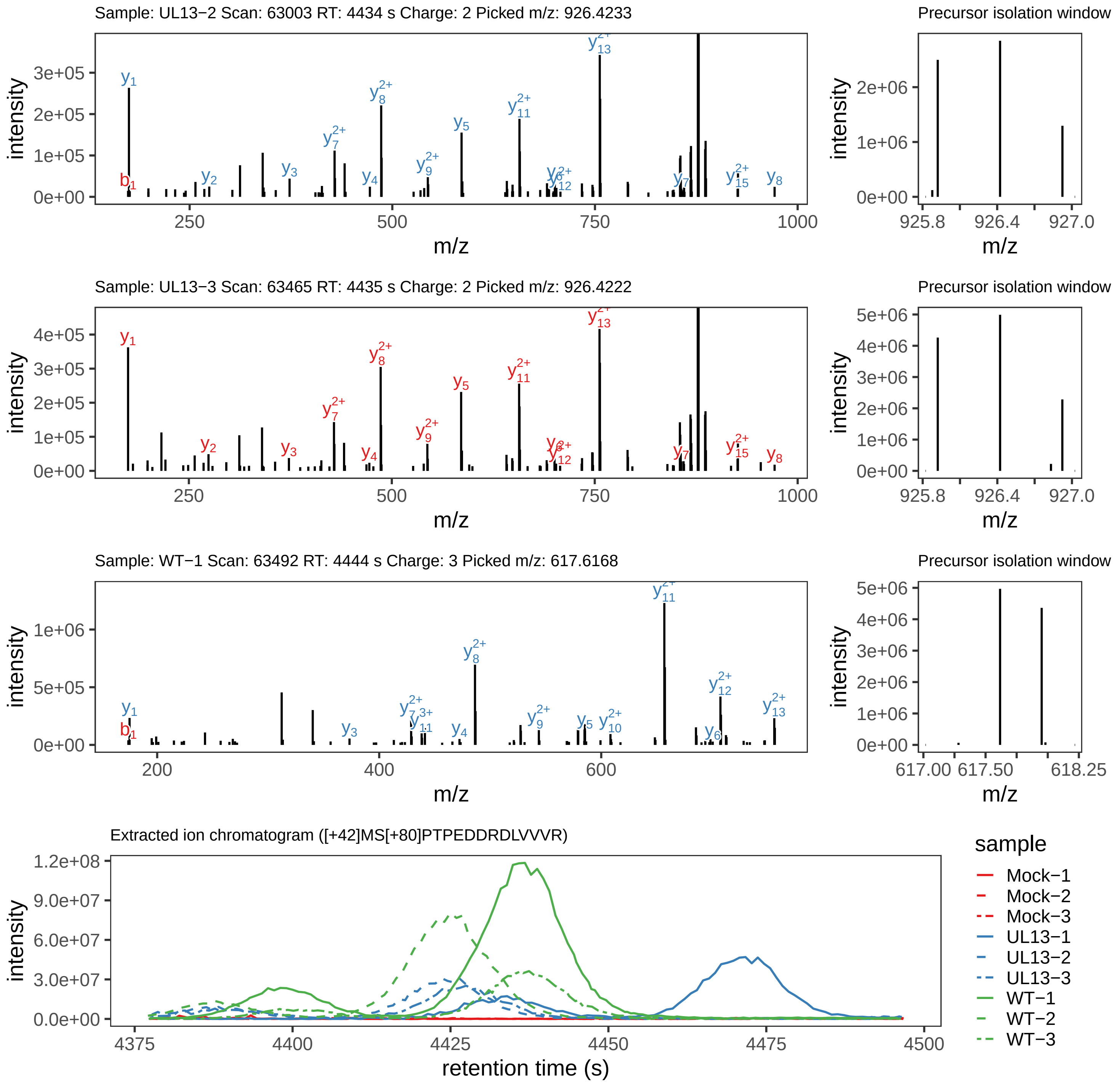

Supplement: Supplemental File 5 [file 737029_file08.zip › phosphopeptide_validation_figs_20240531/MSPTPEDDRDLVVVR/[+42]MS[+80]PTPEDDRDLVVVR.phospho.pdf]

### Peptide [+42]MSPTPEDDRDLVVVR

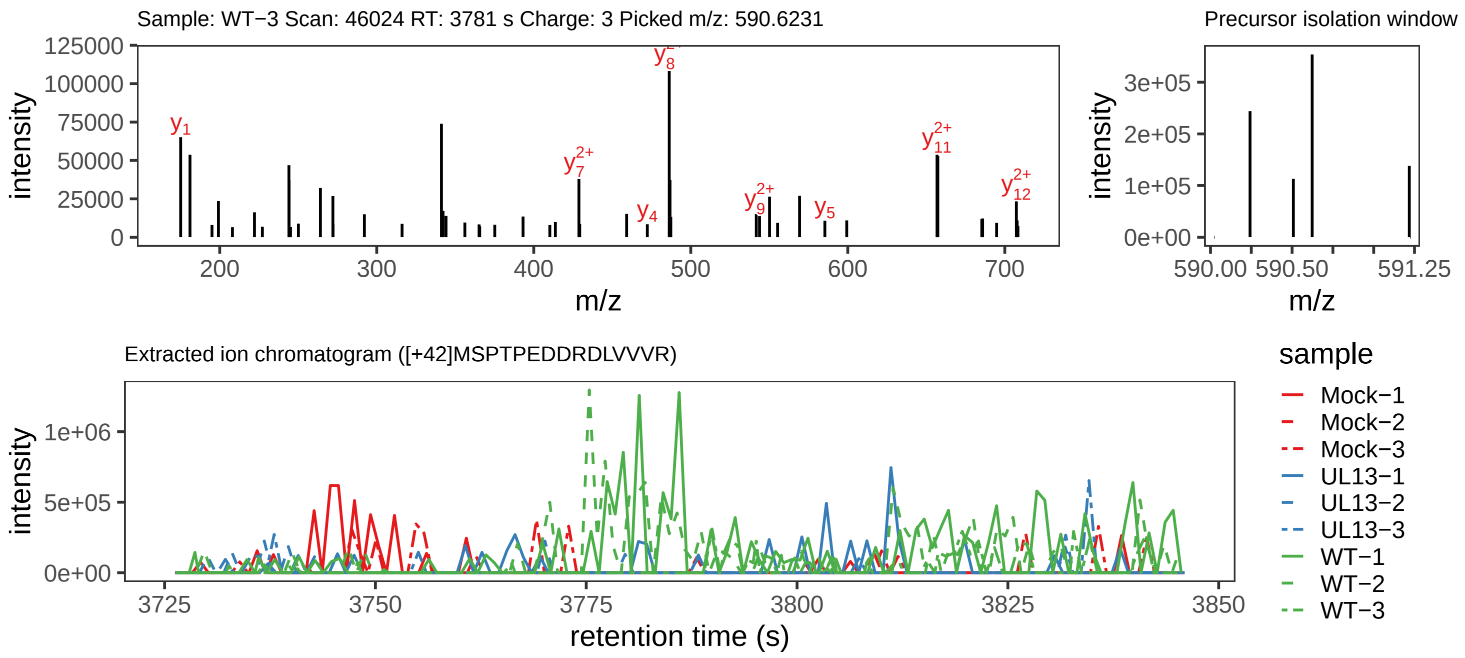

Supplement: Supplemental File 5 [file 737029_file08.zip › phosphopeptide_validation_figs_20240531/MSPTPEDDRDLVVVR/[+42]MSPTPEDDRDLVVVR.phospho.pdf]

### Peptide M[+42]SPT[+80]PEDDRDLVVVR

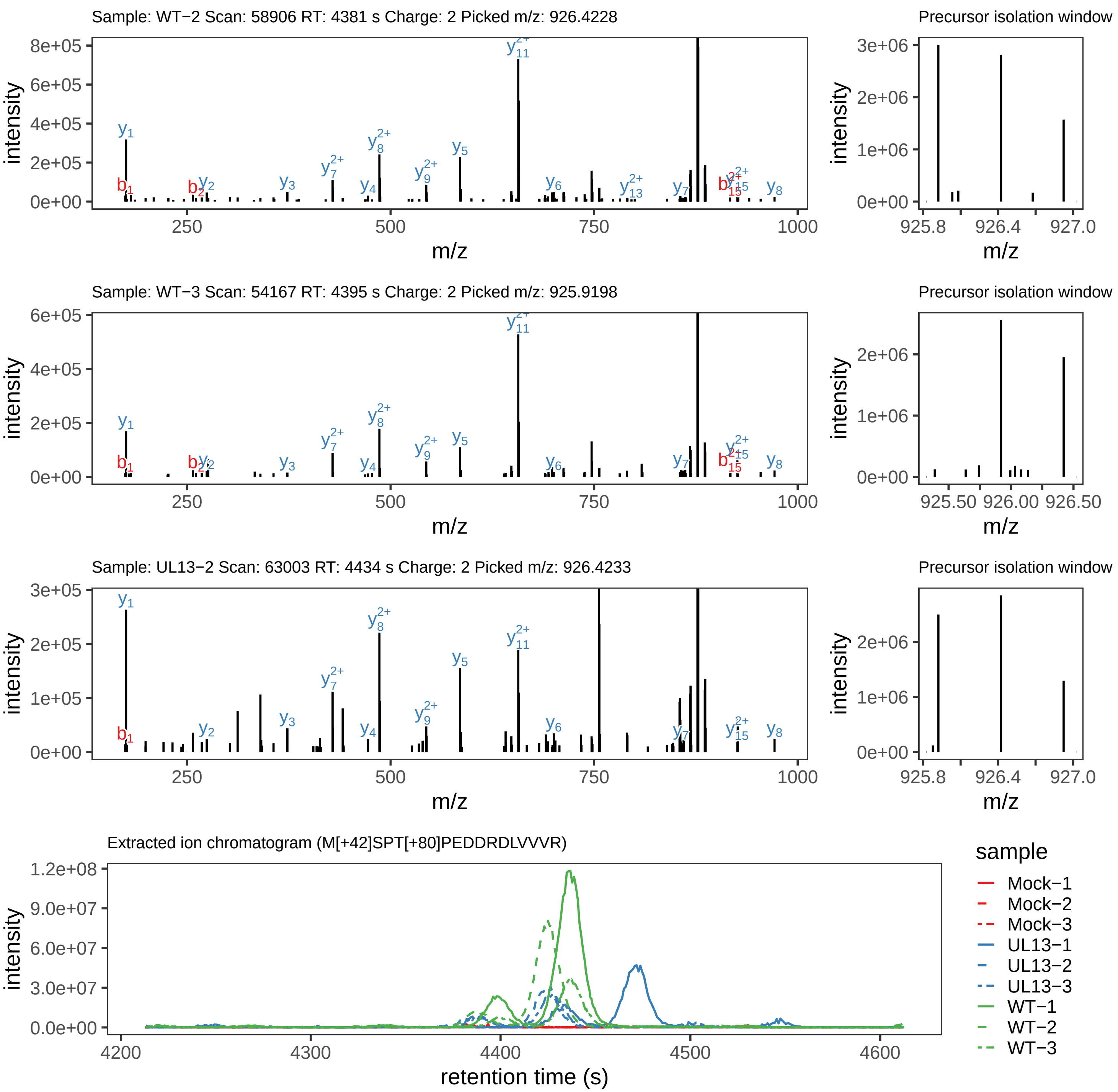

Supplement: Supplemental File 5 [file 737029_file08.zip › phosphopeptide_validation_figs_20240531/MSPTPEDDRDLVVVR/M[+42]SPT[+80]PEDDRDLVVVR.phospho.pdf]

### Peptide M[+16]SPT[+80]PEDDRDLVVVR

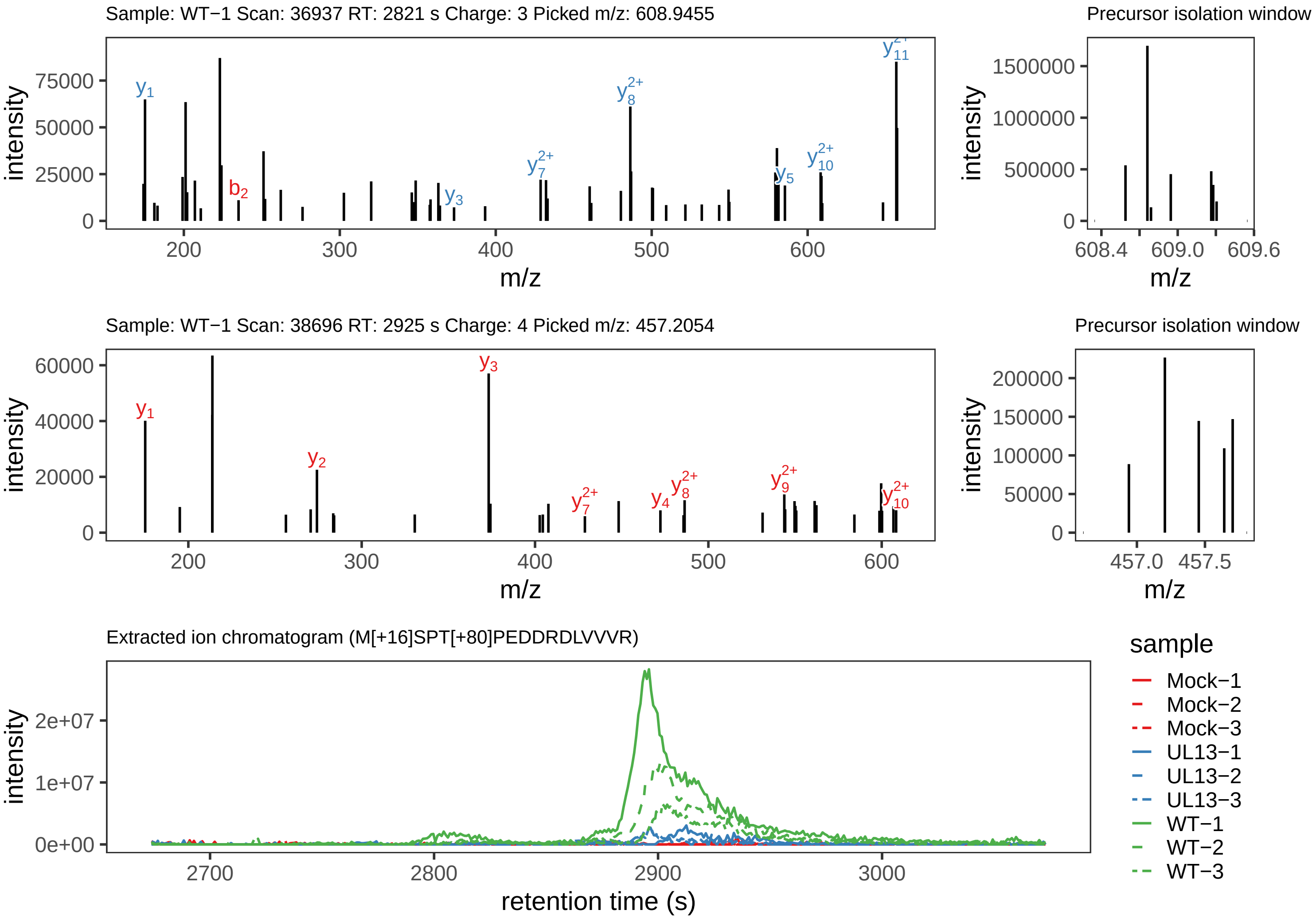

Supplement: Supplemental File 5 [file 737029_file08.zip › phosphopeptide_validation_figs_20240531/MSPTPEDDRDLVVVR/M[+16]SPT[+80]PEDDRDLVVVR.phospho.pdf]

### Peptide [+42]MS[+80]PT[+80]PEDDRDLVVVR

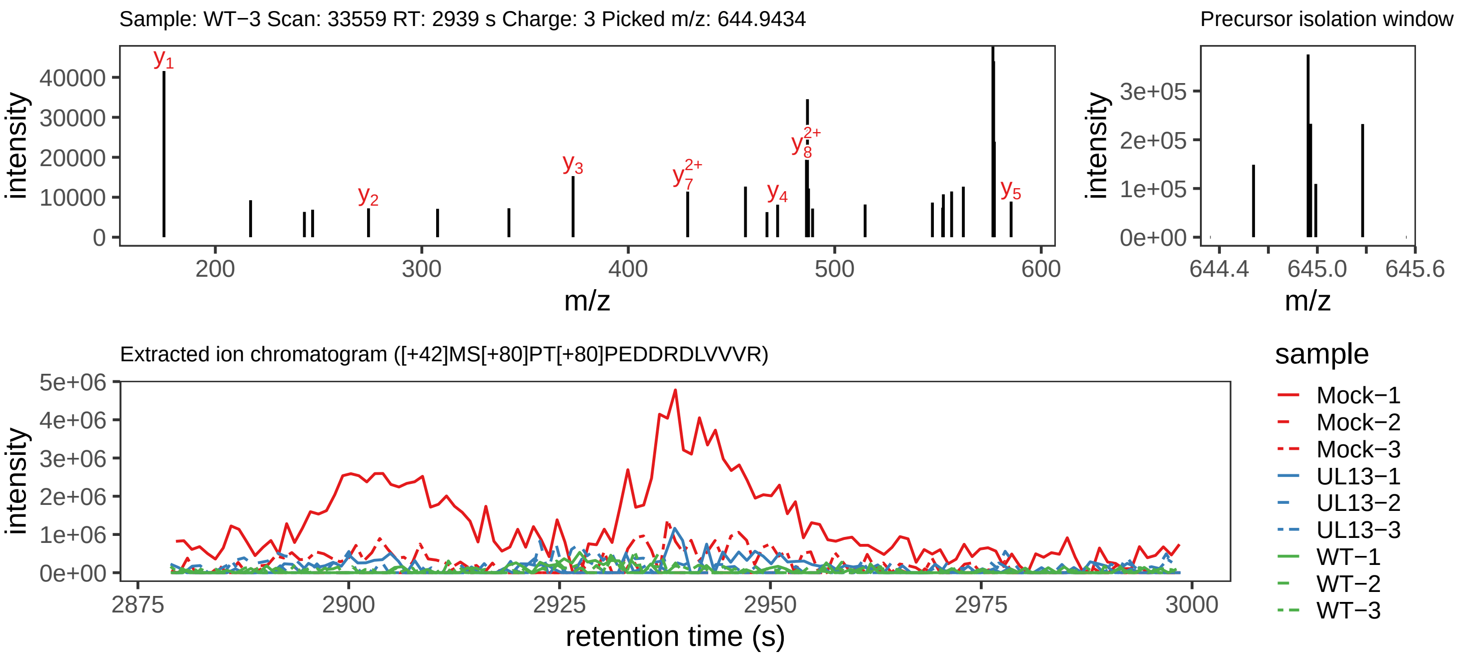

Supplement: Supplemental File 5 [file 737029_file08.zip › phosphopeptide_validation_figs_20240531/MSPTPEDDRDLVVVR/[+42]MS[+80]PT[+80]PEDDRDLVVVR.phospho.pdf]

### Peptide MSPT[+80]PEDDRDLVVVR

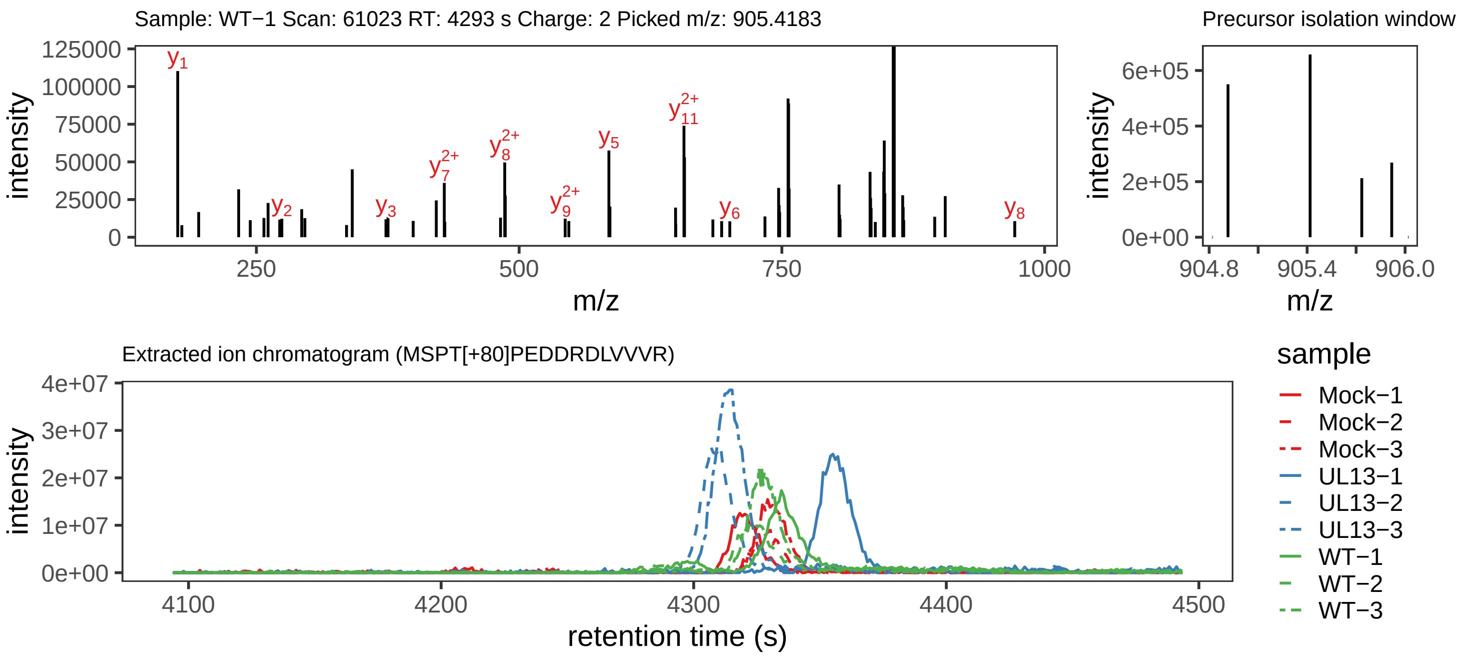

Supplement: Supplemental File 5 [file 737029_file08.zip › phosphopeptide_validation_figs_20240531/MSPTPEDDRDLVVVR/MSPT[+80]PEDDRDLVVVR.phospho.pdf]

### Peptide M[+42]S[+80]PTPEDDRDLVVVR

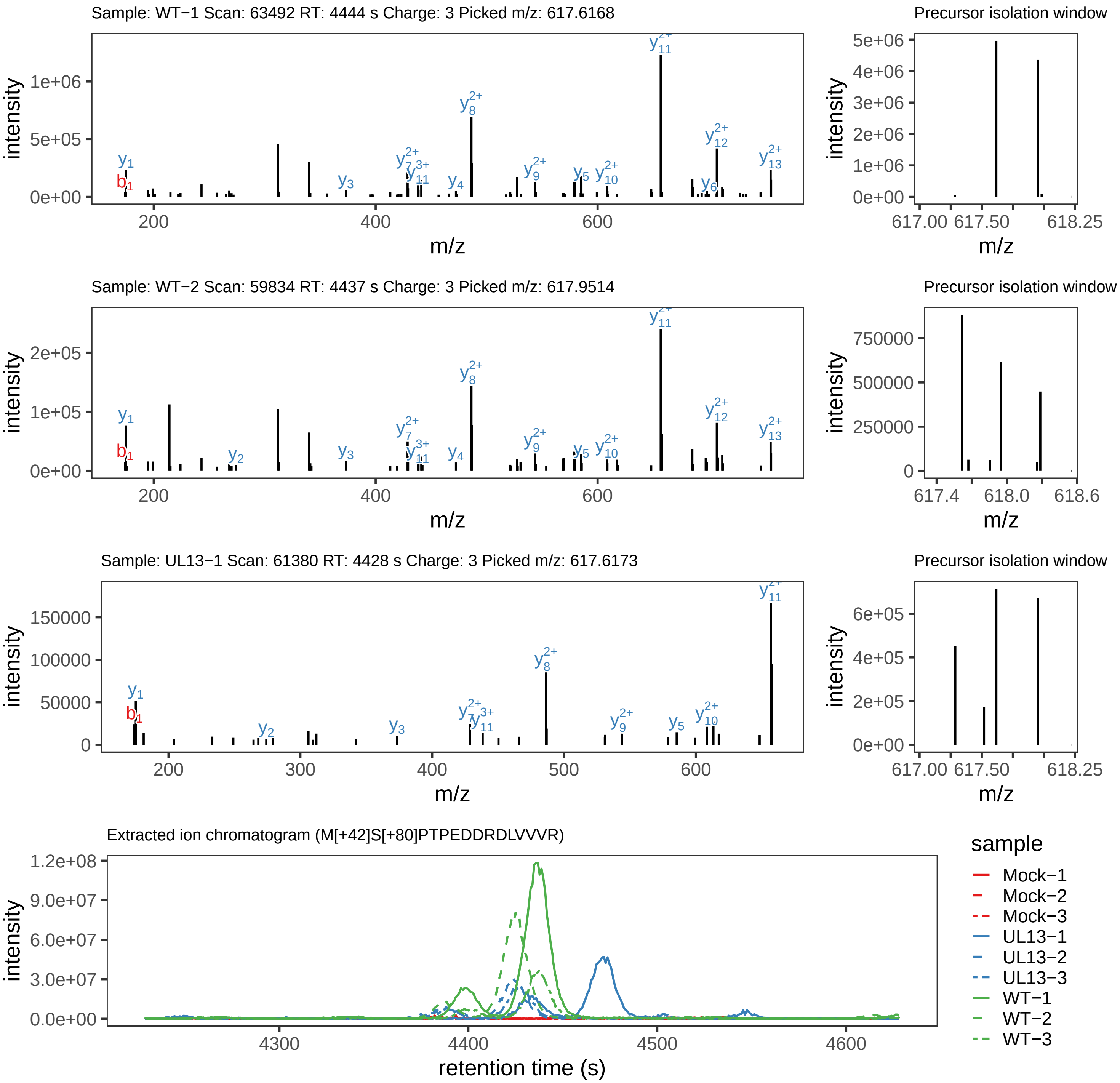

Supplement: Supplemental File 5 [file 737029_file08.zip › phosphopeptide_validation_figs_20240531/MSPTPEDDRDLVVVR/M[+42]S[+80]PTPEDDRDLVVVR.phospho.pdf]

### Peptide M[+16]S[+80]PT[+80]PEDDRDLVVVR

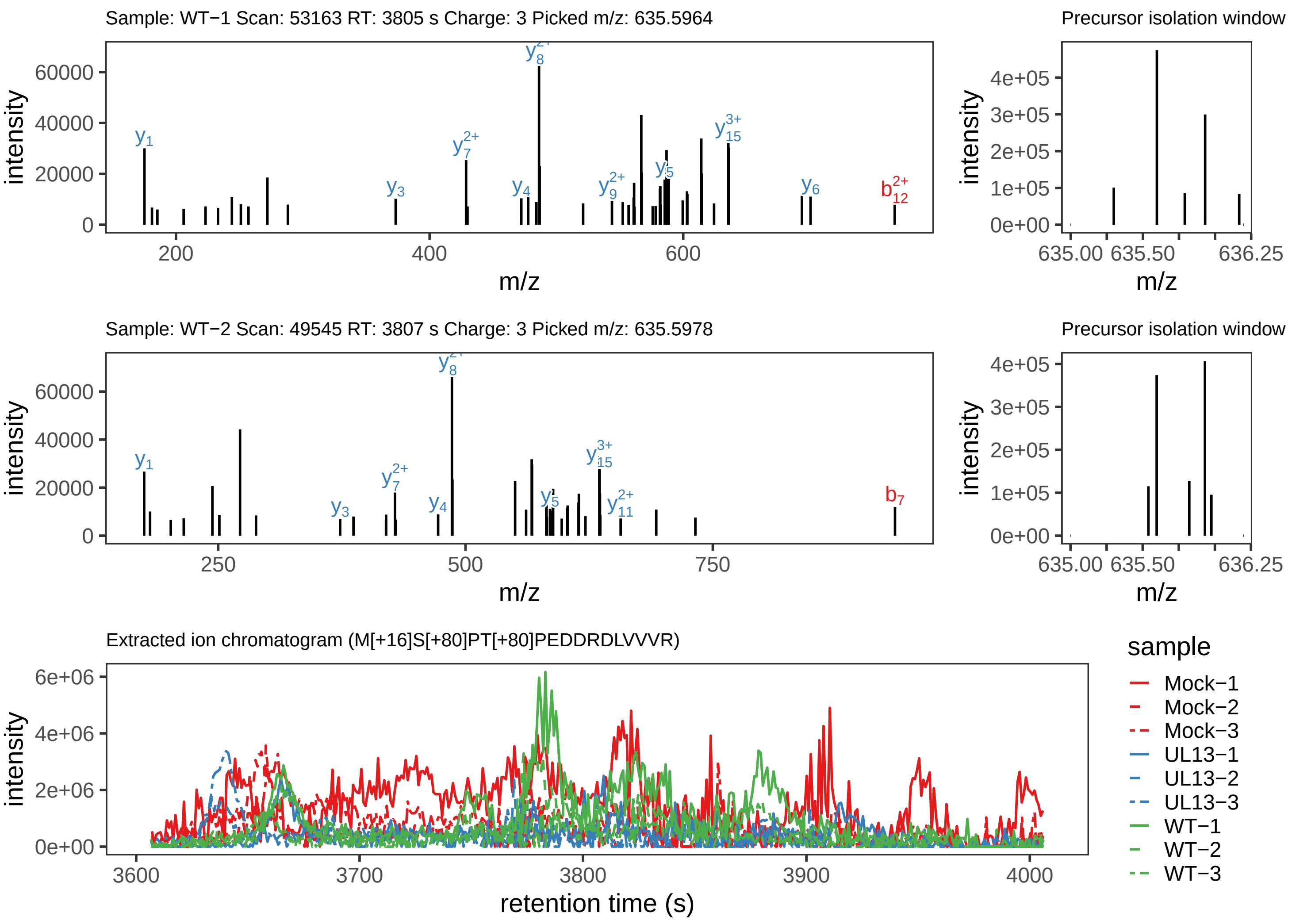

Supplement: Supplemental File 5 [file 737029_file08.zip › phosphopeptide_validation_figs_20240531/MSPTPEDDRDLVVVR/M[+16]S[+80]PT[+80]PEDDRDLVVVR.phospho.pdf]

### Peptide DDLVQPTDLGQ[+1]PSTHEVITC[+57]TSR

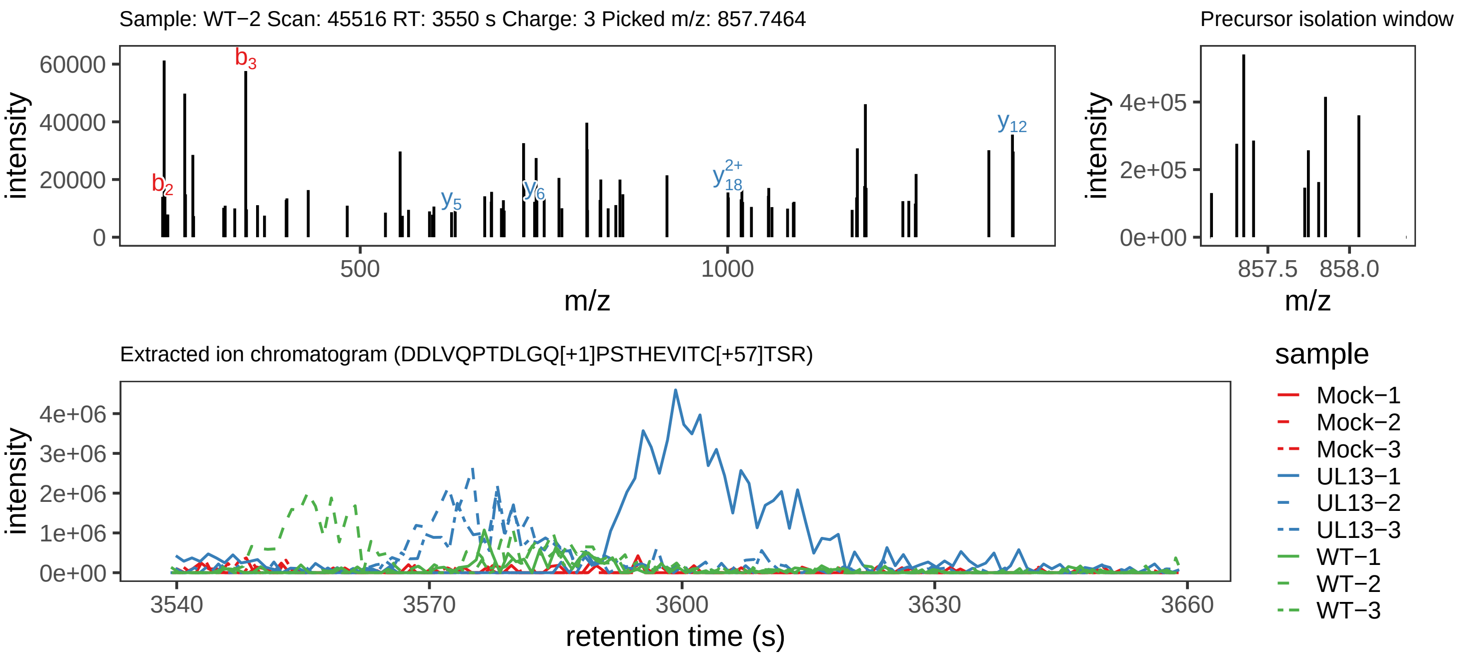

Supplement: Supplemental File 5 [file 737029_file08.zip › phosphopeptide_validation_figs_20240531/DDLVQPTDLGQPSTHEVITCTSR/DDLVQPTDLGQ[+1]PSTHEVITC[+57]TSR.phospho.pdf]

### Peptide DDLVQPTDLGQPST[+80]HEVITC[+57]TSR

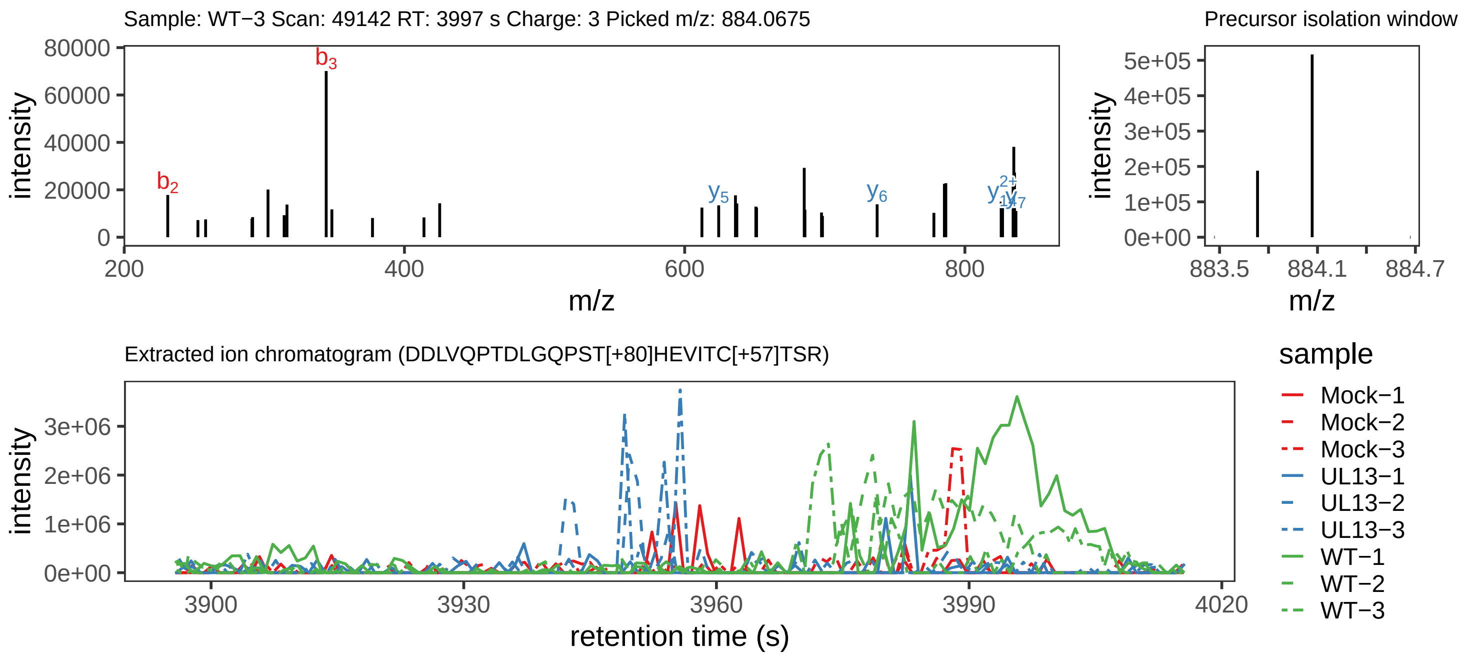

Supplement: Supplemental File 5 [file 737029_file08.zip › phosphopeptide_validation_figs_20240531/DDLVQPTDLGQPSTHEVITCTSR/DDLVQPTDLGQPST[+80]HEVITC[+57]TSR.phospho.pdf]

### Peptide DDLVQPTDLGQPS[+80]THEVITC[+57]TSR

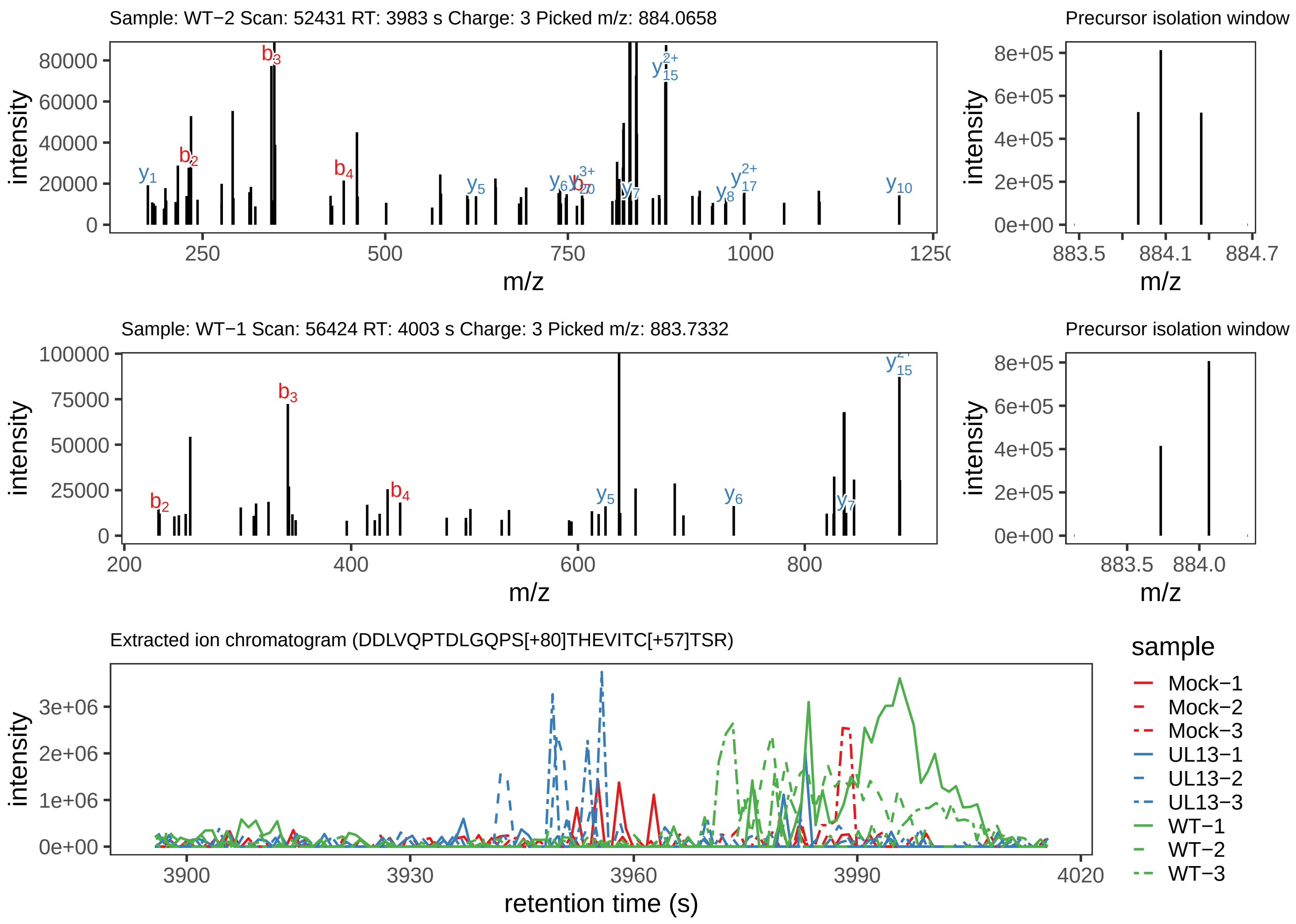

Supplement: Supplemental File 5 [file 737029_file08.zip › phosphopeptide_validation_figs_20240531/DDLVQPTDLGQPSTHEVITCTSR/DDLVQPTDLGQPS[+80]THEVITC[+57]TSR.phospho.pdf]

### Peptide DDLVQPTDLGQ[+1]PST[+80]HEVITC[+57]TSR

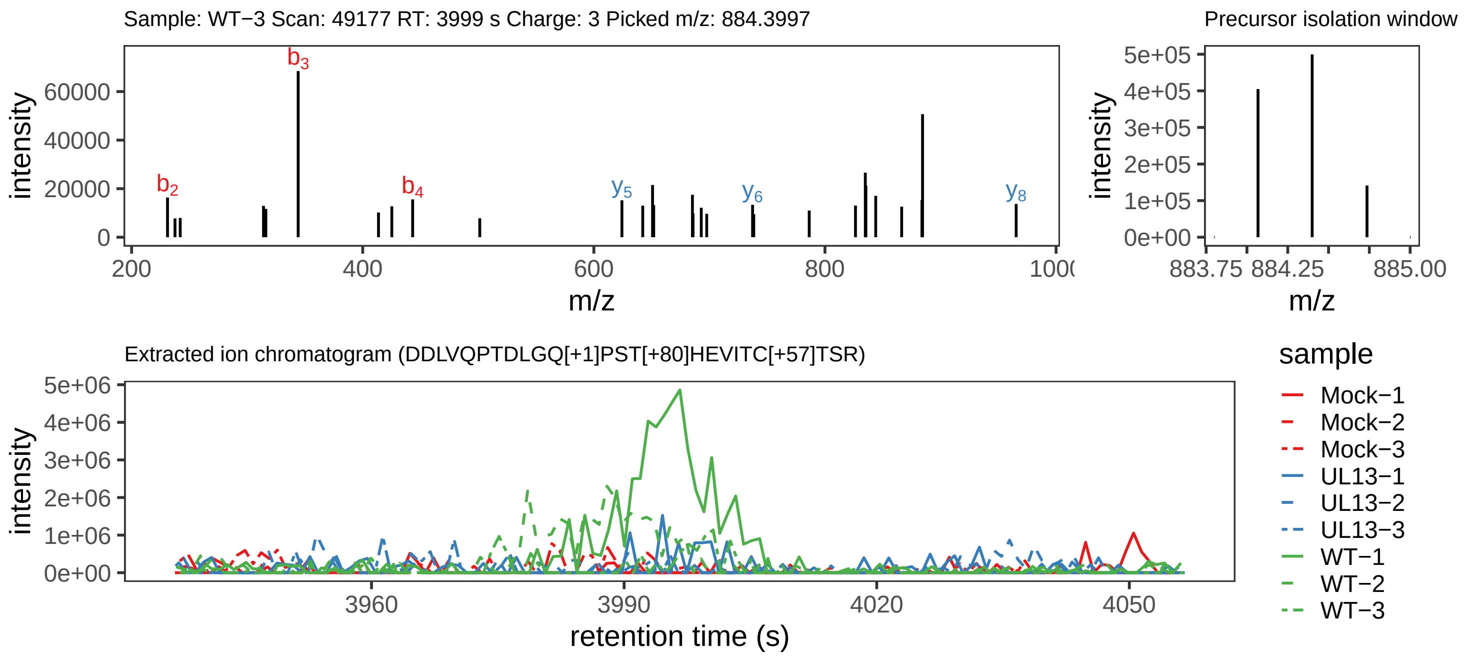

Supplement: Supplemental File 5 [file 737029_file08.zip › phosphopeptide_validation_figs_20240531/DDLVQPTDLGQPSTHEVITCTSR/DDLVQPTDLGQ[+1]PST[+80]HEVITC[+57]TSR.phospho.pdf]

### Peptide DDLVQPTDLGQPSTHEVITC[+57]TSR

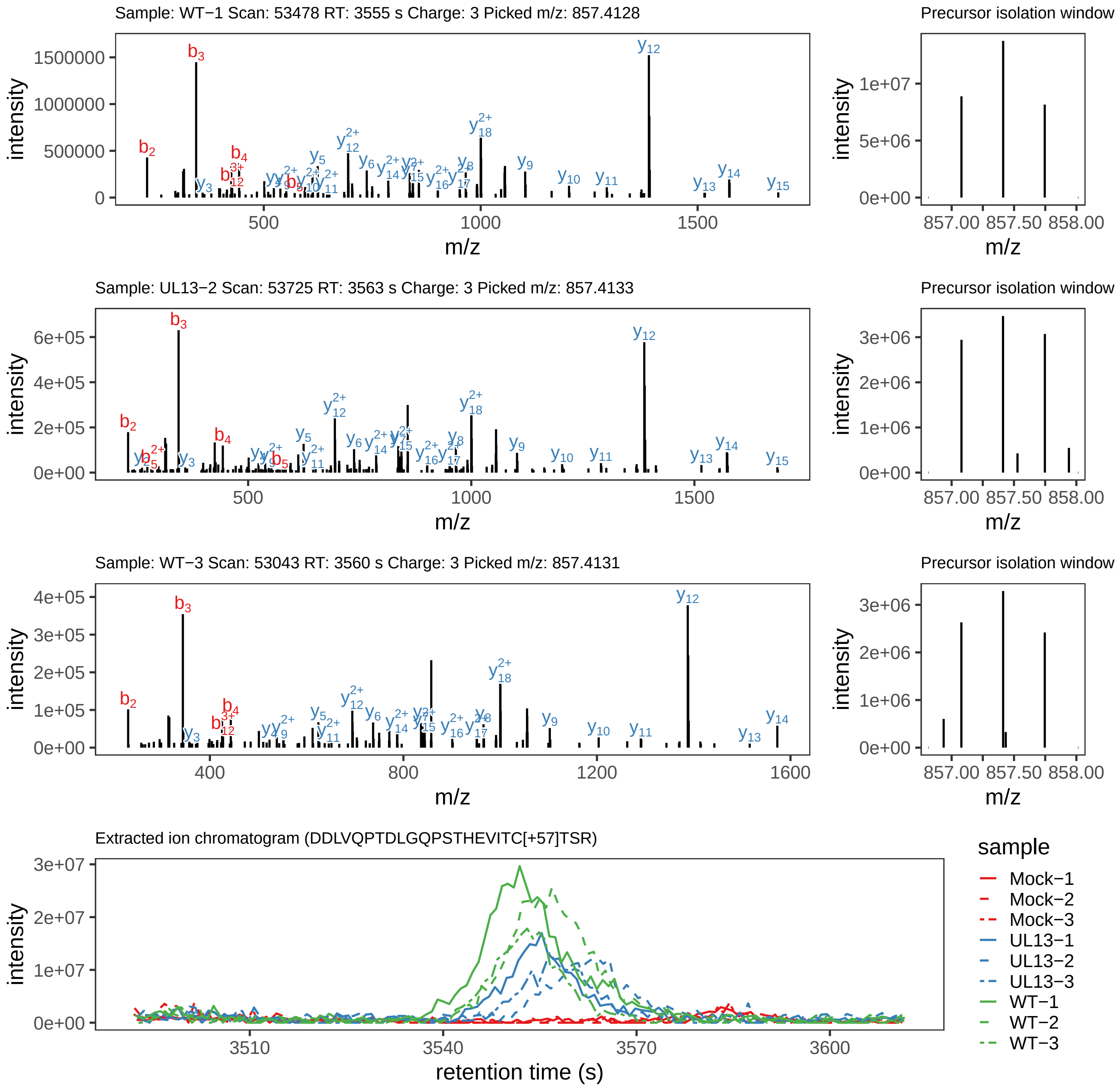

Supplement: Supplemental File 5 [file 737029_file08.zip › phosphopeptide_validation_figs_20240531/DDLVQPTDLGQPSTHEVITCTSR/DDLVQPTDLGQPSTHEVITC[+57]TSR.global.pdf]

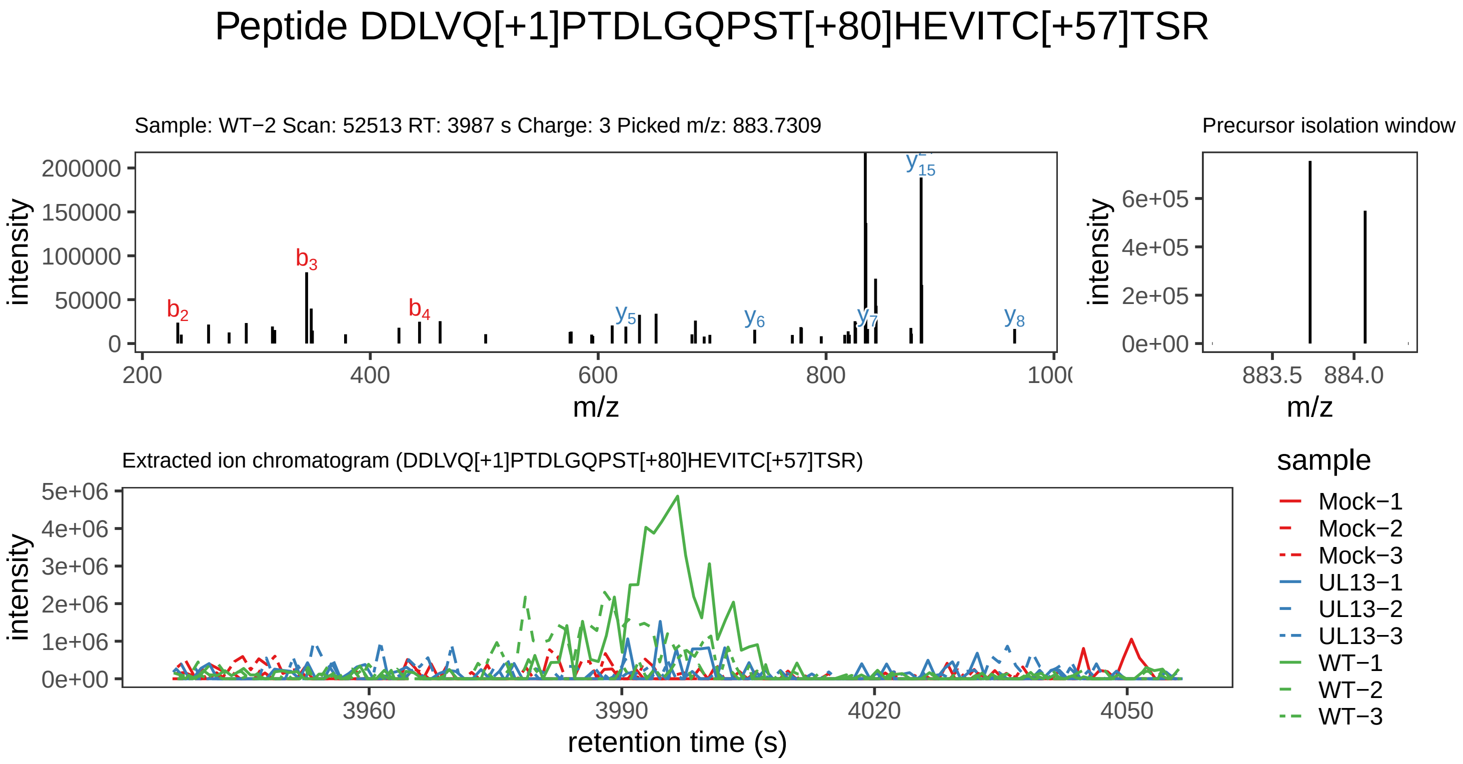

Supplement: Supplemental File 5 [file 737029_file08.zip › phosphopeptide_validation_figs_20240531/DDLVQPTDLGQPSTHEVITCTSR/DDLVQ[+1]PTDLGQPST[+80]HEVITC[+57]TSR.phospho.pdf]

### Peptide RNDFHS[+80]PSSR

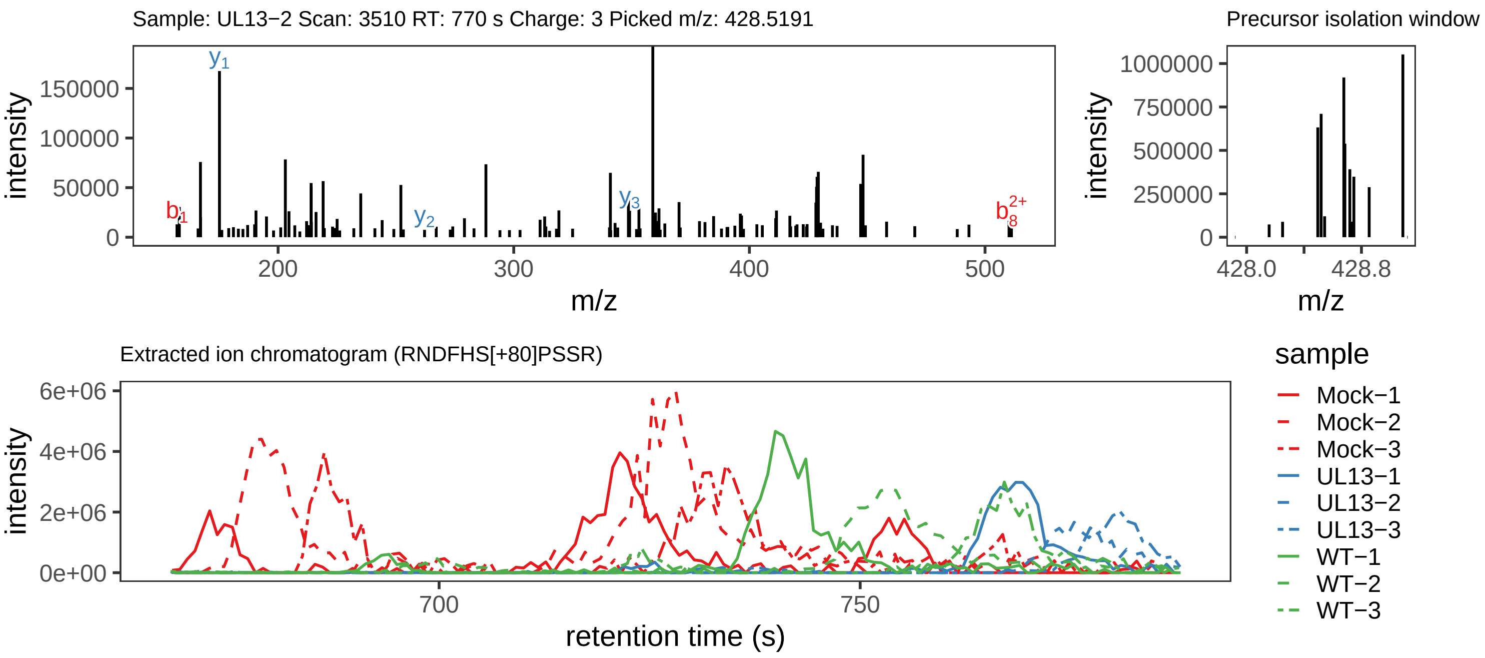

Supplement: Supplemental File 5 [file 737029_file08.zip › phosphopeptide_validation_figs_20240531/RNDFHSPSSR/RNDFHS[+80]PSSR.global.pdf]

### Peptide RN[+1]DFHS[+80]PSSR

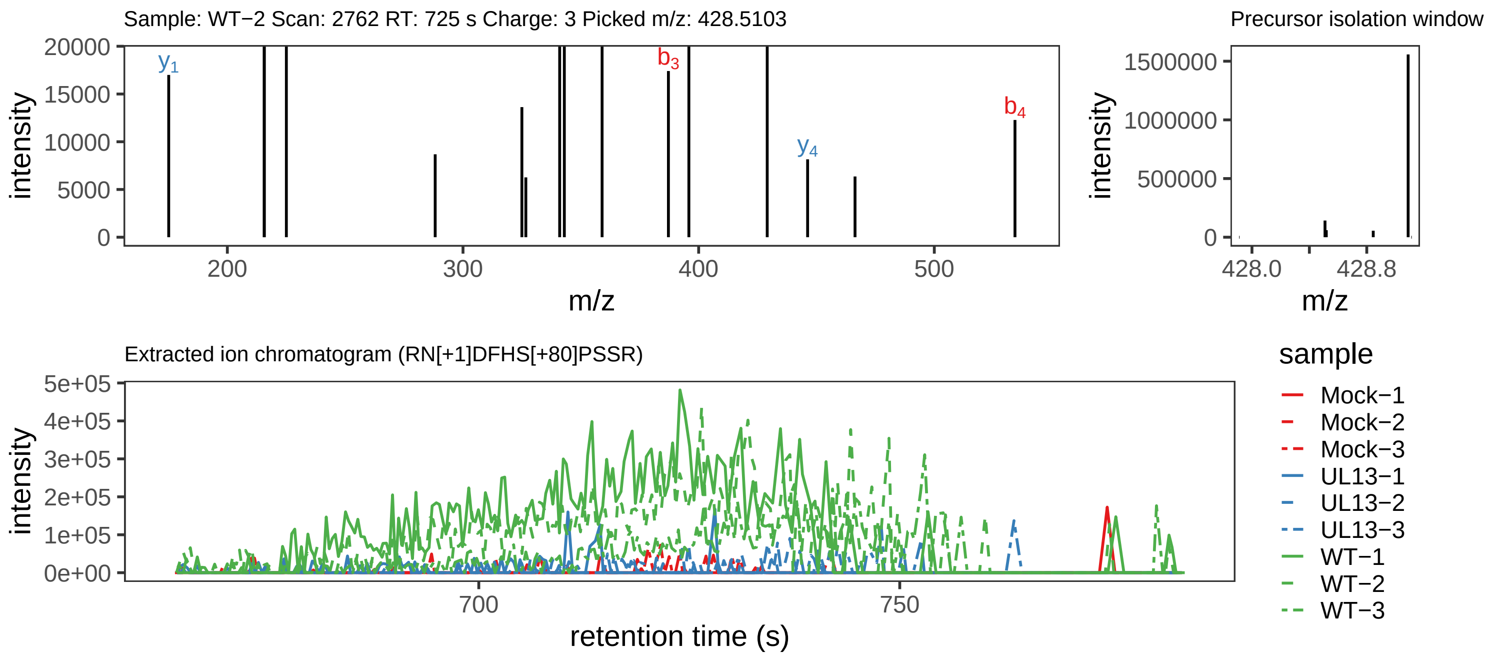

Supplement: Supplemental File 5 [file 737029_file08.zip › phosphopeptide_validation_figs_20240531/RNDFHSPSSR/RN[+1]DFHS[+80]PSSR.phospho.pdf]

### Peptide RNDFHS[+80]PSSR

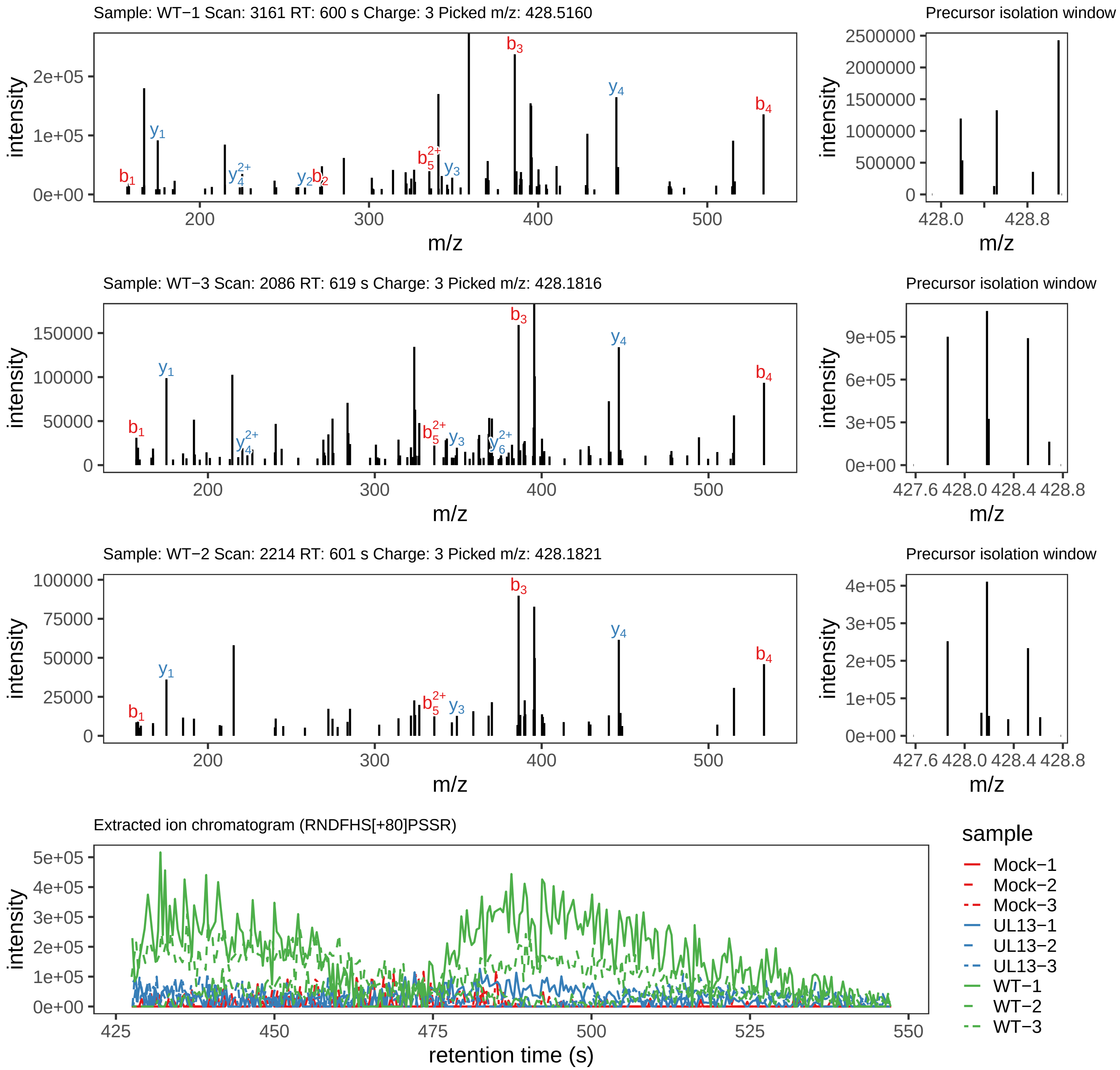

Supplement: Supplemental File 5 [file 737029_file08.zip › phosphopeptide_validation_figs_20240531/RNDFHSPSSR/RNDFHS[+80]PSSR.phospho.pdf]

### Peptide VELSTLLR

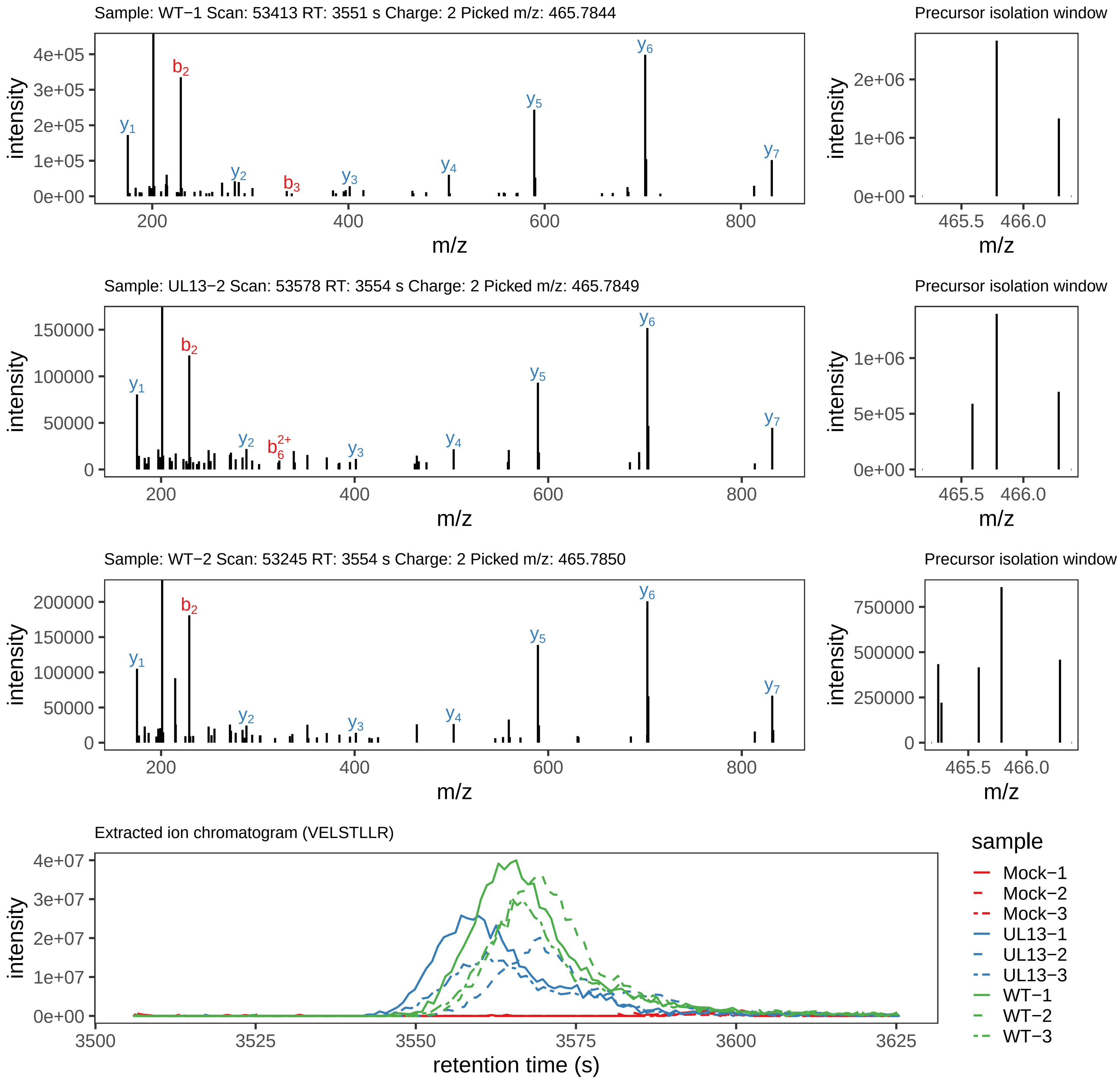

Supplement: Supplemental File 5 [file 737029_file08.zip › phosphopeptide_validation_figs_20240531/VELSTLLR/VELSTLLR.global.pdf]

### Peptide VELSTLLR

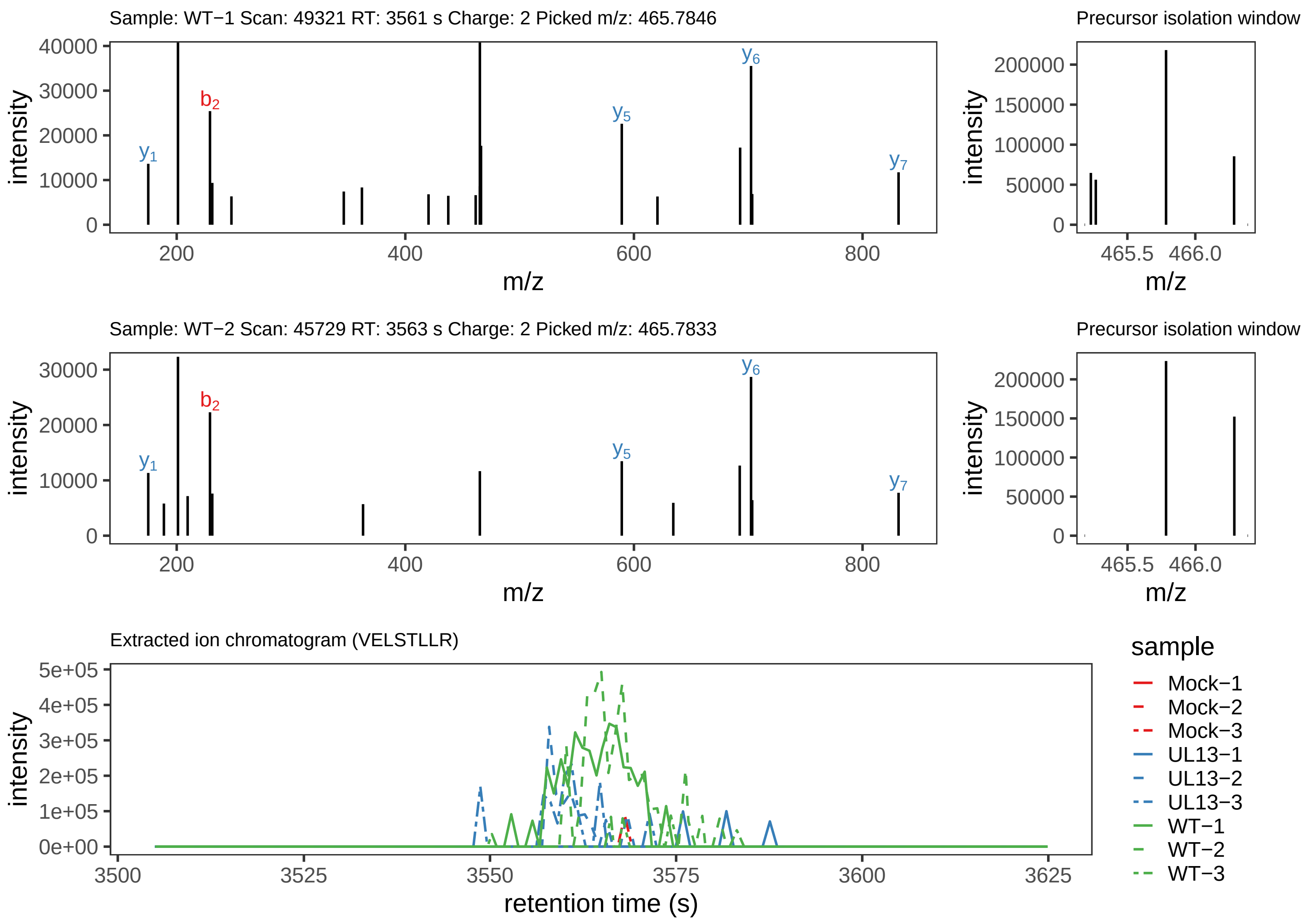

Supplement: Supplemental File 5 [file 737029_file08.zip › phosphopeptide_validation_figs_20240531/VELSTLLR/VELSTLLR.phospho.pdf]

### Peptide SLQM[+16]FILC[+57]R

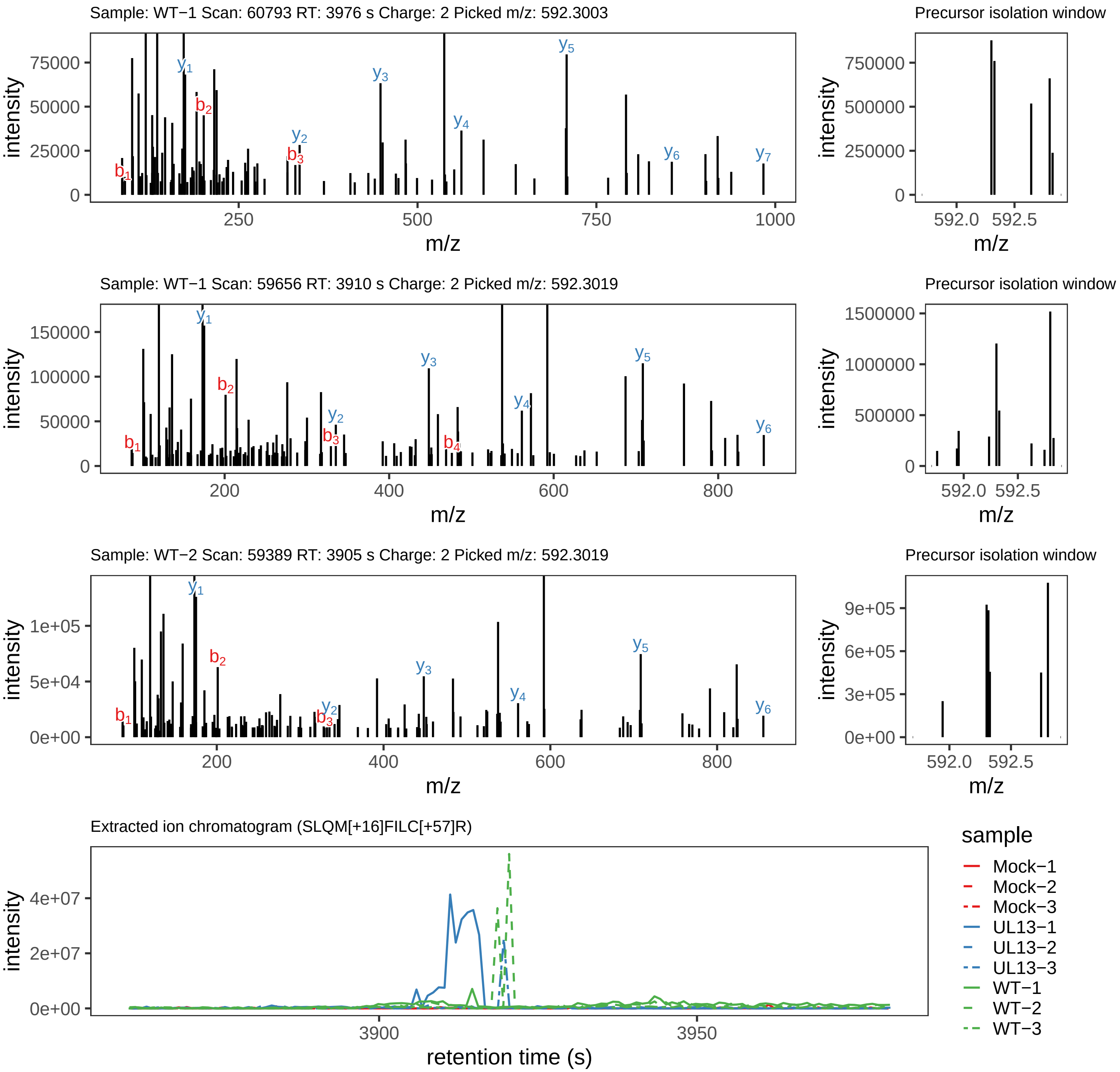

Supplement: Supplemental File 5 [file 737029_file08.zip › phosphopeptide_validation_figs_20240531/SLQMFILCR/SLQM[+16]FILC[+57]R.global.pdf]

### Peptide S[+80]LQM[+16]FILC[+57]R

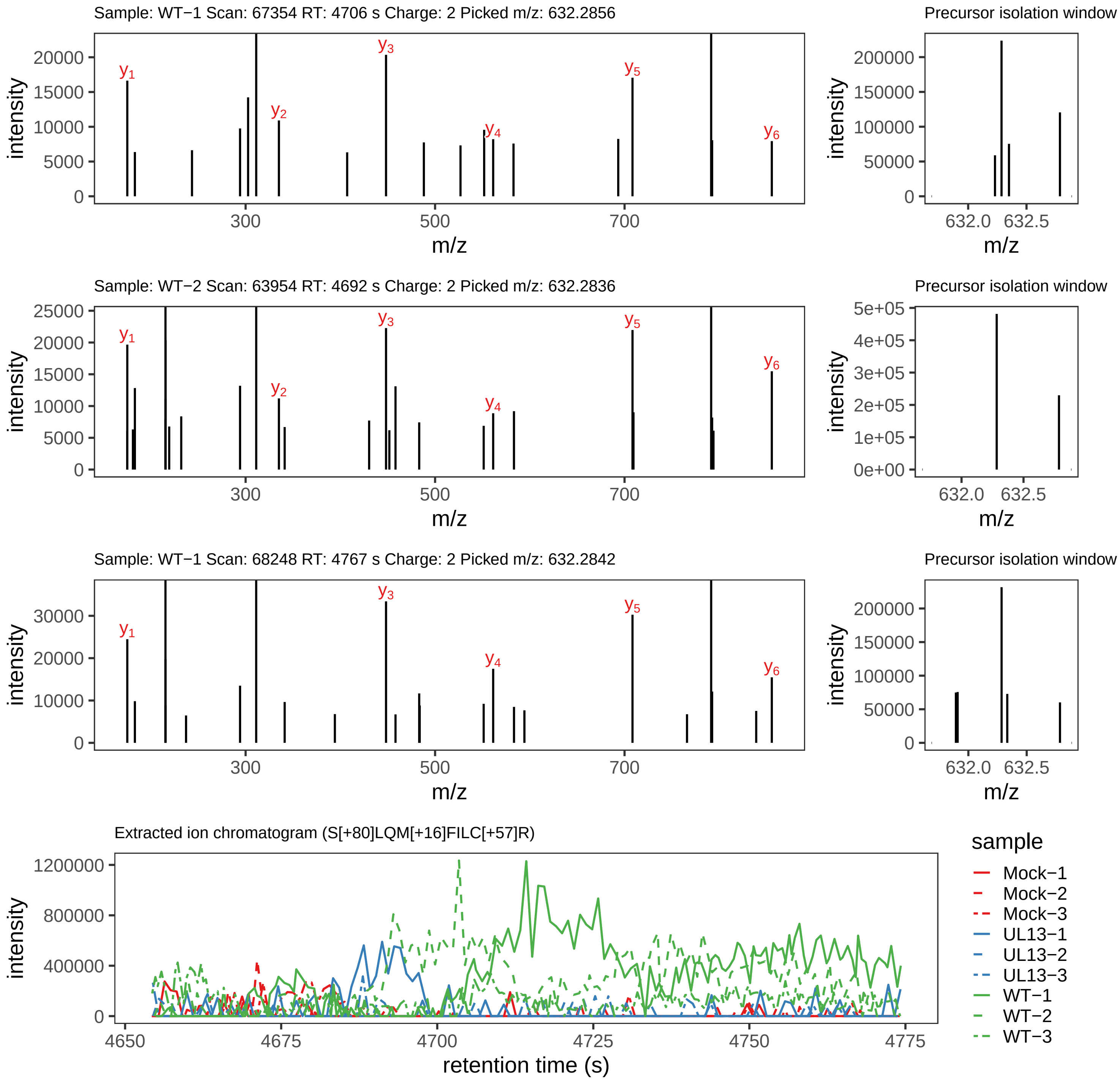

Supplement: Supplemental File 5 [file 737029_file08.zip › phosphopeptide_validation_figs_20240531/SLQMFILCR/S[+80]LQM[+16]FILC[+57]R.phospho.pdf]

### Peptide SLQMFILC[+57]R

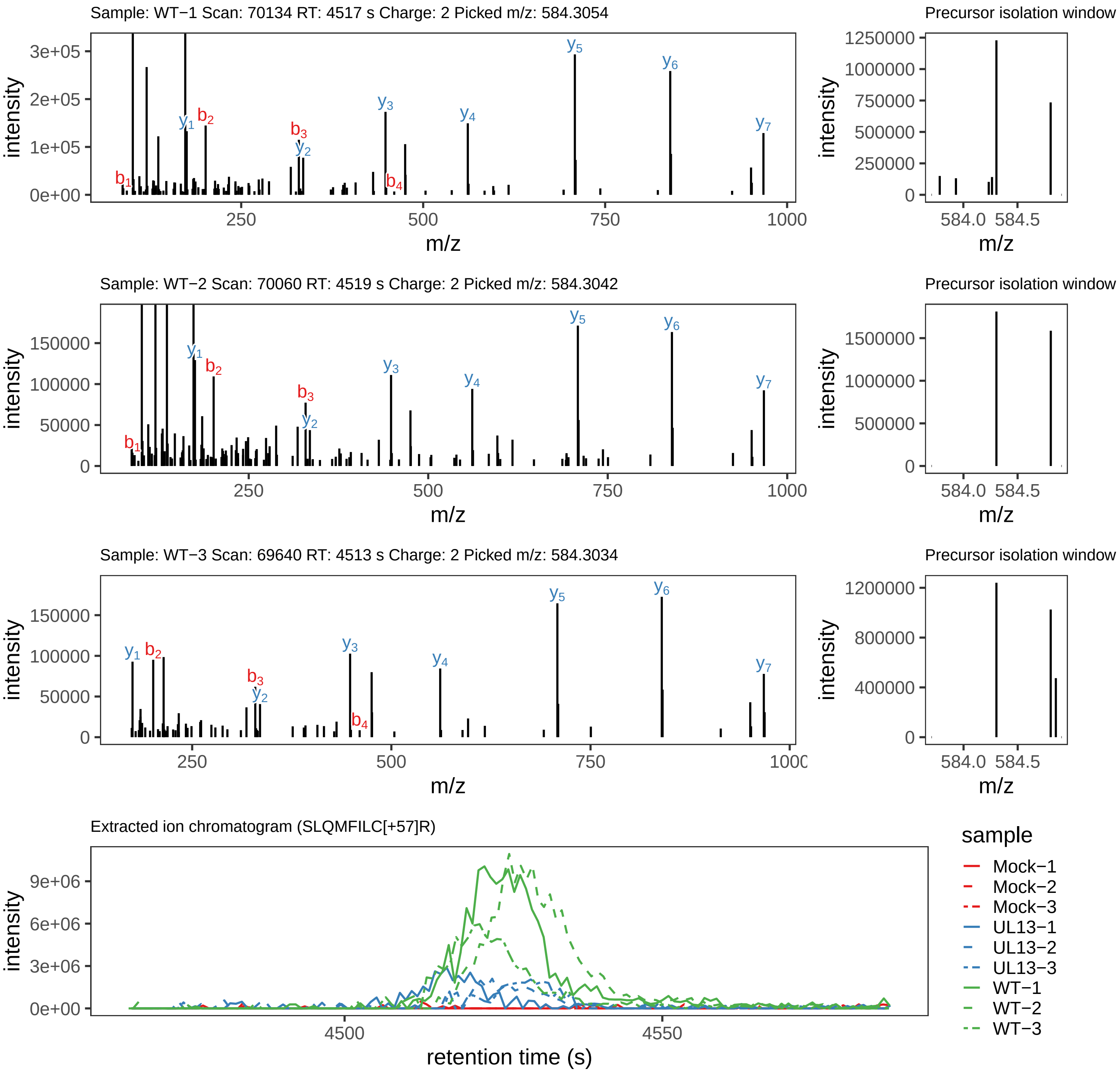

Supplement: Supplemental File 5 [file 737029_file08.zip › phosphopeptide_validation_figs_20240531/SLQMFILCR/SLQMFILC[+57]R.global.pdf]

### Peptide S[+80]LQMFILC[+57]R

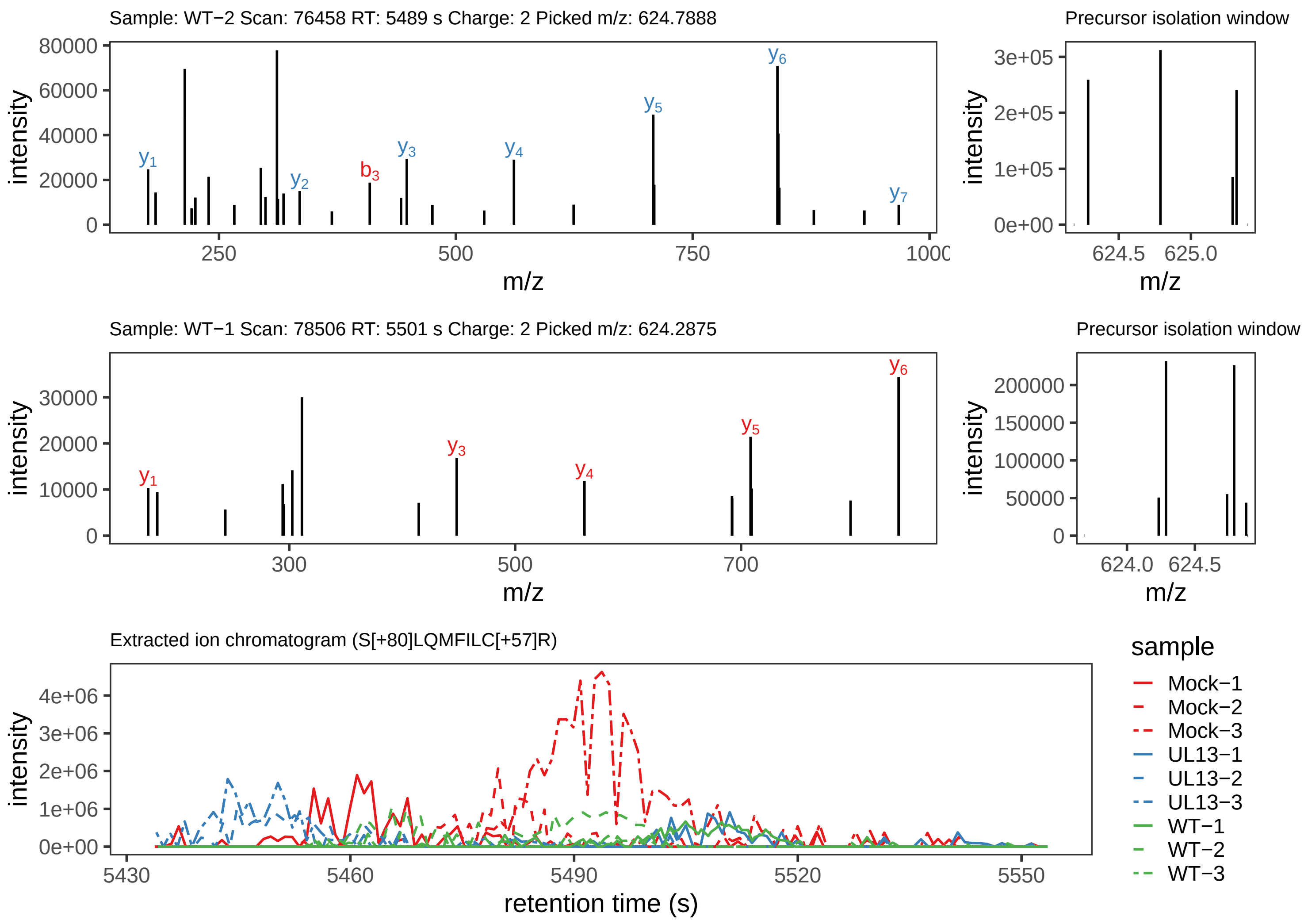

Supplement: Supplemental File 5 [file 737029_file08.zip › phosphopeptide_validation_figs_20240531/SLQMFILCR/S[+80]LQMFILC[+57]R.phospho.pdf]

### Peptide ETYVRHS[+80]PR

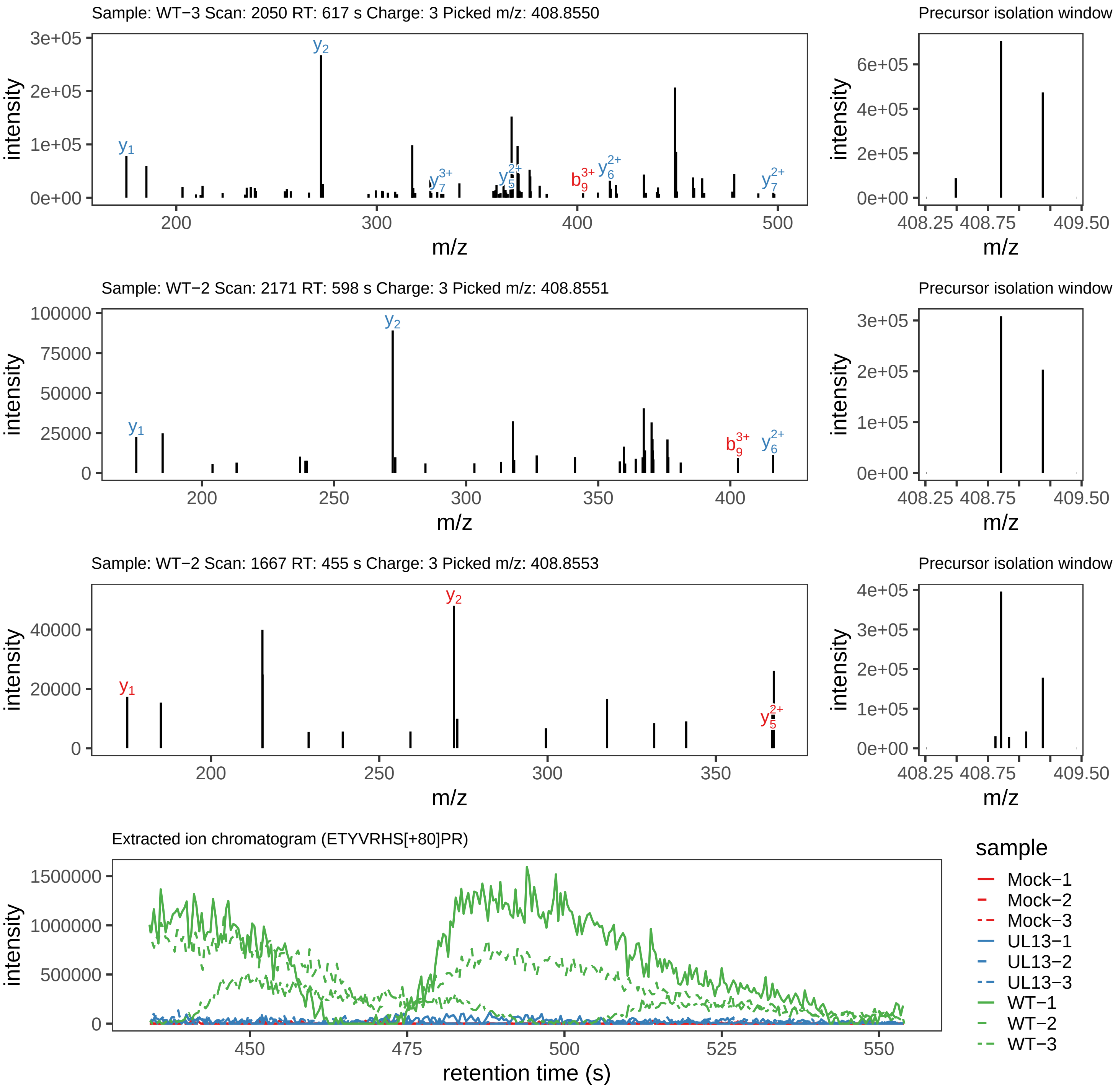

Supplement: Supplemental File 5 [file 737029_file08.zip › phosphopeptide_validation_figs_20240531/ETYVRHSPR/ETYVRHS[+80]PR.phospho.pdf]

### Peptide ET[+80]YVRHSPR

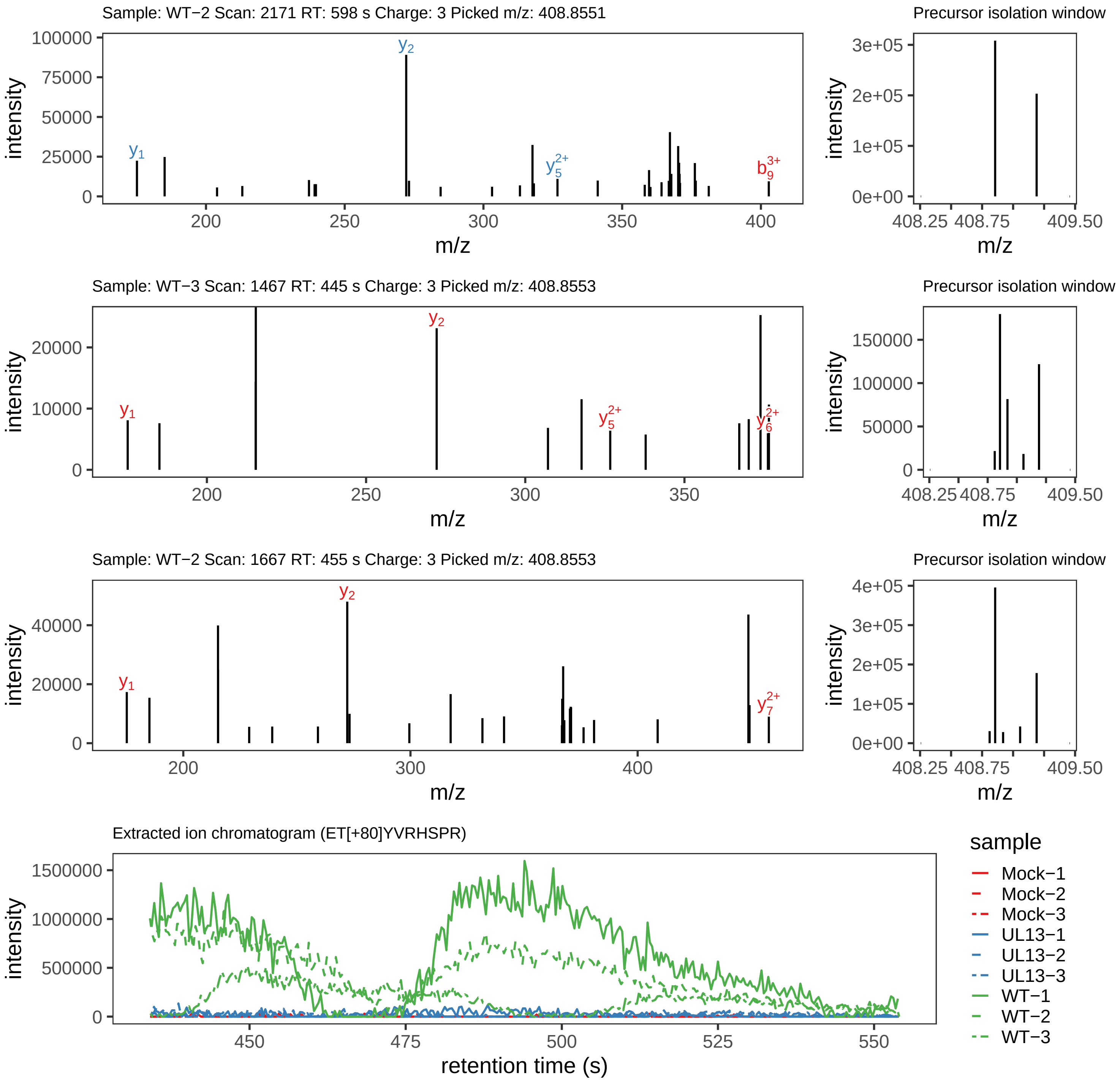

Supplement: Supplemental File 5 [file 737029_file08.zip › phosphopeptide_validation_figs_20240531/ETYVRHSPR/ET[+80]YVRHSPR.phospho.pdf]

### Peptide GDAQIFENSTIHTMRDPMASAAR

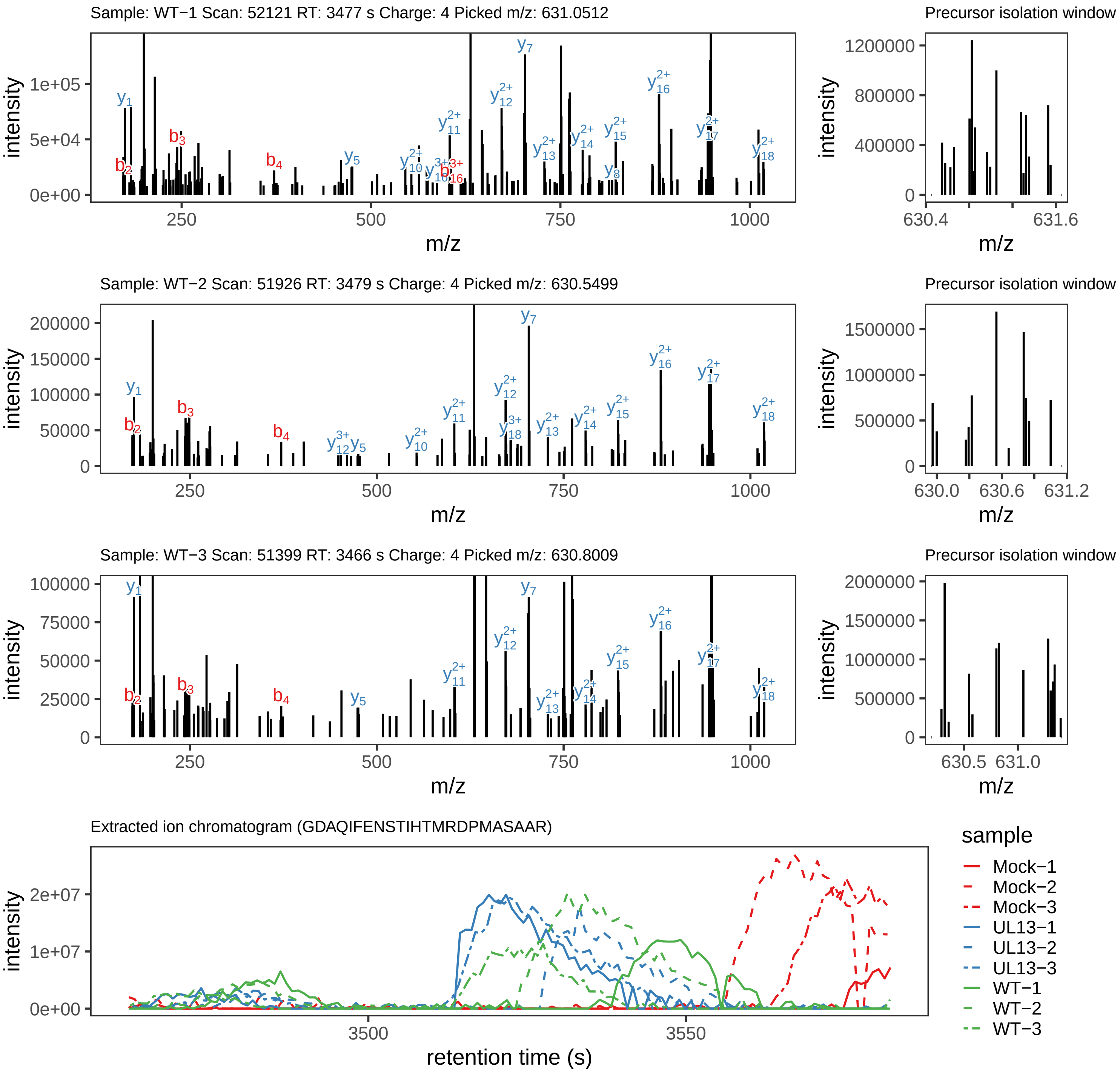

Supplement: Supplemental File 5 [file 737029_file08.zip › phosphopeptide_validation_figs_20240531/GDAQIFENSTIHTMRDPMASAAR/GDAQIFENSTIHTMRDPMASAAR.global.pdf]

### Peptide VS[+80]RNVFITGK

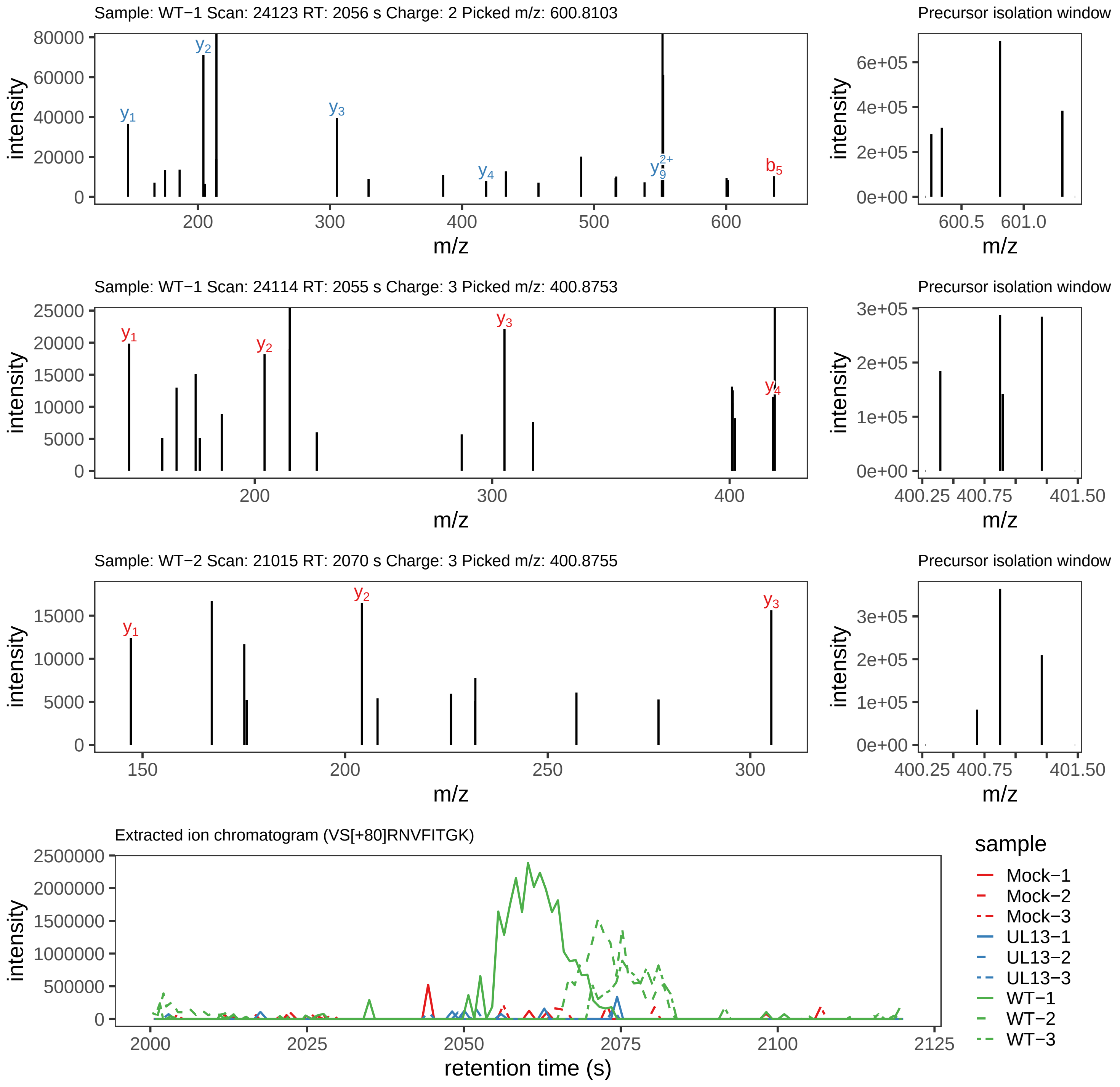

Supplement: Supplemental File 5 [file 737029_file08.zip › phosphopeptide_validation_figs_20240531/VSRNVFITGK/VS[+80]RNVFITGK.phospho.pdf]

### Peptide ILGSPPN[+1]PLKPPEIEPPQMS[+80]STPGR

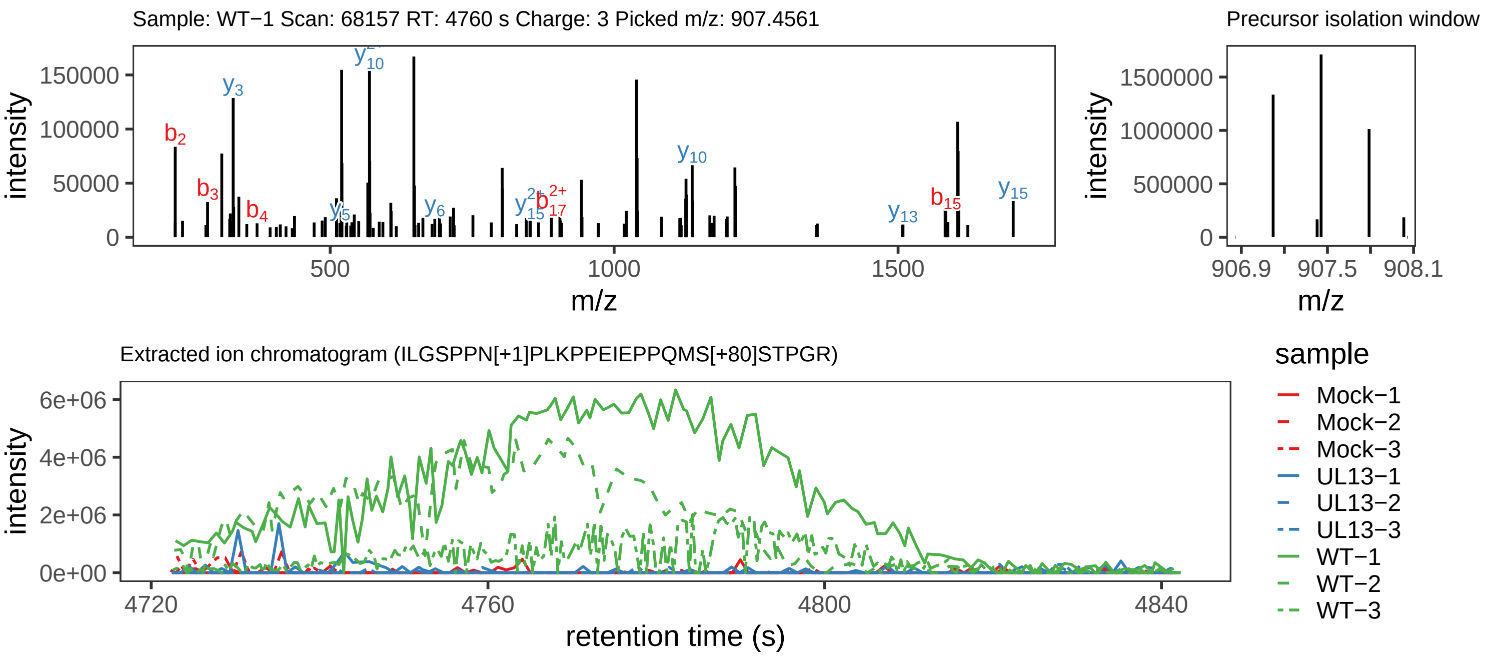

Supplement: Supplemental File 5 [file 737029_file08.zip › phosphopeptide_validation_figs_20240531/ILGSPPNPLKPPEIEPPQMSSTPGR/ILGSPPN[+1]PLKPPEIEPPQMS[+80]STPGR.phospho.pdf]

### Peptide ILGSPPN[+1]PLKPPEIEPPQM[+16]S[+80]STPGR

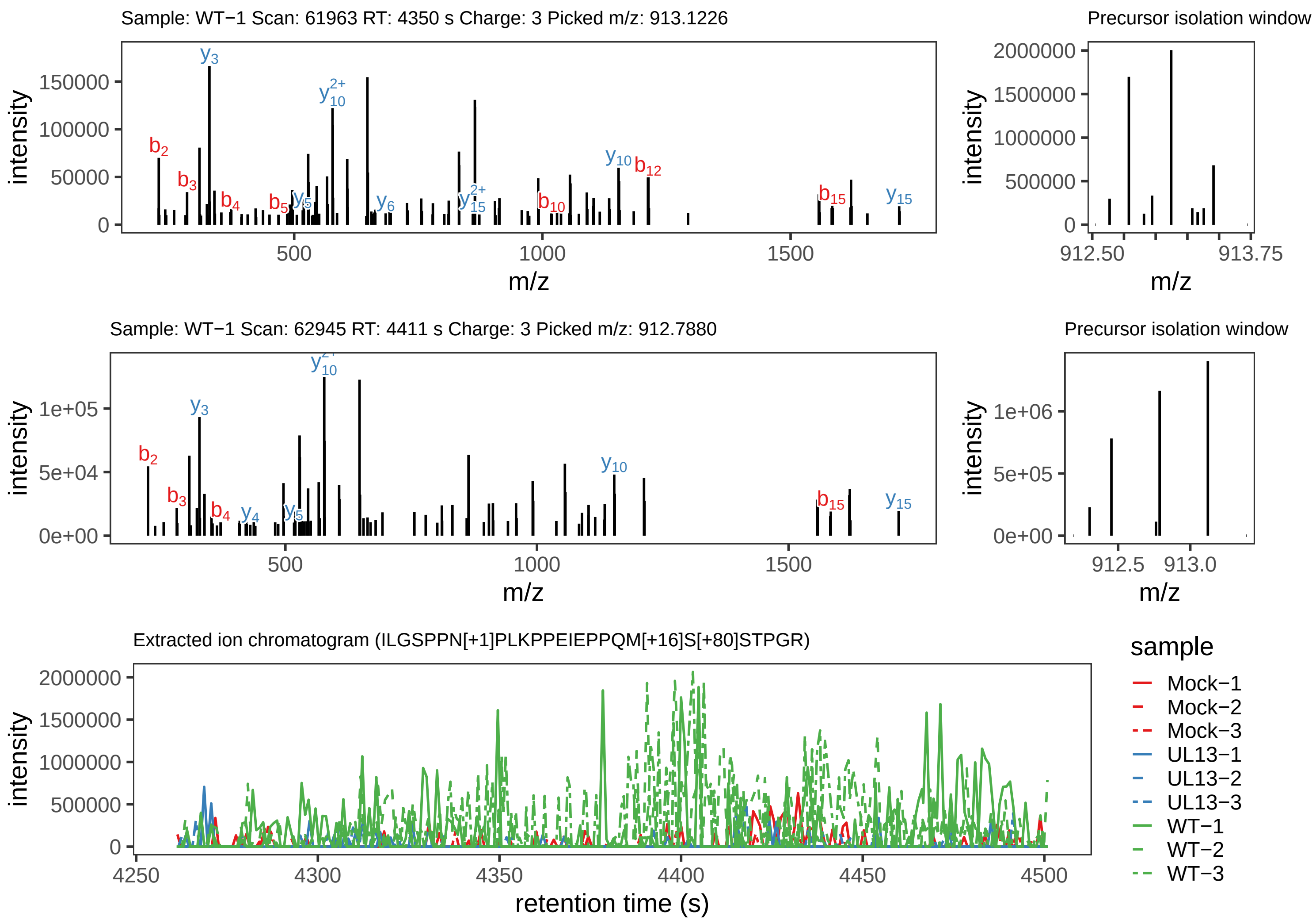

Supplement: Supplemental File 5 [file 737029_file08.zip › phosphopeptide_validation_figs_20240531/ILGSPPNPLKPPEIEPPQMSSTPGR/ILGSPPN[+1]PLKPPEIEPPQM[+16]S[+80]STPGR.phospho.pdf]
