## Supplemental Tables for "Conserved herpesvirus protein kinase (CHPK)-mediated phosphorylation of viral proteins associated with nucleocytoplasmic trafficking during natural infection"

**S1 Table. Summary of relative infection level in feather follicles used in the experimental plan.**

| Group | Sample | Clinical Notes | UL47eGFP[a] | DPI[b] |
|---|---|---|---|---|
| Mock | U1 | No clinical signs or gross lesions | - | 21 |
| Mock[c] | U2 | No clinical signs or gross lesions | - | 31 |
| Mock[c] | U3 | No clinical signs or gross lesions | - | 31 |
| Mock | U4 | No clinical signs or gross lesions | - | 31 |
| Mock[c] | U5 | No clinical signs or gross lesions | - | 35 |
| Mock | U6 | No clinical signs or gross lesions | - | 35 |
| vCHPKwt[c] | 41 | Spleen enlarged | +++ | 23 |
| vCHPKwt | 42 | Spleen enlarged | ++ | 23 |
| vCHPKwt[c] | 43 | Liver, spleen, and kidney tumors | +++ | 24 |
| vCHPKwt | 44 | Breast, liver, ovary, kidney, and spleen tumors | ++ | 28 |
| vCHPKwt | 45 | Leg paralysis; liver, spleen, and kidney tumors | ++ | 31 |
| vCHPKwt[c] | 46 | Liver, spleen, kidney, and ovary tumors | +++ | 24 |
| vΔCHPK[c] | 81 | Testicular tumors | +++ | 22 |
| vΔCHPK[c] | 82 | Yellow liver | +++ | 23 |
| vΔCHPK[c] | 83 | Spleen tumors | +++ | 24 |
| vΔCHPK | 84 | Spleen tumors | ++ | 31 |
| vΔCHPK | 85 | Kidney tumors | ++ | 31 |

[a]Relative level of fluorescence in feather follicles (see images below)

[b]Day post-infection

[c]Samples used for mass-spectrometry.

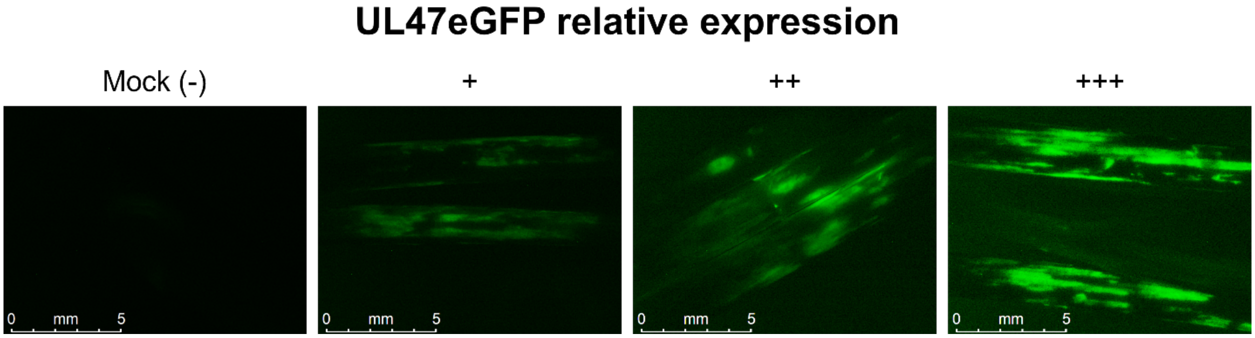

**S2 Table. Primers used to generate mutations in expression plasmids.**

| Construct[a] | Primer Name[b] | Sequence (5'- 3') |
|---|---|---|
| pcUS10ΔNLS1 | DEL_NLS1_US10For | TCTAGCAGGAGTGTGCAACT |
| | DEL_NLS1_US10Rev | CATGGCGGCAAGGGCA |
| pcUS10ΔNLS2 | DEL_NLS2_US10For | TCGTCTCGTTTGTGGGGT |
| | DEL_NLS2_US10Rev | TAGCGCAACATGTTCCCC |
| pcUS10ΔNLS3 | DEL_NLS3_US10For | ATGTATTGGTGTTGTCTTGGACA |
| | DEL_NLS3_US10Rev | TGTCGGGTGGAATGGCGA |
| pcUS10ΔNES | DEL_NES_US10For | GCCAGGACCATATTAACCGC |
| | DEL_NES_US10Rev | GGCAGAAAGTATATCATAACTCTGTTCT |
| pcICP27ΔNLS1/NES1 | DEL_NLS1_UL54For | GTAGATTCTGCATGCATTTCCC |
| | DEL_NLS1_UL54Rev | TGTAGCATATCGACTTGCTGA |
| pcICP27ΔNLS2 | DEL_NLS2_UL54For | CAATATCACAGACGTAATTTTCCG |
| | DEL_NLS2_UL54Rev | TGTCAAATTAGCTAAGCGTAGA |
| pcICP27ΔNES | DEL_NES2_UL54For | TTATCGCGCATTCGTCGTGT |
| | DEL_NES2_UL54Rev | AGATCTTCTTAACATATCCCCCATGG |

[a]Expression construct generated.

[b]Name of the primers.
